# Transcriptome of apple cv. Ametyst in *Rvi6*-effective and *Rvi6*-breaking *Venturia inaequalis* interactions identifies defense candidates

**DOI:** 10.64898/2026.09.15.751701

**Authors:** Sunee Kertbundit, Dimitrij Tyč, Radek Černý, Helena Štorchová, Miloslav Juříček

## Abstract

**Background:** *Rvi6*, introgressed from *Malus floribunda* 821, is the most widely deployed apple scab resistance gene, but *Venturia inaequalis* races overcoming it are widespread. Transcriptomic differences between *Rvi6*-effective and *Rvi6*-breaking interactions in one host genotype remain unresolved.

**Aims:** To compare these interactions in cv. Ametyst and nominate genes for subsequent validation.

**Methods:** Own-rooted Ametyst plants were mock-inoculated or inoculated with an *Rvi6*-avirulent isolate (Rin; incompatible) or an *Rvi6*-virulent isolate (Rola; compatible) and harvested at 4 days post-inoculation (21 mRNA libraries). Expression was quantified using GDDH13 reference-based and *de novo* Trinity strategies, followed by DESeq2, WGCNA and GSEA.

**Results:** Rin vs Rola yielded 1,339 candidate genes (more than two-fold change, FDR < 0.01) in the reference-based analysis. Curation of Rin-induced defense candidates retained 151 genes, 92 of which concentrated in a single co-expression module. Thirty-one also passed the Rin-vs-Rola thresholded test and were prioritized for genetic validation.

**Conclusions:** The incompatible interaction was associated with a coherent defense program and the compatible interaction with a distinct transcriptional state. With one isolate per class, isolate identity and *Rvi6* virulence cannot be separated.

**Highlights:** • Distinct host–isolate responses were resolved in one apple genotype.

• 151 Rin-induced defense candidates, 92 in one co-expression module.

• Candidate genes are targets for future genetic validation, not yet markers.

## Introduction

Apple scab, caused by the ascomycete *Venturia inaequalis*, is among the most economically damaging diseases of apple in temperate production regions, with losses of up to 70% reported under severe, uncontrolled epidemics (MacHardy, 1996; Jha et al., 2009; Bowen et al., 2011).

Under susceptible crops and conducive weather, control relies on repeated fungicide applications through the season, with attendant costs and selection for fungicide resistance (Bowen et al., 2011; Weber et al., 2025). Host resistance is therefore a major breeding objective, and major resistance genes have anchored many apple scab breeding programs (Gessler et al., 2006; Jha et al., 2009; Bus et al., 2011).

Foremost among these is *Rvi6* (formerly *Vf*), introgressed from *Malus floribunda* 821 and widely deployed in scab-resistant cultivars (Vinatzer et al., 2001; Bus et al., 2011). The *Rvi6* locus contains a cluster of Hcr*Vf*/*Vf*a receptor-like genes. *HcrVf2* (*Vfa2*) is sufficient to confer scab resistance when expressed in susceptible ‘Gala’ (Belfanti et al., 2004), whereas native-promoter constructs of *Vfa1*/*HcrVf1* and *Vfa2*/*HcrVf2* produced partial resistance in ‘Galaxy’ and ‘McIntosh’ (Malnoy et al., 2008). The resistance is no longer universally effective: *V. inaequalis* races able to sporulate on *Rvi6* cultivars have been reported from several production areas (Parisi et al., 1993; Bénaouf and Parisi, 2000; Guérin and Le Cam, 2004; Papp et al., 2020). This erosion makes a direct comparison of effective and resistance-breaking interactions biologically and practically relevant.

Transcriptomic studies of the apple–*Venturia inaequalis* pathosystem have cataloged resistance-associated responses (Degenhardt et al., 2005; Gusberti et al., 2013; Masoodi et al., 2022; Švara et al., 2024; Chen et al., 2025). Many designs compare different cultivars, so interaction outcome is entangled with host-genotype differences. Comparing an *Rvi6*-effective and an *Rvi6*-breaking isolate in the same host genotype removes that host-background bias, although isolate identity remains inseparable from virulence on *Rvi6* when only one isolate represents each class. Characterizing apple defense-response transcription and the signaling genes underlying resistance was identified as a priority for this pathosystem by Jha et al. (2009); RNA sequencing now permits this comparison at whole-transcriptome scale.

Our aims were threefold: (i) to compare the incompatible Rin and compatible Rola interactions in cv. Ametyst, which carries *Rvi6*; (ii) to evaluate the response with complementary reference-based (GDDH13) and *de novo* (Trinity) quantification strategies; and (iii) to identify candidate genes as targets for subsequent genetic validation.

## Materials and methods

### Plant material and inoculation

Own-rooted plants of *Malus × domestica* cv. Ametyst, which carries *Rvi6* (formerly *Vf*) introgressed from *M. floribunda* 821, were used. The cultivar (accession UEB 2650/2) was derived from the cross ‘Nela’ × ‘Vistabella’ made in 1987. Plants were propagated clonally by axillary shoot multiplication from *in vitro* shoot cultures of cv. Ametyst. Shoots were multiplied and rooted on agar-solidified half-strength Murashige and Skoog basal medium (Murashige and Skoog, 1962) supplemented with growth regulators for apple as described by Sriskandarajah et al. (1990). All experimental plants were therefore own-rooted clones of a single genotype, and no rootstock contributed to the response. Rooted plantlets were transferred to vermiculite in Magenta GA-7 vessels (Merck, Darmstadt, Germany) and grown for one to two months in a tissue-culture room at 21 °C under a 16-h photoperiod provided by fluorescent lamps, until roots and leaves were well established. They were then potted into a peat-based substrate amended with perlite and sand in 1-l pots and grown on under the same temperature and photoperiod. Plants were staged by leaf production rather than by age. A plant was inoculated once it had formed four to five new leaves after potting, at which point it bore 8–12 leaves in total. Only visually healthy, uniformly developed plants were used.

Conidia were washed from infected leaves of the two source cultivars with distilled water and concentrated by centrifugation at 4 °C; monoconidial isolates were then established following Parker et al. (1995). Conidial suspensions were stored in water at −20 °C, and the inoculum used here was prepared less than one month before inoculation; suspensions were adjusted to 2.5 × 10⁵ conidia ml⁻¹ before use, following Peil et al. (2018). Virulence on *Rvi6* was verified for both isolates before the experiment: Rin did not infect cultivars carrying *Rvi6*, whereas Rola did.

Plants were inoculated by spraying the suspension onto the foliage, and control plants were sprayed with distilled water. All plants were held in closed transparent boxes at 15–17 °C and approximately 100% relative humidity under low-intensity light for 24 h, and thereafter grown at approximately 21 °C under a 16-h light / 8-h dark photoperiod. Leaves were harvested 96 h (4 d) after inoculation, a single time point sampling the established phase of both interactions and matching the later sampling point of Gusberti et al. (2013). Two *V. inaequalis* isolates were used: Rin, isolated from ‘Rubín’ and avirulent on *Rvi6* (incompatible interaction), and Rola, isolated from ‘Rubinola’ and virulent on *Rvi6* (compatible interaction) (Fig. 1; Supplementary Table S6).

**Figure 1.**
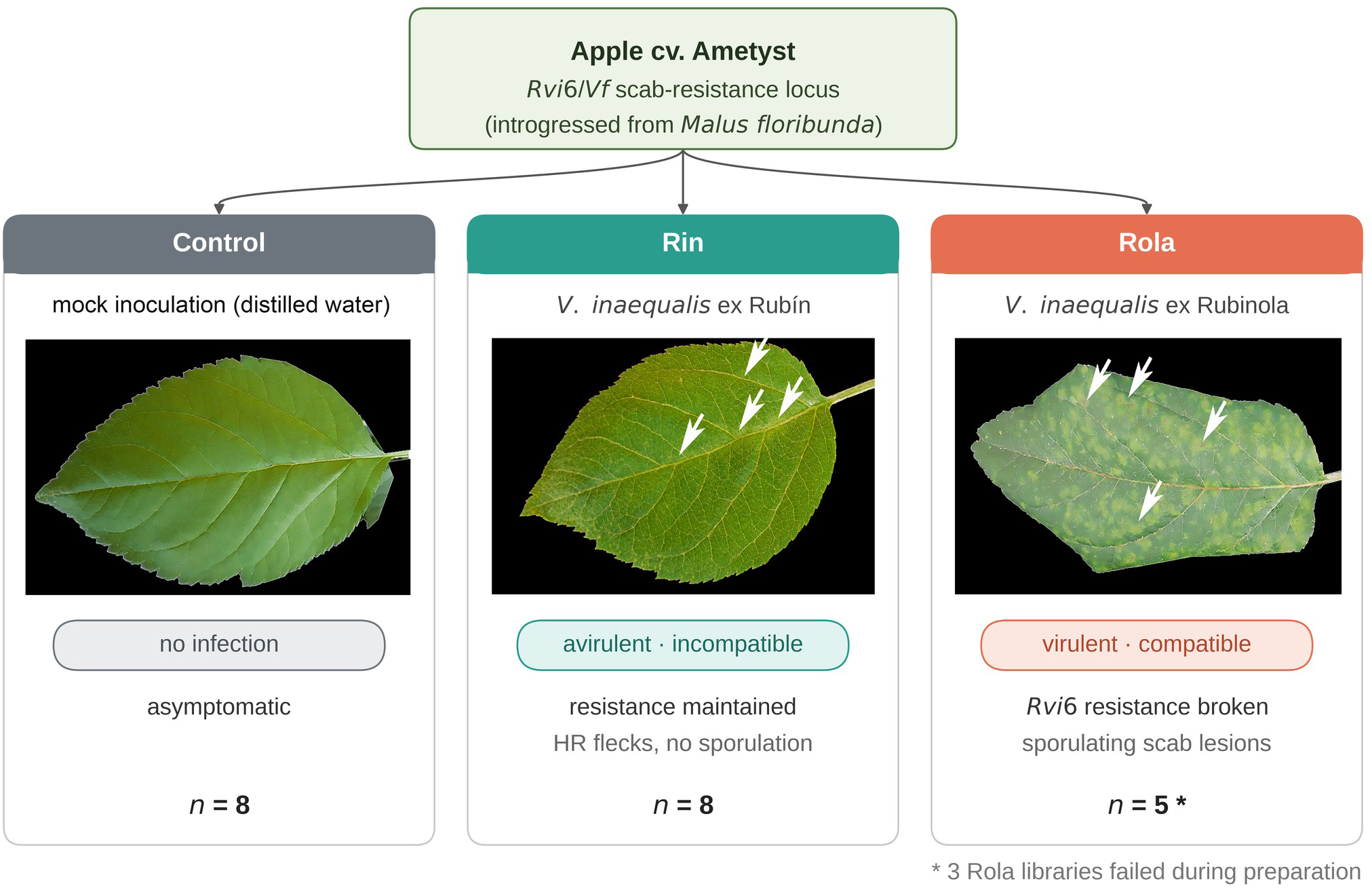
Experimental design and representative disease phenotypes in apple cv. Ametyst. Leaves of *Malus × domestica* ‘Ametyst’, carrying the *Rvi6* (formerly *Vf*) scab-resistance locus introgressed from *Malus floribunda*, were mock-inoculated with distilled water or inoculated with one of two *Venturia inaequalis* isolates. Control — mock-inoculated leaf, asymptomatic. Rin — isolate from ‘Rubín’, avirulent on *Rvi6*: incompatible interaction with sparse hypersensitive-response flecks and no sporulation. Rola — isolate from the *Rvi6*-carrying cultivar ‘Rubinola’, virulent on *Rvi6*: compatible interaction with extensive sporulating lesions. Arrows indicate representative symptoms. All RNA-seq leaves were harvested at 4 dpi; each sample was the three youngest fully expanded leaves of one plant, pooled, one plant per library, so each is an independent biological replicate; n is the number of biological samples. The analysis workflow is shown in Supplementary Fig. S1.

### RNA extraction and sequencing

All leaves were harvested simultaneously at 4 days post inoculation (dpi), and RNA was extracted from all samples at the same time. Each biological sample comprised the three youngest fully expanded leaves from the shoot apex of a single plant, pooled; one plant contributed one library. Leaves were immediately frozen in liquid nitrogen, ground, and total RNA was extracted with the Maxwell RSC Plant RNA Kit on a Maxwell RSC instrument (Promega, Madison, WI, USA; Cat. No. AS1500, www.promega.com). RNA integrity numbers (RINs), determined with an Agilent 2100 Bioanalyzer (Agilent Technologies, Santa Clara, CA, USA; www.agilent.com), ranged from 5.6 to 8.3. RNA was stored at −80 °C until library preparation. Poly(A)-enriched libraries were prepared with an Illumina TruSeq RNA Library Prep Kit (Illumina, San Diego, CA, USA; www.illumina.com) in two batches approximately six months apart, which were subsequently sequenced by different providers. Because library-preparation batch and sequencing provider were completely linked, they constitute a single technical batch factor.

Twenty-four libraries were prepared; 21 were sequenced and analyzed (8 control, 8 Rin and 5 Rola). Three Rola libraries failed during library preparation and were not removed as analytical outliers. Control and Rin were balanced 4/4 between the two batches, whereas Rola was distributed 3 in the first batch and 2 in the second. Libraries were sequenced by Eurofins Genomics (Konstanz, Germany) and by Institute of Applied Biotechnologies (IAB, Prague, Czechia) on Illumina NovaSeq 6000 instruments, paired-end 2 × 150 bp, as unstranded cDNA libraries, yielding 17.5–97.2 million read pairs per library (median 25.2 million; 5.29–29.36 Gb). The ordered sequencing depths were 20× and 50× for the first and second batches, respectively; per-library values are given in Supplementary Table S6.

### Reference-based quantification

Reads were error-corrected with Rcorrector 1.0.4 (Song and Florea, 2015), trimmed with Trim Galore v0.6.10 (Martin, 2011; Krueger, 2023), and aligned to the GDDH13 v1.1 genome from the Genome Database for Rosaceae (Daccord et al., 2017; Jung et al., 2019) using STAR 2.7.11 (Dobin et al., 2013). Fragments were quantified with featureCounts v2.0.0 (Liao et al., 2014) using -p -B -C -s 0 -t exon -g gene_name. A total of 38,467 genes entered the analysis.

### De novo assembly and quantification

A *de novo* transcriptome was assembled with Trinity 2.15.2 (Grabherr et al., 2011; Haas et al., 2013), as in our earlier transcriptome analyses of non-model species (Krüger et al., 2020; Gutiérrez-Larruscain et al., 2021), producing 443,892 transcripts grouped into 249,983 Trinity genes. One representative transcript per Trinity gene was retained. Reads were quantified against this gene-representative index with Salmon 1.10.2 (Patro et al., 2017) using automatic library-type inference, validateMappings, sequence-bias and GC-bias correction; tximport (Soneson et al., 2015) imported length-scaled abundance estimates at gene level. After count filtering, 119,509 genes were tested. Mapping rates were 75.7–79.6%. The lower rate reflects isoform collapsing, which leaves fragments unique to non-representative isoforms unmapped.

### Differential expression

Differential expression was analyzed independently for the reference-based and *de novo* quantification strategies (the two quantification layers) with DESeq2 (Love et al., 2014). The standard Wald test (alpha = 0.05) identified DEGs. A thresholded Wald test (alpha = 0.01, lfcThreshold = 1, altHypothesis = "greaterAbs") tested |log₂FC| ≤ 1 as the null directly rather than by post hoc filtering. Throughout, "DEG" refers to the standard Wald result and "candidate gene" exclusively to the thresholded result. The standard Wald test provides the DEG counts in the first column of Table 1; because the thresholded test is stricter, every candidate gene is also a DEG, but most DEGs are not candidate genes. Three contrasts were evaluated, with Rin vs Rola primary; positive log₂FC denotes higher expression in the first-named group. Benjamini– Hochberg correction was applied to all DESeq2 tests (Benjamini and Hochberg, 1995); GSEA significance is the permutation-based FDR q-value reported by that procedure, not a Benjamini– Hochberg adjustment. Adaptive shrinkage (Stephens, 2017) was applied through lfcShrink for the Rin vs Rola contrast solely to produce the MA plot in Fig. 2C; all tests, candidate definitions and reported fold changes use the unshrunken estimates.

**Figure 2.**
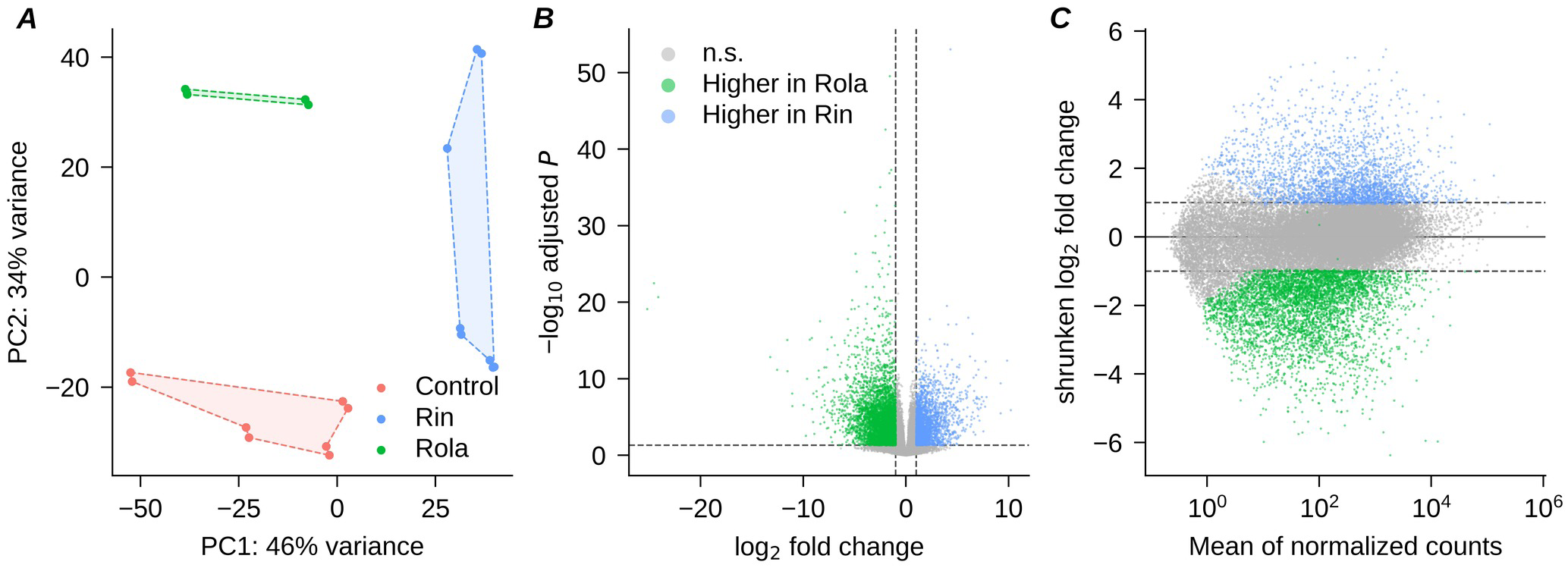
Overview of the Ametyst leaf transcriptome. (***A***) Principal component analysis of variance-stabilized (VST) expression values from the GDDH13 reference-based analysis (PC1 46%, PC2 34% of variance). Each point is one RNA-Seq library, colored by treatment; dashed outlines are group convex hulls (*n* = 8 control, 8 Rin, 5 Rola). (***B***) Volcano plot of the Rin vs Rola contrast; genes with adjusted *P* < 0.05 and |log2 fold change| ≥ 1 are colored by direction (blue, higher in Rin; green, higher in Rola), dashed lines marking these thresholds. (***C***) MA plot of the same contrast, with log2 fold changes shrunken by the ashr method plotted against mean normalized counts. Statistics are from DESeq2 (design ∼ condition, standard Wald test, alpha = 0.05); adjusted *P* values are Benjamini–Hochberg corrected. An equivalent PCA on the *de novo* Trinity transcriptome is shown in Supplementary Fig. S3.

**Table 1.**
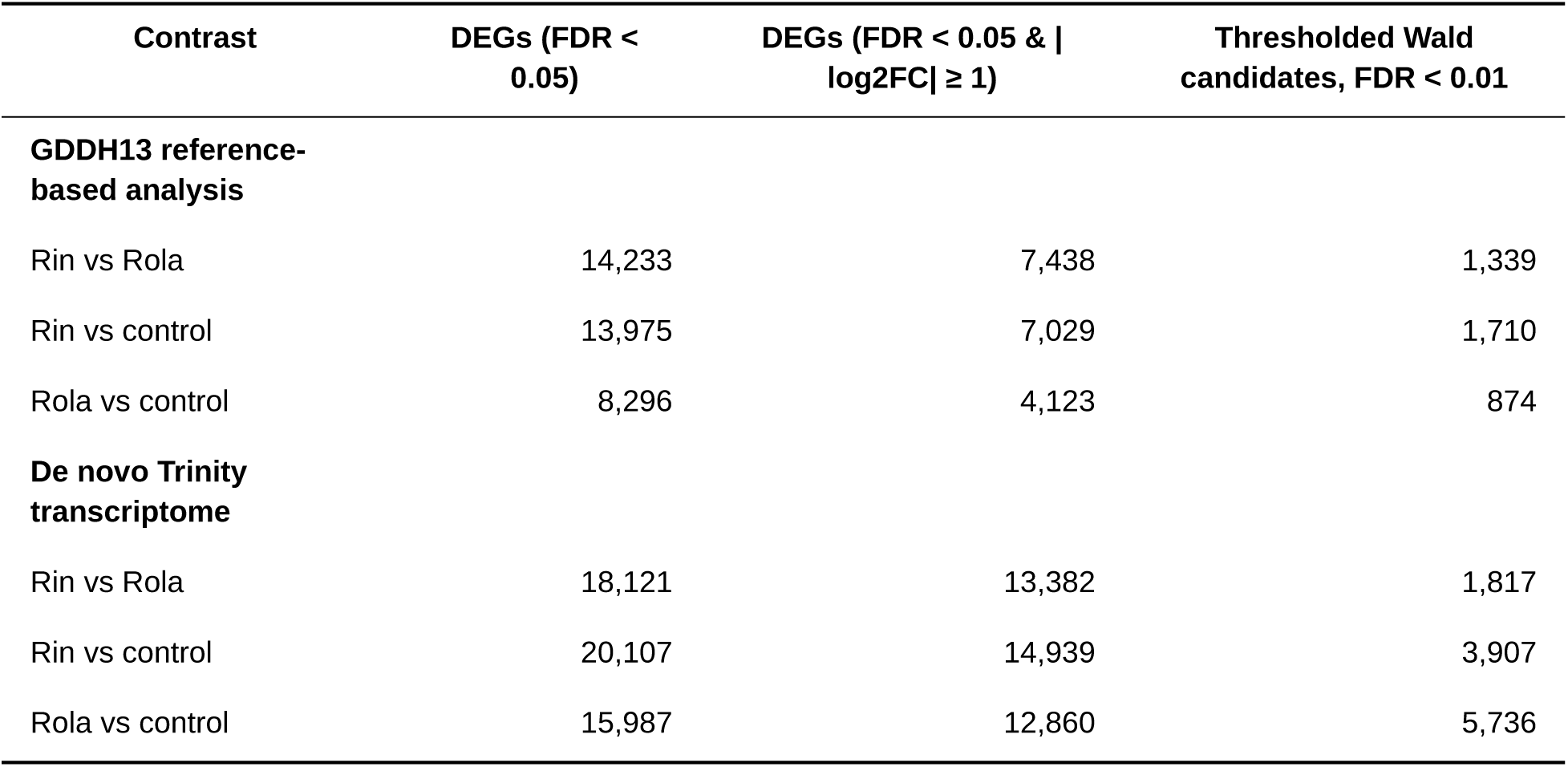
Differentially expressed genes across the three contrasts, GDDH13 reference-based and *de novo* Trinity analyses. Counts from DESeq2 (design ∼ condition) for each quantification strategy. Columns report differentially expressed genes (DEGs) at FDR < 0.05, those additionally passing |log2FC| ≥ 1 (standard Wald test), and the candidate set from the thresholded Wald test (FDR < 0.01, H0: |log2FC| ≤ 1).

The condition-only model is the primary analysis; as a sensitivity analysis, DESeq2 was additionally fitted with the technical batch factor (∼ batch + condition; originally coded as time); effect estimates were essentially unchanged (Pearson r ≥ 0.996 between the two models for every contrast in both quantification layers) and the candidate sets overlapped closely (Jaccard 0.85–0.95 across contrasts and layers), with the batch model returning slightly fewer candidates in every contrast. Of the 151 curated Rin-induced candidates, 140 met their defining criteria unchanged under the batch model; the other 11 are itemized in Supplementary Table S2. Both fits are reported in full in Supplementary Data S13 and S14. Results are reported in Table 1 and Supplementary Table S1.

### Co-expression network analysis

A module is a cluster of genes whose expression is correlated across samples — a statistical grouping, not a predefined metabolic or signaling pathway. Weighted gene co-expression network analysis (WGCNA; Langfelder and Horvath, 2008) was applied to the 10,000 most variable of the 38,467 GDDH13 genes at soft-threshold power 8, producing 18 informative modules plus a grey bin of 97 unassigned genes; 9,903 genes were assigned. Module–trait relationships were estimated as module eigengene correlations (Pearson r with nominal two-sided Student *P* values; BH-adjusted values in Supplementary Table S7). Candidate enrichment was tested against the 9,903 module-assigned genes and, secondarily, against the broader pool of 707 Rin-induced candidates. The latter tests whether the 151-gene set remains concentrated in the green module beyond Rin induction alone; it does not isolate the effect of defense annotation, because selection also used the Rola-vs-control classification and manual curation. OmicsBox hub tagging used kME ≥ 0.8 and *P* < 0.05; because 7,495 of 10,000 genes met this criterion, the tag is not a stringent measure of network centrality.

### Functional annotation and gene-set enrichment

Functional annotation was performed with OmicsBox 4.0.54 using BLASTX first against the *Malus × domestica*/GDDH13 protein set and then against Viridiplantae for sequences without a confident apple hit, together with InterProScan (Jones et al., 2014). Gene Ontology terms were assigned through Blast2GO (Conesa et al., 2005; Götz et al., 2008) and filtered by taxon (Viridiplantae) before final interpretation. RefSeq locus identifiers for GDDH13 genes were assigned by reciprocal protein BLAST against the NCBI *Malus × domestica* protein set. OmicsBox default parameters were used at every step and are listed, with the InterProScan member databases queried, in Supplementary Table S10. InterProScan was run through the OmicsBox CloudIPS service, which queries the InterPro release current at run time; the annotation was produced on 14 July 2026. The reciprocal protein BLAST was run locally with blastp from BLAST+ 2.17.0+ against the NCBI RefSeq protein set of assembly GCF_004115385.1 (ASM411538v1) on 9 July 2026; the expectation value, acceptance thresholds and reciprocal-best-hit flagging are given in Supplementary Table S10.

Functional annotations were reviewed manually for the 355 genes that were induced in Rin relative to the mock control according to the thresholded Wald and did not meet the thresholded-Wald criterion for either induction or repression in Rola relative to the control. All 151 genes judged on this review to have defense-related functions were retained and assigned to functional families by annotation keyword; these families are descriptive groupings of the annotation text, not Gene Ontology categories, and were not tested for enrichment. WGCNA module membership was not used at any stage of this selection.

Gene-set enrichment analysis (GSEA; Subramanian et al., 2005) used the complete DESeq2 Wald-statistic ranking, gene-set sizes of 15–500, 1,000 permutations and FDR *q* < 0.05. Of 673 GDDH13 GO terms tested, 314 were significant (162 Rin-high and 152 Rola-high).

### Cross-layer comparison and *Rvi6* screening

Trinity candidate transcripts were screened against the GDDH13 proteome by blastx from BLAST+ 2.17.0+ (Altschul et al., 1990). A confident GDDH13 protein match required ≥40% amino-acid identity across ≥50 aligned amino-acid residues, summed across high-scoring segment pairs to the best-scoring subject. Query coverage was not used because untranslated regions reduce transcript-level BLASTX coverage. “GDDH13 protein match” therefore denotes sequence similarity rather than orthology, and absence of a hit does not establish absence from the genome. Candidates without a confident match were additionally screened against *HcrVf1*– *HcrVf3* (GenBank AJ297739–AJ297741).

### Screening the *de novo* assembly for non-plant sequence

Because the libraries were prepared from infected leaf tissue, the *de novo* assembly necessarily contains transcripts of *Venturia inaequalis* and of other leaf-associated organisms. The 249,983 gene-representative sequences were screened with NCBI FCS-GX 0.5.5 (Astashyn et al., 2024) against the GX reference database build of 24 January 2023 (3,025,824 sequences, 709 Gbp), with *Malus × domestica* (taxonomy identifier 3750) declared as the expected organism.

Sequences carrying an EXCLUDE call were removed and the four TRIM calls applied. A REVIEW call marks a sequence the tool cannot resolve rather than a confirmed contaminant; the 281 such sequences were also removed, the conservative choice for a public deposit, leaving 150,655 sequences. NCBI FCS-adaptor was then used in eukaryote mode before deposition, and 28 sequences carrying internal Illumina TruSeq read-through adaptor, which FCS-adaptor does not act on because it is not terminal, were repaired on submission screening; the full sequence of cleaning steps and counts is given in Supplementary Table S15; the final cleaned set contained 150,621 sequences. Screening followed rather than preceded the differential-expression analysis, so the calls annotate it rather than repeat it: the taxonomic call was joined to the Trinity gene-level results and the non-plant fraction of each candidate set is reported in Supplementary Tables S5 and S11. No gene was removed before quantification and no reported statistic was recomputed. Reference-based quantification is far less affected because a read must align to the GDDH13 assembly and fall within an annotated gene model to be counted; conserved sequence can still cross-map, so this is a strong filter rather than an absolute one.

### Software and statistics

Software versions and key parameters are listed in Supplementary Table S10. Reporting follows MIAME principles (Brazma et al., 2001): sample metadata are in Supplementary Table S6, and the complete gene-level results for both quantification layers, the Rin-vs-Rola candidate classification and the WGCNA module assignments are provided in Supplementary Tables S1, S5, S8 and S11. The source data, software provenance and reproducibility status of every table are summarized in Supplementary Table S12. The two DESeq2 analysis notebooks are supplied as knitted reports, Supplementary Data S13 (GDDH13 reference-based strategy) and Supplementary Data S14 (Trinity strategy), so that every step from count import to the reported contrasts can be inspected as executed code.

#### Use of AI tools

This manuscript, including the presentation and interpretation of the Results, the Discussion and the Conclusions, was entirely written by the authors without the help of AI. AI-assisted tools were used only as supporting tools for scripting the extraction, restructuring and visualization of author-specified results, preparing preliminary figure and table elements from the original analysis outputs, checking internal consistency between the manuscript, tables and figures, verifying literature citations, and assisting with English-language correction and stylistic editing of author-written text.

## Results

### Disease phenotypes confirm the contrasting interactions

The inoculations produced the expected contrasting phenotypes (Fig. 1). Rin-inoculated plants showed hypersensitive flecking without sporulation, whereas Rola-inoculated plants developed sporulating lesions; mock-treated leaves remained asymptomatic. RNA-seq leaves were harvested at 4 dpi. The expression-analysis workflow is summarized in Supplementary Fig. S1.

### Global transcriptome structure separates the three treatments

Principal component analysis of the reference-based expression profiles separated the three treatment groups, with PC1 and PC2 explaining 46% and 34% of the variance, respectively (Fig. 2A); volcano and MA representations are shown in Fig. 2B,C. The Trinity strategy showed a similar treatment pattern (Supplementary Fig. S3). Clustering of the 500 most variable genes recovered the three treatment groups (Supplementary Fig. S2), and quality-control summaries are provided in Supplementary Fig. S5.

### Differential expression identifies 31 Rin-specific candidates for marker development

Table 1 compares the numbers of DEGs (FDR < 0.05) and the thresholded-Wald candidates for the three contrasts between the reference-based and *de novo* transcriptomes. In the incompatible vs compatible contrast (Rin-vs-Rola), the 1,339 reference-based candidates split into 338 Rin-high and 1,001 Rola-high.

Of the 1,339 reference-based Rin-vs-Rola candidates, 615 (45.9%) also passed the thresholded Rin-vs-control test; within that shared set the two contrasts were closely correlated (Pearson correlation coefficient *r* = 0.98) and all but one of the 615 genes changed in the same direction (Fig. 3). These data demonstrate a strongly concordant shared component between the two contrasts. They do not mean that the entire Rin-vs-Rola difference is Rin-driven: some candidates may also respond to Rola infection, but at a different level than to Rin. To obtain Rin-specific candidates we therefore worked entirely within the reference-based layer: of the 1,710 Rin-vs-control candidates, 707 were induced in Rin with baseMean ≥ 50, and 355 of those did not meet the thresholded-Wald criterion (adjusted *P* < 0.01, |log₂FC| > 1, baseMean ≥ 50) for either induction or repression in the Rola-vs-control contrast. Manual inspection of their annotations identified 151 with defense-related functions.

**Figure 3.**
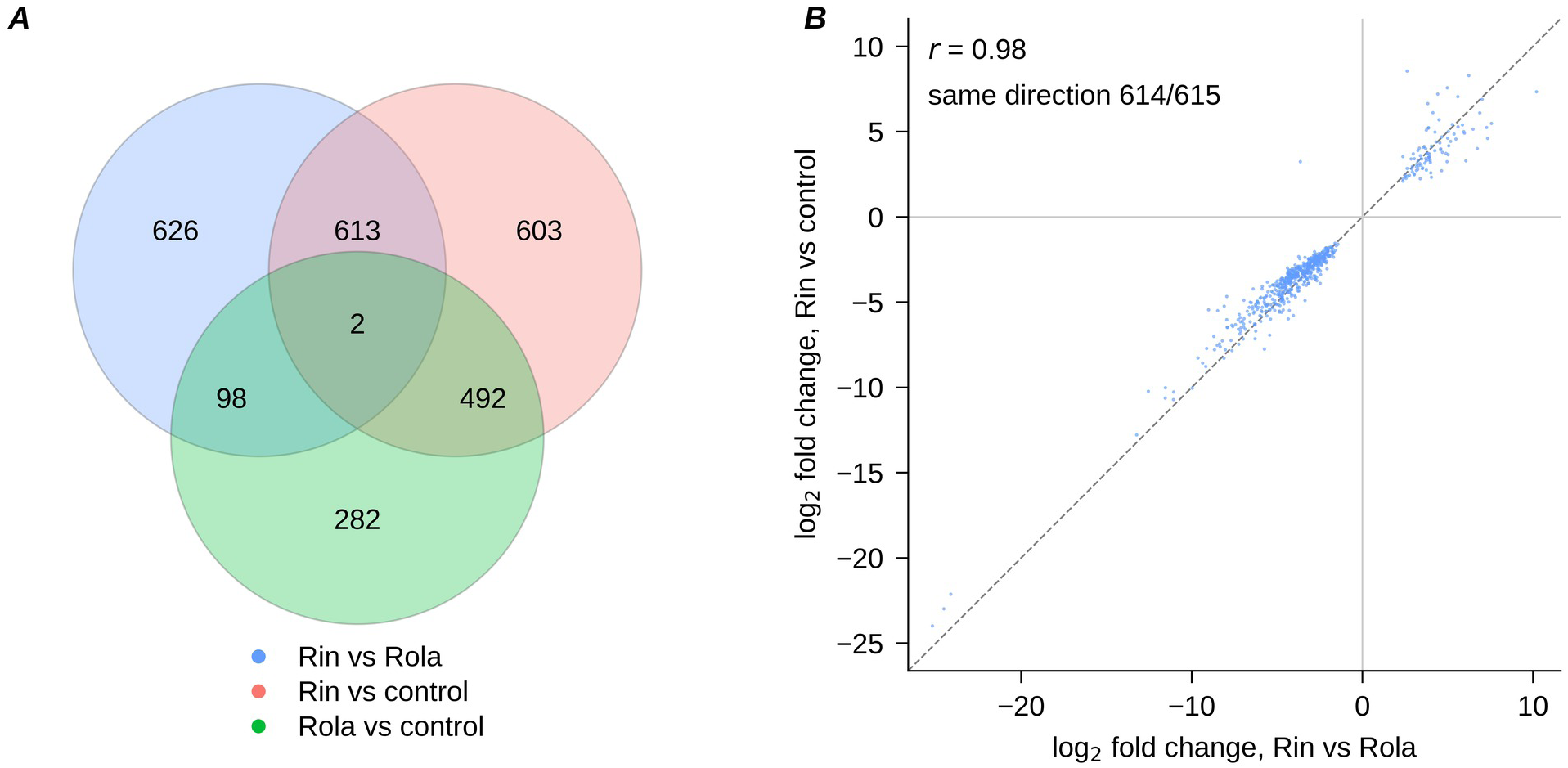
Rin-associated expression changes constitute the major shared component of the Rin-vs-Rola contrast. (A) Overlap of candidate gene sets identified by thresholded Wald tests for Rin vs Rola, Rin vs control, and Rola vs control. (B) For the 615 genes significant in both Rin vs Rola and Rin vs control, log2 fold changes in the two contrasts are plotted against each other; the dashed line denotes equality. Effects were highly concordant (*r* = 0.98), and 614 of 615 genes changed in the same direction. These results identify a strongly Rin-associated component of the transcriptional difference between the incompatible and compatible interactions. Candidate sets were defined using a thresholded Wald test with FDR < 0.01 and the null hypothesis |log2 fold change| ≤ 1; complete counts are given in Table 1. The panel shows the three GDDH13 contrasts only. The 151 curated defense candidates are not a fourth class: they are a subset of the Rin-vs-control induced branch, drawn as a selection funnel in Supplementary Fig. S1. Cross-layer overlap with the Trinity analysis was assessed by BLASTX similarity to GDDH13 proteins (Supplementary Table S5), not by a gene-identifier Venn, because the identifiers are not directly comparable.

These 151 genes are listed with their annotations in Supplementary Table S2, and the selection funnel is drawn in Supplementary Fig. S1. Of these, 130 fell into 11 functional families and 21 remained unclassified (Fig. 4A); the families are assigned by annotation keyword and are descriptive, not Gene Ontology categories.

**Figure 4.**
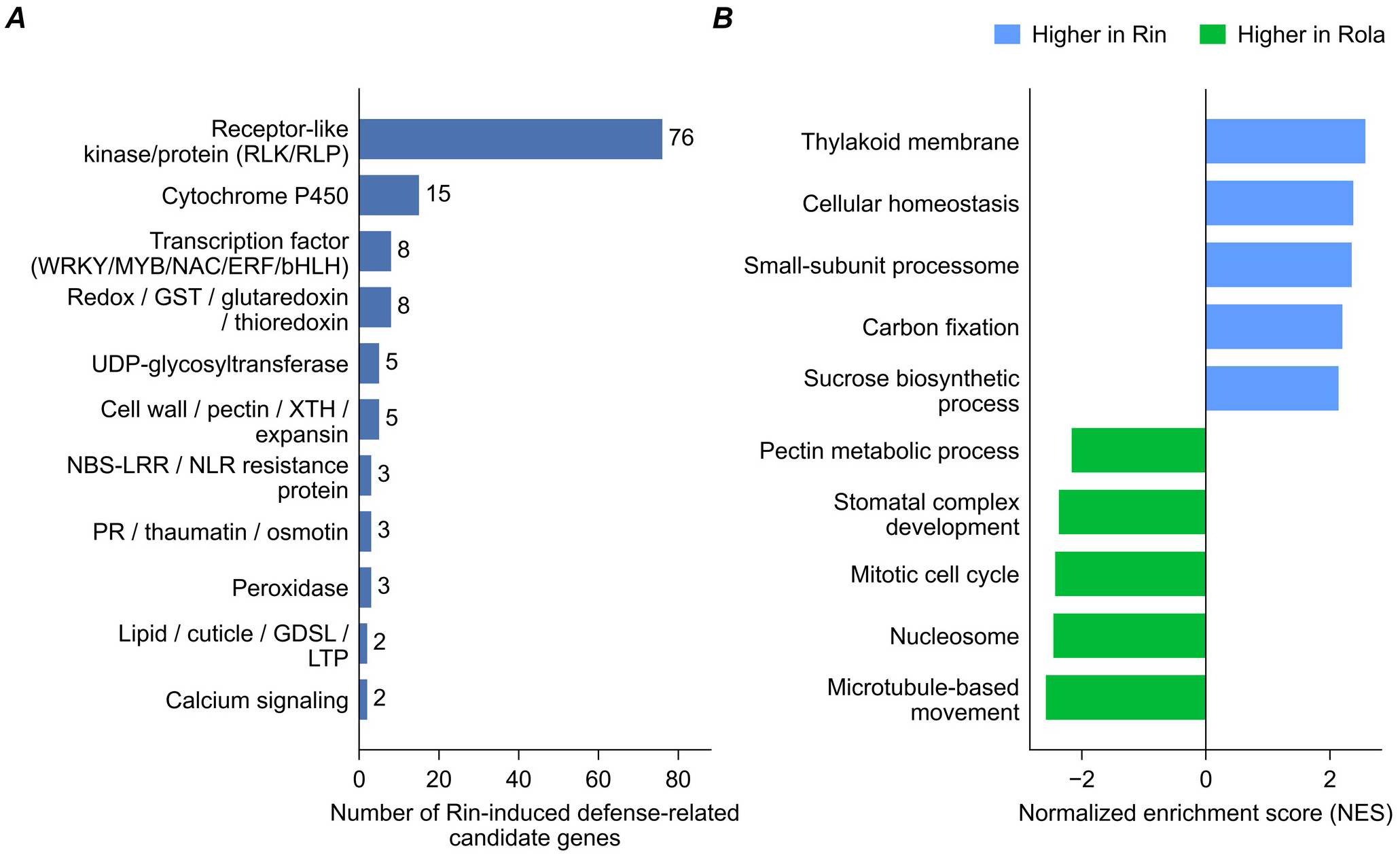
The Rin-induced defense program. (A) 151 curated Rin-induced defense-related candidate genes (Materials and methods) assigned to functional families by annotation keyword; bars give the number of genes per family. Of the 151 candidates, 130 fall into the 11 families shown and 21 remain unclassified. The set is dominated by receptor-like kinases and proteins (76 genes), together with cytochrome P450s (15), defense-related transcription factors (8) and redox enzymes (8), the biological rationale for the candidates in Table 2. Because the set is curated rather than exhaustive, family counts describe the prioritized subset, not the total induced in each family. (B) Ten representative Gene Ontology terms — five higher in Rin, five higher in Rola — enriched by gene-set enrichment analysis (GSEA) of the Rin vs Rola contrast (FDR *q* < 0.05), collapsed to one term per biological theme. Genes were ranked by the DESeq2 Wald statistic; positive normalized enrichment scores (NES) denote enrichment among genes higher in Rin (blue), negative among genes higher in Rola (green). Of 673 terms tested, 314 passed FDR *q* < 0.05 (162 Rin-enriched, 152 Rola-enriched); complete output in Supplementary Table S4. An equivalent GSEA on the *de novo* Trinity transcriptome is shown in Supplementary Fig. S4.

**Table 2.** Prioritized defense-response candidate genes with potential for marker development. The 31 Rin-induced defense-related candidate genes that also pass the thresholded Rin-vs-Rola test (FDR < 0.01, log2FC > 1), ranked by log2 fold change in that contrast; the 151 curated candidates were defined as described in Materials and methods. Positive values denote higher expression in Rin. The log2 fold changes are ordinary DESeq2 estimates shown for description; they do not define membership. RefSeq locus is the NCBI LOC identifier from reciprocal protein BLAST and is not one-to-one. Module is the WGCNA co-expression module (Fig. 5). Adjusted *P* values, mean normalized counts, module membership and hub-tag status are in Supplementary Table S2. Five genes — MD03G1014200, MD07G1240700, MD15G1239300, MD15G1239500 and MD17G1092400 — lose thresholded Rin-vs-Rola significance under the technical-batch model with effect estimates essentially unchanged (Supplementary Data S13).

| GDDH13 gene ID | RefSeq locus | Module | $\log_2$ fold change<br>Rin vs control | $\log_2$ fold change<br>Rin vs Rola | Description |
| --- | --- | --- | --- | --- | --- |
| MD02G1138900 | LOC103407082 | midnightblue | 7.34 | 10.21 | Peroxidase superfamily protein |
| MD16G1091200 | LOC103403007 | red | 5.16 | 6.48 | xyloglucan:xyloglucosyl transferase 33 |
| MD12G1075400 | LOC103423272 | green | 8.30 | 6.23 | Glycosyltransferase family 61 protein |
| MD07G1236500 | LOC103410350 | black | 3.30 | 6.06 | Leucine-rich repeat protein kinase family protein |
| MD16G1116100 | LOC103416681 | salmon | 5.41 | 5.87 | cytochrome P450 |
| MD13G1172100 | LOC103452977 | black | 4.43 | 5.17 | Thioredoxin superfamily protein |
| MD12G1161000 | LOC103450543 | midnightblue | 4.62 | 5.00 | Laccase/Diphenol oxidase family protein |
| MD04G1183000 | LOC103434046 | black | 3.73 | 4.90 | Leucine-rich repeat protein kinase family protein |
| MD11G1054100 | LOC103422644 | black | 4.69 | 4.65 | lipid transfer protein 1 |
| MD03G1014200 | LOC114823840 | green | 4.98 | 4.25 | peroxidase 2 |
| MD03G1077700 | LOC103427128 | red | 2.33 | 4.04 | calmodulin-binding receptor-like cytoplasmic kinase 1 |
| MD07G1236800 | LOC103410350 | black | 3.20 | 3.98 | Leucine-rich repeat protein kinase family protein |
| MD07G1240700 | LOC103420753 | black | 3.51 | 3.88 | Fe superoxide dismutase 2 |
| MD10G1250900 | LOC103422333 | green | 5.22 | 3.87 | wall associated kinase 5 |
| MD04G1181800 | LOC103434049 | salmon | 4.01 | 3.71 | Leucine-rich repeat protein kinase family protein |
| MD17G1092400 | LOC103425937 | black | 3.44 | 3.71 | Peroxidase superfamily protein |
| MD04G1182200 | LOC103408596 | salmon | 4.14 | 3.62 | Leucine-rich repeat protein kinase family protein |
| MD00G1023600 | LOC103432961 | black | 3.41 | 3.48 | Leucine-rich repeat transmembrane protein kinase |
| MD15G1239500 | LOC103424618 | green | 4.04 | 3.33 | Leucine-rich repeat transmembrane protein kinase |
| MD17G1126300 | LOC103405101 | black | 3.33 | 3.26 | UDP-Glycosyltransferase superfamily protein |
| MD15G1239400 | LOC103431980 | green | 3.90 | 3.25 | Leucine-rich repeat transmembrane protein kinase |
| MD01G1172500 | LOC103406468 | black | 2.79 | 3.24 | cytochrome P450 |
| MD06G1032200 | LOC103436877 | black | 2.94 | 3.16 | plant intracellular ras group-related LRR 4 |
| MD13G1103800 | LOC103452391 | black | 2.52 | 3.03 | cytochrome P450 |
| MD15G1239300 | LOC103431980 | green | 3.70 | 2.98 | Leucine-rich repeat transmembrane protein kinase |
| MD13G1103500 | LOC103430668 | black | 2.65 | 2.93 | cytochrome P450 |
| MD00G1183000 | LOC103431980 | green | 3.41 | 2.85 | Leucine-rich repeat transmembrane protein kinase |
| MD11G1066000 | LOC103447339 | midnightblue | 2.41 | 2.67 | Leucine-rich repeat protein kinase family protein |
| MD04G1066300 | LOC103427519 | black | 2.35 | 2.49 | cytochrome P450 |
| MD05G1257000 | LOC103438143 | green | 2.11 | 2.38 | Leucine-rich repeat transmembrane protein kinase |
| MD00G1046700 | LOC103432961 | green | 2.23 | 2.37 | Leucine-rich repeat transmembrane protein kinase |

Receptor-like kinases/proteins were most abundant (76), followed by cytochrome P450s (15), transcription factors (8) and redox enzymes (8). Only 31 of the 151 also passed the thresholded Rin-vs-Rola test in the Rin-high direction. Estimated effects are larger than that test alone conveys: 97 of the 151 have an estimated Rin-vs-Rola log₂ fold change above 1, so for 66 of them an induction above two-fold relative to Rola was not statistically established at the chosen threshold (Supplementary Table S2).

### Prioritized candidates and their genomic positions

The 31 candidates that also pass the thresholded Rin-vs-Rola test, with RefSeq locus and module assignment, are presented in Table 2; coordinates for all 151 are given in Supplementary Table S9. The mapping is not one-to-one: 15 NCBI LOC identifiers were each assigned to two or three GDDH13 genes (34 genes total). Nine clusters contained three or more candidates with successive genes separated by ≤ 200 kb; the largest contained seven green-module genes spanning 311 kb on Chr10, whereas three candidates on Chr15 occurred within 6.3 kb. This is consistent with tandem arrays of RLK/RLP-family expansions, and such groups should be regarded as clustered candidate regions rather than independent genomic signals. Nine candidates sit on Chr00, unanchored scaffolds with no map position.

### Gene-set enrichment reproduces across layers

GSEA in the reference-based strategy returned 314 significant GO terms of 673 tested; ten representative terms, five from each direction, are shown in Fig. 4B, and the complete output is given in Supplementary Table S4. Directionality was concordant across quantification strategies: among the 54 terms significant in both the reference-based and the *de novo* Trinity analyses, Pearson *r* = 0.87 and 50 were enriched in the same direction (Supplementary Fig. S4).

### The candidates concentrate in a single co-expression module

Co-expression analysis resolved 18 informative modules. Ninety-two of the 151 Rin-induced defense candidates fell in the green module — an 8.1-fold enrichment over the 11.3 expected of the 9,903 module-assigned genes (hypergeometric *P* = 1.4 × 10⁻⁶⁵).

The green module showed an eigengene–Rin correlation of *r* = 0.69 and an eigengene–control correlation of *r* = −0.81 (Fig. 5); nominal and BH-adjusted *P* values for all module–trait pairs are given in Supplementary Table S7.

**Figure 5.**
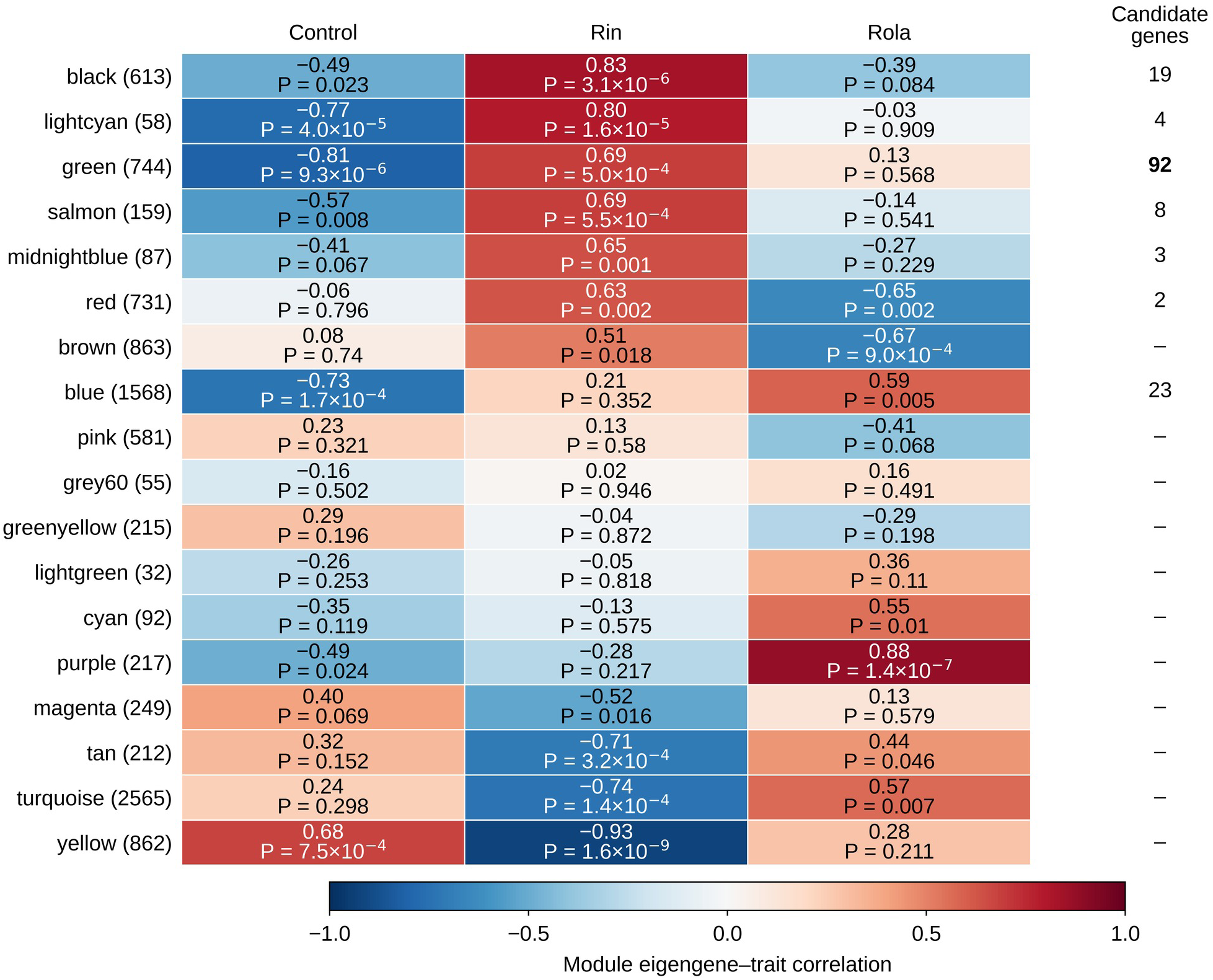
Co-expression modules and their association with treatment. Weighted gene co-expression network analysis (WGCNA) of the 10,000 most variable genes (soft-thresholding power 8) assigned 9,903 genes to 18 informative co-expression modules; a further 97 genes fell into the unassigned grey bin, not plotted. The heatmap shows the Pearson correlation between each module eigengene and the control, Rin and Rola treatments (red, positive; blue, negative), with the nominal two-sided Student *P* value beneath each coefficient; modules are ordered by their correlation with Rin. Benjamini–Hochberg adjusted values across all 54 tests (18 modules × 3 treatments) are given in Supplementary Table S7. Module–trait statistics and the candidate enrichment in the green module are reported in Results. Green-module genes meeting the OmicsBox hub-tag criterion (module membership ≥ 0.8 and module-membership *P* < 0.05) included PR5-like and wall-associated receptor kinases, S-locus lectin kinases, and NBS-LRR/LRR resistance proteins (Supplementary Table S3). The ’Candidate genes’ column gives the number of candidates per module.

Compared with the broader pool of 707 Rin-induced candidates, the final 151-gene set remained 1.8-fold enriched in the green module (92 observed versus 50.8 expected; hypergeometric *P* = 6.1 × 10⁻¹⁵), so the concentration is not merely a consequence of Rin induction. The module itself is defense-related rather than pathway-specific: among its 744 genes, receptor-like kinases and LRR proteins (149, 20% against 7% of network genes) and pathogenesis-related or disease-resistance annotations (53, 7% against 2.5%) are each about 2.8-fold more frequent than in the network as a whole, and cytochrome P450s 2.1-fold; no single metabolic pathway dominates, and the most common single annotation is “protein of unknown function”. Green-module gene-level statistics are listed in Supplementary Table S3. The remaining 59 candidates were spread over six modules with no comparable enrichment; the 23 in the blue module are exactly what its size predicts. Green is only the third most Rin-correlated module (behind black, *r* = 0.83, and lightcyan, *r* = 0.80) — it is where the candidates concentrate, not the top-correlated module — and because the network and the candidate set derive from the same expression data, this represents expression coherence rather than independent validation.

### Cross-layer concordance and the *Rvi6* screen

Trinity gene identifiers have no one-to-one correspondence with GDDH13 gene models, so the two layers were compared through protein-level BLASTX matching rather than by intersecting gene sets. Of 1,817 Trinity Rin-vs-Rola candidates (Supplementary Table S5), 1,154 (64%) had a confident GDDH13 protein match; of these, 815 were candidates in both layers and changed in the same direction, the remaining classes being listed in Supplementary Table S5. The screen identified no additional transcript that could be confidently attributed to the *HcrVf1*–*HcrVf3* introgressed receptor cluster.

### Non-plant sequence in the *de novo* layer

FCS-GX returned an action for 99,332 of the 249,983 Trinity genes, almost all EXCLUDE calls, dominated by animal, fungal and bacterial sequence; the most abundant taxa were *V. inaequalis*, *Homo sapiens* and the greenhouse whitefly *Trialeurodes vaporariorum*, the last consistent with an incidental infestation of the plant room. Counts and median lengths are given in Supplementary Table S15. The calls need caution: the *H. sapiens* sequences are much shorter than the *V. inaequalis* ones, and FCS-GX is calibrated for genome assemblies rather than short transcripts.

The flag is distributed very unevenly across the Trinity candidate classes of the primary contrast (Supplementary Table S5). Of the 815 candidates that were also candidates in the reference-based layer and changed in the same direction, only two genes (0.2%) were flagged. Every Trinity-only class was flagged at between 44% and 66% (Supplementary Table S5), dominated by *V. inaequalis* (97%).

The same gradient accounts for the Trinity candidate totals in Table 1: the flagged fraction rises from about a third in Rin vs Rola to about half in Rin vs control and four fifths in Rola vs control (Supplementary Table S15). Absence of a flag is not sufficient evidence of apple origin; FCS-GX reports only what it can assign to another taxon.

The 6.6-fold excess of Trinity over GDDH13 candidates in the Rola-vs-control contrast (5,736 against 874), set against 1.4-fold and 2.3-fold in the two Rin contrasts, is therefore predominantly caused by fungal transcripts, not by an additional host response detected by the *de novo* layer. Nineteen of the 265 defense-annotated Trinity candidates in the primary contrast are likewise *V. inaequalis* (Supplementary Table S5).

The fungal fraction is itself informative. The assembly therefore holds an incidental dual RNA-seq dataset, and the number of *V. inaequalis* genes meeting the thresholded induction criterion tracks the phenotype: 4,485 in Rola relative to the mock control against 1,868 in Rin, a 2.4-fold difference at 4 dpi. These are counts of fungal genes passing the differential-expression criterion rather than a calibrated measure of fungal biomass, but they are a molecular correlate of the contrasting phenotypes in Fig. 1. The fungal fraction was not analyzed further: the experiment was not designed for it and the libraries were not normalized for fungal biomass.

## Discussion

### What the comparison establishes

The Rin-vs-Rola contrast therefore characterizes two distinct host–isolate interactions in Ametyst. Holding the apple genotype, sampled tissue and harvest time constant removed host-background variation that complicates comparisons among cultivars (Degenhardt et al., 2005; Masoodi et al., 2022; reviewed by Švara et al., 2024).

However, the Rin-vs-Rola contrast did not distinguish virulence on *Rvi6* from all other isolate-specific differences, because only a single isolate represents each interaction phenotype.

Additional *Rvi6*-avirulent and *Rvi6*-virulent isolates would be required to attribute the transcriptomic differences specifically to resistance-breaking ability. Within these limits, the design provides a direct view of transcriptional states associated with incompatible and compatible interactions in the same host genotype.

### The function of the candidate genes

The candidate set is dominated by RLKs and RLPs (76 of 151), consistent with extensive receptor-mediated signaling during an *Rvi6*-effective interaction (Boller and Felix, 2009; Couto and Zipfel, 2016). The 15 cytochrome P450 candidates are consistent with specialized-metabolite metabolism. The eight redox candidates—a superoxide dismutase, two thioredoxins, two glutathione S-transferases, two laccases and an NAD(P)-linked oxidoreductase—point to antioxidant/redox-homeostasis and cell-wall oxidation processes. No respiratory burst oxidase homolog occurs among the 151 candidates, but this does not exclude an early oxidative burst mediated by genes outside the thresholded set or by post-transcriptional regulation (Lamb and Dixon, 1997; Schuler and Werck-Reichhart, 2003). Eight candidates are annotated as transcription factors: WRKY72, WRKY48 and WRKY65, three genes annotated as MYB73 and one as MYB36, and a bHLH protein. WRKY factors form the transcriptional networks that reprogram gene expression during plant immune responses (Eulgem and Somssich, 2007), and MYB proteins act broadly in phenylpropanoid and cell-wall responses. All eight carry log₂ fold changes of 2.3–4.6 against the mock control, yet none of them passes the thresholded Rin-vs-Rola test, so on these data the regulatory step is not itself a discriminating signal between the two interactions. The cross-layer GSEA themes fall within process classes repeatedly reported in apple–*Venturia* studies, including pathogen recognition/signaling, redox homeostasis, photosynthesis, specialized metabolism and cell-wall processes (Švara et al., 2024). Concentration of 92 candidates in one module supports a coherent expression program, however it is not independent validation because candidate selection and network construction use the same expression data. The green module is therefore best interpreted not as a single pathway but as a coordinated Rin-associated defense-expression state, enriched for receptor-like kinases and proteins, pathogenesis-related and disease-resistance annotations, and cytochrome P450s.

### Comparison with *Rvi6* gain-of-function

A direct recent comparator is Chen et al. (2025), who overexpressed *Rvi6* in transgenic ‘Orin’ apple calli and compared them with empty-vector controls. Their design isolates constitutive effects of *Rvi6* in undifferentiated tissue, whereas ours examines the native locus in intact plants infected with isolates that differ in virulence on *Rvi6*. *Rvi6* overexpression enriched plant–pathogen interaction and MAPK pathways and reduced auxin signaling and callus growth. Chen et al. interpreted the plant–pathogen interaction enrichment through a *Cf-9*-related CDPK–Rboh–ROS/Ca²⁺ signaling model. This converges with the strong representation of receptor-like proteins and kinases in our candidate set, but does not establish that the same cascade operates in Ametyst. Their enhanced antioxidant capacity is compatible with our redox-homeostasis candidates, although our data do not exclude an oxidative burst. Rola-high enrichment of auxin-response, cell-cycle and cell-division terms is directionally consistent with the growth–defense trade-off reported in the callus system. Photosynthesis-related genes were downregulated by *Rvi6* overexpression in dark-maintained calli, whereas photosynthesis/thylakoid terms were Rin-high in leaves; tissue and growth-condition differences preclude a direct comparison, and our pattern may instead reflect stronger photosynthetic impairment in the compatible interaction. We did not recover significant MAPK or flavonoid pathway enrichment at *q* < 0.05. Chen et al. defined DEGs with | log₂FC| ≥ 1 and unadjusted *P* < 0.05, whereas our gene-level sets use BH-adjusted thresholds, further limiting direct gene-list comparison.

### Comparison with ontogenic resistance

Our candidate set, dominated by receptor-like kinases and proteins, reflects receptor-mediated signaling in the *Rvi6*-effective interaction, and can be compared with the study of ontogenic resistance by Gusberti et al. (2013), who held cultivar constant but varied leaf age: young susceptible and old ontogenically resistant ‘Golden Delicious’ leaves were sampled after mock treatment or inoculation at 72 and 96 h. Their 96-h sampling matches our 4-dpi time point, although the resistance mechanisms differ. Our samples were young leaves — the three youngest fully expanded — corresponding to their susceptible age class, so the comparison sets *Rvi6*-mediated resistance in young tissue against ontogenic resistance in old tissue. Ontogenic resistance is broadly race-nonspecific and considered durable, but is partial and can wane as tissues senesce; *Rvi6* is race-specific and has been overcome (Švara et al., 2024). Gusberti et al. identified five qRT-PCR-validated candidates: peroxidase 3, lipoxygenase, lipid transfer protein and metallothionein 3-like were constitutively higher in old leaves, whereas enhanced disease susceptibility 1 was lower. These are mainly defense-output, lipid and redox functions. The overlap with our results is partial: three peroxidases occur among our 151 Rin-induced candidates (log₂FC 3.44–7.34 for Rin vs control, the contrast defining the set), ten in the wider Rin-vs-control candidate set, and a lipid-transfer protein 1 is present (log₂FC 4.69 for Rin vs control), whereas lipoxygenase appears only in the wider set. No metallothionein or EDS1 candidate was identified, although annotation and gene-model differences may contribute. The studies therefore suggest partial convergence on oxidative and lipid-associated outputs while differing in resistance context and upstream recognition.

### Phenolic metabolism

Different experimental systems yield different phenolic signatures. Chen et al. (2025) reported upregulation of flavonoid biosynthesis in *Rvi6*-overexpressing calli; Gusberti et al. (2013) found no consistent induction of flavonoid or phenolic-precursor transcripts in ontogenically resistant leaves; and Švara et al. (2025), measuring metabolites in *Rvi6*-transgenic ‘Gala’ lines and diploid/tetraploid cultivar pairs, found procyanidin dimers positively associated with resistance while many other phenolics were lower in *Rvi6*-harboring genotypes. These studies assess different tissues, resistance mechanisms and molecular levels, and mixed-direction changes among branches of phenolic metabolism can obscure aggregate pathway enrichment.

In our data no phenylpropanoid or flavonoid term reached *q* < 0.05, although individual genes changed in both directions: of twelve phenylpropanoid/flavonoid candidates in the Rin-vs-control set, eight were higher in Rin and four in the control, including three chalcone/stilbene synthases (log₂FC 7.39–9.11) and five O-methyltransferases (5.94–7.69); UDP-glycosyltransferases and laccases split almost evenly (7 up/8 down and 5 up/6 down). We detected no differential expression of committed flavan-3-ol branch enzymes. Because transcript abundance does not directly predict metabolite concentration, our data neither support nor contradict a procyanidin-specific mechanism at 4 dpi.

### Pathogenesis-related proteins and the limits of an induction contrast

PR proteins are established components of apple–*Venturia* defense, including apoplastic chitinases, β-1,3-glucanases, thaumatin-like proteins and proteases (Gau et al., 2004; Mohamed et al., 2025). The curated 151-gene set is nevertheless PR-poor: it contains an osmotin, a pathogenesis-related family protein and a thaumatin-superfamily protein, but no chitinase. This reflects the selection criterion rather than an absence of PR induction. PR genes are strongly induced in the incompatible interaction: among the 707 Rin-induced candidates, nineteen carry PR-type annotation, including three chitinases (log₂FC 2.58–5.37 for Rin vs control), eleven MLP-like PR-10 proteins (3.69–8.23) and five thaumatin-, osmotin- or PR-family proteins (2.97–5.29). Almost all of them are induced in the compatible interaction as well: all three chitinases and nine of the eleven PR-10 proteins are classified as common-infection-induced, whereas the three PR-family and thaumatin proteins that are Rin-specific are precisely the three in the curated set. The classical PR arsenal therefore belongs to the shared response to infection rather than to the Rin-specific program, and Fig. 4A describes the latter. Salicylate signaling is a different matter. No salicylic-acid, NPR1 or systemic-acquired-resistance component reaches candidate status in either quantification strategy, and across the 16 SA-related genes in the tested set no absolute log₂ fold change exceeds 1.17. The broad GO category “regulation of defense response” was Rola-high, but such a category includes both positive and negative regulators and is not evidence of stronger defense per se. Gau et al. (2004) found induced apoplastic PR accumulation in susceptible ‘Elstar’ and constitutive PR expression/protein patterns in resistant ‘Remo’. Thus, an induction-based contrast may miss defenses already active before inoculation. Testing that possibility in Ametyst would require baseline comparison with susceptible material; it cannot be resolved from a single-cultivar dataset.

### The compatible interaction

The Rola-high side of the primary contrast (1,001 of 1,339 candidates) admits at least four mutually non-exclusive explanations: active suppression of a Rin-associated program, different infection kinetics sampled at one time point, a qualitatively distinct compatible-interaction program, or isolate-specific effects unrelated to *Rvi6* virulence. One time point and one isolate per interaction class cannot distinguish among these possibilities. A time course including multiple independently collected *Rvi6*-avirulent and *Rvi6*-virulent isolates would be required.

### Two quantification strategies as an analytical cross-check

The high GO-theme directionality concordance (*r* = 0.87) between the reference and *de novo* strategies supports robustness to quantification method, although both use the same biological libraries and are not independent validation datasets. The Trinity screen did not identify an additional defense candidate confidently attributable to *HcrVf1*–*HcrVf3*; it does not establish that all introgressed or Ametyst-specific sequence is represented by GDDH13.

### Non-plant sequence in the *de novo* layer

Screening the gene-representative set after the analysis was complete showed that the two-layer design was self-protecting. Only 2 of 815 (0.2%) same-direction cross-layer candidates were flagged as non-plant by FCS-GX, compared with 44–66% in the Trinity-only classes, almost all of it *V. inaequalis*. A read must align to the GDDH13 assembly and fall within an annotated gene model to be counted, so reference-based quantification strongly filters non-apple sequence — not absolutely, since conserved sequence can cross-map. Cross-layer confirmation therefore acts as a taxonomic filter as well as a consistency check, and the Trinity-only fraction is not a reservoir of Ametyst-specific host sequence invisible to GDDH13. Annotation is affected too: with the Blast2GO taxonomy filter set to Viridiplantae a fungal transcript is assigned to its nearest plant homolog, which is why 19 *V. inaequalis* transcripts carry plant defense-gene names among the Trinity candidates. Screening before annotation would have prevented them in a *de novo* host–pathogen transcriptome.

None of this affects the reference-based results, the 151 curated candidates, the 31 prioritized genes or the co-expression analysis, all of which derive from the reference layer. Finally, the fungal fraction is a result rather than a nuisance: the 2.4-fold difference in the number of *V. inaequalis* genes meeting the thresholded induction criterion, higher in the compatible interaction, is a molecular counterpart to the visual phenotype, and the deposited set and its taxonomic calls preserve it for analysis of the pathogen side.

### Marker outlook

The 31 prioritized genes are targets for marker discovery, not yet the validated markers. A usable breeding marker requires a polymorphism, association with resistance, and validation in germplasm or a segregating population—none of which can be established from 21 libraries of one cultivar. The next step is genomic variant discovery across the genomic regions corresponding to the 151 candidate genes (coordinates in Supplementary Table S9), followed by linkage or association testing. Because single major resistance genes can be overcome, durable breeding strategies increasingly emphasize pyramiding multiple resistance sources and quantitative resistance loci (Jha et al., 2009; Švara et al., 2024). Validated polymorphisms in additional defense-associated loci could eventually contribute to such strategies, but no marker was developed or validated here.

## Conclusion

In cv. Ametyst, the *Rvi6*-effective Rin interaction was associated with a large, coherent defense-related transcriptional response, whereas the Rola interaction showed a distinct expression pattern. Receptor-like kinases and receptor-like proteins dominated the Rin-induced candidate set, and 92 of 151 candidates concentrated in one co-expression module.

The 31 prioritized genes provide a starting point for marker discovery, subject to polymorphism discovery and genetic validation. Interpretation is limited to the two host–isolate combinations studied and should not be generalized to all *Rvi6*-avirulent and *Rvi6*-virulent isolates.

## Abbreviations

DEG: differentially expressed gene;
Dpi: days post-inoculation;
FCS-GX: NCBI Foreign Contamination Screen using the genome cross-species aligner (GX);
FDR: false discovery rate;
GS: gene significance;
GSEA: gene-set enrichment analysis;
kME: module membership;
NES: normalized enrichment score;
RLK: receptor-like kinase;
RLP: receptor-like protein;
WGCNA: weighted gene co-expression network analysis.

## Acknowledgements

This work was supported by the Ministry of Agriculture of the Czech Republic through the National Agency for Agricultural Research, Programme of Applied Research of the Ministry of Agriculture for 2017-2025 (ZEMĚ), project No. QK21010390 ‘Modern breeding using molecular-genetic methods to make selection and practical application of new apple tree varieties with high resistance to significant apple diseases faster and more effective’. We thank Květa Rabochová for technical assistance with the tissue culture work.

## Statements

### Data and materials availability

Raw reads and the processed expression matrices were deposited in the NCBI Gene Expression Omnibus under accession GSE345841 (samples GSM10017734–GSM10017754); the reads are also available from the Sequence Read Archive under BioProject PRJNA1522210 (runs SRR40468427–SRR40468447). The GEO accession holds the complete 249,983-sequence Trinity gene-representative set used for quantification, together with its per-sequence FCS-GX taxonomic calls. The cleaned subset of that set, 150,621 sequences from which non-plant, adaptor and vector sequence has been removed, is archived together with the analysis scripts at Zenodo (https://doi.org/10.5281/zenodo.22707497).

Complete processed gene-level results for both quantification strategies, and sample metadata, are provided in the Supplementary Information: all 38,467 queried GDDH13 genes in the Supplementary Table S1 workbook and all 119,509 queried Trinity genes in the Supplementary Table S11 workbook. The two DESeq2 analysis notebooks are provided in full as knitted reports, Supplementary Data S13 (GDDH13 reference-based strategy) and Supplementary Data S14 (Trinity strategy).

### Biological materials availability

*Malus × domestica* cv. Ametyst (accession UEB 2650/2) is maintained in tissue culture at the Institute of Experimental Botany of the Czech Academy of Sciences and is available from the corresponding author upon reasonable request, subject to applicable institutional and phytosanitary requirements. The *Venturia inaequalis* isolates used in this experiment are no longer available; newly collected isolates from the original host sources would not necessarily be genetically identical to those used in this study.

## Ethical and legal standards compliance

The authors declare that the work did not involve experiments subject to special ethical regulation and complied with applicable institutional and national requirements.

