## Supplementary tables and figures for "Transcriptome of apple cv. Ametyst in *Rvi6*-effective and *Rvi6*-breaking *Venturia inaequalis* interactions identifies defense candidates": 00_Supplementary_contents.pdf

| File | Content |
| --- | --- |
| 01_Supplementary_figure_legends.pdf | Legends to Supplementary Figures S1–S5 |
| 02_SupplementaryFigure_S1_workflow.pdf | Experimental design and expression-analysis workflow |
| 03_SupplementaryFigure_S2_heatmap.pdf | Expression heatmap of the 500 most variable genes |
| 04_SupplementaryFigure_S3_trinity_pca.pdf | PCA based on the de novo Trinity transcriptome |
| 05_SupplementaryFigure_S4_trinity_gsea.pdf | GSEA of the Rin vs Rola contrast, Trinity layer |
| 06_SupplementaryFigure_S5_qc.pdf | Quality control of the RNA-Seq data |
| 07_SupplementaryTable_S1_DE_results.pdf | Table S1 — caption and description |
| 08_SupplementaryTable_S1_DE_results_workbook.xlsx | Table S1 data — all 38,467 queried GDDH13 genes, three contrasts |
| 09_SupplementaryTable_S2_candidates.pdf | The 151 curated Rin-induced defense-related candidate genes |
| 10_SupplementaryTable_S3_green_module.pdf | Candidates of the WGCNA green module |
| 11_SupplementaryTable_S4_GO_terms.pdf | All 673 GO terms tested by GSEA, GDDH13 layer |
| 12_SupplementaryTable_S5_Trinity_vs_GDDH13.pdf | All 1,817 Trinity Rin-vs-Rola candidates, cross-layer classification and FCS-GX calls |
| 13_SupplementaryTable_S6_sample_overview.pdf | Sample metadata for the 21 libraries |
| 14_SupplementaryTable_S7_module_trait.pdf | WGCNA module–trait analysis |
| 15_SupplementaryTable_S8_WGCNA_gene_modules.pdf | Table S8 — caption and description |
| 16_SupplementaryTable_S8_WGCNA_gene_modules.xlsx | Table S8 data — all 10,000 genes in the co-expression network |
| 17_SupplementaryTable_S9_candidate_coordinates.pdf | GDDH13 coordinates of the curated candidates |
| 18_SupplementaryTable_S10_software_versions.pdf | Software versions and key parameters |
| 19_SupplementaryTable_S11_Trinity_all_genes.pdf | Table S11 — caption and description |
| 20_SupplementaryTable_S11_Trinity_all_genes.xlsx | Table S11 data — all 119,509 queried Trinity genes |
| 21_SupplementaryTable_S12_provenance.pdf | Data source and software provenance of every table |
| 22_SupplementaryData_S13_DESeq2_GDDH.pdf | Knitted DESeq2 report, GDDH13 reference-based strategy |
| 23_SupplementaryData_S14_DESeq2_Trinity.pdf | Knitted DESeq2 report, de novo Trinity strategy |
| 24_SupplementaryTable_S15_screening_summary.pdf | FCS-GX and FCS-adaptor contamination screening summary |

Tables and figures are provided as PDF. Three supplementary tables (S1, S8, S11) contain the complete gene-level result sets and are additionally provided as Excel workbooks, since they are data files rather than text or figures.
