## Supplementary tables and figures for "Transcriptome of apple cv. Ametyst in *Rvi6*-effective and *Rvi6*-breaking *Venturia inaequalis* interactions identifies defense candidates": 01_Supplementary_figure_legends.pdf

**Supplementary Figure S1. Experimental design and analysis workflow.** Clonally propagated own-rooted plants of apple cv. Ametyst, which carry the *Rvi6*/*Vf* locus introgressed from *Malus floribunda* 821, were mock-inoculated (control) or inoculated with the *Venturia inaequalis* isolate Rin (from 'Rubín'; avirulent on *Rvi6*, incompatible interaction) or Rola (from 'Rubinola'; virulent on *Rvi6*, compatible interaction). Of 24 libraries prepared, 21 were analyzed (8 control, 8 Rin, 5 Rola); three Rola libraries failed during library preparation (Supplementary Table S6). Reads were quantified by two complementary quantification strategies: against the GDDH13 reference genome (Genome Database for Rosaceae) by STAR alignment and featureCounts (38,467 genes tested), and against a *de novo* Trinity assembly of Ametyst (443,892 transcripts, 249,983 genes) by Salmon quantification and tximport (119,509 genes tested). Each strategy was analyzed separately with DESeq2 under a condition-only design (~ condition); differentially expressed genes were called by the standard Wald test (FDR < 0.05) and the candidate sets by the thresholded Wald test (alpha = 0.01, lfcThreshold = 1, altHypothesis = "greaterAbs"). Three contrasts were extracted, with Rin vs Rola as the primary comparison; positive log2 fold change indicates higher expression in Rin. Downstream analyses comprised WGCNA on the 10,000 most variable genes (power 8, 18 informative modules), functional annotation and GSEA, blastx reconciliation of the 1,817 Trinity candidates against the GDDH13 proteome, and BLAST screening of the candidates against *HcrVf1*–*HcrVf3*, which did not identify additional candidate transcripts uniquely attributable to the *Malus floribunda* introgression. Within the reference-based layer only, the Rin-specific selection proceeded in four steps: of the 1,710 Rin-vs-control candidates, 707 were induced in Rin with baseMean ≥ 50; 355 of those had no thresholded Rola-vs-control call; annotation review retained 151 with defense-related functions, 92 of them in the WGCNA green module; and 31 of the 151 also pass the thresholded Rin-vs-Rola test. Cross-layer reconciliation plays no part in this selection. Those 31 are reported in Table 2; only 9 of them are green-module genes, consistent with the green module reflecting Rin induction more strongly than Rin-versus-Rola discrimination.

**Supplementary Figure S2. Expression heatmap of the 500 most variable genes.** log2(DESeq2-normalized count + 1) expression of the 500 genes with the highest variance across all libraries, of 38,467 quantified, row-scaled to z-scores (red, high; blue, low relative to the gene mean). Rows (genes) and columns (samples) were clustered by correlation distance with average linkage; the color strip above the heatmap marks the treatment of each library and is labeled in place. Sample clustering recovers the three treatment groups as one contiguous block each (*n* = 8 control, 8 Rin, 5 Rola).

**Supplementary Figure S3. Principal component analysis based on the *de novo* Trinity transcriptome.** As Figure 2***A***, but using Salmon quantification against the Ametyst Trinity assembly (PC1 42%, PC2 34%). Both panels apply the DESeq2 plotPCA method to the 500 most variable genes, so the two are directly comparable. The separation of control, Rin and Rola closely reproduces the GDDH13 reference-based analysis, showing that the treatment structure is similar under the two quantification strategies.

**Supplementary Figure S4. GSEA of the Rin vs Rola contrast based on the *de novo* Trinity transcriptome.** (A) As Figure 4B, but using the Ametyst Trinity assembly with OmicsBox GO annotation: ten representative terms — four higher in Rin, six higher in Rola — collapsed to one per biological theme from 355 theme-matched terms significant at FDR $q < 0.05$. (B) Normalized enrichment scores of the 54 GO terms significant at FDR $q < 0.05$ in both quantification layers, plotted against each other; Pearson $r$ = 0.87, with 50 of 54 terms enriched in the same direction. Points are colored according to the direction of enrichment in the Trinity layer; the dashed line denotes equality between layers. The theme-level enrichment pattern is therefore highly concordant with the GDDH13 reference-based analysis. The numbers of significant terms are not comparable between the two layers, because the two OmicsBox exports were pre-filtered differently. NES, normalized enrichment score.

**Supplementary Figure S5. Quality control of the RNA-Seq data.** (A) Sequencing depth per library, expressed as paired-end fragments processed by Salmon (17.5–97.2 million, median 25.1 million; the corresponding median of the raw read pairs reported in Materials and methods is 25.2 million); one fragment corresponds to one read pair. (B) Salmon mapping rate per library against the *de novo* Trinity assembly (75.7–79.6%, median 77.3%). (C) featureCounts fragment assignment per library for the GDDH13 reference-based layer, as proportions of the fragments in each category; assigned fragments account for 86.2–89.6% (median 87.8%), the remainder being multi-mapping, ambiguous or falling outside an annotated feature. featureCounts (v2.0.0) was run with -p -B -C, so fragments were counted rather than individual mates, and only fragments with both ends aligned and non-chimeric were included. Libraries are grouped by treatment. Sample codes: C, control; RIN, Rin; ROL, Rola.
