## Supplementary tables and figures for "Transcriptome of apple cv. Ametyst in *Rvi6*-effective and *Rvi6*-breaking *Venturia inaequalis* interactions identifies defense candidates": 07_SupplementaryTable_S1_DE_results.pdf

**Supplementary Table S1. Differential expression results, GDDH13 reference-based analysis.**

Summary of the complete DESeq2 results (design ~ condition) for the three contrasts — Rin vs Rola, Rin vs control and Rola vs control — provided in full as the accompanying workbook SupplementaryTable_S1_DE_results_workbook.xlsx, one worksheet per contrast. Each worksheet contains all 38,467 quantified genes with the columns gene_id, gene_name, baseMean, log2FoldChange, lfcSE, stat, pvalue, padj and Description, sorted by adjusted P value. Positive log2 fold change denotes higher expression in the first-named group of each contrast. Differentially expressed gene counts are from the standard Wald test; candidate genes are from the thresholded Wald test (H0: |log2FC| ≤ 1, altHypothesis = "greaterAbs"), which tests the fold-change threshold directly and therefore yields a smaller, more conservative set than a post hoc fold-change filter applied to the standard test. Counts are identical to the GDDH13 block of Table 1.

| Contrast | Worksheet | Genes tested | DEGs (FDR < 0.05) | DEGs (FDR < 0.05 & \|log2FC\| ≥ 1) | Thresholded Wald candidates, FDR < 0.01 |
|---|---|---|---|---|---|
| Rin vs Rola | Rin_vs_Rola | 38,467 | 14,233 | 7,438 | 1,339 |
| Rin vs control | Rin_vs_control | 38,467 | 13,975 | 7,029 | 1,710 |
| Rola vs control | Rola_vs_control | 38,467 | 8,296 | 4,123 | 874 |
