## Supplementary tables and figures for "Transcriptome of apple cv. Ametyst in *Rvi6*-effective and *Rvi6*-breaking *Venturia inaequalis* interactions identifies defense candidates": 09_SupplementaryTable_S2_candidates.pdf

**Supplementary Table S2. Rin-induced defense-related candidate genes.**

The 151 genes classified as Rin-induced. Membership was defined in three steps from the Rin-vs-control candidates (thresholded Wald test, H0: |log2FC| ≤ 1; FDR < 0.01): 707 were induced in Rin with baseMean ≥ 50; 355 of these additionally did not meet the thresholded-Wald criterion in either direction for Rola vs control; and manual review of the annotations of those 355 retained 151 genes with defense-related functions. Genes are grouped by functional family and ranked within each family by log2 fold change in the Rin-vs-control contrast. The families are assigned by annotation keyword, exactly as in Fig. 4A; they are descriptive groupings of the annotation text, not Gene Ontology categories, and were not tested for enrichment. Of the 151 genes, 130 fall into the 11 families shown and 21 are unclassified. Module gives the WGCNA co-expression module and kME the module membership; Hub marks genes meeting the OmicsBox hub-tag criterion (kME ≥ 0.8 and module-membership P < 0.05). NCBI LOC is the identifier assigned by reciprocal protein BLAST (— indicates no confident assignment). Statistics are displayed for description rather than to define membership, and are given for both contrasts: Rin vs control (ordinary DESeq2 estimates; all genes listed also pass the standard Wald test at FDR < 0.01 with log2FC > 1) and Rin vs Rola (thresholded Wald test, H0: |log2FC| ≤ 1). Because the thresholded test asks whether induction exceeds two-fold, a high adjusted P value in the Rin vs Rola columns indicates that the gene is not confidently induced beyond that margin relative to Rola, not that expression is equivalent. Positive log2FC denotes higher expression in Rin. The 31 candidates that also pass the thresholded Rin-vs-Rola test are presented in Table 2. Batch-model status records how each gene behaves when the analysis is repeated with the technical batch factor included (~ batch + condition; see Supplementary Data S13): “retained” means the gene still meets the full classification above (140 genes); “threshold crossing” means its adjusted P value moved across 0.01 while the effect estimate stayed essentially unchanged (6 genes, median standard-error inflation 1.02 and largest change in log2FC 0.04); “not evaluable” means independent filtering excluded it under the batch model, so it was neither retained nor rejected (2 genes); and “classification changed” means it acquired a thresholded Rola-vs-control call and is therefore no longer operationally Rin-specific (3 genes). The condition-only model is the primary analysis; the batch-adjusted fit is reported as a robustness analysis.

| Gene ID | NCBI LOC | Functional family | Module | kME | Hub | baseMean | Rin vs control log2FC | Rin vs control padj | Rin vs Rola log2FC | Rin vs Rola padj | Batch-model status | Description |
|---|---|---|---|---|---|---|---|---|---|---|---|---|
| MD05G1207800 | LOC114825146 | NBS-LRR / NLR resistance protein | green | 0.947 | yes | 106 | 6.05 | 5.92e-12 | 1.87 | 1.00e+0 | retained | NB-ARC domain-containing disease resistance protein |
| MD05G1207400 | LOC114825146 | NBS-LRR / NLR resistance protein | green | 0.956 | yes | 165 | 5.78 | 1.01e-10 | 1.76 | 1.00e+0 | retained | NB-ARC domain-containing disease resistance protein |
| MD04G1011700 | LOC103429293 | NBS-LRR / NLR resistance protein | green | 0.969 | yes | 281 | 4.45 | 2.82e-17 | 2.12 | 2.45e-1 | retained | Disease resistance protein (TIR-NBS-LRR class) family |
| MD04G1020400 | LOC103402536 | Receptor-like kinase/protein (RLK/RLP) | green | 0.964 | yes | 339 | 4.94 | 3.01e-12 | 1.57 | 1.00e+0 | retained | disease resistance family protein / LRR family protein |
| MD00G1101700 | LOC103419471 | Receptor-like kinase/protein (RLK/RLP) | green | 0.954 | yes | 249 | 4.40 | 8.40e-13 | 2.40 | 1.80e-1 | retained | cysteine-rich RLK (RECEPTOR-like protein kinase) 26 |

| Gene ID | NCBI LOC | Functional family | Module | kME | Hub | baseMean | Rin vs control<br><br>log2FC | Rin vs control<br><br>padj | Rin vs Rola<br><br>log2FC | Rin vs Rola<br><br>padj | Batch-model status | Description |
|---|---|---|---|---|---|---|---|---|---|---|---|---|
| MD10G1250500 | LOC103422333 | Receptor-like kinase/protein (RLK/RLP) | green | 0.971 | yes | 277 | 4.29 | 7.03e-11 | 1.99 | 6.03e-1 | retained | wall-associated kinase 2 |
| MD10G1212700 | LOC108172485 | Receptor-like kinase/protein (RLK/RLP) | green | 0.907 | yes | 1,042 | 4.16 | 1.16e-9 | 3.36 | 1.33e-2 | retained | Leucine-rich repeat protein kinase family protein |
| MD15G1426700 | LOC103417504 | Receptor-like kinase/protein (RLK/RLP) | green | 0.964 | yes | 2,705 | 4.14 | 3.56e-9 | 1.16 | 1.00e+0 | retained | disease resistance family protein / LRR family protein |
| MD10G1306800 | LOC103446654 | Receptor-like kinase/protein (RLK/RLP) | green | 0.915 | yes | 57 | 4.14 | 1.50e-9 | 1.07 | 1.00e+0 | retained | S-locus lectin protein kinase family protein |
| MD02G1250800 | LOC103408537 | Receptor-like kinase/protein (RLK/RLP) | green | 0.982 | yes | 169 | 4.14 | 1.13e-9 | 1.25 | 1.00e+0 | retained | receptor serine/threonine kinase |
| MD04G1182200 | LOC103408596 | Receptor-like kinase/protein (RLK/RLP) | salmon | 0.862 | yes | 286 | 4.14 | 4.46e-10 | 3.62 | 3.19e-3 | retained | Leucine-rich repeat protein kinase family protein |
| MD10G1248500 | LOC103446854 | Receptor-like kinase/protein (RLK/RLP) | green | 0.956 | yes | 87 | 4.05 | 5.27e-9 | 2.01 | 6.14e-1 | retained | wall-associated kinase 2 |
| MD15G1239500 | LOC103424618 | Receptor-like kinase/protein (RLK/RLP) | green | 0.869 | yes | 225 | 4.04 | 8.72e-10 | 3.33 | 9.85e-3 | retained | Leucine-rich repeat transmembrane protein kinase |
| MD02G1251800 | LOC103408537 | Receptor-like kinase/protein (RLK/RLP) | green | 0.989 | yes | 101 | 4.02 | 1.18e-12 | 1.32 | 1.00e+0 | retained | PR5-like receptor kinase |
| MD04G1181800 | LOC103434049 | Receptor-like kinase/protein (RLK/RLP) | salmon | 0.884 | yes | 639 | 4.01 | 4.54e-10 | 3.71 | 1.48e-3 | retained | Leucine-rich repeat protein kinase family protein |

| Gene ID | NCBI LOC | Functional family | Module | kME | Hub | baseMean | Rin vs control log2FC | Rin vs control padj | Rin vs Rola log2FC | Rin vs Rola padj | Batch-model status | Description |
|---|---|---|---|---|---|---|---|---|---|---|---|---|
| MD15G1239400 | LOC103431980 | Receptor-like kinase/protein (RLK/RLP) | green | 0.882 | yes | 754 | 3.90 | 2.57e-16 | 3.25 | 3.15e-4 | retained | Leucine-rich repeat transmembrane protein kinase |
| MD07G1014100 | LOC103412581 | Receptor-like kinase/protein (RLK/RLP) | blue | 0.829 | yes | 50 | 3.88 | 1.22e-5 | 0.49 | 1.00e+0 | retained | cysteine-rich RLK (RECEPTOR-like protein kinase) 5 |
| MD05G1190200 | LOC103409199 | Receptor-like kinase/protein (RLK/RLP) | green | 0.925 | yes | 109 | 3.87 | 3.20e-7 | 2.24 | 4.90e-1 | retained | Leucine-rich repeat receptor-like protein kinase family protein |
| MD17G1062200 | LOC103409759 | Receptor-like kinase/protein (RLK/RLP) | green | 0.977 | yes | 69 | 3.86 | 4.30e-11 | 0.72 | 1.00e+0 | classification changed | Malectin/receptor-like protein kinase family protein |
| MD02G1253600 | LOC108170170 | Receptor-like kinase/protein (RLK/RLP) | green | 0.963 | yes | 447 | 3.80 | 5.12e-7 | 1.20 | 1.00e+0 | retained | receptor serine/threonine kinase |
| MD10G1251200 | LOC103446854 | Receptor-like kinase/protein (RLK/RLP) | green | 0.977 | yes | 260 | 3.79 | 2.60e-11 | 1.57 | 1.00e+0 | retained | wall-associated kinase 2 |
| MD02G1235600 | LOC114821487 | Receptor-like kinase/protein (RLK/RLP) | green | 0.877 | yes | 87 | 3.76 | 6.67e-8 | 1.85 | 8.15e-1 | retained | PR5-like receptor kinase |
| MD04G1183000 | LOC103434046 | Receptor-like kinase/protein (RLK/RLP) | black | 0.914 | yes | 188 | 3.73 | 9.60e-15 | 4.90 | 8.64e-11 | retained | Leucine-rich repeat protein kinase family protein |
| MD15G1239300 | LOC103431980 | Receptor-like kinase/protein (RLK/RLP) | green | 0.875 | yes | 88 | 3.70 | 2.17e-11 | 2.98 | 8.54e-3 | retained | Leucine-rich repeat transmembrane protein kinase |
| MD14G1207100 | LOC103455550 | Receptor-like kinase/protein (RLK/RLP) | salmon | 0.927 | yes | 65 | 3.68 | 6.65e-7 | 1.44 | 1.00e+0 | retained | cysteine-rich RLK (RECEPTOR-like protein kinase) 25 |

| Gene ID | NCBI LOC | Functional family | Module | kME | Hub | baseMean | Rin vs control log2FC | Rin vs control padj | Rin vs Rola log2FC | Rin vs Rola padj | Batch-model status | Description |
|---|---|---|---|---|---|---|---|---|---|---|---|---|
| MD10G1273200 | LOC103410611 | Receptor-like kinase/protein (RLK/RLP) | blue | 0.946 | yes | 212 | 3.63 | 6.18e-8 | 0.21 | 1.00e+0 | retained | Leucine-rich repeat transmembrane protein kinase |
| MD08G1107900 | LOC103428814 | Receptor-like kinase/protein (RLK/RLP) | green | 0.978 | yes | 783 | 3.49 | 6.54e-18 | 1.69 | 4.88e-1 | retained | wall-associated kinase 2 |
| MD10G1308000 | LOC108174318 | Receptor-like kinase/protein (RLK/RLP) | blue | 0.942 | yes | 380 | 3.45 | 3.86e-9 | 0.58 | 1.00e+0 | retained | S-locus lectin protein kinase family protein |
| MD00G1023600 | LOC103432961 | Receptor-like kinase/protein (RLK/RLP) | black | 0.855 | yes | 4,560 | 3.41 | 2.80e-9 | 3.48 | 1.05e-3 | retained | Leucine-rich repeat transmembrane protein kinase |
| MD00G1183000 | LOC103431980 | Receptor-like kinase/protein (RLK/RLP) | green | 0.883 | yes | 784 | 3.41 | 4.85e-18 | 2.85 | 3.48e-4 | retained | Leucine-rich repeat transmembrane protein kinase |
| MD03G1246400 | LOC103452293 | Receptor-like kinase/protein (RLK/RLP) | blue | 0.899 | yes | 64 | 3.37 | 4.58e-6 | 0.76 | 1.00e+0 | retained | Leucine-rich repeat transmembrane protein kinase |
| MD03G1246600 | LOC103452293 | Receptor-like kinase/protein (RLK/RLP) | green | 0.917 | yes | 351 | 3.33 | 1.25e-6 | 0.63 | 1.00e+0 | retained | Leucine-rich repeat transmembrane protein kinase |
| MD10G1307800 | LOC103446652 | Receptor-like kinase/protein (RLK/RLP) | green | 0.964 | yes | 192 | 3.31 | 1.47e-8 | 1.40 | 1.00e+0 | retained | S-locus lectin protein kinase family protein |
| MD07G1236500 | LOC103410350 | Receptor-like kinase/protein (RLK/RLP) | black | 0.951 | yes | 1,385 | 3.30 | 4.93e-8 | 6.06 | 2.61e-12 | retained | Leucine-rich repeat protein kinase family protein |

| Gene ID | NCBI LOC | Functional family | Module | kME | Hub | baseMean | Rin vs control<br><br>log2FC | Rin vs control<br><br>padj | Rin vs Rola<br><br>log2FC | Rin vs Rola<br><br>padj | Batch-model status | Description |
|---|---|---|---|---|---|---|---|---|---|---|---|---|
| MD10G1214300 | LOC103445917 | Receptor-like kinase/protein (RLK/RLP) | green | 0.968 | yes | 316 | 3.30 | 3.84e-11 | 0.97 | 1.00e+0 | retained | Leucine-rich repeat receptor-like protein kinase family protein |
| MD07G1236800 | LOC103410350 | Receptor-like kinase/protein (RLK/RLP) | black | 0.930 | yes | 664 | 3.20 | 1.02e-17 | 3.98 | 7.67e-11 | retained | Leucine-rich repeat protein kinase family protein |
| MD01G1178700 | LOC103415095 | Receptor-like kinase/protein (RLK/RLP) | green | 0.963 | yes | 187 | 3.20 | 2.57e-8 | 1.25 | 1.00e+0 | retained | disease resistance family protein / LRR family protein |
| MD05G1217100 | LOC103428093 | Receptor-like kinase/protein (RLK/RLP) | green | 0.946 | yes | 64 | 3.16 | 6.32e-13 | 1.23 | 1.00e+0 | retained | S-locus lectin protein kinase family protein |
| MD00G1124500 | LOC103401065 | Receptor-like kinase/protein (RLK/RLP) | blue | 0.950 | yes | 54 | 3.11 | 9.77e-6 | -0.16 | 1.00e+0 | threshold crossing | Leucine-rich repeat receptor-like protein kinase family protein |
| MD10G1251400 | LOC103422333 | Receptor-like kinase/protein (RLK/RLP) | green | 0.975 | yes | 245 | 3.10 | 2.02e-11 | 1.33 | 1.00e+0 | retained | wall-associated kinase 2 |
| MD05G1236100 | LOC103427965 | Receptor-like kinase/protein (RLK/RLP) | green | 0.945 | yes | 52 | 3.09 | 4.17e-10 | 0.83 | 1.00e+0 | retained | disease resistance family protein / LRR family protein |
| MD01G1179800 | LOC103417845 | Receptor-like kinase/protein (RLK/RLP) | green | 0.975 | yes | 303 | 3.09 | 3.50e-8 | 1.46 | 1.00e+0 | retained | disease resistance family protein / LRR family protein |
| MD11G1231100 | LOC103448565 | Receptor-like kinase/protein (RLK/RLP) | green | 0.907 | yes | 1,059 | 3.08 | 3.96e-8 | 1.71 | 8.05e-1 | retained | S-locus lectin protein kinase family protein |

| Gene ID | NCBI LOC | Functional family | Module | kME | Hub | baseMean | Rin vs control log2FC | Rin vs control padj | Rin vs Rola log2FC | Rin vs Rola padj | Batch-model status | Description |
|---|---|---|---|---|---|---|---|---|---|---|---|---|
| MD11G1018800 | LOC103447088 | Receptor-like kinase/protein (RLK/RLP) | blue | 0.966 | yes | 200 | 3.06 | 8.04e-6 | -0.40 | 1.00e+0 | threshold crossing | Leucine-rich receptor-like protein kinase family protein |
| MD00G1170700 | LOC103401065 | Receptor-like kinase/protein (RLK/RLP) | blue | 0.944 | yes | 108 | 3.06 | 2.34e-6 | 0.20 | 1.00e+0 | retained | Leucine-rich repeat receptor-like protein kinase family protein |
| MD10G1312500 | LOC114827806 | Receptor-like kinase/protein (RLK/RLP) | green | 0.967 | yes | 346 | 3.05 | 1.56e-11 | 0.58 | 1.00e+0 | retained | cysteine-rich RLK (RECEPTOR-like protein kinase) 26 |
| MD01G1057400 | LOC103406479 | Receptor-like kinase/protein (RLK/RLP) | blue | 0.915 | yes | 189 | 2.97 | 1.47e-8 | 0.39 | 1.00e+0 | retained | disease resistance family protein / LRR family protein |
| MD01G1150100 | LOC103455178 | Receptor-like kinase/protein (RLK/RLP) | green | 0.783 | | 140 | 2.96 | 1.84e-13 | 0.47 | 1.00e+0 | retained | Leucine-rich repeat protein kinase family protein |
| MD06G1032200 | LOC103436877 | Receptor-like kinase/protein (RLK/RLP) | black | 0.841 | yes | 244 | 2.94 | 1.28e-9 | 3.16 | 6.50e-4 | retained | plant intracellular ras group-related LRR 4 |
| MD00G1062800 | LOC114823514 | Receptor-like kinase/protein (RLK/RLP) | blue | 0.886 | yes | 64 | 2.91 | 1.19e-6 | 0.58 | 1.00e+0 | retained | Leucine-rich repeat protein kinase family protein |
| MD05G1257600 | LOC103424618 | Receptor-like kinase/protein (RLK/RLP) | green | 0.863 | yes | 85 | 2.90 | 3.04e-10 | 2.40 | 3.49e-2 | retained | Leucine-rich repeat transmembrane protein kinase |
| MD10G1215200 | LOC103445917 | Receptor-like kinase/protein (RLK/RLP) | salmon | 0.849 | yes | 2,092 | 2.88 | 1.11e-7 | 2.53 | 5.68e-2 | retained | Leucine-rich repeat receptor-like protein kinase family protein |

| Gene ID | NCBI LOC | Functional family | Module | kME | Hub | baseMean | Rin vs control<br><br>log2FC | Rin vs control<br><br>padj | Rin vs Rola<br><br>log2FC | Rin vs Rola<br><br>padj | Batch-model status | Description |
|---|---|---|---|---|---|---|---|---|---|---|---|---|
| MD09G1145200 | LOC103443146 | Receptor-like kinase/protein (RLK/RLP) | green | 0.879 | yes | 603 | 2.81 | 3.54e-12 | 1.67 | 5.10e-1 | retained | Wall-associated kinase family protein |
| MD01G1057300 | LOC114821571 | Receptor-like kinase/protein (RLK/RLP) | blue | 0.906 | yes | 577 | 2.71 | 6.81e-8 | 0.52 | 1.00e+0 | retained | disease resistance family protein / LRR family protein |
| MD04G1233100 | LOC103434224 | Receptor-like kinase/protein (RLK/RLP) | blue | 0.947 | yes | 467 | 2.67 | 1.32e-8 | 0.07 | 1.00e+0 | retained | PR5-like receptor kinase |
| MD12G1256500 | LOC103414020 | Receptor-like kinase/protein (RLK/RLP) | blue | 0.819 | yes | 51 | 2.66 | 2.95e-9 | 0.25 | 1.00e+0 | retained | Malectin/receptor-like protein kinase family protein |
| MD12G1020000 | LOC103449405 | Receptor-like kinase/protein (RLK/RLP) | green | 0.960 | yes | 483 | 2.65 | 1.89e-8 | 1.06 | 1.00e+0 | retained | disease resistance family protein / LRR family protein |
| MD04G1177400 | LOC114824522 | Receptor-like kinase/protein (RLK/RLP) | green | 0.965 | yes | 143 | 2.65 | 1.44e-7 | 0.96 | 1.00e+0 | retained | Leucine-rich repeat protein kinase family protein |
| MD02G1107300 | LOC103442649 | Receptor-like kinase/protein (RLK/RLP) | lightcyan | 0.877 | yes | 78 | 2.62 | 6.51e-12 | 1.95 | 1.13e-1 | retained | Concanavalin A-like lectin protein kinase family protein |
| MD02G1107100 | LOC103442649 | Receptor-like kinase/protein (RLK/RLP) | lightcyan | 0.926 | yes | 288 | 2.61 | 2.60e-13 | 1.85 | 1.54e-1 | retained | Concanavalin A-like lectin protein kinase family protein |
| MD10G1334600 | LOC114827494 | Receptor-like kinase/protein (RLK/RLP) | green | 0.795 | | 229 | 2.58 | 2.68e-8 | 0.11 | 1.00e+0 | classification changed | Leucine-rich repeat transmembrane protein kinase |

| Gene ID | NCBI LOC | Functional family | Module | kME | Hub | baseMean | Rin vs control<br><br>log2FC | Rin vs control<br><br>padj | Rin vs Rola<br><br>log2FC | Rin vs Rola<br><br>padj | Batch-model status | Description |
|---|---|---|---|---|---|---|---|---|---|---|---|---|
| MD01G1172800 | LOC114824914 | Receptor-like kinase/protein (RLK/RLP) | green | 0.971 | yes | 861 | 2.49 | 1.24e-12 | 0.60 | 1.00e+0 | retained | disease resistance family protein / LRR family protein |
| MD01G1064600 | LOC103406169 | Receptor-like kinase/protein (RLK/RLP) | blue | 0.938 | yes | 495 | 2.48 | 8.34e-8 | 0.27 | 1.00e+0 | retained | disease resistance family protein / LRR family protein |
| MD05G1214700 | LOC103436186 | Receptor-like kinase/protein (RLK/RLP) | green | 0.978 | yes | 309 | 2.47 | 5.02e-11 | 1.03 | 1.00e+0 | retained | S-locus lectin protein kinase family protein |
| MD11G1268200 | LOC103452293 | Receptor-like kinase/protein (RLK/RLP) | green | 0.952 | yes | 436 | 2.44 | 9.76e-9 | 1.30 | 1.00e+0 | retained | Leucine-rich repeat transmembrane protein kinase |
| MD11G1066000 | LOC103447339 | Receptor-like kinase/protein (RLK/RLP) | midnightblue | 0.942 | yes | 331 | 2.41 | 3.06e-8 | 2.67 | 4.35e-3 | retained | Leucine-rich repeat protein kinase family protein |
| MD00G1082100 | LOC103412691 | Receptor-like kinase/protein (RLK/RLP) | blue | 0.904 | yes | 557 | 2.34 | 1.08e-7 | -0.04 | 1.00e+0 | threshold crossing | Leucine-rich receptor-like protein kinase family protein |
| MD03G1077700 | LOC103427128 | Receptor-like kinase/protein (RLK/RLP) | red | 0.841 | yes | 86 | 2.33 | 4.31e-8 | 4.04 | 3.01e-8 | retained | calmodulin-binding receptor-like cytoplasmic kinase 1 |
| MD13G1096000 | LOC103452004 | Receptor-like kinase/protein (RLK/RLP) | green | 0.897 | yes | 864 | 2.29 | 1.89e-10 | 1.40 | 9.48e-1 | retained | cysteine-rich RLK (RECEPTOR-like protein kinase) 2 |
| MD14G1186800 | LOC108172934 | Receptor-like kinase/protein (RLK/RLP) | green | 0.961 | yes | 476 | 2.27 | 1.39e-9 | 0.63 | 1.00e+0 | retained | disease resistance family protein / LRR family protein |

| Gene ID | NCBI LOC | Functional family | Module | kME | Hub | baseMean | Rin vs control<br><br>log2FC | Rin vs control<br><br>padj | Rin vs Rola<br><br>log2FC | Rin vs Rola<br><br>padj | Batch-model status | Description |
|---|---|---|---|---|---|---|---|---|---|---|---|---|
| MD01G1062000 | LOC103403871 | Receptor-like kinase/protein (RLK/RLP) | green | 0.940 | yes | 336 | 2.24 | 2.67e-10 | 0.80 | 1.00e+0 | retained | disease resistance family protein / LRR family protein |
| MD01G1062500 | LOC103417604 | Receptor-like kinase/protein (RLK/RLP) | green | 0.938 | yes | 432 | 2.23 | 3.93e-9 | 0.96 | 1.00e+0 | retained | disease resistance family protein / LRR family protein |
| MD00G1046700 | LOC103432961 | Receptor-like kinase/protein (RLK/RLP) | green | 0.824 | yes | 980 | 2.23 | 5.58e-12 | 2.37 | 1.47e-3 | retained | Leucine-rich repeat transmembrane protein kinase |
| MD05G1213900 | LOC103436186 | Receptor-like kinase/protein (RLK/RLP) | green | 0.962 | yes | 204 | 2.22 | 6.98e-9 | 0.89 | 1.00e+0 | retained | S-locus lectin protein kinase family protein |
| MD14G1186700 | LOC108172934 | Receptor-like kinase/protein (RLK/RLP) | green | 0.938 | yes | 616 | 2.21 | 7.12e-9 | 1.09 | 1.00e+0 | retained | disease resistance family protein / LRR family protein |
| MD05G1257000 | LOC103438143 | Receptor-like kinase/protein (RLK/RLP) | green | 0.803 | yes | 1,204 | 2.11 | 5.88e-9 | 2.38 | 4.85e-3 | threshold crossing | Leucine-rich repeat transmembrane protein kinase |
| MD07G1236200 | LOC103413690 | Receptor-like kinase/protein (RLK/RLP) | black | 0.884 | yes | 99 | 2.02 | 5.18e-11 | 1.98 | 2.67e-2 | retained | Leucine-rich repeat protein kinase family protein |
| MD08G1210000 | LOC103441752 | Receptor-like kinase/protein (RLK/RLP) | green | 0.885 | yes | 1,896 | 1.98 | 6.69e-12 | 0.43 | 1.00e+0 | retained | Leucine-rich receptor-like protein kinase family protein |
| MD17G1063200 | LOC103432089 | Receptor-like kinase/protein (RLK/RLP) | green | 0.940 | yes | 3,616 | 1.89 | 6.21e-12 | 0.96 | 1.00e+0 | retained | Malectin/receptor-like protein kinase family protein |

| Gene ID | NCBI LOC | Functional family | Module | kME | Hub | baseMean | Rin vs control<br><br>log2FC | Rin vs control<br><br>padj | Rin vs Rola<br><br>log2FC | Rin vs Rola<br><br>padj | Batch-model status | Description |
|---|---|---|---|---|---|---|---|---|---|---|---|---|
| MD13G1077900 | LOC103451854 | Transcription factor (WRKY/MYB/NAC/ERF/bHLH) | black | 0.895 | yes | 55 | 4.60 | 7.06e-8 | 3.72 | 2.10e-2 | retained | WRKY DNA-binding protein 72 |
| MD13G1150700 | LOC103452669 | Transcription factor (WRKY/MYB/NAC/ERF/bHLH) | green | 0.851 | yes | 211 | 4.34 | 5.29e-12 | 1.49 | 1.00e+0 | retained | WRKY DNA-binding protein 48 |
| MD08G1092000 | LOC103441079 | Transcription factor (WRKY/MYB/NAC/ERF/bHLH) | green | 0.815 | yes | 2,774 | 3.62 | 6.11e-12 | 1.22 | 1.00e+0 | retained | myb domain protein 73 |
| MD05G1295700 | LOC103434906 | Transcription factor (WRKY/MYB/NAC/ERF/bHLH) | green | 0.891 | yes | 170 | 3.30 | 7.32e-8 | 0.81 | 1.00e+0 | retained | WRKY DNA-binding protein 65 |
| MD04G1092400 | LOC103433108 | Transcription factor (WRKY/MYB/NAC/ERF/bHLH) | blue | 0.964 | yes | 58 | 3.03 | 3.67e-7 | -0.03 | 1.00e+0 | retained | myb domain protein 36 |
| MD02G1179000 | LOC103453725 | Transcription factor (WRKY/MYB/NAC/ERF/bHLH) | salmon | 0.878 | yes | 1,687 | 3.01 | 6.87e-9 | 0.85 | 1.00e+0 | retained | myb domain protein 73 |
| MD15G1288600 | LOC103401412 | Transcription factor (WRKY/MYB/NAC/ERF/bHLH) | green | 0.883 | yes | 1,934 | 2.43 | 5.04e-12 | 0.44 | 1.00e+0 | retained | myb domain protein 73 |

| Gene ID | NCBI LOC | Functional family | Module | kME | Hub | baseMean | Rin vs control log2FC | Rin vs control padj | Rin vs Rola log2FC | Rin vs Rola padj | Batch-model status | Description |
|---|---|---|---|---|---|---|---|---|---|---|---|---|
| MD01G1155000 | LOC103437419 | Transcription factor (WRKY/MYB/NAC/ERF/bHLH) | green | 0.847 | yes | 418 | 2.29 | 1.55e-9 | 0.93 | 1.00e+0 | retained | basic helix-loop-helix (bHLH) DNA-binding superfamily protein |
| MD02G1138900 | LOC103407082 | Peroxidase | midnightblue | 0.976 | yes | 114 | 7.34 | 1.47e-5 | 10.21 | 5.93e-5 | retained | Peroxidase superfamily protein |
| MD03G1014200 | LOC114823840 | Peroxidase | green | 0.873 | yes | 133 | 4.98 | 2.04e-8 | 4.25 | 6.51e-3 | retained | peroxidase 2 |
| MD17G1092400 | LOC103425937 | Peroxidase | black | 0.921 | yes | 1,399 | 3.44 | 6.05e-6 | 3.71 | 7.96e-3 | retained | Peroxidase superfamily protein |
| MD10G1042300 | LOC114827523 | Redox / GST / glutaredoxin / thioredoxin | green | 0.885 | yes | 181 | 5.16 | 4.17e-5 | 3.54 | 2.62e-1 | retained | Laccase/Diphenol oxidase family protein |
| MD12G1161000 | LOC103450543 | Redox / GST / glutaredoxin / thioredoxin | midnightblue | 0.982 | yes | 196 | 4.62 | 1.70e-9 | 5.00 | 4.48e-5 | retained | Laccase/Diphenol oxidase family protein |
| MD13G1172100 | LOC103452977 | Redox / GST / glutaredoxin / thioredoxin | black | 0.919 | yes | 114 | 4.43 | 4.85e-11 | 5.17 | 1.22e-6 | retained | Thioredoxin superfamily protein |
| MD07G1240700 | LOC103420753 | Redox / GST / glutaredoxin / thioredoxin | black | 0.430 | | 3,979 | 3.51 | 1.65e-5 | 3.88 | 8.49e-3 | threshold crossing | Fe superoxide dismutase 2 |
| MD02G1236100 | LOC103409906 | Redox / GST / glutaredoxin / thioredoxin | blue | 0.983 | yes | 760 | 3.05 | 1.19e-7 | 0.21 | 1.00e+0 | retained | glutathione S-transferase TAU 8 |
| MD12G1239800 | LOC103451027 | Redox / GST / glutaredoxin / thioredoxin | blue | 0.892 | yes | 740 | 2.82 | 5.80e-9 | 0.08 | 1.00e+0 | retained | NAD(P)-linked oxidoreductase superfamily protein |
| MD10G1172300 | LOC103445446 | Redox / GST / glutaredoxin / thioredoxin | green | 0.921 | yes | 348 | 2.62 | 1.76e-12 | 0.30 | 1.00e+0 | retained | glutathione S-transferase TAU 8 |

| Gene ID | NCBI LOC | Functional family | Module | kME | Hub | baseMean | Rin vs control log2FC | Rin vs control padj | Rin vs Rola log2FC | Rin vs Rola padj | Batch-model status | Description |
|---|---|---|---|---|---|---|---|---|---|---|---|---|
| MD09G1074300 | LOC103442449 | Redox / GST / glutaredoxin / thioredoxin | green | 0.831 | yes | 60 | 2.52 | 5.44e-8 | 0.44 | 1.00e+0 | retained | Thioredoxin superfamily protein |
| MD17G1250000 | LOC103426173 | PR / thaumatin / osmotin | green | 0.952 | yes | 2,690 | 5.29 | 1.15e-9 | 1.24 | 1.00e+0 | retained | Pathogenesis-related thaumatin superfamily protein |
| MD04G1064400 | LOC103433014 | PR / thaumatin / osmotin | green | 0.899 | yes | 265 | 4.51 | 8.02e-10 | 1.18 | 1.00e+0 | retained | osmotin 34 |
| MD10G1017300 | LOC103436524 | PR / thaumatin / osmotin | green | 0.856 | yes | 772 | 2.97 | 4.56e-8 | 0.42 | 1.00e+0 | retained | pathogenesis-related family protein |
| MD11G1219900 | LOC103428490 | Cytochrome P450 | green | 0.907 | yes | 249 | 5.44 | 2.14e-11 | 1.68 | 1.00e+0 | retained | Cytochrome P450 superfamily protein |
| MD16G1116100 | LOC103416681 | Cytochrome P450 | salmon | 0.921 | yes | 806 | 5.41 | 3.35e-9 | 5.87 | 2.56e-5 | retained | cytochrome P450 |
| MD16G1285900 | LOC103444865 | Cytochrome P450 | salmon | 0.933 | yes | 103 | 4.60 | 9.07e-6 | 4.10 | 3.46e-2 | retained | cytochrome P450 |
| MD15G1028700 | LOC103450491 | Cytochrome P450 | green | 0.866 | yes | 311 | 3.19 | 1.85e-7 | 2.40 | 1.68e-1 | retained | cytochrome P450 |
| MD11G1220600 | LOC103422998 | Cytochrome P450 | green | 0.964 | yes | 404 | 3.10 | 2.45e-8 | 1.07 | 1.00e+0 | retained | Cytochrome P450 superfamily protein |
| MD13G1033900 | LOC103402649 | Cytochrome P450 | green | 0.904 | yes | 64 | 2.80 | 1.92e-8 | 0.98 | 1.00e+0 | retained | Cytochrome P450 superfamily protein |
| MD01G1172500 | LOC103406468 | Cytochrome P450 | black | 0.939 | yes | 103 | 2.79 | 1.61e-9 | 3.24 | 1.98e-4 | retained | cytochrome P450 |
| MD13G1103500 | LOC103430668 | Cytochrome P450 | black | 0.942 | yes | 759 | 2.65 | 4.11e-10 | 2.93 | 4.98e-4 | retained | cytochrome P450 |

| Gene ID | NCBI LOC | Functional family | Module | kME | Hub | baseMean | Rin vs control<br><br>log2FC | Rin vs control<br><br>padj | Rin vs Rola<br><br>log2FC | Rin vs Rola<br><br>padj | Batch-model status | Description |
|---|---|---|---|---|---|---|---|---|---|---|---|---|
| MD15G1033400 | LOC103441496 | Cytochrome P450 | black | 0.900 | yes | 1,178 | 2.62 | 6.17e-12 | 1.58 | 5.95e-1 | retained | cytochrome P450 |
| MD13G1103800 | LOC103452391 | Cytochrome P450 | black | 0.878 | yes | 81 | 2.52 | 1.34e-8 | 3.03 | 5.59e-4 | retained | cytochrome P450 |
| MD15G1033900 | LOC103441496 | Cytochrome P450 | green | 0.894 | yes | 1,076 | 2.37 | 2.98e-18 | 0.54 | 1.00e+0 | retained | cytochrome P450 |
| MD07G1006100 | LOC103409912 | Cytochrome P450 | green | 0.934 | yes | 548 | 2.36 | 2.10e-9 | 0.16 | 1.00e+0 | retained | cytochrome P450 |
| MD04G1066300 | LOC103427519 | Cytochrome P450 | black | 0.925 | yes | 520 | 2.35 | 2.05e-10 | 2.49 | 2.60e-3 | retained | cytochrome P450 |
| MD06G1164000 | LOC103411574 | Cytochrome P450 | green | 0.905 | yes | 661 | 2.32 | 3.13e-10 | 1.31 | 1.00e+0 | retained | cytochrome P450 |
| MD07G1220800 | LOC103432823 | Cytochrome P450 | green | 0.762 | | 638 | 1.90 | 4.20e-23 | 0.80 | 1.00e+0 | retained | Cytochrome P450 superfamily protein |
| MD16G1091200 | LOC103403007 | Cell wall / pectin / XTH / expansin | red | 0.831 | yes | 1,546 | 5.16 | 2.61e-14 | 6.48 | 2.47e-11 | retained | xyloglucan:xyloglucosyl transferase 33 |
| MD00G1105300 | LOC103439438 | Cell wall / pectin / XTH / expansin | blue | 0.639 | | 1,780 | 3.93 | 1.27e-6 | 1.34 | 1.00e+0 | retained | Plant invertase/pectin methylesterase inhibitor superfamily protein |
| MD13G1237300 | LOC103445478 | Cell wall / pectin / XTH / expansin | green | 0.838 | yes | 390 | 3.86 | 2.59e-5 | 1.62 | 1.00e+0 | not evaluable | xyloglucan endotransglycosylase 6 |
| MD16G1014000 | LOC103429442 | Cell wall / pectin / XTH / expansin | black | 0.868 | yes | 5,992 | 2.48 | 3.76e-10 | 2.37 | 1.32e-2 | retained | xyloglucan endotransglucosylase/hydrolase 28 |
| MD06G1027300 | LOC103436866 | Cell wall / pectin / XTH / expansin | black | 0.912 | yes | 679 | 2.44 | 1.21e-7 | 2.15 | 1.17e-1 | retained | xyloglucan endotransglucosylase/hydrolase 30 |

| Gene ID | NCBI LOC | Functional family | Module | kME | Hub | baseMean | Rin vs control log2FC | Rin vs control padj | Rin vs Rola log2FC | Rin vs Rola padj | Batch-model status | Description |
|---|---|---|---|---|---|---|---|---|---|---|---|---|
| MD11G1054100 | LOC103422644 | Lipid / cuticle / GDSL / LTP | black | 0.958 | yes | 10,183 | 4.69 | 2.04e-6 | 4.65 | 5.66e-3 | retained | lipid transfer protein 1 |
| MD11G1075900 | LOC103412826 | Lipid / cuticle / GDSL / LTP | green | 0.943 | yes | 121 | 3.35 | 8.83e-10 | 1.30 | 1.00e+0 | retained | non-specific phospholipase C1 |
| MD17G1014200 | LOC103404209 | Calcium signaling | blue | 0.899 | yes | 89 | 3.71 | 8.75e-6 | 0.71 | 1.00e+0 | retained | calmodulin-like 38 |
| MD07G1164200 | LOC103439466 | Calcium signaling | salmon | 0.928 | yes | 1,439 | 3.24 | 6.20e-7 | 2.31 | 2.84e-1 | retained | Calcium-binding endonuclease/exonuclease/phosphatase family |
| MD12G1075400 | LOC103423272 | UDP-glycosyltransferase | green | 0.860 | yes | 84 | 8.30 | 9.19e-17 | 6.23 | 1.32e-6 | retained | Glycosyltransferase family 61 protein |
| MD09G1142500 | LOC103410840 | UDP-glycosyltransferase | blue | 0.962 | yes | 241 | 5.68 | 1.70e-7 | 0.87 | 1.00e+0 | classification changed | UDP-Glycosyltransferase superfamily protein |
| MD17G1126300 | LOC103405101 | UDP-glycosyltransferase | black | 0.967 | yes | 239 | 3.33 | 2.00e-8 | 3.26 | 4.67e-3 | retained | UDP-Glycosyltransferase superfamily protein |
| MD16G1016800 | LOC103444588 | UDP-glycosyltransferase | lightcyan | 0.933 | yes | 2,095 | 2.52 | 1.04e-7 | 0.95 | 1.00e+0 | retained | UDP-Glycosyltransferase / trehalose-phosphatase family protein |
| MD09G1064700 | LOC103429051 | UDP-glycosyltransferase | lightcyan | 0.943 | yes | 134 | 2.43 | 1.06e-10 | 1.70 | 3.58e-1 | retained | UDP-Glycosyltransferase superfamily protein |
| MD08G1048100 | LOC103413415 | Unclassified | blue | 0.884 | yes | 51 | 5.52 | 4.95e-7 | 1.64 | 1.00e+0 | retained | glutamate receptor 2.2 |
| MD10G1250900 | LOC103422333 | Unclassified | green | 0.872 | yes | 790 | 5.22 | 8.68e-22 | 3.87 | 4.40e-5 | retained | wall associated kinase 5 |

| Gene ID | NCBI LOC | Functional family | Module | kME | Hub | baseMean | Rin vs control<br><br>log2FC | Rin vs control<br><br>padj | Rin vs Rola<br><br>log2FC | Rin vs Rola<br><br>padj | Batch-model status | Description |
|---|---|---|---|---|---|---|---|---|---|---|---|---|
| MD15G1218000 | LOC103400938 | Unclassified | green | 0.873 | yes | 129 | 5.20 | 3.14e-9 | 1.66 | 1.00e+0 | not evaluable | Protein kinase superfamily protein |
| MD15G1046200 | LOC103418317 | Unclassified | green | 0.925 | yes | 113 | 4.76 | 1.07e-12 | 1.69 | 9.97e-1 | retained | glutamate receptor 2.8 |
| MD02G1091900 | LOC114819647 | Unclassified | green | 0.788 | | 1,233 | 4.52 | 1.01e-11 | 1.45 | 1.00e+0 | retained | Protein kinase superfamily protein |
| MD10G1250000 | LOC108171771 | Unclassified | green | 0.912 | yes | 133 | 4.38 | 1.12e-8 | 2.45 | 3.34e-1 | retained | wall associated kinase 5 |
| MD01G1213000 | LOC103439830 | Unclassified | green | 0.924 | yes | 209 | 3.95 | 1.52e-8 | 0.95 | 1.00e+0 | retained | plastidic pyruvate kinase beta subunit 1 |
| MD09G1130600 | LOC103443070 | Unclassified | green | 0.954 | yes | 321 | 3.94 | 8.75e-9 | 1.96 | 6.76e-1 | retained | receptor like protein 6 |
| MD01G1059700 | LOC103407534 | Unclassified | green | 0.950 | yes | 1,463 | 3.60 | 7.96e-13 | 1.21 | 1.00e+0 | retained | Protein kinase superfamily protein |
| MD09G1255200 | LOC103405757 | Unclassified | green | 0.909 | yes | 57 | 3.41 | 3.83e-7 | 0.77 | 1.00e+0 | retained | Protein kinase superfamily protein |
| MD03G1192900 | LOC103448830 | Unclassified | green | 0.878 | yes | 64 | 3.33 | 3.00e-6 | 1.57 | 1.00e+0 | retained | receptor like protein 2 |
| MD06G1232600 | LOC103438110 | Unclassified | green | 0.803 | yes | 792 | 3.25 | 1.67e-17 | 0.89 | 1.00e+0 | retained | MAP kinase kinase 9 |
| MD05G1305700 | LOC103408934 | Unclassified | green | 0.940 | yes | 523 | 3.06 | 2.78e-6 | 1.08 | 1.00e+0 | retained | glutamate receptor 2.7 |
| MD10G1226100 | LOC103445978 | Unclassified | green | 0.956 | yes | 3,710 | 2.76 | 1.84e-13 | 1.33 | 1.00e+0 | retained | VACUOLAR SORTING RECEPTOR 6 |
| MD04G1233200 | LOC103434225 | Unclassified | blue | 0.989 | yes | 866 | 2.59 | 2.85e-7 | -0.11 | 1.00e+0 | retained | Protein kinase superfamily protein |

| Gene ID | NCBI LOC | Functional family | Module | kME | Hub | baseMean | Rin vs control log2FC | Rin vs control padj | Rin vs Rola log2FC | Rin vs Rola padj | Batch-model status | Description |
|---|---|---|---|---|---|---|---|---|---|---|---|---|
| MD09G1070100 | LOC103442602 | Unclassified | green | 0.964 | yes | 1,138 | 2.53 | 1.46e-7 | 0.86 | 1.00e+0 | retained | glutamate receptor 3.4 |
| MD15G1146100 | LOC103456338 | Unclassified | green | 0.904 | yes | 179 | 2.41 | 2.19e-11 | 1.30 | 1.00e+0 | retained | Integrin-linked protein kinase family |
| MD13G1203800 | LOC103401970 | Unclassified | blue | 0.973 | yes | 885 | 2.40 | 2.11e-7 | 0.03 | 1.00e+0 | threshold crossing | Protein kinase superfamily protein |
| MD10G1249500 | LOC103446854 | Unclassified | green | 0.936 | yes | 390 | 2.33 | 3.43e-13 | 1.45 | 7.04e-1 | retained | wall associated kinase 5 |
| MD07G1249700 | LOC103439597 | Unclassified | green | 0.834 | yes | 1,170 | 2.27 | 1.30e-14 | 0.67 | 1.00e+0 | retained | U-box domain-containing protein kinase family protein |
| MD17G1034800 | LOC103404344 | Unclassified | green | 0.903 | yes | 2,315 | 1.53 | 8.32e-70 | 0.67 | 1.00e+0 | retained | formate dehydrogenase |
