## Supplementary tables and figures for "Transcriptome of apple cv. Ametyst in *Rvi6*-effective and *Rvi6*-breaking *Venturia inaequalis* interactions identifies defense candidates": 10_SupplementaryTable_S3_green_module.pdf

**Supplementary Table S3. Green-module genes among the Rin-induced defense-related candidate genes.**

The 92 candidates from Supplementary Table S2 assigned to the green WGCNA co-expression module — the module in which the candidate genes are most strongly concentrated — ranked by module membership (kME). kME P is the *P* value of the module-membership correlation; Hub marks genes meeting the OmicsBox hub-tag criterion, kME ≥ 0.8 and module-membership *P* < 0.05 (88 of 92); r (Rin) is the gene–trait correlation with the Rin treatment. Statistics are given for both the Rin vs control and Rin vs Rola contrasts, as defined for Supplementary Table S2. Positive log2FC denotes higher expression in Rin. Genes with high module membership — including PR5-like and wall-associated receptor kinases, S-locus lectin kinases, and NBS-LRR/LRR resistance proteins — represent prioritized co-expression candidates.

| Gene ID | NCBI LOC | kME | kME P | Hub | r (Rin) | baseMean | Rin vs control | | Rin vs Rola | | Description |
|---|---|---|---|---|---|---|---|---|---|---|---|
| | | | | | | | log2FC | padj | log2FC | padj | |
| MD02G1251800 | LOC103408537 | 0.989 | 3.04e-17 | yes | 0.70 | 101 | 4.02 | 1.18e-12 | 1.32 | 1.00e+00 | PR5-like receptor kinase |
| MD02G1250800 | LOC103408537 | 0.982 | 3.46e-15 | yes | 0.64 | 169 | 4.14 | 1.13e-09 | 1.25 | 1.00e+00 | receptor serine/threonine kinase |
| MD08G1107900 | LOC103428814 | 0.978 | 1.78e-14 | yes | 0.79 | 783 | 3.49 | 6.54e-18 | 1.69 | 4.88e-01 | wall-associated kinase 2 |
| MD05G1214700 | LOC103436186 | 0.978 | 2.45e-14 | yes | 0.71 | 309 | 2.47 | 5.02e-11 | 1.03 | 1.00e+00 | S-locus lectin protein kinase family protein |
| MD10G1251200 | LOC103446854 | 0.977 | 2.71e-14 | yes | 0.73 | 260 | 3.79 | 2.60e-11 | 1.57 | 1.00e+00 | wall-associated kinase 2 |
| MD17G1062200 | LOC103409759 | 0.977 | 3.16e-14 | yes | 0.62 | 69 | 3.86 | 4.30e-11 | 0.72 | 1.00e+00 | Malectin/receptor-like protein kinase family protein |
| MD10G1251400 | LOC103422333 | 0.975 | 6.00e-14 | yes | 0.74 | 245 | 3.10 | 2.02e-11 | 1.33 | 1.00e+00 | wall-associated kinase 2 |
| MD01G1179800 | LOC103417845 | 0.975 | 6.23e-14 | yes | 0.69 | 303 | 3.09 | 3.50e-08 | 1.46 | 1.00e+00 | disease resistance family protein / LRR family protein |
| MD01G1172800 | LOC114824914 | 0.971 | 2.85e-13 | yes | 0.62 | 861 | 2.49 | 1.24e-12 | 0.60 | 1.00e+00 | disease resistance family protein / LRR family protein |
| MD10G1250500 | LOC103422333 | 0.971 | 3.08e-13 | yes | 0.71 | 277 | 4.29 | 7.03e-11 | 1.99 | 6.03e-01 | wall-associated kinase 2 |
| MD04G1011700 | LOC103429293 | 0.969 | 5.99e-13 | yes | 0.76 | 281 | 4.45 | 2.82e-17 | 2.12 | 2.45e-01 | Disease resistance protein (TIR-NBS-LRR class) family |
| MD10G1214300 | LOC103445917 | 0.968 | 6.70e-13 | yes | 0.60 | 316 | 3.30 | 3.84e-11 | 0.97 | 1.00e+00 | Leucine-rich repeat receptor-like protein kinase family protein |
| MD10G1312500 | LOC114827806 | 0.967 | 8.69e-13 | yes | 0.56 | 346 | 3.05 | 1.56e-11 | 0.58 | 1.00e+00 | cysteine-rich RLK (RECEPTOR-like protein kinase) 26 |
| MD04G1177400 | LOC114824522 | 0.965 | 1.67e-12 | yes | 0.66 | 143 | 2.65 | 1.44e-07 | 0.96 | 1.00e+00 | Leucine-rich repeat protein kinase family protein |
| MD11G1220600 | LOC103422998 | 0.964 | 2.08e-12 | yes | 0.59 | 404 | 3.10 | 2.45e-08 | 1.07 | 1.00e+00 | Cytochrome P450 superfamily protein |
| MD09G1070100 | LOC103442602 | 0.964 | 2.15e-12 | yes | 0.56 | 1138 | 2.53 | 1.46e-07 | 0.86 | 1.00e+00 | glutamate receptor 3.4 |
| MD10G1307800 | LOC103446652 | 0.964 | 2.20e-12 | yes | 0.70 | 192 | 3.31 | 1.47e-08 | 1.40 | 1.00e+00 | S-locus lectin protein kinase family protein |

| Gene ID | NCBI LOC | kME | kME P | Hub | r (Rin) | baseMean | Rin vs control | | Rin vs Rola | | Description |
|---|---|---|---|---|---|---|---|---|---|---|---|
| MD15G1426700 | LOC103417504 | 0.964 | 2.27e-12 | yes | 0.52 | 2705 | 4.14 | 3.56e-09 | 1.16 | 1.00e+00 | disease resistance family protein / LRR family protein |
| MD04G1020400 | LOC103402536 | 0.964 | 2.35e-12 | yes | 0.59 | 339 | 4.94 | 3.01e-12 | 1.57 | 1.00e+00 | disease resistance family protein / LRR family protein |
| MD02G1253600 | LOC108170170 | 0.963 | 2.85e-12 | yes | 0.61 | 447 | 3.80 | 5.12e-07 | 1.20 | 1.00e+00 | receptor serine/threonine kinase |
| MD01G1178700 | LOC103415095 | 0.963 | 3.07e-12 | yes | 0.59 | 187 | 3.20 | 2.57e-08 | 1.25 | 1.00e+00 | disease resistance family protein / LRR family protein |
| MD05G1213900 | LOC103436186 | 0.962 | 3.52e-12 | yes | 0.64 | 204 | 2.22 | 6.98e-09 | 0.89 | 1.00e+00 | S-locus lectin protein kinase family protein |
| MD14G1186800 | LOC108172934 | 0.961 | 4.60e-12 | yes | 0.67 | 476 | 2.27 | 1.39e-09 | 0.63 | 1.00e+00 | disease resistance family protein / LRR family protein |
| MD12G1020000 | LOC103449405 | 0.960 | 5.52e-12 | yes | 0.64 | 483 | 2.65 | 1.89e-08 | 1.06 | 1.00e+00 | disease resistance family protein / LRR family protein |
| MD10G1226100 | LOC103445978 | 0.956 | 1.49e-11 | yes | 0.78 | 3710 | 2.76 | 1.84e-13 | 1.33 | 1.00e+00 | VACUOLAR SORTING RECEPTOR 6 |
| MD05G1207400 | LOC114825146 | 0.956 | 1.58e-11 | yes | 0.55 | 165 | 5.78 | 1.01e-10 | 1.76 | 1.00e+00 | NB-ARC domain-containing disease resistance protein |
| MD10G1248500 | LOC103446854 | 0.956 | 1.58e-11 | yes | 0.72 | 87 | 4.05 | 5.27e-09 | 2.01 | 6.14e-01 | wall-associated kinase 2 |
| MD00G1101700 | LOC103419471 | 0.954 | 2.02e-11 | yes | 0.82 | 249 | 4.40 | 8.40e-13 | 2.40 | 1.80e-01 | cysteine-rich RLK (RECEPTOR-like protein kinase) 26 |
| MD09G1130600 | LOC103443070 | 0.954 | 2.29e-11 | yes | 0.66 | 321 | 3.94 | 8.75e-09 | 1.96 | 6.76e-01 | receptor like protein 6 |
| MD17G1250000 | LOC103426173 | 0.952 | 3.01e-11 | yes | 0.54 | 2690 | 5.29 | 1.15e-09 | 1.24 | 1.00e+00 | Pathogenesis-related thaumatin superfamily protein |
| MD11G1268200 | LOC103452293 | 0.952 | 3.25e-11 | yes | 0.71 | 436 | 2.44 | 9.76e-09 | 1.30 | 1.00e+00 | Leucine-rich repeat transmembrane protein kinase |
| MD01G1059700 | LOC103407534 | 0.950 | 4.44e-11 | yes | 0.59 | 1463 | 3.60 | 7.96e-13 | 1.21 | 1.00e+00 | Protein kinase superfamily protein |
| MD05G1207800 | LOC114825146 | 0.947 | 7.80e-11 | yes | 0.56 | 106 | 6.05 | 5.92e-12 | 1.87 | 1.00e+00 | NB-ARC domain-containing disease resistance protein |
| MD05G1217100 | LOC103428093 | 0.946 | 9.47e-11 | yes | 0.74 | 64 | 3.16 | 6.32e-13 | 1.23 | 1.00e+00 | S-locus lectin protein kinase family protein |
| MD05G1236100 | LOC103427965 | 0.945 | 1.21e-10 | yes | 0.68 | 52 | 3.09 | 4.17e-10 | 0.83 | 1.00e+00 | disease resistance family protein / LRR family protein |
| MD11G1075900 | LOC103412826 | 0.943 | 1.68e-10 | yes | 0.60 | 121 | 3.35 | 8.83e-10 | 1.30 | 1.00e+00 | non-specific phospholipase C1 |
| MD01G1062000 | LOC103403871 | 0.940 | 2.37e-10 | yes | 0.68 | 336 | 2.24 | 2.67e-10 | 0.80 | 1.00e+00 | disease resistance family protein / LRR family protein |
| MD17G1063200 | LOC103432089 | 0.940 | 2.51e-10 | yes | 0.76 | 3616 | 1.89 | 6.21e-12 | 0.96 | 1.00e+00 | Malectin/receptor-like protein kinase family protein |

| Gene ID | NCBI LOC | kME | kME P | Hub | r (Rin) | baseMean | Rin vs control | | Rin vs Rola | | Description |
|---|---|---|---|---|---|---|---|---|---|---|---|
| MD05G1305700 | LOC103408934 | 0.940 | 2.72e-10 | yes | 0.51 | 523 | 3.06 | 2.78e-06 | 1.08 | 1.00e+00 | glutamate receptor 2.7 |
| MD01G1062500 | LOC103417604 | 0.938 | 3.39e-10 | yes | 0.70 | 432 | 2.23 | 3.93e-09 | 0.96 | 1.00e+00 | disease resistance family protein / LRR family protein |
| MD14G1186700 | LOC108172934 | 0.938 | 3.66e-10 | yes | 0.74 | 616 | 2.21 | 7.12e-09 | 1.09 | 1.00e+00 | disease resistance family protein / LRR family protein |
| MD10G1249500 | LOC103446854 | 0.936 | 4.83e-10 | yes | 0.83 | 390 | 2.33 | 3.43e-13 | 1.45 | 7.04e-01 | wall associated kinase 5 |
| MD07G1006100 | LOC103409912 | 0.934 | 6.45e-10 | yes | 0.48 | 548 | 2.36 | 2.10e-09 | 0.16 | 1.00e+00 | cytochrome P450 |
| MD15G1046200 | LOC103418317 | 0.925 | 1.93e-09 | yes | 0.59 | 113 | 4.76 | 1.07e-12 | 1.69 | 9.97e-01 | glutamate receptor 2.8 |
| MD05G1190200 | LOC103409199 | 0.925 | 1.98e-09 | yes | 0.67 | 109 | 3.87 | 3.20e-07 | 2.24 | 4.90e-01 | Leucine-rich repeat receptor-like protein kinase family protein |
| MD01G1213000 | LOC103439830 | 0.924 | 2.19e-09 | yes | 0.56 | 209 | 3.95 | 1.52e-08 | 0.95 | 1.00e+00 | plastidic pyruvate kinase beta subunit 1 |
| MD10G1172300 | LOC103445446 | 0.921 | 3.36e-09 | yes | 0.56 | 348 | 2.62 | 1.76e-12 | 0.30 | 1.00e+00 | glutathione S-transferase TAU 8 |
| MD03G1246600 | LOC103452293 | 0.917 | 4.95e-09 | yes | 0.43 | 351 | 3.33 | 1.25e-06 | 0.63 | 1.00e+00 | Leucine-rich repeat transmembrane protein kinase |
| MD10G1306800 | LOC103446654 | 0.915 | 6.53e-09 | yes | 0.54 | 57 | 4.14 | 1.50e-09 | 1.07 | 1.00e+00 | S-locus lectin protein kinase family protein |
| MD10G1250000 | LOC108171771 | 0.912 | 8.63e-09 | yes | 0.80 | 133 | 4.38 | 1.12e-08 | 2.45 | 3.34e-01 | wall associated kinase 5 |
| MD09G1255200 | LOC103405757 | 0.909 | 1.23e-08 | yes | 0.46 | 57 | 3.41 | 3.83e-07 | 0.77 | 1.00e+00 | Protein kinase superfamily protein |
| MD11G1231100 | LOC103448565 | 0.907 | 1.45e-08 | yes | 0.71 | 1059 | 3.08 | 3.96e-08 | 1.71 | 8.05e-01 | S-locus lectin protein kinase family protein |
| MD10G1212700 | LOC108172485 | 0.907 | 1.48e-08 | yes | 0.83 | 1042 | 4.16 | 1.16e-09 | 3.36 | 1.33e-02 | Leucine-rich repeat protein kinase family protein |
| MD11G1219900 | LOC103428490 | 0.907 | 1.50e-08 | yes | 0.55 | 249 | 5.44 | 2.14e-11 | 1.68 | 1.00e+00 | Cytochrome P450 superfamily protein |
| MD06G1164000 | LOC103411574 | 0.905 | 1.73e-08 | yes | 0.80 | 661 | 2.32 | 3.13e-10 | 1.31 | 1.00e+00 | cytochrome P450 |
| MD15G1146100 | LOC103456338 | 0.904 | 1.84e-08 | yes | 0.81 | 179 | 2.41 | 2.19e-11 | 1.30 | 1.00e+00 | Integrin-linked protein kinase family |
| MD13G1033900 | LOC103402649 | 0.904 | 1.88e-08 | yes | 0.60 | 64 | 2.80 | 1.92e-08 | 0.98 | 1.00e+00 | Cytochrome P450 superfamily protein |
| MD17G1034800 | LOC103404344 | 0.903 | 2.15e-08 | yes | 0.85 | 2315 | 1.53 | 8.32e-70 | 0.67 | 1.00e+00 | formate dehydrogenase |
| MD04G1064400 | LOC103433014 | 0.899 | 3.07e-08 | yes | 0.52 | 265 | 4.51 | 8.02e-10 | 1.18 | 1.00e+00 | osmotin 34 |
| MD13G1096000 | LOC103452004 | 0.897 | 3.66e-08 | yes | 0.79 | 864 | 2.29 | 1.89e-10 | 1.40 | 9.48e-01 | cysteine-rich RLK (RECEPTOR-like protein kinase) 2 |
| MD15G1033900 | LOC103441496 | 0.894 | 4.75e-08 | yes | 0.69 | 1076 | 2.37 | 2.98e-18 | 0.54 | 1.00e+00 | cytochrome P450 |
| MD05G1295700 | LOC103434906 | 0.891 | 5.97e-08 | yes | 0.49 | 170 | 3.30 | 7.32e-08 | 0.81 | 1.00e+00 | WRKY DNA-binding protein 65 |

| Gene ID | NCBI LOC | kME | kME P | Hub | r (Rin) | baseMean | Rin vs control | | Rin vs Rola | | Description |
|---|---|---|---|---|---|---|---|---|---|---|---|
| MD10G1042300 | LOC114827523 | 0.885 | 9.92e-08 | yes | 0.60 | 181 | 5.16 | 4.17e-05 | 3.54 | 2.62e-01 | Laccase/Diphenol oxidase family protein |
| MD08G1210000 | LOC103441752 | 0.885 | 1.03e-07 | yes | 0.62 | 1896 | 1.98 | 6.69e-12 | 0.43 | 1.00e+00 | Leucine-rich receptor-like protein kinase family protein |
| MD00G1183000 | LOC103431980 | 0.883 | 1.16e-07 | yes | 0.90 | 784 | 3.41 | 4.85e-18 | 2.85 | 3.48e-04 | Leucine-rich repeat transmembrane protein kinase |
| MD15G1288600 | LOC103401412 | 0.883 | 1.19e-07 | yes | 0.61 | 1934 | 2.43 | 5.04e-12 | 0.44 | 1.00e+00 | myb domain protein 73 |
| MD15G1239400 | LOC103431980 | 0.882 | 1.24e-07 | yes | 0.90 | 754 | 3.90 | 2.57e-16 | 3.25 | 3.15e-04 | Leucine-rich repeat transmembrane protein kinase |
| MD09G1145200 | LOC103443146 | 0.879 | 1.61e-07 | yes | 0.80 | 603 | 2.81 | 3.54e-12 | 1.67 | 5.10e-01 | Wall-associated kinase family protein |
| MD03G1192900 | LOC103448830 | 0.878 | 1.73e-07 | yes | 0.51 | 64 | 3.33 | 3.00e-06 | 1.57 | 1.00e+00 | receptor like protein 2 |
| MD02G1235600 | LOC114821487 | 0.877 | 1.84e-07 | yes | 0.57 | 87 | 3.76 | 6.67e-08 | 1.85 | 8.15e-01 | PR5-like receptor kinase |
| MD15G1239300 | LOC103431980 | 0.875 | 2.14e-07 | yes | 0.87 | 88 | 3.70 | 2.17e-11 | 2.98 | 8.54e-03 | Leucine-rich repeat transmembrane protein kinase |
| MD15G1218000 | LOC103400938 | 0.873 | 2.43e-07 | yes | 0.47 | 129 | 5.20 | 3.14e-09 | 1.66 | 1.00e+00 | Protein kinase superfamily protein |
| MD03G1014200 | LOC114823840 | 0.873 | 2.43e-07 | yes | 0.83 | 133 | 4.98 | 2.04e-08 | 4.25 | 6.51e-03 | peroxidase 2 |
| MD10G1250900 | LOC103422333 | 0.872 | 2.60e-07 | yes | 0.92 | 790 | 5.22 | 8.68e-22 | 3.87 | 4.40e-05 | wall associated kinase 5 |
| MD15G1239500 | LOC103424618 | 0.869 | 3.14e-07 | yes | 0.86 | 225 | 4.04 | 8.72e-10 | 3.33 | 9.85e-03 | Leucine-rich repeat transmembrane protein kinase |
| MD15G1028700 | LOC103450491 | 0.866 | 3.94e-07 | yes | 0.65 | 311 | 3.19 | 1.85e-07 | 2.40 | 1.68e-01 | cytochrome P450 |
| MD05G1257600 | LOC103424618 | 0.863 | 4.88e-07 | yes | 0.82 | 85 | 2.90 | 3.04e-10 | 2.40 | 3.49e-02 | Leucine-rich repeat transmembrane protein kinase |
| MD12G1075400 | LOC103423272 | 0.860 | 5.63e-07 | yes | 0.86 | 84 | 8.30 | 9.19e-17 | 6.23 | 1.32e-06 | Glycosyltransferase family 61 protein |
| MD10G1017300 | LOC103436524 | 0.856 | 7.27e-07 | yes | 0.57 | 772 | 2.97 | 4.56e-08 | 0.42 | 1.00e+00 | pathogenesis-related family protein |
| MD13G1150700 | LOC103452669 | 0.851 | 1.00e-06 | yes | 0.61 | 211 | 4.34 | 5.29e-12 | 1.49 | 1.00e+00 | WRKY DNA-binding protein 48 |
| MD01G1155000 | LOC103437419 | 0.847 | 1.28e-06 | yes | 0.66 | 418 | 2.29 | 1.55e-09 | 0.93 | 1.00e+00 | basic helix-loop-helix (bHLH) DNA-binding superfamily protein |
| MD13G1237300 | LOC103445478 | 0.838 | 2.13e-06 | yes | 0.36 | 390 | 3.86 | 2.59e-05 | 1.62 | 1.00e+00 | xyloglucan endotransglycosylase 6 |
| MD07G1249700 | LOC103439597 | 0.834 | 2.70e-06 | yes | 0.67 | 1170 | 2.27 | 1.30e-14 | 0.67 | 1.00e+00 | U-box domain-containing protein kinase family protein |
| MD09G1074300 | LOC103442449 | 0.831 | 3.08e-06 | yes | 0.48 | 60 | 2.52 | 5.44e-08 | 0.44 | 1.00e+00 | Thioredoxin superfamily protein |
| MD00G1046700 | LOC103432961 | 0.824 | 4.44e-06 | yes | 0.88 | 980 | 2.23 | 5.58e-12 | 2.37 | 1.47e-03 | Leucine-rich repeat transmembrane protein kinase |

| Gene ID | NCBI LOC | kME | kME P | Hub | r (Rin) | baseMean | Rin vs control | | Rin vs Rola | | Description |
|---|---|---|---|---|---|---|---|---|---|---|---|
| MD08G1092000 | LOC103441079 | 0.815 | 6.96e-06 | yes | 0.59 | 2774 | 3.62 | 6.11e-12 | 1.22 | 1.00e+00 | myb domain protein 73 |
| MD06G1232600 | LOC103438110 | 0.803 | 1.16e-05 | yes | 0.64 | 792 | 3.25 | 1.67e-17 | 0.89 | 1.00e+00 | MAP kinase kinase 9 |
| MD05G1257000 | LOC103438143 | 0.803 | 1.17e-05 | yes | 0.83 | 1204 | 2.11 | 5.88e-09 | 2.38 | 4.85e-03 | Leucine-rich repeat transmembrane protein kinase |
| MD10G1334600 | LOC114827494 | 0.795 | 1.63e-05 | | 0.51 | 229 | 2.58 | 2.68e-08 | 0.11 | 1.00e+00 | Leucine-rich repeat transmembrane protein kinase |
| MD02G1091900 | LOC114819647 | 0.788 | 2.23e-05 | | 0.58 | 1233 | 4.52 | 1.01e-11 | 1.45 | 1.00e+00 | Protein kinase superfamily protein |
| MD01G1150100 | LOC103455178 | 0.783 | 2.72e-05 | | 0.61 | 140 | 2.96 | 1.84e-13 | 0.47 | 1.00e+00 | Leucine-rich repeat protein kinase family protein |
| MD07G1220800 | LOC103432823 | 0.762 | 6.03e-05 | | 0.80 | 638 | 1.90 | 4.20e-23 | 0.80 | 1.00e+00 | Cytochrome P450 superfamily protein |
