## Supplementary tables and figures for "Transcriptome of apple cv. Ametyst in *Rvi6*-effective and *Rvi6*-breaking *Venturia inaequalis* interactions identifies defense candidates": 11_SupplementaryTable_S4_GO_terms.pdf

**Supplementary Table S4. Gene Ontology terms tested by GSEA for the Rin vs Rola contrast (GDDH13 reference-based analysis).**

Complete GSEA output for all 673 Gene Ontology terms tested, grouped by enrichment outcome (Rin-enriched, Rola-enriched, not significant) and ranked within each group by normalized enrichment score (NES). Genes were ranked by the DESeq2 Wald statistic for the Rin vs Rola contrast, so a positive NES denotes enrichment among genes expressed more highly in the Rin interaction and a negative NES enrichment among genes expressed more highly in Rola. Gene-set size was restricted to 15–500 genes. Of the 673 terms, 314 passed FDR q < 0.05 (162 Rin-enriched, 152 Rola-enriched); terms not reaching this threshold are listed in the final block for completeness. ES, enrichment score; NES, normalized enrichment score; FDR q, false discovery rate q-value; Leading edge, the tags/list/signal percentages reported by GSEA. Curated representative terms, collapsed to one per biological theme, are plotted in Fig. 4B. Values shown as 0 were below the numerical or permutation resolution of the GSEA output (228 nominal P and 22 FDR q values).

| GO ID | GO name | Category | Size | ES | NES | Nominal P | FDR q | Leading edge |
|---|---|---|---|---|---|---|---|---|
| **Enriched in the Rin interaction — positive NES (FDR *q* < 0.05; *n* = 162)** | | | | | | | | |
| GO:0019867 | outer membrane | Cellular Component | 270 | 0.496 | 2.589 | 0 | 0 | tags=49%, list=22%, signal=63% |
| GO:0042651 | thylakoid membrane | Cellular Component | 198 | 0.519 | 2.572 | 0 | 0 | tags=57%, list=24%, signal=74% |
| GO:0034357 | photosynthetic membrane | Cellular Component | 211 | 0.517 | 2.567 | 0 | 0 | tags=54%, list=22%, signal=69% |
| GO:0009535 | chloroplast thylakoid membrane | Cellular Component | 184 | 0.518 | 2.556 | 0 | 0 | tags=57%, list=24%, signal=74% |
| GO:0055035 | plastid thylakoid membrane | Cellular Component | 184 | 0.518 | 2.552 | 0 | 0 | tags=57%, list=24%, signal=74% |
| GO:0031968 | organelle outer membrane | Cellular Component | 270 | 0.496 | 2.549 | 0 | 0 | tags=49%, list=22%, signal=63% |
| GO:0015979 | photosynthesis | Biological Process | 141 | 0.525 | 2.487 | 0 | 0 | tags=57%, list=25%, signal=75% |
| GO:0009534 | chloroplast thylakoid | Cellular Component | 256 | 0.480 | 2.480 | 0 | 0 | tags=50%, list=21%, signal=62% |
| GO:0031976 | plastid thylakoid | Cellular Component | 256 | 0.480 | 2.441 | 0 | 1.68e-04 | tags=50%, list=21%, signal=62% |
| GO:0009579 | thylakoid | Cellular Component | 281 | 0.472 | 2.428 | 0 | 1.51e-04 | tags=49%, list=21%, signal=61% |
| GO:0019684 | photosynthesis, light reaction | Biological Process | 66 | 0.586 | 2.420 | 0 | 1.37e-04 | tags=59%, list=19%, signal=73% |
| GO:0019725 | cellular homeostasis | Biological Process | 216 | 0.470 | 2.377 | 0 | 1.26e-04 | tags=53%, list=25%, signal=70% |
| GO:0032040 | small-subunit processome | Cellular Component | 61 | 0.574 | 2.351 | 0 | 1.16e-04 | tags=56%, list=27%, signal=76% |
| GO:0005730 | nucleolus | Cellular Component | 268 | 0.455 | 2.348 | 0 | 1.08e-04 | tags=53%, list=27%, signal=72% |

| GO ID | GO name | Category | Size | ES | NES | Nominal P | FDR q | Leading edge |
|---|---|---|---|---|---|---|---|---|
| GO:0042170 | plastid membrane | Cellular Component | 298 | 0.449 | 2.325 | 0 | 1.01e-04 | tags=48%, list=24%, signal=63% |
| GO:0009070 | serine family amino acid biosynthetic process | Biological Process | 29 | 0.659 | 2.278 | 0 | 1.91e-04 | tags=59%, list=19%, signal=72% |
| GO:0045454 | cell redox homeostasis | Biological Process | 102 | 0.506 | 2.276 | 0 | 1.79e-04 | tags=65%, list=27%, signal=89% |
| GO:0030686 | 90S preribosome | Cellular Component | 38 | 0.596 | 2.274 | 0 | 1.69e-04 | tags=66%, list=27%, signal=90% |
| GO:0030684 | preribosome | Cellular Component | 102 | 0.500 | 2.261 | 0 | 1.60e-04 | tags=53%, list=27%, signal=73% |
| GO:0030490 | maturation of SSU-rRNA | Biological Process | 55 | 0.564 | 2.260 | 0 | 1.52e-04 | tags=51%, list=23%, signal=66% |
| GO:0009526 | plastid envelope | Cellular Component | 350 | 0.418 | 2.251 | 0 | 1.45e-04 | tags=45%, list=24%, signal=58% |
| GO:0006364 | rRNA processing | Biological Process | 216 | 0.442 | 2.210 | 0 | 7.53e-04 | tags=43%, list=23%, signal=56% |
| GO:0016072 | rRNA metabolic process | Biological Process | 222 | 0.442 | 2.202 | 0 | 7.84e-04 | tags=50%, list=28%, signal=69% |
| GO:0015977 | carbon fixation | Biological Process | 21 | 0.688 | 2.202 | 0 | 7.51e-04 | tags=48%, list=6%, signal=50% |
| GO:0009521 | photosystem | Cellular Component | 60 | 0.532 | 2.199 | 0 | 7.81e-04 | tags=53%, list=23%, signal=70% |
| GO:0071806 | protein transmembrane transport | Biological Process | 65 | 0.514 | 2.185 | 0 | 0.0011 | tags=46%, list=25%, signal=62% |
| GO:0006383 | transcription by RNA polymerase III | Biological Process | 23 | 0.663 | 2.177 | 0 | 0.0010 | tags=57%, list=19%, signal=69% |
| GO:0005681 | spliceosomal complex | Cellular Component | 138 | 0.445 | 2.151 | 0 | 0.0015 | tags=47%, list=28%, signal=65% |
| GO:0044743 | protein transmembrane import into intracellular organelle | Biological Process | 37 | 0.575 | 2.145 | 0 | 0.0015 | tags=54%, list=25%, signal=72% |
| GO:0005986 | sucrose biosynthetic process | Biological Process | 18 | 0.713 | 2.139 | 0 | 0.0017 | tags=67%, list=14%, signal=78% |
| GO:0042592 | homeostatic process | Biological Process | 287 | 0.411 | 2.136 | 0 | 0.0018 | tags=46%, list=23%, signal=59% |
| GO:0000462 | maturation of SSU-rRNA from tricistronic rRNA transcript (SSU-rRNA, 5.8S rRNA, LSU-rRNA) | Biological Process | 38 | 0.571 | 2.118 | 0 | 0.0021 | tags=53%, list=23%, signal=69% |
| GO:0015035 | protein-disulfide reductase activity | Molecular Function | 37 | 0.574 | 2.106 | 0 | 0.0024 | tags=70%, list=30%, signal=100% |

| GO ID | GO name | Category | Size | ES | NES | Nominal P | FDR q | Leading edge |
|---|---|---|---|---|---|---|---|---|
| GO:0016667 | oxidoreductase activity, acting on a sulfur group of donors | Molecular Function | 87 | 0.482 | 2.105 | 0 | 0.0024 | tags=57%, list=25%, signal=76% |
| GO:0030515 | snoRNA binding | Molecular Function | 21 | 0.657 | 2.087 | 0 | 0.0026 | tags=71%, list=23%, signal=93% |
| GO:0042254 | ribosome biogenesis | Biological Process | 319 | 0.388 | 2.065 | 0 | 0.0033 | tags=42%, list=27%, signal=58% |
| GO:0019682 | glyceraldehyde-3-phosphate metabolic process | Biological Process | 16 | 0.686 | 2.058 | 0 | 0.0037 | tags=63%, list=13%, signal=72% |
| GO:0019203 | carbohydrate phosphatase activity | Molecular Function | 39 | 0.551 | 2.053 | 0.0025 | 0.0040 | tags=41%, list=14%, signal=48% |
| GO:0071011 | precatalytic spliceosome | Cellular Component | 38 | 0.541 | 2.048 | 0 | 0.0041 | tags=61%, list=35%, signal=93% |
| GO:0065002 | intracellular protein transmembrane transport | Biological Process | 42 | 0.546 | 2.042 | 0 | 0.0044 | tags=50%, list=25%, signal=66% |
| GO:0009523 | photosystem II | Cellular Component | 40 | 0.540 | 2.034 | 0 | 0.0048 | tags=55%, list=20%, signal=69% |
| GO:0042274 | ribosomal small subunit biogenesis | Biological Process | 94 | 0.454 | 2.033 | 0 | 0.0048 | tags=43%, list=27%, signal=58% |
| GO:0055082 | intracellular chemical homeostasis | Biological Process | 114 | 0.442 | 2.033 | 0 | 0.0046 | tags=46%, list=23%, signal=59% |
| GO:0006544 | glycine metabolic process | Biological Process | 20 | 0.651 | 2.032 | 0 | 0.0046 | tags=75%, list=27%, signal=102% |
| GO:0016226 | iron-sulfur cluster assembly | Biological Process | 32 | 0.557 | 2.027 | 0 | 0.0048 | tags=59%, list=17%, signal=71% |
| GO:0043574 | peroxisomal transport | Biological Process | 18 | 0.663 | 2.020 | 0 | 0.0051 | tags=61%, list=23%, signal=79% |
| GO:0006541 | L-glutamine metabolic process | Biological Process | 45 | 0.530 | 2.018 | 0 | 0.0050 | tags=47%, list=17%, signal=56% |
| GO:0015919 | peroxisomal membrane transport | Biological Process | 18 | 0.663 | 2.017 | 0 | 0.0050 | tags=61%, list=23%, signal=79% |
| GO:0009368 | endopeptidase Clp complex | Cellular Component | 16 | 0.687 | 2.015 | 0 | 0.0051 | tags=44%, list=7%, signal=47% |
| GO:0022613 | ribonucleoprotein complex biogenesis | Biological Process | 416 | 0.370 | 2.009 | 0 | 0.0054 | tags=39%, list=28%, signal=53% |
| GO:0006352 | DNA-templated transcription initiation | Biological Process | 72 | 0.475 | 1.998 | 0 | 0.0061 | tags=63%, list=33%, signal=93% |
| GO:0000375 | RNA splicing, via transesterification reactions | Biological Process | 215 | 0.395 | 1.988 | 0 | 0.0067 | tags=48%, list=31%, signal=69% |

| GO ID | GO name | Category | Size | ES | NES | Nominal P | FDR q | Leading edge |
|---|---|---|---|---|---|---|---|---|
| GO:0031163 | metallo-sulfur cluster assembly | Biological Process | 32 | 0.557 | 1.986 | 0 | 0.0068 | tags=59%, list=17%, signal=71% |
| GO:0006367 | transcription initiation at RNA polymerase II promoter | Biological Process | 36 | 0.542 | 1.983 | 0 | 0.0070 | tags=75%, list=33%, signal=112% |
| GO:0000377 | RNA splicing, via transesterification reactions with bulged adenosine as nucleophile | Biological Process | 214 | 0.395 | 1.981 | 0 | 0.0069 | tags=48%, list=31%, signal=69% |
| GO:0180051 | translation factor activity | Molecular Function | 136 | 0.415 | 1.978 | 0 | 0.0069 | tags=37%, list=24%, signal=48% |
| GO:0006826 | iron ion transport | Biological Process | 28 | 0.578 | 1.972 | 0 | 0.0073 | tags=57%, list=27%, signal=78% |
| GO:0006873 | intracellular monoatomic ion homeostasis | Biological Process | 102 | 0.437 | 1.970 | 0 | 0.0074 | tags=44%, list=23%, signal=57% |
| GO:0008380 | RNA splicing | Biological Process | 236 | 0.386 | 1.967 | 0 | 0.0075 | tags=46%, list=30%, signal=65% |
| GO:0000398 | mRNA splicing, via spliceosome | Biological Process | 204 | 0.392 | 1.964 | 0 | 0.0079 | tags=48%, list=31%, signal=69% |
| GO:0003746 | translation elongation factor activity | Molecular Function | 35 | 0.543 | 1.958 | 0 | 0.0084 | tags=63%, list=33%, signal=94% |
| GO:0009767 | photosynthetic electron transport chain | Biological Process | 20 | 0.628 | 1.957 | 0 | 0.0084 | tags=65%, list=23%, signal=84% |
| GO:0006399 | tRNA metabolic process | Biological Process | 215 | 0.388 | 1.939 | 0 | 0.0098 | tags=47%, list=31%, signal=67% |
| GO:0030003 | intracellular monoatomic cation homeostasis | Biological Process | 101 | 0.435 | 1.934 | 0 | 0.0102 | tags=44%, list=23%, signal=57% |
| GO:0000959 | mitochondrial RNA metabolic process | Biological Process | 17 | 0.645 | 1.930 | 0.0069 | 0.0106 | tags=59%, list=24%, signal=77% |
| GO:0000422 | autophagy of mitochondrion | Biological Process | 18 | 0.631 | 1.928 | 0 | 0.0106 | tags=44%, list=12%, signal=51% |
| GO:0036452 | ESCRT complex | Cellular Component | 20 | 0.612 | 1.928 | 0.0024 | 0.0105 | tags=55%, list=24%, signal=73% |
| GO:0009836 | fruit ripening, climacteric | Biological Process | 19 | 0.619 | 1.912 | 0.0047 | 0.0120 | tags=37%, list=13%, signal=42% |
| GO:0072662 | protein localization to peroxisome | Biological Process | 16 | 0.640 | 1.901 | 0.0062 | 0.0134 | tags=56%, list=23%, signal=73% |
| GO:0019318 | hexose metabolic process | Biological Process | 80 | 0.437 | 1.891 | 0 | 0.0147 | tags=44%, list=20%, signal=54% |
| GO:0006094 | gluconeogenesis | Biological Process | 38 | 0.507 | 1.887 | 0 | 0.0149 | tags=50%, list=18%, signal=61% |

| GO ID | GO name | Category | Size | ES | NES | Nominal P | FDR q | Leading edge |
|---|---|---|---|---|---|---|---|---|
| GO:0098771 | inorganic ion homeostasis | Biological Process | 94 | 0.424 | 1.884 | 0 | 0.0152 | tags=31%, list=16%, signal=37% |
| GO:0072663 | establishment of protein localization to peroxisome | Biological Process | 16 | 0.640 | 1.877 | 0.0023 | 0.0163 | tags=56%, list=23%, signal=73% |
| GO:0048878 | chemical homeostasis | Biological Process | 174 | 0.385 | 1.875 | 0 | 0.0163 | tags=41%, list=23%, signal=53% |
| GO:0009765 | photosynthesis, light harvesting | Biological Process | 33 | 0.510 | 1.870 | 0.0024 | 0.0169 | tags=55%, list=19%, signal=67% |
| GO:0009835 | fruit ripening | Biological Process | 20 | 0.591 | 1.869 | 0.0046 | 0.0169 | tags=35%, list=13%, signal=40% |
| GO:0005744 | TIM23 mitochondrial import inner membrane translocase complex | Cellular Component | 15 | 0.640 | 1.868 | 0.0024 | 0.0168 | tags=67%, list=28%, signal=93% |
| GO:0005985 | sucrose metabolic process | Biological Process | 37 | 0.506 | 1.861 | 0 | 0.0176 | tags=43%, list=15%, signal=51% |
| GO:0019319 | hexose biosynthetic process | Biological Process | 38 | 0.507 | 1.854 | 0 | 0.0185 | tags=50%, list=18%, signal=61% |
| GO:0005996 | monosaccharide metabolic process | Biological Process | 106 | 0.412 | 1.851 | 0 | 0.0191 | tags=42%, list=20%, signal=53% |
| GO:0034755 | iron ion transmembrane transport | Biological Process | 16 | 0.633 | 1.850 | 0.0047 | 0.0191 | tags=63%, list=22%, signal=81% |
| GO:0015036 | disulfide oxidoreductase activity | Molecular Function | 50 | 0.469 | 1.842 | 0.0027 | 0.0201 | tags=64%, list=30%, signal=91% |
| GO:0042802 | identical protein binding | Molecular Function | 62 | 0.449 | 1.840 | 0 | 0.0203 | tags=40%, list=22%, signal=52% |
| GO:0006091 | generation of precursor metabolites and energy | Biological Process | 339 | 0.353 | 1.839 | 0 | 0.0203 | tags=39%, list=23%, signal=50% |
| GO:0000041 | transition metal ion transport | Biological Process | 70 | 0.434 | 1.837 | 0 | 0.0204 | tags=31%, list=12%, signal=36% |
| GO:0006006 | glucose metabolic process | Biological Process | 67 | 0.437 | 1.836 | 0 | 0.0203 | tags=42%, list=20%, signal=52% |
| GO:0046351 | disaccharide biosynthetic process | Biological Process | 33 | 0.519 | 1.835 | 0 | 0.0203 | tags=42%, list=14%, signal=49% |
| GO:0006081 | aldehyde metabolic process | Biological Process | 52 | 0.452 | 1.827 | 0 | 0.0217 | tags=46%, list=19%, signal=57% |
| GO:0055080 | monoatomic cation homeostasis | Biological Process | 117 | 0.399 | 1.827 | 0 | 0.0215 | tags=42%, list=23%, signal=54% |
| GO:0005666 | RNA polymerase III complex | Cellular Component | 25 | 0.559 | 1.825 | 0.0049 | 0.0217 | tags=44%, list=18%, signal=54% |

| GO ID | GO name | Category | Size | ES | NES | Nominal P | FDR q | Leading edge |
|---|---|---|---|---|---|---|---|---|
| GO:0005741 | mitochondrial outer membrane | Cellular Component | 56 | 0.449 | 1.820 | 0.0027 | 0.0226 | tags=54%, list=31%, signal=78% |
| GO:7770058 | mitochondrial protein import pathway | Biological Process | 38 | 0.502 | 1.819 | 0 | 0.0225 | tags=58%, list=32%, signal=86% |
| GO:0050308 | sugar-phosphatase activity | Molecular Function | 31 | 0.504 | 1.816 | 0.0025 | 0.0229 | tags=42%, list=14%, signal=49% |
| GO:0050801 | monoatomic ion homeostasis | Biological Process | 123 | 0.386 | 1.812 | 0 | 0.0236 | tags=42%, list=23%, signal=55% |
| GO:0019252 | starch biosynthetic process | Biological Process | 17 | 0.599 | 1.810 | 0.0070 | 0.0236 | tags=71%, list=28%, signal=99% |
| GO:0006397 | mRNA processing | Biological Process | 263 | 0.350 | 1.808 | 0 | 0.0236 | tags=43%, list=30%, signal=61% |
| GO:0005984 | disaccharide metabolic process | Biological Process | 53 | 0.442 | 1.808 | 0 | 0.0234 | tags=36%, list=15%, signal=42% |
| GO:0009654 | photosystem II oxygen evolving complex | Cellular Component | 23 | 0.553 | 1.804 | 0.0048 | 0.0240 | tags=57%, list=20%, signal=70% |
| GO:0007031 | peroxisome organization | Biological Process | 38 | 0.474 | 1.803 | 0.0050 | 0.0240 | tags=47%, list=23%, signal=61% |
| GO:0005982 | starch metabolic process | Biological Process | 30 | 0.509 | 1.801 | 0.0073 | 0.0240 | tags=57%, list=26%, signal=77% |
| GO:0140053 | mitochondrial gene expression | Biological Process | 41 | 0.480 | 1.798 | 0.0050 | 0.0248 | tags=54%, list=33%, signal=80% |
| GO:0000428 | DNA-directed RNA polymerase complex | Cellular Component | 115 | 0.389 | 1.794 | 0 | 0.0253 | tags=57%, list=35%, signal=87% |
| GO:0071214 | cellular response to abiotic stimulus | Biological Process | 47 | 0.456 | 1.794 | 0.0026 | 0.0251 | tags=51%, list=24%, signal=67% |
| GO:0005732 | sno(s)RNA-containing ribonucleoprotein complex | Cellular Component | 21 | 0.564 | 1.793 | 0 | 0.0251 | tags=90%, list=40%, signal=149% |
| GO:0160215 | deacylase activity | Molecular Function | 53 | 0.441 | 1.792 | 0 | 0.0250 | tags=47%, list=30%, signal=67% |
| GO:0006754 | ATP biosynthetic process | Biological Process | 28 | 0.514 | 1.790 | 0.0045 | 0.0254 | tags=82%, list=41%, signal=140% |
| GO:0104004 | cellular response to environmental stimulus | Biological Process | 47 | 0.456 | 1.788 | 0.0025 | 0.0256 | tags=51%, list=24%, signal=67% |
| GO:0045259 | proton-transporting ATP synthase complex | Cellular Component | 18 | 0.572 | 1.787 | 0.0071 | 0.0256 | tags=89%, list=41%, signal=151% |
| GO:0097525 | spliceosomal snRNP complex | Cellular Component | 67 | 0.423 | 1.787 | 0.0026 | 0.0255 | tags=49%, list=35%, signal=75% |

| GO ID | GO name | Category | Size | ES | NES | Nominal P | FDR q | Leading edge |
|---|---|---|---|---|---|---|---|---|
| GO:0015986 | proton motive force-driven ATP synthesis | Biological Process | 28 | 0.514 | 1.784 | 0.0121 | 0.0259 | tags=82%, list=41%, signal=140% |
| GO:0008186 | ATP-dependent activity, acting on RNA | Molecular Function | 97 | 0.393 | 1.781 | 0 | 0.0262 | tags=44%, list=22%, signal=57% |
| GO:0006515 | protein quality control for misfolded or incompletely synthesized proteins | Biological Process | 19 | 0.573 | 1.777 | 0.0072 | 0.0270 | tags=37%, list=7%, signal=40% |
| GO:0048511 | rhythmic process | Biological Process | 19 | 0.569 | 1.768 | 0.0024 | 0.0294 | tags=58%, list=21%, signal=73% |
| GO:0071013 | catalytic step 2 spliceosome | Cellular Component | 58 | 0.445 | 1.766 | 0 | 0.0298 | tags=45%, list=30%, signal=64% |
| GO:0003724 | RNA helicase activity | Molecular Function | 97 | 0.393 | 1.765 | 0 | 0.0300 | tags=44%, list=22%, signal=57% |
| GO:0046364 | monosaccharide biosynthetic process | Biological Process | 45 | 0.469 | 1.763 | 0 | 0.0303 | tags=47%, list=18%, signal=57% |
| GO:0031072 | heat shock protein binding | Molecular Function | 37 | 0.476 | 1.760 | 0 | 0.0306 | tags=49%, list=27%, signal=66% |
| GO:0005684 | U2-type spliceosomal complex | Cellular Component | 45 | 0.454 | 1.757 | 0.0026 | 0.0315 | tags=31%, list=14%, signal=36% |
| GO:0009312 | oligosaccharide biosynthetic process | Biological Process | 37 | 0.478 | 1.757 | 0.0051 | 0.0314 | tags=38%, list=14%, signal=44% |
| GO:0032543 | mitochondrial translation | Biological Process | 34 | 0.476 | 1.753 | 0.0025 | 0.0322 | tags=56%, list=33%, signal=83% |
| GO:0000460 | maturation of 5.8S rRNA | Biological Process | 26 | 0.525 | 1.753 | 0.0025 | 0.0320 | tags=62%, list=26%, signal=83% |
| GO:0046915 | transition metal ion transmembrane transporter activity | Molecular Function | 56 | 0.436 | 1.751 | 0.0052 | 0.0323 | tags=34%, list=12%, signal=39% |
| GO:0006879 | intracellular iron ion homeostasis | Biological Process | 19 | 0.561 | 1.749 | 0.0045 | 0.0325 | tags=32%, list=11%, signal=36% |
| GO:0004176 | ATP-dependent peptidase activity | Molecular Function | 40 | 0.469 | 1.744 | 0 | 0.0336 | tags=45%, list=20%, signal=56% |
| GO:0006414 | translational elongation | Biological Process | 54 | 0.439 | 1.741 | 0 | 0.0344 | tags=59%, list=38%, signal=95% |
| GO:0051117 | ATPase binding | Molecular Function | 20 | 0.563 | 1.738 | 0.0071 | 0.0349 | tags=45%, list=16%, signal=54% |
| GO:0042803 | protein homodimerization activity | Molecular Function | 37 | 0.477 | 1.738 | 0.0057 | 0.0349 | tags=51%, list=29%, signal=72% |
| GO:0030150 | protein import into mitochondrial matrix | Biological Process | 22 | 0.545 | 1.733 | 0.0104 | 0.0360 | tags=55%, list=27%, signal=75% |

| GO ID | GO name | Category | Size | ES | NES | Nominal P | FDR q | Leading edge |
|---|---|---|---|---|---|---|---|---|
| GO:0055088 | lipid homeostasis | Biological Process | 32 | 0.484 | 1.732 | 0.0125 | 0.0360 | tags=41%, list=13%, signal=47% |
| GO:0000097 | sulfur amino acid biosynthetic process | Biological Process | 15 | 0.596 | 1.732 | 0.0092 | 0.0357 | tags=53%, list=18%, signal=65% |
| GO:0009707 | chloroplast outer membrane | Cellular Component | 29 | 0.494 | 1.728 | 0.0024 | 0.0369 | tags=31%, list=10%, signal=35% |
| GO:0042558 | pteridine-containing compound metabolic process | Biological Process | 32 | 0.483 | 1.723 | 0.0046 | 0.0383 | tags=44%, list=19%, signal=54% |
| GO:0035097 | histone methyltransferase complex | Cellular Component | 17 | 0.568 | 1.723 | 0.0046 | 0.0380 | tags=59%, list=25%, signal=79% |
| GO:0016036 | cellular response to phosphate starvation | Biological Process | 18 | 0.560 | 1.717 | 0.0044 | 0.0395 | tags=44%, list=21%, signal=57% |
| GO:0004197 | cysteine-type endopeptidase activity | Molecular Function | 34 | 0.471 | 1.715 | 0.0026 | 0.0398 | tags=24%, list=4%, signal=24% |
| GO:0170038 | proteinogenic amino acid biosynthetic process | Biological Process | 118 | 0.371 | 1.715 | 0 | 0.0396 | tags=36%, list=19%, signal=44% |
| GO:0140101 | catalytic activity, acting on a tRNA | Molecular Function | 167 | 0.351 | 1.714 | 0 | 0.0397 | tags=42%, list=30%, signal=60% |
| GO:0047134 | protein-disulfide reductase [NAD(P)H] activity | Molecular Function | 16 | 0.584 | 1.711 | 0.0149 | 0.0404 | tags=63%, list=21%, signal=79% |
| GO:1905354 | exoribonuclease complex | Cellular Component | 15 | 0.595 | 1.710 | 0.0278 | 0.0403 | tags=67%, list=28%, signal=93% |
| GO:0015295 | solute:proton symporter activity | Molecular Function | 19 | 0.550 | 1.707 | 0.0023 | 0.0412 | tags=47%, list=19%, signal=58% |
| GO:0046933 | proton-transporting ATP synthase activity, rotational mechanism | Molecular Function | 17 | 0.573 | 1.707 | 0.0051 | 0.0412 | tags=88%, list=41%, signal=150% |
| GO:0015078 | proton transmembrane transporter activity | Molecular Function | 161 | 0.350 | 1.706 | 0 | 0.0411 | tags=35%, list=23%, signal=45% |
| GO:0008033 | tRNA processing | Biological Process | 114 | 0.371 | 1.702 | 0.0033 | 0.0421 | tags=48%, list=31%, signal=69% |
| GO:0010411 | xyloglucan metabolic process | Biological Process | 37 | 0.458 | 1.700 | 0.0027 | 0.0424 | tags=19%, list=4%, signal=20% |
| GO:0034708 | methyltransferase complex | Cellular Component | 24 | 0.511 | 1.696 | 0.0101 | 0.0435 | tags=42%, list=17%, signal=50% |
| GO:0016071 | mRNA metabolic process | Biological Process | 352 | 0.318 | 1.695 | 0 | 0.0437 | tags=39%, list=29%, signal=55% |
| GO:0051536 | iron-sulfur cluster binding | Molecular Function | 104 | 0.370 | 1.693 | 0 | 0.0442 | tags=34%, list=18%, signal=41% |

| GO ID | GO name | Category | Size | ES | NES | Nominal P | FDR q | Leading edge |
|---|---|---|---|---|---|---|---|---|
| GO:0009527 | plastid outer membrane | Cellular Component | 29 | 0.494 | 1.688 | 0.0122 | 0.0456 | tags=31%, list=10%, signal=35% |
| GO:0046653 | tetrahydrofolate metabolic process | Biological Process | 26 | 0.500 | 1.686 | 0.0026 | 0.0462 | tags=42%, list=14%, signal=49% |
| GO:0007623 | circadian rhythm | Biological Process | 19 | 0.569 | 1.685 | 0.0047 | 0.0460 | tags=58%, list=21%, signal=73% |
| GO:0005777 | peroxisome | Cellular Component | 118 | 0.361 | 1.682 | 0 | 0.0469 | tags=37%, list=20%, signal=46% |
| GO:0051087 | protein-folding chaperone binding | Molecular Function | 75 | 0.395 | 1.682 | 0.0029 | 0.0468 | tags=39%, list=25%, signal=51% |
| GO:0016180 | snRNA processing | Biological Process | 17 | 0.557 | 1.682 | 0.0238 | 0.0465 | tags=76%, list=38%, signal=123% |
| GO:0009145 | purine nucleoside triphosphate biosynthetic process | Biological Process | 35 | 0.460 | 1.681 | 0.0079 | 0.0464 | tags=80%, list=41%, signal=136% |
| GO:0006450 | regulation of translational fidelity | Biological Process | 18 | 0.553 | 1.680 | 0.0128 | 0.0466 | tags=78%, list=38%, signal=126% |
| GO:0016671 | oxidoreductase activity, acting on a sulfur group of donors, disulfide as acceptor | Molecular Function | 21 | 0.519 | 1.677 | 0.0121 | 0.0477 | tags=67%, list=24%, signal=87% |
| GO:0006099 | tricarboxylic acid cycle | Biological Process | 51 | 0.427 | 1.673 | 0.0026 | 0.0487 | tags=37%, list=18%, signal=45% |
| GO:0042579 | microbody | Cellular Component | 118 | 0.361 | 1.673 | 0 | 0.0485 | tags=37%, list=20%, signal=46% |
| GO:0170036 | import into the mitochondrion | Biological Process | 30 | 0.482 | 1.673 | 0.0051 | 0.0484 | tags=57%, list=28%, signal=79% |
| GO:0170039 | proteinogenic amino acid metabolic process | Biological Process | 224 | 0.330 | 1.672 | 0 | 0.0482 | tags=33%, list=19%, signal=41% |
| GO:0016073 | snRNA metabolic process | Biological Process | 18 | 0.548 | 1.670 | 0.0223 | 0.0488 | tags=72%, list=38%, signal=116% |
| GO:0009206 | purine ribonucleoside triphosphate biosynthetic process | Biological Process | 35 | 0.460 | 1.669 | 0.0026 | 0.0491 | tags=80%, list=41%, signal=136% |
| **Enriched in the Rola interaction — negative NES (FDR *q* < 0.05; *n* = 152)** | | | | | | | | |
| GO:0007018 | microtubule-based movement | Biological Process | 55 | -0.697 | -2.579 | 0 | 0 | tags=67%, list=18%, signal=82% |
| GO:0003777 | microtubule motor activity | Molecular Function | 55 | -0.697 | -2.550 | 0 | 0 | tags=67%, list=18%, signal=82% |
| GO:0120544 | polypeptide conformation or assembly isomerase activity | Molecular Function | 65 | -0.659 | -2.502 | 0 | 0 | tags=62%, list=18%, signal=75% |

| GO ID | GO name | Category | Size | ES | NES | Nominal P | FDR q | Leading edge |
|---|---|---|---|---|---|---|---|---|
| GO:0015631 | tubulin binding | Molecular Function | 142 | -0.573 | -2.477 | 0 | 0 | tags=54%, list=20%, signal=67% |
| GO:0008017 | microtubule binding | Molecular Function | 132 | -0.574 | -2.461 | 0 | 0 | tags=55%, list=20%, signal=68% |
| GO:0000786 | nucleosome | Cellular Component | 78 | -0.629 | -2.457 | 0 | 0 | tags=59%, list=20%, signal=73% |
| GO:0003774 | cytoskeletal motor activity | Molecular Function | 67 | -0.643 | -2.442 | 0 | 0 | tags=60%, list=18%, signal=73% |
| GO:0000278 | mitotic cell cycle | Biological Process | 129 | -0.571 | -2.430 | 0 | 0 | tags=50%, list=18%, signal=61% |
| GO:0007059 | chromosome segregation | Biological Process | 88 | -0.589 | -2.401 | 0 | 0 | tags=47%, list=15%, signal=55% |
| GO:0022402 | cell cycle process | Biological Process | 247 | -0.516 | -2.384 | 0 | 0 | tags=44%, list=17%, signal=53% |
| GO:0010374 | stomatal complex development | Biological Process | 37 | -0.700 | -2.369 | 0 | 0 | tags=68%, list=19%, signal=83% |
| GO:0030527 | structural constituent of chromatin | Molecular Function | 73 | -0.603 | -2.349 | 0 | 0 | tags=59%, list=20%, signal=73% |
| GO:1903047 | mitotic cell cycle process | Biological Process | 107 | -0.573 | -2.329 | 0 | 0 | tags=52%, list=18%, signal=64% |
| GO:0032993 | protein-DNA complex | Cellular Component | 122 | -0.551 | -2.325 | 0 | 0 | tags=48%, list=20%, signal=59% |
| GO:0098813 | nuclear chromosome segregation | Biological Process | 56 | -0.618 | -2.293 | 0 | 4.64e-05 | tags=52%, list=15%, signal=61% |
| GO:0007049 | cell cycle | Biological Process | 291 | -0.490 | -2.288 | 0 | 4.35e-05 | tags=42%, list=18%, signal=51% |
| GO:0008092 | cytoskeletal protein binding | Molecular Function | 238 | -0.493 | -2.260 | 0 | 4.10e-05 | tags=46%, list=20%, signal=57% |
| GO:0099512 | supramolecular fiber | Cellular Component | 116 | -0.530 | -2.224 | 0 | 7.68e-05 | tags=51%, list=20%, signal=63% |
| GO:0099513 | polymeric cytoskeletal fiber | Cellular Component | 116 | -0.530 | -2.217 | 0 | 7.28e-05 | tags=51%, list=20%, signal=63% |
| GO:0090627 | plant epidermal cell differentiation | Biological Process | 61 | -0.580 | -2.207 | 0 | 1.05e-04 | tags=49%, list=21%, signal=62% |
| GO:0016837 | carbon-oxygen lyase activity, acting on polysaccharides | Molecular Function | 19 | -0.747 | -2.200 | 0 | 1.00e-04 | tags=58%, list=14%, signal=67% |
| GO:0099081 | supramolecular polymer | Cellular Component | 116 | -0.530 | -2.190 | 0 | 9.55e-05 | tags=51%, list=20%, signal=63% |

| GO ID | GO name | Category | Size | ES | NES | Nominal P | FDR q | Leading edge |
|---|---|---|---|---|---|---|---|---|
| GO:0005874 | microtubule | Cellular Component | 111 | -0.528 | -2.186 | 0 | 9.14e-05 | tags=50%, list=20%, signal=63% |
| GO:0000226 | microtubule cytoskeleton organization | Biological Process | 106 | -0.526 | -2.167 | 0 | 2.08e-04 | tags=49%, list=18%, signal=60% |
| GO:0045488 | pectin metabolic process | Biological Process | 65 | -0.571 | -2.162 | 0 | 2.56e-04 | tags=43%, list=17%, signal=52% |
| GO:0009733 | response to auxin | Biological Process | 168 | -0.484 | -2.144 | 0 | 3.01e-04 | tags=49%, list=23%, signal=64% |
| GO:0090698 | post-embryonic plant morphogenesis | Biological Process | 34 | -0.633 | -2.140 | 0 | 2.90e-04 | tags=53%, list=19%, signal=65% |
| GO:0071365 | cellular response to auxin stimulus | Biological Process | 86 | -0.531 | -2.139 | 0 | 3.04e-04 | tags=51%, list=22%, signal=65% |
| GO:0042973 | glucan endo-1,3-beta-D-glucosidase activity | Molecular Function | 65 | -0.555 | -2.121 | 0 | 3.69e-04 | tags=52%, list=22%, signal=67% |
| GO:0030570 | pectate lyase activity | Molecular Function | 16 | -0.747 | -2.114 | 0 | 4.28e-04 | tags=50%, list=7%, signal=53% |
| GO:0044772 | mitotic cell cycle phase transition | Biological Process | 49 | -0.575 | -2.113 | 0 | 4.14e-04 | tags=45%, list=14%, signal=52% |
| GO:0044786 | cell cycle DNA replication | Biological Process | 37 | -0.622 | -2.112 | 0 | 4.02e-04 | tags=65%, list=21%, signal=82% |
| GO:0009734 | auxin-activated signaling pathway | Biological Process | 85 | -0.525 | -2.110 | 0 | 3.89e-04 | tags=51%, list=22%, signal=64% |
| GO:0007010 | cytoskeleton organization | Biological Process | 217 | -0.460 | -2.105 | 0 | 3.99e-04 | tags=41%, list=19%, signal=50% |
| GO:0015630 | microtubule cytoskeleton | Cellular Component | 204 | -0.467 | -2.104 | 0 | 4.08e-04 | tags=46%, list=22%, signal=59% |
| GO:0007017 | microtubule-based process | Biological Process | 179 | -0.471 | -2.086 | 0 | 4.96e-04 | tags=43%, list=18%, signal=52% |
| GO:0048869 | cellular developmental process | Biological Process | 166 | -0.475 | -2.085 | 0 | 4.82e-04 | tags=49%, list=27%, signal=67% |
| GO:0045490 | pectin catabolic process | Biological Process | 42 | -0.589 | -2.084 | 0 | 4.70e-04 | tags=48%, list=17%, signal=57% |
| GO:0010103 | stomatal complex morphogenesis | Biological Process | 28 | -0.649 | -2.078 | 0 | 5.30e-04 | tags=61%, list=19%, signal=75% |
| GO:0030154 | cell differentiation | Biological Process | 162 | -0.472 | -2.075 | 0 | 5.34e-04 | tags=49%, list=27%, signal=66% |
| GO:0051052 | regulation of DNA metabolic process | Biological Process | 47 | -0.575 | -2.074 | 0 | 5.21e-04 | tags=64%, list=22%, signal=82% |

| GO ID | GO name | Category | Size | ES | NES | Nominal P | FDR q | Leading edge |
|---|---|---|---|---|---|---|---|---|
| GO:0045229 | external encapsulating structure organization | Biological Process | 90 | -0.512 | -2.065 | 0 | 6.45e-04 | tags=40%, list=20%, signal=50% |
| GO:0090558 | plant epidermis development | Biological Process | 83 | -0.516 | -2.052 | 0 | 7.63e-04 | tags=45%, list=20%, signal=56% |
| GO:0044770 | cell cycle phase transition | Biological Process | 65 | -0.529 | -2.050 | 0 | 7.46e-04 | tags=52%, list=23%, signal=67% |
| GO:0071555 | cell wall organization | Biological Process | 87 | -0.511 | -2.047 | 0 | 7.45e-04 | tags=40%, list=20%, signal=50% |
| GO:0006275 | regulation of DNA replication | Biological Process | 27 | -0.633 | -2.046 | 0 | 7.29e-04 | tags=67%, list=21%, signal=84% |
| GO:0046982 | protein heterodimerization activity | Molecular Function | 121 | -0.479 | -2.040 | 0 | 7.74e-04 | tags=38%, list=16%, signal=45% |
| GO:0051301 | cell division | Biological Process | 68 | -0.531 | -2.034 | 0 | 8.47e-04 | tags=37%, list=14%, signal=43% |
| GO:0007051 | spindle organization | Biological Process | 57 | -0.545 | -2.028 | 0 | 9.17e-04 | tags=42%, list=15%, signal=50% |
| GO:0005819 | spindle | Cellular Component | 61 | -0.540 | -2.023 | 0 | 9.55e-04 | tags=56%, list=25%, signal=74% |
| GO:0051015 | actin filament binding | Molecular Function | 41 | -0.579 | -2.017 | 0 | 0.0011 | tags=59%, list=24%, signal=77% |
| GO:0140014 | mitotic nuclear division | Biological Process | 39 | -0.584 | -2.010 | 0 | 0.0014 | tags=49%, list=14%, signal=57% |
| GO:0000819 | sister chromatid segregation | Biological Process | 41 | -0.570 | -2.008 | 0 | 0.0014 | tags=46%, list=15%, signal=54% |
| GO:0009609 | response to symbiotic bacterium | Biological Process | 24 | -0.648 | -2.004 | 0 | 0.0015 | tags=46%, list=15%, signal=54% |
| GO:0010052 | guard cell differentiation | Biological Process | 24 | -0.642 | -1.990 | 0 | 0.0018 | tags=58%, list=19%, signal=72% |
| GO:0000280 | nuclear division | Biological Process | 90 | -0.495 | -1.989 | 0 | 0.0018 | tags=40%, list=14%, signal=47% |
| GO:0031109 | microtubule polymerization or depolymerization | Biological Process | 20 | -0.677 | -1.986 | 0.0017 | 0.0019 | tags=65%, list=17%, signal=78% |
| GO:0010564 | regulation of cell cycle process | Biological Process | 76 | -0.508 | -1.985 | 0 | 0.0018 | tags=51%, list=22%, signal=65% |
| GO:0000070 | mitotic sister chromatid segregation | Biological Process | 38 | -0.566 | -1.951 | 0 | 0.0029 | tags=47%, list=14%, signal=55% |
| GO:0007052 | mitotic spindle organization | Biological Process | 28 | -0.604 | -1.946 | 0 | 0.0031 | tags=50%, list=13%, signal=58% |

| GO ID | GO name | Category | Size | ES | NES | Nominal P | FDR q | Leading edge |
|---|---|---|---|---|---|---|---|---|
| GO:0005856 | cytoskeleton | Cellular Component | 357 | -0.408 | -1.946 | 0 | 0.0031 | tags=39%, list=22%, signal=50% |
| GO:0048507 | meristem development | Biological Process | 55 | -0.522 | -1.944 | 0 | 0.0031 | tags=29%, list=10%, signal=32% |
| GO:0033260 | nuclear DNA replication | Biological Process | 18 | -0.679 | -1.936 | 0 | 0.0034 | tags=72%, list=21%, signal=91% |
| GO:0005618 | cell wall | Cellular Component | 30 | -0.582 | -1.929 | 0 | 0.0038 | tags=37%, list=10%, signal=41% |
| GO:0006261 | DNA-templated DNA replication | Biological Process | 113 | -0.460 | -1.926 | 0 | 0.0038 | tags=44%, list=25%, signal=59% |
| GO:0030312 | external encapsulating structure | Cellular Component | 35 | -0.569 | -1.926 | 0 | 0.0038 | tags=49%, list=21%, signal=62% |
| GO:0048532 | anatomical structure arrangement | Biological Process | 19 | -0.657 | -1.925 | 0 | 0.0037 | tags=37%, list=10%, signal=41% |
| GO:0090626 | plant epidermis morphogenesis | Biological Process | 40 | -0.552 | -1.914 | 0.0016 | 0.0043 | tags=48%, list=19%, signal=59% |
| GO:0051276 | chromosome organization | Biological Process | 120 | -0.454 | -1.908 | 0 | 0.0047 | tags=44%, list=22%, signal=57% |
| GO:0051225 | spindle assembly | Biological Process | 40 | -0.552 | -1.903 | 0 | 0.0049 | tags=73%, list=31%, signal=105% |
| GO:2000022 | regulation of jasmonic acid mediated signaling pathway | Biological Process | 22 | -0.622 | -1.900 | 0 | 0.0051 | tags=77%, list=26%, signal=105% |
| GO:0009608 | response to symbiont | Biological Process | 34 | -0.566 | -1.899 | 0 | 0.0051 | tags=38%, list=15%, signal=45% |
| GO:0009505 | plant-type cell wall | Cellular Component | 30 | -0.582 | -1.893 | 0.0016 | 0.0056 | tags=37%, list=10%, signal=41% |
| GO:0007062 | sister chromatid cohesion | Biological Process | 23 | -0.617 | -1.885 | 0.0035 | 0.0061 | tags=52%, list=16%, signal=62% |
| GO:0097435 | supramolecular fiber organization | Biological Process | 106 | -0.457 | -1.882 | 0 | 0.0062 | tags=42%, list=20%, signal=52% |
| GO:1902850 | microtubule cytoskeleton organization involved in mitosis | Biological Process | 27 | -0.593 | -1.878 | 0 | 0.0064 | tags=48%, list=13%, signal=56% |
| GO:0033044 | regulation of chromosome organization | Biological Process | 20 | -0.633 | -1.870 | 0 | 0.0071 | tags=60%, list=22%, signal=76% |
| GO:0045814 | negative regulation of gene expression, epigenetic | Biological Process | 30 | -0.582 | -1.865 | 0 | 0.0075 | tags=53%, list=21%, signal=67% |
| GO:0015926 | glucosidase activity | Molecular Function | 108 | -0.443 | -1.862 | 0 | 0.0076 | tags=44%, list=22%, signal=57% |

| GO ID | GO name | Category | Size | ES | NES | Nominal P | FDR q | Leading edge |
|---|---|---|---|---|---|---|---|---|
| GO:0009664 | plant-type cell wall organization | Biological Process | 38 | -0.538 | -1.859 | 0.0016 | 0.0078 | tags=39%, list=15%, signal=47% |
| GO:0010016 | shoot system morphogenesis | Biological Process | 52 | -0.504 | -1.837 | 0.0016 | 0.0102 | tags=42%, list=20%, signal=53% |
| GO:0007346 | regulation of mitotic cell cycle | Biological Process | 43 | -0.512 | -1.823 | 0 | 0.0119 | tags=42%, list=17%, signal=50% |
| GO:0008422 | beta-glucosidase activity | Molecular Function | 91 | -0.454 | -1.818 | 0 | 0.0123 | tags=45%, list=22%, signal=57% |
| GO:0006633 | fatty acid biosynthetic process | Biological Process | 119 | -0.421 | -1.796 | 0 | 0.0156 | tags=26%, list=14%, signal=30% |
| GO:0072686 | mitotic spindle | Cellular Component | 20 | -0.610 | -1.794 | 0.0050 | 0.0159 | tags=60%, list=25%, signal=80% |
| GO:0048638 | regulation of developmental growth | Biological Process | 25 | -0.571 | -1.793 | 0.0050 | 0.0158 | tags=40%, list=14%, signal=46% |
| GO:0000775 | chromosome, centromeric region | Cellular Component | 26 | -0.556 | -1.779 | 0.0068 | 0.0182 | tags=42%, list=17%, signal=51% |
| GO:0016835 | carbon-oxygen lyase activity | Molecular Function | 104 | -0.432 | -1.776 | 0 | 0.0187 | tags=34%, list=18%, signal=41% |
| GO:1901987 | regulation of cell cycle phase transition | Biological Process | 49 | -0.492 | -1.768 | 0 | 0.0199 | tags=49%, list=22%, signal=62% |
| GO:0099080 | supramolecular complex | Cellular Component | 198 | -0.393 | -1.766 | 0 | 0.0199 | tags=31%, list=15%, signal=37% |
| GO:0090329 | regulation of DNA-templated DNA replication | Biological Process | 19 | -0.612 | -1.766 | 0 | 0.0199 | tags=63%, list=21%, signal=80% |
| GO:0032875 | regulation of DNA endoreduplication | Biological Process | 16 | -0.635 | -1.766 | 0.0017 | 0.0197 | tags=63%, list=16%, signal=75% |
| GO:0000272 | polysaccharide catabolic process | Biological Process | 95 | -0.434 | -1.763 | 0.0015 | 0.0199 | tags=34%, list=17%, signal=41% |
| GO:0051726 | regulation of cell cycle | Biological Process | 110 | -0.425 | -1.759 | 0 | 0.0205 | tags=35%, list=17%, signal=43% |
| GO:0042555 | MCM complex | Cellular Component | 15 | -0.650 | -1.758 | 0.0017 | 0.0204 | tags=73%, list=31%, signal=106% |
| GO:0040029 | epigenetic regulation of gene expression | Biological Process | 50 | -0.489 | -1.758 | 0.0032 | 0.0204 | tags=46%, list=21%, signal=58% |
| GO:0099402 | plant organ development | Biological Process | 210 | -0.387 | -1.757 | 0 | 0.0204 | tags=26%, list=15%, signal=30% |
| GO:0051783 | regulation of nuclear division | Biological Process | 16 | -0.635 | -1.744 | 0.0053 | 0.0234 | tags=50%, list=14%, signal=58% |

| GO ID | GO name | Category | Size | ES | NES | Nominal P | FDR q | Leading edge |
|---|---|---|---|---|---|---|---|---|
| GO:0050793 | regulation of developmental process | Biological Process | 132 | -0.406 | -1.741 | 0 | 0.0237 | tags=27%, list=14%, signal=31% |
| GO:0007015 | actin filament organization | Biological Process | 74 | -0.441 | -1.739 | 0.0031 | 0.0242 | tags=39%, list=21%, signal=50% |
| GO:0042023 | DNA endoreduplication | Biological Process | 18 | -0.602 | -1.736 | 0.0101 | 0.0247 | tags=56%, list=16%, signal=66% |
| GO:0048367 | shoot system development | Biological Process | 173 | -0.393 | -1.735 | 0 | 0.0248 | tags=36%, list=20%, signal=45% |
| GO:0033043 | regulation of organelle organization | Biological Process | 57 | -0.461 | -1.730 | 0 | 0.0258 | tags=37%, list=18%, signal=45% |
| GO:0016102 | diterpenoid biosynthetic process | Biological Process | 21 | -0.576 | -1.728 | 0.0016 | 0.0262 | tags=52%, list=19%, signal=64% |
| GO:0090567 | reproductive shoot system development | Biological Process | 66 | -0.451 | -1.727 | 0 | 0.0264 | tags=27%, list=11%, signal=30% |
| GO:0005694 | chromosome | Cellular Component | 373 | -0.363 | -1.726 | 0 | 0.0262 | tags=34%, list=21%, signal=43% |
| GO:1901990 | regulation of mitotic cell cycle phase transition | Biological Process | 33 | -0.514 | -1.718 | 0.0101 | 0.0282 | tags=48%, list=22%, signal=62% |
| GO:0008810 | cellulase activity | Molecular Function | 15 | -0.625 | -1.715 | 0.0055 | 0.0287 | tags=60%, list=18%, signal=73% |
| GO:0070652 | HAUS complex | Cellular Component | 15 | -0.624 | -1.711 | 0.0017 | 0.0296 | tags=73%, list=30%, signal=105% |
| GO:0048767 | root hair elongation | Biological Process | 19 | -0.593 | -1.707 | 0.0071 | 0.0307 | tags=42%, list=14%, signal=49% |
| GO:0016298 | lipase activity | Molecular Function | 68 | -0.446 | -1.702 | 0 | 0.0320 | tags=49%, list=27%, signal=67% |
| GO:0016838 | carbon-oxygen lyase activity, acting on phosphates | Molecular Function | 21 | -0.562 | -1.698 | 0.0106 | 0.0329 | tags=52%, list=23%, signal=68% |
| GO:0048364 | root development | Biological Process | 81 | -0.430 | -1.697 | 0.0015 | 0.0329 | tags=31%, list=15%, signal=36% |
| GO:0000785 | chromatin | Cellular Component | 199 | -0.378 | -1.695 | 0 | 0.0331 | tags=37%, list=21%, signal=47% |
| GO:0031347 | regulation of defense response | Biological Process | 66 | -0.438 | -1.695 | 0.0031 | 0.0329 | tags=50%, list=29%, signal=70% |
| GO:0120569 | phospholipase activity | Molecular Function | 44 | -0.473 | -1.684 | 0.0078 | 0.0362 | tags=52%, list=27%, signal=72% |
| GO:0022622 | root system development | Biological Process | 82 | -0.422 | -1.683 | 0.0032 | 0.0362 | tags=30%, list=15%, signal=36% |

| GO ID | GO name | Category | Size | ES | NES | Nominal P | FDR q | Leading edge |
|---|---|---|---|---|---|---|---|---|
| GO:0070161 | anchoring junction | Cellular Component | 73 | -0.425 | -1.678 | 0.0047 | 0.0376 | tags=34%, list=17%, signal=41% |
| GO:0043531 | ADP binding | Molecular Function | 416 | -0.346 | -1.673 | 0 | 0.0389 | tags=43%, list=32%, signal=62% |
| GO:0010054 | trichoblast differentiation | Biological Process | 22 | -0.541 | -1.672 | 0.0133 | 0.0392 | tags=36%, list=14%, signal=42% |
| GO:0009653 | anatomical structure morphogenesis | Biological Process | 184 | -0.374 | -1.672 | 0 | 0.0389 | tags=30%, list=21%, signal=38% |
| GO:0072330 | monocarboxylic acid biosynthetic process | Biological Process | 141 | -0.388 | -1.670 | 0 | 0.0390 | tags=40%, list=27%, signal=55% |
| GO:0048589 | developmental growth | Biological Process | 68 | -0.436 | -1.669 | 0.0016 | 0.0390 | tags=34%, list=21%, signal=43% |
| GO:0030036 | actin cytoskeleton organization | Biological Process | 79 | -0.424 | -1.669 | 0 | 0.0390 | tags=38%, list=21%, signal=48% |
| GO:0048437 | floral organ development | Biological Process | 28 | -0.521 | -1.668 | 0.0117 | 0.0390 | tags=29%, list=10%, signal=32% |
| GO:0055044 | symplast | Cellular Component | 73 | -0.425 | -1.666 | 0.0047 | 0.0392 | tags=34%, list=17%, signal=41% |
| GO:0009611 | response to wounding | Biological Process | 25 | -0.537 | -1.663 | 0.0067 | 0.0401 | tags=68%, list=27%, signal=93% |
| GO:0009506 | plasmodesma | Cellular Component | 73 | -0.425 | -1.662 | 0.0016 | 0.0401 | tags=34%, list=17%, signal=41% |
| GO:0098687 | chromosomal region | Cellular Component | 37 | -0.492 | -1.661 | 0.0050 | 0.0402 | tags=38%, list=17%, signal=45% |
| GO:0030029 | actin filament-based process | Biological Process | 87 | -0.416 | -1.660 | 0.0015 | 0.0401 | tags=37%, list=20%, signal=46% |
| GO:0004553 | hydrolase activity, hydrolyzing O-glycosyl compounds | Molecular Function | 306 | -0.351 | -1.657 | 0 | 0.0412 | tags=27%, list=15%, signal=32% |
| GO:2000280 | regulation of root development | Biological Process | 16 | -0.600 | -1.653 | 0.0124 | 0.0426 | tags=44%, list=14%, signal=51% |
| GO:0048764 | trichoblast maturation | Biological Process | 22 | -0.541 | -1.650 | 0.0147 | 0.0431 | tags=36%, list=14%, signal=42% |
| GO:0030054 | cell junction | Cellular Component | 73 | -0.425 | -1.649 | 0.0031 | 0.0432 | tags=34%, list=17%, signal=41% |
| GO:0048765 | root hair cell differentiation | Biological Process | 22 | -0.541 | -1.648 | 0.0105 | 0.0434 | tags=36%, list=14%, signal=42% |
| GO:0061572 | actin filament bundle organization | Biological Process | 15 | -0.596 | -1.648 | 0.0160 | 0.0435 | tags=80%, list=24%, signal=106% |

| GO ID | GO name | Category | Size | ES | NES | Nominal P | FDR q | Leading edge |
|---|---|---|---|---|---|---|---|---|
| GO:0007389 | pattern specification process | Biological Process | 30 | -0.506 | -1.644 | 0.0134 | 0.0446 | tags=30%, list=13%, signal=34% |
| GO:0080147 | root hair cell development | Biological Process | 22 | -0.541 | -1.643 | 0.0164 | 0.0447 | tags=36%, list=14%, signal=42% |
| GO:0030261 | chromosome condensation | Biological Process | 19 | -0.560 | -1.643 | 0.0160 | 0.0445 | tags=58%, list=22%, signal=74% |
| GO:0007264 | small GTPase-mediated signal transduction | Biological Process | 32 | -0.495 | -1.642 | 0.0102 | 0.0443 | tags=38%, list=16%, signal=45% |
| GO:0009933 | meristem structural organization | Biological Process | 18 | -0.580 | -1.639 | 0.0139 | 0.0456 | tags=33%, list=10%, signal=37% |
| GO:0005911 | cell-cell junction | Cellular Component | 73 | -0.425 | -1.635 | 0.0015 | 0.0466 | tags=34%, list=17%, signal=41% |
| GO:0001727 | lipid kinase activity | Molecular Function | 32 | -0.487 | -1.635 | 0.0095 | 0.0463 | tags=41%, list=21%, signal=51% |
| GO:1902679 | negative regulation of RNA biosynthetic process | Biological Process | 65 | -0.421 | -1.634 | 0.0031 | 0.0463 | tags=40%, list=25%, signal=53% |
| GO:0052742 | phosphatidylinositol kinase activity | Molecular Function | 20 | -0.555 | -1.630 | 0.0243 | 0.0476 | tags=45%, list=19%, signal=56% |
| GO:0009606 | tropism | Biological Process | 34 | -0.479 | -1.630 | 0.0048 | 0.0474 | tags=47%, list=22%, signal=60% |
| GO:0016101 | diterpenoid metabolic process | Biological Process | 27 | -0.515 | -1.630 | 0.0185 | 0.0472 | tags=44%, list=19%, signal=55% |
| GO:0006270 | DNA replication initiation | Biological Process | 37 | -0.472 | -1.628 | 0.0066 | 0.0476 | tags=62%, list=31%, signal=91% |
| GO:0005388 | P-type calcium transporter activity | Molecular Function | 21 | -0.551 | -1.624 | 0.0139 | 0.0489 | tags=67%, list=37%, signal=105% |
| GO:0048469 | cell maturation | Biological Process | 22 | -0.541 | -1.624 | 0.0140 | 0.0488 | tags=36%, list=14%, signal=42% |
| GO:0051017 | actin filament bundle assembly | Biological Process | 15 | -0.596 | -1.624 | 0.0205 | 0.0487 | tags=80%, list=24%, signal=106% |
| GO:0006260 | DNA replication | Biological Process | 167 | -0.368 | -1.623 | 0 | 0.0487 | tags=37%, list=25%, signal=49% |
| **Not significantly enriched (FDR q ≥ 0.05; *n* = 359)** | | | | | | | | |
| GO:0009311 | oligosaccharide metabolic process | Biological Process | 59 | 0.401 | 1.665 | 0.0026 | 0.0505 | tags=32%, list=15%, signal=38% |
| GO:0051540 | metal cluster binding | Molecular Function | 104 | 0.370 | 1.664 | 0.0029 | 0.0506 | tags=34%, list=18%, signal=41% |

| GO ID | GO name | Category | Size | ES | NES | Nominal P | FDR q | Leading edge |
|---|---|---|---|---|---|---|---|---|
| GO:0008652 | amino acid biosynthetic process | Biological Process | 132 | 0.358 | 1.661 | 0 | 0.0514 | tags=35%, list=19%, signal=43% |
| GO:0016778 | diphosphotransferase activity | Molecular Function | 15 | 0.576 | 1.660 | 0.0217 | 0.0514 | tags=40%, list=16%, signal=47% |
| GO:0140223 | general transcription initiation factor activity | Molecular Function | 16 | 0.553 | 1.650 | 0.0140 | 0.0550 | tags=75%, list=33%, signal=112% |
| GO:0001522 | pseudouridine synthesis | Biological Process | 20 | 0.531 | 1.650 | 0.0173 | 0.0551 | tags=45%, list=24%, signal=59% |
| GO:0009522 | photosystem I | Cellular Component | 19 | 0.533 | 1.650 | 0.0196 | 0.0548 | tags=58%, list=35%, signal=89% |
| GO:0016668 | oxidoreductase activity, acting on a sulfur group of donors, NAD(P) as acceptor | Molecular Function | 21 | 0.521 | 1.649 | 0.0213 | 0.0545 | tags=57%, list=21%, signal=72% |
| GO:0047617 | fatty acyl-CoA hydrolase activity | Molecular Function | 21 | 0.519 | 1.649 | 0.0285 | 0.0544 | tags=52%, list=27%, signal=72% |
| GO:1901259 | chloroplast rRNA processing | Biological Process | 15 | 0.560 | 1.648 | 0.0205 | 0.0545 | tags=40%, list=10%, signal=44% |
| GO:0016591 | RNA polymerase II, holoenzyme | Cellular Component | 73 | 0.387 | 1.647 | 0 | 0.0545 | tags=58%, list=35%, signal=89% |
| GO:0003713 | transcription coactivator activity | Molecular Function | 27 | 0.488 | 1.647 | 0.0132 | 0.0544 | tags=48%, list=26%, signal=65% |
| GO:0035999 | tetrahydrofolate interconversion | Biological Process | 19 | 0.514 | 1.636 | 0.0180 | 0.0584 | tags=47%, list=14%, signal=55% |
| GO:0016051 | carbohydrate biosynthetic process | Biological Process | 191 | 0.322 | 1.634 | 0 | 0.0591 | tags=30%, list=18%, signal=36% |
| GO:0061919 | process utilizing autophagic mechanism | Biological Process | 63 | 0.386 | 1.633 | 0.0080 | 0.0592 | tags=41%, list=24%, signal=54% |
| GO:0008643 | carbohydrate transport | Biological Process | 61 | 0.398 | 1.628 | 0.0029 | 0.0612 | tags=31%, list=16%, signal=37% |
| GO:0106074 | aminoacyl-tRNA metabolism involved in translational fidelity | Biological Process | 16 | 0.547 | 1.627 | 0.0167 | 0.0617 | tags=56%, list=30%, signal=80% |
| GO:1990542 | mitochondrial transmembrane transport | Biological Process | 35 | 0.458 | 1.626 | 0.0257 | 0.0614 | tags=51%, list=28%, signal=71% |
| GO:0071482 | cellular response to light stimulus | Biological Process | 38 | 0.429 | 1.624 | 0.0075 | 0.0624 | tags=58%, list=29%, signal=82% |
| GO:0006066 | alcohol metabolic process | Biological Process | 87 | 0.371 | 1.623 | 0.0029 | 0.0623 | tags=30%, list=14%, signal=35% |
| GO:0046165 | alcohol biosynthetic process | Biological Process | 32 | 0.460 | 1.623 | 0.0288 | 0.0621 | tags=41%, list=12%, signal=46% |

| GO ID | GO name | Category | Size | ES | NES | Nominal P | FDR q | Leading edge |
|---|---|---|---|---|---|---|---|---|
| GO:0015291 | secondary active transmembrane transporter activity | Molecular Function | 189 | 0.322 | 1.619 | 0 | 0.0637 | tags=39%, list=23%, signal=51% |
| GO:0005736 | RNA polymerase I complex | Cellular Component | 19 | 0.517 | 1.615 | 0.0284 | 0.0652 | tags=37%, list=18%, signal=45% |
| GO:0006672 | ceramide metabolic process | Biological Process | 27 | 0.472 | 1.615 | 0.0167 | 0.0649 | tags=33%, list=12%, signal=38% |
| GO:0006413 | translational initiation | Biological Process | 94 | 0.356 | 1.611 | 0 | 0.0664 | tags=31%, list=23%, signal=40% |
| GO:0050997 | quaternary ammonium group binding | Molecular Function | 15 | 0.554 | 1.610 | 0.0319 | 0.0665 | tags=67%, list=20%, signal=84% |
| GO:0015293 | symporter activity | Molecular Function | 22 | 0.498 | 1.610 | 0.0314 | 0.0665 | tags=45%, list=19%, signal=56% |
| GO:0015144 | carbohydrate transmembrane transporter activity | Molecular Function | 60 | 0.389 | 1.609 | 0.0029 | 0.0662 | tags=37%, list=22%, signal=47% |
| GO:0006914 | autophagy | Biological Process | 63 | 0.386 | 1.607 | 0.0056 | 0.0672 | tags=41%, list=24%, signal=54% |
| GO:0006760 | folic acid-containing compound metabolic process | Biological Process | 30 | 0.463 | 1.598 | 0.0103 | 0.0713 | tags=37%, list=14%, signal=42% |
| GO:0004222 | metalloendopeptidase activity | Molecular Function | 48 | 0.414 | 1.595 | 0.0106 | 0.0728 | tags=38%, list=20%, signal=47% |
| GO:0002161 | aminoacyl-tRNA deacylase activity | Molecular Function | 16 | 0.547 | 1.595 | 0.0228 | 0.0724 | tags=56%, list=30%, signal=80% |
| GO:0120114 | Sm-like protein family complex | Cellular Component | 84 | 0.366 | 1.593 | 0.0058 | 0.0732 | tags=49%, list=36%, signal=76% |
| GO:0030532 | small nuclear ribonucleoprotein complex | Cellular Component | 75 | 0.369 | 1.591 | 0.0115 | 0.0735 | tags=47%, list=35%, signal=71% |
| GO:0009451 | RNA modification | Biological Process | 367 | 0.298 | 1.589 | 0 | 0.0740 | tags=39%, list=31%, signal=56% |
| GO:0051604 | protein maturation | Biological Process | 328 | 0.300 | 1.584 | 0 | 0.0765 | tags=32%, list=25%, signal=42% |
| GO:0034702 | monoatomic ion channel complex | Cellular Component | 28 | 0.461 | 1.582 | 0 | 0.0777 | tags=61%, list=35%, signal=93% |
| GO:0006839 | mitochondrial transport | Biological Process | 55 | 0.394 | 1.579 | 0.0026 | 0.0790 | tags=53%, list=32%, signal=78% |
| GO:0000463 | maturation of LSU-rRNA from tricistronic rRNA transcript (SSU-rRNA, 5.8S rRNA, LSU-rRNA) | Biological Process | 20 | 0.501 | 1.576 | 0.0453 | 0.0802 | tags=55%, list=32%, signal=80% |
| GO:0016485 | protein processing | Biological Process | 28 | 0.460 | 1.576 | 0.0213 | 0.0799 | tags=36%, list=18%, signal=44% |

| GO ID | GO name | Category | Size | ES | NES | Nominal P | FDR q | Leading edge |
|---|---|---|---|---|---|---|---|---|
| GO:0010257 | NADH dehydrogenase complex assembly | Biological Process | 20 | 0.497 | 1.576 | 0.0239 | 0.0796 | tags=40%, list=24%, signal=53% |
| GO:0000096 | sulfur amino acid metabolic process | Biological Process | 28 | 0.451 | 1.572 | 0.0199 | 0.0814 | tags=50%, list=19%, signal=62% |
| GO:0034703 | cation channel complex | Cellular Component | 28 | 0.461 | 1.572 | 0.0263 | 0.0812 | tags=61%, list=35%, signal=93% |
| GO:0006457 | protein folding | Biological Process | 210 | 0.312 | 1.572 | 0 | 0.0810 | tags=31%, list=25%, signal=42% |
| GO:0009142 | nucleoside triphosphate biosynthetic process | Biological Process | 44 | 0.402 | 1.568 | 0.0054 | 0.0831 | tags=75%, list=41%, signal=128% |
| GO:0003743 | translation initiation factor activity | Molecular Function | 88 | 0.351 | 1.567 | 0.0062 | 0.0828 | tags=32%, list=24%, signal=42% |
| GO:0071478 | cellular response to radiation | Biological Process | 39 | 0.420 | 1.566 | 0.0099 | 0.0833 | tags=56%, list=29%, signal=80% |
| GO:0022900 | electron transport chain | Biological Process | 98 | 0.353 | 1.565 | 0.0060 | 0.0834 | tags=32%, list=24%, signal=41% |
| GO:0000470 | maturation of LSU-rRNA | Biological Process | 31 | 0.444 | 1.554 | 0.0284 | 0.0897 | tags=58%, list=33%, signal=86% |
| GO:0016762 | xyloglucan:xyloglucosyl transferase activity | Molecular Function | 27 | 0.461 | 1.554 | 0.0290 | 0.0894 | tags=26%, list=8%, signal=28% |
| GO:0170033 | L-amino acid metabolic process | Biological Process | 204 | 0.310 | 1.552 | 0 | 0.0906 | tags=32%, list=19%, signal=39% |
| GO:0004812 | aminoacyl-tRNA ligase activity | Molecular Function | 76 | 0.358 | 1.551 | 0.0029 | 0.0902 | tags=42%, list=30%, signal=60% |
| GO:0007005 | mitochondrion organization | Biological Process | 105 | 0.350 | 1.551 | 0.0033 | 0.0899 | tags=42%, list=32%, signal=61% |
| GO:0051139 | metal cation:proton antiporter activity | Molecular Function | 33 | 0.434 | 1.551 | 0.0179 | 0.0897 | tags=48%, list=23%, signal=63% |
| GO:0006418 | tRNA aminoacylation for protein translation | Biological Process | 76 | 0.358 | 1.548 | 0 | 0.0910 | tags=42%, list=30%, signal=60% |
| GO:0140828 | metal cation:monoatomic cation antiporter activity | Molecular Function | 35 | 0.423 | 1.547 | 0.0280 | 0.0915 | tags=46%, list=23%, signal=59% |
| GO:0043038 | amino acid activation | Biological Process | 79 | 0.360 | 1.545 | 0.0057 | 0.0921 | tags=42%, list=30%, signal=59% |
| GO:0008237 | metallopeptidase activity | Molecular Function | 87 | 0.352 | 1.544 | 0.0031 | 0.0923 | tags=34%, list=23%, signal=44% |
| GO:0170034 | L-amino acid biosynthetic process | Biological Process | 106 | 0.337 | 1.538 | 0 | 0.0954 | tags=33%, list=18%, signal=40% |

| GO ID | GO name | Category | Size | ES | NES | Nominal P | FDR q | Leading edge |
|---|---|---|---|---|---|---|---|---|
| GO:1990204 | oxidoreductase complex | Cellular Component | 118 | 0.328 | 1.536 | 0.0031 | 0.0967 | tags=43%, list=28%, signal=60% |
| GO:0006002 | fructose 6-phosphate metabolic process | Biological Process | 29 | 0.449 | 1.535 | 0.0254 | 0.0967 | tags=62%, list=30%, signal=89% |
| GO:0009201 | ribonucleoside triphosphate biosynthetic process | Biological Process | 44 | 0.402 | 1.533 | 0.0103 | 0.0975 | tags=75%, list=41%, signal=128% |
| GO:0016604 | nuclear body | Cellular Component | 35 | 0.423 | 1.531 | 0.0177 | 0.0990 | tags=57%, list=25%, signal=76% |
| GO:1902555 | endoribonuclease complex | Cellular Component | 17 | 0.506 | 1.531 | 0.0413 | 0.0986 | tags=65%, list=38%, signal=104% |
| GO:0009408 | response to heat | Biological Process | 81 | 0.355 | 1.529 | 0.0151 | 0.0989 | tags=36%, list=21%, signal=45% |
| GO:0015294 | solute:monoatomic cation symporter activity | Molecular Function | 21 | 0.480 | 1.526 | 0.0374 | 0.1010 | tags=43%, list=19%, signal=53% |
| GO:0016859 | cis-trans isomerase activity | Molecular Function | 83 | 0.351 | 1.525 | 0.0058 | 0.1011 | tags=39%, list=24%, signal=50% |
| GO:0043039 | tRNA aminoacylation | Biological Process | 79 | 0.360 | 1.524 | 0.0089 | 0.1012 | tags=42%, list=30%, signal=59% |
| GO:0006520 | amino acid metabolic process | Biological Process | 378 | 0.285 | 1.522 | 0 | 0.1022 | tags=32%, list=22%, signal=41% |
| GO:0055074 | calcium ion homeostasis | Biological Process | 41 | 0.401 | 1.520 | 0.0284 | 0.1033 | tags=34%, list=16%, signal=41% |
| GO:0016875 | ligase activity, forming carbon-oxygen bonds | Molecular Function | 76 | 0.358 | 1.517 | 0 | 0.1047 | tags=42%, list=30%, signal=60% |
| GO:0009668 | plastid membrane organization | Biological Process | 16 | 0.522 | 1.517 | 0.0337 | 0.1047 | tags=56%, list=22%, signal=72% |
| GO:0009570 | chloroplast stroma | Cellular Component | 171 | 0.311 | 1.516 | 0.0035 | 0.1044 | tags=36%, list=21%, signal=45% |
| GO:0005654 | nucleoplasm | Cellular Component | 247 | 0.301 | 1.514 | 0 | 0.1057 | tags=43%, list=29%, signal=60% |
| GO:0016289 | acyl-CoA hydrolase activity | Molecular Function | 24 | 0.456 | 1.512 | 0.0370 | 0.1068 | tags=46%, list=27%, signal=63% |
| GO:0008138 | protein tyrosine/serine/threonine phosphatase activity | Molecular Function | 24 | 0.451 | 1.511 | 0.0319 | 0.1067 | tags=29%, list=14%, signal=34% |
| GO:0016811 | hydrolase activity, acting on carbon-nitrogen (but not peptide) bonds, in linear amides | Molecular Function | 53 | 0.381 | 1.510 | 0.0231 | 0.1077 | tags=34%, list=17%, signal=41% |
| GO:0072594 | establishment of protein localization to organelle | Biological Process | 165 | 0.312 | 1.508 | 0.0036 | 0.1082 | tags=43%, list=33%, signal=64% |

| GO ID | GO name | Category | Size | ES | NES | Nominal P | FDR q | Leading edge |
|---|---|---|---|---|---|---|---|---|
| GO:0006400 | tRNA modification | Biological Process | 88 | 0.346 | 1.504 | 0.0028 | 0.1108 | tags=36%, list=23%, signal=47% |
| GO:0006874 | intracellular calcium ion homeostasis | Biological Process | 41 | 0.401 | 1.504 | 0.0341 | 0.1105 | tags=34%, list=16%, signal=41% |
| GO:0031647 | regulation of protein stability | Biological Process | 19 | 0.476 | 1.501 | 0.0376 | 0.1123 | tags=16%, list=4%, signal=16% |
| GO:0140168 | nuclear ribonucleoprotein granule | Cellular Component | 31 | 0.416 | 1.501 | 0.0257 | 0.1120 | tags=58%, list=25%, signal=77% |
| GO:0030880 | RNA polymerase complex | Cellular Component | 130 | 0.315 | 1.497 | 0 | 0.1143 | tags=52%, list=35%, signal=79% |
| GO:0097526 | spliceosomal tri-snRNP complex | Cellular Component | 38 | 0.409 | 1.496 | 0.0263 | 0.1147 | tags=53%, list=36%, signal=82% |
| GO:0019209 | kinase activator activity | Molecular Function | 16 | 0.494 | 1.494 | 0.0394 | 0.1162 | tags=38%, list=10%, signal=42% |
| GO:0055029 | nuclear DNA-directed RNA polymerase complex | Cellular Component | 82 | 0.346 | 1.493 | 0.0113 | 0.1160 | tags=55%, list=35%, signal=84% |
| GO:0042273 | ribosomal large subunit biogenesis | Biological Process | 73 | 0.349 | 1.493 | 0.0216 | 0.1157 | tags=52%, list=33%, signal=78% |
| GO:0015386 | potassium:proton antiporter activity | Molecular Function | 15 | 0.508 | 1.493 | 0.0562 | 0.1153 | tags=60%, list=23%, signal=77% |
| GO:0050821 | protein stabilization | Biological Process | 15 | 0.514 | 1.488 | 0.0639 | 0.1191 | tags=20%, list=4%, signal=21% |
| GO:0034219 | carbohydrate transmembrane transport | Biological Process | 54 | 0.382 | 1.486 | 0.0245 | 0.1201 | tags=35%, list=22%, signal=45% |
| GO:0000387 | spliceosomal snRNP assembly | Biological Process | 22 | 0.451 | 1.485 | 0.0258 | 0.1204 | tags=45%, list=35%, signal=70% |
| GO:0009532 | plastid stroma | Cellular Component | 175 | 0.299 | 1.484 | 0 | 0.1203 | tags=35%, list=21%, signal=44% |
| GO:0005778 | peroxisomal membrane | Cellular Component | 35 | 0.400 | 1.482 | 0.0280 | 0.1215 | tags=46%, list=23%, signal=59% |
| GO:0004725 | protein tyrosine phosphatase activity | Molecular Function | 36 | 0.399 | 1.480 | 0.0286 | 0.1226 | tags=31%, list=14%, signal=35% |
| GO:0016597 | amino acid binding | Molecular Function | 22 | 0.454 | 1.479 | 0.0402 | 0.1236 | tags=32%, list=15%, signal=37% |
| GO:0007034 | vacuolar transport | Biological Process | 74 | 0.349 | 1.477 | 0.0139 | 0.1247 | tags=51%, list=35%, signal=78% |
| GO:0009251 | glucan catabolic process | Biological Process | 18 | 0.467 | 1.472 | 0.0587 | 0.1277 | tags=22%, list=4%, signal=23% |

| GO ID | GO name | Category | Size | ES | NES | Nominal P | FDR q | Leading edge |
|---|---|---|---|---|---|---|---|---|
| GO:0008324 | monoatomic cation transmembrane transporter activity | Molecular Function | 339 | 0.280 | 1.472 | 0 | 0.1279 | tags=32%, list=23%, signal=41% |
| GO:0005759 | mitochondrial matrix | Cellular Component | 112 | 0.320 | 1.470 | 0.0094 | 0.1289 | tags=31%, list=21%, signal=39% |
| GO:0032981 | mitochondrial respiratory chain complex I assembly | Biological Process | 19 | 0.475 | 1.470 | 0.0696 | 0.1285 | tags=37%, list=24%, signal=49% |
| GO:0009657 | plastid organization | Biological Process | 78 | 0.341 | 1.467 | 0.0169 | 0.1301 | tags=49%, list=30%, signal=69% |
| GO:0031903 | microbody membrane | Cellular Component | 35 | 0.400 | 1.465 | 0.0446 | 0.1314 | tags=46%, list=23%, signal=59% |
| GO:0035770 | ribonucleoprotein granule | Cellular Component | 68 | 0.347 | 1.464 | 0.0145 | 0.1322 | tags=50%, list=29%, signal=70% |
| GO:0006730 | one-carbon metabolic process | Biological Process | 26 | 0.428 | 1.463 | 0.0480 | 0.1326 | tags=38%, list=14%, signal=45% |
| GO:0015075 | monoatomic ion transmembrane transporter activity | Molecular Function | 397 | 0.271 | 1.462 | 0 | 0.1326 | tags=31%, list=23%, signal=40% |
| GO:0043021 | ribonucleoprotein complex binding | Molecular Function | 68 | 0.348 | 1.459 | 0.0248 | 0.1351 | tags=41%, list=27%, signal=56% |
| GO:0022834 | ligand-gated channel activity | Molecular Function | 25 | 0.434 | 1.453 | 0.0576 | 0.1399 | tags=40%, list=24%, signal=53% |
| GO:0070566 | adenylyltransferase activity | Molecular Function | 27 | 0.422 | 1.448 | 0.0653 | 0.1438 | tags=44%, list=25%, signal=60% |
| GO:0033108 | mitochondrial respiratory chain complex assembly | Biological Process | 44 | 0.379 | 1.445 | 0.0358 | 0.1459 | tags=36%, list=24%, signal=48% |
| GO:0070585 | protein localization to mitochondrion | Biological Process | 29 | 0.412 | 1.444 | 0.0400 | 0.1466 | tags=52%, list=29%, signal=72% |
| GO:0015145 | monosaccharide transmembrane transporter activity | Molecular Function | 21 | 0.450 | 1.442 | 0.0649 | 0.1482 | tags=29%, list=15%, signal=34% |
| GO:0005682 | U5 snRNP | Cellular Component | 18 | 0.478 | 1.441 | 0.0634 | 0.1484 | tags=50%, list=40%, signal=83% |
| GO:0007006 | mitochondrial membrane organization | Biological Process | 23 | 0.438 | 1.438 | 0.0578 | 0.1514 | tags=61%, list=37%, signal=96% |
| GO:0098798 | mitochondrial protein-containing complex | Cellular Component | 105 | 0.312 | 1.435 | 0.0259 | 0.1539 | tags=42%, list=29%, signal=59% |
| GO:0046540 | U4/U6 x U5 tri-snRNP complex | Cellular Component | 30 | 0.408 | 1.434 | 0.0771 | 0.1540 | tags=53%, list=35%, signal=82% |
| GO:0019693 | ribose phosphate metabolic process | Biological Process | 188 | 0.288 | 1.434 | 0 | 0.1534 | tags=34%, list=22%, signal=43% |

| GO ID | GO name | Category | Size | ES | NES | Nominal P | FDR q | Leading edge |
|---|---|---|---|---|---|---|---|---|
| GO:0009414 | response to water deprivation | Biological Process | 66 | 0.343 | 1.433 | 0.0424 | 0.1540 | tags=39%, list=21%, signal=49% |
| GO:0015980 | energy derivation by oxidation of organic compounds | Biological Process | 225 | 0.283 | 1.432 | 0 | 0.1538 | tags=34%, list=23%, signal=44% |
| GO:0015749 | monosaccharide transmembrane transport | Biological Process | 21 | 0.450 | 1.431 | 0.0655 | 0.1546 | tags=29%, list=15%, signal=34% |
| GO:0031667 | response to nutrient levels | Biological Process | 59 | 0.342 | 1.431 | 0.0184 | 0.1540 | tags=34%, list=24%, signal=44% |
| GO:0016854 | racemase and epimerase activity | Molecular Function | 32 | 0.389 | 1.431 | 0.0502 | 0.1538 | tags=41%, list=17%, signal=49% |
| GO:0043022 | ribosome binding | Molecular Function | 47 | 0.374 | 1.429 | 0.0300 | 0.1547 | tags=40%, list=24%, signal=53% |
| GO:0005685 | U1 snRNP | Cellular Component | 24 | 0.431 | 1.428 | 0.0571 | 0.1556 | tags=42%, list=35%, signal=64% |
| GO:0009415 | response to water | Biological Process | 66 | 0.343 | 1.428 | 0.0248 | 0.1553 | tags=39%, list=21%, signal=49% |
| GO:0072528 | pyrimidine-containing compound biosynthetic process | Biological Process | 47 | 0.369 | 1.425 | 0.0421 | 0.1573 | tags=32%, list=18%, signal=39% |
| GO:0006740 | NADPH regeneration | Biological Process | 30 | 0.402 | 1.423 | 0.0539 | 0.1593 | tags=37%, list=17%, signal=44% |
| GO:0015276 | ligand-gated monoatomic ion channel activity | Molecular Function | 25 | 0.434 | 1.421 | 0.0639 | 0.1603 | tags=40%, list=24%, signal=53% |
| GO:0003755 | peptidyl-prolyl cis-trans isomerase activity | Molecular Function | 76 | 0.331 | 1.418 | 0.0249 | 0.1632 | tags=38%, list=24%, signal=50% |
| GO:0022821 | solute:potassium antiporter activity | Molecular Function | 16 | 0.493 | 1.417 | 0.0736 | 0.1635 | tags=56%, list=23%, signal=73% |
| GO:0015252 | proton channel activity | Molecular Function | 18 | 0.467 | 1.416 | 0.0764 | 0.1641 | tags=61%, list=34%, signal=92% |
| GO:0042546 | cell wall biogenesis | Biological Process | 83 | 0.325 | 1.414 | 0.0268 | 0.1650 | tags=27%, list=15%, signal=31% |
| GO:0016860 | intramolecular oxidoreductase activity | Molecular Function | 54 | 0.358 | 1.411 | 0.0318 | 0.1677 | tags=44%, list=20%, signal=55% |
| GO:0006360 | transcription by RNA polymerase I | Biological Process | 26 | 0.420 | 1.410 | 0.0754 | 0.1678 | tags=46%, list=23%, signal=60% |
| GO:0006098 | pentose-phosphate shunt | Biological Process | 30 | 0.402 | 1.410 | 0.0643 | 0.1682 | tags=37%, list=17%, signal=44% |
| GO:0016861 | intramolecular oxidoreductase activity, interconverting aldoses and ketoses | Molecular Function | 22 | 0.439 | 1.408 | 0.0704 | 0.1698 | tags=50%, list=17%, signal=60% |

| GO ID | GO name | Category | Size | ES | NES | Nominal P | FDR q | Leading edge |
|---|---|---|---|---|---|---|---|---|
| GO:0042364 | water-soluble vitamin biosynthetic process | Biological Process | 41 | 0.374 | 1.407 | 0.0566 | 0.1695 | tags=37%, list=19%, signal=45% |
| GO:0001101 | response to acid chemical | Biological Process | 70 | 0.332 | 1.406 | 0.0220 | 0.1703 | tags=39%, list=21%, signal=48% |
| GO:0022618 | protein-RNA complex assembly | Biological Process | 150 | 0.289 | 1.404 | 0.0134 | 0.1715 | tags=37%, list=35%, signal=57% |
| GO:0000966 | RNA 5'-end processing | Biological Process | 16 | 0.476 | 1.395 | 0.0903 | 0.1808 | tags=63%, list=32%, signal=92% |
| GO:0005253 | monoatomic anion channel activity | Molecular Function | 19 | 0.457 | 1.395 | 0.0885 | 0.1803 | tags=63%, list=29%, signal=89% |
| GO:0045333 | cellular respiration | Biological Process | 210 | 0.281 | 1.395 | 0.0039 | 0.1800 | tags=33%, list=23%, signal=42% |
| GO:0016810 | hydrolase activity, acting on carbon-nitrogen (but not peptide) bonds | Molecular Function | 109 | 0.307 | 1.393 | 0.0157 | 0.1809 | tags=25%, list=13%, signal=28% |
| GO:0006188 | IMP biosynthetic process | Biological Process | 24 | 0.416 | 1.392 | 0.0819 | 0.1820 | tags=33%, list=17%, signal=40% |
| GO:0015368 | calcium:monoatomic cation antiporter activity | Molecular Function | 17 | 0.455 | 1.390 | 0.0848 | 0.1838 | tags=41%, list=16%, signal=49% |
| GO:0070279 | vitamin B6 binding | Molecular Function | 81 | 0.317 | 1.388 | 0.0093 | 0.1846 | tags=37%, list=21%, signal=47% |
| GO:0003727 | single-stranded RNA binding | Molecular Function | 22 | 0.424 | 1.388 | 0.0754 | 0.1848 | tags=50%, list=26%, signal=67% |
| GO:0065003 | protein-containing complex assembly | Biological Process | 387 | 0.257 | 1.387 | 0 | 0.1850 | tags=32%, list=26%, signal=42% |
| GO:0004386 | helicase activity | Molecular Function | 137 | 0.290 | 1.387 | 0.0069 | 0.1846 | tags=36%, list=22%, signal=47% |
| GO:0000209 | protein polyubiquitination | Biological Process | 49 | 0.350 | 1.386 | 0.0533 | 0.1845 | tags=45%, list=30%, signal=64% |
| GO:0043094 | metabolic compound salvage | Biological Process | 44 | 0.364 | 1.386 | 0.0761 | 0.1850 | tags=34%, list=16%, signal=40% |
| GO:0006163 | purine nucleotide metabolic process | Biological Process | 221 | 0.277 | 1.385 | 0.0038 | 0.1851 | tags=28%, list=18%, signal=33% |
| GO:0000049 | tRNA binding | Molecular Function | 37 | 0.373 | 1.379 | 0.0697 | 0.1913 | tags=46%, list=31%, signal=66% |
| GO:0030170 | pyridoxal phosphate binding | Molecular Function | 81 | 0.317 | 1.379 | 0.0142 | 0.1913 | tags=37%, list=21%, signal=47% |
| GO:1902600 | proton transmembrane transport | Biological Process | 145 | 0.290 | 1.378 | 0.0074 | 0.1916 | tags=43%, list=30%, signal=61% |

| GO ID | GO name | Category | Size | ES | NES | Nominal P | FDR q | Leading edge |
|---|---|---|---|---|---|---|---|---|
| GO:0016791 | phosphatase activity | Molecular Function | 308 | 0.261 | 1.377 | 0 | 0.1914 | tags=25%, list=15%, signal=29% |
| GO:0006575 | modified amino acid metabolic process | Biological Process | 110 | 0.301 | 1.377 | 0.0216 | 0.1909 | tags=28%, list=16%, signal=33% |
| GO:0006979 | response to oxidative stress | Biological Process | 174 | 0.282 | 1.376 | 0.0211 | 0.1913 | tags=26%, list=18%, signal=32% |
| GO:0031123 | RNA 3'-end processing | Biological Process | 49 | 0.359 | 1.376 | 0.0541 | 0.1913 | tags=53%, list=34%, signal=81% |
| GO:0009117 | nucleotide metabolic process | Biological Process | 265 | 0.266 | 1.375 | 0.0042 | 0.1913 | tags=29%, list=19%, signal=35% |
| GO:0010268 | brassinosteroid homeostasis | Biological Process | 16 | 0.468 | 1.375 | 0.0835 | 0.1907 | tags=44%, list=13%, signal=50% |
| GO:0005771 | multivesicular body | Cellular Component | 20 | 0.435 | 1.372 | 0.0957 | 0.1937 | tags=55%, list=28%, signal=76% |
| GO:0046390 | ribose phosphate biosynthetic process | Biological Process | 97 | 0.311 | 1.372 | 0.0212 | 0.1938 | tags=40%, list=30%, signal=57% |
| GO:0004721 | phosphoprotein phosphatase activity | Molecular Function | 155 | 0.287 | 1.368 | 0.0033 | 0.1977 | tags=25%, list=14%, signal=29% |
| GO:0072350 | tricarboxylic acid metabolic process | Biological Process | 19 | 0.449 | 1.367 | 0.0918 | 0.1983 | tags=53%, list=28%, signal=73% |
| GO:0071826 | protein-RNA complex organization | Biological Process | 157 | 0.281 | 1.367 | 0.0290 | 0.1977 | tags=37%, list=35%, signal=57% |
| GO:0006885 | regulation of pH | Biological Process | 27 | 0.393 | 1.367 | 0.0587 | 0.1978 | tags=63%, list=28%, signal=87% |
| GO:0009060 | aerobic respiration | Biological Process | 199 | 0.275 | 1.365 | 0.0110 | 0.1987 | tags=30%, list=21%, signal=38% |
| GO:0009640 | photomorphogenesis | Biological Process | 26 | 0.405 | 1.364 | 0.0725 | 0.1992 | tags=50%, list=23%, signal=65% |
| GO:0009240 | isopentenyl diphosphate biosynthetic process | Biological Process | 17 | 0.454 | 1.364 | 0.0862 | 0.1987 | tags=41%, list=16%, signal=49% |
| GO:0015369 | calcium:proton antiporter activity | Molecular Function | 16 | 0.463 | 1.364 | 0.1233 | 0.1984 | tags=44%, list=16%, signal=52% |
| GO:0046490 | isopentenyl diphosphate metabolic process | Biological Process | 17 | 0.454 | 1.363 | 0.1023 | 0.1994 | tags=41%, list=16%, signal=49% |
| GO:0005261 | monoatomic cation channel activity | Molecular Function | 54 | 0.339 | 1.361 | 0.0744 | 0.2007 | tags=44%, list=26%, signal=60% |
| GO:0051537 | 2 iron, 2 sulfur cluster binding | Molecular Function | 34 | 0.380 | 1.358 | 0.0969 | 0.2036 | tags=26%, list=11%, signal=30% |

| GO ID | GO name | Category | Size | ES | NES | Nominal P | FDR q | Leading edge |
|---|---|---|---|---|---|---|---|---|
| GO:0051156 | glucose 6-phosphate metabolic process | Biological Process | 37 | 0.368 | 1.355 | 0.0920 | 0.2066 | tags=32%, list=14%, signal=38% |
| GO:0002098 | tRNA wobble uridine modification | Biological Process | 21 | 0.420 | 1.353 | 0.0909 | 0.2084 | tags=62%, list=35%, signal=95% |
| GO:0009658 | chloroplast organization | Biological Process | 55 | 0.334 | 1.353 | 0.0708 | 0.2084 | tags=51%, list=30%, signal=73% |
| GO:0016160 | amylase activity | Molecular Function | 32 | 0.374 | 1.352 | 0.0771 | 0.2083 | tags=47%, list=27%, signal=64% |
| GO:0072521 | purine-containing compound metabolic process | Biological Process | 303 | 0.261 | 1.352 | 0 | 0.2081 | tags=27%, list=19%, signal=33% |
| GO:0009259 | ribonucleotide metabolic process | Biological Process | 181 | 0.275 | 1.352 | 0.0105 | 0.2081 | tags=33%, list=22%, signal=42% |
| GO:0120013 | lipid transfer activity | Molecular Function | 16 | 0.448 | 1.350 | 0.1136 | 0.2099 | tags=56%, list=30%, signal=81% |
| GO:0045239 | tricarboxylic acid cycle heteromeric enzyme complex | Cellular Component | 18 | 0.446 | 1.349 | 0.0990 | 0.2096 | tags=67%, list=30%, signal=95% |
| GO:0016607 | nuclear speck | Cellular Component | 29 | 0.392 | 1.343 | 0.1134 | 0.2178 | tags=55%, list=25%, signal=74% |
| GO:0044042 | glucan metabolic process | Biological Process | 123 | 0.286 | 1.342 | 0.0407 | 0.2180 | tags=27%, list=20%, signal=33% |
| GO:0019842 | vitamin binding | Molecular Function | 118 | 0.291 | 1.340 | 0.0387 | 0.2193 | tags=36%, list=21%, signal=45% |
| GO:0019898 | extrinsic component of membrane | Cellular Component | 41 | 0.360 | 1.337 | 0.0791 | 0.2230 | tags=37%, list=20%, signal=45% |
| GO:0042908 | xenobiotic transport | Biological Process | 71 | 0.313 | 1.335 | 0.0480 | 0.2248 | tags=27%, list=13%, signal=31% |
| GO:0051453 | regulation of intracellular pH | Biological Process | 16 | 0.459 | 1.334 | 0.1100 | 0.2263 | tags=63%, list=23%, signal=81% |
| GO:0034976 | response to endoplasmic reticulum stress | Biological Process | 51 | 0.339 | 1.333 | 0.0656 | 0.2257 | tags=29%, list=17%, signal=35% |
| GO:0006812 | monoatomic cation transport | Biological Process | 348 | 0.252 | 1.333 | 0 | 0.2257 | tags=31%, list=23%, signal=39% |
| GO:0120545 | nucleic acid conformation isomerase activity | Molecular Function | 150 | 0.277 | 1.332 | 0.0213 | 0.2263 | tags=35%, list=22%, signal=44% |
| GO:0009636 | response to toxic substance | Biological Process | 248 | 0.260 | 1.331 | 0.0295 | 0.2275 | tags=26%, list=17%, signal=31% |
| GO:0098800 | inner mitochondrial membrane protein complex | Cellular Component | 49 | 0.340 | 1.329 | 0.0590 | 0.2284 | tags=45%, list=28%, signal=62% |

| GO ID | GO name | Category | Size | ES | NES | Nominal P | FDR q | Leading edge |
|---|---|---|---|---|---|---|---|---|
| GO:0051119 | sugar transmembrane transporter activity | Molecular Function | 43 | 0.349 | 1.329 | 0.0687 | 0.2282 | tags=26%, list=15%, signal=30% |
| GO:0032984 | protein-containing complex disassembly | Biological Process | 76 | 0.308 | 1.328 | 0.0533 | 0.2288 | tags=32%, list=18%, signal=38% |
| GO:0016874 | ligase activity | Molecular Function | 306 | 0.255 | 1.328 | 0.0045 | 0.2284 | tags=39%, list=30%, signal=56% |
| GO:0036503 | ERAD pathway | Biological Process | 38 | 0.369 | 1.327 | 0.0979 | 0.2290 | tags=29%, list=17%, signal=35% |
| GO:0002097 | tRNA wobble base modification | Biological Process | 25 | 0.403 | 1.325 | 0.1067 | 0.2312 | tags=56%, list=35%, signal=86% |
| GO:0046173 | polyol biosynthetic process | Biological Process | 21 | 0.419 | 1.325 | 0.1261 | 0.2305 | tags=38%, list=12%, signal=43% |
| GO:0018193 | peptidyl-amino acid modification | Biological Process | 33 | 0.366 | 1.322 | 0.1027 | 0.2345 | tags=39%, list=27%, signal=54% |
| GO:0042910 | xenobiotic transmembrane transporter activity | Molecular Function | 71 | 0.313 | 1.321 | 0.0520 | 0.2342 | tags=27%, list=13%, signal=31% |
| GO:0031969 | chloroplast membrane | Cellular Component | 115 | 0.285 | 1.321 | 0.0340 | 0.2343 | tags=34%, list=24%, signal=44% |
| GO:0019751 | polyol metabolic process | Biological Process | 49 | 0.341 | 1.320 | 0.0819 | 0.2346 | tags=29%, list=13%, signal=33% |
| GO:0003729 | mRNA binding | Molecular Function | 448 | 0.240 | 1.320 | 0 | 0.2341 | tags=31%, list=29%, signal=43% |
| GO:0051539 | 4 iron, 4 sulfur cluster binding | Molecular Function | 43 | 0.341 | 1.318 | 0.1031 | 0.2362 | tags=35%, list=19%, signal=43% |
| GO:0046618 | xenobiotic export from cell | Biological Process | 49 | 0.332 | 1.316 | 0.0829 | 0.2378 | tags=31%, list=13%, signal=35% |
| GO:0009156 | ribonucleoside monophosphate biosynthetic process | Biological Process | 50 | 0.338 | 1.315 | 0.0912 | 0.2383 | tags=34%, list=20%, signal=42% |
| GO:0015297 | antiporter activity | Molecular Function | 127 | 0.280 | 1.313 | 0.0425 | 0.2416 | tags=35%, list=22%, signal=45% |
| GO:0008234 | cysteine-type peptidase activity | Molecular Function | 98 | 0.293 | 1.311 | 0.0426 | 0.2431 | tags=31%, list=24%, signal=40% |
| GO:0098754 | detoxification | Biological Process | 246 | 0.258 | 1.307 | 0.0180 | 0.2481 | tags=26%, list=17%, signal=31% |
| GO:1901679 | nucleotide transmembrane transport | Biological Process | 22 | 0.410 | 1.306 | 0.1165 | 0.2489 | tags=32%, list=15%, signal=37% |
| GO:0050832 | defense response to fungus | Biological Process | 37 | -0.382 | -1.315 | 0.1086 | 0.2457 | tags=49%, list=30%, signal=69% |

| GO ID | GO name | Category | Size | ES | NES | Nominal P | FDR q | Leading edge |
|---|---|---|---|---|---|---|---|---|
| GO:0006801 | superoxide metabolic process | Biological Process | 21 | -0.439 | -1.315 | 0.1353 | 0.2459 | tags=48%, list=24%, signal=63% |
| GO:0048868 | pollen tube development | Biological Process | 34 | -0.389 | -1.316 | 0.1105 | 0.2460 | tags=12%, list=5%, signal=12% |
| GO:0009890 | negative regulation of biosynthetic process | Biological Process | 231 | -0.285 | -1.317 | 0.0247 | 0.2454 | tags=32%, list=26%, signal=43% |
| GO:1903046 | meiotic cell cycle process | Biological Process | 56 | -0.351 | -1.317 | 0.0899 | 0.2461 | tags=32%, list=16%, signal=38% |
| GO:0080134 | regulation of response to stress | Biological Process | 95 | -0.322 | -1.317 | 0.0630 | 0.2462 | tags=43%, list=29%, signal=60% |
| GO:0044550 | secondary metabolite biosynthetic process | Biological Process | 83 | -0.330 | -1.319 | 0.0676 | 0.2437 | tags=37%, list=26%, signal=50% |
| GO:0042761 | very long-chain fatty acid biosynthetic process | Biological Process | 19 | -0.461 | -1.322 | 0.1123 | 0.2411 | tags=21%, list=6%, signal=22% |
| GO:0140013 | meiotic nuclear division | Biological Process | 48 | -0.365 | -1.322 | 0.1053 | 0.2415 | tags=31%, list=16%, signal=37% |
| GO:0019207 | kinase regulator activity | Molecular Function | 56 | -0.362 | -1.326 | 0.0894 | 0.2363 | tags=30%, list=15%, signal=35% |
| GO:0022414 | reproductive process | Biological Process | 339 | -0.278 | -1.326 | 0.0076 | 0.2363 | tags=26%, list=21%, signal=33% |
| GO:0016682 | oxidoreductase activity, acting on diphenols and related substances as donors, oxygen as acceptor | Molecular Function | 37 | -0.388 | -1.327 | 0.0759 | 0.2365 | tags=30%, list=14%, signal=34% |
| GO:0019748 | secondary metabolic process | Biological Process | 135 | -0.308 | -1.327 | 0.0295 | 0.2367 | tags=36%, list=25%, signal=47% |
| GO:0009892 | negative regulation of metabolic process | Biological Process | 249 | -0.287 | -1.331 | 0.0239 | 0.2329 | tags=33%, list=26%, signal=44% |
| GO:0008299 | isoprenoid biosynthetic process | Biological Process | 106 | -0.324 | -1.333 | 0.0602 | 0.2293 | tags=40%, list=24%, signal=52% |
| GO:0060918 | auxin transport | Biological Process | 34 | -0.408 | -1.335 | 0.1059 | 0.2285 | tags=35%, list=18%, signal=43% |
| GO:0009914 | hormone transport | Biological Process | 35 | -0.403 | -1.335 | 0.1081 | 0.2279 | tags=34%, list=18%, signal=42% |
| GO:0030135 | coated vesicle | Cellular Component | 106 | -0.322 | -1.340 | 0.0570 | 0.2221 | tags=43%, list=28%, signal=61% |
| GO:0016881 | acid-amino acid ligase activity | Molecular Function | 26 | -0.424 | -1.341 | 0.1142 | 0.2216 | tags=54%, list=26%, signal=73% |
| GO:0032940 | secretion by cell | Biological Process | 49 | -0.371 | -1.342 | 0.0671 | 0.2206 | tags=33%, list=26%, signal=44% |

| GO ID | GO name | Category | Size | ES | NES | Nominal P | FDR q | Leading edge |
|---|---|---|---|---|---|---|---|---|
| GO:0009812 | flavonoid metabolic process | Biological Process | 31 | -0.406 | -1.343 | 0.0967 | 0.2205 | tags=32%, list=17%, signal=39% |
| GO:0008353 | RNA polymerase II CTD heptapeptide repeat kinase activity | Molecular Function | 26 | -0.423 | -1.344 | 0.0983 | 0.2197 | tags=27%, list=11%, signal=30% |
| GO:0010329 | auxin efflux transmembrane transporter activity | Molecular Function | 19 | -0.473 | -1.353 | 0.1117 | 0.2085 | tags=26%, list=9%, signal=29% |
| GO:0030137 | COPI-coated vesicle | Cellular Component | 24 | -0.445 | -1.354 | 0.0957 | 0.2082 | tags=46%, list=24%, signal=60% |
| GO:0016049 | cell growth | Biological Process | 57 | -0.365 | -1.357 | 0.0692 | 0.2038 | tags=33%, list=27%, signal=45% |
| GO:0048519 | negative regulation of biological process | Biological Process | 363 | -0.286 | -1.360 | 0.0063 | 0.2010 | tags=35%, list=26%, signal=47% |
| GO:0009808 | lignin metabolic process | Biological Process | 35 | -0.399 | -1.360 | 0.0793 | 0.2017 | tags=31%, list=14%, signal=36% |
| GO:0030118 | clathrin coat | Cellular Component | 40 | -0.396 | -1.363 | 0.0695 | 0.1989 | tags=48%, list=26%, signal=64% |
| GO:0000159 | protein phosphatase type 2A complex | Cellular Component | 29 | -0.428 | -1.364 | 0.0929 | 0.1982 | tags=38%, list=16%, signal=45% |
| GO:0006259 | DNA metabolic process | Biological Process | 424 | -0.283 | -1.365 | 0.0025 | 0.1978 | tags=32%, list=25%, signal=42% |
| GO:0030663 | COPI-coated vesicle membrane | Cellular Component | 24 | -0.445 | -1.366 | 0.0846 | 0.1962 | tags=46%, list=24%, signal=60% |
| GO:1901659 | glycosyl compound biosynthetic process | Biological Process | 15 | -0.506 | -1.367 | 0.1091 | 0.1955 | tags=47%, list=12%, signal=53% |
| GO:0048523 | negative regulation of cellular process | Biological Process | 337 | -0.289 | -1.368 | 0.0110 | 0.1955 | tags=35%, list=26%, signal=47% |
| GO:0030234 | enzyme regulator activity | Molecular Function | 415 | -0.284 | -1.372 | 0.0047 | 0.1909 | tags=24%, list=17%, signal=29% |
| GO:0005811 | lipid droplet | Cellular Component | 18 | -0.477 | -1.373 | 0.1011 | 0.1903 | tags=61%, list=33%, signal=92% |
| GO:0080161 | auxin transmembrane transporter activity | Molecular Function | 19 | -0.473 | -1.374 | 0.1047 | 0.1907 | tags=26%, list=9%, signal=29% |
| GO:0030126 | COPI vesicle coat | Cellular Component | 23 | -0.449 | -1.375 | 0.0909 | 0.1897 | tags=48%, list=24%, signal=63% |
| GO:0030131 | clathrin adaptor complex | Cellular Component | 22 | -0.461 | -1.375 | 0.0839 | 0.1902 | tags=45%, list=26%, signal=61% |
| GO:0050660 | flavin adenine dinucleotide binding | Molecular Function | 120 | -0.331 | -1.379 | 0.0161 | 0.1862 | tags=22%, list=14%, signal=25% |

| GO ID | GO name | Category | Size | ES | NES | Nominal P | FDR q | Leading edge |
|---|---|---|---|---|---|---|---|---|
| GO:0010015 | root morphogenesis | Biological Process | 40 | -0.395 | -1.380 | 0.0618 | 0.1854 | tags=25%, list=14%, signal=29% |
| GO:0031507 | heterochromatin formation | Biological Process | 18 | -0.481 | -1.380 | 0.1147 | 0.1859 | tags=44%, list=24%, signal=58% |
| GO:0003688 | DNA replication origin binding | Molecular Function | 16 | -0.492 | -1.383 | 0.1005 | 0.1832 | tags=63%, list=31%, signal=91% |
| GO:0016052 | carbohydrate catabolic process | Biological Process | 196 | -0.310 | -1.386 | 0.0068 | 0.1805 | tags=28%, list=17%, signal=34% |
| GO:0010104 | regulation of ethylene-activated signaling pathway | Biological Process | 15 | -0.509 | -1.388 | 0.1168 | 0.1789 | tags=33%, list=6%, signal=36% |
| GO:0051304 | chromosome separation | Biological Process | 16 | -0.502 | -1.391 | 0.0886 | 0.1755 | tags=38%, list=14%, signal=44% |
| GO:0048468 | cell development | Biological Process | 47 | -0.390 | -1.395 | 0.0684 | 0.1721 | tags=30%, list=21%, signal=37% |
| GO:0009617 | response to bacterium | Biological Process | 91 | -0.346 | -1.397 | 0.0209 | 0.1703 | tags=32%, list=19%, signal=39% |
| GO:0005839 | proteasome core complex | Cellular Component | 40 | -0.402 | -1.400 | 0.0470 | 0.1671 | tags=68%, list=41%, signal=114% |
| GO:0016114 | terpenoid biosynthetic process | Biological Process | 69 | -0.363 | -1.401 | 0.0257 | 0.1665 | tags=42%, list=24%, signal=55% |
| GO:0005881 | cytoplasmic microtubule | Cellular Component | 22 | -0.469 | -1.403 | 0.0839 | 0.1647 | tags=27%, list=9%, signal=30% |
| GO:0045489 | pectin biosynthetic process | Biological Process | 16 | -0.509 | -1.406 | 0.0829 | 0.1629 | tags=56%, list=33%, signal=83% |
| GO:0010558 | negative regulation of macromolecule biosynthetic process | Biological Process | 225 | -0.308 | -1.406 | 0.0027 | 0.1635 | tags=33%, list=26%, signal=44% |
| GO:0009147 | pyrimidine nucleoside triphosphate metabolic process | Biological Process | 20 | -0.486 | -1.406 | 0.0683 | 0.1635 | tags=30%, list=8%, signal=33% |
| GO:0008081 | phosphoric diester hydrolase activity | Molecular Function | 38 | -0.412 | -1.411 | 0.0530 | 0.1593 | tags=50%, list=27%, signal=69% |
| GO:0070297 | regulation of phosphorelay signal transduction system | Biological Process | 15 | -0.509 | -1.411 | 0.0810 | 0.1593 | tags=33%, list=6%, signal=36% |
| GO:0000139 | Golgi membrane | Cellular Component | 302 | -0.301 | -1.412 | 0.0027 | 0.1592 | tags=32%, list=24%, signal=43% |
| GO:0051321 | meiotic cell cycle | Biological Process | 60 | -0.373 | -1.415 | 0.0441 | 0.1564 | tags=35%, list=17%, signal=42% |
| GO:0009630 | gravitropism | Biological Process | 16 | -0.519 | -1.416 | 0.0796 | 0.1558 | tags=38%, list=18%, signal=46% |

| GO ID | GO name | Category | Size | ES | NES | Nominal P | FDR q | Leading edge |
|---|---|---|---|---|---|---|---|---|
| GO:0033612 | receptor serine/threonine kinase binding | Molecular Function | 39 | -0.414 | -1.416 | 0.0348 | 0.1562 | tags=36%, list=19%, signal=44% |
| GO:0004784 | superoxide dismutase activity | Molecular Function | 18 | -0.496 | -1.417 | 0.0927 | 0.1566 | tags=56%, list=27%, signal=77% |
| GO:0030981 | cortical microtubule cytoskeleton | Cellular Component | 18 | -0.499 | -1.418 | 0.0752 | 0.1554 | tags=28%, list=9%, signal=31% |
| GO:0048827 | phyllome development | Biological Process | 71 | -0.365 | -1.418 | 0.0281 | 0.1558 | tags=28%, list=15%, signal=33% |
| GO:0005507 | copper ion binding | Molecular Function | 94 | -0.355 | -1.421 | 0.0136 | 0.1528 | tags=30%, list=14%, signal=34% |
| GO:0009694 | jasmonic acid metabolic process | Biological Process | 22 | -0.474 | -1.423 | 0.0684 | 0.1519 | tags=27%, list=12%, signal=31% |
| GO:0000160 | phosphorelay signal transduction system | Biological Process | 113 | -0.340 | -1.424 | 0.0186 | 0.1512 | tags=28%, list=15%, signal=33% |
| GO:0009813 | flavonoid biosynthetic process | Biological Process | 29 | -0.440 | -1.427 | 0.0617 | 0.1489 | tags=34%, list=17%, signal=42% |
| GO:0016721 | oxidoreductase activity, acting on superoxide radicals as acceptor | Molecular Function | 18 | -0.496 | -1.427 | 0.0646 | 0.1495 | tags=56%, list=27%, signal=77% |
| GO:1902531 | regulation of intracellular signal transduction | Biological Process | 33 | -0.428 | -1.431 | 0.0682 | 0.1464 | tags=36%, list=18%, signal=44% |
| GO:0060560 | developmental growth involved in morphogenesis | Biological Process | 49 | -0.391 | -1.433 | 0.0396 | 0.1444 | tags=29%, list=21%, signal=36% |
| GO:0051253 | negative regulation of RNA metabolic process | Biological Process | 75 | -0.369 | -1.436 | 0.0341 | 0.1422 | tags=36%, list=25%, signal=48% |
| GO:0006887 | exocytosis | Biological Process | 40 | -0.411 | -1.439 | 0.0273 | 0.1400 | tags=38%, list=26%, signal=51% |
| GO:0010605 | negative regulation of macromolecule metabolic process | Biological Process | 240 | -0.315 | -1.439 | 0.0026 | 0.1401 | tags=34%, list=26%, signal=45% |
| GO:0006631 | fatty acid metabolic process | Biological Process | 201 | -0.317 | -1.445 | 0.0079 | 0.1352 | tags=34%, list=27%, signal=47% |
| GO:0016645 | oxidoreductase activity, acting on the CH-NH group of donors | Molecular Function | 26 | -0.462 | -1.446 | 0.0417 | 0.1342 | tags=27%, list=7%, signal=29% |
| GO:0010315 | auxin export across the plasma membrane | Biological Process | 18 | -0.502 | -1.450 | 0.0524 | 0.1315 | tags=28%, list=9%, signal=30% |
| GO:0030863 | cortical cytoskeleton | Cellular Component | 21 | -0.479 | -1.452 | 0.0714 | 0.1297 | tags=33%, list=15%, signal=39% |
| GO:0140994 | RNA polymerase II CTD heptapeptide repeat modifying activity | Molecular Function | 34 | -0.436 | -1.453 | 0.0271 | 0.1293 | tags=26%, list=11%, signal=30% |

| GO ID | GO name | Category | Size | ES | NES | Nominal P | FDR q | Leading edge |
|---|---|---|---|---|---|---|---|---|
| GO:0006022 | aminoglycan metabolic process | Biological Process | 23 | -0.474 | -1.454 | 0.0634 | 0.1291 | tags=26%, list=8%, signal=28% |
| GO:0019914 | cyclin-dependent protein kinase regulator activity | Molecular Function | 26 | -0.466 | -1.455 | 0.0509 | 0.1290 | tags=31%, list=7%, signal=33% |
| GO:0055028 | cortical microtubule | Cellular Component | 18 | -0.499 | -1.457 | 0.0511 | 0.1279 | tags=28%, list=9%, signal=31% |
| GO:0007127 | meiosis I | Biological Process | 34 | -0.435 | -1.457 | 0.0556 | 0.1282 | tags=32%, list=16%, signal=38% |
| GO:0016538 | cyclin-dependent protein serine/threonine kinase regulator activity | Molecular Function | 26 | -0.466 | -1.458 | 0.0444 | 0.1282 | tags=31%, list=7%, signal=33% |
| GO:0031349 | positive regulation of defense response | Biological Process | 27 | -0.458 | -1.458 | 0.0542 | 0.1286 | tags=52%, list=28%, signal=72% |
| GO:0000776 | kinetochore | Cellular Component | 16 | -0.534 | -1.460 | 0.0614 | 0.1272 | tags=56%, list=25%, signal=75% |
| GO:0016799 | hydrolase activity, hydrolyzing N-glycosyl compounds | Molecular Function | 96 | -0.360 | -1.462 | 0.0106 | 0.1262 | tags=57%, list=36%, signal=90% |
| GO:0010928 | regulation of auxin mediated signaling pathway | Biological Process | 26 | -0.470 | -1.463 | 0.0639 | 0.1254 | tags=27%, list=9%, signal=29% |
| GO:0030662 | coated vesicle membrane | Cellular Component | 76 | -0.376 | -1.467 | 0.0203 | 0.1231 | tags=54%, list=32%, signal=79% |
| GO:0030117 | membrane coat | Cellular Component | 85 | -0.366 | -1.470 | 0.0123 | 0.1209 | tags=44%, list=26%, signal=59% |
| GO:0048475 | coated membrane | Cellular Component | 85 | -0.366 | -1.471 | 0.0162 | 0.1204 | tags=44%, list=26%, signal=59% |
| GO:0000724 | double-strand break repair via homologous recombination | Biological Process | 74 | -0.379 | -1.474 | 0.0301 | 0.1185 | tags=30%, list=16%, signal=35% |
| GO:0000155 | phosphorelay sensor kinase activity | Molecular Function | 23 | -0.495 | -1.480 | 0.0467 | 0.1142 | tags=26%, list=6%, signal=28% |
| GO:0000779 | condensed chromosome, centromeric region | Cellular Component | 16 | -0.534 | -1.484 | 0.0587 | 0.1106 | tags=56%, list=25%, signal=75% |
| GO:0003779 | actin binding | Molecular Function | 85 | -0.374 | -1.484 | 0.0134 | 0.1111 | tags=38%, list=21%, signal=48% |
| GO:0009629 | response to gravity | Biological Process | 18 | -0.520 | -1.485 | 0.0762 | 0.1109 | tags=39%, list=18%, signal=48% |
| GO:0030120 | vesicle coat | Cellular Component | 62 | -0.397 | -1.487 | 0.0188 | 0.1103 | tags=58%, list=32%, signal=85% |
| GO:0030599 | pectinesterase activity | Molecular Function | 29 | -0.456 | -1.487 | 0.0383 | 0.1108 | tags=38%, list=19%, signal=47% |

| GO ID | GO name | Category | Size | ES | NES | Nominal P | FDR q | Leading edge |
|---|---|---|---|---|---|---|---|---|
| GO:1903293 | phosphatase complex | Cellular Component | 34 | -0.442 | -1.489 | 0.0339 | 0.1096 | tags=38%, list=16%, signal=45% |
| GO:0048438 | floral whorl development | Biological Process | 15 | -0.540 | -1.489 | 0.0661 | 0.1101 | tags=27%, list=8%, signal=29% |
| GO:0006302 | double-strand break repair | Biological Process | 98 | -0.363 | -1.489 | 0.0177 | 0.1101 | tags=30%, list=16%, signal=35% |
| GO:0051983 | regulation of chromosome segregation | Biological Process | 17 | -0.521 | -1.497 | 0.0407 | 0.1046 | tags=35%, list=15%, signal=41% |
| GO:0070001 | aspartic-type peptidase activity | Molecular Function | 60 | -0.400 | -1.497 | 0.0163 | 0.1048 | tags=18%, list=11%, signal=21% |
| GO:0010014 | meristem initiation | Biological Process | 16 | -0.549 | -1.500 | 0.0502 | 0.1031 | tags=31%, list=10%, signal=35% |
| GO:0008287 | protein serine/threonine phosphatase complex | Cellular Component | 34 | -0.442 | -1.504 | 0.0219 | 0.1008 | tags=38%, list=16%, signal=45% |
| GO:0003002 | regionalization | Biological Process | 24 | -0.492 | -1.505 | 0.0277 | 0.1002 | tags=25%, list=13%, signal=29% |
| GO:0071949 | FAD binding | Molecular Function | 56 | -0.409 | -1.505 | 0.0147 | 0.1005 | tags=23%, list=9%, signal=25% |
| GO:0004190 | aspartic-type endopeptidase activity | Molecular Function | 60 | -0.400 | -1.510 | 0.0138 | 0.0979 | tags=18%, list=11%, signal=21% |
| GO:0033045 | regulation of sister chromatid segregation | Biological Process | 15 | -0.554 | -1.511 | 0.0363 | 0.0972 | tags=40%, list=15%, signal=47% |
| GO:0016775 | phosphotransferase activity, nitrogenous group as acceptor | Molecular Function | 23 | -0.495 | -1.511 | 0.0258 | 0.0976 | tags=26%, list=6%, signal=28% |
| GO:0071669 | plant-type cell wall organization or biogenesis | Biological Process | 70 | -0.391 | -1.521 | 0.0061 | 0.0906 | tags=43%, list=24%, signal=56% |
| GO:0000217 | DNA secondary structure binding | Molecular Function | 34 | -0.449 | -1.522 | 0.0299 | 0.0902 | tags=59%, list=29%, signal=82% |
| GO:0000725 | recombinational repair | Biological Process | 79 | -0.389 | -1.525 | 0.0139 | 0.0886 | tags=29%, list=16%, signal=35% |
| GO:0009908 | flower development | Biological Process | 61 | -0.409 | -1.528 | 0.0079 | 0.0869 | tags=25%, list=11%, signal=27% |
| GO:0010053 | root epidermal cell differentiation | Biological Process | 25 | -0.489 | -1.529 | 0.0273 | 0.0872 | tags=32%, list=14%, signal=37% |
| GO:0120543 | macromolecular conformation isomerase activity | Molecular Function | 216 | -0.336 | -1.531 | 0 | 0.0862 | tags=32%, list=18%, signal=39% |
| GO:0004673 | protein histidine kinase activity | Molecular Function | 23 | -0.495 | -1.533 | 0.0299 | 0.0854 | tags=26%, list=6%, signal=28% |

| GO ID | GO name | Category | Size | ES | NES | Nominal P | FDR q | Leading edge |
|---|---|---|---|---|---|---|---|---|
| GO:0071369 | cellular response to ethylene stimulus | Biological Process | 63 | -0.406 | -1.534 | 0.0155 | 0.0847 | tags=40%, list=19%, signal=49% |
| GO:0061809 | NAD+ nucleosidase activity, cyclic ADP-ribose generating | Molecular Function | 64 | -0.406 | -1.539 | 0.0132 | 0.0820 | tags=56%, list=36%, signal=88% |
| GO:0009873 | ethylene-activated signaling pathway | Biological Process | 63 | -0.406 | -1.542 | 0.0047 | 0.0806 | tags=40%, list=19%, signal=49% |
| GO:0051128 | regulation of cellular component organization | Biological Process | 81 | -0.388 | -1.545 | 0.0090 | 0.0792 | tags=37%, list=24%, signal=48% |
| GO:0140414 | phosphopantetheine-dependent carrier activity | Molecular Function | 16 | -0.567 | -1.547 | 0.0361 | 0.0782 | tags=38%, list=23%, signal=48% |
| GO:0009524 | phragmoplast | Cellular Component | 21 | -0.522 | -1.550 | 0.0330 | 0.0765 | tags=48%, list=16%, signal=57% |
| GO:0005085 | guanyl-nucleotide exchange factor activity | Molecular Function | 34 | -0.463 | -1.553 | 0.0205 | 0.0749 | tags=32%, list=16%, signal=39% |
| GO:0004650 | polygalacturonase activity | Molecular Function | 30 | -0.472 | -1.555 | 0.0239 | 0.0742 | tags=23%, list=5%, signal=25% |
| GO:0000347 | THO complex | Cellular Component | 20 | -0.525 | -1.558 | 0.0358 | 0.0726 | tags=45%, list=19%, signal=56% |
| GO:0002229 | defense response to oomycetes | Biological Process | 26 | -0.497 | -1.561 | 0.0239 | 0.0713 | tags=42%, list=19%, signal=52% |
| GO:0000036 | acyl carrier activity | Molecular Function | 16 | -0.567 | -1.567 | 0.0244 | 0.0687 | tags=38%, list=23%, signal=48% |
| GO:0009755 | hormone-mediated signaling pathway | Biological Process | 339 | -0.330 | -1.568 | 0 | 0.0681 | tags=34%, list=22%, signal=43% |
| GO:0003682 | chromatin binding | Molecular Function | 127 | -0.366 | -1.569 | 0.0043 | 0.0681 | tags=38%, list=22%, signal=49% |
| GO:0002239 | response to oomycetes | Biological Process | 26 | -0.497 | -1.569 | 0.0153 | 0.0683 | tags=42%, list=19%, signal=52% |
| GO:0071554 | cell wall organization or biogenesis | Biological Process | 177 | -0.353 | -1.573 | 0.0014 | 0.0665 | tags=30%, list=17%, signal=36% |
| GO:0044620 | ACP phosphopantetheine attachment site binding | Molecular Function | 16 | -0.567 | -1.574 | 0.0324 | 0.0666 | tags=38%, list=23%, signal=48% |
| GO:0006325 | chromatin organization | Biological Process | 195 | -0.350 | -1.576 | 0.0014 | 0.0659 | tags=35%, list=24%, signal=46% |
| GO:0051192 | prosthetic group binding | Molecular Function | 16 | -0.567 | -1.576 | 0.0372 | 0.0662 | tags=38%, list=23%, signal=48% |
| GO:0042545 | cell wall modification | Biological Process | 27 | -0.497 | -1.580 | 0.0162 | 0.0646 | tags=44%, list=19%, signal=55% |

| GO ID | GO name | Category | Size | ES | NES | Nominal P | FDR q | Leading edge |
|---|---|---|---|---|---|---|---|---|
| GO:0040008 | regulation of growth | Biological Process | 51 | -0.434 | -1.584 | 0.0155 | 0.0627 | tags=27%, list=14%, signal=32% |
| GO:0040007 | growth | Biological Process | 100 | -0.388 | -1.587 | 0.0014 | 0.0613 | tags=34%, list=24%, signal=44% |
| GO:0003678 | DNA helicase activity | Molecular Function | 35 | -0.478 | -1.587 | 0.0188 | 0.0616 | tags=29%, list=11%, signal=32% |
| GO:0006338 | chromatin remodeling | Biological Process | 160 | -0.358 | -1.591 | 0.0015 | 0.0602 | tags=36%, list=24%, signal=47% |
| GO:0009610 | response to symbiotic fungus | Biological Process | 24 | -0.510 | -1.591 | 0.0308 | 0.0605 | tags=25%, list=13%, signal=29% |
| GO:0070828 | heterochromatin organization | Biological Process | 21 | -0.533 | -1.592 | 0.0141 | 0.0605 | tags=52%, list=24%, signal=69% |
| GO:0032870 | cellular response to hormone stimulus | Biological Process | 343 | -0.334 | -1.593 | 0 | 0.0602 | tags=34%, list=22%, signal=43% |
| GO:0071495 | cellular response to endogenous stimulus | Biological Process | 344 | -0.334 | -1.593 | 0 | 0.0605 | tags=34%, list=22%, signal=43% |
| GO:0045892 | negative regulation of DNA-templated transcription | Biological Process | 65 | -0.421 | -1.594 | 0.0048 | 0.0607 | tags=40%, list=25%, signal=53% |
| GO:0032451 | demethylase activity | Molecular Function | 25 | -0.515 | -1.595 | 0.0171 | 0.0604 | tags=52%, list=28%, signal=73% |
| GO:0004620 | glycerophospholipase activity | Molecular Function | 45 | -0.458 | -1.607 | 0.0097 | 0.0546 | tags=51%, list=27%, signal=70% |
| GO:0048285 | organelle fission | Biological Process | 121 | -0.377 | -1.608 | 0 | 0.0544 | tags=32%, list=14%, signal=37% |
| GO:0000793 | condensed chromosome | Cellular Component | 37 | -0.477 | -1.611 | 0.0066 | 0.0531 | tags=38%, list=17%, signal=45% |
| GO:0045934 | negative regulation of nucleobase-containing compound metabolic process | Biological Process | 90 | -0.396 | -1.611 | 0.0015 | 0.0533 | tags=40%, list=25%, signal=54% |
| GO:0003680 | minor groove of adenine-thymine-rich DNA binding | Molecular Function | 30 | -0.492 | -1.618 | 0.0116 | 0.0506 | tags=63%, list=29%, signal=89% |
