## Supplementary tables and figures for "Transcriptome of apple cv. Ametyst in *Rvi6*-effective and *Rvi6*-breaking *Venturia inaequalis* interactions identifies defense candidates": 12_SupplementaryTable_S5_Trinity_vs_GDDH13.pdf

**Supplementary Table S5. Concordance and classification of *de novo* Trinity candidates against the GDDH13 reference.**

All 1,817 Trinity Rin-vs-Rola candidate genes (thresholded Wald test, FDR < 0.01, |log2FC| > 1) were aligned to the GDDH13 proteome by BLASTX and classified according to their relationship with the reference-based analysis; summary counts for both quantification strategies are given in Table 1. A confident GDDH13 protein match required ≥ 40% amino-acid identity across ≥ 50 aligned residues, summed across high-scoring segment pairs to the best-scoring GDDH13 subject; query coverage was deliberately not used as a criterion, because for BLASTX the coverage of a transcript is deflated by its untranslated regions. A candidate was classified as having no GDDH13 protein hit only when no BLASTX alignment was detected. Of the 1,817 candidates, 1,154 (64%) had a confident GDDH13 protein match and 815 were concordant candidates in the same direction, while 274 had a partial hit below the match-confidence threshold — short or divergent alignments typical of fragmentary *de novo* assemblies — and 389 had no hit, dominated by unannotated transcripts and low-complexity glycine-rich and cell-wall proteins. None of the four defense-annotated candidates without a GDDH13 protein hit showed meaningful similarity to *HcrVf1–HcrVf3* (BLASTX; GenBank AJ297739–AJ297741). Trinity log2FC and padj are from the Trinity Rin vs Rola contrast and GDDH13 log2FC and padj from the corresponding reference-based contrast; positive log2FC denotes higher expression in Rin. Def., defense-related annotation, marked 'yes' where present and left blank otherwise; NA in the GDDH13 columns, no confident GDDH13 protein match — either no BLASTX alignment at all (389 candidates) or a hit below the match-confidence threshold (274); NA in the GDDH13 log2FC and padj columns alone, a confident match whose gene is not among the 38,467 genes tested in the reference-based analysis (13 candidates). Within each class, genes are ranked by absolute Trinity log2 fold change.

The final column gives the result of contamination screening of the gene-representative set with FCS-GX (Materials and methods), performed after the differential-expression analysis: the taxon assigned to sequences flagged as non-plant, and an em dash for sequences not flagged. 612 of the 1,817 candidates (33.7 %) were flagged, 594 of them as *Venturia inaequalis*. The flag is concentrated in the Trinity-only classes: 2 of 815 concordant candidates (0.2 %) against 44–66 % of every class without a matching GDDH13 candidate.

| Trinity gene ID | Direction | Trinity | | GDDH13 protein match | | | Def. | Description | FCS-GX |
|---|---|---|---|---|---|---|---|---|---|
| | | log2FC | padj | gene ID | log2FC | padj | | | call |
| **Concordant — confident GDDH13 protein match, candidate in both analyses, same direction (n = 815)** | | | | | | | | | |
| TRINITY_DN180346_c0_g1 | Rola-high | -12.66 | 2.29e-09 | MD05G1159700 | -12.56 | 6.67e-12 | yes | putative lipid-transfer protein DIR1 | — |
| TRINITY_DN13739_c0_g1 | Rola-high | -11.49 | 1.92e-07 | MD04G1234300 | -11.55 | 8.60e-16 | yes | GDSL esterase/lipase At5g33370-like | — |
| TRINITY_DN7844_c0_g2 | Rola-high | -11.14 | 8.46e-09 | MD00G1174400 | -11.55 | 1.01e-11 | | palmitoyl-monogalactosyldiacylglycerol delta-7 desaturase, chloroplastic-like | — |
| TRINITY_DN250_c0_g1 | Rola-high | -10.64 | 2.92e-06 | MD09G1154000 | -9.98 | 2.78e-07 | | protodermal factor 1-like | — |
| TRINITY_DN1804_c0_g2 | Rola-high | -10.50 | 5.02e-12 | MD04G1234300 | -11.55 | 8.60e-16 | yes | GDSL esterase/lipase At5g33370-like | — |
| TRINITY_DN13503_c0_g1 | Rola-high | -9.78 | 1.76e-05 | MD14G1081900 | -11.07 | 8.32e-09 | | protein GAST1 isoform X2 | — |
| TRINITY_DN80465_c0_g1 | Rola-high | -9.50 | 3.09e-11 | MD17G1079900 | -9.18 | 4.88e-16 | | putative cell wall protein | — |
| TRINITY_DN12867_c0_g1 | Rola-high | -9.42 | 1.40e-07 | MD12G1027900 | -9.64 | 1.02e-10 | | uncharacterized protein | — |
| TRINITY_DN9958_c0_g1 | Rin-high | 9.24 | 2.05e-07 | MD07G1266700 | 9.21 | 4.05e-10 | | uncharacterized protein | — |
| TRINITY_DN8846_c0_g2 | Rola-high | -9.23 | 2.95e-03 | MD15G1400200 | -24.53 | 3.29e-23 | yes | GDSL esterase/lipase At1g71691 | — |
| TRINITY_DN7610_c1_g2 | Rola-high | -9.09 | 2.22e-05 | MD17G1044600 | -9.02 | 8.76e-09 | | glycerol-3-phosphate acyltransferase RAM2-like | — |
| TRINITY_DN12454_c0_g1 | Rola-high | -9.09 | 1.01e-11 | MD16G1088600 | -9.36 | 7.91e-16 | yes | MLP-like protein 31 | — |

| Trinity gene ID | Direction | Trinity | | GDDH13 protein match | | | Def. | Description | FCS-GX |
|---|---|---|---|---|---|---|---|---|---|
| | | log2FC | padj | gene ID | log2FC | padj | | | call |
| TRINITY_DN14928_c1_g1 | Rola-high | -8.60 | 6.18e-09 | MD09G1193200 | -8.51 | 8.56e-12 | | protein HOTHEAD | — |
| TRINITY_DN11464_c0_g1 | Rola-high | -8.58 | 1.30e-05 | MD08G1221300 | -7.29 | 9.79e-08 | | protein SODIUM POTASSIUM ROOT DEFECTIVE 2-like isoform X2 | — |
| TRINITY_DN13792_c2_g1 | Rola-high | -8.51 | 4.84e-03 | MD01G1210800 | -24.13 | 2.14e-21 | | ABC transporter G family member 4 | — |
| TRINITY_DN18924_c1_g1 | Rola-high | -8.49 | 8.58e-12 | MD04G1030500 | -8.38 | 2.95e-18 | yes | putative lipid-transfer protein DIR1 | — |
| TRINITY_DN19474_c0_g1 | Rola-high | -8.17 | 5.77e-07 | MD11G1207300 | -3.26 | 6.18e-05 | yes | putative lipid-transfer protein DIR1 | — |
| TRINITY_DN9096_c0_g2 | Rola-high | -8.14 | 3.06e-04 | MD08G1215700 | -8.21 | 5.50e-07 | yes | GDSL esterase/lipase At1g33811 | — |
| TRINITY_DN8522_c0_g1 | Rin-high | 8.11 | 4.79e-07 | MD06G1089700 | 7.56 | 8.50e-08 | yes | myb-related protein 308-like | — |
| TRINITY_DN1692_c0_g1 | Rola-high | -8.07 | 5.51e-05 | MD08G1227900 | -7.96 | 4.97e-05 | | fatty acid elongase 3-like | — |
| TRINITY_DN13485_c0_g1 | Rola-high | -7.97 | 8.25e-04 | MD04G1028300 | -8.04 | 1.10e-05 | | TPD1 protein homolog 1-like | — |
| TRINITY_DN8592_c0_g1 | Rin-high | 7.73 | 8.26e-06 | MD13G1003800 | 7.84 | 2.46e-08 | | chaperone protein dnaJ C76, chloroplastic | — |
| TRINITY_DN18127_c0_g2 | Rola-high | -7.67 | 5.16e-03 | MD05G1185200 | -7.56 | 2.58e-04 | | FT-interacting protein 3 | — |
| TRINITY_DN8201_c1_g1 | Rola-high | -7.60 | 6.41e-12 | MD10G1171100 | -7.64 | 1.48e-14 | yes | GDSL esterase/lipase At5g33370-like | — |
| TRINITY_DN57958_c1_g1 | Rin-high | 7.57 | 4.76e-07 | MD16G1032300 | 7.02 | 3.81e-08 | | senescence-specific cysteine protease SAG12-like | — |
| TRINITY_DN318_c0_g1 | Rin-high | 7.50 | 5.65e-04 | MD05G1081700 | 7.07 | 1.03e-06 | | cold-regulated protein 27 | — |
| TRINITY_DN13056_c0_g1 | Rola-high | -7.49 | 7.16e-03 | MD15G1161300 | -7.22 | 1.08e-05 | | uncharacterized methyltransferase At2g41040, chloroplastic-like isoform X1 | — |
| TRINITY_DN19276_c0_g1 | Rola-high | -7.34 | 4.50e-04 | MD06G1058600 | -7.16 | 1.32e-06 | yes | probable pectinesterase 53 | — |
| TRINITY_DN434_c0_g1 | Rin-high | 7.29 | 4.11e-07 | MD07G1299100 | 7.78 | 7.19e-11 | | two-component response regulator-like APRR5 | — |
| TRINITY_DN91469_c0_g1 | Rola-high | -7.22 | 1.66e-04 | MD04G1031400 | -6.25 | 4.62e-08 | | hypothetical protein OIU76_008293 | — |
| TRINITY_DN10557_c0_g1 | Rin-high | 7.16 | 6.25e-07 | MD04G1194600 | 6.73 | 4.63e-13 | | universal stress protein A-like protein isoform X1 | — |
| TRINITY_DN3477_c0_g1 | Rin-high | 7.14 | 2.45e-05 | MD16G1059700 | 6.35 | 2.25e-05 | yes | cytochrome P450 78A5-like | — |
| TRINITY_DN3180_c0_g2 | Rola-high | -7.13 | 2.16e-04 | MD03G1141400 | -7.15 | 3.73e-07 | | alcohol-forming fatty acyl-CoA reductase-like | — |
| TRINITY_DN6115_c0_g1 | Rin-high | 7.13 | 6.54e-06 | MD15G1082300 | 7.38 | 2.40e-08 | | cold-regulated protein 27 | — |
| TRINITY_DN50177_c0_g1 | Rola-high | -7.11 | 6.34e-06 | MD09G1144100 | -7.04 | 3.70e-09 | | DNA damage-repair/toleration protein DRT100-like | — |

| Trinity gene ID | Direction | Trinity | | GDDH13 protein match | | | Def. | Description | FCS-GX |
|---|---|---|---|---|---|---|---|---|---|
| | | log2FC | padj | gene ID | log2FC | padj | | | call |
| TRINITY_DN956_c1_g2 | Rin-high | 7.11 | 1.26e-04 | MD15G1082300 | 7.38 | 2.40e-08 | | cold-regulated protein 27-like isoform X1 | — |
| TRINITY_DN32998_c1_g1 | Rola-high | -7.05 | 1.64e-05 | MD06G1234500 | -6.58 | 2.50e-08 | | zinc-finger homeodomain protein 11-like | — |
| TRINITY_DN31077_c0_g1 | Rola-high | -6.96 | 2.95e-04 | MD03G1291200 | -7.12 | 1.21e-10 | yes | endoglucanase 17-like | — |
| TRINITY_DN35795_c0_g1 | Rin-high | 6.88 | 7.35e-05 | MD01G1228200 | 6.65 | 3.89e-09 | | two-component response regulator-like APRR5 | — |
| TRINITY_DN15390_c0_g1 | Rola-high | -6.81 | 1.73e-05 | MD17G1079900 | -9.18 | 4.88e-16 | | putative cell wall protein | — |
| TRINITY_DN182589_c0_g1 | Rin-high | 6.77 | 3.50e-05 | MD04G1140600 | 6.71 | 1.42e-07 | | 18.1 kDa class I heat shock protein-like | — |
| TRINITY_DN36141_c0_g1 | Rin-high | 6.73 | 1.68e-04 | MD05G1329700 | 5.90 | 8.28e-06 | yes | G-type lectin S-receptor-like serine/threonine-protein kinase SD1-29 | — |
| TRINITY_DN34535_c0_g3 | Rola-high | -6.72 | 5.02e-12 | MD09G1075500 | -5.69 | 1.68e-12 | yes | leucine-rich repeat extensin-like protein 5 isoform X2 | — |
| TRINITY_DN27098_c0_g1 | Rin-high | 6.72 | 9.00e-06 | MD01G1228200 | 6.65 | 3.89e-09 | | two-component response regulator-like APRR5 | — |
| TRINITY_DN28602_c0_g1 | Rola-high | -6.69 | 1.43e-03 | MD04G1031400 | -6.25 | 4.62e-08 | | hypothetical protein SUGI_0424310 | — |
| TRINITY_DN12121_c0_g1 | Rola-high | -6.67 | 6.25e-05 | MD04G1031500 | -6.44 | 1.21e-06 | | extensin-2-like isoform X1 | — |
| TRINITY_DN64355_c0_g1 | Rin-high | 6.64 | 1.08e-06 | MD05G1028500 | 6.58 | 7.19e-11 | | protein argonaute 18-like | — |
| TRINITY_DN14202_c0_g1 | Rola-high | -6.57 | 2.71e-05 | MD04G1131000 | -6.52 | 2.97e-06 | yes | group IIc WRKY transcription factor | — |
| TRINITY_DN3264_c0_g1 | Rin-high | 6.56 | 3.70e-04 | MD09G1276000 | 7.15 | 4.76e-08 | | microtubule-destabilizing protein 60 | — |
| TRINITY_DN8324_c0_g3 | Rola-high | -6.53 | 2.23e-05 | MD15G1095700 | -6.38 | 2.01e-08 | | U-box domain-containing protein 26 | — |
| TRINITY_DN23518_c1_g1 | Rola-high | -6.53 | 6.43e-05 | MD00G1115000 | -6.01 | 8.42e-08 | | uncharacterized protein | — |
| TRINITY_DN4008_c0_g2 | Rin-high | 6.49 | 3.33e-04 | MD13G1118500 | 6.51 | 2.47e-06 | yes | cytochrome P450 78A5-like | — |
| TRINITY_DN6594_c1_g1 | Rola-high | -6.48 | 1.60e-19 | MD05G1179500 | -6.95 | 2.67e-07 | | probable pectate lyase 8 | — |
| TRINITY_DN7443_c0_g1 | Rola-high | -6.46 | 3.27e-07 | MD02G1194700 | -6.41 | 1.13e-10 | yes | probable hexokinase-like 2 protein | — |
| TRINITY_DN50217_c0_g1 | Rola-high | -6.44 | 3.34e-09 | MD07G1135500 | -5.94 | 2.20e-15 | yes | cytochrome P450 86A8 | — |
| TRINITY_DN26566_c0_g1 | Rola-high | -6.43 | 5.94e-03 | MD10G1298300 | -6.58 | 6.46e-05 | yes | polyphenol oxidase latent form, chloroplastic-like | — |
| TRINITY_DN27579_c0_g2 | Rin-high | 6.41 | 2.19e-03 | MD06G1089700 | 7.56 | 8.50e-08 | yes | myb-related protein 308-like | — |
| TRINITY_DN2744_c0_g1 | Rin-high | 6.39 | 2.81e-09 | MD13G1025600 | 6.24 | 1.25e-13 | | uncharacterized protein | — |

| Trinity gene ID | Direction | Trinity | | GDDH13 protein match | | | Def. | Description | FCS-GX |
| --- | --- | --- | --- | --- | --- | --- | --- | --- | --- |
| | | log2FC | padj | gene ID | log2FC | padj | | | call |
| TRINITY_DN24046_c0_g1 | Rin-high | 6.37 | 1.86e-03 | MD01G1228200 | 6.65 | 3.89e-09 | | two-component response regulator-like APRR5 | — |
| TRINITY_DN48524_c0_g1 | Rin-high | 6.35 | 3.55e-04 | MD16G1116100 | 5.87 | 3.67e-08 | | geraniol 8-hydroxylase-like | — |
| TRINITY_DN2583_c0_g1 | Rola-high | -6.35 | 9.81e-07 | MD04G1031400 | -6.25 | 4.62e-08 | | extensin-2-like isoform X1 | — |
| TRINITY_DN24806_c0_g1 | Rin-high | 6.33 | 8.57e-11 | MD00G1192900 | 4.06 | 1.40e-12 | yes | probable LRR receptor-like serine/threonine-protein kinase At3g47570 isoform X2 | — |
| TRINITY_DN30682_c0_g2 | Rin-high | 6.27 | 4.64e-07 | MD07G1236800 | 3.98 | 3.09e-20 | yes | probable LRR receptor-like serine/threonine-protein kinase At3g47570 isoform X1 | — |
| TRINITY_DN1897_c0_g1 | Rin-high | 6.27 | 6.25e-10 | MD16G1091200 | 6.48 | 1.29e-16 | yes | probable xyloglucan endotransglucosylase/hydrolase protein 33 | — |
| TRINITY_DN30910_c0_g1 | Rola-high | -6.26 | 4.77e-03 | MD17G1078700 | -6.17 | 1.77e-04 | | hypothetical protein DVH24_028323 | — |
| TRINITY_DN3734_c0_g1 | Rin-high | 6.22 | 2.20e-04 | MD13G1172200 | 5.98 | 5.66e-07 | yes | monothiol glutaredoxin-S10 | — |
| TRINITY_DN92642_c0_g1 | Rola-high | -6.21 | 5.30e-05 | MD06G1066600 | -6.74 | 9.96e-07 | | acid beta-fructofuranosidase 2, vacuolar | — |
| TRINITY_DN13025_c0_g1 | Rola-high | -6.21 | 1.19e-08 | MD12G1259000 | -6.30 | 8.27e-13 | | hypothetical protein DVH24_023738 | — |
| TRINITY_DN33549_c0_g1 | Rola-high | -6.20 | 1.15e-06 | MD12G1200200 | -5.42 | 9.51e-10 | | probable L-cysteine desulfhydrase, chloroplastic | — |
| TRINITY_DN8543_c0_g2 | Rola-high | -6.20 | 2.02e-04 | MD03G1291200 | -7.12 | 1.21e-10 | yes | endoglucanase 17-like | — |
| TRINITY_DN8144_c1_g1 | Rola-high | -6.18 | 1.19e-03 | MD02G1190900 | -5.77 | 1.17e-05 | | 1-aminocyclopropane-1-carboxylate synthase | — |
| TRINITY_DN890_c0_g2 | Rin-high | 6.16 | 1.78e-03 | MD16G1191800 | 1.05 | 8.22e-03 | | hypothetical protein DVH24_022554 | — |
| TRINITY_DN35489_c0_g2 | Rola-high | -6.14 | 2.15e-04 | MD05G1269800 | -6.11 | 1.48e-06 | | putative germin-like protein 2-1 | — |
| TRINITY_DN36803_c0_g1 | Rola-high | -6.14 | 1.46e-05 | MD10G1288000 | -6.28 | 9.48e-09 | | MLO-like protein 6 | — |
| TRINITY_DN21546_c0_g1 | Rin-high | 6.13 | 1.01e-06 | MD06G1210200 | 5.86 | 9.28e-09 | | protein CHUP1, chloroplastic-like | — |
| TRINITY_DN42745_c0_g1 | Rola-high | -6.10 | 2.65e-03 | MD16G1188800 | -5.88 | 3.36e-04 | | uncharacterized protein | — |
| TRINITY_DN8093_c1_g3 | Rola-high | -6.07 | 1.42e-03 | MD14G1113300 | -6.08 | 1.41e-05 | | protein LURP-one-related 17-like | — |
| TRINITY_DN10616_c0_g2 | Rola-high | -6.06 | 1.64e-07 | MD03G1058700 | -6.93 | 4.29e-15 | | transcription factor SCREAM2-like isoform X1 | — |
| TRINITY_DN477_c8_g3 | Rin-high | 6.03 | 7.99e-10 | MD07G1236200 | 1.98 | 3.88e-08 | yes | probable LRR receptor-like serine/threonine-protein kinase At3g47570 isoform X1 | — |

| Trinity gene ID | Direction | Trinity | | GDDH13 protein match | | | Def. | Description | FCS-GX call |
|---|---|---|---|---|---|---|---|---|---|
| | | log2FC | padj | gene ID | log2FC | padj | | | |
| TRINITY_DN30762_c3_g1 | Rin-high | 5.98 | 5.23e-03 | MD08G1005500 | 4.69 | 2.74e-06 | | uncharacterized protein | — |
| TRINITY_DN228717_c0_g1 | Rola-high | -5.97 | 2.48e-05 | MD10G1036400 | -5.89 | 6.43e-08 | | 3-ketoacyl-CoA synthase 13-like | — |
| TRINITY_DN31390_c0_g1 | Rola-high | -5.97 | 1.91e-03 | MD13G1036800 | -2.83 | 2.53e-07 | yes | probable leucine-rich repeat receptor-like protein kinase At1g68400 | — |
| TRINITY_DN2044_c0_g1 | Rin-high | 5.95 | 1.67e-05 | MD15G1253900 | 6.05 | 1.11e-09 | | dehydrin Xero 1 | — |
| TRINITY_DN18168_c1_g1 | Rola-high | -5.93 | 1.11e-05 | MD15G1108300 | -6.00 | 1.62e-08 | yes | peroxidase 16-like | — |
| TRINITY_DN8435_c0_g1 | Rin-high | 5.92 | 5.60e-05 | MD14G1015900 | 5.96 | 2.54e-08 | | heat stress transcription factor A-3-like | — |
| TRINITY_DN62397_c0_g2 | Rola-high | -5.89 | 2.26e-03 | MD09G1118600 | -5.76 | 3.55e-04 | | glycine-rich protein-like | — |
| TRINITY_DN9496_c0_g1 | Rola-high | -5.88 | 2.81e-06 | MD08G1036400 | -4.99 | 1.75e-04 | yes | pathogenesis-related thaumatin-like protein 3.5 | — |
| TRINITY_DN12742_c0_g1 | Rola-high | -5.83 | 1.07e-08 | MD17G1190800 | -5.62 | 3.15e-13 | | sulfite exporter TauE/SafE family protein 3-like | — |
| TRINITY_DN8277_c4_g2 | Rola-high | -5.82 | 2.65e-06 | MD17G1259400 | -4.18 | 1.59e-14 | yes | cytochrome P450 736A117-like | — |
| TRINITY_DN7016_c0_g2 | Rola-high | -5.80 | 3.96e-03 | MD08G1030700 | -6.86 | 7.77e-05 | | zinc-finger homeodomain protein 2-like | — |
| TRINITY_DN25461_c0_g1 | Rola-high | -5.79 | 1.52e-06 | MD12G1049900 | -5.77 | 1.26e-10 | yes | GDSL esterase/lipase At5g45670-like | — |
| TRINITY_DN52873_c0_g1 | Rola-high | -5.79 | 3.73e-03 | MD06G1231300 | -6.29 | 7.14e-06 | | WUSCHEL-related homeobox 3-like | — |
| TRINITY_DN10215_c2_g1 | Rola-high | -5.78 | 2.38e-08 | MD10G1293600 | -5.80 | 1.04e-10 | | hypothetical protein DVH24_017234 | — |
| TRINITY_DN7183_c0_g1 | Rin-high | 5.74 | 4.59e-05 | MD07G1175000 | 5.99 | 5.11e-09 | | hypothetical protein DVH24_023959 | — |
| TRINITY_DN24616_c1_g1 | Rola-high | -5.72 | 6.55e-04 | MD06G1177500 | -5.70 | 6.10e-07 | | spermidine synthase 1-like | — |
| TRINITY_DN15097_c0_g1 | Rin-high | 5.67 | 2.88e-06 | MD00G1175200 | 6.84 | 4.03e-08 | | strychnine-10-hydroxylase-like | — |
| TRINITY_DN47279_c0_g1 | Rola-high | -5.65 | 7.86e-03 | MD13G1036800 | -2.83 | 2.53e-07 | yes | probable leucine-rich repeat receptor-like protein kinase At1g68400 | — |
| TRINITY_DN10967_c0_g1 | Rola-high | -5.64 | 3.64e-05 | MD14G1010300 | -5.67 | 8.62e-08 | yes | peroxidase 57-like | — |
| TRINITY_DN66456_c0_g1 | Rola-high | -5.64 | 3.82e-03 | MD05G1198800 | -5.32 | 2.78e-04 | yes | ethylene-responsive transcription factor ERF098-like | — |
| TRINITY_DN18323_c1_g1 | Rin-high | 5.63 | 1.54e-06 | MD04G1195900 | 5.24 | 1.48e-08 | | hypothetical protein DVH24_015128 | — |
| TRINITY_DN11137_c0_g1 | Rola-high | -5.62 | 9.00e-05 | MD00G1036800 | -5.48 | 1.91e-07 | | ABC transporter G family member 1-like | — |

| Trinity gene ID | Direction | Trinity | | GDDH13 protein match | | | Def. | Description | FCS-GX |
|---|---|---|---|---|---|---|---|---|---|
| | | log2FC | padj | gene ID | log2FC | padj | | | call |
| TRINITY_DN14471_c0_g1 | Rola-high | -5.61 | 9.55e-05 | MD15G1077000 | -5.27 | 1.83e-08 | | probable isoaspartyl peptidase/L-asparaginase 2 | — |
| TRINITY_DN5734_c1_g1 | Rola-high | -5.60 | 7.63e-03 | MD12G1196300 | -5.44 | 6.60e-05 | | protein SCARECROW-like | — |
| TRINITY_DN24450_c0_g1 | Rola-high | -5.59 | 1.36e-07 | MD05G1208600 | -4.98 | 7.53e-20 | | interactor of constitutive active ROPs 4-like | — |
| TRINITY_DN6439_c0_g1 | Rola-high | -5.59 | 7.94e-08 | MD08G1232300 | -7.96 | 3.38e-12 | | cocosin 1-like isoform X2 | — |
| TRINITY_DN8504_c1_g1 | Rola-high | -5.57 | 7.48e-06 | MD15G1049300 | -2.10 | 2.71e-09 | yes | LRR receptor-like serine/threonine-protein kinase ERECTA isoform X2 | — |
| TRINITY_DN6089_c0_g1 | Rola-high | -5.55 | 8.51e-07 | MD05G1113700 | -5.65 | 7.19e-11 | | auxin-induced protein 15A-like | — |
| TRINITY_DN52364_c0_g1 | Rola-high | -5.55 | 8.39e-04 | MD07G1041500 | -7.16 | 6.61e-08 | | squalene monooxygenase SE1-like | — |
| TRINITY_DN44460_c0_g1 | Rin-high | 5.51 | 2.71e-05 | MD05G1231700 | 5.57 | 4.57e-08 | yes | G-type lectin S-receptor-like serine/threonine-protein kinase At4g27290 isoform X1 | — |
| TRINITY_DN4764_c2_g1 | Rola-high | -5.50 | 3.97e-04 | MD12G1181900 | -4.79 | 6.98e-04 | | short-chain dehydrogenase reductase 2a-like | — |
| TRINITY_DN77883_c0_g1 | Rola-high | -5.47 | 4.26e-07 | MD09G1075500 | -5.69 | 1.68e-12 | yes | leucine-rich repeat extensin-like protein 5 isoform X1 | — |
| TRINITY_DN75769_c0_g1 | Rola-high | -5.47 | 6.67e-09 | MD02G1126900 | -5.57 | 4.99e-15 | yes | GDSL esterase/lipase EXL3-like | — |
| TRINITY_DN10085_c0_g3 | Rin-high | 5.46 | 2.09e-08 | MD16G1022200 | 5.22 | 6.31e-09 | yes | pectinesterase/pectinesterase inhibitor PPE8B | — |
| TRINITY_DN21964_c0_g1 | Rin-high | 5.45 | 8.15e-03 | MD01G1158600 | 3.75 | 3.60e-05 | | dehydration-responsive element-binding protein (DREB2) | — |
| TRINITY_DN55272_c0_g2 | Rola-high | -5.42 | 5.42e-05 | MD15G1106200 | -7.95 | 6.56e-10 | | UDP-glycosyltransferase 90A1-like | — |
| TRINITY_DN18970_c0_g1 | Rola-high | -5.40 | 8.94e-03 | MD06G1120700 | -5.54 | 1.06e-04 | yes | endoglucanase CX-like isoform X1 | — |
| TRINITY_DN16226_c0_g1 | Rin-high | 5.39 | 2.72e-03 | MD08G1062400 | 5.27 | 3.31e-05 | | TRINITY_DN16226_c0_g1 | — |
| TRINITY_DN1786_c0_g1 | Rola-high | -5.37 | 1.02e-03 | MD04G1173400 | -4.38 | 8.63e-06 | yes | expansin-A4-like | — |
| TRINITY_DN14530_c0_g1 | Rola-high | -5.35 | 2.88e-03 | MD09G1201800 | -3.87 | 2.83e-07 | | uncharacterized protein | — |
| TRINITY_DN2105_c2_g2 | Rola-high | -5.34 | 1.45e-07 | MD00G1093600 | -6.95 | 1.79e-10 | yes | GDSL esterase/lipase At5g45670-like | — |
| TRINITY_DN107802_c0_g1 | Rin-high | 5.34 | 1.74e-04 | MD11G1082600 | 5.33 | 8.43e-08 | | uncharacterized protein | — |
| TRINITY_DN25598_c0_g1 | Rola-high | -5.32 | 2.99e-03 | MD13G1170400 | -6.25 | 1.91e-06 | | protein LATERAL BRANCHING OXIDOREDUCTASE 1 | — |
| TRINITY_DN33783_c0_g1 | Rola-high | -5.32 | 5.00e-03 | MD09G1120000 | -5.65 | 1.33e-04 | | exopolygalacturonase-like | — |

| Trinity gene ID | Direction | Trinity | | GDDH13 protein match | | | Def. | Description | FCS-GX |
|---|---|---|---|---|---|---|---|---|---|
| | | log2FC | padj | gene ID | log2FC | padj | | | call |
| TRINITY_DN5158_c0_g1 | Rin-high | 5.31 | 9.83e-08 | MD10G1045100 | 7.33 | 2.13e-07 | | protein DETOXIFICATION 54 | — |
| TRINITY_DN29308_c0_g1 | Rola-high | -5.31 | 6.25e-03 | MD10G1243900 | -5.29 | 1.25e-04 | | berberine bridge enzyme-like 7 | — |
| TRINITY_DN38491_c0_g1 | Rin-high | 5.31 | 7.00e-04 | MD06G1198400 | 5.21 | 1.71e-06 | | uncharacterized protein At5g01610-like | — |
| TRINITY_DN36467_c0_g1 | Rola-high | -5.30 | 1.51e-03 | MD03G1189700 | -3.70 | 1.16e-05 | | kinesin-like protein KIN-12F | — |
| TRINITY_DN63417_c2_g1 | Rin-high | 5.29 | 2.57e-04 | MD11G1082600 | 5.33 | 8.43e-08 | | uncharacterized protein | — |
| TRINITY_DN9514_c1_g1 | Rola-high | -5.27 | 1.43e-03 | MD04G1051700 | -3.92 | 1.09e-05 | | hypothetical protein DVH24_001124 | — |
| TRINITY_DN105859_c0_g1 | Rin-high | 5.26 | 8.07e-06 | MD03G1081900 | 4.29 | 3.08e-11 | | UDP-glycosyltransferase 76E2-like | — |
| TRINITY_DN8699_c0_g1 | Rola-high | -5.25 | 4.37e-03 | MD03G1021500 | -4.56 | 2.23e-06 | | beta-glucosidase 12-like | — |
| TRINITY_DN63417_c1_g1 | Rin-high | 5.25 | 6.21e-04 | MD11G1082600 | 5.33 | 8.43e-08 | | uncharacterized protein | — |
| TRINITY_DN68_c0_g3 | Rola-high | -5.25 | 1.04e-03 | MD16G1160600 | -5.16 | 4.31e-09 | | major allergen Mal d 1.04 | — |
| TRINITY_DN9730_c0_g1 | Rola-high | -5.24 | 3.80e-08 | MD13G1023500 | -8.71 | 4.08e-09 | | hypothetical protein DVH24_040337 | — |
| TRINITY_DN21199_c0_g1 | Rola-high | -5.22 | 5.70e-04 | MD04G1091200 | -4.22 | 1.43e-09 | yes | xyloglucan endotransglucosylase/hydrolase protein 31-like | — |
| TRINITY_DN166788_c0_g1 | Rin-high | 5.21 | 8.09e-03 | MD11G1082600 | 5.33 | 8.43e-08 | | uncharacterized protein | — |
| TRINITY_DN28717_c0_g2 | Rola-high | -5.21 | 3.85e-08 | MD16G1107400 | -5.39 | 4.57e-17 | | very-long-chain 3-oxoacyl-CoA reductase 1-like | — |
| TRINITY_DN1589_c3_g1 | Rola-high | -5.20 | 5.47e-03 | MD17G1227000 | -4.59 | 7.67e-07 | | tubby-like protein 8 | — |
| TRINITY_DN17491_c0_g1 | Rola-high | -5.20 | 6.33e-05 | MD11G1014700 | -5.13 | 1.15e-07 | | ferritin-like catalase Nec2 | — |
| TRINITY_DN25803_c0_g1 | Rola-high | -5.18 | 5.03e-08 | MD05G1208600 | -4.98 | 7.53e-20 | | interactor of constitutive active ROPs 4-like | — |
| TRINITY_DN38927_c0_g1 | Rola-high | -5.17 | 1.24e-03 | MD15G1078000 | -4.24 | 1.28e-06 | yes | probable inactive receptor kinase At5g67200 | — |
| TRINITY_DN41892_c0_g1 | Rola-high | -5.16 | 5.78e-05 | MD11G1027400 | -4.70 | 2.85e-07 | | histone H3.2 | — |
| TRINITY_DN181094_c0_g1 | Rola-high | -5.10 | 7.36e-06 | MD11G1138800 | -5.19 | 5.09e-09 | | uncharacterized protein | — |
| TRINITY_DN13102_c0_g1 | Rola-high | -5.09 | 4.71e-03 | MD13G1130100 | -3.76 | 1.91e-06 | | histone-lysine N-methyltransferase, H3 lysine-9 specific SUVH4-like isoform X2 | — |
| TRINITY_DN17921_c0_g1 | Rola-high | -5.09 | 4.17e-04 | MD15G1077000 | -5.27 | 1.83e-08 | | probable isoaspartyl peptidase/L-asparaginase 2 | — |

| Trinity gene ID | Direction | Trinity | | GDDH13 protein match | | | Def. | Description | FCS-GX |
|---|---|---|---|---|---|---|---|---|---|
| | | log2FC | padj | gene ID | log2FC | padj | | | call |
| TRINITY_DN38982_c0_g1 | Rola-high | -5.08 | 5.59e-05 | MD07G1039600 | -5.12 | 5.74e-08 | | protein NRT1/ PTR FAMILY 5.4-like | — |
| TRINITY_DN68707_c0_g1 | Rin-high | 5.07 | 6.12e-06 | MD05G1257600 | 2.40 | 7.99e-06 | yes | probable LRR receptor-like serine/threonine-protein kinase At1g56130 | — |
| TRINITY_DN64099_c0_g1 | Rola-high | -5.07 | 1.74e-03 | MD13G1007700 | -4.86 | 2.52e-12 | | delta(24)-sterol reductase | — |
| TRINITY_DN11052_c0_g1 | Rola-high | -5.05 | 5.95e-10 | MD03G1068800 | -7.25 | 3.83e-16 | yes | putative serine/threonine-protein kinase-like protein CCR3 | — |
| TRINITY_DN2113_c0_g1 | Rin-high | 5.05 | 4.36e-05 | MD10G1270400 | 4.84 | 1.28e-07 | | alpha,alpha-trehalose-phosphate synthase [UDP-forming] 1-like isoform X1 | — |
| TRINITY_DN30907_c0_g1 | Rola-high | -5.04 | 3.63e-03 | MD08G1198000 | -4.00 | 1.75e-06 | | kinesin-like protein KIN-4C isoform X1 | — |
| TRINITY_DN11822_c0_g2 | Rola-high | -5.03 | 1.53e-07 | MD05G1348500 | -6.99 | 1.22e-07 | | L-ascorbate oxidase homolog | — |
| TRINITY_DN10437_c0_g1 | Rola-high | -5.03 | 6.01e-13 | MD09G1084600 | -5.25 | 5.34e-21 | | allene oxide cyclase, chloroplastic-like | — |
| TRINITY_DN14711_c0_g2 | Rin-high | 5.02 | 5.55e-03 | MD05G1329700 | 5.90 | 8.28e-06 | yes | G-type lectin S-receptor-like serine/threonine-protein kinase At1g61550 | — |
| TRINITY_DN8605_c0_g1 | Rola-high | -5.01 | 8.94e-03 | MD07G1153000 | -5.14 | 1.06e-04 | yes | laccase-6 | — |
| TRINITY_DN58011_c0_g1 | Rola-high | -5.00 | 3.17e-04 | MD07G1240000 | -4.89 | 7.22e-06 | | hypothetical protein DVH24_024483 | — |
| TRINITY_DN213219_c0_g1 | Rin-high | 4.98 | 4.58e-03 | MD11G1082600 | 5.33 | 8.43e-08 | | uncharacterized protein | — |
| TRINITY_DN29307_c0_g1 | Rola-high | -4.97 | 1.35e-04 | MD17G1212400 | -4.99 | 2.75e-07 | yes | receptor like protein 29-like | — |
| TRINITY_DN618_c0_g1 | Rin-high | 4.96 | 1.23e-04 | MD15G1032500 | 4.92 | 9.75e-07 | yes | calcium uniporter protein 2, mitochondrial | — |
| TRINITY_DN54237_c0_g1 | Rola-high | -4.96 | 6.17e-03 | MD10G1169100 | -5.95 | 1.73e-32 | | probable pectate lyase 8 | — |
| TRINITY_DN23019_c0_g1 | Rola-high | -4.95 | 9.87e-08 | MD01G1064900 | -4.31 | 1.84e-15 | yes | cytochrome P450 86A8 | — |
| TRINITY_DN29245_c0_g1 | Rola-high | -4.95 | 5.52e-05 | MD09G1156200 | -5.67 | 3.65e-08 | yes | receptor like protein 29-like | — |
| TRINITY_DN8202_c0_g2 | Rin-high | 4.94 | 1.88e-03 | MD02G1118000 | 5.05 | 1.03e-06 | | dehydrodolichyl diphosphate synthase CPT3-like | — |
| TRINITY_DN6030_c0_g3 | Rola-high | -4.93 | 2.64e-03 | MD00G1033700 | -5.37 | 2.74e-05 | | auxin-responsive protein SAUR50-like | — |
| TRINITY_DN44233_c0_g1 | Rola-high | -4.93 | 6.92e-03 | MD07G1259500 | -2.51 | 4.10e-06 | yes | LRR receptor-like serine/threonine-protein kinase ERECTA isoform X1 | — |
| TRINITY_DN22499_c0_g2 | Rola-high | -4.92 | 9.69e-04 | MD14G1212400 | -3.44 | 3.84e-06 | | replication protein A 70 kDa DNA-binding subunit B-like | — |
| TRINITY_DN8912_c0_g1 | Rola-high | -4.91 | 1.13e-03 | MD13G1191500 | -4.21 | 7.32e-05 | | thermospermine synthase ACAULIS5-like | — |

| Trinity gene ID | Direction | Trinity |  | GDDH13 protein match |  |  | Def. | Description | FCS-GX |
| --- | --- | --- | --- | --- | --- | --- | --- | --- | --- |
|  |  | log2FC | padj | gene ID | log2FC | padj |  |  | call |
| TRINITY_DN11371_c0_g1 | Rola-high | -4.91 | 6.93e-05 | MD06G1231200 | -5.42 | 2.73e-08 | yes | thaumatin-like protein 1 | — |
| TRINITY_DN41034_c0_g1 | Rola-high | -4.91 | 5.17e-03 | MD11G1119000 | -4.59 | 3.69e-05 | yes | laccase-17-like | — |
| TRINITY_DN18958_c0_g1 | Rola-high | -4.90 | 1.55e-04 | MD03G1273800 | -4.89 | 2.73e-07 |  | histone H2AX | — |
| TRINITY_DN21342_c0_g2 | Rola-high | -4.90 | 3.34e-09 | MD03G1169600 | -4.91 | 2.59e-16 |  | protein GAST1-like | — |
| TRINITY_DN27223_c1_g1 | Rola-high | -4.89 | 3.20e-03 | MD02G1183900 | -4.11 | 1.70e-07 |  | glucan endo-1,3-beta-glucosidase 12 | — |
| TRINITY_DN31316_c0_g1 | Rin-high | 4.89 | 6.40e-03 | MD08G1005500 | 4.69 | 2.74e-06 |  | hypothetical protein DVH24_020062 | — |
| TRINITY_DN20739_c2_g1 | Rola-high | -4.87 | 1.33e-03 | MD03G1215800 | -4.71 | 1.67e-05 |  | probable indole-3-acetic acid-amido synthetase GH3.1 | — |
| TRINITY_DN49975_c0_g1 | Rola-high | -4.87 | 3.04e-06 | MD15G1286200 | -4.85 | 1.50e-09 |  | uncharacterized acetyltransferase At3g50280-like | — |
| TRINITY_DN2026_c2_g1 | Rola-high | -4.86 | 2.51e-08 | MD17G1048600 | -4.78 | 6.05e-15 |  | uncharacterized protein | — |
| TRINITY_DN4055_c0_g1 | Rola-high | -4.86 | 1.06e-06 | MD07G1255800 | -5.14 | 4.30e-10 |  | 21 kDa protein-like | — |
| TRINITY_DN51592_c0_g2 | Rola-high | -4.84 | 2.55e-03 | MD16G1235500 | -4.74 | 1.75e-05 |  | 9-cis-epoxycarotenoid dioxygenase NCED2, chloroplastic-like | — |
| TRINITY_DN105273_c0_g1 | Rola-high | -4.83 | 3.21e-04 | MD02G1308000 | -4.14 | 9.11e-08 |  | DNA topoisomerase 2 isoform X2 | — |
| TRINITY_DN41656_c0_g1 | Rola-high | -4.82 | 1.25e-06 | MD00G1134000 | -4.69 | 5.02e-10 |  | 21 kDa protein-like | — |
| TRINITY_DN1459_c1_g1 | Rola-high | -4.82 | 7.61e-14 | MD17G1103100 | -4.81 | 9.81e-25 | yes | endoglucanase 11-like | — |
| TRINITY_DN34677_c0_g1 | Rola-high | -4.81 | 1.43e-04 | MD11G1027400 | -4.70 | 2.85e-07 |  | histone H3.2 | — |
| TRINITY_DN7809_c0_g1 | Rin-high | 4.80 | 1.07e-07 | MD16G1095000 | 2.09 | 1.88e-03 |  | galactinol synthase 1 | — |
| TRINITY_DN6671_c0_g2 | Rola-high | -4.79 | 2.14e-07 | MD09G1250100 | -4.87 | 1.46e-13 | yes | expansin-B3-like | — |
| TRINITY_DN880_c2_g1 | Rin-high | 4.78 | 2.71e-06 | MD05G1301600 | 4.92 | 7.34e-11 |  | heavy metal-associated isoprenylated plant protein 28-like | — |
| TRINITY_DN5441_c2_g1 | Rola-high | -4.78 | 1.51e-03 | MD11G1119000 | -4.59 | 3.69e-05 | yes | laccase-17-like | — |
| TRINITY_DN118173_c0_g1 | Rola-high | -4.78 | 4.21e-05 | MD09G1199000 | -4.78 | 1.69e-08 |  | PREDICTED: CARUB_v10012353mg | n/a |
| TRINITY_DN8241_c0_g1 | Rola-high | -4.76 | 5.27e-10 | MD15G1182100 | -4.71 | 2.24e-17 |  | probable protein ABIL5 | — |
| TRINITY_DN7127_c3_g1 | Rola-high | -4.74 | 4.67e-03 | MD13G1123800 | -3.09 | 6.91e-07 |  | hypothetical protein DVH24_025134 | — |

| Trinity gene ID | Direction | Trinity | | GDDH13 protein match | | | Def. | Description | FCS-GX |
|---|---|---|---|---|---|---|---|---|---|
| | | log2FC | padj | gene ID | log2FC | padj | | | call |
| TRINITY_DN9770_c1_g1 | Rola-high | -4.74 | 2.51e-04 | MD11G1146700 | -5.56 | 3.74e-09 | yes | xyloglucan endotransglucosylase/hydrolase protein 31-like | — |
| TRINITY_DN2017_c0_g1 | Rola-high | -4.74 | 7.03e-05 | MD08G1064300 | -5.23 | 4.56e-08 | | protein IQ-DOMAIN 5-like isoform X1 | — |
| TRINITY_DN221981_c0_g1 | Rola-high | -4.71 | 2.03e-05 | MD04G1174000 | -4.64 | 6.67e-09 | yes | lipid transfer protein precursor | — |
| TRINITY_DN34943_c0_g1 | Rin-high | 4.70 | 1.94e-04 | MD03G1267900 | 4.68 | 2.97e-07 | | adenylate isopentenyltransferase | — |
| TRINITY_DN237426_c0_g1 | Rola-high | -4.70 | 5.56e-04 | MD17G1259400 | -4.18 | 1.59e-14 | yes | cytochrome P450 736A117-like | — |
| TRINITY_DN46216_c0_g1 | Rola-high | -4.69 | 1.16e-04 | MD13G1231000 | -3.86 | 1.09e-09 | | probable pectate lyase 8 | — |
| TRINITY_DN58695_c0_g1 | Rola-high | -4.69 | 8.85e-03 | MD15G1434400 | -4.74 | 2.82e-05 | | protein TIFY 3B-like | — |
| TRINITY_DN838_c0_g1 | Rin-high | 4.69 | 4.45e-04 | MD16G1093600 | 4.76 | 6.52e-07 | yes | serine/threonine-protein kinase BLUS1-like isoform X2 | — |
| TRINITY_DN6176_c0_g1 | Rin-high | 4.68 | 4.56e-03 | MD15G1200400 | 4.57 | 5.27e-06 | | nudix hydrolase 8-like | — |
| TRINITY_DN60422_c0_g2 | Rin-high | 4.68 | 2.77e-05 | MD07G1236200 | 1.98 | 3.88e-08 | yes | probable LRR receptor-like serine/threonine-protein kinase At3g47570 isoform X2 | — |
| TRINITY_DN18888_c0_g1 | Rola-high | -4.68 | 3.34e-09 | MD13G1191400 | -5.32 | 8.13e-20 | | uncharacterized protein C24B11.05-like | — |
| TRINITY_DN108137_c0_g2 | Rola-high | -4.68 | 5.65e-03 | MD12G1219500 | -3.28 | 2.54e-04 | yes | putative disease resistance RPP13-like protein 1 isoform X5 | — |
| TRINITY_DN14756_c0_g1 | Rin-high | 4.68 | 7.11e-03 | MD11G1054100 | 4.65 | 5.02e-05 | yes | non-specific lipid-transfer protein 1-like | — |
| TRINITY_DN8848_c0_g1 | Rola-high | -4.68 | 1.88e-04 | MD11G1027400 | -4.70 | 2.85e-07 | | histone H3.2 | — |
| TRINITY_DN40485_c0_g2 | Rola-high | -4.68 | 2.23e-04 | MD15G1420300 | -4.49 | 4.27e-08 | | TRINITY_DN40485_c0_g2 | — |
| TRINITY_DN13876_c0_g2 | Rin-high | 4.65 | 5.05e-03 | MD00G1011500 | 4.83 | 5.27e-07 | | ankyrin repeat-containing protein ITN1-like isoform X1 | — |
| TRINITY_DN15562_c0_g1 | Rola-high | -4.65 | 7.10e-04 | MD08G1003500 | -4.41 | 5.10e-06 | | protein ALTERED PHOSPHATE STARVATION RESPONSE 1-like | — |
| TRINITY_DN15502_c1_g1 | Rola-high | -4.63 | 8.41e-03 | MD17G1061000 | -4.18 | 4.40e-05 | | phenolic glucoside malonyltransferase 1-like | — |
| TRINITY_DN8330_c0_g2 | Rola-high | -4.63 | 1.44e-05 | MD05G1348500 | -6.99 | 1.22e-07 | | L-ascorbate oxidase homolog | — |
| TRINITY_DN6608_c0_g1 | Rola-high | -4.63 | 1.33e-04 | MD06G1103000 | -4.73 | 5.15e-08 | | F-box/kelch-repeat protein At3g06240-like | — |
| TRINITY_DN43406_c0_g1 | Rin-high | 4.62 | 7.27e-03 | MD04G1182200 | 3.62 | 3.10e-06 | yes | leucine-rich repeat receptor protein kinase HPCA1-like | — |

| Trinity gene ID | Direction | Trinity | | GDDH13 protein match | | | Def. | Description | FCS-GX |
|---|---|---|---|---|---|---|---|---|---|
| | | log2FC | padj | gene ID | log2FC | padj | | | call |
| TRINITY_DN13109_c0_g1 | Rola-high | -4.61 | 2.75e-03 | MD11G1291000 | -4.05 | 2.58e-05 | | E3 ubiquitin-protein ligase SP1-like | — |
| TRINITY_DN40881_c0_g1 | Rola-high | -4.60 | 1.06e-03 | MD17G1045500 | -3.48 | 1.38e-03 | | AAA-ATPase ASD, mitochondrial-like | — |
| TRINITY_DN3142_c1_g2 | Rin-high | 4.57 | 3.57e-04 | MD15G1027000 | 4.39 | 5.32e-07 | | protein RADIALIS-like 3 | — |
| TRINITY_DN43001_c0_g1 | Rin-high | 4.57 | 6.29e-03 | MD10G1034200 | 4.14 | 5.11e-06 | | protein GIGANTEA-like | — |
| TRINITY_DN3562_c1_g2 | Rola-high | -4.56 | 4.37e-06 | MD02G1315200 | -4.94 | 1.51e-13 | yes | leucine-rich repeat receptor-like protein | — |
| TRINITY_DN3562_c1_g1 | Rola-high | -4.55 | 7.22e-06 | MD07G1006700 | -4.33 | 1.59e-09 | yes | leucine-rich repeat receptor-like protein | — |
| TRINITY_DN81108_c0_g1 | Rin-high | 4.55 | 5.42e-05 | MD11G1109100 | 4.17 | 1.73e-08 | | metallothionein-like protein type 2 | — |
| TRINITY_DN15397_c0_g2 | Rola-high | -4.55 | 3.13e-04 | MD09G1049500 | -4.00 | 5.66e-07 | | tubulin alpha chain isoform X2 | — |
| TRINITY_DN1119_c2_g3 | Rin-high | 4.55 | 1.42e-03 | MD15G1139800 | 3.91 | 1.13e-05 | | ammonium transporter 2-like | — |
| TRINITY_DN3971_c0_g1 | Rola-high | -4.53 | 5.16e-04 | MD04G1000500 | -4.47 | 7.01e-07 | yes | leucine-rich repeat extensin-like protein 4 | — |
| TRINITY_DN50530_c0_g1 | Rola-high | -4.52 | 2.69e-04 | MD11G1200000 | -4.46 | 7.71e-07 | | classical arabinogalactan protein 9-like | — |
| TRINITY_DN7042_c0_g1 | Rola-high | -4.52 | 9.53e-06 | MD14G1194100 | -4.56 | 2.34e-10 | | protein NRT1/ PTR FAMILY 4.6-like | — |
| TRINITY_DN10525_c0_g1 | Rola-high | -4.51 | 5.75e-08 | MD01G1064900 | -4.31 | 1.84e-15 | yes | cytochrome P450 86A8 | — |
| TRINITY_DN31016_c0_g1 | Rola-high | -4.50 | 2.94e-04 | MD17G1155800 | -3.19 | 2.54e-06 | | ATPase 10, plasma membrane-type-like | — |
| TRINITY_DN13232_c0_g2 | Rola-high | -4.50 | 7.27e-03 | MD02G1001900 | -4.51 | 6.45e-05 | | 2-alkenal reductase (NADP(+)-dependent)-like | — |
| TRINITY_DN179957_c0_g3 | Rin-high | 4.49 | 5.55e-03 | MD11G1109100 | 4.17 | 1.73e-08 | | metallothionein-like protein type 2 | — |
| TRINITY_DN2969_c1_g2 | Rola-high | -4.49 | 8.42e-04 | MD10G1067800 | -4.00 | 2.48e-10 | | beta-galactosidase 5-like isoform X1 | — |
| TRINITY_DN122722_c0_g1 | Rola-high | -4.48 | 3.46e-03 | MD13G1256400 | -4.46 | 1.35e-05 | | hydroxyproline O-galactosyltransferase GALT3 isoform X1 | — |
| TRINITY_DN35002_c0_g1 | Rola-high | -4.47 | 5.60e-03 | MD00G1014900 | -4.67 | 2.32e-05 | yes | putative laccase-9 | — |
| TRINITY_DN47845_c0_g1 | Rola-high | -4.47 | 1.70e-04 | MD04G1005400 | -4.45 | 1.08e-07 | | glycine-rich cell wall structural protein-like isoform X4 | — |
| TRINITY_DN2144_c7_g1 | Rola-high | -4.47 | 4.88e-04 | MD03G1052400 | -5.01 | 9.06e-11 | yes | expansin-A4-like precursor | — |
| TRINITY_DN23208_c0_g1 | Rola-high | -4.47 | 2.40e-05 | MD13G1171400 | -7.13 | 2.82e-05 | yes | proline-rich receptor-like protein kinase PERK1 isoform X2 | — |
| TRINITY_DN35821_c0_g1 | Rin-high | 4.45 | 7.52e-04 | MD05G1196600 | 4.50 | 5.69e-07 | | patellin-4 isoform X2 | — |

| Trinity gene ID | Direction | Trinity | | GDDH13 protein match | | | Def. | Description | FCS-GX |
|---|---|---|---|---|---|---|---|---|---|
| | | log2FC | padj | gene ID | log2FC | padj | | | call |
| TRINITY_DN23280_c0_g1 | Rin-high | 4.45 | 5.51e-05 | MD06G1163600 | 4.48 | 3.25e-13 | yes | cytochrome P450 736A117-like | — |
| TRINITY_DN44001_c0_g1 | Rola-high | -4.45 | 9.55e-05 | MD04G1234700 | -8.10 | 1.11e-09 | yes | GDSL esterase/lipase LTL1-like | — |
| TRINITY_DN48092_c0_g1 | Rola-high | -4.44 | 4.37e-06 | MD10G1125200 | -4.16 | 4.78e-09 | | probable aquaporin PIP2-8 | — |
| TRINITY_DN6326_c0_g1 | Rola-high | -4.44 | 1.85e-03 | MD07G1053200 | -2.67 | 7.12e-05 | yes | probable pectinesterase/pectinesterase inhibitor 47 | — |
| TRINITY_DN27757_c0_g1 | Rola-high | -4.44 | 4.95e-03 | MD02G1127900 | -3.97 | 1.06e-06 | | protein POLLENLESS 3-LIKE 2-like | — |
| TRINITY_DN63582_c0_g1 | Rola-high | -4.43 | 1.90e-03 | MD12G1188600 | -1.50 | 8.14e-05 | yes | probably inactive leucine-rich repeat receptor-like protein kinase IMK2 | — |
| TRINITY_DN8804_c0_g1 | Rola-high | -4.43 | 3.42e-04 | MD11G1026900 | -4.59 | 3.28e-07 | | histone H3.2 | — |
| TRINITY_DN104418_c0_g1 | Rin-high | 4.43 | 7.48e-05 | MD04G1052600 | 4.68 | 1.51e-09 | yes | expansin-like B1 | — |
| TRINITY_DN48433_c0_g1 | Rola-high | -4.43 | 6.72e-04 | MD09G1049500 | -4.00 | 5.66e-07 | | tubulin alpha-2 chain | — |
| TRINITY_DN3005_c0_g2 | Rin-high | 4.42 | 3.09e-04 | MD16G1022200 | 5.22 | 6.31e-09 | yes | pectinesterase | — |
| TRINITY_DN29862_c0_g1 | Rola-high | -4.41 | 3.86e-06 | MD15G1134300 | -4.40 | 2.82e-10 | | ABC transporter G family member 5 | — |
| TRINITY_DN14750_c0_g1 | Rin-high | 4.40 | 3.88e-10 | MD13G1101300 | 4.13 | 8.05e-18 | yes | RAF-like serine/threonine-protein kinase PRAF | — |
| TRINITY_DN44332_c0_g1 | Rin-high | 4.40 | 6.21e-05 | MD04G1195900 | 5.24 | 1.48e-08 | | hypothetical protein DVH24_015128 | — |
| TRINITY_DN157494_c0_g1 | Rola-high | -4.40 | 3.78e-06 | MD02G1315200 | -4.94 | 1.51e-13 | yes | leucine-rich repeat receptor-like protein | — |
| TRINITY_DN21266_c0_g1 | Rola-high | -4.40 | 1.94e-03 | MD16G1080900 | -3.13 | 3.97e-05 | yes | LEAF RUST 10 DISEASE-RESISTANCEUS RECEPTOR-LIKE PROTEIN KINASE-like 1.5 | — |
| TRINITY_DN39718_c0_g1 | Rola-high | -4.38 | 2.34e-03 | MD03G1068100 | -5.39 | 3.84e-06 | | beta-glucosidase 13-like | — |
| TRINITY_DN8381_c0_g2 | Rola-high | -4.38 | 2.23e-03 | MD02G1146800 | -4.83 | 3.64e-05 | | GATA transcription factor 19-like | — |
| TRINITY_DN4116_c0_g1 | Rola-high | -4.38 | 2.01e-07 | MD04G1147900 | -4.98 | 1.27e-10 | | BAHD acyltransferase DCR | — |
| TRINITY_DN2543_c0_g1 | Rola-high | -4.35 | 8.71e-06 | MD02G1100200 | -4.13 | 7.54e-10 | | auxin-responsive protein SAUR76-like | — |
| TRINITY_DN82969_c0_g1 | Rola-high | -4.35 | 1.32e-04 | MD03G1284600 | -4.32 | 6.65e-08 | | ferritin-like catalase Nec2 | — |
| TRINITY_DN80973_c0_g1 | Rin-high | 4.35 | 9.76e-04 | MD09G1221600 | 3.87 | 7.88e-09 | | uncharacterized protein | — |
| TRINITY_DN1805_c0_g1 | Rola-high | -4.34 | 1.10e-03 | MD13G1213100 | -4.39 | 5.21e-06 | yes | ethylene-responsive transcription factor 1B-like | — |

| Trinity gene ID | Direction | Trinity | | GDDH13 protein match | | | Def. | Description | FCS-GX |
|---|---|---|---|---|---|---|---|---|---|
| | | log2FC | padj | gene ID | log2FC | padj | | | call |
| TRINITY_DN40790_c0_g1 | Rola-high | -4.34 | 5.87e-03 | MD16G1197800 | -4.31 | 3.28e-07 | | protein tesmin/TSO1-like CXC 2 isoform X2 | — |
| TRINITY_DN12526_c0_g1 | Rola-high | -4.33 | 2.49e-06 | MD15G1172600 | -4.93 | 1.01e-07 | | AP2 domain class transcription factor | — |
| TRINITY_DN35651_c0_g1 | Rola-high | -4.33 | 4.33e-04 | MD13G1161700 | -2.44 | 5.83e-04 | | major strawberry allergen Fra a 1.07-like | — |
| TRINITY_DN8602_c0_g1 | Rola-high | -4.32 | 5.52e-03 | MD04G1221900 | -5.66 | 6.91e-10 | | trihelix transcription factor GTL2-like | — |
| TRINITY_DN12042_c0_g1 | Rola-high | -4.32 | 9.06e-04 | MD09G1113100 | -3.94 | 1.15e-04 | yes | endoglucanase 11-like | — |
| TRINITY_DN28031_c0_g1 | Rola-high | -4.32 | 6.12e-03 | MD06G1004900 | -4.57 | 6.34e-06 | yes | NAC domain-containing protein 92-like isoform X1 | — |
| TRINITY_DN18201_c0_g1 | Rin-high | 4.31 | 1.52e-03 | MD16G1022200 | 5.22 | 6.31e-09 | | hypothetical protein DVH24_007574 | — |
| TRINITY_DN7750_c0_g1 | Rola-high | -4.31 | 5.52e-05 | MD07G1301700 | -4.94 | 2.15e-05 | | heavy metal-associated isoprenylated plant protein 7-like | — |
| TRINITY_DN39041_c0_g1 | Rola-high | -4.30 | 9.92e-03 | MD13G1197800 | -4.21 | 1.10e-06 | | protein tesmin/TSO1-like CXC 2 | — |
| TRINITY_DN11444_c0_g1 | Rola-high | -4.29 | 1.12e-04 | MD13G1052500 | -4.19 | 4.53e-08 | | protein STRICTOSIDINE SYNTHASE-LIKE 10-like | — |
| TRINITY_DN16772_c1_g4 | Rola-high | -4.29 | 3.52e-07 | MD15G1044100 | -4.24 | 1.36e-12 | | aspartyl protease family protein At5g10770-like | — |
| TRINITY_DN11672_c1_g2 | Rola-high | -4.28 | 7.90e-03 | MD09G1022800 | -3.67 | 3.09e-06 | | hypothetical protein DVH24_031698 | — |
| TRINITY_DN8474_c0_g1 | Rin-high | 4.28 | 2.05e-04 | MD17G1030400 | 4.14 | 8.15e-08 | | chaperone protein dnaJ C76, chloroplastic isoform X2 | — |
| TRINITY_DN14893_c0_g2 | Rola-high | -4.28 | 9.76e-04 | MD04G1201800 | -4.30 | 3.64e-06 | | histone H2A | — |
| TRINITY_DN12294_c1_g1 | Rola-high | -4.28 | 2.98e-03 | MD05G1060000 | -3.82 | 4.15e-06 | | protein BREAST CANCER SUSCEPTIBILITY 1 homolog isoform X2 | — |
| TRINITY_DN8632_c0_g1 | Rola-high | -4.28 | 6.53e-05 | MD16G1112600 | -5.26 | 1.29e-09 | | remorin 4.1 isoform X3 | — |
| TRINITY_DN21977_c0_g1 | Rola-high | -4.28 | 7.32e-13 | MD04G1148900 | -2.99 | 3.73e-27 | | glycerol-3-phosphate acyltransferase RAM2-like | — |
| TRINITY_DN55118_c0_g1 | Rola-high | -4.27 | 4.89e-03 | MD17G1090100 | -4.49 | 9.74e-06 | | tricyclene synthase EBOS, chloroplastic-like isoform X2 | — |
| TRINITY_DN25940_c0_g1 | Rin-high | 4.26 | 5.70e-03 | MD10G1034200 | 4.14 | 5.11e-06 | | protein GIGANTEA-like | — |
| TRINITY_DN2284_c1_g1 | Rola-high | -4.26 | 4.84e-04 | MD15G1243100 | -4.46 | 3.22e-07 | | protein POLLENLESS 3-LIKE 2-like | — |
| TRINITY_DN14085_c0_g1 | Rola-high | -4.26 | 4.16e-04 | MD01G1124600 | -4.58 | 3.08e-07 | | histone H4 | — |

| Trinity gene ID | Direction | Trinity | | GDDH13 protein match | | | Def. | Description | FCS-GX |
|---|---|---|---|---|---|---|---|---|---|
| | | log2FC | padj | gene ID | log2FC | padj | | | call |
| TRINITY_DN23445_c0_g1 | Rola-high | -4.26 | 1.13e-03 | MD02G1183900 | -4.11 | 1.70e-07 | | glucan endo-1,3-beta-glucosidase 12 | — |
| TRINITY_DN9368_c0_g1 | Rin-high | 4.26 | 6.50e-03 | MD11G1300300 | 3.80 | 5.92e-04 | yes | monothiol glutaredoxin-S6-like | — |
| TRINITY_DN33556_c0_g1 | Rin-high | 4.25 | 1.33e-05 | MD03G1227300 | 4.63 | 2.88e-11 | | histone H3.2 | — |
| TRINITY_DN937_c1_g1 | Rola-high | -4.25 | 2.03e-09 | MD15G1207800 | -3.82 | 5.85e-11 | | pathogen-associated molecular patterns-induced protein A70-like | — |
| TRINITY_DN2726_c0_g1 | Rin-high | 4.25 | 8.90e-04 | MD00G1011500 | 4.83 | 5.27e-07 | | ankyrin repeat-containing protein ITN1-like | — |
| TRINITY_DN41791_c0_g3 | Rin-high | 4.25 | 2.73e-03 | MD10G1270400 | 4.84 | 1.28e-07 | | alpha,alpha-trehalose-phosphate synthase [UDP-forming] 1-like isoform X1 | — |
| TRINITY_DN26924_c0_g1 | Rola-high | -4.25 | 1.91e-03 | MD11G1012800 | -3.56 | 2.85e-06 | | neurofilament heavy polypeptide-like isoform X1 | — |
| TRINITY_DN38314_c0_g1 | Rola-high | -4.24 | 3.78e-06 | MD01G1188800 | -4.15 | 5.66e-10 | | subtilisin-like protease SBT1.3 | — |
| TRINITY_DN72165_c0_g1 | Rola-high | -4.24 | 5.22e-08 | MD12G1162100 | -3.89 | 9.22e-23 | | glycerol-3-phosphate 2-O-acyltransferase 6-like | — |
| TRINITY_DN37378_c0_g1 | Rola-high | -4.24 | 5.47e-03 | MD10G1260700 | -4.84 | 1.42e-05 | | protein TIFY 5A-like | — |
| TRINITY_DN4764_c1_g1 | Rola-high | -4.23 | 8.78e-04 | MD16G1197800 | -4.31 | 3.28e-07 | | protein tesmin/TSO1-like CXC 2 isoform X2 | — |
| TRINITY_DN38131_c0_g1 | Rola-high | -4.22 | 2.24e-06 | MD16G1048900 | -4.19 | 7.64e-11 | | aspartyl protease family protein 2 | — |
| TRINITY_DN6180_c0_g1 | Rola-high | -4.22 | 1.95e-04 | MD17G1265300 | -4.20 | 1.46e-08 | yes | peroxidase P7-like | — |
| TRINITY_DN13801_c0_g1 | Rola-high | -4.22 | 7.42e-05 | MD04G1201400 | -4.50 | 1.52e-08 | | formin-like protein 11 | — |
| TRINITY_DN20914_c0_g1 | Rola-high | -4.20 | 8.42e-03 | MD11G1119000 | -4.59 | 3.69e-05 | | hypothetical protein DVH24_006030 | — |
| TRINITY_DN32751_c0_g1 | Rola-high | -4.19 | 5.93e-07 | MD08G1071900 | -4.24 | 6.61e-15 | yes | LRR receptor-like serine/threonine-protein kinase ERECTA isoform X1 | — |
| TRINITY_DN15776_c0_g1 | Rin-high | 4.19 | 8.74e-03 | MD06G1096000 | 3.04 | 4.11e-05 | | aluminum-activated malate transporter 9-like | — |
| TRINITY_DN4444_c0_g2 | Rola-high | -4.19 | 1.63e-04 | MD11G1107600 | -4.11 | 1.07e-07 | | glucan endo-1,3-beta-glucosidase | — |
| TRINITY_DN13631_c1_g1 | Rola-high | -4.19 | 8.55e-04 | MD03G1251300 | -4.16 | 1.17e-06 | | histone-lysine N-methyltransferase ATX2-like isoform X1 | — |
| TRINITY_DN17339_c0_g1 | Rola-high | -4.16 | 4.82e-03 | MD13G1186500 | -3.03 | 4.34e-07 | yes | mitogen-activated protein kinase kinase kinase NPK1-like | — |
| TRINITY_DN502_c1_g1 | Rola-high | -4.16 | 9.65e-05 | MD11G1189300 | -5.76 | 4.95e-11 | | zinc finger CCCH domain-containing protein 53-like isoform X1 | — |

| Trinity gene ID | Direction | Trinity | | GDDH13 protein match | | | Def. | Description | FCS-GX |
|---|---|---|---|---|---|---|---|---|---|
| | | log2FC | padj | gene ID | log2FC | padj | | | call |
| TRINITY_DN6489_c0_g1 | Rola-high | -4.15 | 8.03e-07 | MD16G1095400 | -3.88 | 6.04e-16 | | polygalacturonase At1g48100 | — |
| TRINITY_DN46517_c0_g1 | Rola-high | -4.14 | 1.94e-03 | MD15G1234900 | -4.27 | 2.11e-06 | | uncharacterized protein | — |
| TRINITY_DN1433_c4_g1 | Rola-high | -4.14 | 1.00e-04 | MD05G1306500 | -4.17 | 2.31e-08 | yes | peroxidase 41-like | — |
| TRINITY_DN223830_c0_g1 | Rola-high | -4.13 | 8.89e-03 | MD09G1285100 | -4.10 | 5.53e-05 | | stachyose synthase | — |
| TRINITY_DN46550_c0_g1 | Rola-high | -4.13 | 9.01e-04 | MD03G1068200 | -3.76 | 9.52e-03 | | beta-glucosidase 13-like | — |
| TRINITY_DN22894_c0_g1 | Rin-high | 4.11 | 4.60e-03 | MD10G1034200 | 4.14 | 5.11e-06 | | protein GIGANTEA-like | — |
| TRINITY_DN24392_c1_g1 | Rola-high | -4.10 | 8.67e-03 | MD17G1045500 | -3.48 | 1.38e-03 | | AAA-ATPase At3g28580-like | — |
| TRINITY_DN13803_c0_g2 | Rola-high | -4.10 | 6.75e-04 | MD15G1040900 | -4.15 | 8.89e-07 | | early nodulin-like protein 15 | — |
| TRINITY_DN23707_c0_g1 | Rola-high | -4.10 | 9.88e-03 | MD11G1147000 | -4.14 | 3.37e-05 | | protein NUCLEAR FUSION DEFECTIVE 4-like | — |
| TRINITY_DN1045_c0_g1 | Rin-high | 4.10 | 5.67e-03 | MD13G1279400 | 6.96 | 6.51e-07 | | amino acid transporter AVT1D-like isoform X1 | — |
| TRINITY_DN7203_c2_g1 | Rola-high | -4.10 | 1.08e-04 | MD14G1048900 | -4.45 | 7.82e-10 | | IQ domain-containing protein IQM6 | — |
| TRINITY_DN27390_c0_g1 | Rola-high | -4.10 | 5.86e-03 | MD12G1196300 | -5.44 | 6.60e-05 | | protein SCARECROW-like | — |
| TRINITY_DN3757_c0_g1 | Rin-high | 4.09 | 1.21e-04 | MD10G1056200 | 2.75 | 4.91e-10 | | MADS-box transcription factor 23-like | — |
| TRINITY_DN5206_c0_g1 | Rin-high | 4.09 | 5.00e-03 | MD03G1059200 | 4.43 | 6.98e-06 | yes | peroxidase 4-like | — |
| TRINITY_DN5575_c0_g1 | Rola-high | -4.09 | 3.57e-05 | MD17G1011000 | -4.12 | 1.49e-09 | | uncharacterized protein | — |
| TRINITY_DN14946_c0_g1 | Rola-high | -4.08 | 8.67e-03 | MD15G1218800 | -3.77 | 5.76e-07 | | subtilisin-like protease SBT1.8 | — |
| TRINITY_DN45551_c0_g1 | Rin-high | 4.08 | 3.90e-05 | MD08G1058900 | 4.17 | 1.69e-09 | | brassinosteroid-related acyltransferase 1-like | — |
| TRINITY_DN10570_c0_g3 | Rola-high | -4.08 | 4.89e-04 | MD10G1239200 | -3.88 | 1.21e-06 | | hypothetical protein DVH24_022142 | — |
| TRINITY_DN10786_c0_g1 | Rola-high | -4.07 | 1.82e-03 | MD05G1039300 | -3.98 | 6.01e-06 | yes | GDSL esterase/lipase At1g29670-like | — |
| TRINITY_DN21660_c0_g1 | Rin-high | 4.06 | 2.32e-03 | MD13G1182800 | 4.24 | 7.89e-07 | | adenylate isopentenyltransferase | — |
| TRINITY_DN30803_c0_g1 | Rola-high | -4.06 | 2.62e-03 | MD13G1216600 | -3.19 | 1.81e-08 | | KIN14B-interacting protein At4g14310 | — |
| TRINITY_DN1855_c0_g1 | Rola-high | -4.05 | 4.48e-03 | MD15G1190600 | -4.35 | 1.86e-05 | | rac-like GTP-binding protein ARAC7 | — |
| TRINITY_DN5367_c0_g1 | Rola-high | -4.05 | 7.68e-06 | MD17G1049300 | -7.27 | 3.37e-06 | | transcription factor PRE6-like | — |

| Trinity gene ID | Direction | Trinity | | GDDH13 protein match | | | Def. | Description | FCS-GX |
|---|---|---|---|---|---|---|---|---|---|
| | | log2FC | padj | gene ID | log2FC | padj | | | call |
| TRINITY_DN2387_c1_g1 | Rola-high | -4.04 | 1.33e-04 | MD10G1122000 | -4.19 | 8.85e-05 | | endo-1,4-beta-xylanase 5-like | — |
| TRINITY_DN4995_c0_g1 | Rola-high | -4.03 | 2.64e-03 | MD17G1123400 | -3.91 | 3.84e-06 | | uncharacterized protein | — |
| TRINITY_DN33195_c0_g1 | Rola-high | -4.03 | 5.35e-03 | MD17G1129800 | -4.64 | 8.33e-06 | | uncharacterized protein | — |
| TRINITY_DN7210_c1_g1 | Rola-high | -4.02 | 1.56e-06 | MD07G1228100 | -3.20 | 8.34e-06 | yes | probable leucine-rich repeat receptor-like protein kinase IMK3 | — |
| TRINITY_DN7666_c0_g1 | Rola-high | -4.02 | 1.61e-04 | MD12G1032000 | -2.61 | 8.00e-05 | | B3 domain-containing protein Os03g0120900-like | — |
| TRINITY_DN3187_c0_g2 | Rola-high | -4.02 | 2.25e-09 | MD15G1076900 | -3.93 | 1.49e-18 | | hypothetical protein DVH24_012498 | — |
| TRINITY_DN32239_c0_g1 | Rola-high | -4.01 | 4.56e-05 | MD09G1200500 | -3.67 | 2.63e-09 | | acid phosphatase 1-like | — |
| TRINITY_DN6843_c0_g1 | Rola-high | -4.01 | 3.45e-03 | MD15G1066500 | -4.77 | 1.15e-05 | | kinesin-like protein NACK1 | — |
| TRINITY_DN24878_c0_g1 | Rola-high | -4.00 | 1.74e-04 | MD01G1039400 | -4.02 | 2.63e-08 | | uncharacterized protein At4g38062-like | — |
| TRINITY_DN2093_c0_g2 | Rola-high | -3.99 | 4.68e-04 | MD15G1099400 | -3.45 | 1.44e-08 | | protein ALTERED PHOSPHATE STARVATION RESPONSE 1-like | — |
| TRINITY_DN7833_c0_g1 | Rola-high | -3.99 | 5.77e-07 | MD07G1277100 | -2.55 | 1.08e-03 | | protein DETOXIFICATION 35-like | — |
| TRINITY_DN4232_c0_g1 | Rola-high | -3.98 | 5.44e-12 | MD03G1214100 | -3.78 | 1.66e-20 | | putative anthocyanidin reductase isoform X1 | — |
| TRINITY_DN11592_c0_g1 | Rola-high | -3.98 | 1.83e-04 | MD04G1234400 | -4.23 | 4.44e-07 | yes | GDSL esterase/lipase LTL1-like | — |
| TRINITY_DN15016_c0_g2 | Rola-high | -3.98 | 9.73e-03 | MD09G1075500 | -5.69 | 1.68e-12 | yes | leucine-rich repeat extensin-like protein 5 isoform X1 | — |
| TRINITY_DN26634_c0_g1 | Rola-high | -3.98 | 7.73e-03 | MD15G1248100 | -1.29 | 9.88e-05 | | kinesin-like protein KIN-7E | — |
| TRINITY_DN50606_c0_g2 | Rola-high | -3.96 | 1.51e-04 | MD05G1345800 | -1.02 | 8.73e-07 | yes | peroxidase 42-like | — |
| TRINITY_DN21409_c0_g1 | Rola-high | -3.96 | 2.19e-03 | MD15G1172300 | -4.28 | 2.70e-06 | yes | phospholipase D alpha 1 | — |
| TRINITY_DN18936_c0_g1 | Rin-high | 3.95 | 6.04e-17 | MD05G1266200 | 3.74 | 3.08e-15 | | berberine bridge enzyme-like 8 | — |
| TRINITY_DN3135_c0_g1 | Rola-high | -3.95 | 1.10e-03 | MD05G1029500 | -3.98 | 6.40e-07 | | uncharacterized protein | — |
| TRINITY_DN3559_c2_g1 | Rola-high | -3.95 | 7.28e-03 | MD11G1176500 | -3.21 | 2.04e-06 | | kinesin-like protein KIN-12E isoform X2 | — |
| TRINITY_DN11846_c0_g1 | Rola-high | -3.95 | 2.39e-04 | MD02G1011100 | -3.89 | 8.49e-08 | yes | class V chitinase | — |
| TRINITY_DN13651_c0_g1 | Rola-high | -3.95 | 9.50e-03 | MD16G1167900 | -3.92 | 5.86e-05 | | condensin-2 complex subunit CAP-D3 | — |

| Trinity gene ID | Direction | Trinity | | GDDH13 protein match | | | Def. | Description | FCS-GX |
|---|---|---|---|---|---|---|---|---|---|
| | | log2FC | padj | gene ID | log2FC | padj | | | call |
| TRINITY_DN46579_c0_g1 | Rola-high | -3.94 | 4.26e-03 | MD10G1175000 | -3.28 | 6.32e-07 | | pleiotropic drug resistance protein 1-like | — |
| TRINITY_DN3984_c0_g1 | Rola-high | -3.94 | 2.57e-04 | MD07G1049400 | -3.25 | 4.25e-06 | | aspartyl protease AED3-like | — |
| TRINITY_DN11227_c0_g1 | Rola-high | -3.94 | 1.27e-03 | MD14G1099800 | -3.96 | 8.19e-07 | | condensin-1 complex subunit CAP-D2 | — |
| TRINITY_DN11592_c4_g1 | Rola-high | -3.93 | 9.25e-05 | MD12G1252500 | -3.73 | 6.83e-09 | yes | GDSL esterase/lipase At5g33370-like | — |
| TRINITY_DN12886_c0_g1 | Rola-high | -3.93 | 5.37e-09 | MD07G1269200 | -3.83 | 5.77e-18 | | (-)-germacrene D synthase-like | — |
| TRINITY_DN3147_c0_g1 | Rola-high | -3.92 | 6.96e-03 | MD12G1076000 | -3.78 | 2.14e-05 | | kinesin-like protein KIN-14L isoform X1 | — |
| TRINITY_DN7110_c2_g1 | Rola-high | -3.91 | 4.28e-03 | MD04G1205400 | -4.10 | 1.23e-05 | yes | putative disease resistance RPP13-like protein 1 isoform X5 | — |
| TRINITY_DN21789_c0_g1 | Rola-high | -3.91 | 7.95e-06 | MD15G1049300 | -2.10 | 2.71e-09 | yes | LRR receptor-like serine/threonine-protein kinase ERECTA isoform X1 | — |
| TRINITY_DN23466_c0_g1 | Rin-high | 3.91 | 1.73e-05 | MD13G1233400 | 3.38 | 4.73e-10 | | FCS-Like Zinc finger 14-like | — |
| TRINITY_DN5864_c0_g1 | Rola-high | -3.90 | 3.71e-05 | MD10G1067800 | -4.00 | 2.48e-10 | | beta-galactosidase 5-like isoform X1 | — |
| TRINITY_DN7067_c0_g2 | Rin-high | 3.90 | 1.26e-04 | MD05G1320800 | 5.43 | 2.54e-05 | yes | polyphenol oxidase latent form, chloroplastic-like | — |
| TRINITY_DN31606_c0_g1 | Rola-high | -3.89 | 1.38e-03 | MD03G1098800 | -1.29 | 3.53e-03 | | probable histone H2B.1 | — |
| TRINITY_DN4975_c0_g1 | Rola-high | -3.89 | 3.33e-07 | MD11G1307100 | -3.81 | 8.29e-15 | yes | endoglucanase 17-like | — |
| TRINITY_DN3552_c0_g1 | Rola-high | -3.88 | 2.10e-06 | MD17G1259400 | -4.18 | 1.59e-14 | yes | cytochrome P450 736A117-like | — |
| TRINITY_DN58161_c0_g1 | Rin-high | 3.87 | 5.97e-03 | MD05G1321000 | 3.90 | 8.42e-07 | | COP1-interacting protein 7-like | — |
| TRINITY_DN30113_c0_g1 | Rola-high | -3.87 | 4.41e-07 | MD12G1202600 | -3.83 | 2.15e-14 | yes | GDSL esterase/lipase At1g09390 | — |
| TRINITY_DN27579_c0_g1 | Rola-high | -3.86 | 2.45e-06 | MD05G1088500 | -3.99 | 5.33e-13 | yes | receptor protein kinase-like protein At4g34220 | — |
| TRINITY_DN22585_c0_g1 | Rola-high | -3.86 | 4.49e-03 | MD08G1198000 | -4.00 | 1.75e-06 | | kinesin-like protein KIN-4C isoform X6 | — |
| TRINITY_DN16538_c0_g1 | Rola-high | -3.86 | 9.36e-05 | MD02G1087700 | -3.61 | 1.25e-08 | | transcription factor MYB1-like | — |
| TRINITY_DN8751_c0_g1 | Rola-high | -3.86 | 3.27e-04 | MD13G1123800 | -3.09 | 6.91e-07 | | hypothetical protein DVH24_041080 | — |
| TRINITY_DN50035_c0_g3 | Rin-high | 3.86 | 8.87e-03 | MD09G1051800 | 4.02 | 5.45e-05 | | ATP-dependent zinc metalloprotease FTSH 2, chloroplastic | — |
| TRINITY_DN38328_c0_g1 | Rola-high | -3.85 | 9.19e-04 | MD12G1000200 | -3.84 | 4.74e-07 | | syntaxin-related protein KNOLLE | — |
| TRINITY_DN1430_c2_g2 | Rola-high | -3.84 | 5.76e-03 | MD02G1132500 | -3.90 | 1.72e-07 | | hypothetical protein DVH24_018508 | — |

| Trinity gene ID | Direction | Trinity | | GDDH13 protein match | | | Def. | Description | FCS-GX |
|---|---|---|---|---|---|---|---|---|---|
| | | log2FC | padj | gene ID | log2FC | padj | | | call |
| TRINITY_DN17077_c0_g1 | Rin-high | 3.83 | 2.33e-03 | MD07G1096700 | 3.95 | 2.58e-06 | | uncharacterized protein At1g28695-like | — |
| TRINITY_DN1119_c2_g1 | Rin-high | 3.83 | 3.63e-03 | MD15G1139800 | 3.91 | 1.13e-05 | | ammonium transporter 2-like | — |
| TRINITY_DN923_c1_g1 | Rola-high | -3.83 | 1.48e-05 | MD05G1116900 | -3.87 | 1.98e-10 | | pollen-specific protein-like At4g18596 | — |
| TRINITY_DN35021_c0_g1 | Rin-high | 3.83 | 2.13e-03 | MD01G1172500 | 3.24 | 2.33e-09 | yes | cytochrome P450 714A1-like | — |
| TRINITY_DN26697_c0_g1 | Rola-high | -3.83 | 7.59e-05 | MD06G1065700 | -3.78 | 7.44e-11 | | axial regulator YABBY 1-like | — |
| TRINITY_DN5448_c2_g1 | Rola-high | -3.83 | 1.18e-08 | MD07G1269200 | -3.83 | 5.77e-18 | | (-)-germacrene D synthase-like | — |
| TRINITY_DN10411_c0_g1 | Rola-high | -3.82 | 8.51e-03 | MD06G1225500 | -3.60 | 3.55e-06 | | replication protein A 70 kDa DNA-binding subunit B-like | — |
| TRINITY_DN96695_c0_g2 | Rola-high | -3.82 | 5.40e-04 | MD00G1139900 | -4.69 | 9.94e-17 | yes | probable lysophospholipase BODYGUARD 3 isoform X3 | — |
| TRINITY_DN9388_c0_g1 | Rola-high | -3.82 | 1.06e-03 | MD02G1265300 | -3.25 | 5.11e-10 | yes | ethylene-responsive transcription factor WRI1-like | — |
| TRINITY_DN9850_c0_g1 | Rola-high | -3.82 | 2.27e-03 | MD03G1106600 | -6.15 | 2.22e-07 | yes | laccase-17-like | — |
| TRINITY_DN71466_c0_g1 | Rola-high | -3.82 | 6.37e-05 | MD15G1232200 | -3.82 | 1.11e-09 | | ankyrin repeat-containing protein At5g02620-like | — |
| TRINITY_DN9343_c0_g1 | Rin-high | 3.82 | 2.64e-03 | MD04G1029600 | 5.22 | 3.91e-08 | yes | probable pectinesterase/pectinesterase inhibitor 35 | — |
| TRINITY_DN11475_c0_g1 | Rin-high | 3.80 | 1.21e-03 | MD05G1231100 | 3.10 | 1.63e-06 | yes | G-type lectin S-receptor-like serine/threonine-protein kinase At4g27290 isoform X1 | — |
| TRINITY_DN7460_c0_g2 | Rola-high | -3.80 | 1.56e-04 | MD17G1154500 | -3.77 | 3.39e-10 | yes | pectin methylesterase | — |
| TRINITY_DN15406_c0_g1 | Rola-high | -3.79 | 3.03e-06 | MD11G1046300 | -3.77 | 8.59e-13 | | hypothetical protein DVH24_041817 | — |
| TRINITY_DN6123_c2_g1 | Rin-high | 3.79 | 1.43e-03 | MD12G1083400 | 3.85 | 2.63e-07 | | uncharacterized protein LOC126593932 | — |
| TRINITY_DN21855_c0_g1 | Rola-high | -3.79 | 7.27e-03 | MD00G1153400 | -3.58 | 4.23e-03 | | melianol synthase CYP71BQ5-like | — |
| TRINITY_DN179627_c0_g1 | Rola-high | -3.79 | 4.45e-03 | MD07G1068200 | -3.72 | 9.55e-06 | | zinc finger protein BRUTUS-like isoform X1 | — |
| TRINITY_DN13561_c0_g1 | Rin-high | 3.78 | 1.12e-04 | MD05G1266200 | 3.74 | 3.08e-15 | | berberine bridge enzyme-like 8 | — |
| TRINITY_DN21712_c0_g1 | Rola-high | -3.78 | 9.00e-03 | MD12G1215100 | -3.36 | 4.79e-08 | | formin-like protein 5 | — |
| TRINITY_DN15514_c0_g2 | Rin-high | 3.78 | 1.10e-03 | MD07G1275100 | 3.69 | 2.50e-09 | | silicon efflux transporter LSI2-like | — |
| TRINITY_DN12192_c0_g1 | Rola-high | -3.78 | 2.53e-03 | MD14G1063100 | -4.27 | 3.61e-06 | | protein POLYCHOME-like | — |

| Trinity gene ID | Direction | Trinity | | GDDH13 protein match | | | Def. | Description | FCS-GX |
|---|---|---|---|---|---|---|---|---|---|
| | | log2FC | padj | gene ID | log2FC | padj | | | call |
| TRINITY_DN2878_c2_g1 | Rola-high | -3.77 | 3.52e-03 | MD08G1010800 | -2.93 | 1.86e-06 | | kinesin-like protein KIN-4C isoform X1 | — |
| TRINITY_DN144725_c0_g1 | Rola-high | -3.77 | 7.44e-03 | MD09G1261500 | -3.84 | 8.60e-06 | | uncharacterized protein | — |
| TRINITY_DN5163_c0_g1 | Rola-high | -3.77 | 1.53e-03 | MD13G1152800 | -3.95 | 3.84e-07 | yes | peroxidase 5-like | — |
| TRINITY_DN10016_c0_g1 | Rola-high | -3.75 | 2.49e-04 | MD05G1118600 | -3.25 | 3.29e-09 | | auxin transporter-like protein 2 | — |
| TRINITY_DN33088_c1_g1 | Rola-high | -3.75 | 2.64e-03 | MD09G1049500 | -4.00 | 5.66e-07 | | tubulin alpha chain isoform X1 | — |
| TRINITY_DN29106_c0_g1 | Rola-high | -3.75 | 9.19e-03 | MD15G1350900 | -4.44 | 1.55e-06 | | universal stress protein PHOS32-like | — |
| TRINITY_DN34978_c0_g1 | Rola-high | -3.74 | 4.73e-03 | MD17G1236100 | -3.57 | 4.53e-06 | yes | mitotic checkpoint serine/threonine-protein kinase BUB1 | — |
| TRINITY_DN18864_c0_g1 | Rin-high | 3.74 | 4.52e-03 | MD04G1182200 | 3.62 | 3.10e-06 | yes | leucine-rich repeat receptor protein kinase HPCA1-like | — |
| TRINITY_DN43671_c0_g3 | Rin-high | 3.74 | 2.09e-05 | MD07G1236500 | 6.06 | 1.04e-18 | yes | probable LRR receptor-like serine/threonine-protein kinase At3g47570 isoform X1 | — |
| TRINITY_DN118714_c0_g1 | Rola-high | -3.73 | 3.54e-06 | MD13G1265800 | -3.77 | 1.33e-12 | yes | pathogenesis-related protein 1-like | — |
| TRINITY_DN52317_c0_g1 | Rin-high | 3.73 | 1.61e-04 | MD12G1026200 | 4.54 | 1.53e-10 | | putative polyol transporter 1 | — |
| TRINITY_DN5371_c0_g2 | Rola-high | -3.72 | 9.48e-04 | MD02G1308000 | -4.14 | 9.11e-08 | | DNA topoisomerase 2 isoform X2 | — |
| TRINITY_DN65481_c0_g1 | Rola-high | -3.70 | 5.11e-03 | MD13G1132300 | -6.62 | 3.14e-05 | | indole-3-acetic acid-amido synthetase GH3.17-like | — |
| TRINITY_DN20319_c0_g1 | Rola-high | -3.70 | 2.34e-03 | MD16G1098400 | -3.77 | 8.19e-07 | | L-tryptophan--pyruvate aminotransferase 1-like | — |
| TRINITY_DN1140_c0_g1 | Rola-high | -3.69 | 1.94e-03 | MD17G1019000 | -3.65 | 4.77e-06 | | protein WVD2-like 7 isoform X1 | — |
| TRINITY_DN10096_c0_g1 | Rola-high | -3.69 | 4.73e-03 | MD15G1121200 | -4.06 | 1.31e-05 | | CRC domain-containing protein TSO1-like isoform X2 | — |
| TRINITY_DN3168_c1_g2 | Rola-high | -3.68 | 2.15e-04 | MD04G1148900 | -2.99 | 3.73e-27 | | glycerol-3-phosphate acyltransferase RAM2-like | — |
| TRINITY_DN57297_c0_g1 | Rin-high | 3.68 | 6.12e-06 | MD15G1028700 | 2.40 | 8.60e-04 | | hypothetical protein DVH24_012880 | — |
| TRINITY_DN6251_c0_g1 | Rola-high | -3.68 | 2.61e-03 | MD02G1132500 | -3.90 | 1.72e-07 | | hypothetical protein DVH24_018508 | — |
| TRINITY_DN54761_c0_g1 | Rin-high | 3.68 | 2.58e-03 | MD07G1206400 | 2.86 | 2.53e-07 | | cucumisin isoform X3 | — |
| TRINITY_DN5441_c0_g1 | Rola-high | -3.67 | 1.20e-03 | MD15G1385300 | -3.48 | 5.29e-07 | | kinesin-like protein KIN-4C isoform X1 | — |

| Trinity gene ID | Direction | Trinity | | GDDH13 protein match | | | Def. | Description | FCS-GX |
| --- | --- | --- | --- | --- | --- | --- | --- | --- | --- |
| | | log2FC | padj | gene ID | log2FC | padj | | | call |
| TRINITY_DN10831_c0_g1 | Rin-high | 3.66 | 5.16e-03 | MD08G1153100 | 3.81 | 4.92e-05 | | flotillin-like protein 4 | — |
| TRINITY_DN15429_c0_g1 | Rola-high | -3.66 | 1.58e-03 | MD15G1288300 | -3.02 | 2.69e-03 | | uncharacterized protein | — |
| TRINITY_DN3340_c1_g1 | Rola-high | -3.65 | 2.99e-03 | MD12G1261500 | -3.71 | 1.69e-06 | | uncharacterized protein | — |
| TRINITY_DN8128_c2_g1 | Rola-high | -3.64 | 2.16e-05 | MD15G1292900 | -3.58 | 6.78e-10 | | subtilisin-like protease SBT1.7 | — |
| TRINITY_DN24568_c0_g1 | Rola-high | -3.64 | 4.58e-03 | MD14G1084000 | -3.64 | 7.19e-06 | | uncharacterized protein | — |
| TRINITY_DN3467_c1_g2 | Rola-high | -3.63 | 8.36e-03 | MD10G1046600 | -3.49 | 8.77e-06 | | FHA domain-containing protein PS1-like | — |
| TRINITY_DN15307_c0_g1 | Rola-high | -3.63 | 7.41e-03 | MD04G1166400 | -3.32 | 1.03e-06 | yes | probably inactive leucine-rich repeat receptor-like protein kinase IMK2 | — |
| TRINITY_DN2607_c0_g1 | Rola-high | -3.63 | 5.13e-06 | MD05G1243100 | -4.63 | 2.24e-10 | | uncharacterized protein | — |
| TRINITY_DN30908_c0_g1 | Rola-high | -3.62 | 2.01e-03 | MD03G1052400 | -5.01 | 9.06e-11 | yes | expansin-A4-like precursor | — |
| TRINITY_DN8241_c0_g2 | Rola-high | -3.62 | 2.48e-03 | MD02G1043600 | -3.69 | 7.57e-07 | | probable protein ABIL5 | — |
| TRINITY_DN297_c0_g2 | Rola-high | -3.62 | 3.99e-03 | MD11G1298300 | -2.57 | 7.66e-06 | | classical arabinogalactan protein 26-like | — |
| TRINITY_DN14400_c0_g1 | Rola-high | -3.62 | 3.24e-03 | MD05G1047500 | -3.57 | 5.18e-06 | | uncharacterized protein At4g38062-like | — |
| TRINITY_DN21027_c0_g1 | Rola-high | -3.62 | 8.00e-03 | MD07G1288100 | -3.67 | 1.12e-05 | | protein DEFECTIVE IN MERISTEM SILENCING 3-like | — |
| TRINITY_DN10715_c0_g1 | Rola-high | -3.61 | 2.67e-03 | MD09G1085300 | -3.70 | 1.08e-06 | | uncharacterized protein | — |
| TRINITY_DN8825_c0_g1 | Rin-high | 3.61 | 1.35e-03 | MD15G1282600 | 3.39 | 7.81e-04 | yes | calmodulin-like protein 30 | — |
| TRINITY_DN52724_c0_g1 | Rola-high | -3.61 | 3.15e-03 | MD17G1155800 | -3.19 | 2.54e-06 | | ATPase 10, plasma membrane-type-like | — |
| TRINITY_DN23286_c0_g1 | Rola-high | -3.61 | 6.65e-03 | MD08G1202900 | -2.94 | 2.95e-06 | | kinesin-like protein KIN-14R | — |
| TRINITY_DN10111_c0_g1 | Rola-high | -3.61 | 7.17e-04 | MD09G1083800 | -3.25 | 1.72e-04 | | small polypeptide DEVIL 4-like | — |
| TRINITY_DN56816_c0_g1 | Rola-high | -3.60 | 4.84e-03 | MD09G1201800 | -3.87 | 2.83e-07 | | uncharacterized protein | — |
| TRINITY_DN6981_c1_g1 | Rin-high | 3.60 | 1.51e-04 | MD05G1065500 | 3.66 | 2.50e-07 | | protein RADIALIS-like 3 | — |
| TRINITY_DN28611_c0_g1 | Rola-high | -3.60 | 1.43e-03 | MD10G1254200 | -1.22 | 2.39e-07 | | 187-kDa microtubule-associated protein AIR9-like | — |
| TRINITY_DN12191_c0_g1 | Rola-high | -3.60 | 6.59e-03 | MD11G1022200 | -4.21 | 3.45e-05 | | UDP-glycosyltransferase 92A1-like | — |

| Trinity gene ID | Direction | Trinity | | GDDH13 protein match | | | Def. | Description | FCS-GX |
|---|---|---|---|---|---|---|---|---|---|
| | | log2FC | padj | gene ID | log2FC | padj | | | call |
| TRINITY_DN43772_c0_g1 | Rola-high | -3.59 | 7.28e-03 | MD10G1329000 | -3.59 | 1.70e-06 | | EH domain-containing protein 2-like | — |
| TRINITY_DN9342_c0_g2 | Rola-high | -3.59 | 3.54e-06 | MD02G1087900 | -3.36 | 1.68e-12 | | transcription factor MYB1-like | — |
| TRINITY_DN27473_c0_g1 | Rola-high | -3.59 | 3.35e-03 | MD17G1155800 | -3.19 | 2.54e-06 | | ATPase 10, plasma membrane-type-like | — |
| TRINITY_DN49561_c0_g1 | Rin-high | 3.58 | 1.67e-03 | MD00G1046700 | 2.37 | 3.20e-10 | yes | probable LRR receptor-like serine/threonine-protein kinase At1g56130 | — |
| TRINITY_DN34730_c0_g1 | Rola-high | -3.58 | 5.11e-03 | MD11G1206000 | -3.10 | 3.10e-05 | | kinesin-like protein KIN-12B | — |
| TRINITY_DN5156_c0_g2 | Rola-high | -3.57 | 5.46e-03 | MD13G1099500 | -3.52 | 1.45e-05 | | probable protein phosphatase 2C 25 | — |
| TRINITY_DN8030_c1_g1 | Rola-high | -3.57 | 6.14e-04 | MD15G1300900 | -2.48 | 4.58e-07 | | growth-regulating factor 1 isoform X3 | — |
| TRINITY_DN6313_c0_g1 | Rola-high | -3.57 | 2.55e-03 | MD10G1311800 | -3.75 | 9.33e-07 | | kinesin-like protein KIN-14C | — |
| TRINITY_DN4592_c0_g1 | Rin-high | 3.57 | 6.76e-03 | MD17G1092400 | 3.71 | 2.38e-05 | yes | peroxidase 3-like | — |
| TRINITY_DN19658_c0_g3 | Rola-high | -3.57 | 1.45e-04 | MD02G1108700 | -3.51 | 1.11e-08 | | 3-hydroxy-3-methylglutaryl coenzyme A reductase 2-B-like | — |
| TRINITY_DN107038_c0_g1 | Rola-high | -3.56 | 7.09e-05 | MD17G1246800 | -3.85 | 2.85e-10 | | tryptophan synthase alpha chain-like | — |
| TRINITY_DN20648_c0_g1 | Rola-high | -3.56 | 9.44e-03 | MD05G1223200 | -4.03 | 8.73e-06 | | uncharacterized protein | — |
| TRINITY_DN45705_c0_g1 | Rin-high | 3.56 | 9.91e-03 | MD09G1144400 | 3.51 | 2.73e-05 | | F-box/kelch-repeat protein At3g06240-like isoform X2 | — |
| TRINITY_DN2493_c0_g1 | Rola-high | -3.55 | 7.84e-03 | MD09G1017800 | -3.61 | 2.12e-05 | | glycosyltransferase family 92 protein RCOM_0530710 | — |
| TRINITY_DN3562_c2_g1 | Rola-high | -3.55 | 2.45e-04 | MD03G1069000 | -3.66 | 2.01e-09 | yes | phosphoinositide phospholipase C 4-like | — |
| TRINITY_DN4376_c0_g1 | Rola-high | -3.54 | 1.38e-06 | MD07G1054100 | -3.74 | 5.93e-14 | yes | ethylene-responsive transcription factor WRI1-like | — |
| TRINITY_DN44513_c0_g1 | Rola-high | -3.54 | 3.45e-04 | MD17G1164400 | -3.96 | 4.51e-04 | | jasmonate ZIM domain-containing protein 1-like | — |
| TRINITY_DN41773_c0_g1 | Rola-high | -3.54 | 1.59e-03 | MD15G1021100 | -3.29 | 7.80e-16 | | beta-galactosidase 5 | — |
| TRINITY_DN5370_c0_g1 | Rola-high | -3.54 | 1.63e-03 | MD09G1201700 | -3.18 | 1.34e-06 | | uncharacterized protein | — |
| TRINITY_DN17416_c0_g1 | Rola-high | -3.53 | 8.68e-04 | MD04G1119500 | -3.26 | 1.97e-07 | | kinesin-like protein KIN-5B | — |
| TRINITY_DN7242_c0_g1 | Rola-high | -3.53 | 5.36e-03 | MD13G1130100 | -3.76 | 1.91e-06 | | histone-lysine N-methyltransferase, H3 lysine-9 specific SUVH4-like isoform X2 | — |

| Trinity gene ID | Direction | Trinity | | GDDH13 protein match | | | Def. | Description | FCS-GX |
|---|---|---|---|---|---|---|---|---|---|
| | | log2FC | padj | gene ID | log2FC | padj | | | call |
| TRINITY_DN1572_c1_g1 | Rola-high | -3.53 | 3.88e-10 | MD11G1161800 | -3.22 | 3.35e-27 | | fatty acyl-CoA reductase 3-like | — |
| TRINITY_DN11436_c0_g1 | Rola-high | -3.52 | 5.61e-03 | MD09G1270700 | -3.56 | 5.99e-06 | | hypothetical protein DVH24_035390 | — |
| TRINITY_DN16856_c0_g2 | Rin-high | 3.52 | 5.77e-04 | MD15G1099500 | 3.26 | 2.90e-07 | | group 2 truncated hemoglobin 3-2 | — |
| TRINITY_DN3558_c0_g1 | Rin-high | 3.52 | 2.24e-03 | MD16G1054800 | 3.66 | 1.18e-07 | | uncharacterized protein | — |
| TRINITY_DN51837_c0_g1 | Rola-high | -3.51 | 8.28e-03 | MD07G1117200 | -2.51 | 2.91e-09 | | axial regulator YABBY 1-like isoform X1 | — |
| TRINITY_DN27278_c0_g1 | Rola-high | -3.51 | 4.48e-03 | MD02G1196800 | -3.56 | 2.28e-06 | | subtilisin-like protease SBT1.7 | — |
| TRINITY_DN12409_c0_g1 | Rola-high | -3.51 | 1.46e-03 | MD06G1138000 | -2.99 | 4.11e-08 | | HVA22-like protein e isoform X1 | — |
| TRINITY_DN3424_c0_g1 | Rola-high | -3.50 | 1.80e-05 | MD16G1021500 | -3.47 | 1.68e-10 | yes | endoglucanase 17-like | — |
| TRINITY_DN32227_c0_g1 | Rola-high | -3.50 | 6.98e-03 | MD15G1121500 | -3.43 | 1.50e-06 | | RNA-binding protein involved in heterochromatin assembly dri1-like | — |
| TRINITY_DN4576_c0_g1 | Rola-high | -3.49 | 8.29e-10 | MD10G1195100 | -3.07 | 7.23e-20 | | interactor of constitutive active ROPs 4-like | — |
| TRINITY_DN9371_c0_g2 | Rola-high | -3.49 | 6.19e-03 | MD01G1124600 | -4.58 | 3.08e-07 | | histone H4 | — |
| TRINITY_DN12260_c0_g1 | Rola-high | -3.49 | 8.06e-05 | MD10G1286100 | -3.50 | 3.02e-10 | yes | GDSL esterase/lipase At5g45950 | — |
| TRINITY_DN3663_c0_g1 | Rola-high | -3.48 | 1.23e-03 | MD13G1069000 | -4.09 | 6.24e-07 | | methyltransferase FGSG_00040 | — |
| TRINITY_DN7737_c1_g1 | Rola-high | -3.48 | 2.41e-05 | MD06G1198900 | -2.88 | 2.17e-12 | | hypothetical protein DVH24_008911 | — |
| TRINITY_DN60615_c0_g1 | Rin-high | 3.47 | 6.43e-04 | MD05G1065500 | 3.66 | 2.50e-07 | | protein RADIALIS-like 4 | — |
| TRINITY_DN27864_c0_g2 | Rin-high | 3.47 | 5.51e-04 | MD13G1233400 | 3.38 | 4.73e-10 | | FCS-Like Zinc finger 14-like | — |
| TRINITY_DN102542_c0_g1 | Rola-high | -3.46 | 1.45e-03 | MD10G1118600 | -3.37 | 4.29e-07 | | proline-rich protein 4-like | — |
| TRINITY_DN14675_c0_g1 | Rin-high | 3.46 | 6.84e-03 | MD16G1199500 | 2.39 | 2.35e-05 | | acireductone dioxygenase 2 | — |
| TRINITY_DN47521_c0_g4 | Rola-high | -3.46 | 5.97e-03 | MD14G1085100 | -3.35 | 2.70e-06 | | kinesin-like protein KIN-7O | — |
| TRINITY_DN29755_c0_g1 | Rola-high | -3.46 | 6.58e-05 | MD11G1178000 | -2.51 | 1.40e-12 | | uncharacterized protein | — |
| TRINITY_DN8030_c0_g1 | Rola-high | -3.44 | 6.48e-05 | MD16G1158800 | -5.07 | 2.38e-10 | | transcription factor MYB111 | — |
| TRINITY_DN3559_c0_g1 | Rola-high | -3.44 | 3.07e-03 | MD03G1160900 | -3.68 | 2.62e-06 | | kinesin-like protein KIN-12E | — |
| TRINITY_DN4311_c0_g1 | Rola-high | -3.44 | 3.79e-03 | MD05G1191900 | -3.63 | 4.66e-06 | | protein JASON-like | — |

| Trinity gene ID | Direction | Trinity | | GDDH13 protein match | | | Def. | Description | FCS-GX |
| --- | --- | --- | --- | --- | --- | --- | --- | --- | --- |
| | | log2FC | padj | gene ID | log2FC | padj | | | call |
| TRINITY_DN1597_c0_g1 | Rin-high | 3.44 | 1.00e-05 | MD01G1055200 | 3.05 | 1.69e-10 | | uncharacterized protein | — |
| TRINITY_DN38814_c0_g1 | Rola-high | -3.43 | 2.42e-04 | MD13G1094100 | -3.30 | 1.94e-07 | | polygalacturonase At1g48100 | — |
| TRINITY_DN9290_c0_g1 | Rola-high | -3.43 | 1.37e-03 | MD04G1129000 | -3.92 | 2.36e-07 | | kinesin-like protein KIN-10B isoform X1 | — |
| TRINITY_DN2719_c2_g1 | Rola-high | -3.42 | 2.69e-07 | MD01G1143600 | -2.10 | 9.29e-12 | | UDP-glycosyltransferase 76F1-like | — |
| TRINITY_DN38533_c1_g1 | Rin-high | 3.41 | 2.53e-04 | MD00G1023600 | 3.48 | 2.08e-07 | yes | probable LRR receptor-like serine/threonine-protein kinase At1g56130 | — |
| TRINITY_DN58970_c0_g1 | Rola-high | -3.40 | 3.46e-03 | MD14G1119800 | -3.24 | 3.22e-10 | yes | probable inactive receptor kinase At4g23740 | — |
| TRINITY_DN41655_c0_g1 | Rola-high | -3.39 | 8.28e-03 | MD12G1140400 | -3.05 | 2.43e-04 | | uncharacterized protein | — |
| TRINITY_DN3276_c0_g1 | Rola-high | -3.39 | 7.60e-03 | MD14G1212400 | -3.44 | 3.84e-06 | | replication protein A 70 kDa DNA-binding subunit B-like | — |
| TRINITY_DN59200_c0_g1 | Rin-high | 3.38 | 1.33e-04 | MD11G1160200 | 3.34 | 6.42e-09 | yes | MLP-like protein 423 | — |
| TRINITY_DN7490_c0_g1 | Rola-high | -3.38 | 3.83e-03 | MD15G1233100 | -2.97 | 1.98e-05 | | G2/mitotic-specific cyclin-2-like | — |
| TRINITY_DN3549_c0_g1 | Rin-high | 3.38 | 9.25e-03 | MD09G1285700 | 3.39 | 8.93e-06 | | uncharacterized protein | — |
| TRINITY_DN10798_c2_g1 | Rola-high | -3.38 | 4.29e-05 | MD05G1160900 | -3.16 | 8.58e-08 | | uncharacterized protein | — |
| TRINITY_DN185563_c0_g1 | Rola-high | -3.37 | 1.32e-04 | MD05G1113600 | -3.47 | 2.52e-10 | | auxin-induced protein 15A-like | — |
| TRINITY_DN26153_c0_g1 | Rola-high | -3.37 | 2.26e-03 | MD10G1118600 | -3.37 | 4.29e-07 | | proline-rich protein 4-like | — |
| TRINITY_DN1478_c1_g1 | Rola-high | -3.36 | 3.57e-03 | MD12G1108300 | -3.11 | 9.35e-06 | | probable pectate lyase 12 | — |
| TRINITY_DN6512_c0_g1 | Rola-high | -3.36 | 1.24e-03 | MD13G1216300 | -3.71 | 5.20e-07 | | kinesin-like protein KIN-5B | — |
| TRINITY_DN528_c0_g1 | Rola-high | -3.35 | 9.40e-04 | MD02G1062300 | -3.37 | 9.11e-03 | | small polypeptide DEVIL 10-like | — |
| TRINITY_DN7910_c0_g1 | Rola-high | -3.35 | 6.43e-05 | MD16G1184300 | -2.84 | 7.00e-10 | | heavy metal-associated isoprenylated plant protein 34-like | — |
| TRINITY_DN7009_c0_g2 | Rola-high | -3.34 | 1.21e-03 | MD10G1152100 | -3.28 | 2.90e-07 | | uncharacterized protein At5g48480 | — |
| TRINITY_DN24496_c0_g1 | Rin-high | 3.34 | 6.07e-03 | MD08G1107900 | 1.69 | 5.61e-04 | yes | wall-associated receptor kinase-like 9 | — |
| TRINITY_DN7227_c0_g1 | Rola-high | -3.33 | 2.01e-07 | MD12G1240300 | -3.21 | 1.06e-15 | | perakine reductase-like | — |
| TRINITY_DN153_c2_g2 | Rola-high | -3.33 | 4.45e-03 | MD14G1146900 | -3.32 | 2.21e-06 | | E3 ubiquitin-protein ligase ORTHRUS 2 | — |

| Trinity gene ID | Direction | Trinity | | GDDH13 protein match | | | Def. | Description | FCS-GX |
|---|---|---|---|---|---|---|---|---|---|
| | | log2FC | padj | gene ID | log2FC | padj | | | call |
| TRINITY_DN18350_c0_g1 | Rola-high | -3.33 | 7.83e-03 | MD16G1208400 | -3.44 | 5.35e-06 | | transcription factor bHLH92-like | — |
| TRINITY_DN4234_c0_g4 | Rola-high | -3.32 | 2.71e-03 | MD01G1158800 | -3.47 | 4.29e-08 | | cation/H(+) antiporter 15-like | — |
| TRINITY_DN3453_c0_g1 | Rin-high | 3.32 | 1.26e-03 | MD10G1338600 | 3.44 | 2.10e-07 | | uncharacterized protein | — |
| TRINITY_DN16492_c0_g1 | Rola-high | -3.32 | 2.53e-05 | MD05G1128000 | -3.41 | 5.18e-12 | | acetylserotonin O-methyltransferase-like | — |
| TRINITY_DN4164_c1_g1 | Rola-high | -3.31 | 4.17e-03 | MD03G1022800 | -2.32 | 1.08e-04 | | G2/mitotic-specific cyclin S13-7-like | — |
| TRINITY_DN54903_c0_g1 | Rola-high | -3.31 | 3.95e-03 | MD12G1040200 | -2.71 | 3.59e-13 | yes | receptor protein kinase TMK1-like | — |
| TRINITY_DN29867_c0_g1 | Rola-high | -3.30 | 2.40e-06 | MD11G1156200 | -3.35 | 2.68e-17 | | glucomannan 4-beta-mannosyltransferase 9-like | — |
| TRINITY_DN13164_c0_g1 | Rin-high | 3.30 | 2.04e-03 | MD01G1158900 | 4.96 | 2.60e-10 | yes | laccase-12-like | — |
| TRINITY_DN68970_c0_g1 | Rola-high | -3.30 | 1.93e-03 | MD08G1024300 | -2.72 | 1.31e-09 | | BEL1-like homeodomain protein 11 | — |
| TRINITY_DN17795_c0_g1 | Rola-high | -3.30 | 1.57e-04 | MD02G1067000 | -3.36 | 1.42e-11 | yes | probable pectinesterase 53 | — |
| TRINITY_DN13236_c3_g1 | Rola-high | -3.30 | 8.89e-03 | MD01G1124600 | -4.58 | 3.08e-07 | | histone H4 | — |
| TRINITY_DN9716_c0_g1 | Rin-high | 3.29 | 9.14e-05 | MD04G1100000 | 3.33 | 2.60e-10 | | fasciclin-like arabinogalactan protein 12 | — |
| TRINITY_DN2645_c0_g1 | Rola-high | -3.29 | 1.26e-04 | MD06G1073500 | -3.35 | 3.60e-07 | | scarecrow-like protein 28 | — |
| TRINITY_DN12844_c0_g1 | Rin-high | 3.29 | 6.55e-03 | MD17G1153400 | 4.36 | 4.31e-10 | | FCS-Like Zinc finger 17-like | — |
| TRINITY_DN1310_c0_g1 | Rola-high | -3.29 | 1.66e-03 | MD17G1049700 | -3.62 | 2.76e-07 | | tubulin alpha chain isoform X1 | — |
| TRINITY_DN21248_c2_g1 | Rola-high | -3.28 | 9.90e-03 | MD14G1188500 | -3.30 | 1.04e-05 | | formin-like protein 4 | — |
| TRINITY_DN13918_c1_g1 | Rola-high | -3.28 | 2.58e-03 | MD04G1066800 | -3.22 | 8.75e-10 | | scarecrow-like protein 28 | — |
| TRINITY_DN11412_c1_g1 | Rola-high | -3.27 | 1.49e-03 | MD13G1094100 | -3.30 | 1.94e-07 | | polygalacturonase At1g48100 | — |
| TRINITY_DN6546_c0_g1 | Rola-high | -3.27 | 7.73e-03 | MD01G1182300 | -3.61 | 2.97e-06 | | structural maintenance of chromosomes protein 3-like isoform X1 | — |
| TRINITY_DN10047_c0_g1 | Rin-high | 3.27 | 3.85e-04 | MD02G1124700 | 3.93 | 1.25e-07 | yes | peroxidase 64 | — |
| TRINITY_DN2243_c0_g1 | Rola-high | -3.26 | 4.54e-03 | MD08G1202900 | -2.94 | 2.95e-06 | | kinesin-like protein KIN-14R | — |
| TRINITY_DN2310_c0_g1 | Rola-high | -3.26 | 2.28e-03 | MD12G1093900 | -3.33 | 1.91e-07 | | G2/mitotic-specific cyclin S13-7-like isoform X1 | — |
| TRINITY_DN10984_c0_g1 | Rola-high | -3.26 | 4.27e-09 | MD05G1128600 | -3.24 | 9.70e-23 | yes | protein kinase STUNTED isoform X2 | — |

| Trinity gene ID | Direction | Trinity | | GDDH13 protein match | | | Def. | Description | FCS-GX |
|---|---|---|---|---|---|---|---|---|---|
| | | log2FC | padj | gene ID | log2FC | padj | | | call |
| TRINITY_DN45351_c0_g1 | Rin-high | 3.25 | 8.62e-04 | MD04G1183100 | 2.49 | 2.66e-05 | yes | leucine-rich repeat receptor protein kinase HPCA1-like | — |
| TRINITY_DN29977_c0_g1 | Rin-high | 3.25 | 8.04e-03 | MD14G1066000 | 2.93 | 2.76e-06 | | carboxyl-terminal-processing peptidase 3, chloroplastic-like isoform X2 | — |
| TRINITY_DN389_c0_g1 | Rola-high | -3.25 | 5.87e-03 | MD07G1184000 | -2.68 | 8.98e-05 | | cyclic nucleotide-gated ion channel 1-like isoform X2 | — |
| TRINITY_DN12475_c0_g1 | Rin-high | 3.25 | 2.33e-03 | MD15G1239200 | 2.15 | 7.29e-03 | yes | probable LRR receptor-like serine/threonine-protein kinase At1g56140 | — |
| TRINITY_DN10570_c0_g2 | Rola-high | -3.24 | 9.79e-03 | MD05G1260300 | -3.02 | 2.78e-05 | | uncharacterized protein | — |
| TRINITY_DN4620_c0_g1 | Rin-high | 3.24 | 3.48e-05 | MD16G1158300 | 3.94 | 8.20e-11 | | probable beta-D-xylosidase 2 | — |
| TRINITY_DN20164_c0_g1 | Rin-high | 3.24 | 7.45e-03 | MD13G1275500 | 2.90 | 1.01e-04 | | gamma-glutamyl peptidase 5-like | — |
| TRINITY_DN11989_c1_g1 | Rin-high | 3.23 | 5.00e-03 | MD03G1006600 | 3.46 | 2.63e-06 | | probable polyamine transporter At3g13620 | — |
| TRINITY_DN24827_c0_g1 | Rin-high | 3.23 | 2.25e-03 | MD10G1186000 | 3.25 | 5.32e-07 | | WAT1-related protein At4g08300 | — |
| TRINITY_DN50368_c0_g4 | Rin-high | 3.23 | 1.70e-03 | MD17G1126300 | 3.26 | 2.46e-06 | | 7-deoxyloganetin glucosyltransferase-like | — |
| TRINITY_DN55460_c0_g1 | Rin-high | 3.23 | 2.89e-04 | MD11G1314000 | 3.15 | 1.88e-10 | | calvin cycle protein CP12-1, chloroplastic-like | — |
| TRINITY_DN12415_c0_g1 | Rin-high | 3.23 | 1.42e-03 | MD11G1315600 | 9.84 | 4.14e-13 | | universal stress protein A-like protein | — |
| TRINITY_DN10626_c1_g1 | Rola-high | -3.23 | 3.24e-03 | MD04G1147100 | -2.43 | 1.82e-06 | | protein OCTOPUS | — |
| TRINITY_DN10259_c0_g1 | Rola-high | -3.22 | 4.41e-03 | MD13G1232000 | -3.19 | 1.47e-06 | | TORTIFOLIA1-like protein 2 | — |
| TRINITY_DN9274_c0_g1 | Rin-high | 3.22 | 5.97e-03 | MD10G1025300 | 2.59 | 8.71e-08 | | WAT1-related protein At1g43650-like | — |
| TRINITY_DN7009_c0_g1 | Rola-high | -3.21 | 1.62e-04 | MD05G1159600 | -3.10 | 4.59e-09 | | uncharacterized protein At5g48480 | — |
| TRINITY_DN9135_c1_g1 | Rin-high | 3.21 | 1.64e-03 | MD07G1081200 | 4.85 | 5.99e-07 | | hypothetical protein DVH24_013782 | — |
| TRINITY_DN25634_c1_g1 | Rola-high | -3.20 | 4.54e-03 | MD15G1389100 | -3.14 | 1.15e-07 | | uncharacterized protein | — |
| TRINITY_DN26275_c0_g1 | Rin-high | 3.20 | 5.29e-03 | MD06G1091600 | 2.90 | 2.50e-06 | yes | endoglucanase 17-like | — |
| TRINITY_DN3180_c0_g1 | Rola-high | -3.20 | 6.98e-09 | MD11G1161800 | -3.22 | 3.35e-27 | | fatty acyl-CoA reductase 3-like | — |
| TRINITY_DN36069_c0_g1 | Rola-high | -3.20 | 2.46e-04 | MD15G1144900 | -2.94 | 1.86e-10 | yes | probable inactive receptor kinase At2g26730 | — |
| TRINITY_DN7210_c0_g1 | Rola-high | -3.19 | 5.42e-03 | MD01G1159600 | -4.41 | 2.83e-13 | yes | probably inactive leucine-rich repeat receptor-like protein kinase IMK2 | — |

| Trinity gene ID | Direction | Trinity | | GDDH13 protein match | | | Def. | Description | FCS-GX |
|---|---|---|---|---|---|---|---|---|---|
| | | log2FC | padj | gene ID | log2FC | padj | | | call |
| TRINITY_DN29142_c0_g1 | Rin-high | 3.19 | 1.98e-03 | MD09G1008100 | 3.16 | 2.75e-09 | | ACT domain-containing protein ACR6-like isoform X2 | — |
| TRINITY_DN16921_c0_g1 | Rin-high | 3.19 | 3.94e-04 | MD07G1236200 | 1.98 | 3.88e-08 | yes | probable LRR receptor-like serine/threonine-protein kinase At3g47570 isoform X2 | — |
| TRINITY_DN21646_c0_g1 | Rola-high | -3.19 | 4.37e-06 | MD11G1268700 | -4.41 | 3.80e-08 | | protein trichome birefringence-like 41 | — |
| TRINITY_DN9583_c1_g1 | Rola-high | -3.18 | 8.85e-03 | MD03G1170200 | -3.11 | 3.87e-06 | | ubiquitin-conjugating enzyme E2 20-like | — |
| TRINITY_DN372_c0_g3 | Rola-high | -3.18 | 9.52e-03 | MD01G1124600 | -4.58 | 3.08e-07 | | histone H4 | — |
| TRINITY_DN6594_c1_g2 | Rola-high | -3.17 | 2.92e-06 | MD16G1236000 | -3.18 | 7.50e-16 | | probable pectate lyase 8 | — |
| TRINITY_DN8116_c0_g1 | Rin-high | 3.17 | 2.54e-03 | MD12G1181200 | 3.04 | 5.99e-08 | yes | F-box/LRR-repeat protein 3-like | — |
| TRINITY_DN11636_c0_g1 | Rola-high | -3.16 | 8.81e-06 | MD10G1153200 | -3.26 | 5.28e-12 | | hypothetical protein DVH24_028509 | — |
| TRINITY_DN28053_c0_g1 | Rola-high | -3.16 | 9.86e-03 | MD13G1094100 | -3.30 | 1.94e-07 | | polygalacturonase At1g48100 | — |
| TRINITY_DN96967_c0_g1 | Rola-high | -3.16 | 9.92e-03 | MD12G1252500 | -3.73 | 6.83e-09 | yes | GDSL esterase/lipase At5g33370-like | — |
| TRINITY_DN10592_c0_g1 | Rola-high | -3.16 | 1.48e-03 | MD03G1171800 | -2.83 | 2.24e-07 | | zinc finger CCCH domain-containing protein 53-like | — |
| TRINITY_DN2244_c0_g1 | Rola-high | -3.16 | 2.58e-03 | MD01G1233900 | -3.05 | 1.23e-06 | yes | acidic leucine-rich nuclear phosphoprotein 32-related protein-like | — |
| TRINITY_DN1764_c0_g1 | Rola-high | -3.16 | 1.36e-05 | MD16G1236000 | -3.18 | 7.50e-16 | | probable pectate lyase 8 | — |
| TRINITY_DN6230_c0_g1 | Rola-high | -3.15 | 5.81e-03 | MD10G1319300 | -2.80 | 5.32e-06 | | uncharacterized protein | — |
| TRINITY_DN6722_c0_g1 | Rola-high | -3.15 | 5.78e-03 | MD13G1226700 | -3.04 | 1.73e-05 | | uncharacterized protein | — |
| TRINITY_DN4691_c0_g1 | Rola-high | -3.15 | 5.34e-04 | MD10G1121900 | -2.91 | 6.57e-06 | yes | thaumatin-like protein 1 | — |
| TRINITY_DN84104_c0_g2 | Rola-high | -3.15 | 5.51e-05 | MD15G1285000 | -3.08 | 2.03e-10 | | protein RALF-like 34 | — |
| TRINITY_DN5993_c0_g1 | Rola-high | -3.15 | 1.34e-06 | MD07G1125600 | -3.71 | 2.53e-13 | | uncharacterized protein | — |
| TRINITY_DN59703_c0_g1 | Rin-high | 3.15 | 9.24e-03 | MD11G1214800 | 3.02 | 2.17e-05 | yes | GDSL esterase/lipase At5g55050-like | — |
| TRINITY_DN3617_c0_g1 | Rin-high | 3.14 | 4.85e-03 | MD12G1174800 | 3.10 | 2.65e-06 | | nudix hydrolase 15, mitochondrial-like | — |
| TRINITY_DN7045_c0_g1 | Rin-high | 3.14 | 3.17e-04 | MD13G1104100 | 3.43 | 1.17e-05 | yes | cytochrome P450 71B26-like | — |
| TRINITY_DN8153_c0_g1 | Rola-high | -3.14 | 4.87e-03 | MD14G1028400 | -3.02 | 3.10e-06 | | kinesin-like protein KIN-5C | — |

| Trinity gene ID | Direction | Trinity | | GDDH13 protein match | | | Def. | Description | FCS-GX |
|---|---|---|---|---|---|---|---|---|---|
| | | log2FC | padj | gene ID | log2FC | padj | | | call |
| TRINITY_DN20359_c0_g1 | Rin-high | 3.14 | 3.24e-03 | MD05G1260400 | 1.25 | 2.44e-04 | yes | probable LRR receptor-like serine/threonine-protein kinase At1g56130 | — |
| TRINITY_DN5566_c0_g1 | Rola-high | -3.13 | 2.85e-06 | MD05G1080900 | -2.93 | 7.58e-11 | | AP2 domain class transcription factor | — |
| TRINITY_DN5258_c1_g2 | Rola-high | -3.13 | 2.51e-04 | MD10G1300500 | -3.03 | 7.08e-09 | | hypothetical protein DVH24_017152 | — |
| TRINITY_DN80531_c1_g3 | Rola-high | -3.12 | 4.56e-03 | MD10G1004700 | -1.44 | 2.60e-07 | yes | bifunctional aspartokinase/homoserine dehydrogenase 1, chloroplastic-like isoform X2 | Venturia inaequalis |
| TRINITY_DN5414_c0_g2 | Rola-high | -3.12 | 6.79e-03 | MD16G1022400 | -3.16 | 2.40e-05 | | HMG-Y-related protein A-like | — |
| TRINITY_DN2871_c0_g2 | Rola-high | -3.12 | 9.48e-03 | MD10G1073700 | -3.95 | 5.61e-04 | yes | laccase-11-like isoform X1 | — |
| TRINITY_DN10234_c0_g1 | Rola-high | -3.11 | 8.88e-04 | MD09G1200500 | -3.67 | 2.63e-09 | | acid phosphatase 1-like | — |
| TRINITY_DN23501_c0_g1 | Rola-high | -3.11 | 2.28e-03 | MD15G1016500 | -2.53 | 4.69e-09 | yes | probable LRR receptor-like serine/threonine-protein kinase At4g36180 | — |
| TRINITY_DN25171_c0_g1 | Rola-high | -3.10 | 4.54e-03 | MD13G1094100 | -3.30 | 1.94e-07 | | polygalacturonase At1g48100 | — |
| TRINITY_DN33817_c0_g1 | Rola-high | -3.09 | 4.89e-03 | MD15G1017600 | -3.57 | 3.55e-05 | | hypothetical protein DVH24_012957 | — |
| TRINITY_DN11693_c0_g1 | Rola-high | -3.09 | 3.77e-04 | MD03G1246700 | -3.11 | 7.28e-11 | | protein trichome birefringence-like 41 | — |
| TRINITY_DN81623_c0_g1 | Rin-high | 3.09 | 5.51e-04 | MD11G1314000 | 3.15 | 1.88e-10 | | calvin cycle protein CP12-1, chloroplastic-like | — |
| TRINITY_DN18127_c0_g1 | Rola-high | -3.09 | 1.79e-03 | MD12G1012200 | -2.87 | 1.10e-06 | | FT-interacting protein 3 | — |
| TRINITY_DN10562_c2_g3 | Rin-high | 3.09 | 7.31e-03 | MD02G1145800 | 3.21 | 9.45e-07 | | protein RADIALIS-like 4 | — |
| TRINITY_DN5989_c0_g1 | Rola-high | -3.09 | 4.10e-04 | MD00G1063500 | -3.04 | 1.72e-09 | yes | probable receptor-like protein kinase At1g11050 isoform X2 | — |
| TRINITY_DN9865_c0_g1 | Rin-high | 3.09 | 9.59e-03 | MD04G1243800 | 4.64 | 1.01e-03 | | auxin efflux carrier component 5-like | — |
| TRINITY_DN12548_c0_g1 | Rola-high | -3.08 | 1.72e-03 | MD08G1010400 | -2.16 | 2.90e-12 | | ABC transporter B family member 13-like | — |
| TRINITY_DN37771_c0_g1 | Rola-high | -3.07 | 5.29e-03 | MD15G1235100 | -2.88 | 1.15e-06 | | uncharacterized protein | — |
| TRINITY_DN35608_c0_g1 | Rin-high | 3.07 | 2.60e-05 | MD02G1045900 | 2.89 | 4.21e-15 | | probable chlorophyll(ide) b reductase NYC1, chloroplastic | — |
| TRINITY_DN52779_c0_g1 | Rola-high | -3.07 | 8.09e-03 | MD16G1097900 | -2.80 | 5.54e-11 | | protein neprosin-like | — |
| TRINITY_DN60803_c0_g3 | Rola-high | -3.07 | 2.37e-03 | MD15G1017600 | -3.57 | 3.55e-05 | | hypothetical protein DVH24_012957 | — |

| Trinity gene ID | Direction | Trinity | | GDDH13 protein match | | | Def. | Description | FCS-GX |
|---|---|---|---|---|---|---|---|---|---|
| | | log2FC | padj | gene ID | log2FC | padj | | | call |
| TRINITY_DN8356_c0_g1 | Rola-high | -3.07 | 3.45e-03 | MD13G1109700 | -3.10 | 1.28e-06 | | mitotic spindle checkpoint protein MAD2 | — |
| TRINITY_DN1059_c1_g1 | Rola-high | -3.07 | 1.64e-06 | MD17G1286700 | -3.12 | 1.01e-15 | | uncharacterized protein | — |
| TRINITY_DN6233_c0_g2 | Rola-high | -3.06 | 3.23e-03 | MD15G1435800 | -3.04 | 7.38e-08 | | uncharacterized protein | — |
| TRINITY_DN25121_c0_g1 | Rola-high | -3.05 | 5.39e-03 | MD13G1216600 | -3.19 | 1.81e-08 | | KIN14B-interacting protein At4g14310 | — |
| TRINITY_DN39255_c0_g1 | Rin-high | 3.05 | 5.19e-03 | MD04G1181800 | 3.71 | 8.51e-07 | yes | leucine-rich repeat receptor protein kinase HPCA1-like | — |
| TRINITY_DN26941_c0_g1 | Rola-high | -3.05 | 7.12e-03 | MD14G1220100 | -2.83 | 7.52e-06 | | kinesin-like protein KIN-8A isoform X2 | — |
| TRINITY_DN109731_c1_g1 | Rin-high | 3.04 | 9.04e-04 | MD11G1314000 | 3.15 | 1.88e-10 | | calvin cycle protein CP12-1, chloroplastic-like | — |
| TRINITY_DN2275_c0_g1 | Rin-high | 3.04 | 9.85e-04 | MD06G1032200 | 3.16 | 2.06e-08 | yes | disease resistance protein RPV1-like isoform X1 | — |
| TRINITY_DN2129_c0_g4 | Rin-high | 3.03 | 4.41e-03 | MD13G1207100 | 1.46 | 1.66e-04 | yes | calmodulin calcium-dependent NAD kinase-like isoform X1 | — |
| TRINITY_DN10219_c1_g1 | Rola-high | -3.03 | 5.57e-03 | MD10G1329000 | -3.59 | 1.70e-06 | | EH domain-containing protein 2-like | — |
| TRINITY_DN10912_c2_g1 | Rola-high | -3.02 | 4.62e-03 | MD15G1016500 | -2.53 | 4.69e-09 | yes | probable LRR receptor-like serine/threonine-protein kinase At4g36180 | — |
| TRINITY_DN3565_c0_g1 | Rola-high | -3.02 | 9.09e-03 | MD11G1012800 | -3.56 | 2.85e-06 | | neurofilament heavy polypeptide-like isoform X1 | — |
| TRINITY_DN8088_c0_g1 | Rola-high | -3.01 | 6.80e-03 | MD16G1222000 | -2.78 | 2.22e-05 | | KIN14B-interacting protein At4g14310-like | — |
| TRINITY_DN12417_c2_g1 | Rin-high | 3.01 | 8.67e-03 | MD04G1233500 | 3.11 | 2.88e-06 | | ammonium transporter 1 member 2-like | — |
| TRINITY_DN10023_c0_g1 | Rola-high | -3.01 | 4.92e-03 | MD15G1081400 | -2.70 | 6.03e-09 | | hypothetical protein DVH24_012465 | — |
| TRINITY_DN19285_c0_g1 | Rola-high | -3.01 | 2.83e-03 | MD14G1094000 | -2.62 | 3.18e-09 | yes | putative receptor kinase | — |
| TRINITY_DN21170_c0_g1 | Rola-high | -3.01 | 7.26e-03 | MD06G1103000 | -4.73 | 5.15e-08 | | F-box/kelch-repeat protein At3g06240-like | — |
| TRINITY_DN13418_c0_g1 | Rola-high | -3.00 | 8.02e-03 | MD09G1190900 | -3.06 | 1.86e-06 | | protein IQ-DOMAIN 8-like | — |
| TRINITY_DN62391_c1_g1 | Rin-high | 2.99 | 8.51e-03 | MD11G1314000 | 3.15 | 1.88e-10 | | calvin cycle protein CP12-1, chloroplastic-like | — |
| TRINITY_DN4754_c0_g1 | Rola-high | -2.99 | 1.96e-04 | MD05G1070800 | -2.11 | 1.99e-12 | | probable CoA ligase CCL6 isoform X2 | — |
| TRINITY_DN9948_c0_g1 | Rola-high | -2.99 | 5.93e-10 | MD02G1185300 | -2.97 | 2.15e-29 | | BTB/POZ domain-containing protein At1g30440-like | — |

| Trinity gene ID | Direction | Trinity | | GDDH13 protein match | | | Def. | Description | FCS-GX |
|---|---|---|---|---|---|---|---|---|---|
| | | log2FC | padj | gene ID | log2FC | padj | | | call |
| TRINITY_DN896_c0_g1 | Rin-high | 2.99 | 9.45e-03 | MD01G1211500 | 1.67 | 7.26e-05 | | S-type anion channel SLAH2-like isoform X1 | — |
| TRINITY_DN32604_c0_g1 | Rola-high | -2.98 | 4.99e-03 | MD04G1222000 | -2.80 | 9.09e-09 | yes | serine/threonine-protein kinase Nek3 isoform X4 | — |
| TRINITY_DN10693_c0_g2 | Rin-high | 2.98 | 8.89e-06 | MD10G1035700 | 3.05 | 1.98e-14 | | tryptophan aminotransferase-related protein 3-like | — |
| TRINITY_DN55223_c0_g1 | Rola-high | -2.98 | 2.98e-03 | MD10G1240800 | -3.01 | 1.34e-07 | | subtilisin-like protease SBT2.5 isoform X1 | — |
| TRINITY_DN1012_c1_g1 | Rola-high | -2.98 | 2.38e-03 | MD08G1120200 | -1.48 | 5.24e-03 | | protein ALTERED PHOSPHATE STARVATION RESPONSE 1-like | — |
| TRINITY_DN8207_c0_g1 | Rola-high | -2.98 | 8.47e-03 | MD14G1196400 | -2.85 | 4.25e-06 | | uncharacterized protein | — |
| TRINITY_DN10451_c0_g2 | Rola-high | -2.97 | 6.63e-04 | MD06G1022700 | -2.97 | 4.94e-09 | | DNA (cytosine-5)-methyltransferase CMT2 isoform X1 | — |
| TRINITY_DN1509_c0_g1 | Rola-high | -2.97 | 9.84e-04 | MD06G1173600 | -2.85 | 1.43e-08 | | DOF domain class transcription factor | — |
| TRINITY_DN23357_c0_g1 | Rin-high | 2.97 | 3.94e-07 | MD02G1045900 | 2.89 | 4.21e-15 | | probable chlorophyll(ide) b reductase NYC1, chloroplastic | — |
| TRINITY_DN216_c0_g2 | Rola-high | -2.96 | 5.34e-03 | MD06G1142300 | -2.95 | 5.63e-07 | | nucleotide pyrophosphatase/phosphodiesterase-like | — |
| TRINITY_DN9794_c1_g1 | Rin-high | 2.95 | 3.51e-03 | MD01G1199000 | 3.05 | 1.42e-07 | | uncharacterized protein | — |
| TRINITY_DN1314_c1_g1 | Rola-high | -2.95 | 1.51e-03 | MD13G1092000 | -2.41 | 4.79e-06 | | polygalacturonase 1 beta-like protein 3 | — |
| TRINITY_DN11372_c2_g1 | Rola-high | -2.95 | 1.38e-03 | MD05G1116400 | -2.76 | 5.99e-08 | | proline-rich protein 4 | — |
| TRINITY_DN13624_c0_g1 | Rola-high | -2.94 | 3.42e-04 | MD13G1146400 | -2.70 | 1.05e-11 | | probable boron transporter 2 isoform X2 | — |
| TRINITY_DN15047_c0_g1 | Rola-high | -2.94 | 6.89e-03 | MD09G1063100 | -2.87 | 3.88e-05 | | microtubule-destabilizing protein 60-like | — |
| TRINITY_DN62391_c0_g1 | Rin-high | 2.94 | 1.08e-03 | MD11G1314000 | 3.15 | 1.88e-10 | | calvin cycle protein CP12-1, chloroplastic-like | — |
| TRINITY_DN34923_c0_g1 | Rin-high | 2.94 | 4.22e-04 | MD08G1209800 | 3.21 | 5.51e-10 | | protein DETOXIFICATION 48-like | — |
| TRINITY_DN20226_c0_g1 | Rola-high | -2.93 | 3.10e-03 | MD05G1211600 | -2.88 | 3.11e-07 | yes | protein kinase PINOID 2-like | — |
| TRINITY_DN1276_c0_g1 | Rola-high | -2.93 | 1.92e-06 | MD06G1105900 | -2.97 | 1.10e-10 | yes | endoglucanase 11-like | — |
| TRINITY_DN3413_c0_g1 | Rola-high | -2.93 | 7.18e-03 | MD17G1201400 | -3.01 | 7.00e-07 | | uncharacterized protein | — |
| TRINITY_DN11579_c0_g1 | Rin-high | 2.93 | 2.12e-03 | MD14G1132900 | 2.91 | 1.34e-08 | | protein ABIL2-like | — |
| TRINITY_DN6070_c1_g1 | Rola-high | -2.92 | 1.74e-04 | MD08G1051200 | -2.90 | 2.30e-11 | yes | peroxidase 18 | — |

| Trinity gene ID | Direction | Trinity | | GDDH13 protein match | | | Def. | Description | FCS-GX |
|---|---|---|---|---|---|---|---|---|---|
| | | log2FC | padj | gene ID | log2FC | padj | | | call |
| TRINITY_DN22777_c0_g1 | Rola-high | -2.91 | 7.94e-04 | MD05G1242800 | -2.02 | 5.24e-10 | | galactan beta-1,4-galactosyltransferase GALS3-like | — |
| TRINITY_DN8216_c1_g1 | Rin-high | 2.91 | 2.49e-03 | MD01G1088100 | 3.14 | 7.81e-06 | yes | NAC domain-containing protein 43-like | — |
| TRINITY_DN7486_c0_g1 | Rola-high | -2.90 | 4.95e-03 | MD14G1059200 | -2.92 | 5.41e-07 | | uncharacterized protein | — |
| TRINITY_DN42650_c0_g1 | Rin-high | 2.90 | 1.26e-03 | MD07G1206400 | 2.86 | 2.53e-07 | | cucumisin isoform X3 | — |
| TRINITY_DN21864_c0_g1 | Rola-high | -2.90 | 9.21e-03 | MD11G1022700 | -2.23 | 5.67e-15 | | FT-interacting protein 3 | — |
| TRINITY_DN21541_c0_g1 | Rola-high | -2.89 | 3.68e-03 | MD10G1298500 | -3.95 | 1.47e-13 | yes | polyphenol oxidase, chloroplastic-like | — |
| TRINITY_DN21593_c0_g1 | Rola-high | -2.89 | 9.53e-04 | MD01G1143200 | -3.45 | 1.09e-14 | | UDP-glycosyltransferase 76F1-like | — |
| TRINITY_DN7067_c0_g1 | Rola-high | -2.89 | 4.60e-03 | MD10G1298500 | -3.95 | 1.47e-13 | yes | polyphenol oxidase, chloroplastic-like | — |
| TRINITY_DN72522_c0_g1 | Rin-high | 2.89 | 9.75e-03 | MD07G1206400 | 2.86 | 2.53e-07 | | cucumisin isoform X3 | — |
| TRINITY_DN4781_c2_g1 | Rin-high | 2.89 | 4.73e-04 | MD09G1008100 | 3.16 | 2.75e-09 | | ACT domain-containing protein ACR6-like isoform X1 | — |
| TRINITY_DN22036_c0_g1 | Rola-high | -2.89 | 8.93e-03 | MD13G1036800 | -2.83 | 2.53e-07 | yes | putative receptor kinase | — |
| TRINITY_DN36457_c0_g1 | Rola-high | -2.89 | 3.70e-04 | MD06G1100200 | -2.63 | 1.27e-17 | yes | probable inactive receptor kinase At4g23740 | — |
| TRINITY_DN59885_c0_g1 | Rola-high | -2.88 | 3.23e-03 | MD17G1027200 | -2.91 | 9.48e-05 | yes | cyclin-dependent protein kinase inhibitor SMR6-like | — |
| TRINITY_DN10173_c0_g1 | Rola-high | -2.88 | 2.62e-03 | MD06G1183000 | -2.83 | 1.03e-08 | | TORTIFOLIA1-like protein 3 | — |
| TRINITY_DN33996_c0_g1 | Rola-high | -2.88 | 6.28e-04 | MD10G1254200 | -1.22 | 2.39e-07 | | 187-kDa microtubule-associated protein AIR9-like | — |
| TRINITY_DN21566_c0_g1 | Rin-high | 2.88 | 7.78e-04 | MD07G1236800 | 3.98 | 3.09e-20 | yes | probable LRR receptor-like serine/threonine-protein kinase At3g47570 isoform X1 | — |
| TRINITY_DN46716_c0_g1 | Rin-high | 2.88 | 7.10e-04 | MD15G1217900 | 2.74 | 5.24e-11 | | thiol protease aleurain-like | — |
| TRINITY_DN12575_c0_g1 | Rola-high | -2.88 | 6.84e-03 | MD17G1095200 | -2.91 | 6.39e-09 | | uncharacterized protein | — |
| TRINITY_DN8484_c0_g1 | Rola-high | -2.87 | 7.18e-03 | MD14G1154200 | -3.00 | 1.14e-04 | | heavy metal-associated isoprenylated plant protein 7-like | — |
| TRINITY_DN71306_c0_g1 | Rola-high | -2.87 | 2.56e-03 | MD16G1043700 | -2.50 | 3.15e-09 | | alpha-xylosidase 1-like | — |
| TRINITY_DN2973_c0_g2 | Rin-high | 2.86 | 4.56e-03 | MD14G1137900 | 2.76 | 4.18e-07 | yes | NAC domain-containing protein 43-like | — |
| TRINITY_DN11693_c0_g3 | Rola-high | -2.86 | 1.56e-06 | MD03G1246700 | -3.11 | 7.28e-11 | | protein trichome birefringence-like 41 | — |

| Trinity gene ID | Direction | Trinity | | GDDH13 protein match | | | Def. | Description | FCS-GX |
|---|---|---|---|---|---|---|---|---|---|
| | | log2FC | padj | gene ID | log2FC | padj | | | call |
| TRINITY_DN4488_c1_g2 | Rola-high | -2.86 | 2.60e-03 | MD10G1240800 | -3.01 | 1.34e-07 | | subtilisin-like protease SBT2.5 isoform X1 | — |
| TRINITY_DN7515_c0_g1 | Rola-high | -2.83 | 2.56e-03 | MD09G1165600 | -2.82 | 2.72e-08 | | uncharacterized protein | — |
| TRINITY_DN51896_c1_g1 | Rin-high | 2.83 | 9.56e-04 | MD13G1090300 | 4.49 | 1.62e-06 | | probable carotenoid cleavage dioxygenase 4, chloroplastic | — |
| TRINITY_DN9233_c1_g1 | Rola-high | -2.83 | 9.75e-06 | MD11G1156200 | -3.35 | 2.68e-17 | | glucomannan 4-beta-mannosyltransferase 9-like | — |
| TRINITY_DN10325_c0_g1 | Rola-high | -2.83 | 4.84e-03 | MD10G1221000 | -2.90 | 8.19e-08 | | uncharacterized protein | — |
| TRINITY_DN38325_c0_g1 | Rola-high | -2.81 | 4.27e-03 | MD06G1030500 | -2.67 | 1.73e-07 | yes | serine/threonine-protein kinase BSK2 | — |
| TRINITY_DN20681_c0_g2 | Rola-high | -2.81 | 2.00e-04 | MD10G1190400 | -2.92 | 9.91e-07 | yes | cytochrome P450 94A1-like | — |
| TRINITY_DN4823_c0_g1 | Rin-high | 2.81 | 5.45e-04 | MD14G1003600 | 2.75 | 6.73e-12 | | uncharacterized protein | — |
| TRINITY_DN3999_c0_g1 | Rin-high | 2.81 | 9.55e-05 | MD13G1016500 | 3.12 | 1.83e-13 | yes | probable xyloglucan endotransglucosylase/hydrolase protein 30 | — |
| TRINITY_DN4988_c1_g1 | Rola-high | -2.81 | 2.07e-05 | MD03G1259100 | -2.83 | 4.17e-14 | | chlorophyllase-1-like | — |
| TRINITY_DN10953_c0_g1 | Rola-high | -2.81 | 3.20e-03 | MD16G1089300 | -3.14 | 1.86e-08 | | transcription factor PAR2 | — |
| TRINITY_DN890_c1_g1 | Rin-high | 2.80 | 5.67e-03 | MD02G1050800 | 2.10 | 3.62e-05 | | 1-aminocyclopropane-1-carboxylate oxidase homolog 1-like | — |
| TRINITY_DN15641_c0_g1 | Rola-high | -2.80 | 2.55e-03 | MD06G1175800 | -6.97 | 3.81e-06 | | polygalacturonase 1 beta-like protein 3 | — |
| TRINITY_DN11305_c0_g1 | Rola-high | -2.80 | 2.13e-03 | MD11G1084300 | -2.79 | 4.34e-09 | | uncharacterized protein | — |
| TRINITY_DN4920_c0_g1 | Rin-high | 2.79 | 7.29e-04 | MD05G1231700 | 5.57 | 4.57e-08 | yes | G-type lectin S-receptor-like serine/threonine-protein kinase At4g27290 isoform X1 | — |
| TRINITY_DN2029_c0_g1 | Rola-high | -2.79 | 1.61e-05 | MD14G1110300 | -2.81 | 1.87e-14 | | probable polygalacturonase | — |
| TRINITY_DN520_c0_g1 | Rola-high | -2.79 | 1.47e-05 | MD14G1119800 | -3.24 | 3.22e-10 | yes | probable inactive receptor kinase At4g23740 | — |
| TRINITY_DN51896_c0_g1 | Rin-high | 2.79 | 7.12e-04 | MD13G1090300 | 4.49 | 1.62e-06 | | probable carotenoid cleavage dioxygenase 4, chloroplastic | — |
| TRINITY_DN51896_c2_g1 | Rin-high | 2.79 | 1.94e-03 | MD16G1090700 | 2.76 | 1.43e-09 | | probable carotenoid cleavage dioxygenase 4, chloroplastic | — |
| TRINITY_DN612_c2_g1 | Rola-high | -2.79 | 1.16e-03 | MD12G1061300 | -2.91 | 2.49e-10 | | FT-interacting protein 3 | — |
| TRINITY_DN1390_c3_g1 | Rin-high | 2.78 | 4.14e-03 | MD02G1048100 | 2.81 | 8.98e-08 | | hypothetical protein C1H46_014782 | — |

| Trinity gene ID | Direction | Trinity | | GDDH13 protein match | | | Def. | Description | FCS-GX |
|---|---|---|---|---|---|---|---|---|---|
| | | log2FC | padj | gene ID | log2FC | padj | | | call |
| TRINITY_DN10251_c1_g1 | Rola-high | -2.77 | 3.69e-08 | MD07G1136200 | -1.53 | 6.66e-07 | | glycerol-3-phosphate acyltransferase RAM2-like | — |
| TRINITY_DN3818_c0_g1 | Rola-high | -2.77 | 6.72e-04 | MD04G1132500 | -2.75 | 1.48e-09 | yes | thaumatin-like protein | — |
| TRINITY_DN95024_c0_g2 | Rin-high | 2.76 | 4.96e-04 | MD15G1217900 | 2.74 | 5.24e-11 | | thiol protease aleurain-like | — |
| TRINITY_DN53114_c0_g1 | Rola-high | -2.75 | 5.34e-07 | MD17G1256500 | -2.69 | 1.09e-17 | | hypothetical protein DVH24_030924 | — |
| TRINITY_DN6249_c0_g1 | Rola-high | -2.75 | 4.39e-05 | MD03G1298300 | -2.81 | 6.50e-15 | | uncharacterized protein | — |
| TRINITY_DN3497_c0_g2 | Rola-high | -2.75 | 4.82e-03 | MD04G1239700 | -2.34 | 9.25e-10 | | sterol 3-beta-glucosyltransferase UGT80A2-like isoform X1 | — |
| TRINITY_DN4064_c0_g1 | Rola-high | -2.75 | 1.36e-03 | MD08G1017800 | -2.68 | 4.99e-07 | yes | probable LRR receptor-like serine/threonine-protein kinase At4g36180 | — |
| TRINITY_DN13441_c0_g1 | Rola-high | -2.75 | 2.71e-03 | MD15G1155300 | -2.80 | 4.00e-09 | | glycerophosphodiester phosphodiesterase GDPDL3-like | — |
| TRINITY_DN63130_c0_g1 | Rin-high | 2.73 | 5.33e-04 | MD15G1217900 | 2.74 | 5.24e-11 | | thiol protease aleurain-like | — |
| TRINITY_DN2916_c1_g1 | Rin-high | 2.72 | 2.37e-03 | MD08G1002100 | 2.60 | 5.42e-08 | | BTB/POZ domain-containing protein At2g30600 isoform X2 | — |
| TRINITY_DN56200_c0_g1 | Rin-high | 2.71 | 1.74e-04 | MD02G1136000 | 2.56 | 2.30e-09 | yes | cytochrome P450 84A1-like | — |
| TRINITY_DN3213_c0_g1 | Rin-high | 2.70 | 8.57e-04 | MD05G1325900 | 1.21 | 1.11e-05 | | dynein light chain 1, cytoplasmic-like | — |
| TRINITY_DN18346_c0_g1 | Rin-high | 2.70 | 1.18e-03 | MD16G1090700 | 2.76 | 1.43e-09 | | probable carotenoid cleavage dioxygenase 4, chloroplastic | — |
| TRINITY_DN67629_c0_g1 | Rin-high | 2.69 | 1.99e-05 | MD10G1035700 | 3.05 | 1.98e-14 | | tryptophan aminotransferase-related protein 3-like | — |
| TRINITY_DN7445_c0_g2 | Rola-high | -2.69 | 4.56e-07 | MD15G1211600 | -2.30 | 3.37e-19 | | protein ALP1-like | — |
| TRINITY_DN12285_c0_g1 | Rola-high | -2.69 | 9.00e-04 | MD16G1117200 | -2.57 | 8.96e-10 | | uncharacterized protein | — |
| TRINITY_DN9658_c1_g1 | Rin-high | 2.69 | 1.22e-03 | MD03G1081900 | 4.29 | 3.08e-11 | | UDP-glycosyltransferase 76E2-like | — |
| TRINITY_DN3769_c4_g1 | Rola-high | -2.68 | 5.08e-03 | MD13G1036800 | -2.83 | 2.53e-07 | yes | putative receptor kinase | — |
| TRINITY_DN6031_c1_g1 | Rola-high | -2.68 | 2.91e-06 | MD17G1237200 | -2.25 | 4.59e-15 | | wax ester synthase/diacylglycerol acyltransferase 11-like | — |
| TRINITY_DN5062_c0_g1 | Rin-high | 2.68 | 3.90e-03 | MD15G1120300 | 2.65 | 1.71e-07 | | GTP-binding nuclear protein Ran1-like | — |
| TRINITY_DN14010_c1_g1 | Rin-high | 2.67 | 4.56e-04 | MD02G1091800 | 2.62 | 8.51e-07 | | thiol protease aleurain-like | — |

| Trinity gene ID | Direction | Trinity | | GDDH13 protein match | | | Def. | Description | FCS-GX |
| --- | --- | --- | --- | --- | --- | --- | --- | --- | --- |
| | | log2FC | padj | gene ID | log2FC | padj | | | call |
| TRINITY_DN1056_c6_g1 | Rola-high | -2.67 | 9.71e-03 | MD10G1222900 | -2.93 | 9.45e-05 | yes | receptor-like protein EIX2 | — |
| TRINITY_DN12980_c0_g1 | Rola-high | -2.67 | 5.81e-03 | MD02G1216600 | -2.53 | 2.46e-07 | | uncharacterized protein | — |
| TRINITY_DN2261_c0_g2 | Rin-high | 2.66 | 9.25e-06 | MD02G1045900 | 2.89 | 4.21e-15 | | probable chlorophyll(ide) b reductase NYC1, chloroplastic | — |
| TRINITY_DN3574_c0_g1 | Rin-high | 2.66 | 4.26e-04 | MD17G1103300 | 2.66 | 2.70e-10 | | plastidic glucose transporter 4 | — |
| TRINITY_DN18469_c0_g1 | Rin-high | 2.66 | 1.19e-03 | MD16G1090700 | 2.76 | 1.43e-09 | | probable carotenoid cleavage dioxygenase 4, chloroplastic | — |
| TRINITY_DN16318_c0_g1 | Rola-high | -2.65 | 4.60e-03 | MD10G1221400 | -2.05 | 3.72e-12 | yes | receptor-like protein EIX2 | — |
| TRINITY_DN15999_c0_g1 | Rola-high | -2.63 | 8.12e-08 | MD14G1131500 | -2.61 | 4.34e-26 | | 3-oxoacyl-[acyl-carrier-protein] synthase I, chloroplastic-like | — |
| TRINITY_DN37474_c1_g1 | Rola-high | -2.63 | 7.45e-04 | MD13G1197900 | -2.11 | 2.00e-10 | | protein NETWORKED 1A-like | — |
| TRINITY_DN4406_c0_g1 | Rin-high | 2.63 | 5.33e-03 | MD05G1197200 | 2.80 | 4.77e-07 | | uncharacterized protein | — |
| TRINITY_DN734_c3_g2 | Rola-high | -2.62 | 8.68e-04 | MD00G1096700 | -2.61 | 1.27e-10 | | uncharacterized protein | — |
| TRINITY_DN2293_c1_g1 | Rin-high | 2.62 | 2.43e-03 | MD17G1212000 | 2.44 | 2.61e-06 | | protein POST-ILLUMINATION CHLOROPHYLL FLUORESCENCE INCREASE, chloroplastic | — |
| TRINITY_DN6651_c0_g1 | Rin-high | 2.62 | 7.44e-03 | MD12G1171500 | 2.91 | 4.84e-08 | | protein trichome birefringence-like 3 | — |
| TRINITY_DN7681_c0_g2 | Rola-high | -2.62 | 1.58e-04 | MD04G1027600 | -2.66 | 3.80e-14 | yes | probable LRR receptor-like serine/threonine-protein kinase IRK | — |
| TRINITY_DN3569_c0_g1 | Rola-high | -2.61 | 7.52e-04 | MD06G1157800 | -1.97 | 1.10e-07 | | E3 ubiquitin-protein ligase AIRP2-like | — |
| TRINITY_DN2774_c0_g1 | Rola-high | -2.61 | 1.13e-03 | MD14G1234700 | -3.99 | 5.46e-09 | | uncharacterized GPI-anchored protein At4g28100-like | — |
| TRINITY_DN15306_c0_g1 | Rola-high | -2.60 | 2.67e-05 | MD10G1283000 | -2.64 | 1.67e-17 | yes | GDSL esterase/lipase At1g28580-like | — |
| TRINITY_DN27436_c0_g1 | Rola-high | -2.60 | 2.09e-03 | MD14G1217100 | -2.32 | 5.86e-09 | | transcription factor BIM1 isoform X1 | — |
| TRINITY_DN40157_c0_g1 | Rin-high | 2.59 | 3.25e-03 | MD11G1212200 | 2.57 | 4.81e-08 | | hypothetical protein DVH24_003317 | — |
| TRINITY_DN7084_c1_g1 | Rola-high | -2.58 | 5.94e-03 | MD04G1126900 | -2.71 | 3.83e-09 | | hypothetical protein DVH24_026783 | — |
| TRINITY_DN13941_c1_g3 | Rola-high | -2.58 | 9.16e-03 | MD10G1121700 | -2.17 | 8.60e-06 | | auxin transporter-like protein 2 | — |
| TRINITY_DN7154_c0_g1 | Rola-high | -2.58 | 8.98e-04 | MD04G1215900 | -3.11 | 6.47e-07 | | growth-regulating factor 6-like isoform X2 | — |

| Trinity gene ID | Direction | Trinity | | GDDH13 protein match | | | Def. | Description | FCS-GX |
|---|---|---|---|---|---|---|---|---|---|
| | | log2FC | padj | gene ID | log2FC | padj | | | call |
| TRINITY_DN21032_c0_g1 | Rola-high | -2.58 | 2.89e-04 | MD07G1008000 | -2.70 | 4.20e-13 | | phenolic glucoside malonyltransferase 1-like | — |
| TRINITY_DN13747_c0_g1 | Rola-high | -2.57 | 1.16e-05 | MD15G1021100 | -3.29 | 7.80e-16 | | beta-galactosidase 5 | — |
| TRINITY_DN17568_c0_g1 | Rola-high | -2.57 | 1.70e-03 | MD10G1044500 | -2.56 | 6.67e-08 | yes | GDSL esterase/lipase At1g29670-like | — |
| TRINITY_DN2527_c0_g1 | Rola-high | -2.56 | 6.65e-03 | MD00G1006600 | -2.70 | 1.54e-07 | yes | probable leucine-rich repeat receptor-like serine/threonine-protein kinase At3g14840 isoform X2 | — |
| TRINITY_DN5864_c1_g1 | Rola-high | -2.56 | 5.38e-03 | MD10G1146200 | -2.60 | 1.97e-10 | | cytochrome b561 and DOMON domain-containing protein At3g07570-like | — |
| TRINITY_DN15829_c0_g1 | Rin-high | 2.56 | 4.32e-04 | MD14G1003600 | 2.75 | 6.73e-12 | | uncharacterized protein | — |
| TRINITY_DN24992_c0_g1 | Rola-high | -2.56 | 9.38e-03 | MD02G1188100 | -1.98 | 5.96e-09 | | CAMTA domain class transcription factor | — |
| TRINITY_DN2114_c0_g1 | Rin-high | 2.56 | 4.85e-03 | MD15G1264200 | 3.16 | 8.68e-05 | | sulfite exporter TauE/SafE family protein 3-like isoform X2 | — |
| TRINITY_DN1968_c0_g1 | Rola-high | -2.56 | 8.00e-04 | MD17G1129500 | -2.45 | 1.89e-07 | | UDP-glycosyltransferase 71A15-like | — |
| TRINITY_DN19439_c0_g1 | Rola-high | -2.55 | 3.01e-03 | MD03G1090700 | -2.62 | 1.87e-10 | | hypothetical protein DVH24_038635 | — |
| TRINITY_DN10225_c0_g1 | Rola-high | -2.55 | 2.92e-06 | MD04G1233600 | -2.56 | 4.32e-21 | | protein neprosin | — |
| TRINITY_DN70663_c0_g1 | Rin-high | 2.53 | 9.16e-03 | MD17G1212100 | 2.66 | 3.56e-09 | | protein POST-ILLUMINATION CHLOROPHYLL FLUORESCENCE INCREASE, chloroplastic | — |
| TRINITY_DN3926_c0_g1 | Rola-high | -2.53 | 5.39e-03 | MD10G1226800 | -2.35 | 4.16e-07 | | uncharacterized protein | — |
| TRINITY_DN28530_c0_g4 | Rola-high | -2.52 | 6.49e-03 | MD05G1354800 | -1.70 | 1.45e-11 | | uncharacterized protein | — |
| TRINITY_DN60040_c0_g1 | Rola-high | -2.52 | 4.40e-03 | MD13G1105800 | -1.98 | 9.44e-19 | | oxysterol-binding protein-related protein 1D-like | — |
| TRINITY_DN5296_c1_g4 | Rola-high | -2.52 | 3.28e-03 | MD11G1136500 | -2.30 | 1.46e-10 | | protein S40-4-like | — |
| TRINITY_DN12480_c0_g1 | Rola-high | -2.52 | 3.89e-03 | MD16G1083600 | -3.62 | 6.08e-09 | | TORTIFOLIA1-like protein 4 | — |
| TRINITY_DN9186_c1_g1 | Rola-high | -2.51 | 3.84e-03 | MD09G1075500 | -5.69 | 1.68e-12 | yes | leucine-rich repeat extensin-like protein 5 | — |
| TRINITY_DN9357_c1_g1 | Rola-high | -2.50 | 1.22e-03 | MD14G1071100 | -2.75 | 1.29e-10 | yes | leucine-rich repeat receptor-like tyrosine-protein kinase PXC3 | — |
| TRINITY_DN312_c0_g1 | Rola-high | -2.50 | 3.58e-03 | MD02G1258300 | -2.51 | 8.27e-09 | | uncharacterized protein At3g28850-like | — |

| Trinity gene ID | Direction | Trinity | | GDDH13 protein match | | | Def. | Description | FCS-GX |
|---|---|---|---|---|---|---|---|---|---|
| | | log2FC | padj | gene ID | log2FC | padj | | | call |
| TRINITY_DN5486_c0_g2 | Rin-high | 2.50 | 9.99e-03 | MD04G1183100 | 2.49 | 2.66e-05 | yes | leucine-rich repeat receptor protein kinase HPCA1-like | — |
| TRINITY_DN6976_c0_g1 | Rin-high | 2.50 | 2.94e-03 | MD04G1066300 | 2.49 | 7.22e-09 | yes | cytochrome P450 87A3-like | — |
| TRINITY_DN50729_c0_g1 | Rola-high | -2.50 | 9.90e-03 | MD03G1106300 | -2.53 | 3.09e-08 | | putative respiratory burst oxidase homolog protein H isoform X1 | — |
| TRINITY_DN26933_c0_g2 | Rola-high | -2.49 | 7.75e-03 | MD15G1120000 | -1.92 | 3.64e-06 | | BIIDXI-like protein At5g11420 | — |
| TRINITY_DN31025_c0_g1 | Rola-high | -2.48 | 6.25e-10 | MD09G1148500 | -2.50 | 9.27e-36 | yes | GDSL esterase/lipase At2g42990 | — |
| TRINITY_DN58303_c0_g1 | Rin-high | 2.47 | 5.17e-03 | MD04G1113200 | 2.59 | 6.03e-10 | | two-component response regulator-like APRR7 isoform X2 | — |
| TRINITY_DN1037_c1_g1 | Rola-high | -2.46 | 2.21e-03 | MD01G1043100 | -2.50 | 7.57e-12 | | probable mannitol dehydrogenase | — |
| TRINITY_DN24103_c0_g1 | Rola-high | -2.46 | 9.19e-03 | MD05G1338200 | -3.70 | 1.39e-07 | | transcription factor bHLH162-like | — |
| TRINITY_DN46723_c0_g1 | Rola-high | -2.45 | 3.43e-03 | MD17G1054500 | -2.50 | 1.08e-09 | | early nodulin-like protein 17 | — |
| TRINITY_DN54590_c1_g1 | Rola-high | -2.45 | 5.99e-03 | MD16G1049900 | -2.55 | 6.89e-09 | yes | receptor like protein 29-like | — |
| TRINITY_DN30653_c0_g1 | Rola-high | -2.44 | 1.15e-03 | MD11G1157600 | -1.44 | 2.93e-04 | | arogenate dehydratase 3-like | — |
| TRINITY_DN1390_c0_g1 | Rola-high | -2.44 | 5.01e-03 | MD07G1117200 | -2.51 | 2.91e-09 | | axial regulator YABBY 1-like | — |
| TRINITY_DN632_c1_g2 | Rola-high | -2.43 | 6.86e-03 | MD07G1159100 | -2.68 | 2.65e-07 | | uncharacterized protein | — |
| TRINITY_DN84104_c0_g1 | Rola-high | -2.43 | 1.91e-05 | MD02G1173500 | -2.39 | 1.48e-15 | | protein RALF-like 34 | — |
| TRINITY_DN14667_c1_g1 | Rola-high | -2.42 | 2.69e-07 | MD17G1144800 | -2.55 | 4.52e-24 | yes | probable xyloglucan glycosyltransferase 5 | — |
| TRINITY_DN10567_c1_g1 | Rin-high | 2.42 | 9.16e-04 | MD07G1236500 | 6.06 | 1.04e-18 | yes | probable LRR receptor-like serine/threonine-protein kinase At3g47570 isoform X2 | — |
| TRINITY_DN6753_c0_g1 | Rola-high | -2.41 | 1.43e-03 | MD00G1122700 | -1.20 | 4.53e-03 | | uncharacterized protein | — |
| TRINITY_DN13919_c0_g1 | Rola-high | -2.40 | 4.17e-03 | MD15G1232500 | -2.26 | 3.58e-09 | | protein RKD5 isoform X1 | — |
| TRINITY_DN14804_c0_g1 | Rola-high | -2.40 | 6.13e-04 | MD08G1010400 | -2.16 | 2.90e-12 | | ABC transporter B family member 13-like | — |
| TRINITY_DN12426_c0_g1 | Rola-high | -2.39 | 2.30e-03 | MD16G1043700 | -2.50 | 3.15e-09 | | alpha-xylosidase 1-like | — |
| TRINITY_DN48332_c0_g1 | Rin-high | 2.39 | 8.31e-03 | MD07G1083200 | 2.10 | 4.86e-09 | | ent-kaurenoic acid oxidase 1-like | — |
| TRINITY_DN1910_c0_g1 | Rin-high | 2.38 | 1.86e-03 | MD13G1162500 | 2.44 | 3.98e-10 | | probable protein phosphatase 2C 34 | — |

| Trinity gene ID | Direction | Trinity | | GDDH13 protein match | | | Def. | Description | FCS-GX |
|---|---|---|---|---|---|---|---|---|---|
| | | log2FC | padj | gene ID | log2FC | padj | | | call |
| TRINITY_DN10209_c0_g1 | Rola-high | -2.38 | 1.88e-03 | MD07G1251900 | -2.41 | 4.18e-08 | yes | leucine-rich repeat receptor-like protein kinase TDR | — |
| TRINITY_DN280_c0_g1 | Rin-high | 2.37 | 2.51e-03 | MD15G1226500 | 2.51 | 3.27e-07 | yes | probable L-type lectin-domain containing receptor kinase S.5 | — |
| TRINITY_DN5225_c1_g1 | Rin-high | 2.36 | 9.00e-03 | MD16G1238400 | 2.28 | 4.57e-08 | | FCS-Like Zinc finger 14-like | — |
| TRINITY_DN27950_c0_g1 | Rola-high | -2.35 | 6.83e-03 | MD15G1227900 | -1.54 | 3.47e-06 | | 3-hydroxy-3-methylglutaryl coenzyme A reductase | — |
| TRINITY_DN9983_c1_g2 | Rin-high | 2.35 | 5.97e-03 | MD11G1079500 | 3.01 | 9.46e-09 | | sodium-dependent phosphate transport protein 1, chloroplastic | — |
| TRINITY_DN28530_c0_g3 | Rola-high | -2.34 | 6.11e-03 | MD05G1354800 | -1.70 | 1.45e-11 | | uncharacterized protein | — |
| TRINITY_DN15703_c0_g1 | Rin-high | 2.34 | 1.95e-03 | MD07G1236500 | 6.06 | 1.04e-18 | yes | probable LRR receptor-like serine/threonine-protein kinase At3g47570 | — |
| TRINITY_DN32076_c1_g3 | Rin-high | 2.34 | 3.77e-03 | MD16G1023700 | 2.39 | 1.42e-10 | | uncharacterized protein | — |
| TRINITY_DN4123_c2_g1 | Rola-high | -2.33 | 2.66e-03 | MD17G1242600 | -2.00 | 3.29e-09 | yes | expansin-B3-like | — |
| TRINITY_DN55185_c0_g1 | Rola-high | -2.33 | 5.94e-03 | MD04G1025800 | -2.38 | 1.05e-10 | | rho GDP-dissociation inhibitor 1-like | — |
| TRINITY_DN3709_c0_g1 | Rola-high | -2.33 | 2.86e-03 | MD05G1185400 | -3.12 | 8.75e-10 | yes | receptor-like serine/threonine-protein kinase ALE2 isoform X1 | — |
| TRINITY_DN1695_c0_g2 | Rola-high | -2.32 | 5.47e-03 | MD15G1378900 | -2.27 | 2.80e-08 | | protein DETOXIFICATION 27-like | — |
| TRINITY_DN9100_c0_g1 | Rola-high | -2.32 | 5.98e-03 | MD17G1237600 | -5.74 | 1.98e-07 | | wax ester synthase/diacylglycerol acyltransferase 11-like | — |
| TRINITY_DN118_c0_g1 | Rin-high | 2.32 | 1.07e-03 | MD05G1260400 | 1.25 | 2.44e-04 | yes | probable LRR receptor-like serine/threonine-protein kinase At1g56130 | — |
| TRINITY_DN7445_c0_g1 | Rola-high | -2.31 | 1.77e-05 | MD15G1211600 | -2.30 | 3.37e-19 | | protein ALP1-like | — |
| TRINITY_DN9927_c0_g1 | Rola-high | -2.26 | 8.22e-03 | MD05G1170600 | -2.40 | 2.42e-08 | | hypothetical protein DVH24_010362 | — |
| TRINITY_DN494_c0_g1 | Rin-high | 2.23 | 6.83e-03 | MD01G1114400 | 2.13 | 8.38e-09 | | two-component response regulator-like PRR73 | — |
| TRINITY_DN26144_c0_g1 | Rola-high | -2.22 | 7.25e-03 | MD09G1241900 | -2.22 | 6.11e-10 | | protein E6 | — |
| TRINITY_DN10389_c0_g1 | Rin-high | 2.22 | 6.12e-03 | MD15G1384500 | 2.03 | 9.33e-08 | | MADS-box protein SVP-like isoform X1 | — |
| TRINITY_DN37706_c0_g1 | Rin-high | 2.22 | 4.90e-03 | MD17G1037500 | 2.13 | 1.65e-09 | | potassium transporter 6-like | — |

| Trinity gene ID | Direction | Trinity | | GDDH13 protein match | | | Def. | Description | FCS-GX |
|---|---|---|---|---|---|---|---|---|---|
| | | log2FC | padj | gene ID | log2FC | padj | | | call |
| TRINITY_DN4948_c1_g1 | Rola-high | -2.20 | 6.83e-03 | MD02G1201100 | -2.11 | 1.82e-07 | | fasciclin-like arabinogalactan protein 2 | — |
| TRINITY_DN906_c1_g1 | Rola-high | -2.19 | 9.45e-03 | MD11G1178000 | -2.51 | 1.40e-12 | | uncharacterized protein | — |
| TRINITY_DN51_c0_g1 | Rola-high | -2.19 | 1.01e-11 | MD09G1169500 | -2.99 | 7.05e-20 | | uncharacterized protein | — |
| TRINITY_DN19917_c0_g1 | Rin-high | 2.19 | 1.55e-03 | MD08G1007500 | 2.43 | 2.37e-12 | yes | thioredoxin-like protein CDSP32, chloroplastic | — |
| TRINITY_DN3168_c1_g1 | Rola-high | -2.18 | 4.25e-03 | MD04G1148900 | -2.99 | 3.73e-27 | | glycerol-3-phosphate 2-O-acyltransferase 6 | — |
| TRINITY_DN9177_c0_g1 | Rola-high | -2.18 | 3.03e-03 | MD09G1110300 | -2.23 | 1.98e-11 | | 5'-adenylylsulfate reductase 3, chloroplastic-like | — |
| TRINITY_DN10218_c0_g1 | Rola-high | -2.17 | 4.66e-03 | MD03G1090700 | -2.62 | 1.87e-10 | | hypothetical protein DVH24_038635 | — |
| TRINITY_DN32440_c0_g1 | Rola-high | -2.17 | 5.67e-03 | MD17G1215500 | -2.14 | 1.75e-10 | | VQ motif-containing protein 4 | — |
| TRINITY_DN13458_c0_g1 | Rin-high | 2.17 | 7.09e-03 | MD16G1241000 | 1.84 | 3.12e-09 | | isoamylase 1, chloroplastic | — |
| TRINITY_DN17011_c0_g1 | Rin-high | 2.16 | 2.61e-03 | MD17G1210200 | 2.12 | 1.17e-10 | | uncharacterized protein | — |
| TRINITY_DN10565_c0_g2 | Rola-high | -2.16 | 2.87e-03 | MD16G1195800 | -2.07 | 1.11e-09 | | uncharacterized protein | — |
| TRINITY_DN2630_c1_g1 | Rola-high | -2.15 | 6.97e-03 | MD03G1090700 | -2.62 | 1.87e-10 | | hypothetical protein DVH24_038635 | — |
| TRINITY_DN7001_c0_g1 | Rola-high | -2.15 | 3.83e-03 | MD10G1232600 | -2.23 | 1.84e-14 | | protein EXORDIUM-like 7 | — |
| TRINITY_DN1106_c1_g1 | Rin-high | 2.15 | 9.05e-03 | MD05G1121800 | 2.19 | 4.20e-10 | | heavy metal-associated isoprenylated plant protein 26 | — |
| TRINITY_DN7327_c0_g1 | Rola-high | -2.13 | 2.59e-04 | MD12G1139500 | -2.26 | 8.63e-18 | | short-chain dehydrogenase TIC 32 B, chloroplastic-like | — |
| TRINITY_DN20550_c0_g1 | Rin-high | 2.13 | 8.94e-03 | MD09G1176500 | 1.96 | 8.28e-08 | | uncharacterized protein | — |
| TRINITY_DN2836_c0_g2 | Rola-high | -2.12 | 4.54e-04 | MD14G1014300 | -1.24 | 2.32e-03 | | glucomannan 4-beta-mannosyltransferase 9-like | — |
| TRINITY_DN1074_c0_g2 | Rola-high | -2.11 | 3.75e-06 | MD15G1386800 | -2.09 | 7.95e-30 | | protein ECERIFERUM 26-like | — |
| TRINITY_DN1061_c2_g2 | Rin-high | 2.10 | 4.27e-04 | MD08G1238200 | 2.78 | 2.84e-11 | | serine carboxypeptidase-like 45 isoform X1 | — |
| TRINITY_DN2247_c0_g1 | Rola-high | -2.10 | 5.69e-03 | MD02G1136300 | -1.16 | 1.68e-05 | yes | probable leucine-rich repeat receptor-like protein kinase At1g35710 | — |
| TRINITY_DN1764_c4_g1 | Rola-high | -2.10 | 7.45e-03 | MD14G1167100 | -2.08 | 1.41e-09 | | probable pectate lyase 5 | — |
| TRINITY_DN15364_c0_g1 | Rola-high | -2.08 | 1.10e-05 | MD09G1158600 | -1.77 | 5.70e-14 | yes | probable xyloglucan glycosyltransferase 5 | — |

| Trinity gene ID | Direction | Trinity |  | GDDH13 protein match |  |  | Def. | Description | FCS-GX |
|---|---|---|---|---|---|---|---|---|---|
|  |  | log2FC | padj | gene ID | log2FC | padj |  |  | call |
| TRINITY_DN4754_c0_g2 | Rola-high | -2.08 | 5.94e-03 | MD05G1070800 | -2.11 | 1.99e-12 |  | probable CoA ligase CCL6 isoform X1 | — |
| TRINITY_DN82_c0_g1 | Rola-high | -2.08 | 4.75e-04 | MD11G1179700 | -2.05 | 1.58e-12 | yes | non-specific lipid transfer protein GPI-anchored 2-like isoform X2 | — |
| TRINITY_DN6498_c0_g2 | Rola-high | -2.07 | 1.79e-03 | MD03G1133700 | -1.85 | 4.61e-17 |  | glucomannan 4-beta-mannosyltransferase 2-like | — |
| TRINITY_DN2836_c0_g1 | Rola-high | -2.01 | 2.57e-04 | MD11G1156200 | -3.35 | 2.68e-17 |  | glucomannan 4-beta-mannosyltransferase 9-like | — |
| TRINITY_DN2559_c0_g2 | Rola-high | -1.99 | 2.08e-03 | MD15G1206900 | -2.29 | 5.79e-06 |  | probable polyamine oxidase 5 | — |
| TRINITY_DN13807_c0_g1 | Rola-high | -1.95 | 9.17e-03 | MD11G1218200 | -1.89 | 1.37e-09 |  | IQ domain-containing protein IQM4-like | — |
| TRINITY_DN4789_c2_g1 | Rola-high | -1.95 | 3.59e-03 | MD17G1140800 | -2.13 | 9.70e-23 |  | trifunctional UDP-glucose 4,6-dehydratase/UDP-4-keto-6-deoxy-D-glucose 3,5-epimerase/UDP-4-keto-L-rhamnose-reductase RHM1 | — |
| TRINITY_DN22704_c0_g1 | Rola-high | -1.94 | 2.45e-06 | MD13G1087700 | -2.00 | 2.66e-43 |  | aldehyde oxidase GLOX1-like | — |
| TRINITY_DN5921_c0_g1 | Rola-high | -1.91 | 2.60e-04 | MD10G1336500 | -2.30 | 3.39e-25 |  | CASP-like protein 2B1 isoform X2 | — |
| TRINITY_DN6728_c0_g1 | Rola-high | -1.88 | 1.27e-03 | MD15G1323600 | -1.94 | 2.13e-19 |  | uncharacterized protein | — |
| TRINITY_DN2446_c0_g1 | Rola-high | -1.87 | 1.12e-04 | MD12G1082400 | -2.02 | 2.18e-31 |  | glycerol-3-phosphate dehydrogenase [NAD(+)] GPDHC1, cytosolic | — |
| TRINITY_DN5914_c0_g1 | Rin-high | 1.84 | 8.15e-03 | MD09G1183800 | 1.86 | 8.56e-12 |  | transcription factor WER-like | — |
| TRINITY_DN10440_c0_g1 | Rola-high | -1.84 | 3.90e-05 | MD12G1082400 | -2.02 | 2.18e-31 |  | glycerol-3-phosphate dehydrogenase [NAD(+)] GPDHC1, cytosolic | — |
| TRINITY_DN7336_c0_g1 | Rola-high | -1.84 | 4.45e-03 | MD10G1290900 | -1.71 | 4.78e-09 | yes | ethylene-responsive transcription factor 12 | — |
| TRINITY_DN612_c1_g1 | Rola-high | -1.73 | 5.14e-03 | MD03G1021000 | -1.52 | 3.85e-16 |  | FT-interacting protein 3 | — |
| TRINITY_DN26385_c0_g1 | Rola-high | -1.72 | 4.56e-04 | MD01G1152500 | -1.77 | 2.33e-24 | yes | GDSL esterase/lipase WDL1-like isoform X2 | — |
| TRINITY_DN1126_c3_g1 | Rola-high | -1.70 | 2.99e-03 | MD04G1069000 | -1.69 | 3.92e-28 |  | protein HOTHEAD isoform X1 | — |
| TRINITY_DN8741_c0_g1 | Rola-high | -1.62 | 1.58e-04 | MD05G1242800 | -2.02 | 5.24e-10 |  | LOB domain-containing protein 37-like | — |
| TRINITY_DN13723_c0_g1 | Rola-high | -1.58 | 1.56e-05 | MD15G1262600 | -1.58 | 2.87e-50 |  | hypothetical protein DVH24_029871 | — |
| **Discordant — confident GDDH13 protein match, significant in GDDH13 but opposite direction (n = 9)** |  |  |  |  |  |  |  |  |  |

| Trinity gene ID | Direction | Trinity | | GDDH13 protein match | | | Def. | Description | FCS-GX |
|---|---|---|---|---|---|---|---|---|---|
| | | log2FC | padj | gene ID | log2FC | padj | | | call |
| TRINITY_DN28753_c0_g1 | Rola-high | -6.12 | 5.09e-04 | MD16G1265300 | 1.98 | 8.02e-04 | | (R)-mandelonitrile lyase 2-like isoform X1 | — |
| TRINITY_DN5527_c1_g1 | Rin-high | 5.10 | 4.86e-03 | MD17G1125800 | -3.87 | 7.16e-04 | | 7-deoxyloganetin glucosyltransferase-like | — |
| TRINITY_DN34159_c0_g2 | Rin-high | 5.08 | 3.04e-03 | MD17G1125800 | -3.87 | 7.16e-04 | | 7-deoxyloganetin glucosyltransferase-like | — |
| TRINITY_DN220_c0_g1 | Rola-high | -4.94 | 8.39e-09 | MD01G1044100 | 1.08 | 3.42e-04 | | serine hydroxymethyltransferase, mitochondrial | Venturia inaequalis |
| TRINITY_DN18211_c0_g1 | Rola-high | -4.39 | 6.21e-06 | MD04G1096600 | 1.46 | 3.19e-03 | | 29 kDa ribonucleoprotein A, chloroplastic | Venturia inaequalis |
| TRINITY_DN10820_c1_g4 | Rin-high | 3.80 | 2.76e-03 | MD16G1042700 | -2.44 | 1.65e-05 | | ribulose bisphosphate carboxylase small subunit, chloroplastic | — |
| TRINITY_DN36132_c0_g1 | Rola-high | -3.62 | 4.36e-05 | MD04G1153300 | 1.16 | 8.25e-03 | | pyridoxal 5'-phosphate synthase subunit PDX1.3 | Venturia inaequalis |
| TRINITY_DN35270_c0_g3 | Rola-high | -3.47 | 2.13e-03 | MD05G1220800 | 2.59 | 9.65e-06 | | elongation factor 2 isoform X2 | Venturia inaequalis |
| TRINITY_DN39211_c0_g2 | Rin-high | 3.32 | 3.96e-03 | MD10G1212300 | -1.83 | 8.22e-04 | | putative cyclin-B3-1 isoform X3 | — |
| **Confident GDDH13 protein match, not a GDDH13 candidate (Trinity-driven) (n = 330)** | | | | | | | | | |
| TRINITY_DN18390_c0_g1 | Rola-high | -22.72 | 9.81e-10 | MD10G1289300 | -0.45 | 1.72e-05 | | alpha-glucan phosphorylase, H isozyme-like | Trialeurodes vaporariorum |
| TRINITY_DN27044_c0_g1 | Rola-high | -7.87 | 2.03e-05 | MD02G1133600 | -3.12 | NA | | palmitoyl-monogalactosyldiacylglycerol delta-7 desaturase, chloroplastic-like | — |
| TRINITY_DN7926_c0_g2 | Rola-high | -7.61 | 9.01e-06 | MD12G1240900 | 0.56 | 1.43e-01 | | GDP-mannose 3,5-epimerase 2 | — |
| TRINITY_DN2351_c0_g2 | Rola-high | -7.12 | 2.33e-11 | MD15G1002900 | -0.47 | 4.70e-01 | | uncharacterized protein | — |
| TRINITY_DN29847_c0_g2 | Rola-high | -6.87 | 1.08e-04 | MD15G1219500 | -0.38 | 6.82e-01 | | BURP domain protein RD22-like | — |
| TRINITY_DN7285_c0_g1 | Rin-high | 6.79 | 1.51e-04 | MD14G1024500 | -0.98 | 1.04e-04 | | heat shock cognate 70 kDa protein-like | — |
| TRINITY_DN66304_c1_g1 | Rola-high | -6.52 | 2.37e-04 | MD16G1138000 | -0.55 | 1.84e-02 | | pleiotropic drug resistance protein 3-like | — |
| TRINITY_DN119652_c0_g1 | Rola-high | -6.33 | 1.48e-06 | MD14G1105600 | 1.02 | 1.57e-02 | | aminodeoxychorismate synthase, chloroplastic | Venturia inaequalis |
| TRINITY_DN17581_c0_g1 | Rola-high | -6.14 | 3.22e-04 | MD03G1210100 | 0.65 | 4.70e-03 | | thiamine thiazole synthase, chloroplastic | Venturia inaequalis |
| TRINITY_DN29354_c0_g1 | Rola-high | -6.09 | 5.49e-05 | MD17G1179500 | -0.64 | 4.71e-01 | | inositol oxygenase 2-like isoform X2 | Venturia inaequalis |

| Trinity gene ID | Direction | Trinity | | GDDH13 protein match | | | Def. | Description | FCS-GX |
|---|---|---|---|---|---|---|---|---|---|
| | | log2FC | padj | gene ID | log2FC | padj | | | call |
| TRINITY_DN25548_c0_g1 | Rola-high | -5.94 | 2.92e-13 | MD10G1227800 | 0.29 | 8.40e-01 | | fe(2+) transport protein 1-like | Venturia inaequalis |
| TRINITY_DN21389_c0_g2 | Rin-high | 5.88 | 1.48e-03 | MD16G1094900 | -3.45 | 3.38e-02 | | type 2 DNA topoisomerase 6 subunit B-like | — |
| TRINITY_DN61005_c0_g1 | Rola-high | -5.86 | 1.09e-03 | MD09G1162600 | -5.14 | 1.48e-02 | | 23 kDa jasmonate-induced protein-like | — |
| TRINITY_DN179381_c0_g1 | Rola-high | -5.84 | 2.09e-05 | MD07G1111000 | 0.79 | 1.25e-02 | | elongation factor G-2, mitochondrial | Venturia inaequalis |
| TRINITY_DN9192_c1_g1 | Rola-high | -5.79 | 3.48e-07 | MD10G1081500 | -1.45 | 3.64e-01 | yes | putative lipid-transfer protein DIR1 | — |
| TRINITY_DN228367_c0_g1 | Rola-high | -5.66 | 6.79e-08 | MD08G1087300 | 0.22 | 5.71e-01 | | small ribosomal subunit protein uS14z/uS14y/uS14x-like | Venturia inaequalis |
| TRINITY_DN14914_c0_g1 | Rola-high | -5.64 | 3.35e-04 | MD09G1051300 | -2.08 | 1.54e-02 | | gibberellin-regulated protein 6-like | — |
| TRINITY_DN52951_c0_g1 | Rin-high | 5.61 | 9.09e-05 | MD01G1118800 | 2.99 | 2.50e-01 | | 3-ketoacyl-CoA synthase 19 | — |
| TRINITY_DN178882_c0_g1 | Rola-high | -5.60 | 7.11e-05 | MD07G1296500 | 0.11 | 8.45e-01 | | cytochrome b5 | Venturia inaequalis |
| TRINITY_DN1367_c1_g3 | Rola-high | -5.59 | 1.54e-03 | MD10G1282000 | NA | NA | | hypothetical protein DVH24_041908 | — |
| TRINITY_DN34174_c0_g3 | Rola-high | -5.57 | 2.67e-10 | MD13G1090400 | 0.67 | 5.66e-02 | | spermine synthase-like | Venturia inaequalis |
| TRINITY_DN104398_c0_g1 | Rola-high | -5.51 | 7.85e-06 | MD07G1276800 | 0.55 | 3.73e-02 | | rRNA 2'-O-methyltransferase fibrillarin 2-like | Venturia inaequalis |
| TRINITY_DN163055_c0_g1 | Rola-high | -5.47 | 2.68e-03 | MD15G1298700 | -0.59 | 2.55e-01 | | heptahelical transmembrane protein 1-like isoform X2 | Venturia inaequalis |
| TRINITY_DN55425_c0_g1 | Rola-high | -5.45 | 6.00e-03 | MD05G1281900 | -0.36 | 5.98e-02 | | glycolate oxidase 1-like isoform X1 | Venturia inaequalis |
| TRINITY_DN77004_c0_g1 | Rola-high | -5.44 | 5.93e-10 | MD12G1201100 | 0.20 | 7.66e-01 | | S-adenosylmethionine synthase 1 | Venturia inaequalis |
| TRINITY_DN201878_c0_g1 | Rola-high | -5.42 | 7.39e-05 | MD02G1106200 | -0.41 | 1.87e-02 | | N-acylphosphatidylethanolamine synthase-like isoform X2 | Venturia inaequalis |
| TRINITY_DN110105_c0_g1 | Rola-high | -5.41 | 5.59e-03 | MD11G1013400 | NA | NA | | hypothetical protein DVH24_042068 | — |
| TRINITY_DN33059_c0_g1 | Rola-high | -5.36 | 2.92e-06 | MD12G1233200 | 0.84 | 9.58e-02 | | probable nucleolar protein 5-2 | Venturia inaequalis |
| TRINITY_DN48742_c0_g1 | Rola-high | -5.36 | 2.29e-06 | MD16G1014300 | -4.76 | 1.60e-02 | | branchpoint-bridging protein-like isoform X3 | — |
| TRINITY_DN32510_c0_g1 | Rola-high | -5.35 | 6.32e-04 | MD08G1205700 | 0.53 | 3.96e-01 | | uncharacterized protein | — |

| Trinity gene ID | Direction | Trinity | | GDDH13 protein match | | | Def. | Description | FCS-GX |
|---|---|---|---|---|---|---|---|---|---|
| | | log2FC | padj | gene ID | log2FC | padj | | | call |
| TRINITY_DN18268_c0_g1 | Rola-high | -5.30 | 3.93e-12 | MD12G1262900 | -0.15 | 6.69e-01 | | large ribosomal subunit protein eL38z/eL38y-like | Venturia inaequalis |
| TRINITY_DN50407_c0_g1 | Rin-high | 5.28 | 1.06e-06 | MD07G1187800 | -4.50 | 4.77e-02 | | 3-ketoacyl-CoA synthase 19-like | — |
| TRINITY_DN67934_c0_g1 | Rola-high | -5.26 | 1.94e-03 | MD02G1162200 | -0.97 | 5.09e-01 | | methyl jasmonate esterase 1-like | — |
| TRINITY_DN286_c3_g2 | Rola-high | -5.22 | 5.78e-03 | MD11G1150600 | -0.00 | 9.89e-01 | | uncharacterized protein | — |
| TRINITY_DN18916_c0_g3 | Rola-high | -5.16 | 2.98e-05 | MD11G1309600 | 0.04 | 8.77e-01 | | large ribosomal subunit protein eL22z-like | Venturia inaequalis |
| TRINITY_DN33244_c0_g2 | Rola-high | -5.15 | 1.24e-12 | MD09G1160400 | 0.04 | 9.07e-01 | | large ribosomal subunit protein uL30y | Venturia inaequalis |
| TRINITY_DN14350_c0_g1 | Rin-high | 5.14 | 7.72e-03 | MD01G1145300 | -0.39 | 2.96e-02 | | O-fucosyltransferase 28-like | — |
| TRINITY_DN16546_c0_g1 | Rin-high | 5.11 | 4.94e-05 | MD05G1256900 | 0.55 | 2.08e-03 | | protein ROOT HAIR DEFECTIVE 3 homolog 2-like isoform X3 | — |
| TRINITY_DN222530_c0_g1 | Rola-high | -5.10 | 1.47e-05 | MD05G1271100 | -0.06 | 5.40e-01 | | histidinol dehydrogenase, chloroplastic-like | Venturia inaequalis |
| TRINITY_DN48040_c0_g1 | Rola-high | -5.07 | 1.60e-14 | MD13G1281900 | -0.57 | 3.03e-04 | | adenosylhomocysteinase | Venturia inaequalis |
| TRINITY_DN222381_c0_g1 | Rola-high | -5.03 | 2.02e-03 | MD04G1044800 | 0.80 | 1.16e-03 | | 26S rRNA (cytosine-C(5))-methyltransferase NOP2B isoform X2 | Venturia inaequalis |
| TRINITY_DN137319_c0_g1 | Rola-high | -5.00 | 2.71e-05 | MD16G1277600 | 0.24 | 1.71e-01 | | nucleolar GTP-binding protein 1-like | Venturia inaequalis |
| TRINITY_DN31531_c0_g1 | Rola-high | -4.96 | 2.75e-04 | MD12G1244300 | 0.01 | 9.76e-01 | | phosphoribosylaminoimidazole carboxylase, chloroplastic-like isoform X1 | Venturia inaequalis |
| TRINITY_DN52846_c0_g1 | Rola-high | -4.96 | 8.25e-12 | MD11G1111400 | 0.93 | NA | | heat shock cognate 70 kDa protein 2-like | Venturia inaequalis |
| TRINITY_DN73967_c0_g1 | Rola-high | -4.95 | 9.65e-05 | MD03G1249000 | NA | NA | | protein trichome birefringence-like 41 | — |
| TRINITY_DN82247_c0_g1 | Rola-high | -4.91 | 2.82e-09 | MD12G1053800 | 0.08 | 8.52e-01 | | large ribosomal subunit protein eL21z/eL21y | Venturia inaequalis |
| TRINITY_DN25178_c0_g1 | Rola-high | -4.91 | 7.47e-03 | MD02G1307700 | -0.74 | 3.12e-01 | | protein TEEBE-like | — |
| TRINITY_DN139305_c0_g1 | Rola-high | -4.91 | 1.40e-04 | MD14G1220200 | 1.56 | 3.85e-02 | | DEAD-box ATP-dependent RNA helicase 51-like | Venturia inaequalis |
| TRINITY_DN115989_c0_g1 | Rola-high | -4.90 | 7.08e-07 | MD03G1270100 | 0.41 | 3.28e-01 | | elongation factor Tu, mitochondrial | Venturia inaequalis |

| Trinity gene ID | Direction | Trinity | | GDDH13 protein match | | | Def. | Description | FCS-GX |
|---|---|---|---|---|---|---|---|---|---|
| | | log2FC | padj | gene ID | log2FC | padj | | | call |
| TRINITY_DN52846_c0_g3 | Rola-high | -4.89 | 2.33e-11 | MD17G1225700 | -0.11 | 2.39e-01 | | heat shock 70 kDa protein | Venturia inaequalis |
| TRINITY_DN47914_c0_g1 | Rola-high | -4.88 | 7.57e-03 | MD15G1002900 | -0.47 | 4.70e-01 | | uncharacterized protein | — |
| TRINITY_DN28571_c0_g1 | Rola-high | -4.87 | 5.70e-08 | MD00G1055300 | 0.12 | 6.36e-01 | | citrate synthase, mitochondrial isoform X1 | Venturia inaequalis |
| TRINITY_DN1367_c1_g1 | Rola-high | -4.87 | 4.24e-04 | MD10G1282000 | NA | NA | | hypothetical protein DVH24_030185 | — |
| TRINITY_DN38900_c0_g1 | Rola-high | -4.87 | 4.14e-05 | MD10G1323700 | 0.61 | 2.17e-02 | | aminomethyltransferase, mitochondrial | Venturia inaequalis |
| TRINITY_DN63350_c0_g1 | Rola-high | -4.87 | 9.33e-03 | MD05G1109300 | -2.84 | 1.60e-01 | | hypothetical protein DVH24_033193 | — |
| TRINITY_DN13153_c0_g1 | Rola-high | -4.85 | 2.72e-06 | MD14G1149200 | -1.66 | 1.23e-02 | | mitochondrial phosphate carrier protein 2, mitochondrial-like | Venturia inaequalis |
| TRINITY_DN5011_c1_g1 | Rin-high | 4.81 | 1.16e-06 | MD03G1130100 | 0.13 | 8.88e-01 | | uncharacterized protein | — |
| TRINITY_DN4814_c3_g2 | Rin-high | 4.77 | 7.70e-03 | MD14G1095500 | NA | NA | | hypothetical protein DVH24_016646 | — |
| TRINITY_DN8804_c0_g2 | Rola-high | -4.76 | 1.27e-04 | MD05G1308800 | 0.59 | 4.82e-02 | | probable O-methyltransferase 3 | — |
| TRINITY_DN6068_c0_g1 | Rin-high | 4.76 | 4.15e-03 | MD05G1356900 | 1.67 | 1.11e-01 | | hypothetical protein DVH24_011781 | — |
| TRINITY_DN200851_c0_g1 | Rola-high | -4.75 | 9.51e-04 | MD15G1141700 | -0.65 | 2.22e-05 | | methylenetetrahydrofolate reductase (NADH) 2-like | Venturia inaequalis |
| TRINITY_DN223596_c0_g1 | Rola-high | -4.74 | 8.29e-04 | MD05G1158000 | 0.45 | 9.08e-02 | | chaperone protein dnaJ 49-like | Venturia inaequalis |
| TRINITY_DN157289_c0_g1 | Rola-high | -4.71 | 4.37e-04 | MD15G1098800 | 0.76 | 1.91e-01 | | BURP domain protein RD22-like | — |
| TRINITY_DN62919_c0_g1 | Rin-high | 4.69 | 7.11e-03 | MD17G1107600 | 3.31 | NA | yes | triacylglycerol lipase 2-like isoform X1 | — |
| TRINITY_DN257_c2_g1 | Rola-high | -4.66 | 5.49e-03 | MD09G1069500 | -0.43 | 9.45e-02 | | lipoxygenase | — |
| TRINITY_DN2992_c1_g1 | Rola-high | -4.66 | 3.69e-08 | MD11G1112900 | 0.02 | 9.62e-01 | | small ribosomal subunit protein uS8my | Venturia inaequalis |
| TRINITY_DN18365_c0_g1 | Rola-high | -4.65 | 3.60e-06 | MD01G1164700 | -0.20 | 2.22e-01 | yes | superoxide dismutase [Mn], mitochondrial-like isoform X2 | Venturia inaequalis |
| TRINITY_DN16770_c0_g1 | Rola-high | -4.65 | 2.25e-08 | MD03G1100500 | 0.34 | 2.66e-01 | | large ribosomal subunit protein uL23z | Venturia inaequalis |
| TRINITY_DN183148_c0_g1 | Rola-high | -4.62 | 6.34e-11 | MD06G1158400 | 0.23 | 6.10e-01 | | large ribosomal subunit protein uL24z-like | Venturia inaequalis |

| Trinity gene ID | Direction | Trinity | | GDDH13 protein match | | | Def. | Description | FCS-GX |
|---|---|---|---|---|---|---|---|---|---|
| | | log2FC | padj | gene ID | log2FC | padj | | | call |
| TRINITY_DN141167_c0_g1 | Rola-high | -4.60 | 6.58e-04 | MD02G1033700 | 0.70 | 2.08e-02 | | hypothetical protein DVH24_029175 | Venturia inaequalis |
| TRINITY_DN42818_c0_g1 | Rola-high | -4.59 | 3.50e-06 | MD17G1218700 | -0.03 | 8.58e-01 | | isocitrate dehydrogenase [NAD] catalytic subunit 5, mitochondrial-like | Venturia inaequalis |
| TRINITY_DN24548_c1_g1 | Rola-high | -4.58 | 3.76e-11 | MD11G1179000 | -0.45 | 2.40e-02 | | 5-methyltetrahydropteroyltriglutamate--homocysteine methyltransferase | Venturia inaequalis |
| TRINITY_DN19272_c0_g1 | Rola-high | -4.57 | 5.77e-07 | MD03G1012400 | -0.10 | 6.10e-01 | | formate--tetrahydrofolate ligase | Venturia inaequalis |
| TRINITY_DN6445_c1_g2 | Rola-high | -4.57 | 1.28e-06 | MD03G1118000 | -0.18 | 6.64e-01 | | large ribosomal subunit protein uL11-like | Venturia inaequalis |
| TRINITY_DN40677_c1_g1 | Rin-high | 4.57 | 1.61e-05 | MD05G1288400 | -0.93 | 2.06e-01 | | senescence-specific cysteine protease SAG12-like | — |
| TRINITY_DN2491_c3_g1 | Rin-high | 4.55 | 8.89e-04 | MD17G1165700 | -0.86 | 2.86e-02 | yes | protein NSP-INTERACTING KINASE 2-like isoform X1 | — |
| TRINITY_DN20686_c0_g1 | Rola-high | -4.54 | 1.06e-04 | MD15G1437400 | 0.98 | 2.29e-01 | yes | probable leucine-rich repeat receptor-like protein kinase IMK3 | — |
| TRINITY_DN158651_c0_g1 | Rola-high | -4.52 | 5.94e-03 | MD14G1124700 | 0.45 | 2.68e-01 | | MYG1 protein C694.04c-like | Venturia inaequalis |
| TRINITY_DN27328_c0_g1 | Rola-high | -4.50 | 2.02e-09 | MD09G1175600 | 0.16 | 6.96e-01 | | small ribosomal subunit protein uS2-like | Venturia inaequalis |
| TRINITY_DN32987_c0_g1 | Rola-high | -4.50 | 5.60e-06 | MD13G1000500 | -0.14 | 4.29e-01 | yes | pyruvate kinase, cytosolic isozyme-like | Venturia inaequalis |
| TRINITY_DN14867_c0_g1 | Rola-high | -4.49 | 8.28e-05 | MD01G1185400 | 0.89 | 1.70e-01 | | FK506-binding protein | Venturia inaequalis |
| TRINITY_DN56398_c0_g2 | Rola-high | -4.48 | 3.36e-07 | MD10G1241900 | 0.10 | 7.16e-01 | | ribosomal protein L2 | Venturia inaequalis |
| TRINITY_DN44754_c0_g2 | Rola-high | -4.46 | 9.98e-05 | MD09G1007800 | -0.29 | 5.33e-01 | | large ribosomal subunit protein eL8y | Venturia inaequalis |
| TRINITY_DN163435_c0_g1 | Rola-high | -4.44 | 6.18e-10 | MD10G1164200 | 0.10 | 7.90e-01 | | large ribosomal subunit protein uL13w | Venturia inaequalis |
| TRINITY_DN35954_c0_g1 | Rola-high | -4.43 | 2.22e-06 | MD15G1203800 | 0.46 | 1.22e-01 | | probable protein arginine N-methyltransferase 1.2 isoform X1 | Venturia inaequalis |
| TRINITY_DN49617_c0_g1 | Rola-high | -4.43 | 2.23e-07 | MD01G1099100 | -0.20 | 7.76e-01 | | small ribosomal subunit protein uS15 | Venturia inaequalis |
| TRINITY_DN7286_c0_g1 | Rola-high | -4.41 | 2.17e-03 | MD10G1337900 | 0.21 | 5.48e-01 | | UPF0481 protein | Venturia inaequalis |

| Trinity gene ID | Direction | Trinity | | GDDH13 protein match | | | Def. | Description | FCS-GX |
|---|---|---|---|---|---|---|---|---|---|
| | | log2FC | padj | gene ID | log2FC | padj | | | call |
| TRINITY_DN159946_c0_g1 | Rola-high | -4.41 | 1.18e-03 | MD07G1141700 | -0.27 | 3.44e-01 | | phosphomannomutase-like isoform X1 | Venturia inaequalis |
| TRINITY_DN86671_c1_g3 | Rola-high | -4.40 | 5.59e-08 | MD16G1198000 | -0.09 | 7.96e-01 | | small ribosomal subunit protein eS1y-like | Venturia inaequalis |
| TRINITY_DN22073_c0_g1 | Rola-high | -4.39 | 7.48e-05 | MD07G1304700 | -0.16 | 5.45e-01 | | large ribosomal subunit protein eL13x-like | Venturia inaequalis |
| TRINITY_DN30245_c0_g1 | Rola-high | -4.38 | 9.51e-07 | MD10G1170700 | 0.63 | 4.63e-02 | | chaperonin CPN60-2, mitochondrial | Venturia inaequalis |
| TRINITY_DN46767_c0_g1 | Rola-high | -4.37 | 9.49e-07 | MD01G1099700 | 0.42 | 1.36e-02 | | dihydrolipoyllysine-residue acetyltransferase component 1 of pyruvate dehydrogenase complex, mitochondrial-like isoform X2 | Venturia inaequalis |
| TRINITY_DN4242_c2_g1 | Rola-high | -4.37 | 1.29e-04 | MD05G1102100 | 0.39 | 3.43e-01 | | cyclin-D1-1-like | — |
| TRINITY_DN12317_c0_g1 | Rola-high | -4.36 | 2.37e-06 | MD17G1112900 | -0.64 | 2.95e-01 | | glucan endo-1,3-beta-glucosidase 1 | — |
| TRINITY_DN14189_c0_g3 | Rola-high | -4.36 | 5.73e-04 | MD05G1049700 | 0.97 | 1.74e-03 | | heat shock 70 kDa protein, mitochondrial | Venturia inaequalis |
| TRINITY_DN45095_c0_g1 | Rola-high | -4.35 | 3.35e-06 | MD17G1128200 | -0.04 | 9.32e-01 | | large ribosomal subunit protein eL27-like | Venturia inaequalis |
| TRINITY_DN21926_c0_g2 | Rin-high | 4.34 | 4.75e-04 | MD09G1288900 | 2.05 | NA | | protein RADIALIS-like 3 | — |
| TRINITY_DN39809_c0_g1 | Rola-high | -4.34 | 5.72e-03 | MD07G1166600 | -0.92 | 4.49e-03 | | dihydrolipoyllysine-residue acetyltransferase component 1 of pyruvate dehydrogenase complex, mitochondrial-like isoform X2 | Venturia inaequalis |
| TRINITY_DN37627_c0_g3 | Rola-high | -4.34 | 9.94e-07 | MD15G1155000 | -0.03 | 9.37e-01 | | small ribosomal subunit protein eS12-like | Venturia inaequalis |
| TRINITY_DN20950_c0_g1 | Rola-high | -4.34 | 1.87e-03 | MD17G1203500 | NA | NA | | BIIDXI-like protein At5g11420 | — |
| TRINITY_DN44093_c1_g1 | Rola-high | -4.33 | 3.46e-07 | MD11G1125000 | NA | NA | | small ribosomal subunit protein eS21y | Venturia inaequalis |
| TRINITY_DN45228_c0_g1 | Rola-high | -4.33 | 1.58e-07 | MD11G1063600 | 0.28 | 2.54e-01 | | small ribosomal subunit protein eS4 | Venturia inaequalis |
| TRINITY_DN39775_c0_g1 | Rola-high | -4.33 | 8.07e-05 | MD15G1439400 | 0.29 | 8.33e-02 | | leucine--tRNA ligase, cytoplasmic-like | Venturia inaequalis |
| TRINITY_DN80142_c0_g1 | Rola-high | -4.33 | 3.75e-06 | MD03G1111400 | 0.57 | 2.13e-01 | | carbamoyl phosphate synthase arginine-specific large chain, chloroplastic-like | Venturia inaequalis |
| TRINITY_DN46229_c0_g1 | Rola-high | -4.33 | 8.02e-04 | MD00G1089700 | 0.23 | 6.82e-02 | | guanine nucleotide-binding protein-like NSN1 | Venturia inaequalis |

| Trinity gene ID | Direction | Trinity | | GDDH13 protein match | | | Def. | Description | FCS-GX |
|---|---|---|---|---|---|---|---|---|---|
| | | log2FC | padj | gene ID | log2FC | padj | | | call |
| TRINITY_DN15371_c1_g1 | Rola-high | -4.32 | 1.82e-08 | MD15G1060200 | 0.84 | 9.44e-02 | | glycine dehydrogenase (decarboxylating), mitochondrial | Venturia inaequalis |
| TRINITY_DN29378_c0_g1 | Rola-high | -4.32 | 1.75e-08 | MD07G1172600 | -0.77 | 3.33e-04 | yes | adenosine kinase 2 | Venturia inaequalis |
| TRINITY_DN20632_c1_g1 | Rola-high | -4.31 | 1.18e-08 | MD17G1049900 | 0.20 | 5.57e-01 | | small ribosomal subunit protein uS4y | Venturia inaequalis |
| TRINITY_DN24791_c0_g1 | Rola-high | -4.31 | 7.83e-03 | MD11G1306200 | -0.39 | 8.41e-02 | yes | ethylene receptor | — |
| TRINITY_DN4841_c0_g1 | Rola-high | -4.28 | 1.58e-04 | MD07G1256900 | 0.29 | 3.56e-01 | | small ribosomal subunit protein uS10y | Venturia inaequalis |
| TRINITY_DN61110_c0_g1 | Rola-high | -4.28 | 1.52e-08 | MD01G1145400 | 0.35 | 4.13e-01 | | large ribosomal subunit protein eL33w-like | Venturia inaequalis |
| TRINITY_DN11424_c0_g1 | Rola-high | -4.28 | 1.58e-07 | MD11G1160300 | 0.25 | 4.52e-01 | | large ribosomal subunit protein P2y-like | Venturia inaequalis |
| TRINITY_DN25116_c0_g1 | Rola-high | -4.26 | 2.71e-04 | MD02G1035100 | -0.02 | 9.14e-01 | | NADH-cytochrome b5 reductase-like protein | Venturia inaequalis |
| TRINITY_DN136381_c0_g1 | Rola-high | -4.25 | 2.65e-06 | MD07G1188600 | -0.10 | 8.36e-01 | | large ribosomal subunit protein uL29 | Venturia inaequalis |
| TRINITY_DN45026_c0_g1 | Rola-high | -4.24 | 4.28e-03 | MD05G1213600 | -0.80 | 2.90e-02 | | kinesin-like protein KIN-14Q isoform X1 | — |
| TRINITY_DN361_c0_g1 | Rin-high | 4.23 | 1.37e-03 | MD11G1095800 | 0.20 | 6.47e-01 | | transcription factor PCL1-like | — |
| TRINITY_DN2563_c0_g1 | Rola-high | -4.22 | 1.99e-09 | MD07G1302000 | 0.04 | 9.11e-01 | | large ribosomal subunit protein uL6 | Venturia inaequalis |
| TRINITY_DN15654_c0_g2 | Rin-high | 4.22 | 1.73e-05 | MD16G1136100 | 0.79 | 4.23e-03 | yes | CBL-interacting serine/threonine-protein kinase 1-like | — |
| TRINITY_DN21580_c0_g1 | Rola-high | -4.20 | 7.19e-08 | MD04G1023100 | 0.22 | 6.37e-01 | | small ribosomal subunit protein eS8 | Venturia inaequalis |
| TRINITY_DN25048_c0_g1 | Rola-high | -4.19 | 3.10e-05 | MD02G1004200 | 0.10 | 7.00e-01 | | large ribosomal subunit protein uL18 | Venturia inaequalis |
| TRINITY_DN36635_c0_g1 | Rola-high | -4.18 | 5.34e-03 | MD09G1051300 | -2.08 | 1.54e-02 | | gibberellin-regulated protein 6-like | — |
| TRINITY_DN92274_c0_g1 | Rola-high | -4.18 | 3.04e-04 | MD10G1179600 | 0.08 | 5.09e-01 | | succinate--CoA ligase [ADP-forming] subunit beta, mitochondrial | Venturia inaequalis |
| TRINITY_DN25207_c0_g1 | Rola-high | -4.18 | 4.24e-05 | MD11G1191800 | 0.08 | 7.97e-01 | | small ribosomal subunit protein uS5w-like | Venturia inaequalis |
| TRINITY_DN69195_c0_g1 | Rola-high | -4.17 | 5.29e-03 | MD12G1013300 | 0.07 | 6.29e-01 | | NADH--cytochrome b5 reductase 1 | Venturia inaequalis |

| Trinity gene ID | Direction | Trinity | | GDDH13 protein match | | | Def. | Description | FCS-GX |
|---|---|---|---|---|---|---|---|---|---|
| | | log2FC | padj | gene ID | log2FC | padj | | | call |
| TRINITY_DN36099_c0_g1 | Rola-high | -4.16 | 1.45e-04 | MD13G1192900 | -0.08 | 8.42e-01 | | small ribosomal subunit protein uS13z/uS13y/uS13x | Venturia inaequalis |
| TRINITY_DN14430_c1_g1 | Rola-high | -4.16 | 1.45e-03 | MD12G1228300 | -0.73 | 7.56e-02 | | protein BONZAI 3-like isoform X2 | — |
| TRINITY_DN25125_c0_g1 | Rola-high | -4.15 | 1.28e-05 | MD06G1217300 | -0.03 | 9.34e-01 | | small ribosomal subunit protein eS7 | Venturia inaequalis |
| TRINITY_DN35139_c0_g1 | Rola-high | -4.15 | 8.25e-07 | MD11G1307900 | -0.10 | 7.28e-01 | | large ribosomal subunit protein eL18y | Venturia inaequalis |
| TRINITY_DN80823_c0_g1 | Rola-high | -4.14 | 2.80e-03 | MD08G1028900 | -0.28 | 4.74e-02 | | proteasome subunit alpha type-5 | Venturia inaequalis |
| TRINITY_DN9431_c0_g2 | Rola-high | -4.14 | 3.42e-04 | MD03G1164100 | -0.11 | 5.12e-01 | | T-complex protein 1 subunit eta | Venturia inaequalis |
| TRINITY_DN90889_c0_g1 | Rin-high | 4.13 | 1.46e-03 | MD08G1241100 | 0.22 | 4.09e-01 | | uncharacterized protein | — |
| TRINITY_DN35905_c0_g1 | Rin-high | 4.13 | 5.51e-05 | MD11G1094500 | 0.51 | 6.79e-03 | | callose synthase 10-like isoform X1 | — |
| TRINITY_DN61270_c1_g1 | Rola-high | -4.12 | 8.24e-07 | MD08G1153000 | -0.09 | 7.69e-01 | | small ribosomal subunit protein uS17 | Venturia inaequalis |
| TRINITY_DN3027_c0_g1 | Rin-high | 4.11 | 7.71e-04 | MD08G1111300 | 1.15 | 3.93e-01 | | histidine-containing phosphotransfer protein 4-like | — |
| TRINITY_DN43148_c0_g1 | Rola-high | -4.11 | 1.51e-04 | MD07G1178200 | -0.23 | 4.87e-01 | | large ribosomal subunit protein eL36x-like | Venturia inaequalis |
| TRINITY_DN14002_c0_g1 | Rola-high | -4.11 | 1.16e-06 | MD12G1049600 | -0.15 | 3.01e-01 | | small ribosomal subunit protein uS7 | Venturia inaequalis |
| TRINITY_DN13016_c0_g1 | Rola-high | -4.10 | 2.06e-04 | MD06G1043100 | 0.84 | 1.50e-01 | | H/ACA ribonucleoprotein complex subunit 4-like isoform X1 | Venturia inaequalis |
| TRINITY_DN39372_c0_g1 | Rola-high | -4.10 | 3.19e-06 | MD00G1160700 | -0.07 | 8.45e-01 | | small ribosomal subunit protein RACK1 | Venturia inaequalis |
| TRINITY_DN47902_c0_g1 | Rola-high | -4.10 | 1.02e-03 | MD13G1125300 | 0.06 | 7.95e-01 | | pyruvate dehydrogenase E1 component subunit beta-1, mitochondrial | Venturia inaequalis |
| TRINITY_DN33940_c0_g2 | Rola-high | -4.09 | 9.10e-05 | MD03G1008200 | -0.45 | 1.57e-01 | | hypothetical protein DVH24_041777 | — |
| TRINITY_DN199883_c0_g1 | Rola-high | -4.09 | 2.49e-04 | MD12G1083100 | -0.10 | 7.60e-01 | | eukaryotic translation initiation factor 3 subunit E | Venturia inaequalis |
| TRINITY_DN113738_c0_g1 | Rola-high | -4.08 | 5.40e-03 | MD15G1437700 | -0.08 | 7.85e-01 | | phosphoribosylamine--glycine ligase-like isoform X1 | Venturia inaequalis |
| TRINITY_DN32442_c0_g1 | Rola-high | -4.08 | 7.61e-05 | MD03G1201800 | -0.04 | 7.41e-01 | | heat shock 70 kDa protein 15-like | Venturia inaequalis |

| Trinity gene ID | Direction | Trinity | | GDDH13 protein match | | | Def. | Description | FCS-GX |
|---|---|---|---|---|---|---|---|---|---|
| | | log2FC | padj | gene ID | log2FC | padj | | | call |
| TRINITY_DN18510_c0_g1 | Rola-high | -4.08 | 3.43e-03 | MD05G1194800 | 0.60 | 1.51e-01 | | H/ACA ribonucleoprotein complex subunit 3-like protein | Venturia inaequalis |
| TRINITY_DN3490_c1_g1 | Rin-high | 4.07 | 7.04e-03 | MD11G1317300 | NA | NA | | pentatricopeptide repeat-containing protein At2g32630-like isoform X2 | — |
| TRINITY_DN200796_c0_g1 | Rola-high | -4.07 | 3.26e-07 | MD09G1050600 | 0.12 | 7.30e-01 | | large ribosomal subunit protein eL34-like | Venturia inaequalis |
| TRINITY_DN32661_c0_g1 | Rola-high | -4.07 | 7.70e-07 | MD16G1225500 | 0.32 | 3.75e-01 | | large ribosomal subunit protein uL14x/uL14z/uL14y | Venturia inaequalis |
| TRINITY_DN140188_c0_g1 | Rola-high | -4.07 | 5.60e-05 | MD07G1232200 | -0.02 | 8.98e-01 | yes | superoxide dismutase [Mn], mitochondrial | Venturia inaequalis |
| TRINITY_DN64019_c0_g1 | Rin-high | 4.06 | 1.69e-04 | MD02G1053200 | 0.71 | 5.68e-01 | | 1-aminocyclopropane-1-carboxylate oxidase homolog 1-like | — |
| TRINITY_DN43140_c0_g1 | Rola-high | -4.06 | 3.24e-04 | MD05G1251000 | 0.47 | 2.85e-02 | | glycine cleavage system H protein 2, mitochondrial isoform X3 | Venturia inaequalis |
| TRINITY_DN24640_c0_g1 | Rin-high | 4.06 | 4.02e-03 | MD02G1303800 | 0.26 | 4.64e-01 | | phenylacetaldehyde reductase-like | — |
| TRINITY_DN20761_c0_g1 | Rola-high | -4.06 | 5.19e-03 | MD15G1431500 | 0.24 | 5.55e-01 | | large ribosomal subunit protein eL30 | Venturia inaequalis |
| TRINITY_DN43362_c0_g1 | Rin-high | 4.05 | 1.66e-04 | MD05G1266400 | -1.88 | 7.39e-02 | | berberine bridge enzyme-like 8 | — |
| TRINITY_DN188959_c0_g1 | Rola-high | -4.05 | 2.49e-06 | MD09G1235900 | -0.13 | 7.37e-01 | | large ribosomal subunit protein uL5 isoform X1 | Venturia inaequalis |
| TRINITY_DN8436_c1_g1 | Rola-high | -4.05 | 3.58e-03 | MD16G1163500 | 2.95 | NA | | hypothetical protein DVH24_025471 | — |
| TRINITY_DN75307_c0_g1 | Rola-high | -4.04 | 1.66e-07 | MD10G1263500 | -0.05 | 9.09e-01 | | large ribosomal subunit protein eL20y-like | Venturia inaequalis |
| TRINITY_DN25044_c0_g1 | Rola-high | -4.04 | 1.59e-05 | MD17G1057300 | -0.06 | 8.72e-01 | | small ribosomal subunit protein eS19x | Venturia inaequalis |
| TRINITY_DN188942_c0_g1 | Rola-high | -4.02 | 3.46e-05 | MD00G1143000 | 0.04 | 9.16e-01 | | small ribosomal subunit protein uS9-like | Venturia inaequalis |
| TRINITY_DN55940_c0_g3 | Rola-high | -4.02 | 5.71e-03 | MD07G1262800 | 0.42 | 2.46e-02 | | valine--tRNA ligase, mitochondrial 1-like | Venturia inaequalis |
| TRINITY_DN3561_c2_g1 | Rola-high | -4.01 | 6.27e-06 | MD11G1166000 | 0.15 | 6.20e-01 | | small ribosomal subunit protein eS25-like | Venturia inaequalis |
| TRINITY_DN12479_c0_g1 | Rin-high | 4.01 | 7.32e-04 | MD15G1033100 | 0.93 | 6.67e-01 | | hypothetical protein DVH24_012825 | — |
| TRINITY_DN81098_c0_g1 | Rola-high | -4.01 | 2.25e-06 | MD07G1008900 | 0.00 | 9.98e-01 | | ADP,ATP carrier protein 1, mitochondrial-like | Venturia inaequalis |

| Trinity gene ID | Direction | Trinity | | GDDH13 protein match | | | Def. | Description | FCS-GX |
|---|---|---|---|---|---|---|---|---|---|
| | | log2FC | padj | gene ID | log2FC | padj | | | call |
| TRINITY_DN89262_c0_g1 | Rola-high | -4.01 | 3.65e-06 | MD14G1062400 | 0.11 | 8.03e-01 | | large ribosomal subunit protein eL6z-like | Venturia inaequalis |
| TRINITY_DN222517_c0_g1 | Rola-high | -4.00 | 2.09e-04 | MD16G1145800 | 0.53 | 9.59e-03 | | dihydrolipoyl dehydrogenase, mitochondrial | Venturia inaequalis |
| TRINITY_DN17728_c0_g1 | Rola-high | -4.00 | 4.49e-03 | MD11G1217800 | -0.21 | 2.09e-02 | | reactive Intermediate Deaminase A, chloroplastic-like | Venturia inaequalis |
| TRINITY_DN27446_c0_g1 | Rola-high | -4.00 | 1.50e-06 | MD10G1154100 | 0.24 | 4.10e-01 | | large ribosomal subunit protein eL32z | Venturia inaequalis |
| TRINITY_DN28402_c0_g1 | Rola-high | -3.99 | 1.58e-03 | MD10G1111800 | -0.06 | 8.43e-01 | | tubulin alpha chain isoform X1 | — |
| TRINITY_DN37151_c1_g1 | Rola-high | -3.99 | 2.64e-03 | MD10G1263300 | 0.19 | 5.82e-01 | | small ribosomal subunit protein uS11x | Venturia inaequalis |
| TRINITY_DN10237_c0_g1 | Rola-high | -3.99 | 8.37e-07 | MD16G1251600 | -0.06 | 8.82e-01 | | large ribosomal subunit protein uL4 | Venturia inaequalis |
| TRINITY_DN37111_c0_g1 | Rola-high | -3.99 | 9.91e-03 | MD03G1208800 | 0.68 | 1.73e-03 | | tryptophan synthase alpha chain-like | Venturia inaequalis |
| TRINITY_DN20031_c0_g2 | Rola-high | -3.98 | 2.51e-06 | MD11G1123500 | -0.05 | 8.86e-01 | | small ribosomal subunit protein uS12 | Venturia inaequalis |
| TRINITY_DN71947_c0_g3 | Rola-high | -3.98 | 2.92e-06 | MD13G1207000 | 0.16 | 6.23e-01 | | small ribosomal subunit protein uS19x | Venturia inaequalis |
| TRINITY_DN3239_c1_g1 | Rola-high | -3.97 | 3.42e-07 | MD16G1033600 | 0.17 | 4.43e-01 | | ATP synthase subunit beta, mitochondrial | Venturia inaequalis |
| TRINITY_DN4330_c0_g2 | Rin-high | 3.97 | 8.58e-03 | MD14G1095500 | NA | NA | | V-type proton ATPase 16 kDa proteolipid subunit | — |
| TRINITY_DN84104_c0_g3 | Rola-high | -3.97 | 7.88e-07 | MD08G1186900 | -0.02 | 9.58e-01 | | small ribosomal subunit protein eS6 | Venturia inaequalis |
| TRINITY_DN27415_c0_g1 | Rola-high | -3.96 | 8.53e-04 | MD06G1144100 | 0.29 | 2.69e-01 | | argininosuccinate synthase, chloroplastic | Venturia inaequalis |
| TRINITY_DN10956_c0_g1 | Rola-high | -3.96 | 4.99e-03 | MD15G1445000 | -1.80 | NA | | hypothetical protein DVH24_003275, partial | — |
| TRINITY_DN81808_c0_g1 | Rola-high | -3.96 | 8.32e-04 | MD07G1232300 | 0.02 | 9.50e-01 | | small ribosomal subunit protein eS26x-like | Venturia inaequalis |
| TRINITY_DN95056_c0_g1 | Rola-high | -3.95 | 6.62e-06 | MD15G1214900 | 0.42 | 2.77e-01 | | small ribosomal subunit protein uS3x-like | Venturia inaequalis |
| TRINITY_DN40978_c0_g2 | Rola-high | -3.95 | 1.39e-07 | MD01G1148900 | -0.23 | 4.36e-01 | | large ribosomal subunit protein uL10 | Venturia inaequalis |

| Trinity gene ID | Direction | Trinity | | GDDH13 protein match | | | Def. | Description | FCS-GX |
|---|---|---|---|---|---|---|---|---|---|
| | | log2FC | padj | gene ID | log2FC | padj | | | call |
| TRINITY_DN6276_c0_g2 | Rola-high | -3.95 | 1.56e-05 | MD16G1271600 | -0.10 | 8.22e-01 | | V-type proton ATPase subunit B2 | Venturia inaequalis |
| TRINITY_DN60769_c0_g1 | Rin-high | 3.94 | 4.72e-04 | MD02G1104900 | 1.51 | 4.38e-01 | | senescence-specific cysteine protease SAG12-like | — |
| TRINITY_DN201715_c0_g1 | Rola-high | -3.94 | 1.12e-04 | MD08G1063300 | 0.30 | 1.75e-04 | | ATP-dependent zinc metalloprotease FTSH 4, mitochondrial-like | Venturia inaequalis |
| TRINITY_DN48934_c0_g1 | Rola-high | -3.93 | 2.75e-07 | MD15G1435700 | -0.83 | 1.67e-02 | | large ribosomal subunit protein uL3 | Venturia inaequalis |
| TRINITY_DN27331_c0_g1 | Rola-high | -3.93 | 1.39e-05 | MD04G1003600 | -0.98 | 4.22e-02 | | dihydroxy-acid dehydratase, chloroplastic | Venturia inaequalis |
| TRINITY_DN126885_c0_g1 | Rin-high | 3.93 | 7.80e-03 | MD15G1130100 | 0.25 | 8.37e-01 | | flotillin-like protein 4 | — |
| TRINITY_DN35181_c0_g1 | Rola-high | -3.92 | 3.11e-04 | MD03G1111400 | 0.57 | 2.13e-01 | | acetyl-CoA carboxylase 1-like | Venturia inaequalis |
| TRINITY_DN33162_c0_g1 | Rola-high | -3.90 | 6.92e-06 | MD15G1157300 | 0.07 | 8.88e-01 | | large ribosomal subunit protein eL39z/eL39x | Venturia inaequalis |
| TRINITY_DN23524_c0_g1 | Rola-high | -3.90 | 3.59e-05 | MD15G1403200 | -0.07 | 7.44e-01 | | elongation factor 1-gamma isoform X1 | Venturia inaequalis |
| TRINITY_DN27383_c0_g1 | Rola-high | -3.90 | 5.87e-06 | MD14G1246300 | 0.23 | 1.35e-01 | | nascent polypeptide-associated complex subunit beta | Venturia inaequalis |
| TRINITY_DN50481_c0_g1 | Rola-high | -3.90 | 3.95e-09 | MD03G1210400 | -1.33 | 1.44e-01 | | ubiquitin-ribosomal protein eS31 fusion protein | Venturia inaequalis |
| TRINITY_DN31655_c1_g2 | Rola-high | -3.90 | 1.05e-04 | MD17G1026900 | 0.14 | 7.15e-01 | | large ribosomal subunit protein uL15x-like | Venturia inaequalis |
| TRINITY_DN78153_c0_g1 | Rola-high | -3.89 | 9.76e-07 | MD08G1119100 | -0.15 | 6.66e-01 | | large ribosomal subunit protein eL37x | Venturia inaequalis |
| TRINITY_DN49601_c2_g2 | Rola-high | -3.88 | 2.37e-05 | MD01G1171100 | 0.37 | 1.74e-01 | | large ribosomal subunit protein uL16 | Venturia inaequalis |
| TRINITY_DN63226_c0_g1 | Rola-high | -3.88 | 6.44e-07 | MD09G1011700 | -0.03 | 9.42e-01 | | large ribosomal subunit protein uL22y-like | Venturia inaequalis |
| TRINITY_DN46406_c0_g1 | Rola-high | -3.87 | 2.65e-03 | MD04G1013400 | 0.08 | 5.16e-01 | | probable methionine--tRNA ligase isoform X2 | Venturia inaequalis |
| TRINITY_DN12804_c0_g1 | Rola-high | -3.87 | 1.15e-03 | MD09G1194200 | -0.04 | 7.26e-01 | | DEAD-box ATP-dependent RNA helicase 20 isoform X1 | Venturia inaequalis |
| TRINITY_DN5504_c0_g1 | Rola-high | -3.86 | 6.54e-08 | MD08G1231800 | -0.40 | 3.43e-01 | | small ribosomal subunit protein eS30z/eS30y/eS30x | Venturia inaequalis |

| Trinity gene ID | Direction | Trinity | | GDDH13 protein match | | | Def. | Description | FCS-GX |
|---|---|---|---|---|---|---|---|---|---|
| | | log2FC | padj | gene ID | log2FC | padj | | | call |
| TRINITY_DN213365_c0_g1 | Rin-high | 3.86 | 3.67e-03 | MD14G1167800 | 0.22 | 5.48e-01 | | glycine-rich RNA-binding protein GRP1A-like | — |
| TRINITY_DN23001_c0_g1 | Rin-high | 3.86 | 7.80e-03 | MD06G1163800 | -1.07 | 7.76e-02 | yes | cytochrome P450 736A117-like | — |
| TRINITY_DN222028_c0_g1 | Rola-high | -3.85 | 1.14e-04 | MD06G1070000 | -0.36 | 9.79e-02 | | hydroxymethylglutaryl-CoA synthase-like | Venturia inaequalis |
| TRINITY_DN20126_c0_g1 | Rola-high | -3.85 | 1.08e-05 | MD05G1338100 | -0.01 | 9.69e-01 | | eukaryotic translation initiation factor 3 subunit D-like | Venturia inaequalis |
| TRINITY_DN7900_c0_g1 | Rola-high | -3.85 | 3.20e-03 | MD16G1079400 | 0.12 | 8.07e-01 | | homeobox-leucine zipper protein HDG11-like | Venturia inaequalis |
| TRINITY_DN49487_c0_g1 | Rola-high | -3.84 | 2.63e-04 | MD09G1278300 | 0.01 | 9.76e-01 | | UTP--glucose-1-phosphate uridylyltransferase | Venturia inaequalis |
| TRINITY_DN1466_c1_g1 | Rin-high | 3.84 | 4.77e-15 | MD10G1079800 | -0.54 | 1.97e-01 | | putative polyol transporter 1 | — |
| TRINITY_DN227712_c0_g1 | Rola-high | -3.84 | 1.82e-03 | MD13G1013500 | -0.08 | 7.66e-01 | | eukaryotic translation initiation factor 3 subunit I-like | Venturia inaequalis |
| TRINITY_DN55921_c0_g1 | Rola-high | -3.84 | 8.05e-09 | MD14G1022900 | 0.12 | 7.05e-01 | | large ribosomal subunit protein uL1-like | Penicillium olsonii |
| TRINITY_DN5728_c0_g1 | Rin-high | 3.83 | 2.68e-03 | MD10G1281600 | 1.11 | 2.92e-01 | | senescence-specific cysteine protease SAG12-like | — |
| TRINITY_DN28175_c0_g1 | Rola-high | -3.83 | 2.24e-06 | MD04G1062000 | 0.18 | 6.72e-01 | | large ribosomal subunit protein eL15z | Venturia inaequalis |
| TRINITY_DN31573_c1_g1 | Rola-high | -3.83 | 3.33e-07 | MD16G1127800 | 0.52 | 1.94e-01 | | small ribosomal subunit protein eS27y-like | Venturia nashicola |
| TRINITY_DN37096_c0_g4 | Rola-high | -3.83 | 6.53e-05 | MD16G1033600 | 0.17 | 4.43e-01 | | ATP synthase subunit beta, mitochondrial | Venturia inaequalis |
| TRINITY_DN8684_c0_g1 | Rola-high | -3.83 | 1.65e-05 | MD04G1096100 | 0.58 | 2.15e-02 | | isocitrate dehydrogenase [NAD] catalytic subunit 5, mitochondrial-like | Venturia inaequalis |
| TRINITY_DN22410_c0_g1 | Rola-high | -3.82 | 4.72e-04 | MD06G1089600 | 0.07 | 7.82e-01 | | small ribosomal subunit protein eS10z-like | Venturia inaequalis |
| TRINITY_DN10728_c0_g1 | Rola-high | -3.82 | 6.56e-07 | MD08G1034600 | -0.15 | 5.75e-01 | | glucose-6-phosphate isomerase, cytosolic | Venturia inaequalis |
| TRINITY_DN12648_c1_g1 | Rola-high | -3.82 | 2.05e-03 | MD15G1123700 | -1.56 | 1.74e-01 | | EG45-like domain containing protein isoform X1 | Venturia inaequalis |
| TRINITY_DN30193_c0_g3 | Rola-high | -3.81 | 6.06e-04 | MD01G1228000 | -0.44 | 4.01e-01 | | large ribosomal subunit protein P1-like | Venturia inaequalis |
| TRINITY_DN114966_c0_g1 | Rola-high | -3.81 | 4.77e-04 | MD08G1247600 | 0.28 | 2.43e-02 | | DExH-box ATP-dependent RNA helicase DExH10 | Venturia inaequalis |

| Trinity gene ID | Direction | Trinity | | GDDH13 protein match | | | Def. | Description | FCS-GX |
|---|---|---|---|---|---|---|---|---|---|
| | | log2FC | padj | gene ID | log2FC | padj | | | call |
| TRINITY_DN96576_c0_g2 | Rola-high | -3.81 | 1.58e-04 | MD08G1190300 | 0.09 | 7.80e-01 | | small ribosomal subunit protein eS17w-like | Venturia inaequalis |
| TRINITY_DN113720_c0_g1 | Rola-high | -3.80 | 1.12e-03 | MD00G1145600 | 0.27 | 4.34e-01 | | large ribosomal subunit protein eL24 | Venturia inaequalis |
| TRINITY_DN14003_c0_g2 | Rola-high | -3.78 | 5.51e-04 | MD04G1188300 | NA | NA | | uncharacterized acetyltransferase At3g50280-like | — |
| TRINITY_DN29325_c0_g1 | Rola-high | -3.78 | 2.05e-04 | MD13G1106100 | -0.10 | 6.89e-01 | | T-complex protein 1 subunit epsilon | Venturia inaequalis |
| TRINITY_DN158481_c0_g1 | Rola-high | -3.75 | 1.08e-03 | MD05G1325700 | -0.84 | 2.88e-01 | | external alternative NAD(P)H-ubiquinone oxidoreductase B2, mitochondrial-like | Venturia inaequalis |
| TRINITY_DN23984_c0_g1 | Rola-high | -3.75 | 2.08e-03 | MD11G1210600 | 0.18 | 4.29e-01 | | nascent polypeptide-associated complex subunit alpha-like protein 2 | Venturia inaequalis |
| TRINITY_DN70268_c0_g1 | Rola-high | -3.75 | 9.17e-03 | MD06G1028800 | -0.12 | 8.33e-01 | | protein transport protein Sec61 subunit gamma | Venturia inaequalis |
| TRINITY_DN21194_c0_g1 | Rola-high | -3.74 | 4.94e-04 | MD14G1014600 | 0.27 | 3.22e-02 | | enolase 1, chloroplastic | Venturia inaequalis |
| TRINITY_DN23873_c0_g1 | Rola-high | -3.74 | 1.87e-03 | MD08G1134400 | 0.38 | 1.27e-01 | | aspartate--tRNA ligase 2, cytoplasmic-like | Venturia inaequalis |
| TRINITY_DN51368_c0_g1 | Rola-high | -3.73 | 9.65e-03 | MD02G1072600 | 0.04 | 8.92e-01 | | large ribosomal subunit protein eL31 | Venturia inaequalis |
| TRINITY_DN94840_c0_g1 | Rola-high | -3.73 | 4.88e-04 | MD02G1082700 | 0.14 | 3.92e-01 | | eukaryotic initiation factor 4A-8 | Venturia inaequalis |
| TRINITY_DN20722_c0_g2 | Rola-high | -3.73 | 7.92e-04 | MD16G1154000 | -0.24 | 4.97e-01 | | ATP-citrate synthase beta chain protein 2-like | Venturia inaequalis |
| TRINITY_DN201021_c0_g1 | Rola-high | -3.73 | 5.32e-03 | MD05G1003500 | 0.19 | 5.18e-02 | | histidine--tRNA ligase, cytoplasmic | Venturia inaequalis |
| TRINITY_DN7937_c0_g1 | Rin-high | 3.72 | 5.06e-05 | MD16G1085200 | -0.16 | 7.56e-01 | | TRINITY_DN7937_c0_g1 | — |
| TRINITY_DN1893_c0_g2 | Rola-high | -3.72 | 4.83e-03 | MD02G1096200 | 0.73 | 7.09e-06 | | oligouridylate-binding protein 1B-like | Venturia inaequalis |
| TRINITY_DN32610_c0_g1 | Rola-high | -3.71 | 5.32e-04 | MD06G1231700 | 0.09 | 7.27e-01 | | large ribosomal subunit protein eL42 | Venturia inaequalis |
| TRINITY_DN6154_c0_g1 | Rola-high | -3.70 | 7.48e-05 | MD05G1272300 | 0.33 | 1.69e-03 | | probable GTP-binding protein OBGM, mitochondrial | Venturia inaequalis |
| TRINITY_DN1437_c2_g1 | Rola-high | -3.70 | 1.95e-04 | MD15G1375400 | 0.52 | 7.71e-01 | | uncharacterized protein | — |

| Trinity gene ID | Direction | Trinity | | GDDH13 protein match | | | Def. | Description | FCS-GX |
| --- | --- | --- | --- | --- | --- | --- | --- | --- | --- |
| | | log2FC | padj | gene ID | log2FC | padj | | | call |
| TRINITY_DN105811_c0_g1 | Rola-high | -3.69 | 3.41e-05 | MD03G1007400 | 0.72 | 6.31e-02 | | DEAD-box ATP-dependent RNA helicase 37-like | Venturia inaequalis |
| TRINITY_DN6761_c0_g1 | Rola-high | -3.69 | 2.14e-03 | MD02G1062900 | 0.35 | 4.28e-02 | | peroxiredoxin-2F, mitochondrial | Venturia inaequalis |
| TRINITY_DN11592_c0_g2 | Rola-high | -3.69 | 1.48e-04 | MD12G1252600 | -0.38 | 8.42e-01 | yes | GDSL esterase/lipase LTL1-like | — |
| TRINITY_DN3925_c0_g1 | Rin-high | 3.68 | 1.12e-04 | MD11G1246900 | -0.60 | 2.59e-01 | | cytochrome c oxidase copper chaperone 2 | — |
| TRINITY_DN143378_c0_g1 | Rola-high | -3.67 | 1.06e-03 | MD07G1226100 | 0.12 | 5.04e-01 | | eukaryotic translation initiation factor 2 subunit alpha homolog | Venturia inaequalis |
| TRINITY_DN33400_c0_g1 | Rola-high | -3.63 | 4.41e-04 | MD01G1134600 | 0.52 | 1.60e-02 | | triosephosphate isomerase, cytosolic-like | Venturia inaequalis |
| TRINITY_DN16661_c0_g1 | Rola-high | -3.63 | 9.88e-03 | MD15G1195500 | -0.59 | 7.62e-03 | | NADP-dependent malic enzyme | Venturia inaequalis |
| TRINITY_DN22673_c0_g1 | Rola-high | -3.63 | 1.33e-06 | MD02G1012800 | -3.07 | 1.72e-01 | | hypothetical protein DVH24_017651 | — |
| TRINITY_DN178668_c0_g1 | Rola-high | -3.63 | 8.15e-03 | MD15G1067300 | -0.20 | 1.75e-01 | yes | calcium-transporting ATPase 1, endoplasmic reticulum-type-like | Venturia inaequalis |
| TRINITY_DN62072_c0_g1 | Rola-high | -3.62 | 1.82e-03 | MD10G1270900 | NA | NA | yes | GDSL esterase/lipase At1g29660-like | — |
| TRINITY_DN102273_c0_g1 | Rola-high | -3.59 | 4.78e-05 | MD17G1068900 | -0.56 | 4.89e-04 | | 6-phosphogluconate dehydrogenase, decarboxylating 2 | Venturia inaequalis |
| TRINITY_DN48764_c1_g1 | Rola-high | -3.59 | 3.12e-04 | MD03G1103900 | 0.44 | 1.95e-01 | | elongation factor 1-alpha | Venturia inaequalis |
| TRINITY_DN34461_c0_g1 | Rin-high | 3.58 | 9.81e-03 | MD02G1319100 | -0.70 | 9.63e-03 | | callose synthase 3 | — |
| TRINITY_DN223469_c0_g1 | Rola-high | -3.58 | 5.39e-03 | MD07G1067400 | -0.34 | 2.28e-02 | | cell division control protein 2 homolog | Venturia inaequalis |
| TRINITY_DN19277_c0_g1 | Rola-high | -3.57 | 3.61e-03 | MD01G1194600 | -1.72 | 3.59e-02 | | sorbitol dehydrogenase-like | Venturia inaequalis |
| TRINITY_DN67942_c0_g1 | Rola-high | -3.57 | 1.02e-06 | MD07G1005900 | 0.22 | 4.36e-01 | | elongation factor 1-delta-like | Venturia inaequalis |
| TRINITY_DN52116_c0_g2 | Rola-high | -3.56 | 2.39e-03 | MD06G1239800 | 0.45 | 4.84e-03 | | succinate--CoA ligase [ADP-forming] subunit alpha-2, mitochondrial | Venturia inaequalis |
| TRINITY_DN52846_c1_g1 | Rola-high | -3.55 | 2.88e-03 | MD14G1134100 | -0.18 | 5.72e-01 | | luminal-binding protein 5-like | Venturia inaequalis |
| TRINITY_DN23476_c0_g1 | Rola-high | -3.55 | 7.94e-03 | MD06G1108600 | 0.42 | 1.19e-02 | | UDP-glucose 4-epimerase GEPI48 | Venturia inaequalis |
| TRINITY_DN1873_c0_g3 | Rola-high | -3.53 | 1.52e-04 | MD13G1065000 | 0.09 | 8.33e-01 | yes | receptor-like protein kinase | — |

| Trinity gene ID | Direction | Trinity | | GDDH13 protein match | | | Def. | Description | FCS-GX |
|---|---|---|---|---|---|---|---|---|---|
| | | log2FC | padj | gene ID | log2FC | padj | | | call |
| TRINITY_DN9106_c0_g1 | Rin-high | 3.53 | 6.60e-03 | MD15G1232800 | 0.26 | 3.01e-02 | | dehydrodolichyl diphosphate synthase CPT3-like | — |
| TRINITY_DN18475_c0_g1 | Rola-high | -3.53 | 1.40e-03 | MD04G1096100 | 0.58 | 2.15e-02 | | isocitrate dehydrogenase [NAD] catalytic subunit 5, mitochondrial-like | Venturia inaequalis |
| TRINITY_DN3760_c2_g1 | Rola-high | -3.53 | 7.77e-03 | MD04G1242700 | 0.66 | 6.01e-01 | | DNA damage-repair/toleration protein DRT100-like | — |
| TRINITY_DN118465_c0_g1 | Rola-high | -3.52 | 4.54e-04 | MD12G1125600 | 0.41 | 5.65e-01 | yes | L-ascorbate peroxidase 2, cytosolic | Venturia inaequalis |
| TRINITY_DN156870_c0_g1 | Rola-high | -3.51 | 4.57e-03 | MD05G1234200 | 0.13 | 7.14e-01 | yes | phosphoglycerate kinase 3, cytosolic | Venturia inaequalis |
| TRINITY_DN119993_c0_g1 | Rola-high | -3.49 | 4.91e-03 | MD17G1264300 | -0.01 | 9.83e-01 | | TCP domain class transcription factor | Venturia inaequalis |
| TRINITY_DN118015_c0_g1 | Rola-high | -3.48 | 3.95e-03 | MD17G1104900 | 0.19 | 3.19e-01 | | T-complex protein 1 subunit epsilon | Venturia inaequalis |
| TRINITY_DN11625_c1_g2 | Rola-high | -3.48 | 8.93e-03 | MD15G1411500 | -0.61 | 3.15e-03 | | hypothetical protein DVH24_016491 | — |
| TRINITY_DN103818_c0_g1 | Rola-high | -3.48 | 2.57e-03 | MD08G1206800 | -0.09 | 7.93e-01 | | serine hydroxymethyltransferase, mitochondrial | Venturia inaequalis |
| TRINITY_DN52100_c0_g1 | Rola-high | -3.48 | 1.02e-03 | MD04G1220300 | 0.04 | 8.40e-01 | | small ribosomal subunit protein eS24z | Venturia inaequalis |
| TRINITY_DN4701_c1_g1 | Rola-high | -3.46 | 3.67e-04 | MD03G1066600 | -1.74 | 1.67e-02 | | gibberellin 2-beta-dioxygenase 8-like | — |
| TRINITY_DN22544_c0_g1 | Rola-high | -3.44 | 4.46e-03 | MD15G1386600 | 0.02 | 9.10e-01 | | adenylosuccinate synthetase 2, chloroplastic-like | Venturia inaequalis |
| TRINITY_DN7883_c0_g1 | Rin-high | 3.41 | 1.88e-03 | MD03G1114700 | 3.16 | 2.68e-02 | | fasciclin-like arabinogalactan protein 12 | — |
| TRINITY_DN38675_c0_g1 | Rola-high | -3.41 | 2.17e-04 | MD13G1094800 | 0.18 | 5.09e-01 | | ubiquitin-ribosomal protein eS31 fusion protein | Venturia inaequalis |
| TRINITY_DN23851_c0_g1 | Rola-high | -3.40 | 3.05e-03 | MD17G1264900 | -0.83 | 1.13e-02 | | aspartate aminotransferase, cytoplasmic | Venturia inaequalis |
| TRINITY_DN58451_c0_g1 | Rola-high | -3.40 | 4.86e-04 | MD00G1033600 | 0.55 | 3.16e-02 | | ABC transporter F family member 1-like | Venturia inaequalis |
| TRINITY_DN185056_c0_g1 | Rola-high | -3.39 | 5.29e-03 | MD13G1248200 | 0.15 | 3.51e-01 | | lysine--tRNA ligase, chloroplastic/mitochondrial-like | Venturia inaequalis |
| TRINITY_DN9231_c0_g1 | Rin-high | 3.38 | 2.19e-03 | MD15G1434100 | 0.41 | 2.16e-02 | | hypothetical protein DVH24_033146 | — |
| TRINITY_DN21392_c0_g1 | Rola-high | -3.37 | 2.21e-04 | MD00G1140200 | -0.06 | 8.16e-01 | | fumarate hydratase 1, mitochondrial | Venturia inaequalis |

| Trinity gene ID | Direction | Trinity | | GDDH13 protein match | | | Def. | Description | FCS-GX |
|---|---|---|---|---|---|---|---|---|---|
| | | log2FC | padj | gene ID | log2FC | padj | | | call |
| TRINITY_DN67259_c1_g1 | Rola-high | -3.37 | 6.73e-05 | MD10G1058400 | 0.37 | 1.40e-02 | | soluble inorganic pyrophosphatase 6, chloroplastic-like | Venturia inaequalis |
| TRINITY_DN223483_c0_g1 | Rola-high | -3.36 | 6.62e-04 | MD00G1076300 | -0.39 | 3.13e-01 | | calnexin homolog | Venturia inaequalis |
| TRINITY_DN39377_c0_g1 | Rola-high | -3.36 | 5.33e-04 | MD15G1120100 | -0.03 | 9.28e-01 | | GTP-binding nuclear protein Ran-3-like | Venturia inaequalis |
| TRINITY_DN184165_c0_g1 | Rola-high | -3.35 | 1.23e-03 | MD13G1084800 | 0.17 | 2.03e-01 | | pyruvate dehydrogenase E1 component subunit alpha, mitochondrial | Venturia inaequalis |
| TRINITY_DN22662_c2_g1 | Rin-high | 3.35 | 7.79e-05 | MD03G1008200 | -0.45 | 1.57e-01 | | putative reverse transcriptase family member | — |
| TRINITY_DN90120_c0_g1 | Rola-high | -3.35 | 5.40e-03 | MD08G1002200 | 0.10 | 6.85e-01 | | probable U6 snRNA-associated Sm-like protein LSm4 | Venturia inaequalis |
| TRINITY_DN201751_c0_g1 | Rola-high | -3.34 | 2.56e-05 | MD00G1070400 | 0.27 | 6.21e-02 | | ABC transporter F family member 4-like isoform X2 | Venturia inaequalis |
| TRINITY_DN18787_c0_g2 | Rin-high | 3.33 | 2.55e-03 | MD05G1260900 | 0.55 | 1.51e-02 | yes | probable LRR receptor-like serine/threonine-protein kinase At1g56130 | — |
| TRINITY_DN8160_c0_g1 | Rola-high | -3.32 | 1.88e-03 | MD10G1143600 | -2.88 | 1.64e-02 | | AT-hook motif nuclear-localized protein 25-like | — |
| TRINITY_DN30708_c0_g1 | Rola-high | -3.31 | 3.00e-03 | MD09G1128400 | 0.08 | 8.92e-01 | | acetyl-CoA acetyltransferase 1 isoform X2 | Venturia inaequalis |
| TRINITY_DN8921_c1_g1 | Rin-high | 3.31 | 2.47e-03 | MD06G1010300 | -0.49 | 5.96e-01 | | uncharacterized protein | — |
| TRINITY_DN23515_c0_g1 | Rola-high | -3.31 | 2.61e-03 | MD16G1130800 | 0.73 | 7.29e-04 | | eukaryotic peptide chain release factor subunit 1-3-like | Venturia inaequalis |
| TRINITY_DN24613_c0_g1 | Rola-high | -3.30 | 7.33e-04 | MD17G1006600 | 0.14 | 2.77e-01 | | cytochrome b-c1 complex subunit Rieske-4, mitochondrial-like | Venturia inaequalis |
| TRINITY_DN49095_c0_g2 | Rola-high | -3.30 | 1.99e-03 | MD03G1292100 | -0.26 | 2.22e-01 | | large ribosomal subunit protein eL19x | Venturia inaequalis |
| TRINITY_DN11702_c1_g1 | Rin-high | 3.27 | 5.02e-03 | MD08G1205700 | 0.53 | 3.96e-01 | | uncharacterized protein | — |
| TRINITY_DN80043_c0_g1 | Rola-high | -3.26 | 9.86e-03 | MD02G1207200 | 0.86 | 5.93e-05 | | transketolase, chloroplastic | Venturia inaequalis |
| TRINITY_DN59080_c0_g1 | Rola-high | -3.25 | 7.02e-03 | MD01G1108900 | -0.32 | 7.32e-02 | | chorismate synthase, chloroplastic-like | Venturia inaequalis |
| TRINITY_DN115555_c0_g1 | Rola-high | -3.22 | 2.61e-03 | MD11G1073900 | -0.09 | 5.25e-01 | | peptidyl-prolyl cis-trans isomerase CYP19-4 | Venturia inaequalis |
| TRINITY_DN95418_c0_g1 | Rola-high | -3.21 | 3.87e-03 | MD02G1008000 | 0.10 | 5.33e-01 | | eukaryotic translation initiation factor 5A-2-like | Venturia inaequalis |

| Trinity gene ID | Direction | Trinity | | GDDH13 protein match | | | Def. | Description | FCS-GX |
|---|---|---|---|---|---|---|---|---|---|
| | | log2FC | padj | gene ID | log2FC | padj | | | call |
| TRINITY_DN113537_c0_g1 | Rola-high | -3.21 | 5.32e-03 | MD06G1047900 | 0.06 | 8.35e-01 | | T-complex protein 1 subunit alpha | Venturia inaequalis |
| TRINITY_DN37919_c0_g1 | Rola-high | -3.19 | 9.91e-03 | MD09G1192000 | -0.87 | 1.66e-02 | | nucleosome assembly protein 1-1 | Venturia inaequalis |
| TRINITY_DN9968_c1_g3 | Rola-high | -3.14 | 4.19e-03 | MD04G1159700 | -0.64 | 4.17e-01 | yes | receptor-like protein 33 isoform X2 | — |
| TRINITY_DN27316_c0_g1 | Rola-high | -3.12 | 5.18e-03 | MD13G1123100 | 0.18 | 4.54e-01 | | photosynthetic NDH subunit of lumenal location 5, chloroplastic | Venturia inaequalis |
| TRINITY_DN18925_c0_g1 | Rola-high | -3.11 | 6.97e-03 | MD06G1087700 | -0.04 | 8.39e-01 | | serine/arginine-rich splicing factor SR45a-like isoform X1 | Venturia inaequalis |
| TRINITY_DN13576_c2_g1 | Rola-high | -3.09 | 7.92e-04 | MD05G1161500 | NA | NA | | AUGMIN subunit 3-like | — |
| TRINITY_DN17239_c0_g1 | Rola-high | -3.05 | 7.32e-03 | MD11G1280500 | 0.37 | 2.02e-01 | | biotin carboxylase 1, chloroplastic | Venturia inaequalis |
| TRINITY_DN202224_c0_g1 | Rola-high | -3.05 | 2.21e-03 | MD11G1259400 | 0.08 | 8.31e-01 | | malate dehydrogenase, glyoxysomal-like | Venturia inaequalis |
| TRINITY_DN7705_c4_g1 | Rola-high | -2.98 | 9.09e-05 | MD00G1144800 | 0.22 | 6.48e-01 | | uncharacterized protein | — |
| TRINITY_DN160677_c0_g1 | Rola-high | -2.97 | 6.19e-03 | MD01G1140100 | 1.79 | 1.69e-02 | | uncharacterized protein At2g39795, mitochondrial-like isoform X1 | Venturia inaequalis |
| TRINITY_DN43871_c0_g1 | Rola-high | -2.95 | 1.16e-03 | MD07G1309200 | -0.35 | 4.30e-01 | | cellulose synthase-like protein D3 | — |
| TRINITY_DN3563_c3_g1 | Rola-high | -2.94 | 2.90e-03 | MD06G1145200 | 0.53 | 1.67e-01 | | nucleolar protein 56-like | Venturia inaequalis |
| TRINITY_DN44846_c0_g1 | Rin-high | 2.92 | 6.71e-06 | MD16G1101600 | 0.85 | 6.16e-01 | | protein DOWNY MILDEW RESISTANCE 6-like | — |
| TRINITY_DN4353_c0_g2 | Rola-high | -2.90 | 5.98e-03 | MD13G1088500 | -0.28 | 2.67e-02 | | WAT1-related protein At3g28070-like isoform X2 | — |
| TRINITY_DN17221_c0_g1 | Rola-high | -2.86 | 7.86e-03 | MD09G1259500 | 0.06 | 8.16e-01 | | eukaryotic translation initiation factor 2 subunit beta | Venturia inaequalis |
| TRINITY_DN13881_c0_g1 | Rola-high | -2.83 | 4.64e-05 | MD16G1024900 | -2.11 | NA | yes | MLP-like protein 328 | — |
| TRINITY_DN7829_c0_g3 | Rola-high | -2.77 | 3.13e-03 | MD17G1017300 | 0.06 | 8.17e-01 | | glyceraldehyde-3-phosphate dehydrogenase, cytosolic-like | — |
| TRINITY_DN22546_c0_g1 | Rola-high | -2.76 | 3.04e-03 | MD11G1131300 | 0.60 | 3.00e-04 | | elongation factor 1-alpha | Penicillium brevicompactum |
| TRINITY_DN38741_c0_g1 | Rola-high | -2.74 | 7.56e-03 | MD16G1163400 | NA | NA | | hypothetical protein DVH24_026282 | — |
| TRINITY_DN5584_c0_g1 | Rin-high | 2.71 | 4.18e-03 | MD08G1179200 | 0.55 | 6.70e-01 | yes | probable LRR receptor-like serine/threonine-protein kinase At1g56130 | — |

| Trinity gene ID | Direction | Trinity | | GDDH13 protein match | | | Def. | Description | FCS-GX call |
|---|---|---|---|---|---|---|---|---|---|
| | | log2FC | padj | gene ID | log2FC | padj | | | |
| TRINITY_DN3459_c0_g1 | Rola-high | -2.67 | 4.44e-03 | MD11G1210800 | -0.65 | 6.56e-02 | | glucan endo-1,3-beta-glucosidase 14 | — |
| TRINITY_DN23082_c0_g1 | Rola-high | -2.64 | 1.70e-03 | MD09G1014600 | -0.72 | 5.53e-02 | yes | probable inactive leucine-rich repeat receptor-like protein kinase At3g03770 | — |
| TRINITY_DN1842_c2_g1 | Rin-high | 2.60 | 3.23e-03 | MD14G1171300 | 0.88 | 2.71e-03 | | uncharacterized protein | — |
| TRINITY_DN7984_c0_g1 | Rin-high | 2.59 | 2.99e-03 | MD10G1251400 | 1.33 | 1.85e-02 | yes | putative wall-associated receptor kinase-like 16 | — |
| TRINITY_DN11845_c0_g1 | Rola-high | -2.59 | 8.69e-03 | MD13G1212600 | -1.79 | 4.24e-02 | | RING-H2 finger protein ATL16-like | — |
| TRINITY_DN16781_c0_g1 | Rola-high | -2.58 | 6.43e-03 | MD04G1183700 | -3.73 | 2.73e-02 | | hypothetical protein DVH24_015014 | — |
| TRINITY_DN35492_c0_g1 | Rin-high | 2.55 | 5.97e-03 | MD10G1251400 | 1.33 | 1.85e-02 | yes | putative wall-associated receptor kinase-like 16 | — |
| TRINITY_DN4982_c0_g1 | Rola-high | -2.54 | 1.85e-03 | MD17G1007800 | 0.04 | 9.34e-01 | | pentatricopeptide repeat-containing protein At3g26782, mitochondrial-like | — |
| TRINITY_DN575_c0_g1 | Rola-high | -2.33 | 8.21e-03 | MD11G1017400 | -0.51 | 8.19e-02 | | enoyl-ACP reductase | — |
| TRINITY_DN171_c1_g1 | Rin-high | 2.32 | 4.48e-03 | MD05G1013200 | -0.40 | 5.06e-01 | | alcohol dehydrogenase | — |
| TRINITY_DN3235_c0_g1 | Rin-high | 2.25 | 7.87e-03 | MD01G1230100 | 0.91 | 2.42e-02 | yes | uncharacterized calcium-binding protein At1g02270-like | — |
| TRINITY_DN7981_c0_g1 | Rola-high | -2.24 | 5.69e-03 | MD08G1229500 | 1.31 | 2.15e-01 | | replication protein A 70 kDa DNA-binding subunit E-like isoform X1 | — |
| TRINITY_DN1874_c1_g1 | Rin-high | 2.19 | 3.55e-04 | MD08G1197300 | 0.92 | 9.75e-04 | | MADS-box protein AGL24 | — |
| TRINITY_DN25037_c0_g1 | Rin-high | 2.14 | 8.62e-03 | MD10G1251400 | 1.33 | 1.85e-02 | yes | putative wall-associated receptor kinase-like 16 | — |
| TRINITY_DN23459_c1_g3 | Rola-high | -2.14 | 1.10e-03 | MD11G1219600 | 0.52 | 3.40e-02 | | polyadenylate-binding protein RBP45C-like | — |
| TRINITY_DN931_c0_g1 | Rola-high | -2.11 | 9.50e-03 | MD15G1315000 | 0.61 | 9.00e-02 | | mitochondrial uncoupling protein 5-like | — |
| TRINITY_DN8783_c0_g1 | Rola-high | -1.99 | 1.08e-05 | MD06G1015400 | -0.16 | 2.41e-01 | yes | cytochrome P450 77A3-like | — |
| **GDDH13 hit present but below the match-confidence threshold (fragmentary or divergent) (n = 274)** | | | | | | | | | |
| TRINITY_DN4249_c0_g2 | Rola-high | -25.88 | 7.99e-30 | NA | NA | NA | | protein GAST1 isoform X2 | — |
| TRINITY_DN68535_c0_g1 | Rin-high | 8.28 | 1.07e-05 | NA | NA | NA | | two-component response regulator-like APRR5 | — |

| Trinity gene ID | Direction | Trinity | | GDDH13 protein match | | | Def. | Description | FCS-GX |
|---|---|---|---|---|---|---|---|---|---|
| | | log2FC | padj | gene ID | log2FC | padj | | | call |
| TRINITY_DN101312_c0_g1 | Rin-high | 8.15 | 5.14e-06 | NA | NA | NA | yes | probable xyloglucan endotransglucosylase/hydrolase protein 6 precursor | — |
| TRINITY_DN20127_c0_g1 | Rola-high | -6.99 | 3.42e-04 | NA | NA | NA | | low affinity inorganic phosphate transporter 1-like | Venturia inaequalis |
| TRINITY_DN5588_c0_g1 | Rola-high | -6.87 | 2.91e-10 | NA | NA | NA | | glycine-rich cell wall structural protein | — |
| TRINITY_DN201753_c0_g1 | Rola-high | -6.81 | 6.82e-07 | NA | NA | NA | | alanine--glyoxylate aminotransferase 2 homolog 2, mitochondrial-like | Venturia inaequalis |
| TRINITY_DN1391_c1_g2 | Rin-high | 6.62 | 3.82e-03 | NA | NA | NA | | hypothetical protein DVH24_017063 | — |
| TRINITY_DN5530_c0_g1 | Rola-high | -6.62 | 6.12e-06 | NA | NA | NA | | glycine-rich cell wall structural protein 1-like | — |
| TRINITY_DN4376_c2_g1 | Rin-high | 6.43 | 1.73e-03 | NA | NA | NA | | uncharacterized protein | — |
| TRINITY_DN50014_c0_g2 | Rola-high | -6.40 | 2.10e-03 | NA | NA | NA | | hypothetical protein DVH24_001586 | — |
| TRINITY_DN222698_c0_g1 | Rola-high | -6.28 | 3.37e-06 | NA | NA | NA | | hypothetical protein CRYUN_Cryun02cG0058300 | Venturia inaequalis |
| TRINITY_DN178681_c0_g1 | Rola-high | -6.23 | 1.78e-07 | NA | NA | NA | | alpha-L-fucosidase 2-like isoform X1 | Venturia inaequalis |
| TRINITY_DN92343_c0_g1 | Rola-high | -6.19 | 4.89e-04 | NA | NA | NA | | zinc-finger homeodomain protein 2-like | — |
| TRINITY_DN25028_c0_g3 | Rin-high | 6.19 | 1.45e-03 | NA | NA | NA | yes | G-type lectin S-receptor-like serine/threonine-protein kinase At4g27290 isoform X2 | — |
| TRINITY_DN10914_c1_g1 | Rola-high | -6.19 | 9.45e-03 | NA | NA | NA | | palmitoyl-monogalactosyldiacylglycerol delta-7 desaturase, chloroplastic-like | — |
| TRINITY_DN203138_c0_g1 | Rola-high | -6.11 | 6.48e-11 | NA | NA | NA | | uncharacterized protein | Venturia inaequalis |
| TRINITY_DN4211_c2_g1 | Rin-high | 6.08 | 1.39e-04 | NA | NA | NA | | MADS-box transcription factor 14-like | — |
| TRINITY_DN40621_c0_g1 | Rola-high | -6.01 | 4.40e-03 | NA | NA | NA | | unnamed protein product | Venturia inaequalis |
| TRINITY_DN163652_c0_g1 | Rola-high | -5.99 | 1.69e-06 | NA | NA | NA | | probable carboxylesterase 6 | Venturia inaequalis |
| TRINITY_DN24373_c0_g1 | Rola-high | -5.88 | 1.65e-04 | NA | NA | NA | | pyrroline-5-carboxylate reductase-like | Venturia inaequalis |
| TRINITY_DN12070_c0_g1 | Rin-high | 5.86 | 1.03e-05 | NA | NA | NA | | hypothetical protein DVH24_015128 | — |

| Trinity gene ID | Direction | Trinity | | GDDH13 protein match | | | Def. | Description | FCS-GX |
|---|---|---|---|---|---|---|---|---|---|
| | | log2FC | padj | gene ID | log2FC | padj | | | call |
| TRINITY_DN11659_c2_g1 | Rola-high | -5.85 | 1.28e-06 | NA | NA | NA | | hypothetical protein DVH24_007882 | — |
| TRINITY_DN42178_c0_g1 | Rin-high | 5.83 | 7.49e-05 | NA | NA | NA | | cold-regulated protein 27-like isoform X1 | — |
| TRINITY_DN6774_c0_g1 | Rola-high | -5.83 | 7.94e-08 | NA | NA | NA | | Prothoracicostatic peptide | — |
| TRINITY_DN54389_c0_g1 | Rola-high | -5.82 | 2.10e-06 | NA | NA | NA | | hypothetical protein DVH24_039153 | — |
| TRINITY_DN168270_c0_g1 | Rola-high | -5.67 | 3.70e-04 | NA | NA | NA | yes | expansin-A1-like | — |
| TRINITY_DN30195_c0_g1 | Rola-high | -5.65 | 2.99e-03 | NA | NA | NA | | exopolygalacturonase clone GBGA483-like | Venturia inaequalis |
| TRINITY_DN51515_c0_g1 | Rin-high | 5.65 | 3.24e-06 | NA | NA | NA | yes | probable LRR receptor-like serine/threonine-protein kinase At3g47570 isoform X3 | — |
| TRINITY_DN157273_c0_g1 | Rola-high | -5.63 | 4.88e-07 | NA | NA | NA | | uncharacterized protein | Venturia inaequalis |
| TRINITY_DN23701_c0_g1 | Rola-high | -5.62 | 2.28e-03 | NA | NA | NA | yes | cytochrome P450 704C1-like | Venturia inaequalis |
| TRINITY_DN109338_c2_g1 | Rin-high | 5.57 | 3.59e-04 | NA | NA | NA | | uncharacterized protein | — |
| TRINITY_DN8040_c5_g1 | Rola-high | -5.56 | 1.16e-05 | NA | NA | NA | | hypothetical protein DVH24_022237 | — |
| TRINITY_DN9439_c0_g1 | Rola-high | -5.49 | 2.82e-04 | NA | NA | NA | | ferric reduction oxidase 7, chloroplastic-like | Venturia inaequalis |
| TRINITY_DN109338_c0_g1 | Rin-high | 5.44 | 2.60e-03 | NA | NA | NA | | uncharacterized protein | — |
| TRINITY_DN29417_c0_g1 | Rola-high | -5.43 | 7.52e-04 | NA | NA | NA | | ribosome biogenesis regulatory protein homolog | Venturia inaequalis |
| TRINITY_DN107095_c1_g1 | Rin-high | 5.38 | 1.81e-03 | NA | NA | NA | | uncharacterized protein | — |
| TRINITY_DN15618_c0_g1 | Rola-high | -5.35 | 1.39e-06 | NA | NA | NA | | nicotianamine synthase-like | Venturia inaequalis |
| TRINITY_DN52542_c0_g1 | Rola-high | -5.35 | 2.40e-07 | NA | NA | NA | | uncharacterized protein | Venturia inaequalis |
| TRINITY_DN25564_c0_g1 | Rola-high | -5.33 | 3.16e-03 | NA | NA | NA | | uncharacterized protein | Venturia inaequalis |
| TRINITY_DN160541_c0_g1 | Rola-high | -5.29 | 9.65e-05 | NA | NA | NA | | delta(24)-sterol reductase | Venturia inaequalis |
| TRINITY_DN87194_c0_g1 | Rola-high | -5.27 | 1.03e-04 | NA | NA | NA | | extensin isoform X1 | — |

| Trinity gene ID | Direction | Trinity | | GDDH13 protein match | | | Def. | Description | FCS-GX |
|---|---|---|---|---|---|---|---|---|---|
| | | log2FC | padj | gene ID | log2FC | padj | | | call |
| TRINITY_DN35790_c0_g1 | Rola-high | -5.26 | 1.75e-04 | NA | NA | NA | | adenylosuccinate lyase | Venturia inaequalis |
| TRINITY_DN181442_c0_g1 | Rola-high | -5.21 | 1.82e-08 | NA | NA | NA | | clustered mitochondria protein-like isoform X1 | Venturia inaequalis |
| TRINITY_DN64541_c0_g2 | Rola-high | -5.19 | 6.73e-05 | NA | NA | NA | | uncharacterized protein | Venturia inaequalis |
| TRINITY_DN12515_c0_g1 | Rola-high | -5.19 | 4.60e-05 | NA | NA | NA | | protein tesmin/TSO1-like CXC 2 | — |
| TRINITY_DN24687_c0_g1 | Rola-high | -5.17 | 3.54e-06 | NA | NA | NA | | heat shock 70 kDa protein | Venturia inaequalis |
| TRINITY_DN222868_c0_g1 | Rola-high | -5.16 | 2.09e-04 | NA | NA | NA | | protein LATERAL BRANCHING OXIDOREDUCTASE 1-like | Venturia inaequalis |
| TRINITY_DN228582_c0_g1 | Rola-high | -5.14 | 8.63e-07 | NA | NA | NA | | phenylacetaldehyde synthase | Venturia inaequalis |
| TRINITY_DN223455_c0_g1 | Rola-high | -5.13 | 4.52e-03 | NA | NA | NA | | probable inorganic phosphate transporter 1-7 isoform X1 | Venturia inaequalis |
| TRINITY_DN35593_c0_g2 | Rola-high | -5.12 | 2.57e-03 | NA | NA | NA | | growth-regulating factor 1-like | — |
| TRINITY_DN20639_c0_g1 | Rin-high | 5.09 | 5.70e-03 | NA | NA | NA | | 7-deoxyloganetin glucosyltransferase-like | — |
| TRINITY_DN209907_c0_g1 | Rola-high | -5.09 | 9.53e-04 | NA | NA | NA | | gamma-glutamyl peptidase 5-like | Venturia inaequalis |
| TRINITY_DN28642_c0_g1 | Rola-high | -5.07 | 2.23e-03 | NA | NA | NA | | uncharacterized protein | Venturia inaequalis |
| TRINITY_DN98586_c0_g1 | Rola-high | -5.02 | 2.21e-03 | NA | NA | NA | | UDP-galactose/UDP-glucose transporter 3 | Venturia inaequalis |
| TRINITY_DN48088_c0_g1 | Rola-high | -5.01 | 1.51e-03 | NA | NA | NA | | ureidoglycolate hydrolase | Venturia inaequalis |
| TRINITY_DN55342_c0_g1 | Rin-high | 5.01 | 5.51e-03 | NA | NA | NA | | probable mannitol dehydrogenase | Cystobasidiaceae sp. HBUAS51001 |
| TRINITY_DN42071_c0_g1 | Rin-high | 4.99 | 7.84e-03 | NA | NA | NA | | zinc finger protein CONSTANS-LIKE 9-like | — |
| TRINITY_DN6295_c0_g2 | Rin-high | 4.98 | 1.74e-04 | NA | NA | NA | | hypothetical protein DVH24_004049 | — |
| TRINITY_DN136870_c0_g1 | Rola-high | -4.95 | 1.23e-03 | NA | NA | NA | | bifunctional 3-dehydroquinate dehydratase/shikimate dehydrogenase, chloroplastic-like | Venturia inaequalis |

| Trinity gene ID | Direction | Trinity | | GDDH13 protein match | | | Def. | Description | FCS-GX |
|---|---|---|---|---|---|---|---|---|---|
| | | log2FC | padj | gene ID | log2FC | padj | | | call |
| TRINITY_DN27178_c0_g1 | Rin-high | 4.90 | 6.92e-03 | NA | NA | NA | yes | probable LRR receptor-like serine/threonine-protein kinase At1g56130 | — |
| TRINITY_DN25271_c0_g1 | Rola-high | -4.90 | 3.16e-03 | NA | NA | NA | | uncharacterized protein | Venturia inaequalis |
| TRINITY_DN28859_c0_g1 | Rola-high | -4.88 | 4.87e-06 | NA | NA | NA | | eukaryotic translation initiation factor 3 subunit F-like | Venturia inaequalis |
| TRINITY_DN7030_c1_g1 | Rola-high | -4.87 | 3.31e-03 | NA | NA | NA | | hypothetical protein DVH24_025358 | — |
| TRINITY_DN20883_c0_g1 | Rola-high | -4.87 | 9.06e-04 | NA | NA | NA | | fatty acid amide hydrolase | Venturia inaequalis |
| TRINITY_DN13955_c0_g1 | Rin-high | 4.84 | 4.60e-03 | NA | NA | NA | yes | probable pectinesterase/pectinesterase inhibitor 35 | — |
| TRINITY_DN23879_c2_g2 | Rola-high | -4.84 | 1.24e-04 | NA | NA | NA | | probable histone H2B.1 | — |
| TRINITY_DN6378_c0_g1 | Rin-high | 4.84 | 2.65e-03 | NA | NA | NA | | protein VERNALIZATION 1 | — |
| TRINITY_DN56817_c0_g1 | Rola-high | -4.82 | 2.47e-03 | NA | NA | NA | | pumilio homolog 24 | Venturia inaequalis |
| TRINITY_DN12220_c0_g1 | Rola-high | -4.79 | 1.16e-06 | NA | NA | NA | | heavy metal-associated isoprenylated plant protein 32-like | — |
| TRINITY_DN21077_c0_g1 | Rola-high | -4.78 | 2.14e-03 | NA | NA | NA | | gamma-aminobutyrate transaminase 1, mitochondrial-like isoform X1 | Venturia inaequalis |
| TRINITY_DN225048_c0_g1 | Rola-high | -4.77 | 1.16e-03 | NA | NA | NA | | folylpolyglutamate synthase-like | Venturia inaequalis |
| TRINITY_DN26748_c0_g1 | Rola-high | -4.76 | 5.81e-03 | NA | NA | NA | | probable inactive purple acid phosphatase 16 | Venturia inaequalis |
| TRINITY_DN30519_c0_g1 | Rola-high | -4.73 | 1.52e-03 | NA | NA | NA | | nucleolar complex-associated protein 2 | Venturia inaequalis |
| TRINITY_DN222072_c0_g1 | Rola-high | -4.71 | 3.35e-04 | NA | NA | NA | | amidase 1-like | Venturia inaequalis |
| TRINITY_DN1437_c5_g1 | Rin-high | 4.71 | 4.46e-03 | NA | NA | NA | | uncharacterized protein | — |
| TRINITY_DN36105_c0_g1 | Rola-high | -4.68 | 8.53e-05 | NA | NA | NA | | probable aldo-keto reductase 1 | Venturia inaequalis |
| TRINITY_DN26402_c0_g1 | Rola-high | -4.66 | 8.64e-03 | NA | NA | NA | | hypothetical protein BT93_L0645 | Venturia inaequalis |
| TRINITY_DN39701_c0_g1 | Rola-high | -4.65 | 1.26e-03 | NA | NA | NA | | MFS transporter, ACS family, DAL5 transporter family protein | Venturia inaequalis |

| Trinity gene ID | Direction | Trinity | | GDDH13 protein match | | | Def. | Description | FCS-GX |
|---|---|---|---|---|---|---|---|---|---|
| | | log2FC | padj | gene ID | log2FC | padj | | | call |
| TRINITY_DN107876_c0_g1 | Rola-high | -4.64 | 1.06e-04 | NA | NA | NA | | 3-oxoacyl-[acyl-carrier-protein] reductase 4-like | Venturia inaequalis |
| TRINITY_DN73413_c0_g1 | Rola-high | -4.63 | 8.94e-04 | NA | NA | NA | | mannosyl-oligosaccharide 1,2-alpha-mannosidase MNS1-like | Venturia inaequalis |
| TRINITY_DN51321_c0_g1 | Rola-high | -4.63 | 5.36e-03 | NA | NA | NA | | protein NRT1/ PTR FAMILY 5.4-like | Venturia inaequalis |
| TRINITY_DN23879_c1_g1 | Rola-high | -4.62 | 5.34e-04 | NA | NA | NA | | probable histone H2B.1 | — |
| TRINITY_DN40731_c0_g1 | Rola-high | -4.61 | 1.07e-03 | NA | NA | NA | | L-ascorbate oxidase-like | Venturia inaequalis |
| TRINITY_DN114317_c0_g1 | Rola-high | -4.60 | 4.65e-03 | NA | NA | NA | | NAP1-related protein 2-like | Venturia inaequalis |
| TRINITY_DN1367_c1_g4 | Rola-high | -4.59 | 4.15e-04 | NA | NA | NA | | hypothetical protein DVH24_028176 | — |
| TRINITY_DN28619_c0_g1 | Rola-high | -4.59 | 1.98e-03 | NA | NA | NA | | tRNAse Z TRZ4, mitochondrial-like isoform X2 | Venturia inaequalis |
| TRINITY_DN38374_c0_g1 | Rola-high | -4.59 | 2.08e-03 | NA | NA | NA | | uncharacterized protein | Venturia inaequalis |
| TRINITY_DN4680_c0_g1 | Rin-high | 4.58 | 3.04e-04 | NA | NA | NA | | hypothetical protein DVH24_005801 | — |
| TRINITY_DN25649_c0_g1 | Rin-high | 4.55 | 2.37e-04 | NA | NA | NA | | hypothetical protein DVH24_020741 | — |
| TRINITY_DN203081_c0_g1 | Rola-high | -4.53 | 5.78e-03 | NA | NA | NA | | cystathionine gamma-synthase 1, chloroplastic-like | Venturia inaequalis |
| TRINITY_DN18871_c0_g1 | Rola-high | -4.53 | 6.41e-05 | NA | NA | NA | | large ribosomal subunit protein eL28z-like | Venturia inaequalis |
| TRINITY_DN19787_c0_g1 | Rola-high | -4.52 | 4.51e-04 | NA | NA | NA | yes | glutathione hydrolase 3 | Venturia inaequalis |
| TRINITY_DN28624_c0_g1 | Rola-high | -4.52 | 2.94e-05 | NA | NA | NA | | uncharacterized protein | Venturia inaequalis |
| TRINITY_DN81304_c0_g1 | Rola-high | -4.51 | 4.86e-06 | NA | NA | NA | | mannan endo-1,4-beta-mannosidase 7 | Venturia inaequalis |
| TRINITY_DN46256_c0_g1 | Rola-high | -4.48 | 8.68e-04 | NA | NA | NA | | pescadillo homolog | Venturia inaequalis |
| TRINITY_DN41905_c0_g1 | Rola-high | -4.46 | 3.01e-05 | NA | NA | NA | | alpha-aminoadipic semialdehyde synthase | Venturia inaequalis |
| TRINITY_DN201001_c0_g1 | Rola-high | -4.46 | 6.31e-04 | NA | NA | NA | | protein ILITYHIA-like | Venturia inaequalis |

| Trinity gene ID | Direction | Trinity |  | GDDH13 protein match |  |  | Def. | Description | FCS-GX |
| --- | --- | --- | --- | --- | --- | --- | --- | --- | --- |
|  |  | log2FC | padj | gene ID | log2FC | padj |  |  | call |
| TRINITY_DN26185_c4_g1 | Rola-high | -4.45 | 3.56e-03 | NA | NA | NA |  | hypothetical protein DVH24_000167 | — |
| TRINITY_DN179929_c0_g1 | Rola-high | -4.44 | 7.48e-05 | NA | NA | NA |  | large ribosomal subunit protein eL29z-like | Venturia inaequalis |
| TRINITY_DN32524_c0_g1 | Rola-high | -4.44 | 1.74e-04 | NA | NA | NA | yes | triacylglycerol lipase 2-like | Venturia inaequalis |
| TRINITY_DN238413_c0_g1 | Rin-high | 4.43 | 1.71e-03 | NA | NA | NA |  | protein RADIALIS-like 4 | — |
| TRINITY_DN14840_c0_g1 | Rin-high | 4.42 | 5.31e-04 | NA | NA | NA |  | phytyl ester synthase 1, chloroplastic-like isoform X1 | — |
| TRINITY_DN7427_c0_g1 | Rin-high | 4.41 | 5.51e-05 | NA | NA | NA | yes | probable glutathione S-transferase | — |
| TRINITY_DN18224_c0_g1 | Rola-high | -4.40 | 1.80e-04 | NA | NA | NA |  | mannosyl-oligosaccharide 1,2-alpha-mannosidase MNS1-like | Venturia inaequalis |
| TRINITY_DN48461_c0_g1 | Rola-high | -4.39 | 2.62e-05 | NA | NA | NA |  | adenine phosphoribosyltransferase 1-like | Venturia inaequalis |
| TRINITY_DN158314_c0_g1 | Rola-high | -4.38 | 6.33e-05 | NA | NA | NA |  | la-related protein 1C | Venturia inaequalis |
| TRINITY_DN19020_c0_g1 | Rola-high | -4.37 | 2.61e-07 | NA | NA | NA | yes | putative pectinesterase 11 | Venturia inaequalis |
| TRINITY_DN201479_c0_g1 | Rola-high | -4.37 | 2.51e-03 | NA | NA | NA |  | exocyst complex component SEC15A-like | Venturia inaequalis |
| TRINITY_DN29390_c0_g1 | Rola-high | -4.36 | 2.29e-06 | NA | NA | NA | yes | pectinesterase 2-like | Venturia inaequalis |
| TRINITY_DN30496_c0_g1 | Rola-high | -4.36 | 4.42e-03 | NA | NA | NA |  | peter Pan-like protein isoform X1 | Venturia inaequalis |
| TRINITY_DN61847_c0_g1 | Rin-high | 4.36 | 9.91e-03 | NA | NA | NA |  | uncharacterized protein | — |
| TRINITY_DN2509_c1_g2 | Rin-high | 4.34 | 1.58e-03 | NA | NA | NA | yes | peroxidase A2-like | — |
| TRINITY_DN164377_c0_g1 | Rola-high | -4.32 | 1.10e-03 | NA | NA | NA |  | protein HEAT STRESS TOLERANT DWD 1 | Venturia inaequalis |
| TRINITY_DN161115_c0_g1 | Rola-high | -4.30 | 1.86e-03 | NA | NA | NA |  | uncharacterized protein | Venturia inaequalis |
| TRINITY_DN173305_c0_g1 | Rola-high | -4.30 | 5.52e-03 | NA | NA | NA |  | uncharacterized protein | — |
| TRINITY_DN2139_c0_g1 | Rola-high | -4.29 | 2.30e-03 | NA | NA | NA |  | putative clathrin assembly protein At5g57200 isoform X2 | Venturia inaequalis |
| TRINITY_DN221843_c0_g1 | Rola-high | -4.29 | 7.81e-11 | NA | NA | NA |  | ketol-acid reductoisomerase, chloroplastic | Venturia inaequalis |

| Trinity gene ID | Direction | Trinity | | GDDH13 protein match | | | Def. | Description | FCS-GX |
|---|---|---|---|---|---|---|---|---|---|
| | | log2FC | padj | gene ID | log2FC | padj | | | call |
| TRINITY_DN30370_c0_g2 | Rola-high | -4.29 | 6.63e-03 | NA | NA | NA | | hypothetical protein DVH24_036107 | — |
| TRINITY_DN98466_c0_g1 | Rola-high | -4.29 | 5.50e-05 | NA | NA | NA | | ERBB-3 BINDING PROTEIN 1-like | Venturia inaequalis |
| TRINITY_DN229597_c0_g1 | Rola-high | -4.28 | 8.40e-07 | NA | NA | NA | | large ribosomal subunit protein eL14z-like | Venturia inaequalis |
| TRINITY_DN200656_c0_g1 | Rola-high | -4.27 | 2.70e-03 | NA | NA | NA | | solanesyl diphosphate synthase 3, chloroplastic/mitochondrial-like | Venturia inaequalis |
| TRINITY_DN61170_c0_g1 | Rola-high | -4.26 | 3.40e-04 | NA | NA | NA | | branched-chain-amino-acid aminotransferase 2, chloroplastic-like | Venturia inaequalis |
| TRINITY_DN62_c1_g1 | Rin-high | 4.26 | 3.54e-03 | NA | NA | NA | | uncharacterized protein | — |
| TRINITY_DN14919_c1_g1 | Rin-high | 4.25 | 6.48e-04 | NA | NA | NA | | cucumisin isoform X3 | — |
| TRINITY_DN223149_c0_g1 | Rola-high | -4.24 | 2.56e-03 | NA | NA | NA | | hypothetical protein DVH24_016666 | Venturia inaequalis |
| TRINITY_DN26749_c0_g1 | Rola-high | -4.21 | 6.65e-04 | NA | NA | NA | | ornithine decarboxylase | Venturia inaequalis |
| TRINITY_DN51327_c0_g1 | Rola-high | -4.20 | 1.34e-03 | NA | NA | NA | | unnamed protein product | Venturia inaequalis |
| TRINITY_DN158194_c0_g1 | Rola-high | -4.19 | 1.48e-03 | NA | NA | NA | | very-long-chain 3-oxoacyl-CoA reductase 1-like | Venturia inaequalis |
| TRINITY_DN9811_c1_g2 | Rin-high | 4.18 | 5.23e-03 | NA | NA | NA | | polyprotein (retrotrasposon protein) | — |
| TRINITY_DN34602_c0_g1 | Rola-high | -4.18 | 5.34e-03 | NA | NA | NA | | mechanosensitive ion channel protein 10-like | Venturia inaequalis |
| TRINITY_DN41983_c0_g1 | Rola-high | -4.18 | 4.97e-03 | NA | NA | NA | | amidase 1-like | Venturia inaequalis |
| TRINITY_DN10858_c1_g1 | Rola-high | -4.17 | 4.75e-05 | NA | NA | NA | yes | peroxidase 42-like | — |
| TRINITY_DN2832_c9_g1 | Rin-high | 4.16 | 8.67e-05 | NA | NA | NA | | metallothionein-like protein type 2 | — |
| TRINITY_DN34596_c0_g1 | Rola-high | -4.16 | 8.42e-03 | NA | NA | NA | | EG45-like domain containing protein | Venturia inaequalis |
| TRINITY_DN27869_c0_g2 | Rin-high | 4.16 | 2.45e-03 | NA | NA | NA | | heat shock 70 kDa protein 1 | — |
| TRINITY_DN40324_c0_g1 | Rola-high | -4.12 | 6.35e-06 | NA | NA | NA | | mitochondrial outer membrane protein porin 2-like | Venturia inaequalis |
| TRINITY_DN39352_c0_g1 | Rola-high | -4.12 | 6.30e-04 | NA | NA | NA | | glyoxylate/hydroxypyruvate reductase A HPR2-like | Venturia inaequalis |

| Trinity gene ID | Direction | Trinity |  | GDDH13 protein match |  |  | Def. | Description | FCS-GX call |
|---|---|---|---|---|---|---|---|---|---|
|  |  | log2FC | padj | gene ID | log2FC | padj |  |  |  |
| TRINITY_DN29240_c0_g1 | Rola-high | -4.10 | 5.56e-04 | NA | NA | NA |  | probable N-acetyl-gamma-glutamyl-phosphate reductase, chloroplastic | Venturia inaequalis |
| TRINITY_DN96913_c0_g1 | Rin-high | 4.09 | 1.47e-03 | NA | NA | NA |  | metallothionein-like protein | — |
| TRINITY_DN17546_c0_g2 | Rola-high | -4.08 | 6.97e-03 | NA | NA | NA |  | probable indole-3-pyruvate monooxygenase YUCCA8 | Venturia inaequalis |
| TRINITY_DN181707_c0_g1 | Rola-high | -4.08 | 2.09e-03 | NA | NA | NA | yes | mitochondrial import receptor subunit TOM40-1-like | Venturia inaequalis |
| TRINITY_DN52338_c0_g1 | Rola-high | -4.06 | 2.98e-04 | NA | NA | NA |  | glycine-rich protein 23-like | — |
| TRINITY_DN47857_c0_g1 | Rola-high | -4.05 | 2.25e-05 | NA | NA | NA |  | eukaryotic translation initiation factor 3 subunit C-like | Venturia inaequalis |
| TRINITY_DN37260_c0_g1 | Rola-high | -4.05 | 6.86e-04 | NA | NA | NA |  | isocitrate dehydrogenase [NAD] catalytic subunit 5, mitochondrial-like | Venturia inaequalis |
| TRINITY_DN9875_c0_g1 | Rola-high | -4.05 | 9.60e-03 | NA | NA | NA |  | hypothetical protein DVH24_016800 | — |
| TRINITY_DN23951_c0_g1 | Rola-high | -4.04 | 3.23e-03 | NA | NA | NA |  | ent-copalyl diphosphate synthase 1-like isoform X2 | — |
| TRINITY_DN13255_c0_g1 | Rin-high | 4.03 | 2.43e-09 | NA | NA | NA |  | O-fucosyltransferase 20-like | — |
| TRINITY_DN32612_c0_g1 | Rola-high | -4.03 | 4.24e-04 | NA | NA | NA |  | mitochondrial carnitine/acylcarnitine carrier-like protein | Venturia inaequalis |
| TRINITY_DN5011_c1_g2 | Rin-high | 4.02 | 4.25e-04 | NA | NA | NA |  | uncharacterized protein | — |
| TRINITY_DN160213_c0_g1 | Rola-high | -4.00 | 8.70e-03 | NA | NA | NA |  | ribosomal protein L2 | Venturia inaequalis |
| TRINITY_DN36126_c0_g1 | Rola-high | -3.99 | 1.78e-03 | NA | NA | NA |  | hypothetical protein DVH24_002192 | Venturia inaequalis |
| TRINITY_DN34677_c0_g2 | Rola-high | -3.99 | 4.91e-03 | NA | NA | NA |  | histone H3.2 | — |
| TRINITY_DN31622_c0_g1 | Rola-high | -3.99 | 8.23e-04 | NA | NA | NA |  | deoxyhypusine hydroxylase-B-like | Venturia inaequalis |
| TRINITY_DN31053_c0_g2 | Rola-high | -3.98 | 6.39e-07 | NA | NA | NA |  | ABC transporter F family member 3-like | Venturia inaequalis |
| TRINITY_DN223097_c0_g1 | Rola-high | -3.98 | 6.33e-03 | NA | NA | NA |  | DEAD-box ATP-dependent RNA helicase 31-like | Venturia inaequalis |
| TRINITY_DN26635_c3_g3 | Rola-high | -3.98 | 8.29e-03 | NA | NA | NA | yes | endoglucanase 17-like | — |
| TRINITY_DN8177_c0_g1 | Rola-high | -3.97 | 4.74e-06 | NA | NA | NA |  | 29 kDa ribonucleoprotein A, chloroplastic | Venturia inaequalis |

| Trinity gene ID | Direction | Trinity | | GDDH13 protein match | | | Def. | Description | FCS-GX |
|---|---|---|---|---|---|---|---|---|---|
| | | log2FC | padj | gene ID | log2FC | padj | | | call |
| TRINITY_DN49292_c0_g2 | Rola-high | -3.97 | 8.62e-04 | NA | NA | NA | | small ribosomal subunit protein eS28-like | Venturia inaequalis |
| TRINITY_DN24531_c0_g1 | Rola-high | -3.97 | 4.09e-03 | NA | NA | NA | | asparagine--tRNA ligase, cytoplasmic 1-like | Venturia inaequalis |
| TRINITY_DN117000_c0_g1 | Rola-high | -3.96 | 9.55e-03 | NA | NA | NA | | glucan endo-1,3-beta-glucosidase-like | Venturia inaequalis |
| TRINITY_DN24070_c0_g1 | Rola-high | -3.96 | 8.85e-03 | NA | NA | NA | | prolyl 4-hydroxylase 1 isoform X1 | Venturia inaequalis |
| TRINITY_DN21847_c0_g1 | Rola-high | -3.95 | 1.18e-04 | NA | NA | NA | | eukaryotic translation initiation factor 5A-2 | Venturia inaequalis |
| TRINITY_DN37284_c0_g1 | Rola-high | -3.95 | 4.54e-03 | NA | NA | NA | | 3-hydroxyisobutyryl-CoA hydrolase 1 isoform X1 | Venturia inaequalis |
| TRINITY_DN86_c0_g1 | Rola-high | -3.94 | 2.61e-04 | NA | NA | NA | yes | protein kinase and PP2C-like domain-containing protein isoform X2 | Venturia inaequalis |
| TRINITY_DN20065_c0_g1 | Rola-high | -3.93 | 4.35e-05 | NA | NA | NA | yes | GDSL esterase/lipase At3g48460 | — |
| TRINITY_DN17501_c1_g1 | Rola-high | -3.91 | 6.12e-03 | NA | NA | NA | | flavonoid 3'-hydroxylase | Venturia inaequalis |
| TRINITY_DN51555_c0_g1 | Rola-high | -3.91 | 8.26e-07 | NA | NA | NA | | 3-oxoacyl-[acyl-carrier-protein] synthase II, chloroplastic-like | Venturia inaequalis |
| TRINITY_DN15550_c0_g1 | Rola-high | -3.91 | 1.12e-03 | NA | NA | NA | | probable pectate lyase 5 | — |
| TRINITY_DN40610_c0_g2 | Rola-high | -3.90 | 3.92e-03 | NA | NA | NA | | ribonuclease TUDOR 2 | Venturia inaequalis |
| TRINITY_DN22522_c0_g1 | Rin-high | 3.90 | 2.04e-03 | NA | NA | NA | | aquaporin TIP2-1 | Venturia inaequalis |
| TRINITY_DN15514_c0_g1 | Rin-high | 3.90 | 7.54e-03 | NA | NA | NA | | silicon efflux transporter LSI2-like | — |
| TRINITY_DN81504_c0_g2 | Rin-high | 3.90 | 5.83e-03 | NA | NA | NA | | metallothionein-like protein | — |
| TRINITY_DN181804_c0_g1 | Rola-high | -3.89 | 6.18e-03 | NA | NA | NA | | ribosome production factor 2 homolog | Venturia inaequalis |
| TRINITY_DN4932_c0_g1 | Rin-high | 3.89 | 2.71e-05 | NA | NA | NA | | major latex allergen Hev b 5-like | — |
| TRINITY_DN43694_c0_g1 | Rola-high | -3.87 | 6.35e-06 | NA | NA | NA | | translationally-controlled tumor protein homolog | Venturia inaequalis |
| TRINITY_DN80939_c0_g1 | Rola-high | -3.87 | 7.77e-03 | NA | NA | NA | | putative acyl-activating enzyme 19 isoform X1 | Venturia inaequalis |

| Trinity gene ID | Direction | Trinity | | GDDH13 protein match | | | Def. | Description | FCS-GX call |
|---|---|---|---|---|---|---|---|---|---|
| | | log2FC | padj | gene ID | log2FC | padj | | | |
| TRINITY_DN16317_c0_g1 | Rola-high | -3.85 | 3.81e-03 | NA | NA | NA | | beta-xylosidase/alpha-L-arabinofuranosidase 1-like | Venturia inaequalis |
| TRINITY_DN23500_c0_g1 | Rola-high | -3.84 | 2.92e-06 | NA | NA | NA | | ATP synthase gamma chain, chloroplastic | Venturia inaequalis |
| TRINITY_DN7668_c2_g1 | Rin-high | 3.84 | 2.42e-09 | NA | NA | NA | yes | RAF-like serine/threonine-protein kinase PRAF | — |
| TRINITY_DN137072_c0_g1 | Rola-high | -3.83 | 5.23e-05 | NA | NA | NA | | carbamoyl phosphate synthase small chain, chloroplastic | Venturia inaequalis |
| TRINITY_DN55123_c0_g1 | Rola-high | -3.80 | 9.60e-03 | NA | NA | NA | | probable carboxylesterase 18 | Venturia inaequalis |
| TRINITY_DN47897_c0_g1 | Rola-high | -3.79 | 5.06e-05 | NA | NA | NA | | eukaryotic translation initiation factor 3 subunit B-like | Venturia inaequalis |
| TRINITY_DN30638_c0_g1 | Rola-high | -3.79 | 6.33e-03 | NA | NA | NA | | mitochondrial uncoupling protein 5-like | Venturia inaequalis |
| TRINITY_DN15170_c0_g1 | Rin-high | 3.78 | 6.92e-03 | NA | NA | NA | | uncharacterized protein | — |
| TRINITY_DN29157_c0_g1 | Rola-high | -3.78 | 5.39e-03 | NA | NA | NA | yes | cytochrome P450 76A2-like | Venturia inaequalis |
| TRINITY_DN49646_c0_g1 | Rola-high | -3.77 | 2.38e-04 | NA | NA | NA | | uncharacterized protein | Venturia inaequalis |
| TRINITY_DN137634_c0_g1 | Rola-high | -3.77 | 4.99e-03 | NA | NA | NA | | CBS domain-containing protein CBSCBSPB3-like | Venturia inaequalis |
| TRINITY_DN17541_c1_g1 | Rola-high | -3.77 | 1.79e-03 | NA | NA | NA | | protein INCREASED PETAL GROWTH ANISOTROPY 1-like | — |
| TRINITY_DN95024_c0_g3 | Rin-high | 3.75 | 4.77e-03 | NA | NA | NA | | thiol protease aleurain-like | — |
| TRINITY_DN23131_c0_g1 | Rola-high | -3.74 | 1.12e-05 | NA | NA | NA | | predicted protein | Venturia inaequalis |
| TRINITY_DN116791_c0_g1 | Rola-high | -3.72 | 6.00e-03 | NA | NA | NA | | organelle RRM domain-containing protein 2, mitochondrial-like | Venturia inaequalis |
| TRINITY_DN409_c0_g2 | Rola-high | -3.71 | 2.55e-03 | NA | NA | NA | | hypothetical protein DVH24_023579 | — |
| TRINITY_DN16381_c0_g1 | Rola-high | -3.70 | 4.16e-04 | NA | NA | NA | | THO complex subunit 4A-like | Venturia inaequalis |
| TRINITY_DN21742_c0_g1 | Rola-high | -3.69 | 3.68e-03 | NA | NA | NA | yes | class V chitinase CHIT5a-like | Venturia inaequalis |
| TRINITY_DN51659_c0_g1 | Rola-high | -3.69 | 2.54e-03 | NA | NA | NA | | large ribosomal subunit protein mL46-like | Venturia inaequalis |

| Trinity gene ID | Direction | Trinity log2FC | Trinity padj | GDDH13 protein match gene ID | GDDH13 log2FC | GDDH13 padj | Def. | Description | FCS-GX call |
|---|---|---|---|---|---|---|---|---|---|
| TRINITY_DN179528_c0_g1 | Rola-high | -3.69 | 4.45e-03 | NA | NA | NA | | (S)-coclaurine N-methyltransferase | Venturia inaequalis |
| TRINITY_DN38190_c0_g1 | Rola-high | -3.67 | 6.33e-03 | NA | NA | NA | yes | endoglucanase 24-like | — |
| TRINITY_DN73259_c0_g1 | Rola-high | -3.65 | 5.34e-03 | NA | NA | NA | | rRNA biogenesis protein RRP5 | Venturia inaequalis |
| TRINITY_DN39634_c0_g1 | Rola-high | -3.65 | 7.56e-03 | NA | NA | NA | | importin beta-like SAD2 | Venturia inaequalis |
| TRINITY_DN37007_c0_g1 | Rola-high | -3.64 | 3.88e-03 | NA | NA | NA | | ATP synthase subunit delta', mitochondrial | Venturia inaequalis |
| TRINITY_DN29436_c0_g2 | Rola-high | -3.64 | 3.72e-03 | NA | NA | NA | | transposon Tf2-1 polyprotein | — |
| TRINITY_DN199925_c0_g1 | Rola-high | -3.61 | 2.51e-04 | NA | NA | NA | | protein RNA-directed DNA methylation 3-like | Venturia inaequalis |
| TRINITY_DN7210_c2_g1 | Rola-high | -3.59 | 1.86e-03 | NA | NA | NA | yes | probably inactive leucine-rich repeat receptor-like protein kinase IMK2 | — |
| TRINITY_DN114517_c0_g1 | Rola-high | -3.59 | 7.03e-05 | NA | NA | NA | | S-adenosylmethionine decarboxylase proenzyme-like | Venturia inaequalis |
| TRINITY_DN48488_c0_g1 | Rola-high | -3.59 | 5.48e-03 | NA | NA | NA | | bifunctional dethiobiotin synthetase/7,8-diamino-pelargonic acid aminotransferase, mitochondrial | Venturia inaequalis |
| TRINITY_DN17387_c0_g1 | Rola-high | -3.58 | 2.17e-03 | NA | NA | NA | | eukaryotic translation initiation factor 3 subunit H | Venturia inaequalis |
| TRINITY_DN3991_c0_g1 | Rin-high | 3.57 | 9.75e-03 | NA | NA | NA | | BAG family molecular chaperone regulator 2-like | — |
| TRINITY_DN53216_c0_g1 | Rola-high | -3.57 | 2.40e-04 | NA | NA | NA | | mitochondrial carrier protein MTM1-like | Venturia inaequalis |
| TRINITY_DN12093_c0_g1 | Rola-high | -3.56 | 2.47e-03 | NA | NA | NA | | haloacid dehalogenase-like hydrolase domain-containing protein Sgpp isoform X2 | Venturia inaequalis |
| TRINITY_DN10633_c0_g2 | Rola-high | -3.55 | 8.09e-06 | NA | NA | NA | | hypothetical protein DVH24_011124 | — |
| TRINITY_DN43224_c0_g1 | Rola-high | -3.55 | 3.12e-04 | NA | NA | NA | | eukaryotic translation initiation factor 3 subunit K-like | Venturia inaequalis |
| TRINITY_DN24232_c0_g1 | Rola-high | -3.54 | 8.32e-03 | NA | NA | NA | | root phototropism protein 2-like | — |
| TRINITY_DN3724_c1_g2 | Rola-high | -3.54 | 5.08e-03 | NA | NA | NA | | proline-rich protein 4-like | — |
| TRINITY_DN61869_c0_g1 | Rin-high | 3.51 | 2.32e-03 | NA | NA | NA | | zinc finger protein CONSTANS-LIKE 9-like | — |

| Trinity gene ID | Direction | Trinity | | GDDH13 protein match | | | Def. | Description | FCS-GX |
|---|---|---|---|---|---|---|---|---|---|
| | | log2FC | padj | gene ID | log2FC | padj | | | call |
| TRINITY_DN205507_c0_g1 | Rola-high | -3.47 | 3.85e-03 | NA | NA | NA | | vacuolar protein-sorting-associated protein 11 homolog | Venturia inaequalis |
| TRINITY_DN135963_c0_g1 | Rola-high | -3.46 | 2.02e-03 | NA | NA | NA | | heat shock 70 kDa protein 16-like | Venturia inaequalis |
| TRINITY_DN8187_c0_g2 | Rola-high | -3.45 | 2.77e-04 | NA | NA | NA | | hypothetical protein DVH24_010841 | — |
| TRINITY_DN28483_c1_g1 | Rola-high | -3.44 | 1.82e-03 | NA | NA | NA | | protein disulfide-isomerase-like | Venturia inaequalis |
| TRINITY_DN10324_c0_g2 | Rola-high | -3.42 | 4.47e-06 | NA | NA | NA | yes | thaumatin-like protein 1 | — |
| TRINITY_DN203508_c0_g1 | Rola-high | -3.41 | 9.00e-03 | NA | NA | NA | | coatomer subunit delta-like | Venturia inaequalis |
| TRINITY_DN160046_c0_g1 | Rola-high | -3.39 | 3.30e-03 | NA | NA | NA | | ATP synthase subunit O, mitochondrial | Venturia inaequalis |
| TRINITY_DN12313_c0_g1 | Rola-high | -3.39 | 7.96e-05 | NA | NA | NA | | 28 kDa ribonucleoprotein, chloroplastic-like | Venturia inaequalis |
| TRINITY_DN15092_c0_g1 | Rola-high | -3.38 | 8.62e-03 | NA | NA | NA | | ATP-citrate synthase alpha chain protein 1 | Venturia inaequalis |
| TRINITY_DN199916_c0_g1 | Rola-high | -3.36 | 8.44e-03 | NA | NA | NA | | beta-galactosidase 8-like | Venturia inaequalis |
| TRINITY_DN158996_c0_g1 | Rola-high | -3.35 | 5.87e-03 | NA | NA | NA | | nuclear transport factor 2-like isoform X2 | Venturia inaequalis |
| TRINITY_DN21015_c0_g1 | Rola-high | -3.34 | 2.51e-03 | NA | NA | NA | | granule-bound starch synthase 1, chloroplastic/amyloplastic-like isoform X1 | Venturia inaequalis |
| TRINITY_DN4022_c0_g1 | Rin-high | 3.31 | 3.04e-04 | NA | NA | NA | | hypothetical protein DVH24_028168 | — |
| TRINITY_DN135397_c0_g1 | Rola-high | -3.30 | 6.70e-04 | NA | NA | NA | | eukaryotic translation initiation factor 3 subunit A-like | Venturia inaequalis |
| TRINITY_DN44395_c0_g1 | Rola-high | -3.30 | 2.99e-03 | NA | NA | NA | | aminopeptidase M1-like isoform X1 | Venturia inaequalis |
| TRINITY_DN23157_c0_g2 | Rola-high | -3.28 | 3.56e-03 | NA | NA | NA | | callose synthase 3 | Venturia inaequalis |
| TRINITY_DN44586_c0_g1 | Rola-high | -3.26 | 7.95e-03 | NA | NA | NA | | protein transport protein SEC31 homolog B-like | Venturia inaequalis |
| TRINITY_DN114551_c0_g1 | Rola-high | -3.25 | 5.94e-03 | NA | NA | NA | | pentatricopeptide repeat-containing protein At4g20090-like isoform X1 | Venturia inaequalis |
| TRINITY_DN49491_c1_g1 | Rola-high | -3.23 | 5.72e-08 | NA | NA | NA | | uncharacterized mitochondrial protein AtMg00810-like | — |

| Trinity gene ID | Direction | Trinity | | GDDH13 protein match | | | Def. | Description | FCS-GX |
|---|---|---|---|---|---|---|---|---|---|
| | | log2FC | padj | gene ID | log2FC | padj | | | call |
| TRINITY_DN4318_c0_g2 | Rola-high | -3.20 | 3.10e-03 | NA | NA | NA | | 8-hydroxygeraniol dehydrogenase-like | — |
| TRINITY_DN103806_c0_g1 | Rola-high | -3.19 | 5.69e-03 | NA | NA | NA | | 3-isopropylmalate dehydratase large subunit, chloroplastic | Venturia inaequalis |
| TRINITY_DN143816_c0_g1 | Rola-high | -3.19 | 1.58e-03 | NA | NA | NA | | co-chaperone protein p23-2 | Venturia inaequalis |
| TRINITY_DN179664_c0_g1 | Rola-high | -3.19 | 1.06e-03 | NA | NA | NA | | acetylornithine aminotransferase, mitochondrial-like | Venturia inaequalis |
| TRINITY_DN383_c2_g1 | Rola-high | -3.17 | 2.58e-03 | NA | NA | NA | | hypothetical protein DVH24_013523 | — |
| TRINITY_DN21675_c0_g1 | Rin-high | 3.14 | 3.22e-03 | NA | NA | NA | | hypothetical protein DVH24_026834 | — |
| TRINITY_DN40636_c0_g1 | Rola-high | -3.13 | 7.10e-03 | NA | NA | NA | | eukaryotic translation initiation factor 4G-like isoform X1 | Venturia inaequalis |
| TRINITY_DN46258_c0_g1 | Rola-high | -3.11 | 1.92e-03 | NA | NA | NA | | aconitate hydratase 1 | Venturia inaequalis |
| TRINITY_DN19532_c0_g2 | Rola-high | -3.08 | 3.11e-04 | NA | NA | NA | | hypothetical protein DVH24_032896 | — |
| TRINITY_DN23466_c0_g3 | Rin-high | 3.08 | 3.21e-05 | NA | NA | NA | | FCS-Like Zinc finger 10-like | — |
| TRINITY_DN32565_c0_g1 | Rola-high | -3.00 | 2.33e-03 | NA | NA | NA | | delta(12)-fatty-acid desaturase FAD2 | Venturia inaequalis |
| TRINITY_DN1244_c0_g1 | Rin-high | 3.00 | 6.75e-03 | NA | NA | NA | | TRINITY_DN1244_c0_g1 | — |
| TRINITY_DN157384_c0_g1 | Rola-high | -2.95 | 4.51e-03 | NA | NA | NA | | palmitoyl-monogalactosyldiacylglycerol delta-7 desaturase, chloroplastic-like | Venturia inaequalis |
| TRINITY_DN17014_c0_g1 | Rola-high | -2.95 | 1.44e-03 | NA | NA | NA | | fatty acid elongase 3-like | Venturia inaequalis |
| TRINITY_DN182193_c0_g1 | Rola-high | -2.92 | 3.23e-03 | NA | NA | NA | | translocon at the outer membrane of chloroplasts 64 | Venturia inaequalis |
| TRINITY_DN2409_c0_g3 | Rola-high | -2.89 | 1.46e-03 | NA | NA | NA | yes | leucine-rich repeat receptor-like protein | — |
| TRINITY_DN25556_c0_g1 | Rin-high | 2.88 | 4.45e-03 | NA | NA | NA | | sugar transport protein 7 | Venturia inaequalis |
| TRINITY_DN9074_c0_g1 | Rola-high | -2.88 | 1.54e-03 | NA | NA | NA | | nuclear intron maturase 1, mitochondrial | Venturia inaequalis |
| TRINITY_DN23031_c0_g1 | Rin-high | 2.81 | 6.92e-03 | NA | NA | NA | | pentatricopeptide repeat-containing protein At5g01110 | — |
| TRINITY_DN13959_c0_g1 | Rola-high | -2.80 | 6.48e-04 | NA | NA | NA | yes | leucine-rich repeat receptor-like protein | — |

| Trinity gene ID | Direction | Trinity |  | GDDH13 protein match |  |  | Def. | Description | FCS-GX call |
|---|---|---|---|---|---|---|---|---|---|
|  |  | log2FC | padj | gene ID | log2FC | padj |  |  |  |
| TRINITY_DN65578_c0_g1 | Rola-high | -2.80 | 8.34e-03 | NA | NA | NA |  | 2-isopropylmalate synthase 1, chloroplastic-like | Venturia inaequalis |
| TRINITY_DN1340_c3_g1 | Rola-high | -2.80 | 1.88e-03 | NA | NA | NA |  | hypothetical protein DVH24_005067 | — |
| TRINITY_DN24134_c1_g1 | Rola-high | -2.80 | 8.76e-03 | NA | NA | NA |  | 8-hydroxygeraniol dehydrogenase-like | — |
| TRINITY_DN1994_c0_g1 | Rola-high | -2.80 | 3.56e-03 | NA | NA | NA | yes | pectin methylesterase | — |
| TRINITY_DN16367_c0_g1 | Rola-high | -2.77 | 2.09e-03 | NA | NA | NA |  | (R)-mandelonitrile lyase 3-like | Venturia inaequalis |
| TRINITY_DN27260_c0_g1 | Rola-high | -2.74 | 8.79e-04 | NA | NA | NA |  | flocculation protein FLO11-like | — |
| TRINITY_DN32229_c0_g1 | Rin-high | 2.72 | 7.92e-04 | NA | NA | NA |  | two-component response regulator-like APRR7 isoform X2 | — |
| TRINITY_DN8258_c2_g1 | Rin-high | 2.71 | 5.10e-03 | NA | NA | NA |  | glucan endo-1,3-beta-glucosidase 14-like isoform X3 | — |
| TRINITY_DN33262_c0_g1 | Rola-high | -2.70 | 9.62e-04 | NA | NA | NA | yes | GDSL esterase/lipase EXL3-like | — |
| TRINITY_DN9087_c0_g1 | Rin-high | 2.66 | 2.39e-04 | NA | NA | NA |  | MADS-box protein SVP-like isoform X1 | — |
| TRINITY_DN20929_c0_g1 | Rola-high | -2.61 | 1.76e-03 | NA | NA | NA |  | heavy metal-associated isoprenylated plant protein 7-like | — |
| TRINITY_DN3274_c0_g2 | Rin-high | 2.60 | 8.89e-03 | NA | NA | NA |  | hypothetical protein DVH24_012087 | — |
| TRINITY_DN234_c2_g1 | Rin-high | 2.44 | 5.97e-03 | NA | NA | NA |  | uncharacterized protein | — |
| TRINITY_DN27143_c0_g1 | Rola-high | -2.39 | 7.52e-03 | NA | NA | NA |  | OVATE family protein | — |
| TRINITY_DN34085_c0_g1 | Rola-high | -2.30 | 4.89e-03 | NA | NA | NA |  | uncharacterized protein | — |
| TRINITY_DN34109_c0_g1 | Rola-high | -2.29 | 7.49e-03 | NA | NA | NA |  | galactan beta-1,4-galactosyltransferase GALS3 | — |
| TRINITY_DN13351_c2_g1 | Rin-high | 2.26 | 9.66e-03 | NA | NA | NA |  | hypothetical protein DVH24_006652 | — |
| TRINITY_DN263_c0_g1 | Rin-high | 2.18 | 5.25e-03 | NA | NA | NA |  | uncharacterized protein | — |
| TRINITY_DN211561_c0_g1 | Rola-high | -2.18 | 8.88e-03 | NA | NA | NA |  | probable CoA ligase CCL6 | — |
| TRINITY_DN1473_c0_g2 | Rola-high | -2.16 | 8.54e-03 | NA | NA | NA |  | auxin-induced protein 15A-like | — |
| TRINITY_DN27976_c0_g1 | Rin-high | 1.92 | 5.02e-03 | NA | NA | NA |  | transcription factor DIVARICATA-like | — |

| Trinity gene ID | Direction | Trinity |  | GDDH13 protein match |  |  | Def. | Description | FCS-GX |
|---|---|---|---|---|---|---|---|---|---|
|  |  | log2FC | padj | gene ID | log2FC | padj |  |  | call |
| **No GDDH13 hit — plant/defense annotation (n = 4)** |  |  |  |  |  |  |  |  |  |
| TRINITY_DN36341_c0_g1 | Rola-high | -5.25 | 1.01e-06 | NA | NA | NA | yes | choline kinase | Venturia inaequalis |
| TRINITY_DN86948_c0_g1 | Rin-high | 5.25 | 9.91e-03 | NA | NA | NA | yes | LRR receptor-like serine/threonine-protein kinase GSO2 | — |
| TRINITY_DN138679_c0_g1 | Rola-high | -4.06 | 9.89e-03 | NA | NA | NA | yes | calcium-dependent phosphotriesterase | Venturia inaequalis |
| TRINITY_DN7235_c0_g1 | Rola-high | -3.73 | 1.69e-04 | NA | NA | NA | yes | leucine-rich repeat extensin-like protein 3 | — |
| **No GDDH13 hit — fungal/pathogen-like annotation (n = 0)** |  |  |  |  |  |  |  |  |  |
| **None.** |  |  |  |  |  |  |  |  |  |
| **No GDDH13 hit — no informative annotation (n = 385)** |  |  |  |  |  |  |  |  |  |
| TRINITY_DN36061_c1_g1 | Rola-high | -25.59 | 3.16e-25 | NA | NA | NA |  | protein PELPK2-like | — |
| TRINITY_DN11246_c0_g1 | Rola-high | -23.39 | 7.81e-07 | NA | NA | NA |  | TRINITY_DN11246_c0_g1 | Trialeurodes vaporariorum |
| TRINITY_DN139024_c0_g1 | Rola-high | -23.05 | 1.56e-06 | NA | NA | NA |  | TRINITY_DN139024_c0_g1 | Trialeurodes vaporariorum |
| TRINITY_DN6336_c0_g1 | Rola-high | -22.92 | 1.28e-06 | NA | NA | NA |  | TRINITY_DN6336_c0_g1 | Trialeurodes vaporariorum |
| TRINITY_DN229006_c0_g1 | Rola-high | -22.73 | 2.37e-06 | NA | NA | NA |  | TRINITY_DN229006_c0_g1 | Trialeurodes vaporariorum |
| TRINITY_DN21772_c0_g1 | Rola-high | -22.34 | 1.88e-06 | NA | NA | NA |  | TRINITY_DN21772_c0_g1 | Trialeurodes vaporariorum |
| TRINITY_DN81325_c0_g2 | Rola-high | -21.34 | 3.93e-06 | NA | NA | NA |  | TRINITY_DN81325_c0_g2 | Trialeurodes vaporariorum |
| TRINITY_DN138657_c0_g1 | Rola-high | -21.10 | 7.80e-06 | NA | NA | NA |  | TRINITY_DN138657_c0_g1 | Trialeurodes vaporariorum |
| TRINITY_DN3521_c0_g1 | Rola-high | -12.89 | 9.31e-04 | NA | NA | NA |  | TRINITY_DN3521_c0_g1 | Trialeurodes vaporariorum |
| TRINITY_DN5274_c0_g1 | Rola-high | -10.58 | 2.43e-05 | NA | NA | NA |  | glycine-rich cell wall structural protein-like | — |
| TRINITY_DN104814_c0_g1 | Rola-high | -9.66 | 3.39e-08 | NA | NA | NA |  | putative cell wall protein | — |
| TRINITY_DN40755_c0_g1 | Rin-high | 8.99 | 2.43e-03 | NA | NA | NA |  | hypothetical protein DVH24_042242 | — |

| Trinity gene ID | Direction | Trinity | | GDDH13 protein match | | | Def. | Description | FCS-GX call |
|---|---|---|---|---|---|---|---|---|---|
| | | log2FC | padj | gene ID | log2FC | padj | | | |
| TRINITY_DN39714_c1_g1 | Rola-high | -8.93 | 9.02e-04 | NA | NA | NA | | hypothetical protein DVH24_038767 | — |
| TRINITY_DN5274_c1_g1 | Rola-high | -8.26 | 1.10e-03 | NA | NA | NA | | TRINITY_DN5274_c1_g1 | — |
| TRINITY_DN21285_c1_g1 | Rin-high | 7.66 | 4.36e-03 | NA | NA | NA | | TRINITY_DN21285_c1_g1 | — |
| TRINITY_DN59053_c0_g1 | Rola-high | -7.58 | 2.47e-03 | NA | NA | NA | | TRINITY_DN59053_c0_g1 | — |
| TRINITY_DN168797_c0_g1 | Rola-high | -7.58 | 1.50e-05 | NA | NA | NA | | glycine-rich protein 5 | — |
| TRINITY_DN24576_c0_g1 | Rola-high | -7.47 | 5.44e-12 | NA | NA | NA | | TRINITY_DN24576_c0_g1 | Venturia inaequalis |
| TRINITY_DN33503_c0_g1 | Rola-high | -7.43 | 2.15e-03 | NA | NA | NA | | TRINITY_DN33503_c0_g1 | Venturia inaequalis |
| TRINITY_DN10526_c0_g1 | Rola-high | -7.42 | 2.26e-08 | NA | NA | NA | | TRINITY_DN10526_c0_g1 | Venturia inaequalis |
| TRINITY_DN8735_c0_g1 | Rola-high | -7.39 | 2.92e-06 | NA | NA | NA | | TRINITY_DN8735_c0_g1 | Venturia inaequalis |
| TRINITY_DN1769_c0_g2 | Rola-high | -7.17 | 1.55e-04 | NA | NA | NA | | TRINITY_DN1769_c0_g2 | — |
| TRINITY_DN15559_c1_g2 | Rin-high | 7.09 | 2.05e-03 | NA | NA | NA | | TRINITY_DN15559_c1_g2 | — |
| TRINITY_DN24683_c0_g1 | Rin-high | 7.08 | 2.52e-04 | NA | NA | NA | | TRINITY_DN24683_c0_g1 | — |
| TRINITY_DN9456_c0_g1 | Rola-high | -6.91 | 6.98e-03 | NA | NA | NA | | hypothetical protein SELMODRAFT_407912 | Venturia inaequalis |
| TRINITY_DN28541_c0_g2 | Rin-high | 6.90 | 3.71e-04 | NA | NA | NA | | TRINITY_DN28541_c0_g2 | — |
| TRINITY_DN135606_c0_g1 | Rola-high | -6.82 | 7.22e-06 | NA | NA | NA | | hypothetical protein R6Q59_009881 | Venturia inaequalis |
| TRINITY_DN138530_c0_g1 | Rola-high | -6.78 | 1.32e-05 | NA | NA | NA | | TRINITY_DN138530_c0_g1 | — |
| TRINITY_DN114441_c0_g1 | Rin-high | 6.76 | 1.31e-07 | NA | NA | NA | | TRINITY_DN114441_c0_g1 | Venturia inaequalis |
| TRINITY_DN113352_c0_g1 | Rola-high | -6.74 | 1.72e-05 | NA | NA | NA | | TRINITY_DN113352_c0_g1 | Venturia inaequalis |
| TRINITY_DN32851_c0_g1 | Rin-high | 6.69 | 1.17e-04 | NA | NA | NA | | TRINITY_DN32851_c0_g1 | Venturia inaequalis |
| TRINITY_DN14166_c0_g1 | Rola-high | -6.65 | 6.79e-08 | NA | NA | NA | | TRINITY_DN14166_c0_g1 | Venturia inaequalis |

| Trinity gene ID | Direction | Trinity | | GDDH13 protein match | | | Def. | Description | FCS-GX |
|---|---|---|---|---|---|---|---|---|---|
| | | log2FC | padj | gene ID | log2FC | padj | | | call |
| TRINITY_DN56766_c0_g2 | Rola-high | -6.58 | 3.07e-05 | NA | NA | NA | | TRINITY_DN56766_c0_g2 | Venturia inaequalis |
| TRINITY_DN16321_c0_g1 | Rola-high | -6.57 | 6.12e-06 | NA | NA | NA | | TRINITY_DN16321_c0_g1 | Venturia inaequalis |
| TRINITY_DN19148_c0_g1 | Rola-high | -6.55 | 1.31e-07 | NA | NA | NA | | TRINITY_DN19148_c0_g1 | — |
| TRINITY_DN29527_c0_g3 | Rola-high | -6.53 | 3.70e-05 | NA | NA | NA | | TRINITY_DN29527_c0_g3 | Venturia inaequalis |
| TRINITY_DN157_c0_g1 | Rola-high | -6.53 | 1.23e-06 | NA | NA | NA | | TRINITY_DN157_c0_g1 | Venturia inaequalis |
| TRINITY_DN60234_c0_g2 | Rin-high | 6.53 | 4.86e-04 | NA | NA | NA | | uncharacterized protein | — |
| TRINITY_DN61969_c0_g1 | Rin-high | 6.52 | 3.13e-04 | NA | NA | NA | | TRINITY_DN61969_c0_g1 | — |
| TRINITY_DN10249_c0_g1 | Rola-high | -6.51 | 1.56e-06 | NA | NA | NA | | TRINITY_DN10249_c0_g1 | Venturia inaequalis |
| TRINITY_DN37295_c0_g1 | Rin-high | 6.47 | 2.77e-03 | NA | NA | NA | | hypothetical protein DVH24_004313 | — |
| TRINITY_DN116487_c0_g1 | Rola-high | -6.45 | 6.31e-10 | NA | NA | NA | | TRINITY_DN116487_c0_g1 | Venturia inaequalis |
| TRINITY_DN19239_c0_g1 | Rola-high | -6.40 | 1.62e-03 | NA | NA | NA | | TRINITY_DN19239_c0_g1 | Venturia inaequalis |
| TRINITY_DN8348_c0_g2 | Rola-high | -6.38 | 2.92e-06 | NA | NA | NA | | TRINITY_DN8348_c0_g2 | Venturia inaequalis |
| TRINITY_DN40974_c0_g1 | Rola-high | -6.38 | 3.88e-06 | NA | NA | NA | | hypothetical protein TorRG33x02_043960 | — |
| TRINITY_DN83917_c0_g1 | Rola-high | -6.34 | 2.06e-03 | NA | NA | NA | | TRINITY_DN83917_c0_g1 | Venturia inaequalis |
| TRINITY_DN11641_c0_g1 | Rola-high | -6.29 | 4.48e-05 | NA | NA | NA | | phenolic acid decarboxilase | Venturia inaequalis |
| TRINITY_DN1562_c0_g1 | Rola-high | -6.24 | 3.78e-04 | NA | NA | NA | | TRINITY_DN1562_c0_g1 | Venturia inaequalis |
| TRINITY_DN3989_c2_g5 | Rola-high | -6.24 | 7.14e-03 | NA | NA | NA | | hypothetical protein DVH24_009784 | — |
| TRINITY_DN64650_c0_g1 | Rola-high | -6.21 | 7.42e-05 | NA | NA | NA | | TRINITY_DN64650_c0_g1 | Venturia inaequalis |
| TRINITY_DN13709_c1_g1 | Rola-high | -6.19 | 1.03e-04 | NA | NA | NA | | hypothetical protein DVH24_000929 | — |
| TRINITY_DN3133_c2_g1 | Rola-high | -6.16 | 4.46e-07 | NA | NA | NA | | transcription factor PAR2-like | — |

| Trinity gene ID | Direction | Trinity |  | GDDH13 protein match |  |  | Def. | Description | FCS-GX call |
|---|---|---|---|---|---|---|---|---|---|
|  |  | log2FC | padj | gene ID | log2FC | padj |  |  |  |
| TRINITY_DN28602_c0_g2 | Rola-high | -6.13 | 3.41e-03 | NA | NA | NA |  | TRINITY_DN28602_c0_g2 | — |
| TRINITY_DN16783_c0_g1 | Rin-high | 6.06 | 5.78e-05 | NA | NA | NA |  | TRINITY_DN16783_c0_g1 | — |
| TRINITY_DN12121_c1_g1 | Rola-high | -6.05 | 3.11e-04 | NA | NA | NA |  | TRINITY_DN12121_c1_g1 | — |
| TRINITY_DN42241_c0_g1 | Rin-high | 6.00 | 5.55e-04 | NA | NA | NA |  | hypothetical protein C1H46_018590 | — |
| TRINITY_DN7837_c0_g1 | Rola-high | -6.00 | 9.86e-03 | NA | NA | NA |  | hypothetical chloroplast RF1 | Venturia inaequalis |
| TRINITY_DN4253_c0_g1 | Rola-high | -5.98 | 1.08e-06 | NA | NA | NA |  | TRINITY_DN4253_c0_g1 | Venturia inaequalis |
| TRINITY_DN2658_c0_g1 | Rola-high | -5.98 | 4.14e-03 | NA | NA | NA |  | TRINITY_DN2658_c0_g1 | — |
| TRINITY_DN5413_c0_g2 | Rola-high | -5.98 | 2.84e-03 | NA | NA | NA |  | hypothetical protein DVH24_005859 | — |
| TRINITY_DN99474_c0_g1 | Rola-high | -5.97 | 6.62e-03 | NA | NA | NA |  | TRINITY_DN99474_c0_g1 | Venturia inaequalis |
| TRINITY_DN157392_c0_g1 | Rola-high | -5.95 | 6.34e-05 | NA | NA | NA |  | MFS transporter, ACS family, DAL5 transporter family protein | Venturia inaequalis |
| TRINITY_DN6344_c4_g1 | Rola-high | -5.90 | 5.89e-07 | NA | NA | NA |  | TRINITY_DN6344_c4_g1 | — |
| TRINITY_DN6138_c0_g1 | Rola-high | -5.89 | 5.69e-03 | NA | NA | NA |  | TRINITY_DN6138_c0_g1 | Venturia inaequalis |
| TRINITY_DN35136_c0_g1 | Rola-high | -5.88 | 1.13e-04 | NA | NA | NA |  | MAG: major facilitator superfamily domain-containing protein | Venturia inaequalis |
| TRINITY_DN17880_c0_g1 | Rin-high | 5.87 | 6.88e-03 | NA | NA | NA |  | hypothetical protein DVH24_021445 | — |
| TRINITY_DN38795_c0_g2 | Rola-high | -5.87 | 1.08e-03 | NA | NA | NA |  | TRINITY_DN38795_c0_g2 | Venturia inaequalis |
| TRINITY_DN205903_c0_g1 | Rola-high | -5.84 | 5.94e-03 | NA | NA | NA |  | TRINITY_DN205903_c0_g1 | Venturia inaequalis |
| TRINITY_DN3157_c0_g1 | Rola-high | -5.82 | 1.95e-03 | NA | NA | NA |  | TRINITY_DN3157_c0_g1 | Venturia inaequalis |
| TRINITY_DN224519_c0_g1 | Rin-high | 5.82 | 2.64e-03 | NA | NA | NA |  | TRINITY_DN224519_c0_g1 | — |
| TRINITY_DN11996_c0_g1 | Rola-high | -5.82 | 6.77e-06 | NA | NA | NA |  | heavy metal-associated isoprenylated plant protein 32-like | — |
| TRINITY_DN159311_c0_g1 | Rola-high | -5.82 | 7.35e-05 | NA | NA | NA |  | TRINITY_DN159311_c0_g1 | Venturia inaequalis |

| Trinity gene ID | Direction | Trinity | | GDDH13 protein match | | | Def. | Description | FCS-GX call |
|---|---|---|---|---|---|---|---|---|---|
| | | log2FC | padj | gene ID | log2FC | padj | | | |
| TRINITY_DN30971_c0_g1 | Rola-high | -5.79 | 4.54e-04 | NA | NA | NA | | TRINITY_DN30971_c0_g1 | Venturia inaequalis |
| TRINITY_DN65212_c0_g1 | Rola-high | -5.78 | 1.80e-05 | NA | NA | NA | | TRINITY_DN65212_c0_g1 | Venturia inaequalis |
| TRINITY_DN17126_c0_g1 | Rin-high | 5.76 | 6.28e-05 | NA | NA | NA | | protein SRC2-like isoform X2 | — |
| TRINITY_DN98698_c0_g1 | Rola-high | -5.74 | 7.75e-04 | NA | NA | NA | | hypothetical protein MRB53_038860 | Venturia inaequalis |
| TRINITY_DN1199_c0_g1 | Rola-high | -5.70 | 4.60e-12 | NA | NA | NA | | TRINITY_DN1199_c0_g1 | Venturia inaequalis |
| TRINITY_DN70102_c0_g1 | Rola-high | -5.69 | 1.08e-08 | NA | NA | NA | | TRINITY_DN70102_c0_g1 | Venturia inaequalis |
| TRINITY_DN38343_c0_g1 | Rola-high | -5.66 | 4.26e-04 | NA | NA | NA | | TRINITY_DN38343_c0_g1 | Venturia inaequalis |
| TRINITY_DN15031_c0_g1 | Rola-high | -5.66 | 3.64e-03 | NA | NA | NA | | unnamed protein product | — |
| TRINITY_DN32088_c0_g1 | Rola-high | -5.65 | 1.01e-11 | NA | NA | NA | | TRINITY_DN32088_c0_g1 | — |
| TRINITY_DN27761_c1_g1 | Rin-high | 5.65 | 3.30e-03 | NA | NA | NA | | hypothetical protein DVH24_001312 | — |
| TRINITY_DN12851_c2_g1 | Rola-high | -5.65 | 1.74e-04 | NA | NA | NA | | hypothetical protein D8674_036854 | — |
| TRINITY_DN13995_c0_g1 | Rin-high | 5.65 | 1.57e-03 | NA | NA | NA | | TRINITY_DN13995_c0_g1 | — |
| TRINITY_DN178675_c0_g1 | Rola-high | -5.64 | 3.67e-05 | NA | NA | NA | | unnamed protein product | Venturia inaequalis |
| TRINITY_DN7401_c0_g1 | Rola-high | -5.63 | 9.13e-04 | NA | NA | NA | | TRINITY_DN7401_c0_g1 | Venturia inaequalis |
| TRINITY_DN59473_c0_g1 | Rin-high | 5.63 | 2.92e-06 | NA | NA | NA | | hypothetical protein DVH24_022328 | — |
| TRINITY_DN8910_c0_g2 | Rin-high | 5.63 | 3.75e-06 | NA | NA | NA | | TRINITY_DN8910_c0_g2 | — |
| TRINITY_DN136747_c0_g1 | Rola-high | -5.60 | 9.96e-03 | NA | NA | NA | | hypothetical protein MRB53_041685 | Venturia inaequalis |
| TRINITY_DN29371_c0_g1 | Rola-high | -5.60 | 9.40e-04 | NA | NA | NA | | TRINITY_DN29371_c0_g1 | Venturia inaequalis |
| TRINITY_DN42933_c1_g3 | Rola-high | -5.60 | 4.47e-03 | NA | NA | NA | | unnamed protein product | — |
| TRINITY_DN7666_c0_g2 | Rola-high | -5.59 | 5.50e-03 | NA | NA | NA | | TRINITY_DN7666_c0_g2 | — |

| Trinity gene ID | Direction | Trinity | | GDDH13 protein match | | | Def. | Description | FCS-GX |
|---|---|---|---|---|---|---|---|---|---|
| | | log2FC | padj | gene ID | log2FC | padj | | | call |
| TRINITY_DN38273_c0_g1 | Rola-high | -5.58 | 4.79e-07 | NA | NA | NA | | hypothetical protein MRB53_036981 | Venturia inaequalis |
| TRINITY_DN14535_c0_g1 | Rola-high | -5.56 | 1.52e-06 | NA | NA | NA | | S-methylmethionine permease | Venturia inaequalis |
| TRINITY_DN52849_c0_g2 | Rola-high | -5.55 | 2.85e-06 | NA | NA | NA | | TRINITY_DN52849_c0_g2 | — |
| TRINITY_DN228843_c0_g1 | Rola-high | -5.54 | 5.71e-04 | NA | NA | NA | | TRINITY_DN228843_c0_g1 | Venturia inaequalis |
| TRINITY_DN36065_c0_g1 | Rin-high | 5.54 | 1.98e-04 | NA | NA | NA | | hypothetical protein CYMTET_49262 | Venturia inaequalis |
| TRINITY_DN57199_c0_g1 | Rola-high | -5.54 | 2.24e-06 | NA | NA | NA | | hypothetical protein WJX84_005158 | Venturia inaequalis |
| TRINITY_DN21544_c0_g1 | Rola-high | -5.53 | 1.61e-05 | NA | NA | NA | | hypothetical protein DVH24_006016 | — |
| TRINITY_DN10824_c0_g1 | Rola-high | -5.53 | 7.71e-03 | NA | NA | NA | | TRINITY_DN10824_c0_g1 | Venturia inaequalis |
| TRINITY_DN8800_c0_g1 | Rin-high | 5.52 | 9.65e-05 | NA | NA | NA | | TRINITY_DN8800_c0_g1 | Venturia inaequalis |
| TRINITY_DN22514_c0_g3 | Rola-high | -5.52 | 5.29e-03 | NA | NA | NA | | hypothetical protein DVH24_035451 | — |
| TRINITY_DN755_c3_g3 | Rola-high | -5.50 | 3.20e-03 | NA | NA | NA | | vestitone reductase-like isoform X1 | — |
| TRINITY_DN47204_c0_g1 | Rola-high | -5.48 | 1.24e-05 | NA | NA | NA | | TRINITY_DN47204_c0_g1 | Venturia inaequalis |
| TRINITY_DN32470_c0_g1 | Rola-high | -5.48 | 2.36e-03 | NA | NA | NA | | hypothetical protein R6Q59_010223 | Venturia inaequalis |
| TRINITY_DN3007_c1_g4 | Rin-high | 5.47 | 3.57e-05 | NA | NA | NA | | hypothetical protein DVH24_019536 | — |
| TRINITY_DN16962_c0_g1 | Rola-high | -5.47 | 1.70e-03 | NA | NA | NA | | TRINITY_DN16962_c0_g1 | Venturia inaequalis |
| TRINITY_DN203070_c0_g1 | Rola-high | -5.46 | 2.94e-04 | NA | NA | NA | | hypothetical protein Dsin_033214 | Venturia inaequalis |
| TRINITY_DN35253_c0_g1 | Rola-high | -5.45 | 1.36e-03 | NA | NA | NA | | predicted protein | Venturia inaequalis |
| TRINITY_DN372_c0_g4 | Rola-high | -5.44 | 2.43e-03 | NA | NA | NA | | hypothetical protein DVH24_023122 | — |
| TRINITY_DN118961_c0_g1 | Rola-high | -5.43 | 5.94e-03 | NA | NA | NA | | TRINITY_DN118961_c0_g1 | Venturia inaequalis |

| Trinity gene ID | Direction | Trinity | | GDDH13 protein match | | | Def. | Description | FCS-GX |
|---|---|---|---|---|---|---|---|---|---|
| | | log2FC | padj | gene ID | log2FC | padj | | | call |
| TRINITY_DN23281_c1_g2 | Rola-high | -5.42 | 2.39e-03 | NA | NA | NA | | hypothetical protein DVH24_004300 | — |
| TRINITY_DN2107_c0_g2 | Rola-high | -5.42 | 1.74e-05 | NA | NA | NA | | TRINITY_DN2107_c0_g2 | Venturia inaequalis |
| TRINITY_DN68_c0_g2 | Rola-high | -5.41 | 1.80e-05 | NA | NA | NA | | TRINITY_DN68_c0_g2 | — |
| TRINITY_DN11452_c1_g1 | Rin-high | 5.39 | 2.68e-03 | NA | NA | NA | | TRINITY_DN11452_c1_g1 | — |
| TRINITY_DN984_c0_g5 | Rola-high | -5.39 | 8.94e-03 | NA | NA | NA | | TRINITY_DN984_c0_g5 | Venturia inaequalis |
| TRINITY_DN37960_c0_g1 | Rin-high | 5.38 | 6.33e-03 | NA | NA | NA | | TRINITY_DN37960_c0_g1 | — |
| TRINITY_DN42341_c0_g1 | Rin-high | 5.38 | 3.57e-05 | NA | NA | NA | | TRINITY_DN42341_c0_g1 | — |
| TRINITY_DN53226_c0_g1 | Rola-high | -5.37 | 1.48e-04 | NA | NA | NA | | hypothetical protein MRB53_039979 | Venturia inaequalis |
| TRINITY_DN222110_c0_g1 | Rola-high | -5.37 | 5.47e-05 | NA | NA | NA | | isoprenylcysteine alpha-carbonyl methylesterase ICME-like | Venturia inaequalis |
| TRINITY_DN10247_c0_g1 | Rin-high | 5.35 | 2.78e-06 | NA | NA | NA | | TRINITY_DN10247_c0_g1 | Venturia inaequalis |
| TRINITY_DN9657_c0_g2 | Rin-high | 5.34 | 2.65e-03 | NA | NA | NA | | hypothetical protein DVH24_028819 | — |
| TRINITY_DN40716_c0_g1 | Rola-high | -5.34 | 1.25e-03 | NA | NA | NA | | TRINITY_DN40716_c0_g1 | Venturia inaequalis |
| TRINITY_DN42654_c1_g2 | Rin-high | 5.33 | 7.29e-04 | NA | NA | NA | | TRINITY_DN42654_c1_g2 | — |
| TRINITY_DN11203_c0_g1 | Rin-high | 5.33 | 2.97e-04 | NA | NA | NA | | unnamed protein product | Venturia inaequalis |
| TRINITY_DN16239_c1_g1 | Rola-high | -5.32 | 5.27e-04 | NA | NA | NA | | TRINITY_DN16239_c1_g1 | Venturia inaequalis |
| TRINITY_DN35762_c0_g1 | Rola-high | -5.31 | 3.76e-03 | NA | NA | NA | | TRINITY_DN35762_c0_g1 | — |
| TRINITY_DN22164_c1_g1 | Rola-high | -5.30 | 8.51e-03 | NA | NA | NA | | TRINITY_DN22164_c1_g1 | — |
| TRINITY_DN6179_c0_g1 | Rola-high | -5.28 | 4.50e-04 | NA | NA | NA | | TRINITY_DN6179_c0_g1 | Filobasidium wieringae |
| TRINITY_DN25087_c0_g1 | Rola-high | -5.28 | 2.73e-04 | NA | NA | NA | | hypothetical protein Dsin_033214 | Venturia inaequalis |
| TRINITY_DN140016_c0_g1 | Rola-high | -5.27 | 2.08e-03 | NA | NA | NA | | hypothetical protein DVH24_019539 | — |

| Trinity gene ID | Direction | Trinity | | GDDH13 protein match | | | Def. | Description | FCS-GX call |
|---|---|---|---|---|---|---|---|---|---|
| | | log2FC | padj | gene ID | log2FC | padj | | | |
| TRINITY_DN30339_c0_g1 | Rola-high | -5.27 | 8.28e-05 | NA | NA | NA | | TRINITY_DN30339_c0_g1 | Venturia inaequalis |
| TRINITY_DN1841_c2_g2 | Rin-high | 5.26 | 2.15e-04 | NA | NA | NA | | TRINITY_DN1841_c2_g2 | — |
| TRINITY_DN23498_c0_g1 | Rin-high | 5.24 | 3.31e-04 | NA | NA | NA | | hypothetical protein DVH24_006748 | — |
| TRINITY_DN222533_c0_g1 | Rola-high | -5.22 | 3.37e-06 | NA | NA | NA | | hypothetical protein BSKO_11484 | Venturia inaequalis |
| TRINITY_DN12444_c0_g1 | Rola-high | -5.22 | 9.34e-05 | NA | NA | NA | | TRINITY_DN12444_c0_g1 | Venturia inaequalis |
| TRINITY_DN32933_c0_g1 | Rola-high | -5.19 | 3.88e-03 | NA | NA | NA | | TRINITY_DN32933_c0_g1 | Venturia inaequalis |
| TRINITY_DN41918_c0_g1 | Rola-high | -5.18 | 1.93e-03 | NA | NA | NA | | TRINITY_DN41918_c0_g1 | Venturia inaequalis |
| TRINITY_DN81618_c0_g1 | Rola-high | -5.12 | 5.40e-08 | NA | NA | NA | | Tripeptidyl-peptidase sed2 | Venturia inaequalis |
| TRINITY_DN8348_c0_g1 | Rola-high | -5.09 | 2.09e-04 | NA | NA | NA | | TRINITY_DN8348_c0_g1 | Venturia inaequalis |
| TRINITY_DN182128_c0_g1 | Rola-high | -5.08 | 7.11e-05 | NA | NA | NA | | TRINITY_DN182128_c0_g1 | Venturia inaequalis |
| TRINITY_DN17512_c0_g2 | Rola-high | -5.08 | 8.67e-03 | NA | NA | NA | | TRINITY_DN17512_c0_g2 | Venturia inaequalis |
| TRINITY_DN17033_c0_g1 | Rola-high | -5.07 | 4.42e-07 | NA | NA | NA | | transcription factor PAR2-like | — |
| TRINITY_DN179701_c0_g1 | Rola-high | -5.06 | 2.45e-03 | NA | NA | NA | | uncharacterized protein LOC9640874 isoform X2 | Venturia inaequalis |
| TRINITY_DN23834_c0_g2 | Rola-high | -5.05 | 4.56e-03 | NA | NA | NA | | TRINITY_DN23834_c0_g2 | Venturia inaequalis |
| TRINITY_DN6425_c0_g1 | Rola-high | -5.04 | 2.97e-03 | NA | NA | NA | | TRINITY_DN6425_c0_g1 | Venturia inaequalis |
| TRINITY_DN121472_c0_g1 | Rola-high | -5.04 | 8.41e-04 | NA | NA | NA | | TRINITY_DN121472_c0_g1 | Venturia inaequalis |
| TRINITY_DN71684_c0_g1 | Rin-high | 5.03 | 4.99e-03 | NA | NA | NA | | TRINITY_DN71684_c0_g1 | — |
| TRINITY_DN29202_c0_g1 | Rin-high | 5.01 | 1.68e-05 | NA | NA | NA | | TRINITY_DN29202_c0_g1 | — |
| TRINITY_DN20128_c0_g1 | Rola-high | -5.01 | 1.82e-03 | NA | NA | NA | | TRINITY_DN20128_c0_g1 | Venturia inaequalis |

| Trinity gene ID | Direction | Trinity | | GDDH13 protein match | | | Def. | Description | FCS-GX |
|---|---|---|---|---|---|---|---|---|---|
| | | log2FC | padj | gene ID | log2FC | padj | | | call |
| TRINITY_DN96438_c0_g1 | Rola-high | -5.01 | 9.16e-03 | NA | NA | NA | | fasciclin-like protein | Venturia inaequalis |
| TRINITY_DN86037_c0_g1 | Rola-high | -5.00 | 2.75e-03 | NA | NA | NA | | hypothetical protein | Venturia inaequalis |
| TRINITY_DN138952_c0_g1 | Rola-high | -5.00 | 3.21e-03 | NA | NA | NA | | unnamed protein product | Venturia aucupariae |
| TRINITY_DN157530_c0_g1 | Rola-high | -5.00 | 4.99e-03 | NA | NA | NA | | TRINITY_DN157530_c0_g1 | Venturia inaequalis |
| TRINITY_DN14188_c0_g1 | Rola-high | -4.99 | 6.40e-05 | NA | NA | NA | | TRINITY_DN14188_c0_g1 | Venturia inaequalis |
| TRINITY_DN9397_c0_g1 | Rola-high | -4.99 | 9.76e-03 | NA | NA | NA | | TRINITY_DN9397_c0_g1 | Venturia inaequalis |
| TRINITY_DN6611_c2_g1 | Rola-high | -4.99 | 2.04e-03 | NA | NA | NA | | TRINITY_DN6611_c2_g1 | — |
| TRINITY_DN15054_c1_g2 | Rola-high | -4.99 | 2.15e-03 | NA | NA | NA | | TRINITY_DN15054_c1_g2 | — |
| TRINITY_DN135314_c0_g1 | Rola-high | -4.99 | 1.59e-03 | NA | NA | NA | | TRINITY_DN135314_c0_g1 | Venturia inaequalis |
| TRINITY_DN12157_c0_g1 | Rola-high | -4.98 | 5.95e-03 | NA | NA | NA | | TRINITY_DN12157_c0_g1 | Venturia inaequalis |
| TRINITY_DN14288_c0_g2 | Rola-high | -4.98 | 1.08e-03 | NA | NA | NA | | TRINITY_DN14288_c0_g2 | Venturia inaequalis |
| TRINITY_DN33242_c0_g1 | Rin-high | 4.98 | 1.34e-04 | NA | NA | NA | | hypothetical protein DVH24_039675 | — |
| TRINITY_DN114659_c0_g1 | Rola-high | -4.97 | 3.20e-03 | NA | NA | NA | | TRINITY_DN114659_c0_g1 | Venturia inaequalis |
| TRINITY_DN93979_c0_g1 | Rola-high | -4.97 | 1.74e-04 | NA | NA | NA | | TRINITY_DN93979_c0_g1 | — |
| TRINITY_DN43296_c0_g1 | Rola-high | -4.96 | 2.25e-05 | NA | NA | NA | | TRINITY_DN43296_c0_g1 | Venturia inaequalis |
| TRINITY_DN118460_c0_g1 | Rola-high | -4.96 | 1.74e-04 | NA | NA | NA | | probable polyamine transporter At1g31830 isoform X2 | Venturia inaequalis |
| TRINITY_DN5379_c0_g1 | Rola-high | -4.93 | 3.46e-04 | NA | NA | NA | | TRINITY_DN5379_c0_g1 | Venturia inaequalis |
| TRINITY_DN72248_c0_g1 | Rin-high | 4.92 | 5.33e-07 | NA | NA | NA | | TRINITY_DN72248_c0_g1 | — |
| TRINITY_DN34242_c0_g1 | Rin-high | 4.91 | 2.08e-03 | NA | NA | NA | | TRINITY_DN34242_c0_g1 | Venturia inaequalis |

| Trinity gene ID | Direction | Trinity | | GDDH13 protein match | | | Def. | Description | FCS-GX |
|---|---|---|---|---|---|---|---|---|---|
| | | log2FC | padj | gene ID | log2FC | padj | | | call |
| TRINITY_DN135679_c0_g1 | Rola-high | -4.89 | 6.76e-03 | NA | NA | NA | | TRINITY_DN135679_c0_g1 | Venturia inaequalis |
| TRINITY_DN19504_c1_g2 | Rin-high | 4.89 | 4.98e-04 | NA | NA | NA | | hypothetical protein LUZ63_022753 | Kosakonia radicincitans |
| TRINITY_DN36482_c0_g1 | Rola-high | -4.89 | 6.20e-04 | NA | NA | NA | | hypothetical protein R6Q59_010223 | Venturia inaequalis |
| TRINITY_DN6385_c0_g1 | Rola-high | -4.88 | 2.78e-03 | NA | NA | NA | | TRINITY_DN6385_c0_g1 | Venturia inaequalis |
| TRINITY_DN24001_c0_g1 | Rola-high | -4.88 | 8.62e-03 | NA | NA | NA | | TRINITY_DN24001_c0_g1 | Venturia inaequalis |
| TRINITY_DN14406_c0_g1 | Rola-high | -4.88 | 2.64e-03 | NA | NA | NA | | TRINITY_DN14406_c0_g1 | Venturia inaequalis |
| TRINITY_DN223702_c0_g1 | Rola-high | -4.87 | 1.28e-03 | NA | NA | NA | | hypothetical protein MRB53_040606 | Venturia inaequalis |
| TRINITY_DN12843_c0_g1 | Rola-high | -4.85 | 1.22e-03 | NA | NA | NA | | TRINITY_DN12843_c0_g1 | — |
| TRINITY_DN46686_c0_g1 | Rola-high | -4.85 | 9.49e-04 | NA | NA | NA | | hypothetical protein DVH24_032073 | — |
| TRINITY_DN13169_c0_g1 | Rin-high | 4.82 | 1.73e-07 | NA | NA | NA | | TRINITY_DN13169_c0_g1 | Venturia inaequalis |
| TRINITY_DN17501_c0_g2 | Rola-high | -4.82 | 1.70e-04 | NA | NA | NA | | TRINITY_DN17501_c0_g2 | Venturia inaequalis |
| TRINITY_DN26258_c0_g1 | Rola-high | -4.80 | 3.15e-04 | NA | NA | NA | | TRINITY_DN26258_c0_g1 | Venturia inaequalis |
| TRINITY_DN15853_c0_g1 | Rola-high | -4.78 | 6.29e-03 | NA | NA | NA | | TRINITY_DN15853_c0_g1 | Venturia inaequalis |
| TRINITY_DN17444_c0_g1 | Rola-high | -4.77 | 3.10e-03 | NA | NA | NA | | hypothetical protein ZWY2020_055871 | Venturia inaequalis |
| TRINITY_DN7231_c1_g1 | Rola-high | -4.77 | 9.05e-03 | NA | NA | NA | | TRINITY_DN7231_c1_g1 | — |
| TRINITY_DN24620_c0_g1 | Rola-high | -4.75 | 7.30e-06 | NA | NA | NA | | predicted protein | Venturia inaequalis |
| TRINITY_DN8690_c0_g2 | Rola-high | -4.74 | 7.14e-03 | NA | NA | NA | | TRINITY_DN8690_c0_g2 | Venturia inaequalis |
| TRINITY_DN10716_c0_g1 | Rola-high | -4.74 | 1.32e-04 | NA | NA | NA | | TRINITY_DN10716_c0_g1 | — |
| TRINITY_DN45506_c0_g1 | Rola-high | -4.74 | 9.86e-03 | NA | NA | NA | | unnamed protein product | Venturia inaequalis |

| Trinity gene ID | Direction | Trinity | | GDDH13 protein match | | | Def. Description | FCS-GX |
|---|---|---|---|---|---|---|---|---|
| | | log2FC | padj | gene ID | log2FC | padj | | call |
| TRINITY_DN34280_c1_g1 | Rola-high | -4.73 | 3.68e-03 | NA | NA | NA | TRINITY_DN34280_c1_g1 | Venturia inaequalis |
| TRINITY_DN35835_c0_g1 | Rola-high | -4.73 | 8.28e-03 | NA | NA | NA | unnamed protein product | — |
| TRINITY_DN33001_c0_g2 | Rin-high | 4.72 | 2.22e-04 | NA | NA | NA | hypothetical protein C1H46_030890 | — |
| TRINITY_DN31386_c0_g1 | Rola-high | -4.72 | 2.45e-03 | NA | NA | NA | TRINITY_DN31386_c0_g1 | Venturia inaequalis |
| TRINITY_DN43248_c2_g1 | Rola-high | -4.71 | 8.29e-04 | NA | NA | NA | TRINITY_DN43248_c2_g1 | — |
| TRINITY_DN202568_c0_g1 | Rola-high | -4.70 | 2.27e-03 | NA | NA | NA | hypothetical protein BSKO_11082 | Venturia inaequalis |
| TRINITY_DN43661_c0_g1 | Rola-high | -4.69 | 4.78e-05 | NA | NA | NA | Eukaryotic aspartyl protease family protein | Venturia inaequalis |
| TRINITY_DN31469_c0_g1 | Rola-high | -4.68 | 7.31e-04 | NA | NA | NA | hypothetical protein C1H46_040980 | — |
| TRINITY_DN44515_c0_g2 | Rola-high | -4.67 | 2.66e-03 | NA | NA | NA | unnamed protein product | Venturia inaequalis |
| TRINITY_DN22249_c0_g1 | Rola-high | -4.67 | 1.21e-03 | NA | NA | NA | probable homogentisate phytyltransferase 1, chloroplastic isoform X2 | Venturia inaequalis |
| TRINITY_DN39485_c0_g1 | Rola-high | -4.66 | 2.22e-04 | NA | NA | NA | 36.4 kDa proline-rich protein-like | — |
| TRINITY_DN1152_c0_g2 | Rola-high | -4.65 | 2.28e-04 | NA | NA | NA | hypothetical protein DVH24_039711 | — |
| TRINITY_DN13323_c0_g1 | Rola-high | -4.64 | 1.34e-04 | NA | NA | NA | TRINITY_DN13323_c0_g1 | Venturia inaequalis |
| TRINITY_DN15273_c1_g1 | Rola-high | -4.64 | 8.37e-03 | NA | NA | NA | TRINITY_DN15273_c1_g1 | — |
| TRINITY_DN49199_c0_g1 | Rin-high | 4.63 | 1.94e-03 | NA | NA | NA | MADS-box transcription factor 14-like | — |
| TRINITY_DN5924_c2_g1 | Rin-high | 4.62 | 6.65e-03 | NA | NA | NA | TRINITY_DN5924_c2_g1 | — |
| TRINITY_DN20147_c4_g1 | Rin-high | 4.57 | 2.00e-03 | NA | NA | NA | TRINITY_DN20147_c4_g1 | — |
| TRINITY_DN80318_c0_g1 | Rola-high | -4.56 | 8.53e-04 | NA | NA | NA | unnamed protein product | Venturia inaequalis |
| TRINITY_DN64475_c0_g1 | Rola-high | -4.55 | 2.01e-03 | NA | NA | NA | TRINITY_DN64475_c0_g1 | Venturia inaequalis |
| TRINITY_DN31124_c0_g1 | Rola-high | -4.54 | 4.15e-03 | NA | NA | NA | TRINITY_DN31124_c0_g1 | Venturia inaequalis |

| Trinity gene ID | Direction | Trinity |  | GDDH13 protein match |  |  | Def. | Description | FCS-GX call |
|---|---|---|---|---|---|---|---|---|---|
|  |  | log2FC | padj | gene ID | log2FC | padj |  |  |  |
| TRINITY_DN21956_c1_g1 | Rola-high | -4.54 | 8.26e-06 | NA | NA | NA |  | TRINITY_DN21956_c1_g1 | — |
| TRINITY_DN7940_c4_g1 | Rola-high | -4.54 | 2.33e-03 | NA | NA | NA |  | TRINITY_DN7940_c4_g1 | — |
| TRINITY_DN39384_c0_g1 | Rola-high | -4.54 | 4.49e-03 | NA | NA | NA |  | TRINITY_DN39384_c0_g1 | Venturia inaequalis |
| TRINITY_DN33186_c0_g1 | Rola-high | -4.53 | 1.38e-03 | NA | NA | NA |  | hypothetical protein R6Q59_009818 | Venturia inaequalis |
| TRINITY_DN10733_c0_g1 | Rola-high | -4.53 | 8.37e-03 | NA | NA | NA |  | probable indole-3-pyruvate monooxygenase YUCCA10 | Venturia inaequalis |
| TRINITY_DN15317_c0_g1 | Rola-high | -4.52 | 1.62e-06 | NA | NA | NA |  | Chitin-binding lectin 1 | Venturia inaequalis |
| TRINITY_DN14676_c0_g1 | Rola-high | -4.48 | 3.15e-04 | NA | NA | NA |  | hypothetical protein CBR_g51827 | Venturia inaequalis |
| TRINITY_DN14288_c0_g1 | Rola-high | -4.48 | 3.70e-03 | NA | NA | NA |  | TRINITY_DN14288_c0_g1 | Venturia inaequalis |
| TRINITY_DN13214_c0_g1 | Rola-high | -4.46 | 5.52e-05 | NA | NA | NA |  | TRINITY_DN13214_c0_g1 | — |
| TRINITY_DN3144_c2_g1 | Rola-high | -4.44 | 9.61e-04 | NA | NA | NA |  | TRINITY_DN3144_c2_g1 | — |
| TRINITY_DN12436_c0_g1 | Rola-high | -4.43 | 1.63e-05 | NA | NA | NA |  | TRINITY_DN12436_c0_g1 | Venturia inaequalis |
| TRINITY_DN927_c2_g1 | Rola-high | -4.43 | 9.02e-03 | NA | NA | NA |  | TRINITY_DN927_c2_g1 | Venturia inaequalis |
| TRINITY_DN181680_c0_g1 | Rola-high | -4.42 | 3.28e-03 | NA | NA | NA |  | TRINITY_DN181680_c0_g1 | Venturia inaequalis |
| TRINITY_DN23218_c0_g1 | Rola-high | -4.40 | 1.58e-03 | NA | NA | NA |  | hypothetical protein DVH24_040097 | — |
| TRINITY_DN3907_c15_g1 | Rin-high | 4.40 | 2.46e-07 | NA | NA | NA |  | TRINITY_DN3907_c15_g1 | — |
| TRINITY_DN19301_c0_g1 | Rola-high | -4.38 | 2.85e-06 | NA | NA | NA |  | asparagine synthetase [glutamine-hydrolyzing] 2 | Venturia inaequalis |
| TRINITY_DN25082_c0_g1 | Rola-high | -4.38 | 1.21e-03 | NA | NA | NA |  | hypothetical protein BT93_L5570 | Venturia inaequalis |
| TRINITY_DN40485_c0_g3 | Rola-high | -4.37 | 9.00e-04 | NA | NA | NA |  | TRINITY_DN40485_c0_g3 | — |
| TRINITY_DN6241_c1_g1 | Rola-high | -4.37 | 4.71e-05 | NA | NA | NA |  | TRINITY_DN6241_c1_g1 | — |
| TRINITY_DN4608_c0_g2 | Rin-high | 4.36 | 4.70e-03 | NA | NA | NA |  | TRINITY_DN4608_c0_g2 | — |

| Trinity gene ID | Direction | Trinity | | GDDH13 protein match | | | Def. | Description | FCS-GX |
|---|---|---|---|---|---|---|---|---|---|
| | | log2FC | padj | gene ID | log2FC | padj | | | call |
| TRINITY_DN11375_c0_g1 | Rin-high | 4.36 | 1.14e-03 | NA | NA | NA | | TRINITY_DN11375_c0_g1 | Venturia inaequalis |
| TRINITY_DN7811_c4_g1 | Rola-high | -4.36 | 3.08e-05 | NA | NA | NA | | TRINITY_DN7811_c4_g1 | — |
| TRINITY_DN30771_c0_g1 | Rola-high | -4.34 | 1.71e-03 | NA | NA | NA | | hypothetical protein MRB53_036866 | Venturia inaequalis |
| TRINITY_DN5187_c0_g5 | Rola-high | -4.34 | 2.61e-04 | NA | NA | NA | | TRINITY_DN5187_c0_g5 | — |
| TRINITY_DN204484_c0_g1 | Rola-high | -4.32 | 2.85e-03 | NA | NA | NA | | metal-independent alpha-mannosidase | Venturia inaequalis |
| TRINITY_DN116075_c0_g1 | Rola-high | -4.30 | 1.90e-03 | NA | NA | NA | | TRINITY_DN116075_c0_g1 | Venturia inaequalis |
| TRINITY_DN8374_c1_g1 | Rola-high | -4.30 | 5.00e-03 | NA | NA | NA | | hypothetical protein DVH24_028635 | — |
| TRINITY_DN8848_c2_g1 | Rola-high | -4.29 | 5.97e-03 | NA | NA | NA | | TRINITY_DN8848_c2_g1 | — |
| TRINITY_DN136086_c0_g1 | Rola-high | -4.29 | 1.03e-03 | NA | NA | NA | | TRINITY_DN136086_c0_g1 | Venturia inaequalis |
| TRINITY_DN2203_c0_g1 | Rin-high | 4.28 | 5.09e-08 | NA | NA | NA | | TRINITY_DN2203_c0_g1 | Venturia inaequalis |
| TRINITY_DN11829_c0_g2 | Rola-high | -4.28 | 7.66e-03 | NA | NA | NA | | TRINITY_DN11829_c0_g2 | Venturia inaequalis |
| TRINITY_DN204125_c0_g1 | Rola-high | -4.28 | 7.88e-03 | NA | NA | NA | | TRINITY_DN204125_c0_g1 | Venturia inaequalis |
| TRINITY_DN16602_c0_g1 | Rola-high | -4.27 | 6.35e-03 | NA | NA | NA | | TRINITY_DN16602_c0_g1 | Venturia inaequalis |
| TRINITY_DN25451_c0_g1 | Rin-high | 4.27 | 9.88e-03 | NA | NA | NA | | acyltransferase Pun1-like | — |
| TRINITY_DN40542_c0_g1 | Rola-high | -4.26 | 9.02e-03 | NA | NA | NA | | Protein kri1 | Venturia inaequalis |
| TRINITY_DN223182_c0_g1 | Rola-high | -4.26 | 5.02e-03 | NA | NA | NA | | autophagy protein 22 | Venturia inaequalis |
| TRINITY_DN18380_c0_g1 | Rola-high | -4.24 | 7.25e-03 | NA | NA | NA | | TRINITY_DN18380_c0_g1 | Venturia inaequalis |
| TRINITY_DN2956_c3_g2 | Rola-high | -4.24 | 3.56e-03 | NA | NA | NA | | TRINITY_DN2956_c3_g2 | — |
| TRINITY_DN138107_c0_g1 | Rola-high | -4.24 | 7.73e-03 | NA | NA | NA | | TRINITY_DN138107_c0_g1 | Venturia inaequalis |

| Trinity gene ID | Direction | Trinity | | GDDH13 protein match | | | Def. | Description | FCS-GX |
|---|---|---|---|---|---|---|---|---|---|
| | | log2FC | padj | gene ID | log2FC | padj | | | call |
| TRINITY_DN44745_c1_g1 | Rola-high | -4.24 | 6.19e-03 | NA | NA | NA | | hypothetical protein DVH24_017788, partial | — |
| TRINITY_DN44974_c0_g1 | Rin-high | 4.22 | 1.36e-03 | NA | NA | NA | | TRINITY_DN44974_c0_g1 | Venturia inaequalis |
| TRINITY_DN182971_c0_g1 | Rola-high | -4.22 | 6.43e-04 | NA | NA | NA | | S-methylmethionine permease | Venturia inaequalis |
| TRINITY_DN17409_c0_g3 | Rola-high | -4.22 | 1.81e-04 | NA | NA | NA | | transcription factor PAR2-like | — |
| TRINITY_DN24508_c0_g4 | Rin-high | 4.22 | 1.85e-03 | NA | NA | NA | | hypothetical protein FEM48_ZijujMtG0001300 | — |
| TRINITY_DN40485_c0_g1 | Rola-high | -4.21 | 8.02e-04 | NA | NA | NA | | uncharacterized protein | — |
| TRINITY_DN11737_c0_g1 | Rola-high | -4.21 | 4.80e-04 | NA | NA | NA | | TRINITY_DN11737_c0_g1 | Venturia inaequalis |
| TRINITY_DN12717_c0_g1 | Rola-high | -4.21 | 5.65e-04 | NA | NA | NA | | TRINITY_DN12717_c0_g1 | Venturia inaequalis |
| TRINITY_DN22239_c0_g1 | Rola-high | -4.21 | 1.98e-04 | NA | NA | NA | | hypothetical protein WJX74_004877 | Venturia inaequalis |
| TRINITY_DN114087_c0_g1 | Rola-high | -4.20 | 6.38e-03 | NA | NA | NA | | hypothetical protein | Venturia inaequalis |
| TRINITY_DN13230_c0_g1 | Rin-high | 4.17 | 1.01e-04 | NA | NA | NA | | TRINITY_DN13230_c0_g1 | Venturia inaequalis |
| TRINITY_DN34121_c0_g1 | Rola-high | -4.06 | 4.24e-06 | NA | NA | NA | | TRINITY_DN34121_c0_g1 | Venturia inaequalis |
| TRINITY_DN15738_c0_g1 | Rola-high | -4.04 | 3.58e-03 | NA | NA | NA | | TRINITY_DN15738_c0_g1 | Venturia inaequalis |
| TRINITY_DN203436_c0_g1 | Rola-high | -4.02 | 4.26e-03 | NA | NA | NA | | TRINITY_DN203436_c0_g1 | Venturia inaequalis |
| TRINITY_DN41699_c3_g1 | Rin-high | 4.00 | 1.44e-03 | NA | NA | NA | | hypothetical protein DVH24_032959 | — |
| TRINITY_DN21841_c0_g1 | Rola-high | -3.99 | 6.84e-03 | NA | NA | NA | | TRINITY_DN21841_c0_g1 | Venturia inaequalis |
| TRINITY_DN184819_c0_g1 | Rola-high | -3.98 | 3.84e-04 | NA | NA | NA | | hypothetical protein KP509_35G012400 | Venturia inaequalis |
| TRINITY_DN18686_c0_g1 | Rola-high | -3.97 | 7.94e-03 | NA | NA | NA | | TRINITY_DN18686_c0_g1 | Venturia inaequalis |
| TRINITY_DN161413_c0_g1 | Rola-high | -3.97 | 5.93e-03 | NA | NA | NA | | nitrilotriacetate monooxygenase component a | Venturia inaequalis |

| Trinity gene ID | Direction | Trinity | | GDDH13 protein match | | | Def. | Description | FCS-GX |
|---|---|---|---|---|---|---|---|---|---|
| | | log2FC | padj | gene ID | log2FC | padj | | | call |
| TRINITY_DN156689_c0_g1 | Rola-high | -3.97 | 5.00e-03 | NA | NA | NA | | TRINITY_DN156689_c0_g1 | Venturia inaequalis |
| TRINITY_DN223477_c0_g1 | Rola-high | -3.97 | 5.14e-04 | NA | NA | NA | | TRINITY_DN223477_c0_g1 | Venturia inaequalis |
| TRINITY_DN18658_c0_g1 | Rola-high | -3.96 | 8.83e-05 | NA | NA | NA | | predicted protein | Venturia inaequalis |
| TRINITY_DN24550_c0_g1 | Rola-high | -3.96 | 7.27e-05 | NA | NA | NA | | TRINITY_DN24550_c0_g1 | Venturia inaequalis |
| TRINITY_DN10297_c0_g1 | Rola-high | -3.96 | 5.83e-03 | NA | NA | NA | | TRINITY_DN10297_c0_g1 | Venturia inaequalis |
| TRINITY_DN179261_c0_g1 | Rola-high | -3.96 | 7.88e-03 | NA | NA | NA | | arabinogalactan endo-beta-1,4-galactanase | Venturia inaequalis |
| TRINITY_DN10975_c3_g1 | Rola-high | -3.95 | 5.77e-07 | NA | NA | NA | | TRINITY_DN10975_c3_g1 | — |
| TRINITY_DN159629_c0_g1 | Rin-high | 3.95 | 7.97e-04 | NA | NA | NA | | TRINITY_DN159629_c0_g1 | Venturia inaequalis |
| TRINITY_DN27539_c0_g1 | Rola-high | -3.95 | 2.24e-06 | NA | NA | NA | | ATP synthase F0 subunit 9 | Venturia inaequalis |
| TRINITY_DN30083_c0_g1 | Rola-high | -3.94 | 2.65e-05 | NA | NA | NA | | TRINITY_DN30083_c0_g1 | — |
| TRINITY_DN15770_c0_g1 | Rola-high | -3.94 | 1.66e-04 | NA | NA | NA | | unnamed protein product | Venturia inaequalis |
| TRINITY_DN200371_c0_g1 | Rola-high | -3.93 | 4.33e-05 | NA | NA | NA | | hypothetical protein R6Q59_009932 | Venturia inaequalis |
| TRINITY_DN30134_c0_g1 | Rola-high | -3.92 | 6.97e-03 | NA | NA | NA | | threonine synthase | Venturia inaequalis |
| TRINITY_DN31569_c0_g1 | Rola-high | -3.91 | 1.88e-05 | NA | NA | NA | | TRINITY_DN31569_c0_g1 | Venturia inaequalis |
| TRINITY_DN70055_c1_g1 | Rin-high | 3.91 | 7.84e-03 | NA | NA | NA | | TRINITY_DN70055_c1_g1 | — |
| TRINITY_DN62087_c0_g1 | Rin-high | 3.91 | 7.48e-06 | NA | NA | NA | | tryptophan aminotransferase-related protein 4-like | — |
| TRINITY_DN13062_c0_g1 | Rin-high | 3.91 | 1.81e-04 | NA | NA | NA | | alcohol oxidase | Venturia inaequalis |
| TRINITY_DN183004_c0_g1 | Rola-high | -3.90 | 9.15e-03 | NA | NA | NA | | predicted protein | Venturia inaequalis |
| TRINITY_DN15790_c0_g1 | Rin-high | 3.90 | 5.99e-03 | NA | NA | NA | | TRINITY_DN15790_c0_g1 | — |

| Trinity gene ID | Direction | Trinity | | GDDH13 protein match | | | Def. | Description | FCS-GX |
|---|---|---|---|---|---|---|---|---|---|
| | | log2FC | padj | gene ID | log2FC | padj | | | call |
| TRINITY_DN72763_c0_g1 | Rin-high | 3.89 | 7.86e-05 | NA | NA | NA | | TRINITY_DN72763_c0_g1 | — |
| TRINITY_DN11382_c0_g1 | Rola-high | -3.89 | 1.13e-03 | NA | NA | NA | | TRINITY_DN11382_c0_g1 | Venturia inaequalis |
| TRINITY_DN389_c0_g2 | Rola-high | -3.82 | 3.77e-03 | NA | NA | NA | | TRINITY_DN389_c0_g2 | — |
| TRINITY_DN37097_c0_g1 | Rola-high | -3.81 | 5.53e-03 | NA | NA | NA | | TRINITY_DN37097_c0_g1 | Venturia inaequalis |
| TRINITY_DN19729_c0_g1 | Rin-high | 3.80 | 2.33e-03 | NA | NA | NA | | TRINITY_DN19729_c0_g1 | — |
| TRINITY_DN32232_c0_g1 | Rola-high | -3.80 | 4.26e-03 | NA | NA | NA | | TRINITY_DN32232_c0_g1 | Venturia inaequalis |
| TRINITY_DN73516_c1_g1 | Rin-high | 3.80 | 7.43e-03 | NA | NA | NA | | TRINITY_DN73516_c1_g1 | — |
| TRINITY_DN16620_c0_g1 | Rola-high | -3.80 | 2.99e-04 | NA | NA | NA | | putative glycosidase | Venturia inaequalis |
| TRINITY_DN40927_c0_g2 | Rola-high | -3.80 | 1.13e-04 | NA | NA | NA | | hypothetical protein DVH24_006799 | — |
| TRINITY_DN10024_c0_g1 | Rola-high | -3.78 | 8.94e-03 | NA | NA | NA | | TRINITY_DN10024_c0_g1 | Venturia inaequalis |
| TRINITY_DN229050_c0_g1 | Rola-high | -3.77 | 7.68e-03 | NA | NA | NA | | Chain 1R, Cytochrome b-c1 complex subunit 7 | Venturia inaequalis |
| TRINITY_DN8408_c0_g1 | Rola-high | -3.77 | 9.25e-05 | NA | NA | NA | | endothelin-converting enzyme | Venturia inaequalis |
| TRINITY_DN34915_c0_g1 | Rola-high | -3.76 | 4.59e-03 | NA | NA | NA | | TATA-binding protein-associated factor 2N-like | Venturia inaequalis |
| TRINITY_DN42942_c0_g1 | Rola-high | -3.74 | 6.33e-03 | NA | NA | NA | | beta-galactosidase 17 | Venturia inaequalis |
| TRINITY_DN8910_c0_g1 | Rin-high | 3.73 | 7.69e-04 | NA | NA | NA | | hypothetical protein GBA52_014927 | — |
| TRINITY_DN922_c2_g1 | Rin-high | 3.73 | 6.95e-03 | NA | NA | NA | | TRINITY_DN922_c2_g1 | — |
| TRINITY_DN14862_c1_g2 | Rola-high | -3.72 | 8.69e-04 | NA | NA | NA | | TRINITY_DN14862_c1_g2 | — |
| TRINITY_DN5464_c0_g1 | Rola-high | -3.72 | 3.55e-04 | NA | NA | NA | | TRINITY_DN5464_c0_g1 | Venturia inaequalis |
| TRINITY_DN25585_c0_g2 | Rola-high | -3.70 | 8.21e-03 | NA | NA | NA | | TRINITY_DN25585_c0_g2 | Venturia inaequalis |
| TRINITY_DN10462_c3_g1 | Rin-high | 3.69 | 1.65e-03 | NA | NA | NA | | subtilisin-like protease SBT3.12 | n/a |

| Trinity gene ID | Direction | Trinity | | GDDH13 protein match | | | Def. | Description | FCS-GX |
|---|---|---|---|---|---|---|---|---|---|
| | | log2FC | padj | gene ID | log2FC | padj | | | call |
| TRINITY_DN119461_c0_g1 | Rola-high | -3.69 | 5.56e-03 | NA | NA | NA | | hypothetical protein BT93_L4724 | Venturia inaequalis |
| TRINITY_DN47738_c0_g1 | Rola-high | -3.68 | 1.85e-03 | NA | NA | NA | | hypothetical protein FH972_024069 | Venturia inaequalis |
| TRINITY_DN19725_c0_g1 | Rola-high | -3.68 | 7.33e-04 | NA | NA | NA | | TRINITY_DN19725_c0_g1 | — |
| TRINITY_DN185052_c0_g1 | Rola-high | -3.67 | 8.22e-03 | NA | NA | NA | | TRINITY_DN185052_c0_g1 | Venturia inaequalis |
| TRINITY_DN157159_c0_g1 | Rola-high | -3.66 | 2.68e-03 | NA | NA | NA | | hypothetical protein FH972_023951 | Venturia inaequalis |
| TRINITY_DN7914_c0_g1 | Rola-high | -3.66 | 5.63e-03 | NA | NA | NA | | unnamed protein product | Venturia inaequalis |
| TRINITY_DN19448_c0_g2 | Rola-high | -3.66 | 1.62e-04 | NA | NA | NA | | TRINITY_DN19448_c0_g2 | — |
| TRINITY_DN25848_c0_g1 | Rola-high | -3.65 | 3.15e-04 | NA | NA | NA | | TRINITY_DN25848_c0_g1 | Venturia inaequalis |
| TRINITY_DN44713_c0_g1 | Rin-high | 3.65 | 5.47e-03 | NA | NA | NA | | TRINITY_DN44713_c0_g1 | Venturia inaequalis |
| TRINITY_DN52657_c0_g1 | Rola-high | -3.64 | 2.41e-03 | NA | NA | NA | | cysteine dioxygenase | Venturia inaequalis |
| TRINITY_DN3229_c4_g1 | Rola-high | -3.63 | 4.45e-03 | NA | NA | NA | | hypothetical protein DVH24_016031 | — |
| TRINITY_DN2206_c1_g1 | Rola-high | -3.63 | 6.21e-03 | NA | NA | NA | | TRINITY_DN2206_c1_g1 | — |
| TRINITY_DN15777_c1_g1 | Rin-high | 3.63 | 1.98e-03 | NA | NA | NA | | hypothetical protein DVH24_010417 | — |
| TRINITY_DN158704_c0_g1 | Rola-high | -3.62 | 2.65e-03 | NA | NA | NA | | hypothetical protein MRB53_040252 | Venturia inaequalis |
| TRINITY_DN2348_c0_g2 | Rin-high | 3.61 | 4.03e-03 | NA | NA | NA | | hypothetical protein C1H46_043555 | — |
| TRINITY_DN15817_c0_g1 | Rola-high | -3.60 | 3.62e-03 | NA | NA | NA | | predicted protein | Venturia inaequalis |
| TRINITY_DN5088_c0_g1 | Rin-high | 3.60 | 6.48e-03 | NA | NA | NA | | TRINITY_DN5088_c0_g1 | Venturia inaequalis |
| TRINITY_DN18944_c0_g2 | Rola-high | -3.60 | 6.14e-03 | NA | NA | NA | | TRINITY_DN18944_c0_g2 | Venturia inaequalis |
| TRINITY_DN18269_c0_g1 | Rola-high | -3.57 | 2.93e-04 | NA | NA | NA | | TRINITY_DN18269_c0_g1 | — |

| Trinity gene ID | Direction | Trinity | | GDDH13 protein match | | | Def. | Description | FCS-GX call |
|---|---|---|---|---|---|---|---|---|---|
| | | log2FC | padj | gene ID | log2FC | padj | | | |
| TRINITY_DN4310_c0_g1 | Rin-high | 3.56 | 1.49e-07 | NA | NA | NA | | uncharacterized protein | — |
| TRINITY_DN180618_c0_g1 | Rola-high | -3.54 | 7.50e-03 | NA | NA | NA | | TRINITY_DN180618_c0_g1 | Venturia inaequalis |
| TRINITY_DN36891_c0_g1 | Rola-high | -3.53 | 8.73e-03 | NA | NA | NA | | TRINITY_DN36891_c0_g1 | Venturia inaequalis |
| TRINITY_DN7776_c1_g2 | Rola-high | -3.52 | 7.22e-03 | NA | NA | NA | | protein ALP1-like | — |
| TRINITY_DN26968_c0_g1 | Rin-high | 3.51 | 9.13e-03 | NA | NA | NA | | TRINITY_DN26968_c0_g1 | Venturia inaequalis |
| TRINITY_DN1089_c0_g2 | Rola-high | -3.51 | 5.00e-03 | NA | NA | NA | | TRINITY_DN1089_c0_g2 | — |
| TRINITY_DN1697_c0_g2 | Rola-high | -3.50 | 7.09e-03 | NA | NA | NA | | TRINITY_DN1697_c0_g2 | Venturia inaequalis |
| TRINITY_DN19154_c0_g1 | Rin-high | 3.50 | 4.51e-05 | NA | NA | NA | | TRINITY_DN19154_c0_g1 | — |
| TRINITY_DN10720_c0_g1 | Rola-high | -3.49 | 2.65e-03 | NA | NA | NA | | hypothetical protein MRB53_039381 | Venturia inaequalis |
| TRINITY_DN4384_c0_g1 | Rola-high | -3.48 | 4.88e-05 | NA | NA | NA | | TRINITY_DN4384_c0_g1 | — |
| TRINITY_DN7018_c0_g2 | Rola-high | -3.47 | 2.71e-05 | NA | NA | NA | | TRINITY_DN7018_c0_g2 | — |
| TRINITY_DN2316_c2_g1 | Rola-high | -3.47 | 4.19e-03 | NA | NA | NA | | TRINITY_DN2316_c2_g1 | — |
| TRINITY_DN91906_c0_g1 | Rola-high | -3.46 | 7.49e-03 | NA | NA | NA | | TRINITY_DN91906_c0_g1 | Venturia inaequalis |
| TRINITY_DN6300_c1_g1 | Rin-high | 3.46 | 4.49e-03 | NA | NA | NA | | hypothetical protein DVH24_004300 | — |
| TRINITY_DN114313_c0_g1 | Rola-high | -3.45 | 4.40e-03 | NA | NA | NA | | TRINITY_DN114313_c0_g1 | Venturia inaequalis |
| TRINITY_DN9201_c1_g1 | Rin-high | 3.43 | 2.75e-03 | NA | NA | NA | | TRINITY_DN9201_c1_g1 | — |
| TRINITY_DN36311_c0_g1 | Rola-high | -3.41 | 9.88e-03 | NA | NA | NA | | nuclear pore complex protein NUP160 | Venturia inaequalis |
| TRINITY_DN79927_c0_g1 | Rola-high | -3.40 | 6.97e-03 | NA | NA | NA | | TRINITY_DN79927_c0_g1 | Venturia inaequalis |
| TRINITY_DN39159_c0_g1 | Rola-high | -3.39 | 8.76e-03 | NA | NA | NA | | hypothetical protein DVH24_027315, partial | — |
| TRINITY_DN29137_c0_g1 | Rola-high | -3.37 | 4.99e-03 | NA | NA | NA | | TRINITY_DN29137_c0_g1 | — |

| Trinity gene ID | Direction | Trinity | | GDDH13 protein match | | | Def. | Description | FCS-GX |
|---|---|---|---|---|---|---|---|---|---|
| | | log2FC | padj | gene ID | log2FC | padj | | | call |
| TRINITY_DN22483_c0_g1 | Rola-high | -3.35 | 1.07e-03 | NA | NA | NA | | TRINITY_DN22483_c0_g1 | Venturia inaequalis |
| TRINITY_DN8155_c0_g1 | Rola-high | -3.35 | 1.15e-03 | NA | NA | NA | | hypothetical protein DVH24_022460 | — |
| TRINITY_DN41409_c0_g1 | Rola-high | -3.34 | 2.75e-03 | NA | NA | NA | | extensin-1-like | — |
| TRINITY_DN3885_c0_g2 | Rin-high | 3.33 | 5.46e-03 | NA | NA | NA | | TRINITY_DN3885_c0_g2 | — |
| TRINITY_DN49925_c0_g2 | Rola-high | -3.29 | 5.10e-03 | NA | NA | NA | | hypothetical protein Q3G72_007303 | Venturia inaequalis |
| TRINITY_DN9223_c0_g1 | Rin-high | 3.28 | 1.97e-03 | NA | NA | NA | | TRINITY_DN9223_c0_g1 | Venturia inaequalis |
| TRINITY_DN118764_c0_g1 | Rin-high | 3.26 | 4.45e-03 | NA | NA | NA | | TRINITY_DN118764_c0_g1 | Venturia inaequalis |
| TRINITY_DN12500_c0_g1 | Rola-high | -3.23 | 7.45e-03 | NA | NA | NA | | TRINITY_DN12500_c0_g1 | — |
| TRINITY_DN2099_c2_g1 | Rola-high | -3.19 | 9.23e-03 | NA | NA | NA | | hypothetical protein DVH24_007801 | — |
| TRINITY_DN207497_c0_g1 | Rin-high | 3.15 | 4.88e-04 | NA | NA | NA | | TRINITY_DN207497_c0_g1 | Venturia inaequalis |
| TRINITY_DN222047_c0_g1 | Rola-high | -3.14 | 8.57e-04 | NA | NA | NA | | alpha-N-arabinofuranosidase 2 precursor, putative | Venturia inaequalis |
| TRINITY_DN1769_c0_g3 | Rola-high | -3.13 | 2.96e-03 | NA | NA | NA | | glycine-rich protein 5-like | — |
| TRINITY_DN24262_c0_g1 | Rin-high | 3.09 | 4.82e-03 | NA | NA | NA | | TRINITY_DN24262_c0_g1 | — |
| TRINITY_DN113883_c0_g1 | Rola-high | -3.07 | 7.94e-03 | NA | NA | NA | | lysine-specific histone demethylase 1 homolog 3-like | Venturia inaequalis |
| TRINITY_DN9343_c1_g1 | Rin-high | 3.07 | 8.36e-03 | NA | NA | NA | | TRINITY_DN9343_c1_g1 | — |
| TRINITY_DN5160_c0_g2 | Rola-high | -3.07 | 3.77e-04 | NA | NA | NA | | proline-rich protein 4 | — |
| TRINITY_DN118323_c0_g1 | Rin-high | 3.04 | 5.76e-03 | NA | NA | NA | | TRINITY_DN118323_c0_g1 | Venturia inaequalis |
| TRINITY_DN135926_c0_g1 | Rola-high | -3.04 | 9.84e-04 | NA | NA | NA | | TRINITY_DN135926_c0_g1 | Venturia inaequalis |
| TRINITY_DN6544_c1_g1 | Rola-high | -3.01 | 3.36e-03 | NA | NA | NA | | protein NIM1-INTERACTING 1-like | — |
| TRINITY_DN4933_c0_g1 | Rola-high | -3.00 | 1.02e-04 | NA | NA | NA | | TRINITY_DN4933_c0_g1 | — |

| Trinity gene ID | Direction | Trinity | | GDDH13 protein match | | | Def. | Description | FCS-GX call |
| --- | --- | --- | --- | --- | --- | --- | --- | --- | --- |
| | | log2FC | padj | gene ID | log2FC | padj | | | |
| TRINITY_DN1801_c2_g1 | Rola-high | -2.97 | 3.62e-03 | NA | NA | NA | | TRINITY_DN1801_c2_g1 | — |
| TRINITY_DN6129_c0_g2 | Rin-high | 2.96 | 8.62e-03 | NA | NA | NA | | hypothetical protein DVH24_025398 | — |
| TRINITY_DN20321_c0_g1 | Rola-high | -2.95 | 7.18e-03 | NA | NA | NA | | TRINITY_DN20321_c0_g1 | Venturia inaequalis |
| TRINITY_DN1716_c3_g1 | Rin-high | 2.93 | 9.00e-03 | NA | NA | NA | | TRINITY_DN1716_c3_g1 | — |
| TRINITY_DN34516_c0_g1 | Rola-high | -2.90 | 1.82e-03 | NA | NA | NA | | TRINITY_DN34516_c0_g1 | — |
| TRINITY_DN5952_c1_g1 | Rin-high | 2.89 | 8.64e-03 | NA | NA | NA | | TRINITY_DN5952_c1_g1 | — |
| TRINITY_DN11372_c0_g1 | Rola-high | -2.88 | 1.66e-03 | NA | NA | NA | | proline-rich protein 4 | — |
| TRINITY_DN37425_c0_g1 | Rola-high | -2.87 | 8.37e-03 | NA | NA | NA | | TRINITY_DN37425_c0_g1 | Venturia inaequalis |
| TRINITY_DN7040_c3_g1 | Rin-high | 2.87 | 5.97e-03 | NA | NA | NA | | TRINITY_DN7040_c3_g1 | — |
| TRINITY_DN21724_c0_g1 | Rola-high | -2.82 | 8.93e-03 | NA | NA | NA | | TRINITY_DN21724_c0_g1 | — |
| TRINITY_DN16474_c0_g1 | Rola-high | -2.77 | 1.21e-03 | NA | NA | NA | | cytochrome b-c1 complex subunit 6-1, mitochondrial | Venturia inaequalis |
| TRINITY_DN47_c0_g2 | Rola-high | -2.76 | 8.87e-04 | NA | NA | NA | | TRINITY_DN47_c0_g2 | — |
| TRINITY_DN19275_c3_g1 | Rin-high | 2.72 | 9.65e-03 | NA | NA | NA | | unnamed protein product | — |
| TRINITY_DN52087_c0_g2 | Rin-high | 2.67 | 2.65e-03 | NA | NA | NA | | TRINITY_DN52087_c0_g2 | — |
| TRINITY_DN782_c0_g1 | Rin-high | 2.62 | 3.14e-05 | NA | NA | NA | | TRINITY_DN782_c0_g1 | — |
| TRINITY_DN12215_c0_g1 | Rola-high | -2.60 | 5.86e-04 | NA | NA | NA | | TRINITY_DN12215_c0_g1 | — |
| TRINITY_DN5090_c7_g1 | Rola-high | -2.59 | 3.88e-03 | NA | NA | NA | | TRINITY_DN5090_c7_g1 | — |
| TRINITY_DN35425_c0_g2 | Rola-high | -2.58 | 2.13e-03 | NA | NA | NA | | TRINITY_DN35425_c0_g2 | — |
| TRINITY_DN55368_c0_g2 | Rola-high | -2.39 | 2.11e-03 | NA | NA | NA | | hypothetical protein DVH24_040017 | — |
| TRINITY_DN5611_c0_g2 | Rola-high | -2.26 | 3.20e-03 | NA | NA | NA | | protein NIM1-INTERACTING 3 | — |
| TRINITY_DN368_c1_g2 | Rola-high | -2.11 | 8.27e-03 | NA | NA | NA | | TRINITY_DN368_c1_g2 | — |
