## Supplementary tables and figures for "Transcriptome of apple cv. Ametyst in *Rvi6*-effective and *Rvi6*-breaking *Venturia inaequalis* interactions identifies defense candidates": 13_SupplementaryTable_S6_sample_overview.pdf

**Supplementary Table S6. Per-library sample metadata, preparation batch and sequencing provider.**

Host: apple cultivar Ametyst, carrying the *Rvi6* (formerly *Vf*) scab-resistance locus introgressed from *Malus floribunda*. Leaves were mock-treated with distilled water (Control) or inoculated with one of two *Venturia inaequalis* isolates and sampled at 4 days post-inoculation. One row per sequenced library: 8 control, 8 Rin and 5 Rola, 21 in total. Paired-end reads were aligned to the GDDH13 v1.1 reference genome and counted per gene, and gene-level count data were analyzed with DESeq2 under a condition-only design (~ condition). Libraries were prepared in two batches approximately six months apart and sequenced by different providers; batch and provider are perfectly aliased, so they are represented in the analysis by a single technical batch factor. Batch 1 is the first preparation, comprising 11 libraries sequenced by Eurofins Genomics; batch 2 was prepared about six months later and comprises 10 libraries sequenced by IAB. The Rola group comprises five libraries because three of the eight prepared failed during library preparation, one in the first batch and two in the second, as follows from the 3/2 outcome against a 4/4 preparation (a technical limitation, not outlier removal). Instrument, run and flow-cell identifiers were read directly from the FASTQ headers: batch 1 (Eurofins Genomics) instrument A00604, run 138, flow cell HLCVMDRXX, lane 2; batch 2 (IAB) instrument A00380, run 44, flow cell HLCYTDRXX, lane 1. Both are Illumina NovaSeq 6000 instruments and both runs delivered 151 bp reads, reported here as 2 × 150 bp by convention. The instrument identifiers also confirm the preparation-batch assignment of every library independently of the laboratory records. All 21 libraries were sequenced on Illumina NovaSeq 6000 instruments as paired-end 2 × 150 bp runs, so the platform is stated here rather than repeated in a column. The libraries are deposited in the NCBI Gene Expression Omnibus under series accession GSE345841; the sample accession in the last column identifies each library within it.

| Sample ID | Treatment | Rvi6 interaction | Prep batch | Sequencing provider | Read pairs sequenced | GEO sample accession |
|---|---|---|---|---|---|---|
| C01 | Control | — | 2 | IAB | 65,412,459 | GSM10017734 |
| C02 | Control | — | 2 | IAB | 68,052,319 | GSM10017735 |
| C03 | Control | — | 2 | IAB | 59,845,609 | GSM10017736 |
| C04 | Control | — | 2 | IAB | 54,084,962 | GSM10017737 |
| C05 | Control | — | 1 | Eurofins Genomics | 18,628,652 | GSM10017738 |
| C06 | Control | — | 1 | Eurofins Genomics | 20,238,646 | GSM10017739 |
| C07 | Control | — | 1 | Eurofins Genomics | 18,778,613 | GSM10017740 |
| C08 | Control | — | 1 | Eurofins Genomics | 21,145,880 | GSM10017741 |
| RIN01 | Rin | Incompatible | 2 | IAB | 42,284,722 | GSM10017742 |
| RIN02 | Rin | Incompatible | 2 | IAB | 60,776,586 | GSM10017743 |
| RIN03 | Rin | Incompatible | 2 | IAB | 66,111,129 | GSM10017744 |
| RIN04 | Rin | Incompatible | 2 | IAB | 51,559,902 | GSM10017745 |
| RIN05 | Rin | Incompatible | 1 | Eurofins Genomics | 18,811,235 | GSM10017746 |
| RIN06 | Rin | Incompatible | 1 | Eurofins Genomics | 20,539,020 | GSM10017747 |
| RIN07 | Rin | Incompatible | 1 | Eurofins Genomics | 17,518,676 | GSM10017748 |
| RIN08 | Rin | Incompatible | 1 | Eurofins Genomics | 25,157,781 | GSM10017749 |
| ROL01 | Rola | Compatible | 2 | IAB | 97,215,768 | GSM10017750 |
| ROL02 | Rola | Compatible | 2 | IAB | 68,359,142 | GSM10017751 |
| ROL03 | Rola | Compatible | 1 | Eurofins Genomics | 18,250,742 | GSM10017752 |
| ROL04 | Rola | Compatible | 1 | Eurofins Genomics | 20,251,920 | GSM10017753 |
| ROL05 | Rola | Compatible | 1 | Eurofins Genomics | 20,418,358 | GSM10017754 |
