## Supplementary tables and figures for "Transcriptome of apple cv. Ametyst in *Rvi6*-effective and *Rvi6*-breaking *Venturia inaequalis* interactions identifies defense candidates": 14_SupplementaryTable_S7_module_trait.pdf

**Supplementary Table S7. Module eigengene–trait correlations.**

Pearson correlation between each module eigengene and the control, Rin and Rola treatments across the 21 libraries, with nominal two-sided Student P values and Benjamini–Hochberg adjusted values. Module eigengenes were computed as the first principal component of the module's genes after scaling to mean 0 and unit variance, sign-aligned to the module mean, from the VST matrix. P values were adjusted across all 54 tests (18 modules × 3 treatments) simultaneously. Because each library belongs to exactly one treatment, the three trait indicators are not independent. propVar is the proportion of module variance explained by the eigengene. Of the 54 tests, 25 are significant at nominal $P < 0.05$ and 22 after Benjamini–Hochberg adjustment. These are the values plotted in Fig. 5, where the nominal P values are shown.

| Module | Genes | propVar | Control r | Control P | Control BH | Rin r | Rin P | Rin BH | Rola r | Rola P | Rola BH |
|---|---|---|---|---|---|---|---|---|---|---|---|
| black | 613 | 0.68 | -0.493 | 2.32e-2 | 5.38e-2 | 0.831 | 3.09e-6 | 5.56e-5 | -0.386 | 8.43e-2 | 1.57e-1 |
| lightcyan | 58 | 0.81 | -0.773 | 3.96e-5 | 3.56e-4 | 0.796 | 1.56e-5 | 1.68e-4 | -0.026 | 9.09e-1 | 9.26e-1 |
| green | 744 | 0.72 | -0.808 | 9.34e-6 | 1.26e-4 | 0.692 | 5.04e-4 | 2.69e-3 | 0.132 | 5.68e-1 | 6.52e-1 |
| salmon | 159 | 0.75 | -0.565 | 7.59e-3 | 2.16e-2 | 0.689 | 5.48e-4 | 2.69e-3 | -0.141 | 5.41e-1 | 6.52e-1 |
| midnightblue | 87 | 0.83 | -0.407 | 6.67e-2 | 1.33e-1 | 0.648 | 1.48e-3 | 5.71e-3 | -0.275 | 2.29e-1 | 3.34e-1 |
| red | 731 | 0.72 | -0.060 | 7.96e-1 | 8.60e-1 | 0.626 | 2.41e-3 | 8.13e-3 | -0.645 | 1.59e-3 | 5.72e-3 |
| brown | 863 | 0.73 | 0.077 | 7.40e-1 | 8.16e-1 | 0.510 | 1.81e-2 | 4.44e-2 | -0.670 | 8.96e-4 | 3.72e-3 |
| blue | 1,568 | 0.76 | -0.730 | 1.71e-4 | 1.15e-3 | 0.214 | 3.52e-1 | 4.64e-1 | 0.589 | 4.97e-3 | 1.58e-2 |
| pink | 581 | 0.76 | 0.228 | 3.21e-1 | 4.33e-1 | 0.128 | 5.80e-1 | 6.52e-1 | -0.406 | 6.81e-2 | 1.33e-1 |
| grey60 | 55 | 0.82 | -0.155 | 5.02e-1 | 6.30e-1 | 0.016 | 9.46e-1 | 9.46e-1 | 0.159 | 4.91e-1 | 6.30e-1 |
| greenyellow | 215 | 0.72 | 0.294 | 1.96e-1 | 3.14e-1 | -0.038 | 8.72e-1 | 9.06e-1 | -0.292 | 1.98e-1 | 3.14e-1 |
| lightgreen | 32 | 0.85 | -0.261 | 2.53e-1 | 3.60e-1 | -0.053 | 8.18e-1 | 8.66e-1 | 0.359 | 1.10e-1 | 1.98e-1 |
| cyan | 92 | 0.73 | -0.351 | 1.19e-1 | 2.07e-1 | -0.130 | 5.75e-1 | 6.52e-1 | 0.548 | 1.01e-2 | 2.73e-2 |
| purple | 217 | 0.77 | -0.491 | 2.39e-2 | 5.38e-2 | -0.281 | 2.17e-1 | 3.26e-1 | 0.880 | 1.40e-7 | 3.78e-6 |
| magenta | 249 | 0.75 | 0.405 | 6.89e-2 | 1.33e-1 | -0.517 | 1.63e-2 | 4.19e-2 | 0.128 | 5.79e-1 | 6.52e-1 |
| tan | 212 | 0.84 | 0.324 | 1.52e-1 | 2.57e-1 | -0.710 | 3.15e-4 | 1.89e-3 | 0.440 | 4.62e-2 | 9.98e-2 |
| turquoise | 2,565 | 0.74 | 0.238 | 2.98e-1 | 4.13e-1 | -0.736 | 1.42e-4 | 1.10e-3 | 0.568 | 7.29e-3 | 2.16e-2 |
| yellow | 862 | 0.68 | 0.677 | 7.53e-4 | 3.39e-3 | -0.927 | 1.63e-9 | 8.80e-8 | 0.285 | 2.11e-1 | 3.26e-1 |
