## Supplementary tables and figures for "Transcriptome of apple cv. Ametyst in *Rvi6*-effective and *Rvi6*-breaking *Venturia inaequalis* interactions identifies defense candidates": 15_SupplementaryTable_S8_WGCNA_gene_modules.pdf

**Supplementary Table S8. WGCNA co-expression module assignment for the 10,000 genes analyzed.**

Module membership of every gene included in the weighted gene co-expression network analysis, provided in full as the accompanying workbook SupplementaryTable_S8_WGCNA_gene_modules.xlsx. The network was built on the 10,000 most variable of the 38,467 quantified GDDH13 genes at a soft-thresholding power of 8, yielding 18 informative modules plus a grey bin of 97 genes assigned to no module. Columns are gene_id; module; kME, the module membership of the gene, defined as its correlation with the module eigengene, with the corresponding *P* value; hub_tag, marking genes that meet the OmicsBox hub-tag criterion of kME ≥ 0.8 with kME $P < 0.05$; and gene significance (GS) for each treatment, the correlation of the individual gene with the control, Rin and Rola treatments, each with its *P* value. Genes are ordered with the green module first, the remaining modules alphabetically and grey last, and ranked within each module by kME. Two cautions apply. The hub_tag criterion is met by 7,495 of the 10,000 genes, so it is a broad label rather than evidence that a gene is an exceptionally central network node. Gene significance is a per-gene quantity and is not the same as the module eigengene–trait correlations reported in Supplementary Table S7, which are computed once per module; the two should not be compared directly. Module sizes in this table reproduce those in Supplementary Table S7 exactly, and the module assignments of the 151 Rin-induced defense-related candidate genes agree with Supplementary Table S2 for all 151 genes.
