## Supplementary tables and figures for "Transcriptome of apple cv. Ametyst in *Rvi6*-effective and *Rvi6*-breaking *Venturia inaequalis* interactions identifies defense candidates": 17_SupplementaryTable_S9_candidate_coordinates.pdf

**Supplementary Table S9. Genomic coordinates of the Rin-induced defense-related candidate genes.**

Position in the GDDH13 v1.1 assembly of all 151 candidate genes, ordered by sequence and start coordinate. Coordinates and strand are from the GDDH13 v1.1 annotation; gene length is the span of the annotated gene, including introns. Modules are the WGCNA co-expression modules of Supplementary Table S8 and Fig. 5; RefSeq locus and Description are as defined for Supplementary Table S2. Candidates are distributed across all 18 assembly sequences, most densely on Chr10 (20 genes). 9 candidates lie on Chr00, the bin of unanchored scaffolds, and therefore have no genetic map position; the remaining 142 are anchored to a chromosome. Nine clusters contain three or more candidates with successive genes separated by ≤ 200 kb; the largest contains seven green-module genes spanning 311 kb on Chr10, whereas three genes on Chr15 occur within 6.3 kb, two of which share an NCBI LOC identifier. Such clusters are consistent with tandem duplication, which is common in the receptor-like kinase and protein families prominent in this candidate set, and they should be regarded as clustered candidate regions rather than independent genomic signals. Median gene length is 2,913 bp (range 349–13,888 bp).

| Gene ID | RefSeq locus | Module | Sequence | Start | End | Strand | Length (bp) | Description |
|---|---|---|---|---|---|---|---|---|
| MD00G1023600 | LOC103432961 | black | Chr00 | 3,929,608 | 3,937,138 | + | 7,531 | Leucine-rich repeat transmembrane protein kinase |
| MD00G1046700 | LOC103432961 | green | Chr00 | 8,740,489 | 8,746,522 | - | 6,034 | Leucine-rich repeat transmembrane protein kinase |
| MD00G1062800 | LOC114823514 | blue | Chr00 | 11,567,819 | 11,571,069 | + | 3,251 | Leucine-rich repeat protein kinase family protein |
| MD00G1082100 | LOC103412691 | blue | Chr00 | 16,395,284 | 16,398,003 | + | 2,720 | Leucine-rich receptor-like protein kinase family protein |
| MD00G1101700 | LOC103419471 | green | Chr00 | 21,511,698 | 21,514,756 | - | 3,059 | cysteine-rich RLK (RECEPTOR-like protein kinase) 26 |
| MD00G1105300 | LOC103439438 | blue | Chr00 | 22,004,253 | 22,004,861 | + | 609 | Plant invertase/pectin methylesterase inhibitor superfamily protein |
| MD00G1124500 | LOC103401065 | blue | Chr00 | 26,635,155 | 26,637,513 | + | 2,359 | Leucine-rich repeat receptor-like protein kinase family protein |
| MD00G1170700 | LOC103401065 | blue | Chr00 | 39,257,906 | 39,260,529 | + | 2,624 | Leucine-rich repeat receptor-like protein kinase family protein |
| MD00G1183000 | LOC103431980 | green | Chr00 | 43,283,723 | 43,289,953 | + | 6,231 | Leucine-rich repeat transmembrane protein kinase |
| MD01G1057300 | LOC114821571 | blue | Chr01 | 16,207,098 | 16,208,765 | - | 1,668 | disease resistance family protein / LRR family protein |
| MD01G1057400 | LOC103406479 | blue | Chr01 | 16,208,767 | 16,210,098 | - | 1,332 | disease resistance family protein / LRR family protein |
| MD01G1059700 | LOC103407534 | green | Chr01 | 16,356,823 | 16,364,269 | + | 7,447 | Protein kinase superfamily protein |
| MD01G1062000 | LOC103403871 | green | Chr01 | 16,613,137 | 16,616,734 | + | 3,598 | disease resistance family protein / LRR family protein |
| MD01G1062500 | LOC103417604 | green | Chr01 | 16,645,607 | 16,648,827 | + | 3,221 | disease resistance family protein / LRR family protein |
| MD01G1064600 | LOC103406169 | blue | Chr01 | 16,839,859 | 16,843,110 | + | 3,252 | disease resistance family protein / LRR family protein |
| MD01G1150100 | LOC103455178 | green | Chr01 | 25,876,361 | 25,879,911 | - | 3,551 | Leucine-rich repeat protein kinase family protein |
| MD01G1155000 | LOC103437419 | green | Chr01 | 26,313,651 | 26,315,330 | + | 1,680 | basic helix-loop-helix (bHLH) DNA-binding superfamily protein |
| MD01G1172500 | LOC103406468 | black | Chr01 | 27,451,831 | 27,456,383 | - | 4,553 | cytochrome P450 |
| MD01G1172800 | LOC114824914 | green | Chr01 | 27,476,194 | 27,479,142 | - | 2,949 | disease resistance family protein / LRR family protein |

| Gene ID | RefSeq locus | Module | Sequence | Start | End | Strand | Length (bp) | Description |
|---|---|---|---|---|---|---|---|---|
| MD01G1178700 | LOC103415095 | green | Chr01 | 27,871,991 | 27,874,378 | + | 2,388 | disease resistance family protein / LRR family protein |
| MD01G1179800 | LOC103417845 | green | Chr01 | 27,949,555 | 27,952,692 | + | 3,138 | disease resistance family protein / LRR family protein |
| MD01G1213000 | LOC103439830 | green | Chr01 | 30,633,237 | 30,637,541 | + | 4,305 | plastidic pyruvate kinase beta subunit 1 |
| MD02G1091900 | LOC114819647 | green | Chr02 | 7,298,505 | 7,301,053 | + | 2,549 | Protein kinase superfamily protein |
| MD02G1107100 | LOC103442649 | lightcyan | Chr02 | 8,656,470 | 8,659,345 | - | 2,876 | Concanavalin A-like lectin protein kinase family protein |
| MD02G1107300 | LOC103442649 | lightcyan | Chr02 | 8,659,675 | 8,660,381 | - | 707 | Concanavalin A-like lectin protein kinase family protein |
| MD02G1138900 | LOC103407082 | midnightblue | Chr02 | 11,603,248 | 11,605,294 | - | 2,047 | Peroxidase superfamily protein |
| MD02G1179000 | LOC103453725 | salmon | Chr02 | 15,870,194 | 15,871,135 | + | 942 | myb domain protein 73 |
| MD02G1235600 | LOC114821487 | green | Chr02 | 28,197,968 | 28,199,079 | + | 1,112 | PR5-like receptor kinase |
| MD02G1236100 | LOC103409906 | blue | Chr02 | 28,241,902 | 28,242,900 | - | 999 | glutathione S-transferase TAU 8 |
| MD02G1250800 | LOC103408537 | green | Chr02 | 30,241,693 | 30,242,841 | - | 1,149 | receptor serine/threonine kinase |
| MD02G1251800 | LOC103408537 | green | Chr02 | 30,330,718 | 30,333,189 | - | 2,472 | PR5-like receptor kinase |
| MD02G1253600 | LOC108170170 | green | Chr02 | 30,510,073 | 30,511,379 | - | 1,307 | receptor serine/threonine kinase |
| MD03G1014200 | LOC114823840 | green | Chr03 | 1,111,772 | 1,113,496 | - | 1,725 | peroxidase 2 |
| MD03G1077700 | LOC103427128 | red | Chr03 | 6,296,618 | 6,299,466 | + | 2,849 | calmodulin-binding receptor-like cytoplasmic kinase 1 |
| MD03G1192900 | LOC103448830 | green | Chr03 | 26,496,259 | 26,498,388 | + | 2,130 | receptor like protein 2 |
| MD03G1246400 | LOC103452293 | blue | Chr03 | 33,395,082 | 33,403,641 | + | 8,560 | Leucine-rich repeat transmembrane protein kinase |
| MD03G1246600 | LOC103452293 | green | Chr03 | 33,405,918 | 33,416,188 | + | 10,271 | Leucine-rich repeat transmembrane protein kinase |
| MD04G1011700 | LOC103429293 | green | Chr04 | 1,324,708 | 1,331,182 | - | 6,475 | Disease resistance protein (TIR-NBS-LRR class) family |
| MD04G1020400 | LOC103402536 | green | Chr04 | 2,447,241 | 2,449,712 | + | 2,472 | disease resistance family protein / LRR family protein |
| MD04G1064400 | LOC103433014 | green | Chr04 | 8,673,633 | 8,674,310 | - | 678 | osmotin 34 |
| MD04G1066300 | LOC103427519 | black | Chr04 | 8,844,411 | 8,847,733 | - | 3,323 | cytochrome P450 |
| MD04G1092400 | LOC103433108 | blue | Chr04 | 16,457,220 | 16,459,261 | + | 2,042 | myb domain protein 36 |
| MD04G1177400 | LOC114824522 | green | Chr04 | 26,873,722 | 26,875,667 | + | 1,946 | Leucine-rich repeat protein kinase family protein |
| MD04G1181800 | LOC103434049 | salmon | Chr04 | 27,273,902 | 27,278,398 | - | 4,497 | Leucine-rich repeat protein kinase family protein |

| Gene ID | RefSeq locus | Module | Sequence | Start | End | Strand | Length (bp) | Description |
|---|---|---|---|---|---|---|---|---|
| MD04G1182200 | LOC103408596 | salmon | Chr04 | 27,314,013 | 27,319,578 | - | 5,566 | Leucine-rich repeat protein kinase family protein |
| MD04G1183000 | LOC103434046 | black | Chr04 | 27,404,103 | 27,407,919 | - | 3,817 | Leucine-rich repeat protein kinase family protein |
| MD04G1233100 | LOC103434224 | blue | Chr04 | 31,216,931 | 31,219,976 | - | 3,046 | PR5-like receptor kinase |
| MD04G1233200 | LOC103434225 | blue | Chr04 | 31,220,105 | 31,222,969 | + | 2,865 | Protein kinase superfamily protein |
| MD05G1190200 | LOC103409199 | green | Chr05 | 31,906,037 | 31,909,270 | - | 3,234 | Leucine-rich repeat receptor-like protein kinase family protein |
| MD05G1207400 | LOC114825146 | green | Chr05 | 33,813,808 | 33,816,620 | - | 2,813 | NB-ARC domain-containing disease resistance protein |
| MD05G1207800 | LOC114825146 | green | Chr05 | 33,895,190 | 33,897,766 | - | 2,577 | NB-ARC domain-containing disease resistance protein |
| MD05G1213900 | LOC103436186 | green | Chr05 | 34,522,675 | 34,525,893 | - | 3,219 | S-locus lectin protein kinase family protein |
| MD05G1214700 | LOC103436186 | green | Chr05 | 34,570,863 | 34,573,993 | - | 3,131 | S-locus lectin protein kinase family protein |
| MD05G1217100 | LOC103428093 | green | Chr05 | 34,780,202 | 34,781,645 | + | 1,444 | S-locus lectin protein kinase family protein |
| MD05G1236100 | LOC103427965 | green | Chr05 | 36,755,297 | 36,762,477 | - | 7,181 | disease resistance family protein / LRR family protein |
| MD05G1257000 | LOC103438143 | green | Chr05 | 38,793,726 | 38,801,226 | + | 7,501 | Leucine-rich repeat transmembrane protein kinase |
| MD05G1257600 | LOC103424618 | green | Chr05 | 39,393,044 | 39,399,323 | + | 6,280 | Leucine-rich repeat transmembrane protein kinase |
| MD05G1295700 | LOC103434906 | green | Chr05 | 43,023,949 | 43,025,466 | - | 1,518 | WRKY DNA-binding protein 65 |
| MD05G1305700 | LOC103408934 | green | Chr05 | 43,811,194 | 43,814,581 | + | 3,388 | glutamate receptor 2.7 |
| MD06G1027300 | LOC103436866 | black | Chr06 | 3,351,729 | 3,355,059 | + | 3,331 | xyloglucan endotransglucosylase/hydrolase 30 |
| MD06G1032200 | LOC103436877 | black | Chr06 | 3,863,091 | 3,865,867 | - | 2,777 | plant intracellular ras group-related LRR 4 |
| MD06G1164000 | LOC103411574 | green | Chr06 | 30,471,651 | 30,473,667 | - | 2,017 | cytochrome P450 |
| MD06G1232600 | LOC103438110 | green | Chr06 | 36,351,963 | 36,353,322 | - | 1,360 | MAP kinase kinase 9 |
| MD07G1006100 | LOC103409912 | green | Chr07 | 592,016 | 595,418 | + | 3,403 | cytochrome P450 |
| MD07G1014100 | LOC103412581 | blue | Chr07 | 1,234,291 | 1,248,178 | - | 13,888 | cysteine-rich RLK (RECEPTOR-like protein kinase) 5 |
| MD07G1164200 | LOC103439466 | salmon | Chr07 | 23,889,576 | 23,893,449 | - | 3,874 | Calcium-binding endonuclease/exonuclease/phosphatase family |
| MD07G1220800 | LOC103432823 | green | Chr07 | 29,832,307 | 29,837,736 | + | 5,430 | Cytochrome P450 superfamily protein |
| MD07G1236200 | LOC103413690 | black | Chr07 | 30,917,304 | 30,925,573 | - | 8,270 | Leucine-rich repeat protein kinase family protein |
| MD07G1236500 | LOC103410350 | black | Chr07 | 30,934,003 | 30,938,872 | - | 4,870 | Leucine-rich repeat protein kinase family protein |

| Gene ID | RefSeq locus | Module | Sequence | Start | End | Strand | Length (bp) | Description |
|---|---|---|---|---|---|---|---|---|
| MD07G1236800 | LOC103410350 | black | Chr07 | 30,951,155 | 30,956,021 | - | 4,867 | Leucine-rich repeat protein kinase family protein |
| MD07G1240700 | LOC103420753 | black | Chr07 | 31,204,105 | 31,207,040 | + | 2,936 | Fe superoxide dismutase 2 |
| MD07G1249700 | LOC103439597 | green | Chr07 | 31,853,818 | 31,858,848 | - | 5,031 | U-box domain-containing protein kinase family protein |
| MD08G1048100 | LOC103413415 | blue | Chr08 | 3,694,500 | 3,701,603 | - | 7,104 | glutamate receptor 2.2 |
| MD08G1092000 | LOC103441079 | green | Chr08 | 7,671,741 | 7,672,700 | - | 960 | myb domain protein 73 |
| MD08G1107900 | LOC103428814 | green | Chr08 | 9,481,554 | 9,485,041 | - | 3,488 | wall-associated kinase 2 |
| MD08G1210000 | LOC103441752 | green | Chr08 | 27,338,332 | 27,342,086 | + | 3,755 | Leucine-rich receptor-like protein kinase family protein |
| MD09G1064700 | LOC103429051 | lightcyan | Chr09 | 4,427,983 | 4,430,094 | - | 2,112 | UDP-Glycosyltransferase superfamily protein |
| MD09G1070100 | LOC103442602 | green | Chr09 | 4,822,947 | 4,829,801 | - | 6,855 | glutamate receptor 3.4 |
| MD09G1074300 | LOC103442449 | green | Chr09 | 5,162,693 | 5,164,884 | + | 2,192 | Thioredoxin superfamily protein |
| MD09G1130600 | LOC103443070 | green | Chr09 | 10,050,301 | 10,053,258 | - | 2,958 | receptor like protein 6 |
| MD09G1142500 | LOC103410840 | blue | Chr09 | 11,116,604 | 11,118,334 | + | 1,731 | UDP-Glycosyltransferase superfamily protein |
| MD09G1145200 | LOC103443146 | green | Chr09 | 11,288,129 | 11,290,802 | + | 2,674 | Wall-associated kinase family protein |
| MD09G1255200 | LOC103405757 | green | Chr09 | 32,675,792 | 32,676,664 | - | 873 | Protein kinase superfamily protein |
| MD10G1017300 | LOC103436524 | green | Chr10 | 2,215,926 | 2,218,198 | + | 2,273 | pathogenesis-related family protein |
| MD10G1042300 | LOC114827523 | green | Chr10 | 5,454,484 | 5,464,683 | - | 10,200 | Laccase/Diphenol oxidase family protein |
| MD10G1172300 | LOC103445446 | green | Chr10 | 26,469,318 | 26,470,297 | + | 980 | glutathione S-transferase TAU 8 |
| MD10G1212700 | LOC108172485 | green | Chr10 | 31,121,518 | 31,126,366 | - | 4,849 | Leucine-rich repeat protein kinase family protein |
| MD10G1214300 | LOC103445917 | green | Chr10 | 31,259,494 | 31,262,900 | + | 3,407 | Leucine-rich repeat receptor-like protein kinase family protein |
| MD10G1215200 | LOC103445917 | salmon | Chr10 | 31,360,118 | 31,363,453 | + | 3,336 | Leucine-rich repeat receptor-like protein kinase family protein |
| MD10G1226100 | LOC103445978 | green | Chr10 | 32,337,145 | 32,341,793 | + | 4,649 | VACUOLAR SORTING RECEPTOR 6 |
| MD10G1248500 | LOC103446854 | green | Chr10 | 34,164,259 | 34,170,421 | + | 6,163 | wall-associated kinase 2 |
| MD10G1249500 | LOC103446854 | green | Chr10 | 34,212,764 | 34,214,829 | + | 2,066 | wall associated kinase 5 |
| MD10G1250000 | LOC108171771 | green | Chr10 | 34,277,262 | 34,278,350 | + | 1,089 | wall associated kinase 5 |
| MD10G1250500 | LOC103422333 | green | Chr10 | 34,297,043 | 34,304,303 | + | 7,261 | wall-associated kinase 2 |

| Gene ID | RefSeq locus | Module | Sequence | Start | End | Strand | Length (bp) | Description |
|---|---|---|---|---|---|---|---|---|
| MD10G1250900 | LOC103422333 | green | Chr10 | 34,331,037 | 34,333,115 | + | 2,079 | wall associated kinase 5 |
| MD10G1251200 | LOC103446854 | green | Chr10 | 34,454,093 | 34,459,942 | + | 5,850 | wall-associated kinase 2 |
| MD10G1251400 | LOC103422333 | green | Chr10 | 34,469,819 | 34,474,944 | + | 5,126 | wall-associated kinase 2 |
| MD10G1273200 | LOC103410611 | blue | Chr10 | 36,498,331 | 36,505,277 | - | 6,947 | Leucine-rich repeat transmembrane protein kinase |
| MD10G1306800 | LOC103446654 | green | Chr10 | 39,241,434 | 39,244,765 | + | 3,332 | S-locus lectin protein kinase family protein |
| MD10G1307800 | LOC103446652 | green | Chr10 | 39,297,620 | 39,300,729 | - | 3,110 | S-locus lectin protein kinase family protein |
| MD10G1308000 | LOC108174318 | blue | Chr10 | 39,329,654 | 39,333,182 | + | 3,529 | S-locus lectin protein kinase family protein |
| MD10G1312500 | LOC114827806 | green | Chr10 | 39,694,016 | 39,696,840 | - | 2,825 | cysteine-rich RLK (RECEPTOR-like protein kinase) 26 |
| MD10G1334600 | LOC114827494 | green | Chr10 | 41,195,170 | 41,201,868 | + | 6,699 | Leucine-rich repeat transmembrane protein kinase |
| MD11G1018800 | LOC103447088 | blue | Chr11 | 1,610,715 | 1,613,332 | - | 2,618 | Leucine-rich receptor-like protein kinase family protein |
| MD11G1054100 | LOC103422644 | black | Chr11 | 4,611,387 | 4,612,558 | - | 1,172 | lipid transfer protein 1 |
| MD11G1066000 | LOC103447339 | midnightblue | Chr11 | 5,687,943 | 5,691,859 | + | 3,917 | Leucine-rich repeat protein kinase family protein |
| MD11G1075900 | LOC103412826 | green | Chr11 | 6,457,237 | 6,460,036 | - | 2,800 | non-specific phospholipase C1 |
| MD11G1219900 | LOC103428490 | green | Chr11 | 32,199,734 | 32,201,498 | + | 1,765 | Cytochrome P450 superfamily protein |
| MD11G1220600 | LOC103422998 | green | Chr11 | 32,367,775 | 32,369,713 | + | 1,939 | Cytochrome P450 superfamily protein |
| MD11G1231100 | LOC103448565 | green | Chr11 | 33,606,124 | 33,609,585 | - | 3,462 | S-locus lectin protein kinase family protein |
| MD11G1268200 | LOC103452293 | green | Chr11 | 38,401,010 | 38,410,624 | + | 9,615 | Leucine-rich repeat transmembrane protein kinase |
| MD12G1020000 | LOC103449405 | green | Chr12 | 1,971,622 | 1,974,679 | - | 3,058 | disease resistance family protein / LRR family protein |
| MD12G1075400 | LOC103423272 | green | Chr12 | 9,163,197 | 9,167,383 | + | 4,187 | Glycosyltransferase family 61 protein |
| MD12G1161000 | LOC103450543 | midnightblue | Chr12 | 24,081,135 | 24,084,335 | + | 3,201 | Laccase/Diphenol oxidase family protein |
| MD12G1239800 | LOC103451027 | blue | Chr12 | 31,220,090 | 31,223,601 | - | 3,512 | NAD(P)-linked oxidoreductase superfamily protein |
| MD12G1256500 | LOC103414020 | blue | Chr12 | 32,414,137 | 32,418,548 | + | 4,412 | Malectin/receptor-like protein kinase family protein |
| MD13G1033900 | LOC103402649 | green | Chr13 | 2,376,989 | 2,379,265 | + | 2,277 | Cytochrome P450 superfamily protein |
| MD13G1077900 | LOC103451854 | black | Chr13 | 5,478,407 | 5,482,438 | - | 4,032 | WRKY DNA-binding protein 72 |
| MD13G1096000 | LOC103452004 | green | Chr13 | 6,771,736 | 6,782,546 | - | 10,811 | cysteine-rich RLK (RECEPTOR-like protein kinase) 2 |

| Gene ID | RefSeq locus | Module | Sequence | Start | End | Strand | Length (bp) | Description |
|---|---|---|---|---|---|---|---|---|
| MD13G1103500 | LOC103430668 | black | Chr13 | 7,387,120 | 7,389,501 | + | 2,382 | cytochrome P450 |
| MD13G1103800 | LOC103452391 | black | Chr13 | 7,405,032 | 7,407,460 | + | 2,429 | cytochrome P450 |
| MD13G1150700 | LOC103452669 | green | Chr13 | 11,807,823 | 11,809,934 | - | 2,112 | WRKY DNA-binding protein 48 |
| MD13G1172100 | LOC103452977 | black | Chr13 | 14,090,464 | 14,090,812 | - | 349 | Thioredoxin superfamily protein |
| MD13G1203800 | LOC103401970 | blue | Chr13 | 18,355,102 | 18,356,366 | + | 1,265 | Protein kinase superfamily protein |
| MD13G1237300 | LOC103445478 | green | Chr13 | 24,139,197 | 24,140,931 | - | 1,735 | xyloglucan endotransglycosylase 6 |
| MD14G1186700 | LOC108172934 | green | Chr14 | 27,892,603 | 27,893,783 | - | 1,181 | disease resistance family protein / LRR family protein |
| MD14G1186800 | LOC108172934 | green | Chr14 | 27,893,787 | 27,895,514 | - | 1,728 | disease resistance family protein / LRR family protein |
| MD14G1207100 | LOC103455550 | salmon | Chr14 | 29,404,456 | 29,405,076 | - | 621 | cysteine-rich RLK (RECEPTOR-like protein kinase) 25 |
| MD15G1028700 | LOC103450491 | green | Chr15 | 1,741,377 | 1,743,587 | + | 2,211 | cytochrome P450 |
| MD15G1033400 | LOC103441496 | black | Chr15 | 2,342,858 | 2,344,922 | - | 2,065 | cytochrome P450 |
| MD15G1033900 | LOC103441496 | green | Chr15 | 2,389,764 | 2,391,683 | - | 1,920 | cytochrome P450 |
| MD15G1046200 | LOC103418317 | green | Chr15 | 3,174,460 | 3,178,245 | + | 3,786 | glutamate receptor 2.8 |
| MD15G1146100 | LOC103456338 | green | Chr15 | 10,871,178 | 10,875,219 | + | 4,042 | Integrin-linked protein kinase family |
| MD15G1218000 | LOC103400938 | green | Chr15 | 17,562,593 | 17,564,951 | + | 2,359 | Protein kinase superfamily protein |
| MD15G1239300 | LOC103431980 | green | Chr15 | 19,701,185 | 19,702,629 | + | 1,445 | Leucine-rich repeat transmembrane protein kinase |
| MD15G1239400 | LOC103431980 | green | Chr15 | 19,702,631 | 19,706,174 | + | 3,544 | Leucine-rich repeat transmembrane protein kinase |
| MD15G1239500 | LOC103424618 | green | Chr15 | 19,706,176 | 19,707,471 | + | 1,296 | Leucine-rich repeat transmembrane protein kinase |
| MD15G1288600 | LOC103401412 | green | Chr15 | 26,699,131 | 26,700,069 | + | 939 | myb domain protein 73 |
| MD15G1426700 | LOC103417504 | green | Chr15 | 52,729,199 | 52,733,002 | + | 3,804 | disease resistance family protein / LRR family protein |
| MD16G1014000 | LOC103429442 | black | Chr16 | 1,073,805 | 1,076,717 | - | 2,913 | xyloglucan endotransglucosylase/hydrolase 28 |
| MD16G1016800 | LOC103444588 | lightcyan | Chr16 | 1,251,285 | 1,255,787 | + | 4,503 | UDP-Glycosyltransferase / trehalose-phosphatase family protein |
| MD16G1091200 | LOC103403007 | red | Chr16 | 6,334,488 | 6,336,158 | - | 1,671 | xyloglucan:xyloglucosyl transferase 33 |
| MD16G1116100 | LOC103416681 | salmon | Chr16 | 8,241,332 | 8,243,256 | - | 1,925 | cytochrome P450 |
| MD16G1285900 | LOC103444865 | salmon | Chr16 | 40,864,307 | 40,867,323 | + | 3,017 | cytochrome P450 |

| Gene ID | RefSeq locus | Module | Sequence | Start | End | Strand | Length (bp) | Description |
|---|---|---|---|---|---|---|---|---|
| MD17G1014200 | LOC103404209 | blue | Chr17 | 1,184,927 | 1,185,349 | - | 423 | calmodulin-like 38 |
| MD17G1034800 | LOC103404344 | green | Chr17 | 2,511,187 | 2,513,803 | + | 2,617 | formate dehydrogenase |
| MD17G1062200 | LOC103409759 | green | Chr17 | 5,065,232 | 5,068,366 | - | 3,135 | Malectin/receptor-like protein kinase family protein |
| MD17G1063200 | LOC103432089 | green | Chr17 | 5,167,355 | 5,170,053 | - | 2,699 | Malectin/receptor-like protein kinase family protein |
| MD17G1092400 | LOC103425937 | black | Chr17 | 7,730,960 | 7,732,680 | - | 1,721 | Peroxidase superfamily protein |
| MD17G1126300 | LOC103405101 | black | Chr17 | 11,093,107 | 11,094,650 | - | 1,544 | UDP-Glycosyltransferase superfamily protein |
| MD17G1250000 | LOC103426173 | green | Chr17 | 29,942,908 | 29,944,341 | - | 1,434 | Pathogenesis-related thaumatin superfamily protein |
