## Supplementary tables and figures for "Transcriptome of apple cv. Ametyst in *Rvi6*-effective and *Rvi6*-breaking *Venturia inaequalis* interactions identifies defense candidates": 18_SupplementaryTable_S10_software_versions.pdf

**Supplementary Table S10. Software, versions and key parameters.**

Every tool used in the analysis, in workflow order, with the settings that affect the reported results. Package versions for the R components were taken from the sessionInfo records saved alongside the DESeq2 outputs. Two steps are split across implementations and are listed separately: co-expression modules were constructed in OmicsBox, whereas the module eigengene–trait correlations and P values shown in Fig. 5 were computed by the custom script *wgcna_module_trait.R* from the VST matrix and the OmicsBox module assignments. Gene-set enrichment was likewise run inside OmicsBox; the version given for those rows is the OmicsBox version, and the accompanying references identify the underlying methods. *ashr* shrinkage was applied only to the $\log_2$ fold changes plotted in the MA panel of Fig. 2C; every test, candidate definition and tabulated fold change in this manuscript uses the unshrunken DESeq2 estimates.

| Analysis step | Software | *Version* | Key settings and parameters | Reference |
|---|---|---|---|---|
| Read trimming | *Trim Galore* | 0.6.10 | default settings | Krueger 2023; Martin 2011 |
| Read error correction | *Rcorrector* | 1.0.4 | default settings | Song and Florea 2015 |
| Reference genome | *GDDH13* | v1.1 | obtained from the Genome Database for Rosaceae | Daccord et al. 2017; Jung et al. 2019 |
| Read alignment | *STAR* | 2.7.11 | default settings | Dobin et al. 2013 |
| Fragment counting | *featureCounts (Subread)* | 2.0.0 | -p -B -C -s 0 -t exon -g gene_name. The unstranded setting (-s 0) is correct: the Eurofins run report NG-26891 records an unstranded cDNA library for all 11 libraries of that batch, and the IAB batch was ordered identically apart from sequencing depth | Liao et al. 2014 |
| *De novo* assembly | *Trinity* | 2.15.2 | default settings | Grabherr et al. 2011; Haas et al. 2013 |
| Transcript quantification | *Salmon* | 1.10.2 | automatic library-type inference; validateMappings; sequence-bias and GC-bias correction | Patro et al. 2017 |
| Gene-level import | *tximport* | 1.40.0 | length-scaled abundance estimates | Soneson et al. 2015 |
| Differential expression | *DESeq2* | 1.52.0 | design ~ condition for all reported results; standard Wald test alpha = 0.05; thresholded Wald test alpha = 0.01, lfcThreshold = 1, altHypothesis = "greaterAbs". A ~ batch + condition model (batch = library-preparation batch, aliased with sequencing provider) was fitted as a sensitivity check; ~ condition remains the primary model. Under the batch model the candidate sets change little: 1,339 → 1,248 for Rin vs Rola, 1,710 → 1,649 for Rin vs control and 874 → 859 for Rola vs control, with Pearson r ≥ 0.998 between the two models. The full comparison for both quantification layers is in Supplementary Data S13 and S14. | Love et al. 2014 |
| Multiple-testing correction | *DESeq2 (built-in)* | 1.52.0 | Benjamini–Hochberg, applied to all DESeq2 tests | Benjamini and Hochberg 1995 |
| Fold-change shrinkage | ashr (via DESeq2 lfcShrink) | 2.2-63 | lfcShrink(type = "ashr") for the Rin vs Rola contrast; used only to shrink $\log_2$ fold changes for the MA plot in Fig. 2C. All tests, candidate definitions and tabulated fold changes use the unshrunken DESeq2 results. | Stephens 2017 |

| Analysis step | Software | *Version* | Key settings and parameters | Reference |
|---|---|---|---|---|
| Co-expression network construction | OmicsBox (WGCNA module) | 4.0.54 | 10,000 most variable GDDH13 genes; soft-threshold power 8; 18 informative modules plus a grey bin; hub tag kME ≥ 0.8 and P < 0.05 | Langfelder and Horvath 2008 |
| Module eigengene–trait analysis | *wgcna_module_trait.R* (custom script) | R 4.6.0 | VST matrix; module eigengene = first principal component of the genes of each module after scaling to mean 0 and unit variance, sign-aligned to the module mean; Pearson correlation with the treatment indicators; two-sided Student P; Benjamini–Hochberg adjustment across all 54 tests (18 modules × 3 treatments); grey excluded | Langfelder and Horvath 2008 |
| Functional annotation | *OmicsBox* | 4.0.54 | BLASTX against the *Malus* × *domestica*/GDDH13 protein set, then Viridiplantae for sequences without a confident apple hit; taxon filter Viridiplantae. All OmicsBox defaults: CloudBLAST e-value 1.0E-3, 20 hits per query, word size 6, HSP length cut-off 33, HSP–hit coverage 0 (no filter), SEG low-complexity filter on; Blast2GO annotation cut-off 55, GO weight 5, E-value hit filter 1.0E-6, HSP–hit coverage cut-off 0. GDDH13 CloudBLAST and annotation run 14 Jul 2026 against the NCBI protein database as served by CloudBLAST on that date; nr release not exposed by the service. | Conesa et al. 2005; Götz et al. 2008 |
| Protein-domain annotation | *InterProScan* | via OmicsBox 4.0.54 CloudIPS; run 14 Jul 2026 (CloudIPS queries the InterPro release current at run time, so no local version is pinned) | Default CloudIPS member databases: AntiFam, CDD, FunFam, Gene3D, HAMAP, NCBIfam, PANTHER, PfamA, PIRSF, PIRSR, PRINTS, ProSiteProfiles, SFLD, SUPERFAMILY; ProSitePatterns and Coils disabled. GO terms merged into the Blast2GO annotation. | Jones et al. 2014 |
| Gene-set enrichment | *OmicsBox (GSEA module)* | 4.0.54 | gene-set size 15–500; 1,000 permutations; FDR q < 0.05; ranking by the complete DESeq2 Wald statistic | Subramanian et al. 2005 |
| Cross-strategy protein search | *BLASTX* | 2.17.0+ | confident match ≥ 40 % amino-acid identity over ≥ 50 aligned residues, summed across HSPs to the best-scoring subject; query coverage not used | Altschul et al. 1990 |
| Contamination screening | *FCS-GX (NCBI Foreign Contamination Screen)* | 0.5.5 | GX reference database build 2023-01-24 (3,025,824 sequences, 709 Gbp); --tax-id 3750 (Malus × domestica); run 1 Sep 2026. Sequences with an EXCLUDE call removed, the 4 TRIM calls applied and the 281 REVIEW sequences also removed | Astashyn et al. 2024 |

| Analysis step | Software | *Version* | Key settings and parameters | Reference |
|---|---|---|---|---|
| Adaptor and vector screening | *FCS-adaptor (NCBI Foreign Contamination Screen)* | 0.5.5-linux-amd64-latest | --euk (eukaryotic adaptor set); run 2 Sep 2026 on the FCS-GX-cleaned gene-representative set. 26 sequences excluded, 159 trimmed. A second pass on 2 Sep 2026, after the TSA validator reported VECTOR_MATCH errors, repaired 28 sequences carrying internal TruSeq read-through adaptor that FCS-adaptor does not act on because it is not terminal: 20 trimmed, 8 dropped below the 200 bp TSA minimum | Astashyn et al. 2024 |
| Statistical environment | *R* | 4.6.0 (2026-04-24) | x86_64-apple-darwin20; macOS Tahoe 26.6 (platform recorded in the sessionInfo of the 2 Sep 2026 runs that produced Supplementary Data S13 and S14) | R Core Team (2026). R: A Language and Environment for Statistical Computing. R Foundation for Statistical Computing, Vienna, Austria. https://doi.org/10.32614/R.manuals |
| Supporting R packages | *SummarizedExperiment, limma, ggplot2, pheatmap, dplyr, RColorBrewer* | 1.42.0, 3.68.4, 4.0.3, 1.0.13, 1.2.1, 1.1-3 | data handling and plotting only; no inferential role | — |
