## Supplementary tables and figures for "Transcriptome of apple cv. Ametyst in *Rvi6*-effective and *Rvi6*-breaking *Venturia inaequalis* interactions identifies defense candidates": 19_SupplementaryTable_S11_Trinity_all_genes.pdf

**Supplementary Table S11. Complete *de novo* Trinity gene-level differential-expression results for all 119,509 genes tested.**

Gene-level DESeq2 results for every Trinity gene retained after tximport summarization and pre-DESeq2 count filtering, provided in full as the accompanying workbook SupplementaryTable_S11_Trinity_all_genes.xlsx. One row per gene. Columns give the Trinity gene identifier, the OmicsBox functional description, the mean of normalized counts across all 21 libraries (baseMean), and the standard Wald log2 fold change and Benjamini–Hochberg adjusted P value for each of the three contrasts (Rin vs Rola, Rin vs control, Rola vs control). Positive log2FC denotes higher expression in the first-named group. Blank adjusted P values denote genes filtered out by DESeq2 independent filtering for that contrast; the gene is still listed, so the table covers the complete set of genes queried in the *de novo* strategy. All numeric values are given at full precision. Rounding was deliberately not applied: abbreviating adjusted P values to four significant figures moved one gene (TRINITY_DN30327_c0_g1, adjusted P = 0.0499972) onto exactly 0.05, so recounting adjusted P < 0.05 from the workbook returned 20,106 Rin-vs-control DEGs instead of the 20,107 reported in Table 1, and four-decimal log2 fold changes altered five |log2FC| > 1 calls across the three contrasts. Every threshold count recomputed from this workbook now reproduces the source exactly. Corresponding thresholded-Wald candidate subsets are given in Supplementary Table S5; the complete results for the GDDH13 reference-based strategy are in the Supplementary Table S1 workbook. Two further columns, fcs_gx_action and fcs_gx_top_taxon, give the result of contamination screening of the gene-representative set with FCS-GX (Materials and methods), performed after the differential-expression analysis; they are populated for the 20,340 of these 119,509 genes flagged as non-plant and empty for the remainder. No gene was removed and no statistic was recomputed on the basis of these calls.
