## Supplementary tables and figures for "Transcriptome of apple cv. Ametyst in *Rvi6*-effective and *Rvi6*-breaking *Venturia inaequalis* interactions identifies defense candidates": 21_SupplementaryTable_S12_provenance.pdf

**Supplementary Table S12. Data source and software provenance of every table in this manuscript.**

For each table, the analysis output it derives from and the software that produced that output. The last column records how the table itself is assembled: “script” means it is regenerated from source by BP/Scripts/make_supplementary_tables.py (8 of 17 items), which also re-checks the counts quoted in the manuscript on every run; “manual” means the item was compiled once from the stated source and is not yet regenerated automatically; “knit” means it is produced by knitting the named R Markdown notebook, so it reproduces itself from the code and data it documents. The only step involving biological judgment is the manual annotation-based curation of the candidate list (BP/Scripts/inputs/curated_151_genes.tsv); tables labeled “manual” otherwise involve only transcription, formatting, filtering or deterministic subset selection from the stated analysis outputs. Full details, including the exact filters and sort orders, are in BP/Tables/PROVENANCE.md.

| Table | Source data | Software that produced the source | Table assembled by |
| --- | --- | --- | --- |
| Table 1 | DESeq2 result tables for three contrasts, both quantification strategies | DESeq2 1.52.0 in R 4.6.0 | script |
| Table 2 | Curated candidate list; DESeq2 Rin-vs-control and thresholded Rin-vs-Rola results; WGCNA module assignments; GDDH13 v1.1 coordinates; reciprocal protein-BLAST RefSeq-locus mapping | DESeq2; OmicsBox 4.0.54 (WGCNA); GDDH13 v1.1 annotation; NCBI BLAST+ | script |
| S1 | DESeq2 result tables, GDDH13 strategy, all 38,467 genes tested | DESeq2 1.52.0 | manual |
| S2 | Curated candidate list; DESeq2 results for both contrasts; NCBI reciprocal BLAST; DESeq2_GDDH_batch_sensitivity_{Rin_vs_control,Rola_vs_control}_genes.csv (batch-model status column, recomputed 2 Sep 2026) | DESeq2; OmicsBox 4.0.54 (WGCNA); NCBI BLAST+ | script |
| S3 | S2 restricted to the green module; gene-level WGCNA module membership and trait-correlation values from the OmicsBox export (not the module-eigengene correlations of S7 and Fig. 5); RefSeq-locus mapping inherited from S2 | DESeq2; OmicsBox 4.0.54 (WGCNA); NCBI BLAST+ | manual |
| S4 | GSEA_GDDH.txt, all 673 Gene Ontology terms tested | OmicsBox 4.0.54 (GSEA) | manual |
| S5 | Trinity_vs_GDDH_classified.csv, all 1,817 Trinity candidates | BLASTX (NCBI BLAST+); OmicsBox annotation | manual |
| S6 | sample_metadata_21_libraries.tsv | laboratory records | script |
| S7 | wgcna_module_trait_eigengene.csv | wgcna_module_trait.R in R 4.6.0, from the DESeq2 VST matrix and OmicsBox modules | script |
| S8 | GDH_WGCNA.txt, all 10,000 genes in the network | OmicsBox 4.0.54 (WGCNA) | manual |
| S9 | curated_151_genes.tsv; GDDH13 v1.1 coordinates; reciprocal protein-BLAST RefSeq-locus mapping | GDDH13 v1.1 annotation; NCBI BLAST+ | script |
| S10 | sessionInfo records, analysis scripts, command-line parameters and OmicsBox workflow settings | various, as listed in Supplementary Table S10 | manual |

| Table | Source data | Software that produced the source | Table assembled by |
| --- | --- | --- | --- |
| S11 | DESeq2 result tables, de novo Trinity strategy, all 119,509 genes tested | DESeq2 1.52.0; tximport 1.40.0; Salmon 1.10.2 | script |
| S12 | this table | — | script |
| S13 | knitted analysis report for the GDDH13 reference-based strategy, from BP/Cowork/DESeq2_GDDH.Rmd; includes the section-16 batch-design sensitivity comparison | R Markdown / knitr; DESeq2 1.52.0 in R 4.6.0 | knit |
| S14 | knitted analysis report for the de novo Trinity strategy, from BP/Cowork/DESeq2_Trinity_OmicsBox_corrected.Rmd; includes the section-16 batch-design sensitivity comparison | R Markdown / knitr; DESeq2 1.52.0, tximport 1.40.0 in R 4.6.0 | knit |
| S15 | FCS-GX action report Ametyst_Trinity_TSA.3750.fcs_gx_report.txt; FCS-adaptor report | FCS-GX 0.5.5; FCS-adaptor 0.5.5 | manual |
