## Supplementary tables and figures for "Transcriptome of apple cv. Ametyst in *Rvi6*-effective and *Rvi6*-breaking *Venturia inaequalis* interactions identifies defense candidates": 22_SupplementaryData_S13_DESeq2_GDDH.pdf

### DESeq2 analysis of Ametyst RNA-Seq data against Rosaceae GDDH13

Mila

2026-09-02

#### Table of Contents

#### Analysis overview

This R Markdown document performs the complete DESeq2 analysis of Ametyst RNA-Seq data mapped to the Rosaceae GDDH13 genome annotation.

The biological comparison is between three conditions:

- `control`
- `Rin`, representing the response to the *Venturia inaequalis* isolate from Rubin
- `Rola`, representing the response to the isolate from Rubinola

For the key contrast `Rin` versus `Rola`, positive log2 fold change means higher expression in `Rin`, while negative log2 fold change means higher expression in `Rola`.

All leaves were harvested at the same time, each library from the pooled three youngest fully expanded leaves of one plant, and RNA was isolated in a single run. Library preparation was, however, done in two batches sent to two sequencing providers six months apart, so the reported model is checked against a `~ batch + condition` model in section 16 (Batch-design sensitivity). Section 16 reports, for each contrast, how far the two models agree on effect sizes and on which genes are called. Control and Rin are balanced 4/4 across the batches and Rola is 2 IAB / 3 Eurofins, so the batch term is reasonably determined in every group. Both models are reported; the primary design is:

```
~ condition
```

The annotation parser is adapted for Rosaceae/GDDH13 GTF/GFF attributes, especially `gene_id`, `gene_name`, and `Function`.

### Load packages

The packages loaded here are required for differential expression analysis, data manipulation, plotting, heatmaps, and expression transformations.

```
suppressPackageStartupMessages({
  library(DESeq2)
  library(SummarizedExperiment)
  library(dplyr)
  library(ggplot2)
  library(pheatmap)
  library(limma)
})
```

### 0. Paths and settings

This section defines all input and output paths and the main filtering thresholds used throughout the analysis. Edit base_dir, count_file, gff_file, and annotation_file here if the project is moved to another computer or if the annotation file has a different name.

```
# The project has lived on more than one volume. Resolving the root from a
candidate
# list instead of one hard-coded path is what stops a knit from half-running: the
knit
# of 12 Aug 2026 silently skipped section 16 because base_dir pointed at a volume
that
# was not mounted. Add new locations to the front of this list, do not edit them in.
base_dir_candidates <- c(
  "/Volumes/Data_Disk/Ametyst_re-analysis",
  "/Volumes/Mac_Data/Ametyst_re-analysis",
  getwd(),
  dirname(getwd()),
  dirname(dirname(getwd()))
)
found_root <- base_dir_candidates[dir.exists(file.path(base_dir_candidates,
"GDDH"))]
if (!length(found_root)) {
  stop("Project root not found. Tried:\n  ",
       paste(base_dir_candidates, collapse = "\n  "))
}
base_dir <- found_root[1]
cat("Project root resolved to:", base_dir, "\n")

## Project root resolved to: /Volumes/Data_Disk/Ametyst_re-analysis

metadata_file <- file.path(base_dir, "GDDH/metadata/samples.tsv")

count_file <- file.path(
```

```
  base_dir,
  "GDDH/Counts/featureCounts_gene_counts_unstranded.matrix.tsv"
)

# Change .gtf to .gff here if your file really has .gff extension.
gff_file <- file.path(
  base_dir,
  "GDDH/Annotation/GDDH13_1-1.gff"
)

annotation_file <- file.path(
  base_dir,
  "GDDH/Annotation/GDDH13_gene_annotation.tsv"
)

outdir <- file.path(base_dir, "GDDH/DESeq2")
qcdir <- file.path(outdir, "QC_extra_plots")

dir.create(outdir, recursive = TRUE, showWarnings = FALSE)
dir.create(qcdir, recursive = TRUE, showWarnings = FALSE)

min_total_count <- 10
candidate_baseMean_cutoff <- 50
candidate_padj_cutoff <- 0.05
candidate_abs_lfc_cutoff <- 1

# Batch-design sensitivity analysis (section 16).
# Libraries were prepared in two batches about six months apart and sequenced
by two
# providers, which are perfectly aliased and therefore collapse into one batch
factor.
# The published analysis uses ~ condition; section 16 refits ~ batch + condition on
the
# same genes and reports how much the candidate sets differ, so the choice of
design is
# demonstrated rather than argued. Set FALSE to skip.
run_batch_sensitivity <- TRUE

# Existing project metadata carrying the batch assignment. The Sequencing
column is
# used: EF = Eurofins Genomics = first prep batch, IAB = second, about six
months later.
# The column named Time_Collected in that file is MISNAMED - all leaves were
harvested
# at the same time, so its values track the library-prep batch, not a collection time.
# The file also lists C9 and C10, which belong to a different sample set; only
samples
# present in the metadata above are used, so those two are ignored automatically.
# The knit of 12 Aug 2026 silently skipped section 16 because this single hard-
```

```
coded
# path did not exist on the machine used, and the fallback was a quiet message.
The
# path is now resolved from a candidate list and the outcome is stated loudly, so a
# knitted PDF can never again look complete while the sensitivity analysis is
absent.
batch_metadata_candidates <- c(
  file.path(base_dir, "metadata", "samples_adjusted.tsv"),
  file.path(base_dir, "BP", "metadata", "samples_adjusted.tsv"),
  file.path(dirname(base_dir), "Ametyst_re-analysis", "metadata",
"samples_adjusted.tsv"),
  "/Volumes/Data_Disk/Ametyst_re-analysis/metadata/samples_adjusted.tsv",
  "/Volumes/Mac_Data/Ametyst_re-analysis/metadata/samples_adjusted.tsv",
  file.path(getwd(), "metadata", "samples_adjusted.tsv"),
  file.path(dirname(getwd()), "metadata", "samples_adjusted.tsv")
)
found <- batch_metadata_candidates[file.exists(batch_metadata_candidates)]
batch_metadata_file <- if (length(found)) found[1] else
batch_metadata_candidates[1]
if (length(found)) {
  cat("Batch metadata resolved to:", batch_metadata_file, "\n")
} else {
  cat("No external batch metadata file found; falling back to the inline copy\n")
  cat("embedded below, so section 16 still runs. Paths tried:\n")
  cat(paste0("  ", batch_metadata_candidates, collapse = "\n"), "\n")
}
```

```
## Batch metadata resolved to:
/Volumes/Data_Disk/Ametyst_re-analysis/metadata/samples_adjusted.tsv
```

```
has_vsn <- requireNamespace("vsn", quietly = TRUE)
has_NMF <- requireNamespace("NMF", quietly = TRUE)
has_ashr <- requireNamespace("ashr", quietly = TRUE)
```

### 1. Helper functions

This section defines reusable helper functions. The most important function parses the Rosaceae/GDDH13 GTF or GFF annotation and creates a simple gene annotation table with gene ID, gene name, description, genomic coordinates, and strand. Other helper functions save DESeq2 result tables, write plots to PDF, and generate labelled candidate-gene heatmaps.

```
create_gene_annotation_from_gff <- function(gff_file, annotation_file) {
  if (!file.exists(gff_file)) {
    stop("GFF/GTF file does not exist: ", gff_file)
  }

  message("Creating annotation from GFF/GTF: ", gff_file)
```

```r
gff_lines <- readLines(gff_file, warn = FALSE)
gff_lines <- gff_lines[!grepl("^#", gff_lines) & nzchar(gff_lines)]

parts <- strsplit(gff_lines, "\t", fixed = TRUE)
parts <- parts[vapply(parts, length, integer(1)) >= 9]

if (length(parts) == 0) {
  stop("No valid GFF/GTF records found in: ", gff_file)
}

gff_df <- data.frame(
  seqid      = vapply(parts, `[`, character(1), 1),
  source     = vapply(parts, `[`, character(1), 2),
  type       = vapply(parts, `[`, character(1), 3),
  start      = as.integer(vapply(parts, `[`, character(1), 4)),
  end        = as.integer(vapply(parts, `[`, character(1), 5)),
  score      = vapply(parts, `[`, character(1), 6),
  strand     = vapply(parts, `[`, character(1), 7),
  phase      = vapply(parts, `[`, character(1), 8),
  attributes = vapply(parts, `[`, character(1), 9),
  stringsAsFactors = FALSE
)

extract_attr <- function(x, keys) {
  out <- rep(NA_character_, length(x))

  for (key in keys) {
    # GFF3 style: key=value
    pattern_gff <- paste0("(^|;)[[:space:]]*", key, "=([^;]+)")
    idx <- grepl(pattern_gff, x, perl = TRUE) & is.na(out)
    if (any(idx)) {
      out[idx] <- sub(
        paste0(".*(^|;)[[:space:]]*", key, "=([^;]+).*"),
        "\\2",
        x[idx],
        perl = TRUE
      )
    }

    # GTF style: key "value"
    pattern_gtf <- paste0("(^|;)[[:space:]]*", key, "[[:space:]]+\"([^\"]+)\"")
    idx <- grepl(pattern_gtf, x, perl = TRUE) & is.na(out)
    if (any(idx)) {
      out[idx] <- sub(
        paste0(".*(^|;)[[:space:]]*", key, "[[:space:]]+\"([^\"]+)\".*"),
        "\\2",
        x[idx],
        perl = TRUE
      )
```

```r
    }
  }

  out <- trimws(out)
  out[out == ""] <- NA_character_
  out
}

clean_id <- function(x) {
  x <- sub("^gene:", "", x)
  x <- sub("^mRNA:", "", x)
  x <- sub("^transcript:", "", x)
  x
}

attrs <- gff_df$attributes

gene_id <- extract_attr(attrs, c("gene_id", "ID", "Parent", "gene"))
gene_name <- extract_attr(attrs, c("gene_name", "Name", "gene", "locus_tag"))
description <- extract_attr(attrs, c("Function", "product", "Note", "description"))

gene_id <- clean_id(gene_id)
gene_name <- clean_id(gene_name)

final_gene_id <- ifelse(
  !is.na(gene_name) & gene_name != "",
  gene_name,
  gene_id
)

annot_df <- data.frame(
  gene_id     = final_gene_id,
  raw_gene_id = gene_id,
  gene_name   = gene_name,
  Description = description,
  chr         = gff_df$seqid,
  start       = gff_df$start,
  end         = gff_df$end,
  strand      = gff_df$strand,
  type        = gff_df$type,
  stringsAsFactors = FALSE
)

annot_df <- annot_df[!is.na(annot_df$gene_id) & annot_df$gene_id != "", ]

if (nrow(annot_df) == 0) {
  stop("No gene IDs could be extracted from: ", gff_file)
}
```

```r
# Prefer gene lines if present. Rosaceae GDDH often has only exon/CDS.
if (any(annot_df$type == "gene")) {
  annot_df <- annot_df[annot_df$type == "gene", ]
} else if (any(annot_df$type == "exon")) {
  annot_df <- annot_df[annot_df$type == "exon", ]
}

gene_ids <- unique(annot_df$gene_id)

annotation <- do.call(rbind, lapply(gene_ids, function(gid) {
  x <- annot_df[annot_df$gene_id == gid, , drop = FALSE]

  desc <- x$Description[!is.na(x$Description) & x$Description != ""]
  if (length(desc) == 0) {
    desc <- x$gene_name[1]
  } else {
    desc <- desc[1]
  }

  data.frame(
    gene_id     = gid,
    gene_name   = ifelse(
      is.na(x$gene_name[1]) | x$gene_name[1] == "",
      gid,
      x$gene_name[1]
    ),
    Description = desc,
    chr         = x$chr[1],
    start       = min(x$start, na.rm = TRUE),
    end         = max(x$end, na.rm = TRUE),
    strand      = x$strand[1],
    stringsAsFactors = FALSE
  )
}))

rownames(annotation) <- NULL

annotation <- annotation |>
  dplyr::distinct(gene_id, .keep_all = TRUE) |>
  dplyr::arrange(gene_id) |>
  dplyr::select(gene_id, gene_name, Description, chr, start, end, strand)

write.table(
  annotation,
  annotation_file,
  sep = "\t",
  quote = FALSE,
  row.names = FALSE
)
```

```
  message("Annotation written to: ", annotation_file)
  message("Annotated genes: ", nrow(annotation))

  invisible(annotation)
}

save_deseq_result <- function(res, filename, gene_annot, outdir) {
  df <- as.data.frame(res)
  df$gene_id <- rownames(df)

  df <- df |>
    left_join(gene_annot, by = "gene_id") |>
    mutate(
      gene_name = ifelse(is.na(gene_name) | gene_name == "", gene_id,
gene_name),
      Description = ifelse(is.na(Description) | Description == "", gene_name,
Description)
    ) |>
    select(
      gene_id,
      gene_name,
      baseMean,
      log2FoldChange,
      lfcSE,
      stat,
      pvalue,
      padj,
      Description
    ) |>
    arrange(padj)

  write.csv(df, file.path(outdir, filename), row.names = FALSE)
  return(df)
}

write_png <- function(filename, width, height, expr, res = 150) {
  png(
    filename = filename,
    width = width,
    height = height,
    units = "in",
    res = res,
    type = "cairo-png"
  )
  on.exit(dev.off(), add = TRUE)
  force(expr)
}
```

```
plot_candidate_heatmap <- function(genes, filename, title, vsd_mat, ann, res_df) {
  genes <- intersect(genes, rownames(vsd_mat))

  if (length(genes) < 2) {
    message("Skipping ", filename, ": fewer than 2 genes after intersecting with
VST matrix.")
    return(invisible(NULL))
  }

  mat <- vsd_mat[genes, , drop = FALSE]
  mat_scaled <- t(scale(t(mat)))

  labels <- res_df$Description[match(genes, res_df$gene_id)]
  labels[is.na(labels) | labels == ""] <- res_df$gene_name[match(genes,
res_df$gene_id)]
  labels[is.na(labels) | labels == ""] <- genes
  rownames(mat_scaled) <- make.unique(labels)

  write_png(file.path(qcdir, filename), 10, 9, {
    pheatmap(
      mat_scaled,
      annotation_col = ann,
      show_rownames = TRUE,
      fontsize_row = 5,
      main = title,
      cluster_cols = TRUE,
      cluster_rows = TRUE
    )
  })
}

mean_sd_df <- function(mat, label) {
  data.frame(
    transformation = label,
    mean = rowMeans(mat, na.rm = TRUE),
    sd = apply(mat, 1, sd, na.rm = TRUE)
  )
}

# Display a saved PNG file inside the knitted report.
# This uses explicit markdown image syntax instead of
print(knitr::include_graphics()).
# That avoids printing the internal "knit_image_paths" object in HTML/Word
output.
show_png <- function(path, caption = NULL, width = "100%") {
  if (!file.exists(path)) {
    cat("\n\nMissing PNG file: `", path, "`\n\n", sep = "")
    return(invisible(NULL))
  }
```

```r
  # Use relative paths when possible. This makes the links portable inside
  # the knitted HTML/Word document and avoids ugly absolute paths.
  img_path <- path
  if (requireNamespace("xfun", quietly = TRUE)) {
    img_path <- xfun::relative_path(path, dir = getwd())
  }

  # R Markdown / Pandoc markdown image syntax.
  # The width attribute works in HTML and is usually respected by Word output.
  if (!is.null(caption) && nzchar(caption)) {
    cat("\n\n", "![", caption, "](", img_path, "){width=", width, "}", "\n\n", sep = "")
  } else {
    cat("\n\n", "![](", img_path, "){width=", width, "}", "\n\n", sep = "")
  }

  invisible(img_path)
}

# Display the first rows of a CSV/TSV table inside the knitted report.
show_table_head <- function(path, n = 10, caption = NULL) {
  if (!file.exists(path)) {
    cat("

Missing table file: `", path, "`

", sep = "")
    return(invisible(NULL))
  }

  if (!is.null(caption)) {
    cat("

**", caption, "**

", sep = "")
  }

  ext <- tolower(tools::file_ext(path))

  if (ext == "csv") {
    x <- read.csv(path, stringsAsFactors = FALSE, check.names = FALSE)
  } else {
    x <- read.delim(path, stringsAsFactors = FALSE, check.names = FALSE)
  }

  print(knitr::kable(head(x, n), format = "pipe"))
  invisible(x)
}
```

### 2. Read and clean metadata

This section reads the sample metadata, trims accidental spaces, standardizes condition names, and prepares the condition factor. All leaves were harvested at one time and all RNA extracted together, so no biological batch exists; each library is one plant, sampled as a pool of its three youngest fully expanded leaves. Library preparation and sequencing were not simultaneous: libraries were built in two batches about six months apart and sequenced by two providers, and that factor is perfectly aliased with the time coding in the metadata. Section 16 refits that technical batch term and quantifies the consequence for each contrast: the agreement of the effect estimates, and the overlap of the candidate sets. The batch assignment was corrected on 12 Aug 2026 — ROL3 and ROL4 came from Eurofins, not IAB — so control and Rin are 4/4 and Rola is 2 IAB / 3 Eurofins. Every figure produced by earlier knits described a 1/4 Rola split that never existed and has been discarded. The reported model contains only condition, with the batch model given alongside it. When quoting section 16, take the numbers from the current knit; do not describe the difference as merely minor, and do not read the direction of any asymmetry in the candidate sets as evidence for either model.

```
samples <- read.delim(metadata_file, stringsAsFactors = FALSE, check.names =
FALSE)
samples[] <- lapply(samples, function(x) if (is.character(x)) trimws(x) else x)

if (!"sample" %in% colnames(samples) && "sample_id" %in%
colnames(samples)) {
  samples$sample <- samples$sample_id
}

if (!"sample" %in% colnames(samples)) {
  stop("Metadata must contain column 'sample' or 'sample_id'.")
}

if (!"condition" %in% colnames(samples)) {
  stop("Metadata must contain column 'condition'.")
}

samples$condition[samples$condition == "Rol"] <- "Rola"
samples$condition[samples$condition == "C"] <- "control"
samples$condition[samples$condition == "Control"] <- "control"

samples$condition <- factor(samples$condition, levels = c("control", "Rola",
"Rin"))
samples <- droplevels(samples)

if (any(is.na(samples$condition))) {
```

```
  stop(
    "Some samples have condition values outside expected levels control/Rola/Rin:
\n",
    paste(unique(samples$condition), collapse = ", ")
  )
}

# Attach the library-preparation batch, used only by the sensitivity analysis in
# section 16. The main model remains ~ condition.
normalise_id <- function(x) {
  out <- toupper(trimws(x))
  for (p in c("RIN", "ROL", "C")) {
    hit <- startsWith(out, p)
    num <- suppressWarnings(as.integer(sub(paste0("^", p), "", out[hit])))
    out[hit] <- ifelse(is.na(num), out[hit], paste0(p, num))
  }
  out
}

# Inline fallback, a verbatim copy of metadata/samples_adjusted.tsv
# (created 5 Jun 2026; CORRECTED 12 Aug 2026 — ROL3 and ROL4 were
recorded as IAB
#  but their reads come from Eurofins originals, confirmed by read-ID matching
and
#  by file size, so Rola is 3 Eurofins / 2 IAB, not 1 / 4).
# It is embedded so section 16 runs on any machine, including one that holds the
counts but
# not that metadata file. The external file wins whenever it is present; this copy is
only
# used when it is not, and the report states which source was used. C9 and C10
belong to a
# different sample set and ROL6-ROL8 failed during library preparation; none of
them appear
# in the count matrix, so they are dropped automatically by the match below.
batch_meta_inline <- data.frame(
  sample_id = c(
    "C1", "C2", "C3", "C4", "C5", "C6", "C7", "C8", "C9", "C10", "RIN1", "RIN2",
"RIN3",
    "RIN4", "RIN5", "RIN6", "RIN7", "RIN8", "ROL1", "ROL2", "ROL3", "ROL4",
"ROL5", "ROL6",
    "ROL7", "ROL8"
  ),
  Sequencing = c(
    "IAB", "IAB", "IAB", "IAB", "EF", "EF", "EF", "EF", "EF", "EF", "IAB", "IAB",
"IAB",
    "IAB", "EF", "EF", "EF", "EF", "IAB", "IAB", "EF", "EF", "EF", "EF", "EF", "EF"
  ),
  stringsAsFactors = FALSE
)
```

```
samples$batch <- NA_character_
batch_meta <- NULL
batch_source <- NA_character_
if (file.exists(batch_metadata_file)) {
  cand <- read.delim(batch_metadata_file, stringsAsFactors = FALSE,
check.names = FALSE)
  if (all(c("sample_id", "Sequencing") %in% colnames(cand))) {
    batch_meta <- cand
    batch_source <- paste("external file:", batch_metadata_file)
  } else {
    warning("Batch metadata lacks sample_id/Sequencing columns, using the inline
copy: ",
            batch_metadata_file)
  }
}
if (is.null(batch_meta)) {
  batch_meta <- batch_meta_inline
  batch_source <- "inline copy embedded in this notebook"
}
cat("Library-preparation batch taken from", batch_source, "\n")
```

```
## Library-preparation batch taken from external file:
/Volumes/Data_Disk/Ametyst_re-analysis/metadata/samples_adjusted.tsv
```

```
idx <- match(normalise_id(samples$sample),
normalise_id(batch_meta$sample_id))
provider <- batch_meta$Sequencing[idx]
samples$batch <- dplyr::case_when(
  provider == "EF"  ~ "batch1_Eurofins",
  provider == "IAB" ~ "batch2_IAB",
  TRUE ~ NA_character_
)

# If both sources are available, they must agree, or the provenance is ambiguous.
if (identical(batch_source, paste("external file:", batch_metadata_file))) {
  ref <- batch_meta_inline$Sequencing[
    match(normalise_id(samples$sample),
normalise_id(batch_meta_inline$sample_id))]
  if (!isTRUE(all.equal(provider, ref))) {
    warning("The external batch metadata disagrees with the inline copy in this
notebook. ",
            "Resolve this before reporting the sensitivity analysis.")
  } else {
    cat("External file agrees with the inline copy for all", length(provider),
"libraries.\n")
  }
}
```

```
## External file agrees with the inline copy for all 21 libraries.
```

```r
if (anyNA(samples$batch)) {
  message("No batch assignment for: ",
paste(samples$sample[is.na(samples$batch)], collapse = ", "))
  message("Section 16 will be skipped.")
  samples$batch <- NULL
} else {
  samples$batch <- factor(samples$batch, levels = c("batch1_Eurofins",
"batch2_IAB"))
  cat("\nLibrary-preparation batch by condition:\n")
  print(table(samples$condition, samples$batch))
  cat("\nRola is unbalanced across batches because three of the eight prepared
Rola\n")
  cat("libraries failed during preparation. Rola was prepared 4/4; the surviving\n")
  cat("libraries are 3 in batch 1 and 2 in batch 2, so one failure fell in batch 1\n")
  cat("and two in batch 2. The earlier claim that all three failed in batch 1 was
a\n")
  cat("corollary of the superseded 1/4 assignment and is not consistent with it.\n")

  mm_batch <- model.matrix(~ batch + condition, data = samples)
  cat("\nRank check for ~ batch + condition:\n")
  cat("rank =", qr(mm_batch)$rank, " columns =", ncol(mm_batch), "\n")
  if (qr(mm_batch)$rank == ncol(mm_batch)) {
    cat("Full rank. The batch term IS identifiable; it is estimated from the
balanced\n")
    cat("control and Rin groups. Do not claim otherwise in the manuscript.\n")
  } else {
    cat("Rank deficient, so section 16 cannot run.\n")
  }
}
```

```
##
## Library-preparation batch by condition:
##
##          batch1_Eurofins batch2_IAB
##   control               4          4
##   Rola                  3          2
##   Rin                   4          4
##
## Rola is unbalanced across batches because three of the eight prepared Rola
## libraries failed during preparation. Rola was prepared 4/4; the surviving
## libraries are 3 in batch 1 and 2 in batch 2, so one failure fell in batch 1
## and two in batch 2. The earlier claim that all three failed in batch 1 was a
## corollary of the superseded 1/4 assignment and is not consistent with it.
##
## Rank check for ~ batch + condition:
## rank = 4  columns = 4
## Full rank. The batch term IS identifiable; it is estimated from the balanced
## control and Rin groups. Do not claim otherwise in the manuscript.
```

```r
write.csv(samples, file.path(outdir, "metadata_used_for_DESeq2.csv"), row.names
= FALSE)

cat("\nSamples used:\n")

##
## Samples used:

print(samples)

##    sample_id condition sample           batch
## 1        C01   control    C01      batch2_IAB
## 2        C02   control    C02      batch2_IAB
## 3        C03   control    C03      batch2_IAB
## 4        C04   control    C04      batch2_IAB
## 5        C05   control    C05 batch1_Eurofins
## 6        C06   control    C06 batch1_Eurofins
## 7        C07   control    C07 batch1_Eurofins
## 8        C08   control    C08 batch1_Eurofins
## 9      RIN01       Rin  RIN01      batch2_IAB
## 10     RIN02       Rin  RIN02      batch2_IAB
## 11     RIN03       Rin  RIN03      batch2_IAB
## 12     RIN04       Rin  RIN04      batch2_IAB
## 13     RIN05       Rin  RIN05 batch1_Eurofins
## 14     RIN06       Rin  RIN06 batch1_Eurofins
## 15     RIN07       Rin  RIN07 batch1_Eurofins
## 16     RIN08       Rin  RIN08 batch1_Eurofins
## 17     ROL01      Rola  ROL01      batch2_IAB
## 18     ROL02      Rola  ROL02      batch2_IAB
## 19     ROL03      Rola  ROL03 batch1_Eurofins
## 20     ROL04      Rola  ROL04 batch1_Eurofins
## 21     ROL05      Rola  ROL05 batch1_Eurofins

cat("\nCondition table:\n")

##
## Condition table:

print(table(samples$condition))

##
## control    Rola     Rin
##       8       5       8

mm <- model.matrix(~ condition, data = samples)
cat("\nDesign matrix rank check:\n")

##
## Design matrix rank check:

cat("rank =", qr(mm)$rank, "\n")
```

```
## rank = 3

cat("columns =", ncol(mm), "\n")

## columns = 3

if (qr(mm)$rank < ncol(mm)) {
  stop("Design matrix is not full rank for ~ condition.")
}
```

### 3. Read count matrix

This section reads the featureCounts matrix and extracts the columns corresponding to the samples in the metadata table. It also handles both possible first-column names: gene_id and the default featureCounts name Geneid.

```
counts_df <- read.delim(count_file, check.names = FALSE, stringsAsFactors =
FALSE)

if ("Geneid" %in% colnames(counts_df)) {
  colnames(counts_df)[colnames(counts_df) == "Geneid"] <- "gene_id"
}

if (!"gene_id" %in% colnames(counts_df)) {
  colnames(counts_df)[1] <- "gene_id"
}

rownames(counts_df) <- counts_df$gene_id
counts_df$gene_id <- NULL

missing_samples <- setdiff(samples$sample, colnames(counts_df))
if (length(missing_samples) > 0) {
  stop("Missing samples in count matrix: ", paste(missing_samples, collapse = ",
"))
}

counts_mat <- as.matrix(counts_df[, samples$sample, drop = FALSE])
storage.mode(counts_mat) <- "integer"
stopifnot(all(colnames(counts_mat) == samples$sample))
```

### 4. Create/read gene annotation

This section creates or loads the gene annotation table. If the annotation table already exists, it is reused. If it does not exist, it is rebuilt from the Rosaceae/GDDH13 GTF or GFF file. The required output columns are checked before continuing.

```r
if (!file.exists(annotation_file)) {
  gene_annot <- create_gene_annotation_from_gff(
    gff_file = gff_file,
    annotation_file = annotation_file
  )
} else {
  gene_annot <- read.delim(
    annotation_file,
    stringsAsFactors = FALSE,
    check.names = FALSE
  )
}

required_annot_cols <- c("gene_id", "gene_name", "Description")
missing_annot_cols <- setdiff(required_annot_cols, colnames(gene_annot))

if (length(missing_annot_cols) > 0) {
  stop(
    "Annotation file is missing required columns: ",
    paste(missing_annot_cols, collapse = ", "),
    "\nDelete the old annotation file and rerun:\n",
    annotation_file
  )
}

save_result <- function(res, filename) {
  save_deseq_result(res, filename, gene_annot, outdir)
}
```

### 5. DESeq2 model

This section builds the DESeq2 object, filters out very low-count genes, and runs the differential expression model. The design is `~ condition`, so each contrast compares biological conditions directly.

```r
dds <- DESeqDataSetFromMatrix(
  countData = counts_mat,
  colData = samples,
  design = ~ condition
)

dds <- dds[rowSums(counts(dds)) >= min_total_count, ]
dds <- DESeq(dds)

## estimating size factors

## estimating dispersions

## gene-wise dispersion estimates
```

```
## mean-dispersion relationship

## final dispersion estimates

## fitting model and testing

## -- replacing outliers and refitting for 17 genes
## -- DESeq argument 'minReplicatesForReplace' = 7
## -- original counts are preserved in counts(dds)

## estimating dispersions

## fitting model and testing

saveRDS(dds, file.path(outdir, "dds_GDDH_featureCounts_condition_only.rds"))
writeLines(resultsNames(dds), file.path(outdir, "DESeq2_results_names.txt"))
```

### 6. DESeq2 contrasts and result tables

This section extracts the three main DESeq2 contrasts: Rin versus Rola, Rin versus control, and Rola versus control. The result tables are joined with gene descriptions and saved as CSV files. A compact summary table reports the number of significant genes per contrast.

```
res_Rin_vs_Rola <- results(
  dds,
  contrast = c("condition", "Rin", "Rola"),
  alpha = candidate_padj_cutoff
)

res_Rin_vs_control <- results(
  dds,
  contrast = c("condition", "Rin", "control"),
  alpha = candidate_padj_cutoff
)

res_Rola_vs_control <- results(
  dds,
  contrast = c("condition", "Rola", "control"),
  alpha = candidate_padj_cutoff
)

df_Rin_vs_Rola <- save_deseq_result(
  res_Rin_vs_Rola,
  "DESeq2_GDDH_condition_only_Rin_vs_Rola_with_description.csv",
  gene_annot,
  outdir
)

df_Rin_vs_control <- save_deseq_result(
```

```r
  res_Rin_vs_control,
  "DESeq2_GDDH_condition_only_Rin_vs_control_with_description.csv",
  gene_annot,
  outdir
)

df_Rola_vs_control <- save_deseq_result(
  res_Rola_vs_control,
  "DESeq2_GDDH_condition_only_Rola_vs_control_with_description.csv",
  gene_annot,
  outdir
)

write.csv(df_Rin_vs_Rola, file.path(outdir,
"DESeq2_GDDH_condition_only_Rin_vs_Rola.csv"), row.names = FALSE)
write.csv(df_Rin_vs_control, file.path(outdir,
"DESeq2_GDDH_condition_only_Rin_vs_control.csv"), row.names = FALSE)
write.csv(df_Rola_vs_control, file.path(outdir,
"DESeq2_GDDH_condition_only_Rola_vs_control.csv"), row.names = FALSE)

summary_table <- data.frame(
  contrast = c("Rin_vs_Rola", "Rin_vs_control", "Rola_vs_control"),
  padj_0.05 = c(
    sum(res_Rin_vs_Rola$padj < 0.05, na.rm = TRUE),
    sum(res_Rin_vs_control$padj < 0.05, na.rm = TRUE),
    sum(res_Rola_vs_control$padj < 0.05, na.rm = TRUE)
  ),
  padj_0.05_absLFC1 = c(
    sum(res_Rin_vs_Rola$padj < 0.05 & abs(res_Rin_vs_Rola$log2FoldChange) >=
1, na.rm = TRUE),
    sum(res_Rin_vs_control$padj < 0.05 &
abs(res_Rin_vs_control$log2FoldChange) >= 1, na.rm = TRUE),
    sum(res_Rola_vs_control$padj < 0.05 &
abs(res_Rola_vs_control$log2FoldChange) >= 1, na.rm = TRUE)
  )
)

write.csv(summary_table, file.path(outdir,
"DESeq2_GDDH_condition_only_summary_counts.csv"), row.names = FALSE)
print(summary_table)
```

```
##          contrast padj_0.05 padj_0.05_absLFC1
## 1     Rin_vs_Rola     14233              7438
## 2  Rin_vs_control     13975              7029
## 3 Rola_vs_control      8296              4123
```

### 7. Normalized/transformed matrices

This section exports normalized count matrices and transformed expression matrices. The VST and rlog matrices are used for PCA, heatmaps, sample-distance plots, and other QC visualizations.

```
raw_counts <- counts(dds, normalized = FALSE)
norm_counts <- counts(dds, normalized = TRUE)
log2_raw_counts <- log2(raw_counts + 1)
log2_norm_counts <- log2(norm_counts + 1)

vsd <- vst(dds, blind = FALSE)
vsd_mat <- SummarizedExperiment::assay(vsd)

rld <- rlog(dds, blind = FALSE)
rlog_mat <- SummarizedExperiment::assay(rld)

write.csv(as.data.frame(norm_counts), file.path(outdir,
"DESeq2_GDDH_condition_only_normalized_counts.csv"))
write.csv(as.data.frame(vsd_mat), file.path(qcdir, "VST_expression_matrix.csv"))
write.csv(as.data.frame(rlog_mat), file.path(qcdir,
"rlog_expression_matrix.csv"))

ann <- as.data.frame(colData(dds)[, c("condition"), drop = FALSE])
rownames(ann) <- colnames(vsd_mat)
```

### 8. PCA and sample distance QC

This section performs sample-level QC using PCA and sample-distance heatmaps. PCA helps identify whether samples cluster primarily by condition and whether any samples behave as outliers.

```
write_png(file.path(outdir,
"PCA_GDDH_condition_only_raw_vst_by_condition.png"), 7, 6, {
  print(plotPCA(vsd, intgroup = "condition"))
})

## using ntop=500 top features by variance

sampleDistMatrix <- as.matrix(dist(t(vsd_mat)))
rownames(sampleDistMatrix) <- samples$sample
colnames(sampleDistMatrix) <- samples$sample

write_png(file.path(outdir,
"Sample_distance_heatmap_GDDH_condition_only_raw_vst.png"), 9, 8, {
  pheatmap(sampleDistMatrix, annotation_col = ann, annotation_row = ann)
})
```

```
pca <- prcomp(t(vsd_mat))
percentVar <- round(100 * (pca$sdev^2 / sum(pca$sdev^2)), 1)

n_pc <- min(5, ncol(pca$x))
pca_df <- as.data.frame(pca$x[, seq_len(n_pc), drop = FALSE])
pca_df$sample <- rownames(pca_df)
pca_df$condition <- samples$condition
write.csv(pca_df, file.path(outdir, "PCA_scores_condition_only.csv"), row.names =
FALSE)

for (grp in c("condition")) {
  write_png(file.path(outdir, paste0("PCA_GDDH_condition_only_by_", grp,
".png")), 7, 6, {
    print(
      ggplot(pca_df, aes(.data[["PC1"]], .data[["PC2"]], color = .data[[grp]], label =
sample)) +
        geom_point(size = 3) +
        geom_text(size = 2.5, vjust = -0.6, show.legend = FALSE) +
        xlab(paste0("PC1: ", percentVar[1], "% variance")) +
        ylab(paste0("PC2: ", percentVar[2], "% variance")) +
        coord_fixed() +
        theme_bw()
    )
  })
}
```

### 9. Extra QC: boxplots, mean-SD, correlations

This section generates additional QC plots: count-distribution boxplots, mean-SD relationships, and Pearson correlation heatmaps. These plots help evaluate whether normalization and transformation behave as expected.

```
write_png(file.path(qcdir, "Boxplot_raw_counts.png"), 11, 6, {
  boxplot(raw_counts, las = 2, outline = FALSE, main = "Raw gene counts", ylab =
"Raw counts")
})

write_png(file.path(qcdir, "Boxplot_log2_raw_counts_plus1.png"), 11, 6, {
  boxplot(log2_raw_counts, las = 2, outline = FALSE, main = "log2(raw counts +
1)", ylab = "log2(raw counts + 1)")
})

write_png(file.path(qcdir, "Boxplot_normalized_counts.png"), 11, 6, {
  boxplot(norm_counts, las = 2, outline = FALSE, main = "DESeq2 normalized
counts", ylab = "Normalized counts")
})

write_png(file.path(qcdir, "Boxplot_log2_normalized_counts_plus1.png"), 11, 6, {
```

```r
  boxplot(log2_norm_counts, las = 2, outline = FALSE, main = "log2(DESeq2
normalized counts + 1)", ylab = "log2(normalized counts + 1)")
})

write_png(file.path(qcdir, "Boxplot_VST_counts.png"), 11, 6, {
  boxplot(vsd_mat, las = 2, outline = FALSE, main = "Variance-stabilized counts",
ylab = "VST expression")
})

write_png(file.path(qcdir, "Boxplot_rlog_counts.png"), 11, 6, {
  boxplot(rlog_mat, las = 2, outline = FALSE, main = "rlog-transformed counts",
ylab = "rlog expression")
})

msd <- bind_rows(
  mean_sd_df(log2_raw_counts, "log2 raw counts + 1"),
  mean_sd_df(log2_norm_counts, "log2 normalized counts + 1"),
  mean_sd_df(vsd_mat, "VST"),
  mean_sd_df(rlog_mat, "rlog")
)

write.csv(msd, file.path(qcdir, "Mean_SD_by_transformation.csv"), row.names =
FALSE)

# geom_smooth() is deliberately not used here. ggplot2 resolves the smoother's
# default method through mgcv in StatSmooth$setup_params even when "loess" is
named
# explicitly, and loading mgcv segfaulted during the knit of 2 Sep 2026
# (dyn.load, cause 'memory not mapped') with the Trinity object in memory.
# stat_summary_bin is core ggplot2, loads nothing extra, and shows the same
trend.
# The scatter is drawn from a sample when the gene set is large - 119,509 genes
over
# three transformations is 358,527 overplotted points - while the median line is
# always computed on every gene.
msd_points <- msd
n_per_facet <- 40000
if (nrow(msd_points) > 3 * n_per_facet) {
  set.seed(1)
  keep <- unlist(lapply(
    split(seq_len(nrow(msd_points)), msd_points$transformation),
    function(ix) if (length(ix) > n_per_facet) sample(ix, n_per_facet) else ix
  ))
  msd_points <- msd_points[sort(keep), ]
  message("Mean-SD scatter drawn from ", n_per_facet,
          " genes per transformation; the median line uses all genes.")
}

## Mean-SD scatter drawn from 40000 genes per transformation; the median line
uses all genes.
```

```r
write_png(file.path(qcdir, "Heteroskedasticity_mean_SD_plots.png"), 10, 8, {
  print(
    ggplot(msd_points, aes(x = mean, y = sd)) +
      geom_point(alpha = 0.15, size = 0.4) +
      stat_summary_bin(data = msd, fun = median, geom = "line", bins = 100,
                       colour = "#2166ac", linewidth = 0.8, na.rm = TRUE) +
      facet_wrap(~ transformation, scales = "free") +
      theme_bw() +
      xlab("Gene-wise mean expression") +
      ylab("Gene-wise standard deviation") +
      ggtitle("Mean-SD relationship / heteroskedasticity check")
  )
})

if (has_vsn) {
  write_png(file.path(qcdir, "vsn_meanSdPlot_log2_normalized_counts.png"), 7, 6,
{
    vsn::meanSdPlot(log2_norm_counts, ranks = FALSE)
  })
  write_png(file.path(qcdir, "vsn_meanSdPlot_VST.png"), 7, 6, {
    vsn::meanSdPlot(vsd_mat, ranks = FALSE)
  })
  write_png(file.path(qcdir, "vsn_meanSdPlot_rlog.png"), 7, 6, {
    vsn::meanSdPlot(rlog_mat, ranks = FALSE)
  })
} else {
  message("Package 'vsn' not installed: skipping vsn::meanSdPlot outputs.")
}
```

```
## Package 'vsn' not installed: skipping vsn::meanSdPlot outputs.
```

```r
pearson_cor_vst <- cor(vsd_mat, method = "pearson")
pearson_cor_rlog <- cor(rlog_mat, method = "pearson")

write.csv(pearson_cor_vst, file.path(qcdir,
"Pearson_correlation_VST_matrix.csv"))
write.csv(pearson_cor_rlog, file.path(qcdir,
"Pearson_correlation_rlog_matrix.csv"))

write_png(file.path(qcdir, "Pearson_correlation_heatmap_VST.png"), 9, 8, {
  pheatmap(pearson_cor_vst, annotation_col = ann, annotation_row = ann, main =
"Pearson correlation: VST")
})

write_png(file.path(qcdir, "Pearson_correlation_heatmap_rlog.png"), 9, 8, {
  pheatmap(pearson_cor_rlog, annotation_col = ann, annotation_row = ann, main
= "Pearson correlation: rlog")
})
```

### 10. MA and volcano plots for Rin vs Rola

This section creates MA and volcano plots for the most important contrast, Rin versus Rola. Positive log2 fold change means higher expression in Rin; negative log2 fold change means higher expression in Rola.

```
write_png(file.path(qcdir, "MA_plot_Rin_vs_Rola_DESeq2.png"), 7, 6, {
  DESeq2::plotMA(res_Rin_vs_Rola, ylim = c(-8, 8), main = "MA plot: Rin vs Rola")
})

ma_df <- df_Rin_vs_Rola |>
  mutate(
    significant = !is.na(padj) & padj < candidate_padj_cutoff,
    baseMean_for_plot = ifelse(baseMean <= 0, NA_real_, baseMean)
  )

write.csv(ma_df, file.path(qcdir, "MA_input_Rin_vs_Rola_DESeq2.csv"),
row.names = FALSE)

write_png(file.path(qcdir, "MA_plot_Rin_vs_Rola_custom_ggplot.png"), 7, 6, {
  print(
    ggplot(ma_df, aes(x = baseMean_for_plot, y = log2FoldChange)) +
      geom_point(aes(color = significant), alpha = 0.5, size = 0.6, na.rm = TRUE) +
      scale_x_log10() +
      geom_hline(yintercept = 0, linetype = "dashed") +
      coord_cartesian(ylim = c(-8, 8)) +
      theme_bw() +
      xlab("Mean of normalized counts: baseMean, log10 scale") +
      ylab("log2 fold change: Rin vs Rola") +
      ggtitle("MA plot: Rin vs Rola")
  )
})

if (has_ashr) {
  res_shrunk <- lfcShrink(dds, contrast = c("condition", "Rin", "Rola"), type =
"ashr")
  write_png(file.path(qcdir, "MA_plot_Rin_vs_Rola_LFC_shrunk_ashr.png"), 7, 6, {
    DESeq2::plotMA(res_shrunk, ylim = c(-8, 8), main = "MA plot: Rin vs Rola, LFC
shrinkage (ashr)")
  })
} else {
  message("Package 'ashr' not installed: skipping LFC-shrunk MA plot for the Rin
vs Rola contrast.")
}
```

```
## Package 'ashr' not installed: skipping LFC-shrunk MA plot for the Rin vs Rola
contrast.
```

```
volcano_df <- df_Rin_vs_Rola |>
  mutate(
    direction = case_when(
      !is.na(padj) & padj < candidate_padj_cutoff & log2FoldChange >=
candidate_abs_lfc_cutoff ~ "Rin-high",
      !is.na(padj) & padj < candidate_padj_cutoff & log2FoldChange <=
-candidate_abs_lfc_cutoff ~ "Rola-high",
      TRUE ~ "not significant"
    ),
    neg_log10_padj = -log10(padj),
    neg_log10_padj = ifelse(is.infinite(neg_log10_padj), NA_real_, neg_log10_padj)
  )

write.csv(volcano_df, file.path(qcdir, "Volcano_input_Rin_vs_Rola.csv"),
row.names = FALSE)

write_png(file.path(qcdir, "Volcano_plot_Rin_vs_Rola.png"), 7, 6, {
  print(
    ggplot(volcano_df, aes(x = log2FoldChange, y = neg_log10_padj, color =
direction)) +
      geom_point(alpha = 0.6, size = 0.8, na.rm = TRUE) +
      geom_vline(xintercept = c(-candidate_abs_lfc_cutoff,
candidate_abs_lfc_cutoff), linetype = "dashed") +
      geom_hline(yintercept = -log10(candidate_padj_cutoff), linetype = "dashed")
+
      theme_bw() +
      xlab("log2FC: Rin vs Rola") +
      ylab("-log10 adjusted P value") +
      ggtitle("Volcano plot: Rin vs Rola")
  )
})
```

### 11. Heatmaps

This section generates heatmaps for highly variable genes, significant Rin-versus-Rola genes, and selected candidate genes. Candidate heatmaps use functional descriptions as row labels where available.

```
top_n <- 500
gene_var_vst <- apply(vsd_mat, 1, var)
top_var_genes <- names(sort(gene_var_vst, decreasing = TRUE))
[seq_len(min(top_n, length(gene_var_vst)))]
mat_top_vst_scaled <- t(scale(t(vsd_mat[top_var_genes, , drop = FALSE])))

write_png(file.path(qcdir,
"Heatmap_top500_variable_genes_VST_pheatmap.png"), 9, 10, {
  pheatmap(
    mat_top_vst_scaled,
    annotation_col = ann,
```

```
    show_rownames = FALSE,
    main = "Top 500 variable genes, VST, row-scaled"
  )
})

sig_genes <- volcano_df |>
  filter(!is.na(padj), padj < candidate_padj_cutoff, abs(log2FoldChange) >=
candidate_abs_lfc_cutoff) |>
  arrange(padj) |>
  slice_head(n = 500) |>
  pull(gene_id)

if (length(sig_genes) >= 2) {
  mat_sig_vst_scaled <- t(scale(t(vsd_mat[sig_genes, , drop = FALSE])))
  write_png(file.path(qcdir,
"Heatmap_Rin_vs_Rola_significant_genes_VST_pheatmap.png"), 9, 10, {
    pheatmap(
      mat_sig_vst_scaled,
      annotation_col = ann,
      show_rownames = FALSE,
      main = "Significant Rin vs Rola genes, VST, row-scaled"
    )
  })
} else {
  message("Fewer than 2 significant Rin vs Rola genes: skipping significant-gene
heatmap.")
}

rin_high <- df_Rin_vs_Rola |>
  filter(!is.na(padj), padj < candidate_padj_cutoff,
         log2FoldChange >= candidate_abs_lfc_cutoff,
         baseMean >= candidate_baseMean_cutoff) |>
  arrange(padj)

rola_high <- df_Rin_vs_Rola |>
  filter(!is.na(padj), padj < candidate_padj_cutoff,
         log2FoldChange <= -candidate_abs_lfc_cutoff,
         baseMean >= candidate_baseMean_cutoff) |>
  arrange(padj)

write.csv(rin_high, file.path(outdir,
"Candidate_genes_Rin_high_padj0.05_LFC1_baseMean50.csv"), row.names =
FALSE)
write.csv(rola_high, file.path(outdir,
"Candidate_genes_Rola_high_padj0.05_LFC1_baseMean50.csv"), row.names =
FALSE)

plot_candidate_heatmap(
  head(rin_high$gene_id, 50),
```

```r
  "Heatmap_top50_Rin_high_genes_VST_row_scaled.png",
  "Top 50 Rin-high genes, VST, row-scaled",
  vsd_mat,
  ann,
  df_Rin_vs_Rola
)

plot_candidate_heatmap(
  head(rola_high$gene_id, 50),
  "Heatmap_top50_Rola_high_genes_VST_row_scaled.png",
  "Top 50 Rola-high genes, VST, row-scaled",
  vsd_mat,
  ann,
  df_Rin_vs_Rola
)

plot_candidate_heatmap(
  c(head(rin_high$gene_id, 25), head(rola_high$gene_id, 25)),
  "Heatmap_top25_Rin_high_top25_Rola_high_VST_row_scaled.png",
  "Top 25 Rin-high + top 25 Rola-high genes, VST, row-scaled",
  vsd_mat,
  ann,
  df_Rin_vs_Rola
)

if (has_NMF) {
  ann_colors <- data.frame(
    condition = as.character(colData(dds)$condition)
  )
  rownames(ann_colors) <- colnames(vsd_mat)

  write_png(file.path(qcdir, "aheatmap_top500_variable_genes_VST_scaled.png"),
9, 10, {
    NMF::aheatmap(
      mat_top_vst_scaled,
      annCol = ann_colors,
      Rowv = TRUE,
      Colv = TRUE,
      distfun = "euclidean",
      hclustfun = "complete",
      labRow = NA,
      main = "aheatmap: top 500 variable genes, VST scaled"
    )
  })

  write_png(file.path(qcdir, "aheatmap_Pearson_correlation_VST.png"), 9, 8, {
    NMF::aheatmap(
      pearson_cor_vst,
      annCol = ann_colors,
      annRow = ann_colors,
```

```
    Rowv = TRUE,
    Colv = TRUE,
    distfun = function(x) as.dist(1 - x),
    hclustfun = "complete",
    main = "aheatmap: Pearson correlation, VST"
  )
 })

 if (exists("mat_sig_vst_scaled") && nrow(mat_sig_vst_scaled) >= 2) {
  write_png(file.path(qcdir,
"aheatmap_Rin_vs_Rola_significant_genes_VST_scaled.png"), 9, 10, {
    NMF::aheatmap(
      mat_sig_vst_scaled,
      annCol = ann_colors,
      Rowv = TRUE,
      Colv = TRUE,
      distfun = "euclidean",
      hclustfun = "complete",
      labRow = NA,
      main = "aheatmap: significant Rin vs Rola genes, VST scaled"
    )
  })
 }
} else {
 message("Package 'NMF' not installed: skipping NMF::aheatmap outputs.")
}
```

### 12. Session info

This section records the R session information, including package versions. This is important for reproducibility when publishing or rerunning the analysis later.

```
sink(file.path(outdir, "sessionInfo.txt"))
sessionInfo()
sink()

cat("\nMain analysis done. Results written to:\n")

##
## Main analysis done. Results written to:

cat(outdir, "\n")

## /Volumes/Data_Disk/Ametyst_re-analysis/GDDH/DESeq2

cat("\nExtra QC plots written to:\n")

##
## Extra QC plots written to:
```

```
cat(qcdir, "\n")

## /Volumes/Data_Disk/Ametyst_re-analysis/GDDH/DESeq2/QC_extra_plots
```

### 13. Extra selection: Rin vs Rola, Rin vs control, Rola vs control

This section performs stricter extra selections using lfcThreshold = 1 and adjusted P value cutoff 0.01. These tables are useful for candidate-gene prioritization because the statistical test directly asks whether the absolute log2 fold change is greater than 1.

```
res_Rin_vs_Rola_LFC1 <- results(
  dds,
  contrast = c("condition", "Rin", "Rola"),
  alpha = 0.01,
  lfcThreshold = 1,
  altHypothesis = "greaterAbs"
)

df_Rin_vs_Rola_LFC1 <- save_result(
  res_Rin_vs_Rola_LFC1,
  "DESeq2_GDDH_condition_only_Rin_vs_Rola_lfcThreshold1_with_description.csv"
)

df_Rin_vs_Rola_LFC1_sorted_by_LFC <- df_Rin_vs_Rola_LFC1 |>
  dplyr::arrange(dplyr::desc(log2FoldChange))

write.csv(
  df_Rin_vs_Rola_LFC1_sorted_by_LFC,
  file.path(outdir,
"DESeq2_Rin_vs_Rola_lfcThreshold1_padj0.01_with_description_by_log2FC.csv"),
  row.names = FALSE
)

rin_vs_rola_Rin_high <- df_Rin_vs_Rola_LFC1 |>
  dplyr::filter(!is.na(padj), padj < 0.01, log2FoldChange > 1) |>
  dplyr::arrange(dplyr::desc(log2FoldChange))

rin_vs_rola_Rola_high <- df_Rin_vs_Rola_LFC1 |>
  dplyr::filter(!is.na(padj), padj < 0.01, log2FoldChange < -1) |>
  dplyr::arrange(log2FoldChange)

write.csv(
  rin_vs_rola_Rin_high,
  file.path(outdir,
"Candidate_Rin_vs_Rola_Rin_high_lfcThreshold1_padj0.01_by_log2FC.csv"),
  row.names = FALSE
```

```
)

write.csv(
  rin_vs_rola_Rola_high,
  file.path(outdir,
"Candidate_Rin_vs_Rola_Rola_high_lfcThreshold1_padj0.01_by_log2FC.csv"),
  row.names = FALSE
)

candidate_Rin_vs_Rola_LFC1_by_absLFC <- df_Rin_vs_Rola_LFC1 |>
  dplyr::filter(!is.na(padj), padj < 0.01, abs(log2FoldChange) > 1) |>
  dplyr::mutate(
    direction = dplyr::case_when(
      log2FoldChange > 1  ~ "Rin-high",
      log2FoldChange < -1 ~ "Rola-high",
      TRUE ~ "none"
    )
  ) |>
  dplyr::select(
    gene_id, gene_name, baseMean, log2FoldChange, lfcSE, stat, pvalue, padj,
direction, Description
  ) |>
  dplyr::arrange(dplyr::desc(abs(log2FoldChange)))

write.csv(
  candidate_Rin_vs_Rola_LFC1_by_absLFC,
  file.path(outdir,
"Candidate_Rin_vs_Rola_lfcThreshold1_padj0.01_by_abs_log2FC.csv"),
  row.names = FALSE
)

res_Rin_vs_C_LFC1 <- results(
  dds,
  contrast = c("condition", "Rin", "control"),
  alpha = 0.01,
  lfcThreshold = 1,
  altHypothesis = "greaterAbs"
)

df_Rin_vs_C_LFC1 <- save_result(
  res_Rin_vs_C_LFC1,
  "DESeq2_GDDH_condition_only_Rin_vs_C_lfcThreshold1_with_description.csv"
)

df_Rin_vs_C_LFC1_sorted_by_LFC <- df_Rin_vs_C_LFC1 |>
  dplyr::arrange(dplyr::desc(log2FoldChange))

write.csv(
  df_Rin_vs_C_LFC1_sorted_by_LFC,
```

```
  file.path(outdir,
"DESeq2_Rin_vs_C_lfcThreshold1_padj0.01_with_description_by_log2FC.csv"),
  row.names = FALSE
)

rin_vs_C_Rin_high <- df_Rin_vs_C_LFC1 |>
  dplyr::filter(!is.na(padj), padj < 0.01, log2FoldChange > 1) |>
  dplyr::arrange(dplyr::desc(log2FoldChange))

rin_vs_C_C_high <- df_Rin_vs_C_LFC1 |>
  dplyr::filter(!is.na(padj), padj < 0.01, log2FoldChange < -1) |>
  dplyr::arrange(log2FoldChange)

write.csv(
  rin_vs_C_Rin_high,
  file.path(outdir,
"Candidate_Rin_vs_C_Rin_high_lfcThreshold1_padj0.01_by_log2FC.csv"),
  row.names = FALSE
)

write.csv(
  rin_vs_C_C_high,
  file.path(outdir,
"Candidate_Rin_vs_C_C_high_lfcThreshold1_padj0.01_by_log2FC.csv"),
  row.names = FALSE
)

candidate_Rin_vs_C_LFC1_by_absLFC <- df_Rin_vs_C_LFC1 |>
  dplyr::filter(!is.na(padj), padj < 0.01, abs(log2FoldChange) > 1) |>
  dplyr::mutate(
    direction = dplyr::case_when(
      log2FoldChange > 1  ~ "Rin-high",
      log2FoldChange < -1 ~ "C-high",
      TRUE ~ "none"
    )
  ) |>
  dplyr::select(
    gene_id, gene_name, baseMean, log2FoldChange, lfcSE, stat, pvalue, padj,
direction, Description
  ) |>
  dplyr::arrange(dplyr::desc(abs(log2FoldChange)))

write.csv(
  candidate_Rin_vs_C_LFC1_by_absLFC,
  file.path(outdir,
"Candidate_Rin_vs_C_lfcThreshold1_padj0.01_by_abs_log2FC.csv"),
  row.names = FALSE
)
```

```r
res_Rola_vs_C_LFC1 <- results(
  dds,
  contrast = c("condition", "Rola", "control"),
  alpha = 0.01,
  lfcThreshold = 1,
  altHypothesis = "greaterAbs"
)

df_Rola_vs_C_LFC1 <- save_result(
  res_Rola_vs_C_LFC1,
  "DESeq2_GDDH_condition_only_Rola_vs_C_lfcThreshold1_with_description.csv"
)

df_Rola_vs_C_LFC1_sorted_by_LFC <- df_Rola_vs_C_LFC1 |>
  dplyr::arrange(dplyr::desc(log2FoldChange))

write.csv(
  df_Rola_vs_C_LFC1_sorted_by_LFC,
  file.path(outdir,
"DESeq2_Rola_vs_C_lfcThreshold1_padj0.01_with_description_by_log2FC.csv"),
  row.names = FALSE
)

rola_vs_C_Rola_high <- df_Rola_vs_C_LFC1 |>
  dplyr::filter(!is.na(padj), padj < 0.01, log2FoldChange > 1) |>
  dplyr::arrange(dplyr::desc(log2FoldChange))

rola_vs_C_C_high <- df_Rola_vs_C_LFC1 |>
  dplyr::filter(!is.na(padj), padj < 0.01, log2FoldChange < -1) |>
  dplyr::arrange(log2FoldChange)

write.csv(
  rola_vs_C_Rola_high,
  file.path(outdir,
"Candidate_Rola_vs_C_Rola_high_lfcThreshold1_padj0.01_by_log2FC.csv"),
  row.names = FALSE
)

write.csv(
  rola_vs_C_C_high,
  file.path(outdir,
"Candidate_Rola_vs_C_C_high_lfcThreshold1_padj0.01_by_log2FC.csv"),
  row.names = FALSE
)

candidate_Rola_vs_C_LFC1_by_absLFC <- df_Rola_vs_C_LFC1 |>
  dplyr::filter(!is.na(padj), padj < 0.01, abs(log2FoldChange) > 1) |>
  dplyr::mutate(
    direction = dplyr::case_when(
```

```
    log2FoldChange > 1  ~ "Rola-high",
    log2FoldChange < -1 ~ "C-high",
    TRUE ~ "none"
  )
 ) |>
 dplyr::select(
  gene_id, gene_name, baseMean, log2FoldChange, lfcSE, stat, pvalue, padj,
direction, Description
 ) |>
 dplyr::arrange(dplyr::desc(abs(log2FoldChange)))

write.csv(
 candidate_Rola_vs_C_LFC1_by_absLFC,
 file.path(outdir,
"Candidate_Rola_vs_C_lfcThreshold1_padj0.01_by_abs_log2FC.csv"),
 row.names = FALSE
)
```

### 14. Comparison table and classification

This section combines the three strict contrasts into one comparison table and classifies each gene according to its response pattern. The classes distinguish common infection responses, Rin-specific responses, Rola-specific responses, opposite responses, and genes without strong differential expression.

```
make_compact_result <- function(res, prefix) {
 df <- as.data.frame(res)
 df$gene_id <- rownames(df)

 df |>
  dplyr::select(
   gene_id, baseMean, log2FoldChange, lfcSE, stat, pvalue, padj
  ) |>
  dplyr::rename(
   !!paste0("baseMean_", prefix) := baseMean,
   !!paste0("log2FC_", prefix) := log2FoldChange,
   !!paste0("lfcSE_", prefix) := lfcSE,
   !!paste0("stat_", prefix) := stat,
   !!paste0("pvalue_", prefix) := pvalue,
   !!paste0("padj_", prefix) := padj
  )
}

rin_c <- make_compact_result(res_Rin_vs_C_LFC1, "Rin_vs_C")
rola_c <- make_compact_result(res_Rola_vs_C_LFC1, "Rola_vs_C")
rin_rola <- make_compact_result(res_Rin_vs_Rola_LFC1, "Rin_vs_Rola")
```

```r
comparison <- gene_annot |>
  dplyr::select(gene_id, gene_name, Description) |>
  dplyr::left_join(rin_c, by = "gene_id") |>
  dplyr::left_join(rola_c, by = "gene_id") |>
  dplyr::left_join(rin_rola, by = "gene_id")

is_up <- function(lfc, padj) {
  !is.na(padj) & padj < 0.01 & !is.na(lfc) & lfc > 1
}

is_down <- function(lfc, padj) {
  !is.na(padj) & padj < 0.01 & !is.na(lfc) & lfc < -1
}

comparison <- comparison |>
  dplyr::mutate(
    Rin_up_vs_C = is_up(log2FC_Rin_vs_C, padj_Rin_vs_C),
    Rin_down_vs_C = is_down(log2FC_Rin_vs_C, padj_Rin_vs_C),

    Rola_up_vs_C = is_up(log2FC_Rola_vs_C, padj_Rola_vs_C),
    Rola_down_vs_C = is_down(log2FC_Rola_vs_C, padj_Rola_vs_C),

    class = dplyr::case_when(
      Rin_up_vs_C & Rola_up_vs_C ~ "common_infection_induced",
      Rin_down_vs_C & Rola_down_vs_C ~ "common_infection_repressed",

      Rin_up_vs_C & !Rola_up_vs_C & !Rola_down_vs_C ~ "Rin_specific_induced",
      Rola_up_vs_C & !Rin_up_vs_C & !Rin_down_vs_C ~ "Rola_specific_induced",

      Rin_down_vs_C & !Rola_up_vs_C & !Rola_down_vs_C ~
"Rin_specific_repressed",
      Rola_down_vs_C & !Rin_up_vs_C & !Rin_down_vs_C ~
"Rola_specific_repressed",

      Rin_up_vs_C & Rola_down_vs_C ~ "opposite_Rin_up_Rola_down",
      Rin_down_vs_C & Rola_up_vs_C ~ "opposite_Rin_down_Rola_up",

      TRUE ~ "not_strong_DE"
    )
  )

write.csv(
  comparison,
  file.path(outdir,
"Comparison_Rin_vs_C_Rola_vs_C_Rin_vs_Rola_lfcThreshold1_padj0.01.csv"),
  row.names = FALSE
)

class_summary <- comparison |>
```

```
  dplyr::count(class, sort = TRUE)

write.csv(
  class_summary,
  file.path(outdir, "Comparison_class_summary_lfcThreshold1_padj0.01.csv"),
  row.names = FALSE
)

print(class_summary)
```

```
##                     class     n
## 1           not_strong_DE 50651
## 2   Rin_specific_repressed   708
## 3     Rin_specific_induced   508
## 4 common_infection_induced   494
## 5    Rola_specific_induced   332
## 6  Rola_specific_repressed    48
```

### 15. Scatter plot: similarity of Rin and Rola vs control

This section compares the Rin-versus-control and Rola-versus-control responses directly. The scatter plot and correlation coefficients show how similar the two infection responses are at the log2 fold-change level.

```
plot_df <- comparison |>
  dplyr::filter(
    !is.na(log2FC_Rin_vs_C),
    !is.na(log2FC_Rola_vs_C)
  )

cor_pearson <- cor(
  plot_df$log2FC_Rin_vs_C,
  plot_df$log2FC_Rola_vs_C,
  method = "pearson",
  use = "complete.obs"
)

cor_spearman <- cor(
  plot_df$log2FC_Rin_vs_C,
  plot_df$log2FC_Rola_vs_C,
  method = "spearman",
  use = "complete.obs"
)

png(filename = file.path(outdir, "Scatter_log2FC_Rin_vs_C_vs_Rola_vs_C.png"),
width = 7, height = 7, units = "in", res = 150, type = "cairo-png")
```

```
print(
  ggplot(plot_df, aes(x = log2FC_Rin_vs_C, y = log2FC_Rola_vs_C)) +
    geom_point(alpha = 0.35, size = 0.7) +
    geom_hline(yintercept = 0, linetype = "dashed") +
    geom_vline(xintercept = 0, linetype = "dashed") +
    geom_abline(slope = 1, intercept = 0, linetype = "dotted") +
    coord_cartesian(xlim = c(-10, 10), ylim = c(-10, 10)) +
    theme_bw() +
    xlab("log2FC Rin vs control") +
    ylab("log2FC Rola vs control") +
    ggtitle(
      paste0(
        "Rin and Rola responses vs control\n",
        "Pearson r = ", round(cor_pearson, 3),
        ", Spearman rho = ", round(cor_spearman, 3)
      )
    )
)
```

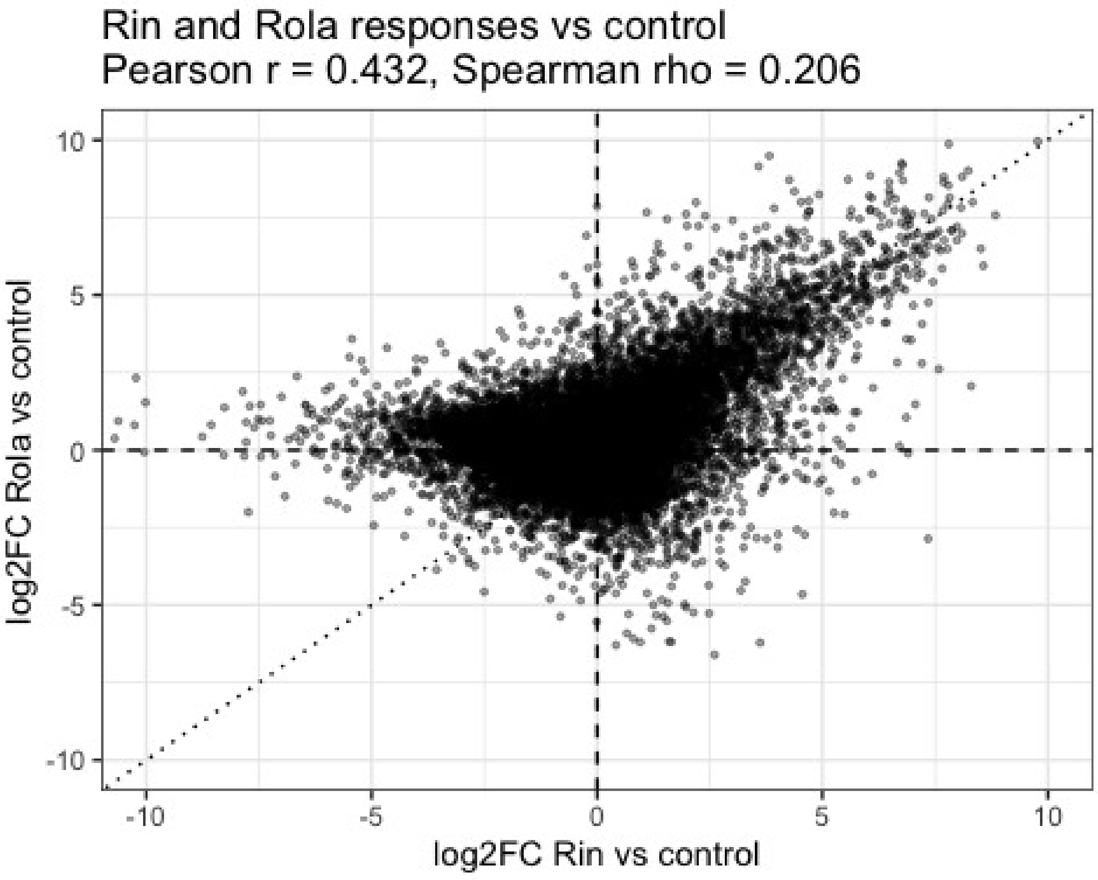


```
dev.off()

## null device
##           1
```

```r
cat("\nFull analysis completed successfully.\n")
```

```
##
## Full analysis completed successfully.
```

### 16. Batch-design sensitivity analysis

Libraries were prepared in two batches about six months apart and sequenced by two providers. Because the prep batch and the provider are perfectly aliased they cannot be separated and are treated as a single batch factor. All leaves were harvested at the same time and all RNA extracted together, so this batch effect is technical, not biological.

Control and Rin are balanced 4/4 across the two batches; Rola is 2 IAB / 3 Eurofins. Three further Rola libraries failed during preparation and were never sequenced. A rank check shows `~ batch + condition` is full rank, so the batch-adjusted condition effects are estimable, and all samples contribute to fitting the batch and condition coefficients jointly.

Note that this assignment was corrected on 12 Aug 2026. ROL3 and ROL4 had been recorded as IAB, which would have made Rola 1 Eurofins / 4 IAB; read-ID matching against the untrimmed originals showed both came from Eurofins files, and the file sizes agree, IAB libraries being roughly four times larger. Every batch-sensitivity result produced before that date used the wrong design and has been discarded.

With Rola at 2/3 the design is close to balanced in every group, so the batch term is reasonably determined throughout and the earlier concern — that the additive-batch assumption could not be checked inside Rola at $n = 1$ — no longer applies. Both models are still reported, but the comparison should now be read as a straightforward robustness check rather than a choice forced by an awkward design.

This section refits `~ batch + condition` on exactly the same filtered genes as the main model and reports, for each contrast, how many candidate genes each design returns, how many they share, and how well the log2 fold changes agree. Quote these numbers in the Methods instead of arguing about the design.

It also writes a per-gene file for each of the three contrasts, carrying baseMean, log2 fold change, lfcSE and adjusted $P$ under both designs. All three are needed, not just Rin vs Rola: the prioritized Table 2 set is defined by a thresholded Rin-vs-Rola call and can be checked from that one file, but the 151 curated genes are defined by the 707 -> 355 -> 151 chain, that is thresholded Rin-vs-control induction with baseMean >= 50 followed by the *absence* of a thresholded Rola-vs-control call in either direction. Asking whether those genes still satisfy their own defining criteria under the batch model therefore requires the Rin-vs-control and Rola-vs-control tables as well.

```r
run_sensitivity <- isTRUE(run_batch_sensitivity) &&
  "batch" %in% colnames(samples) &&
  nlevels(droplevels(samples$batch)) == 2

if (!run_sensitivity) {
  cat(strrep("!", 78), "\n")
  cat("BATCH-DESIGN SENSITIVITY ANALYSIS SKIPPED - NO RESULT IN THIS
REPORT.\n")
  cat("Diagnosis:\n")
  cat("  run_batch_sensitivity is TRUE      :", isTRUE(run_batch_sensitivity), "\n")
  cat("  a 'batch' column reached section 16:", "batch" %in% colnames(samples),
"\n")
  if ("batch" %in% colnames(samples)) {
    cat("  levels of that batch factor       :",
        nlevels(droplevels(as.factor(samples$batch))), "\n")
  } else {
    cat("  batch metadata file used          :", batch_metadata_file, "\n")
    cat("  that file exists                  :", file.exists(batch_metadata_file), "\n")
  }
  cat("The manuscript and Supplementary Table S10 both cite this comparison, so
this\n")
  cat("report is NOT usable as a supplement until the lines above all read TRUE.
\n")
  cat(strrep("!", 78), "\n")
} else {
  # Refit on the same object, so the gene filter and size factors are comparable and
  # the two designs differ only in the model formula.
  dds_batch <- dds
  design(dds_batch) <- ~ batch + condition
  dds_batch <- DESeq(dds_batch)

  saveRDS(dds_batch, file.path(outdir,
"dds_GDDH_featureCounts_batch_plus_condition.rds"))
  writeLines(resultsNames(dds_batch),
             file.path(outdir,
"DESeq2_GDDH_batch_plus_condition_results_names.txt"))

  cat("\nGenes tested under both designs:", nrow(dds_batch), "\n")

  thresholded <- function(object, contrast) {
    r <- results(object, contrast = contrast, alpha = 0.01,
                 lfcThreshold = 1, altHypothesis = "greaterAbs")
    r[!is.na(r$padj), ]
  }

  contrast_list <- list(
```

```
  Rin_vs_Rola     = c("condition", "Rin", "Rola"),
  Rin_vs_control  = c("condition", "Rin", "control"),
  Rola_vs_control = c("condition", "Rola", "control")
)

sens_rows <- list()

for (nm in names(contrast_list)) {
  r_cond  <- thresholded(dds,       contrast_list[[nm]])
  r_batch <- thresholded(dds_batch, contrast_list[[nm]])

  g_cond  <- rownames(r_cond)[r_cond$padj < 0.01]
  g_batch <- rownames(r_batch)[r_batch$padj < 0.01]
  shared  <- intersect(g_cond, g_batch)
  common  <- intersect(rownames(r_cond), rownames(r_batch))

  sens_rows[[nm]] <- data.frame(
    contrast                 = nm,
    genes_with_padj_in_both  = length(common),
    candidates_condition     = length(g_cond),
    candidates_batch         = length(g_batch),
    shared                   = length(shared),
    only_condition           = length(setdiff(g_cond, g_batch)),
    only_batch               = length(setdiff(g_batch, g_cond)),
    jaccard                  = round(length(shared) / length(union(g_cond, g_batch)),
4),
    pearson_log2FC           = round(cor(r_cond[common, "log2FoldChange"],
                                   r_batch[common, "log2FoldChange"]), 4),
    spearman_log2FC          = round(cor(r_cond[common, "log2FoldChange"],
                                   r_batch[common, "log2FoldChange"],
                                   method = "spearman"), 4),
    stringsAsFactors = FALSE
  )

  # Gene-level detail for EVERY contrast, not just Rin vs Rola. The 151-gene set is
  # defined by Rin-vs-control induction with baseMean >= 50 and the ABSENCE
of a
  # thresholded Rola-vs-control call, so testing whether those genes still meet
their
  # own defining criteria under the batch model needs the per-gene tables for all
  # three contrasts. baseMean is carried for the same reason.
  #
  # lfcSE is carried for both models because the standard error is what the extra
  # batch parameter actually changes; comparing it gene by gene shows whether
a
  # candidate was dropped because its effect estimate moved or because its
  # uncertainty grew.
  #
  # Genes evaluable under only one model are KEPT and labelled, not dropped.
```

```
The
  # earlier version intersected on non-NA padj, so candidates the batch model
could
  # not evaluate vanished from the file: for Rin vs Rola the summary counted 340
  # condition-only losses while the per-gene table explained only 313, the missing
27
  # being exactly those cases. A file used to audit disagreements must not hide
any.
  all_ids <- union(rownames(r_cond), rownames(r_batch))
  grab <- function(res, ids, col) {
    out <- rep(NA_real_, length(ids))
    hit <- ids %in% rownames(res)
    out[hit] <- res[ids[hit], col]
    out
  }
  detail <- data.frame(
    contrast           = nm,
    gene_id            = all_ids,
    baseMean_condition = grab(r_cond,  all_ids, "baseMean"),
    log2FC_condition   = grab(r_cond,  all_ids, "log2FoldChange"),
    lfcSE_condition    = grab(r_cond,  all_ids, "lfcSE"),
    padj_condition     = grab(r_cond,  all_ids, "padj"),
    baseMean_batch     = grab(r_batch, all_ids, "baseMean"),
    log2FC_batch       = grab(r_batch, all_ids, "log2FoldChange"),
    lfcSE_batch        = grab(r_batch, all_ids, "lfcSE"),
    padj_batch         = grab(r_batch, all_ids, "padj"),
    stringsAsFactors   = FALSE
  )
  detail$lfcSE_ratio <- detail$lfcSE_batch / detail$lfcSE_condition
  detail$evaluable_condition <- detail$gene_id %in% rownames(r_cond)
  detail$evaluable_batch     <- detail$gene_id %in% rownames(r_batch)
  detail$candidate_condition <- detail$gene_id %in% g_cond
  detail$candidate_batch     <- detail$gene_id %in% g_batch
  detail$agreement <- dplyr::case_when(
    !detail$evaluable_batch     & detail$candidate_condition ~
"condition_only_not_evaluable_in_batch",
    !detail$evaluable_condition & detail$candidate_batch     ~
"batch_only_not_evaluable_in_condition",
    detail$candidate_condition  &  detail$candidate_batch    ~ "both",
    detail$candidate_condition  & !detail$candidate_batch    ~
"only_condition_design",
    !detail$candidate_condition &  detail$candidate_batch    ~
"only_batch_design",
    TRUE ~ "neither"
  )
  detail <- detail |>
    dplyr::left_join(
      gene_annot |> dplyr::select(gene_id, gene_name, Description),
      by = "gene_id"
    ) |>
```

```
    dplyr::filter(agreement != "neither") |>
    dplyr::arrange(agreement, dplyr::desc(abs(log2FC_condition)))

  write.csv(
    detail,
    file.path(outdir, paste0("DESeq2_GDDH_batch_sensitivity_", nm,
"_genes.csv")),
    row.names = FALSE
  )
  cat("\n", nm, ": candidate genes written to the per-gene file: ", nrow(detail),
      " rows\n", sep = "")
  print(table(detail$agreement))
  stopifnot(sum(detail$candidate_condition) == length(g_cond),
            sum(detail$candidate_batch)     == length(g_batch))
}

sensitivity_summary <- do.call(rbind, sens_rows)

write.csv(
  sensitivity_summary,
  file.path(outdir, "DESeq2_GDDH_batch_design_sensitivity_summary.csv"),
  row.names = FALSE
)

cat("\nBatch-design sensitivity, GDDH13 reference-based (featureCounts) layer:
\n")
print(sensitivity_summary)

cat("\nHow to read this. Pearson and Spearman r near 1 mean the two models
agree on\n")
cat("effect sizes and on gene rankings. The candidate columns are a separate
question.\n")
cat("Where the two candidate sets differ, this design cannot say which model
is\n")
cat("right: Rola is 2 IAB / 3 Eurofins, so batch and condition are mildly
correlated\n")
cat("for Rola contrasts, though far less so than the 1/4 split assumed before
the\n")
cat("assignment was corrected on 12 Aug 2026. The design is full rank, so
any\n")
cat("residual effect is imprecision, not confounding in the DESeq2 sense of a\n")
cat("non-identifiable model. A gene called under one model and not\n")
cat("the other may reflect the power an additional term costs, or a call the
batch\n")
cat("term removed because batch partly explained it. Note that the contrast
which\n")
cat("changes least is Rin vs control, where both groups are balanced 4/4, and
the\n")
cat("contrasts that change most both involve Rola. The per-gene table carries
```

```
lfcSE\n")
  cat("for both models and their ratio, which separates the two readings gene by
gene:\n")
  cat("an lfcSE_ratio well above 1 with a near-identical log2FC means the call was
lost\n")
  cat("to added uncertainty, whereas a shifted log2FC at similar lfcSE means the
batch\n")
  cat("term moved the estimate itself. Report both models and the per-gene\n")
  cat("disagreements; do not read the direction of the asymmetry as evidence
either\n")
  cat("way.\n")
}
```

```
## using pre-existing size factors

## estimating dispersions

## found already estimated dispersions, replacing these

## gene-wise dispersion estimates

## mean-dispersion relationship

## final dispersion estimates

## fitting model and testing

##
## Genes tested under both designs: 38467
##
## Rin_vs_Rola: candidate genes written to the per-gene file: 1366 rows
##
##                                both condition_only_not_evaluable_in_batch
##                                1221                                  28
##                     only_batch_design               only_condition_design
##                                  27                                  90
##
## Rin_vs_control: candidate genes written to the per-gene file: 1737 rows
##
##                                both condition_only_not_evaluable_in_batch
##                                1622                                  27
##                     only_batch_design               only_condition_design
##                                  27                                  61
##
## Rola_vs_control: candidate genes written to the per-gene file: 901 rows
##
##                                both condition_only_not_evaluable_in_batch
##                                 832                                  10
##                     only_batch_design               only_condition_design
##                                  27                                  32
##
## Batch-design sensitivity, GDDH13 reference-based (featureCounts) layer:
```

```
##                     contrast genes_with_padj_in_both candidates_condition
## Rin_vs_Rola         Rin_vs_Rola                  36887                1339
## Rin_vs_control   Rin_vs_control                  35397                1710
## Rola_vs_control Rola_vs_control                  37632                 874
##               candidates_batch shared only_condition only_batch jaccard
## Rin_vs_Rola                1248   1221            118         27  0.8939
## Rin_vs_control             1649   1622             88         27  0.9338
## Rola_vs_control             859    832             42         27  0.9234
##               pearson_log2FC spearman_log2FC
## Rin_vs_Rola            0.9989          0.9979
## Rin_vs_control         0.9996          0.9994
## Rola_vs_control        0.9984          0.9966
##
## How to read this. Pearson and Spearman r near 1 mean the two models agree
on
## effect sizes and on gene rankings. The candidate columns are a separate
question.
## Where the two candidate sets differ, this design cannot say which model is
## right: Rola is 2 IAB / 3 Eurofins, so batch and condition are mildly correlated
## for Rola contrasts, though far less so than the 1/4 split assumed before the
## assignment was corrected on 12 Aug 2026. The design is full rank, so any
## residual effect is imprecision, not confounding in the DESeq2 sense of a
## non-identifiable model. A gene called under one model and not
## the other may reflect the power an additional term costs, or a call the batch
## term removed because batch partly explained it. Note that the contrast which
## changes least is Rin vs control, where both groups are balanced 4/4, and the
## contrasts that change most both involve Rola. The per-gene table carries lfcSE
## for both models and their ratio, which separates the two readings gene by
gene:
## an lfcSE_ratio well above 1 with a near-identical log2FC means the call was
lost
## to added uncertainty, whereas a shifted log2FC at similar lfcSE means the
batch
## term moved the estimate itself. Report both models and the per-gene
## disagreements; do not read the direction of the asymmetry as evidence either
## way.
```

### Output overview

After knitting this document, the main output directory is stored in `outdir`, and additional QC plots are stored in `qcdir`.

Important result files include:

- DESeq2 result tables for `Rin_vs_Rola`, `Rin_vs_control`, and `Rola_vs_control`
- candidate-gene tables using adjusted P value and log2 fold-change filters
- comparison and classification tables across the three contrasts

- PCA plots, volcano plots, MA plots, heatmaps, and correlation heatmaps
- sessionInfo.txt for reproducibility
- batch-design sensitivity tables from section 16: DESeq2_GDDH_batch_design_sensitivity_summary.csv and one per-gene file per contrast, DESeq2_GDDH_batch_sensitivity_{Rin_vs_Rola,Rin_vs_control,Rola_vs_control}_genes.csv, each carrying baseMean, log2FC, lfcSE and adjusted *P* under both designs

```
## Main output directory:
##  /Volumes/Data_Disk/Ametyst_re-analysis/GDDH/DESeq2
##
##  [1] "BLAST"
##  [2] "Candidate_genes_Rin_high_padj0.05_LFC1_baseMean50.csv"
##  [3] "Candidate_genes_Rola_high_padj0.05_LFC1_baseMean50.csv"
##  [4] "Candidate_Rin_vs_C_C_high_lfcThreshold1_padj0.01_by_log2FC.csv"
##  [5] "Candidate_Rin_vs_C_lfcThreshold1_padj0.01_by_abs_log2FC.csv"
##  [6] "Candidate_Rin_vs_C_Rin_high_lfcThreshold1_padj0.01_by_log2FC.csv"
##  [7] "Candidate_Rin_vs_Rola_GDDH_with_NCBI_blast.csv"
##  [8]
"Candidate_Rin_vs_Rola_lfcThreshold1_padj0.01_by_abs_log2FC_GDDH.csv"
##  [9] "Candidate_Rin_vs_Rola_lfcThreshold1_padj0.01_by_abs_log2FC.csv"
## [10] "Candidate_Rin_vs_Rola_Rin_high_lfcThreshold1_padj0.01_by_log2FC.csv"
## [11]
"Candidate_Rin_vs_Rola_Rola_high_lfcThreshold1_padj0.01_by_log2FC.csv"
## [12] "Candidate_Rola_vs_C_C_high_lfcThreshold1_padj0.01_by_log2FC.csv"
## [13] "Candidate_Rola_vs_C_lfcThreshold1_padj0.01_by_abs_log2FC.csv"
## [14] "Candidate_Rola_vs_C_Rola_high_lfcThreshold1_padj0.01_by_log2FC.csv"
## [15] "coexpression"
## [16] "Comparison_class_summary_lfcThreshold1_padj0.01.csv"
## [17]
"Comparison_Rin_vs_C_Rola_vs_C_Rin_vs_Rola_lfcThreshold1_padj0.01.csv"
## [18] "dds_GDDH_featureCounts_batch_plus_condition.rds"
## [19] "dds_GDDH_featureCounts_condition_only.rds"
## [20] "DESeq2_GDDH_batch_design_sensitivity_summary.csv"
## [21] "DESeq2_GDDH_batch_plus_condition_results_names.txt"
## [22] "DESeq2_GDDH_batch_sensitivity_Rin_vs_control_genes.csv"
## [23] "DESeq2_GDDH_batch_sensitivity_Rin_vs_Rola_genes.csv"
## [24] "DESeq2_GDDH_batch_sensitivity_Rola_vs_control_genes.csv"
## [25] "DESeq2_GDDH_condition_only_normalized_counts.csv"
## [26]
"DESeq2_GDDH_condition_only_Rin_vs_C_lfcThreshold1_with_description.csv"
## [27] "DESeq2_GDDH_condition_only_Rin_vs_control_with_description.csv"
## [28] "DESeq2_GDDH_condition_only_Rin_vs_control.csv"
## [29]
"DESeq2_GDDH_condition_only_Rin_vs_Rola_lfcThreshold1_with_description.csv"
## [30] "DESeq2_GDDH_condition_only_Rin_vs_Rola_with_description.csv"
## [31] "DESeq2_GDDH_condition_only_Rin_vs_Rola.csv"
## [32]
```

```
"DESeq2_GDDH_condition_only_Rola_vs_C_lfcThreshold1_with_description.csv"
## [33] "DESeq2_GDDH_condition_only_Rola_vs_control_with_description.csv"
## [34] "DESeq2_GDDH_condition_only_Rola_vs_control.csv"
## [35] "DESeq2_GDDH_condition_only_summary_counts.csv"
## [36] "DESeq2_GDDH_condition_only.R"
## [37] "DESeq2_GDDH_files"
## [38] "DESeq2_GDDH.docx"
## [39] "DESeq2_GDDH.pdf"
## [40] "DESeq2_GDDH.Rmd"
## [41] "DESeq2_results_names.txt"
## [42]
"DESeq2_Rin_vs_C_lfcThreshold1_padj0.01_with_description_by_log2FC.csv"
## [43]
"DESeq2_Rin_vs_Rola_lfcThreshold1_padj0.01_with_description_by_log2FC.csv"
## [44]
"DESeq2_Rola_vs_C_lfcThreshold1_padj0.01_with_description_by_log2FC.csv"
## [45] "DESeq2.Rproj"
## [46] "Fig2_PCA_GDDH_ellipse.png"
## [47] "Fig2_PCA_GDDH.pdf"
## [48] "Fig2_PCA_GDDH.png"
## [49] "Fig3_heatmap_top500.png"
## [50] "Fig4_volcano_MA_A_volcano.png"
## [51] "Fig4_volcano_MA_B_MA.png"
## [52] "Fig4_volcano_MA.png"
## [53] "Fig5_overlap.png"
## [54] "Fig6_functional_categories.png"
## [55] "GSEA_rank_GDDH_Rin_vs_Rola_DESeq2_stat_no_header.rnk"
## [56] "make_coexpression_input.R"
## [57] "make_Fig2_PCA.R"
## [58] "make_Fig3_hetmap.R"
## [59] "make_Fig4_volcano_MA.R"
## [60] "make_Fig5_overlap.R"
## [61] "make_Fig6_Categories.R"
## [62] "make_Table3.R"
## [63] "metadata_used_for_DESeq2.csv"
## [64] "PCA_GDDH_condition_only_by_condition.png"
## [65] "PCA_GDDH_condition_only_raw_vst_by_condition.png"
## [66] "PCA_scores_condition_only.csv"
## [67] "QC_extra_plots"
## [68] "Rplots.pdf"
## [69] "Scatter_log2FC_Rin_vs_C_vs_Rola_vs_C.png"
## [70] "sessionInfo.txt"
## [71] "Table3_main.csv"
## [72] "Table3_supplement.csv"

##
## QC output directory:
##  /Volumes/Data_Disk/Ametyst_re-analysis/GDDH/DESeq2/QC_extra_plots
##
##  [1] "aheatmap_Pearson_correlation_VST.png"
```

```
##  [2] "aheatmap_Rin_vs_Rola_significant_genes_VST_scaled.png"
##  [3] "aheatmap_top500_variable_genes_VST_scaled.png"
##  [4] "assemble_QC_figure.py"
##  [5] "Boxplot_log2_normalized_counts_plus1.png"
##  [6] "Boxplot_log2_raw_counts_plus1.png"
##  [7] "Boxplot_normalized_counts.png"
##  [8] "Boxplot_raw_counts.png"
##  [9] "Boxplot_rlog_counts.png"
## [10] "Boxplot_VST_counts.png"
## [11] "Heatmap_Rin_vs_Rola_significant_genes_VST_pheatmap.png"
## [12] "Heatmap_top25_Rin_high_top25_Rola_high_VST_row_scaled.png"
## [13] "Heatmap_top50_Rin_high_genes_VST_row_scaled.png"
## [14] "Heatmap_top50_Rola_high_genes_VST_row_scaled.png"
## [15] "Heteroskedasticity_mean_SD_plots.png"
## [16] "MA_input_Rin_vs_Rola_DESeq2.csv"
## [17] "MA_plot_Rin_vs_Rola_custom_ggplot.png"
## [18] "MA_plot_Rin_vs_Rola_DESeq2.png"
## [19] "MA_plot_Rin_vs_Rola_LFC_shrunk_ashr.png"
## [20] "Mean_SD_by_transformation.csv"
## [21] "Pearson_correlation_heatmap_rlog.png"
## [22] "Pearson_correlation_rlog_matrix.csv"
## [23] "Pearson_correlation_VST_matrix.csv"
## [24] "rlog_expression_matrix.csv"
## [25] "Volcano_input_Rin_vs_Rola.csv"
## [26] "Volcano_plot_Rin_vs_Rola.png"
## [27] "vsn_meanSdPlot_log2_normalized_counts.png"
## [28] "vsn_meanSdPlot_rlog.png"
## [29] "vsn_meanSdPlot_VST.png"
## [30] "VST_expression_matrix.csv"
```

### Report appendix: inline figures and table previews

### Report appendix: inline figures and table previews

This section is intended for the knitted HTML or Word report.
The analysis above writes all figures and result tables to files. The chunks below embed the saved PNG figures directly into the report using markdown image links and show the first 10 rows of each generated CSV table.

### Inline graphical output

The following figures are inserted from the PNG files generated during the analysis. If a figure was skipped because too few genes were available, the report prints a short “missing file” message instead of stopping.

#### Figure 1

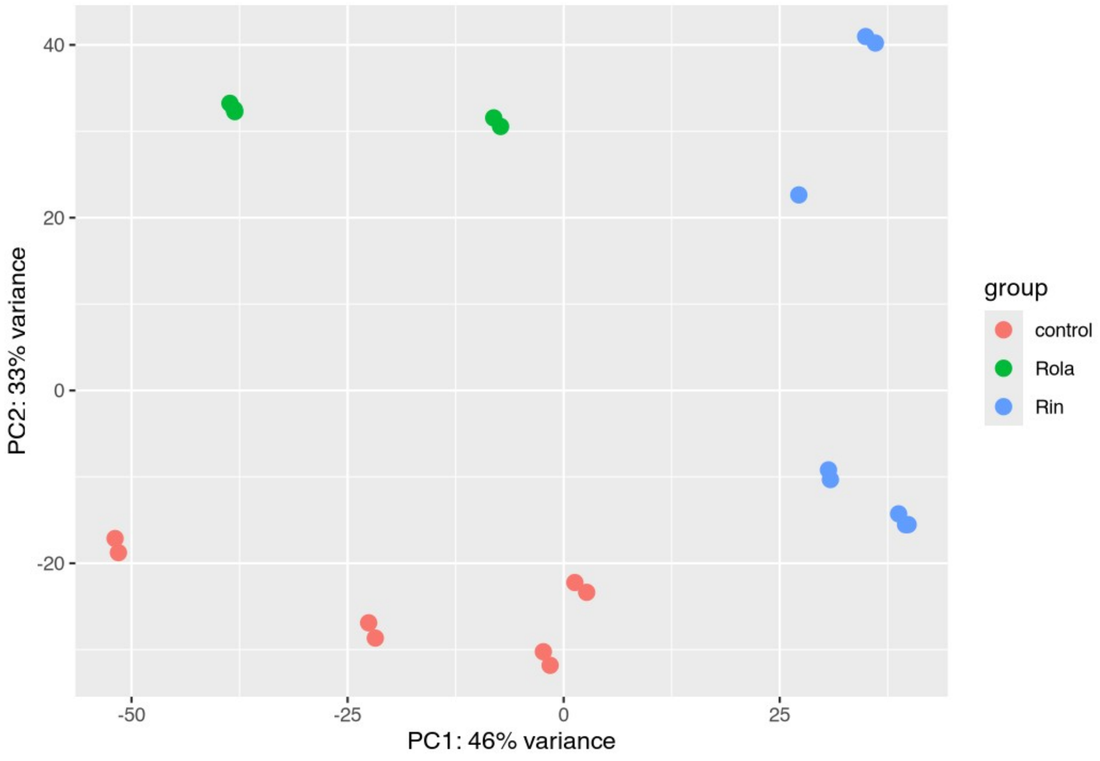


*PCA of VST-transformed expression values, colored by condition.*

#### Figure 2

Missing PNG file:
/Volumes/Data_Disk/Ametyst_re-analysis/GDDH/DESeq2/Sample_distance_heatmap_GDDH_condition_only_raw_vst.png

#### Figure 3

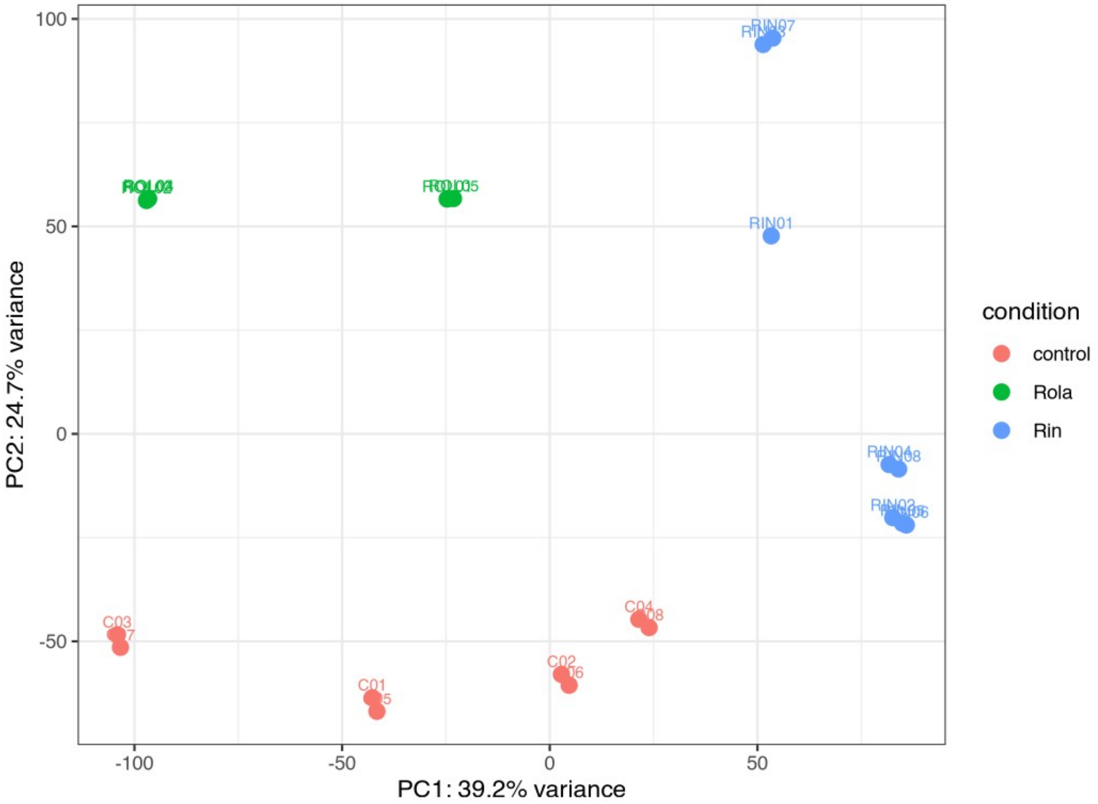


*Manual PCA of VST-transformed expression values, colored by condition.*

#### Figure 4

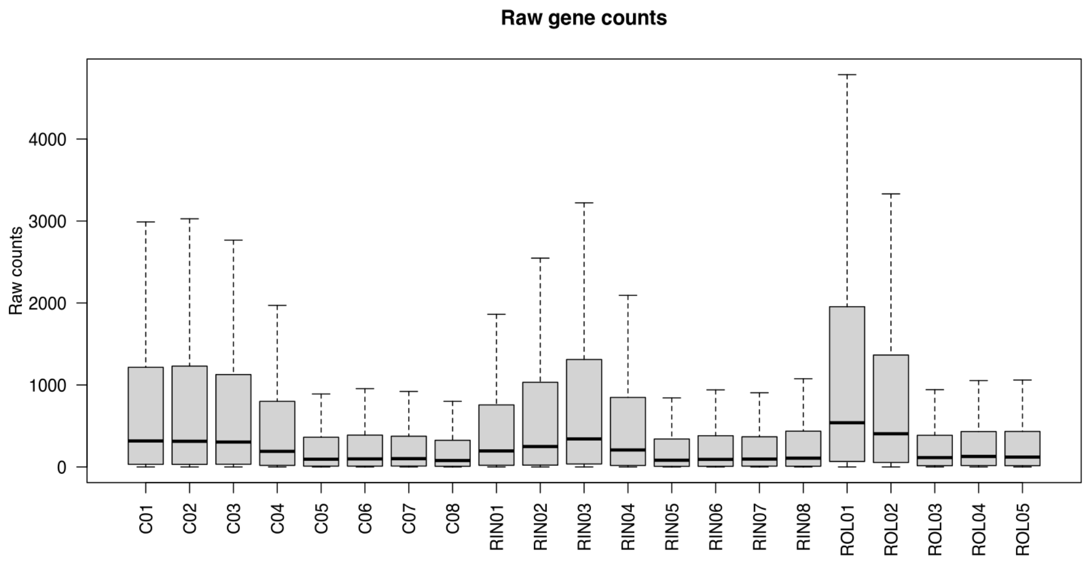


*Distribution of raw gene counts across samples.*

#### Figure 5

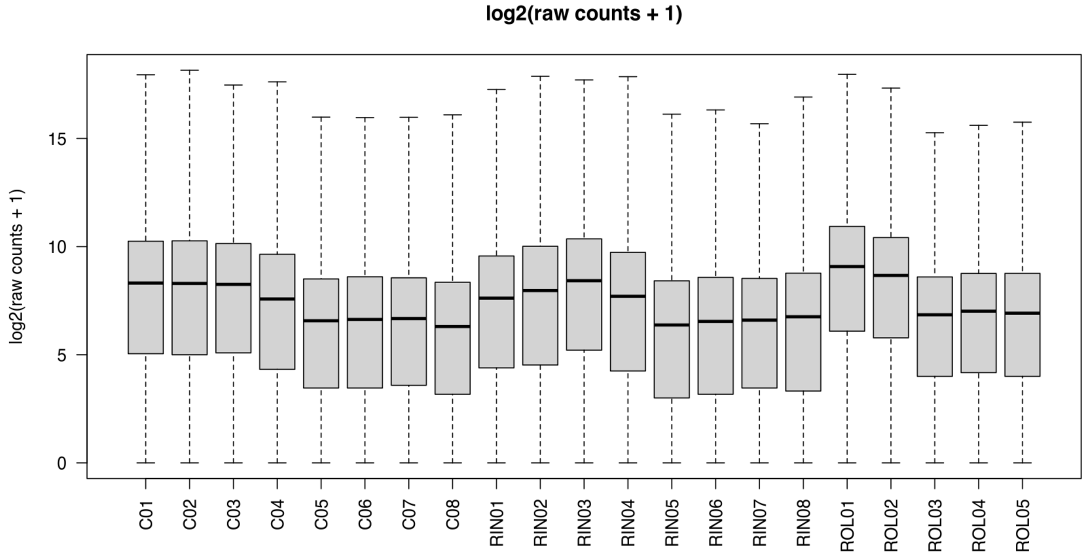


*Distribution of log2(raw counts + 1) across samples.*

#### Figure 6

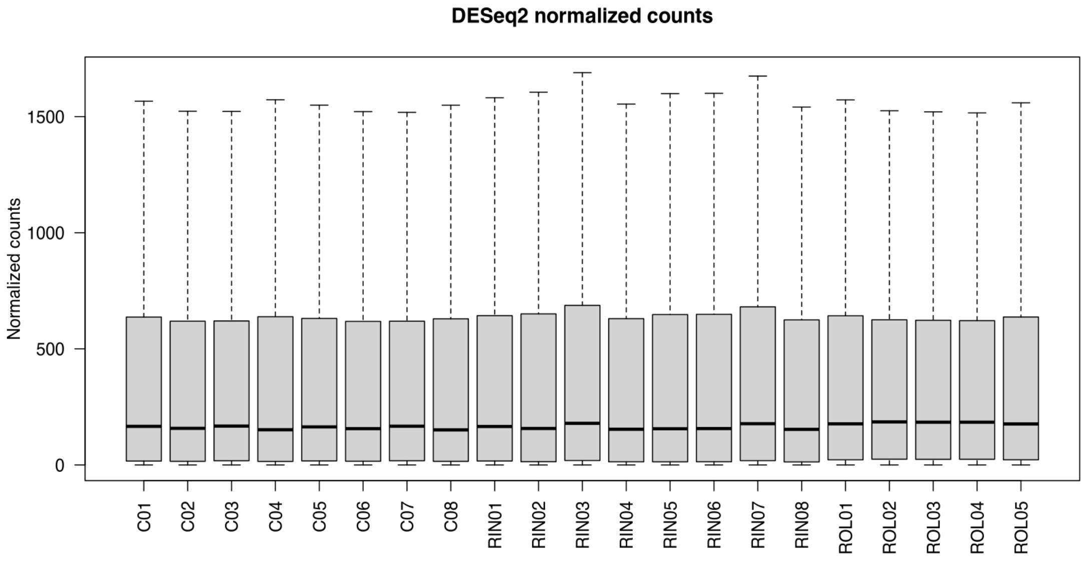


*Distribution of DESeq2 normalized counts across samples.*

#### Figure 7

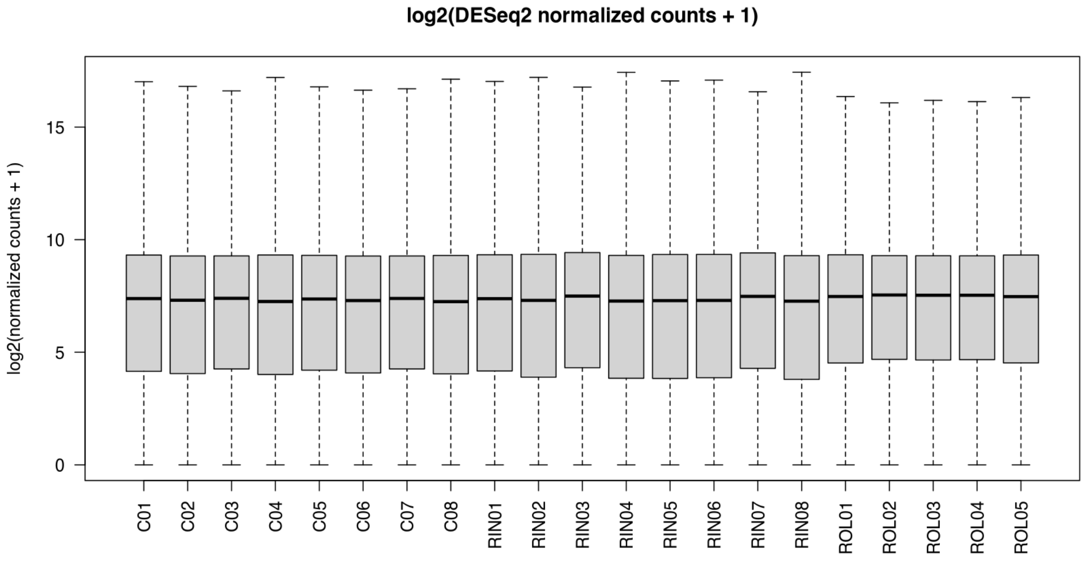


*Distribution of log2(normalized counts + 1) across samples.*

#### Figure 8

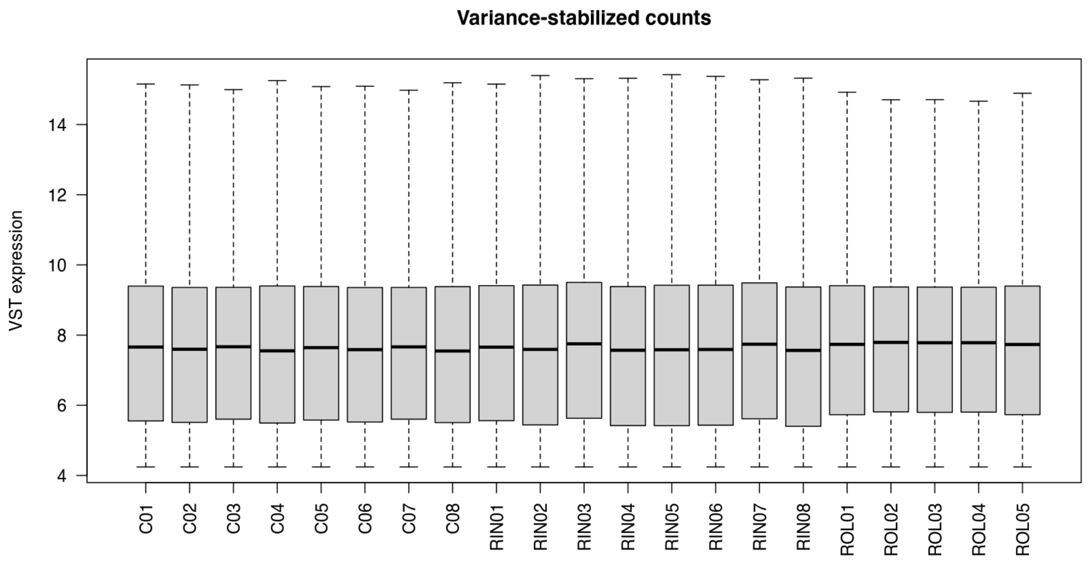


*Distribution of VST-transformed expression values across samples.*

#### Figure 9

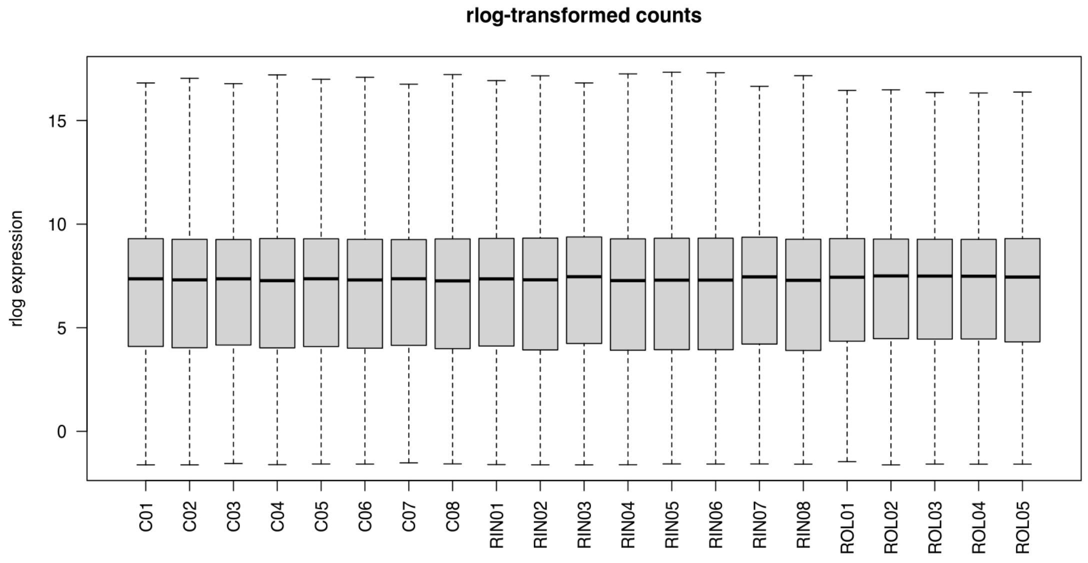


*Distribution of rlog-transformed expression values across samples.*

### Figure 10

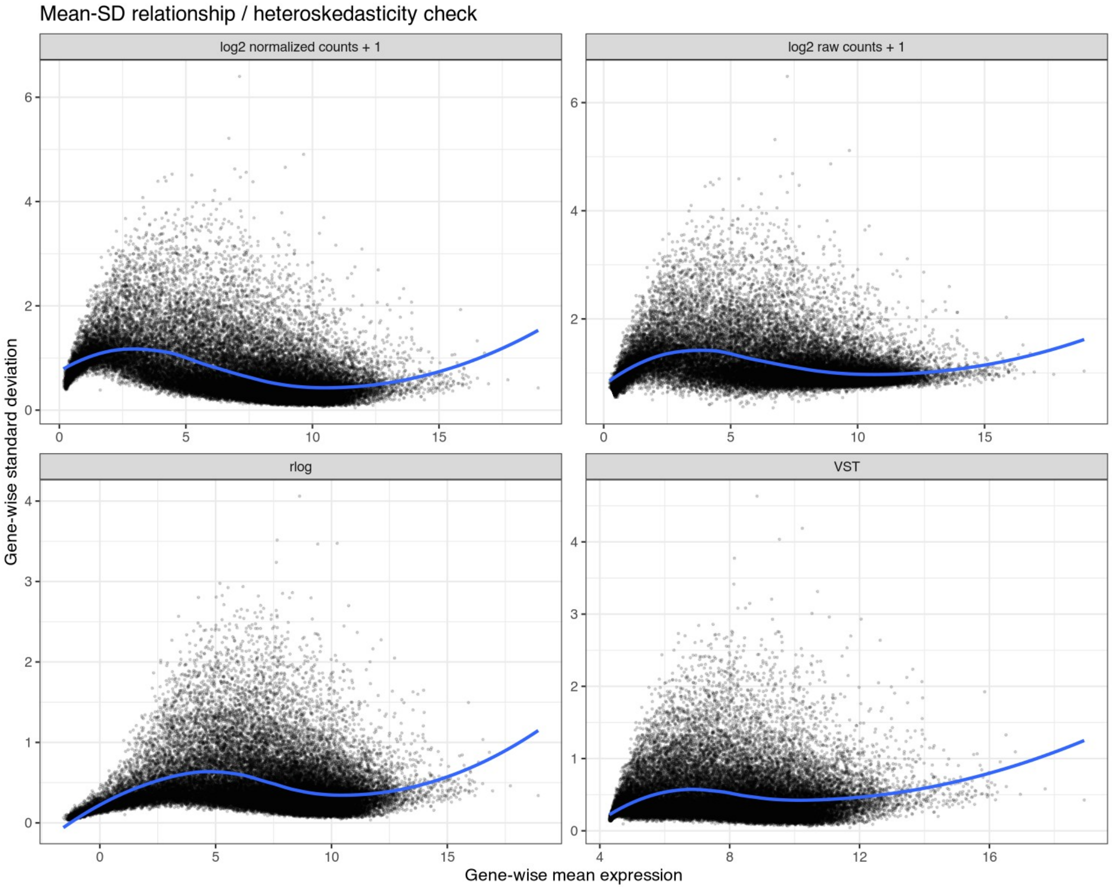


*Mean-SD relationship for raw, normalized, VST, and rlog transformations.*

Figure 11

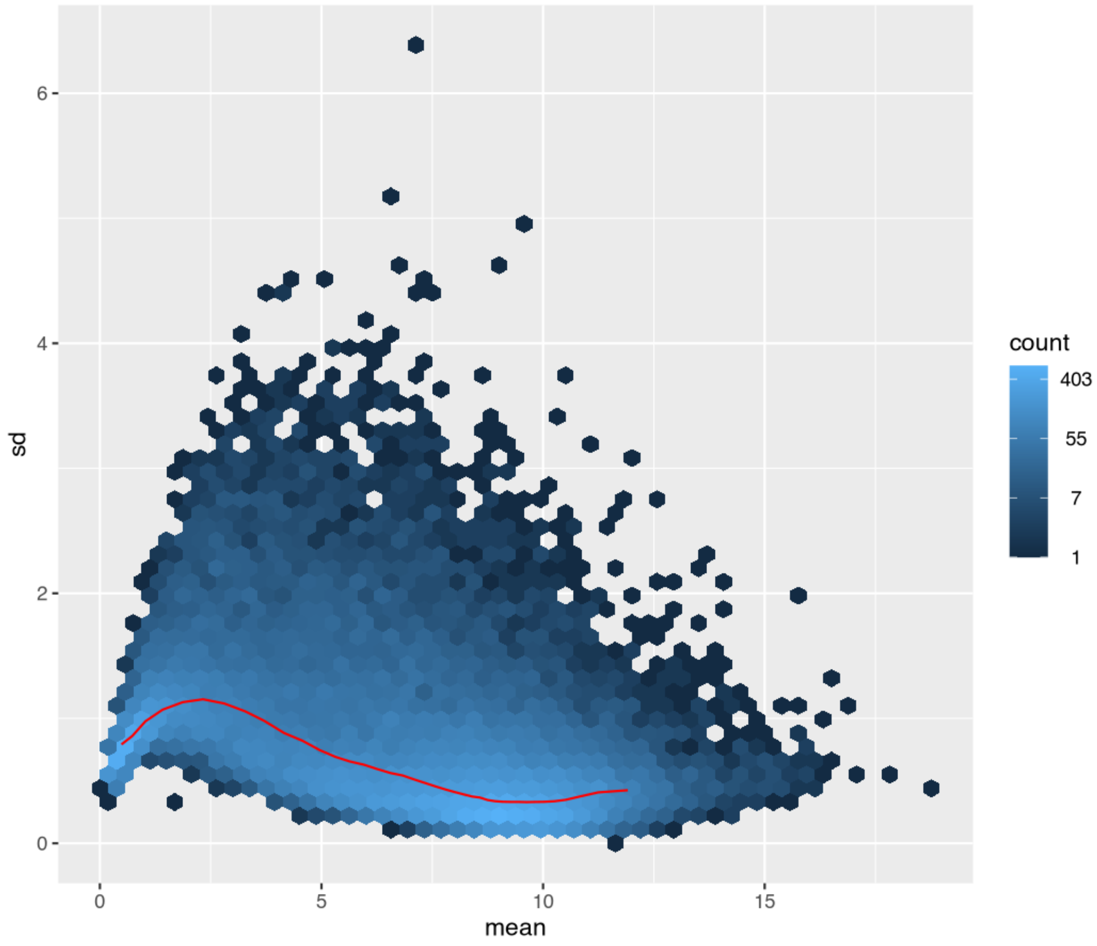


*vsn mean-SD plot for log2 normalized counts.*

#### Figure 12

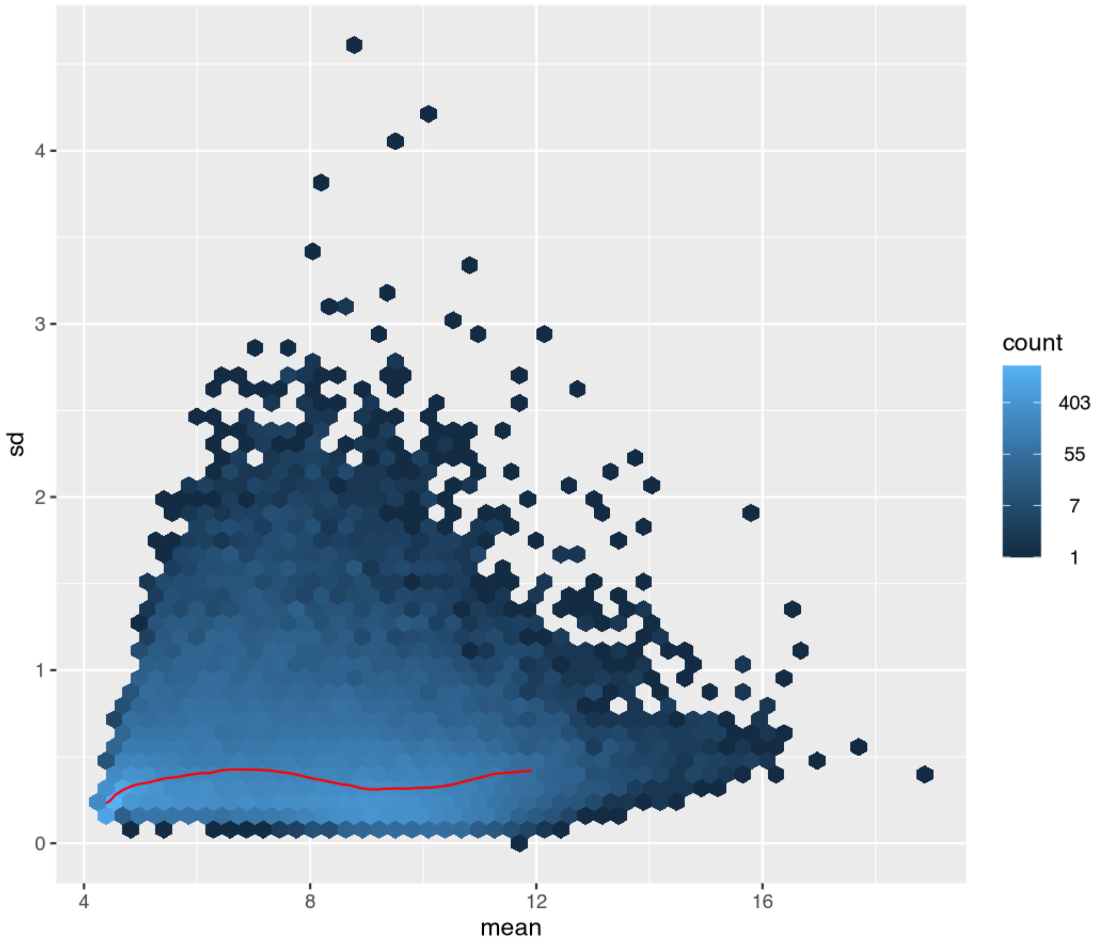


*vsn mean-SD plot for VST values.*

#### Figure 13

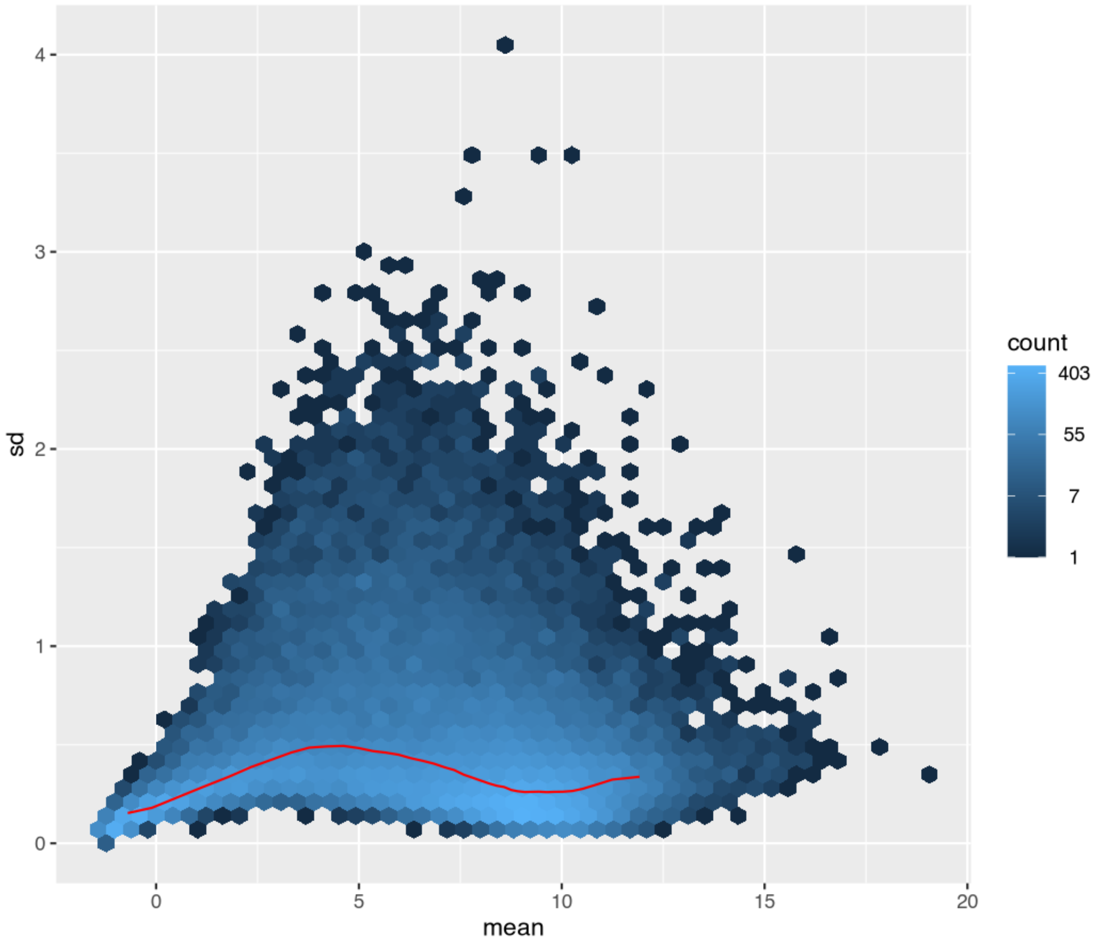


*vsn mean-SD plot for rlog values.*

#### Figure 14

Missing PNG file:
/Volumes/Data_Disk/Ametyst_re-analysis/GDDH/DESeq2/QC_extra_plots/
Pearson_correlation_heatmap_VST.png

#### Figure 15

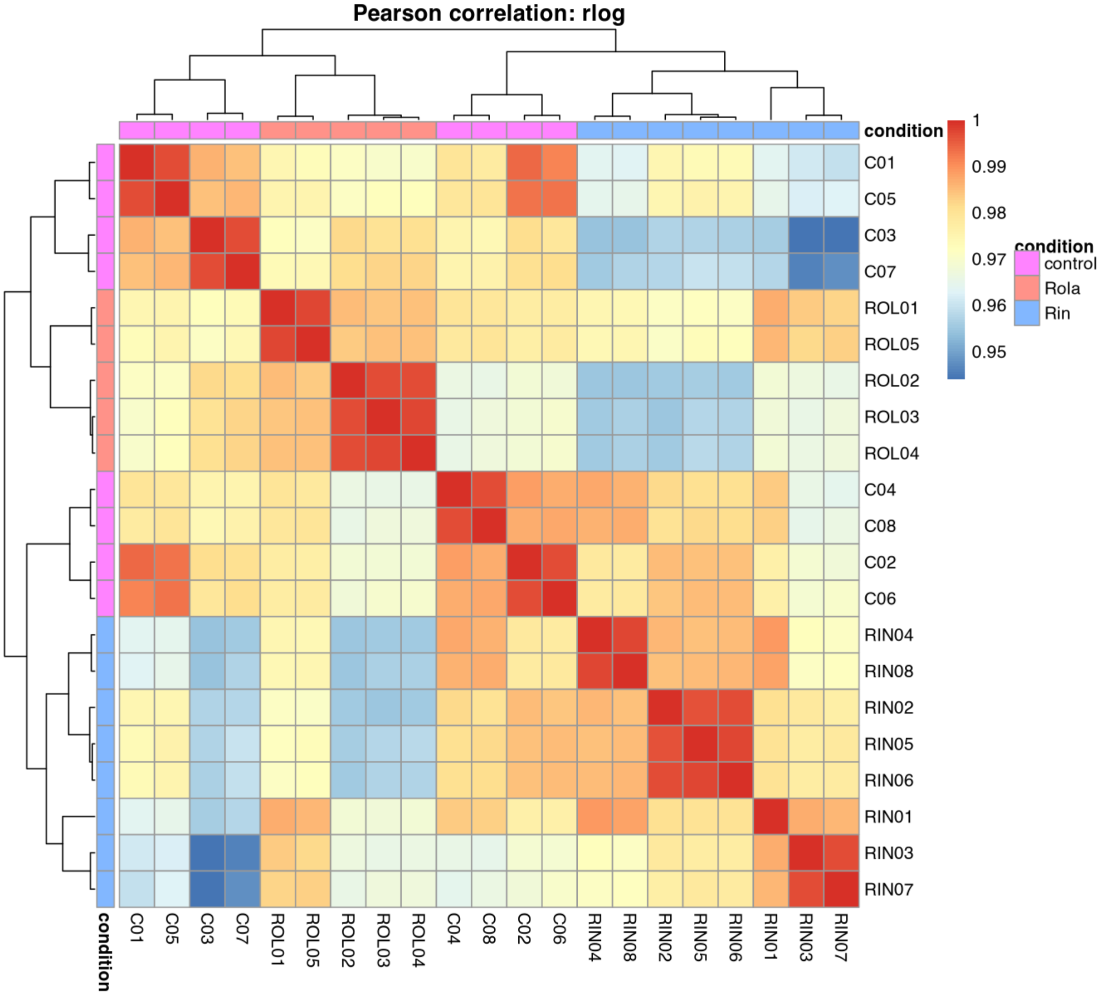


*Pearson correlation heatmap based on rlog-transformed expression values.*

#### Figure 16

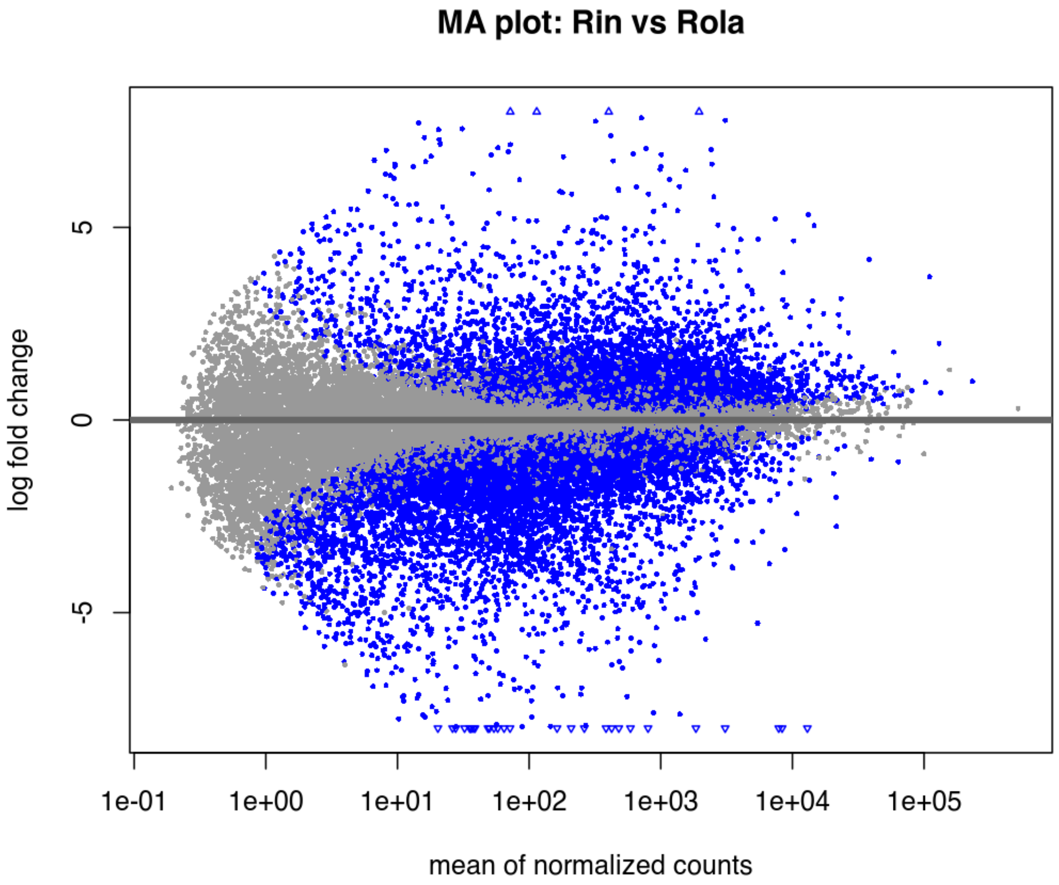


*DESeq2 MA plot for Rin vs Rola.*

#### Figure 17

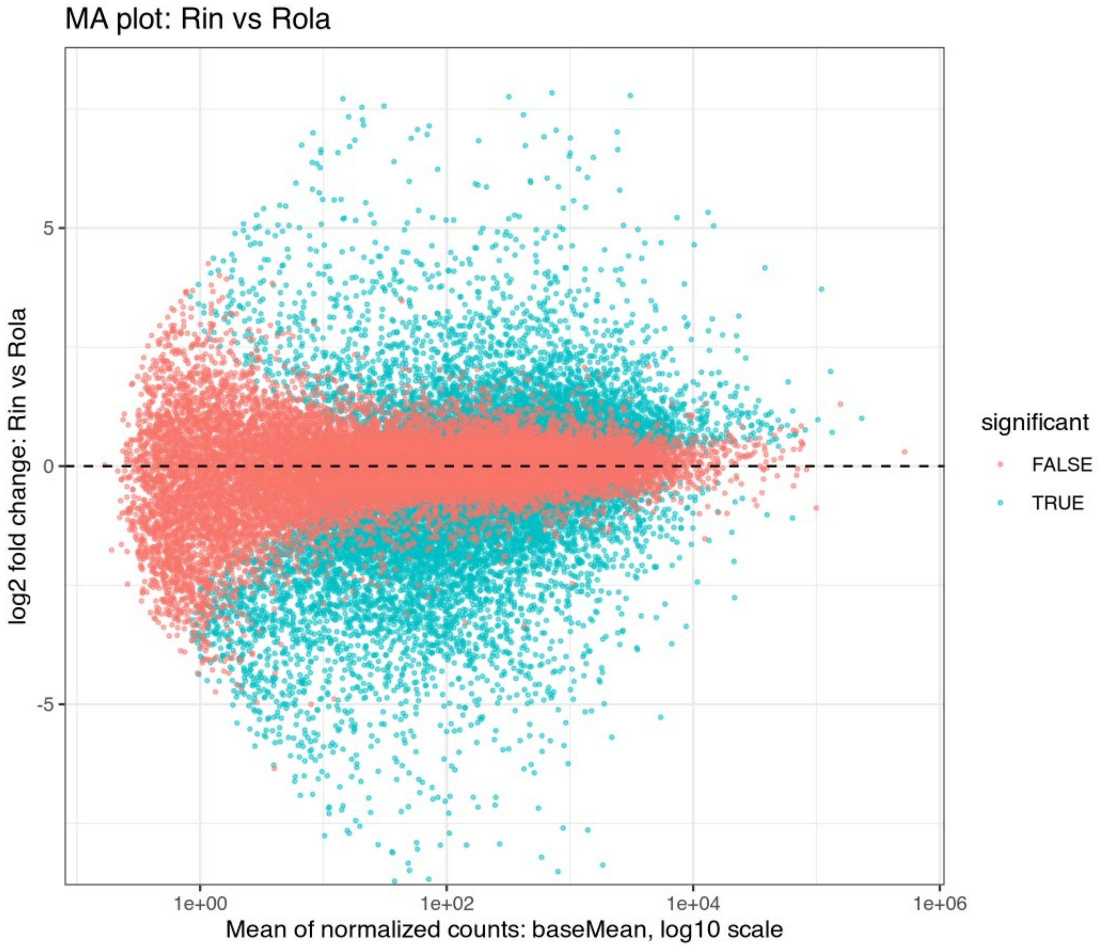


*Custom ggplot MA plot for Rin vs Rola.*

#### Figure 18

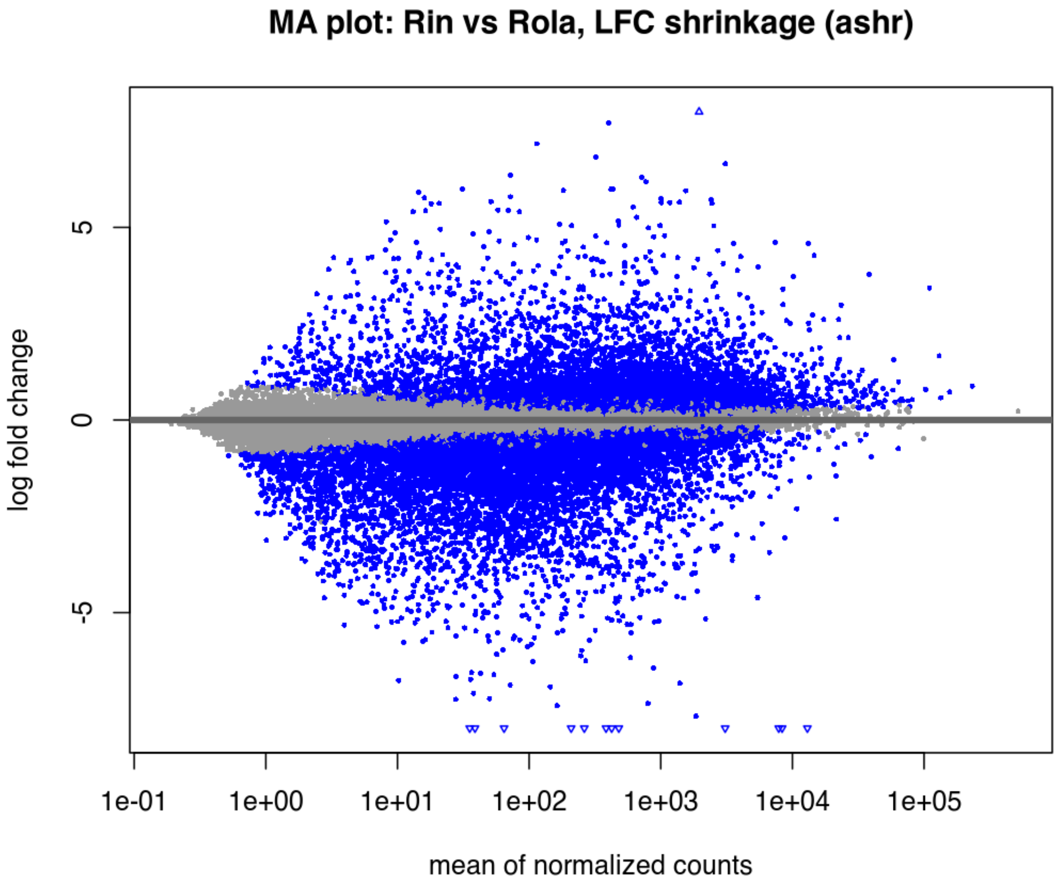


*MA plot using shrunken log2 fold changes for Rin vs Rola, if ashr was available.*

#### Figure 19

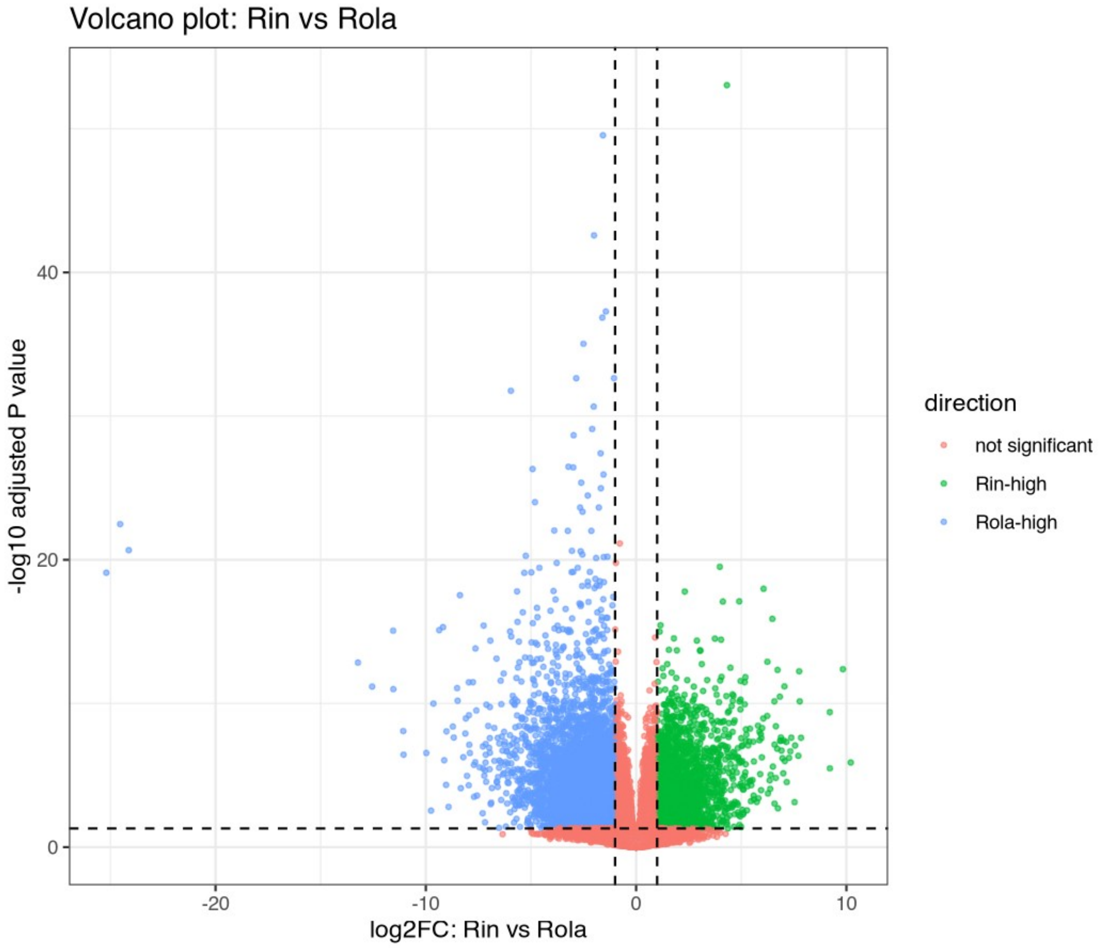


*Volcano plot for Rin vs Rola.*

#### Figure 20

Missing PNG file:
/Volumes/Data_Disk/Ametyst_re-analysis/GDDH/DESeq2/QC_extra_plots/
Heatmap_top500_variable_genes_VST_pheatmap.png

#### Figure 21

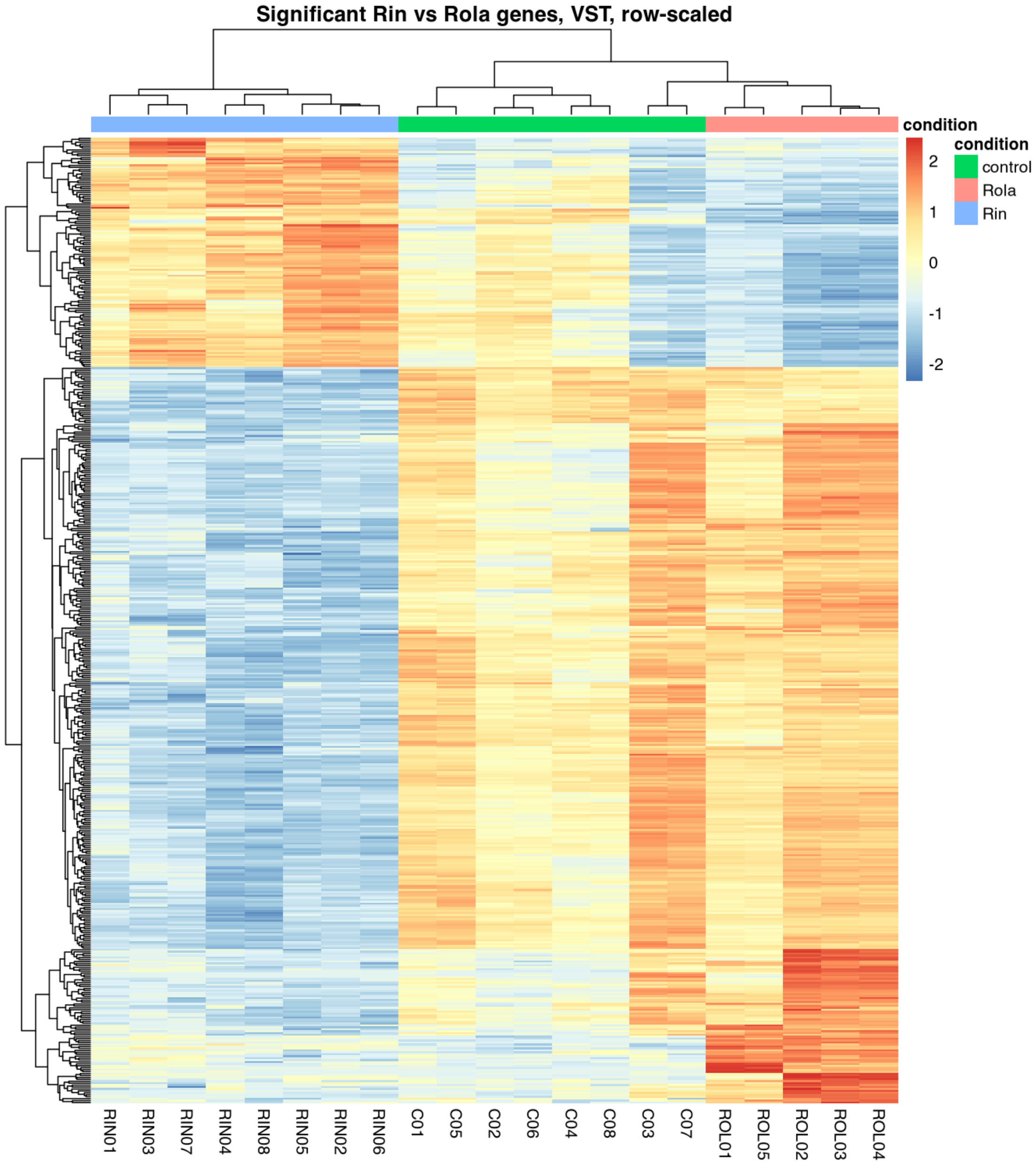


*Heatmap of significant Rin vs Rola genes, if enough genes passed the filter.*

### Figure 22

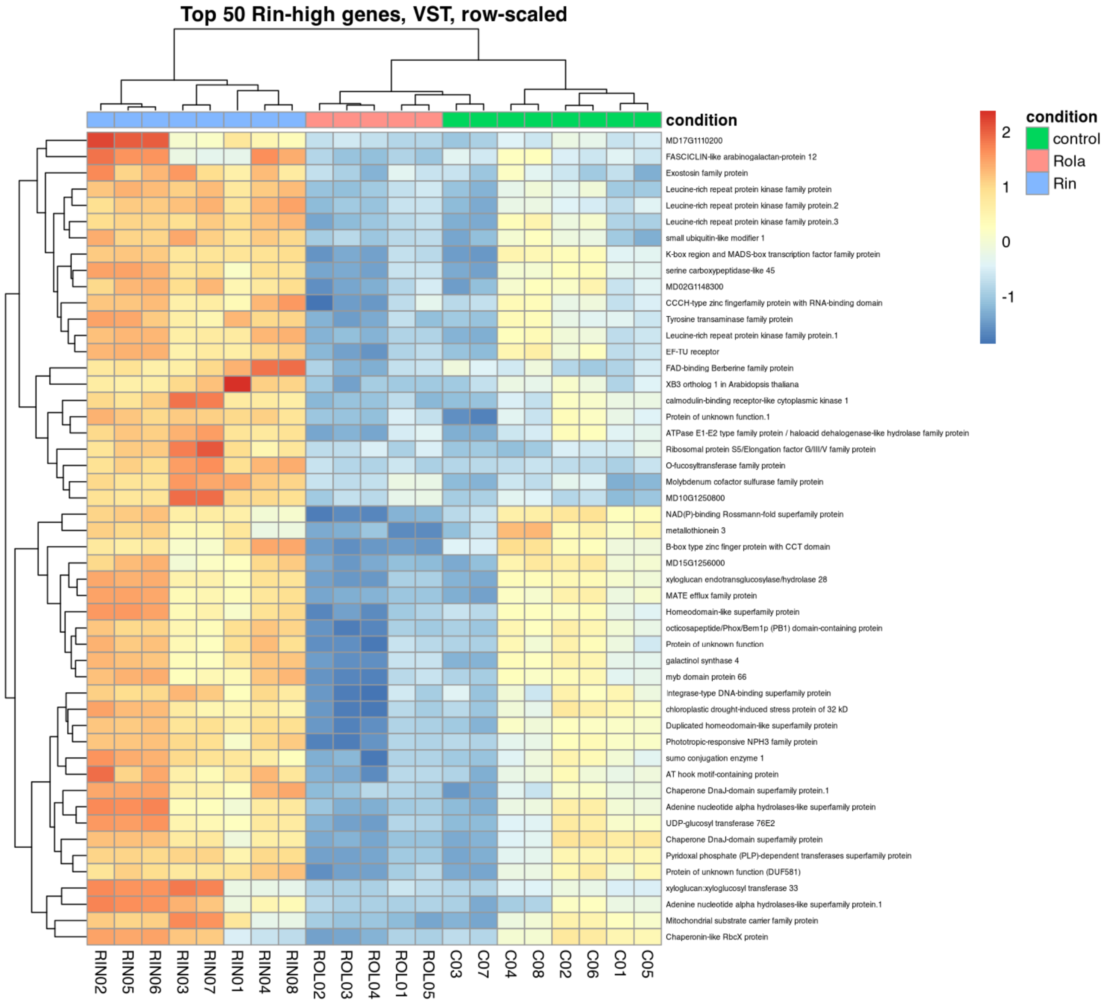


*Heatmap of top Rin-high candidate genes.*

### Figure 23

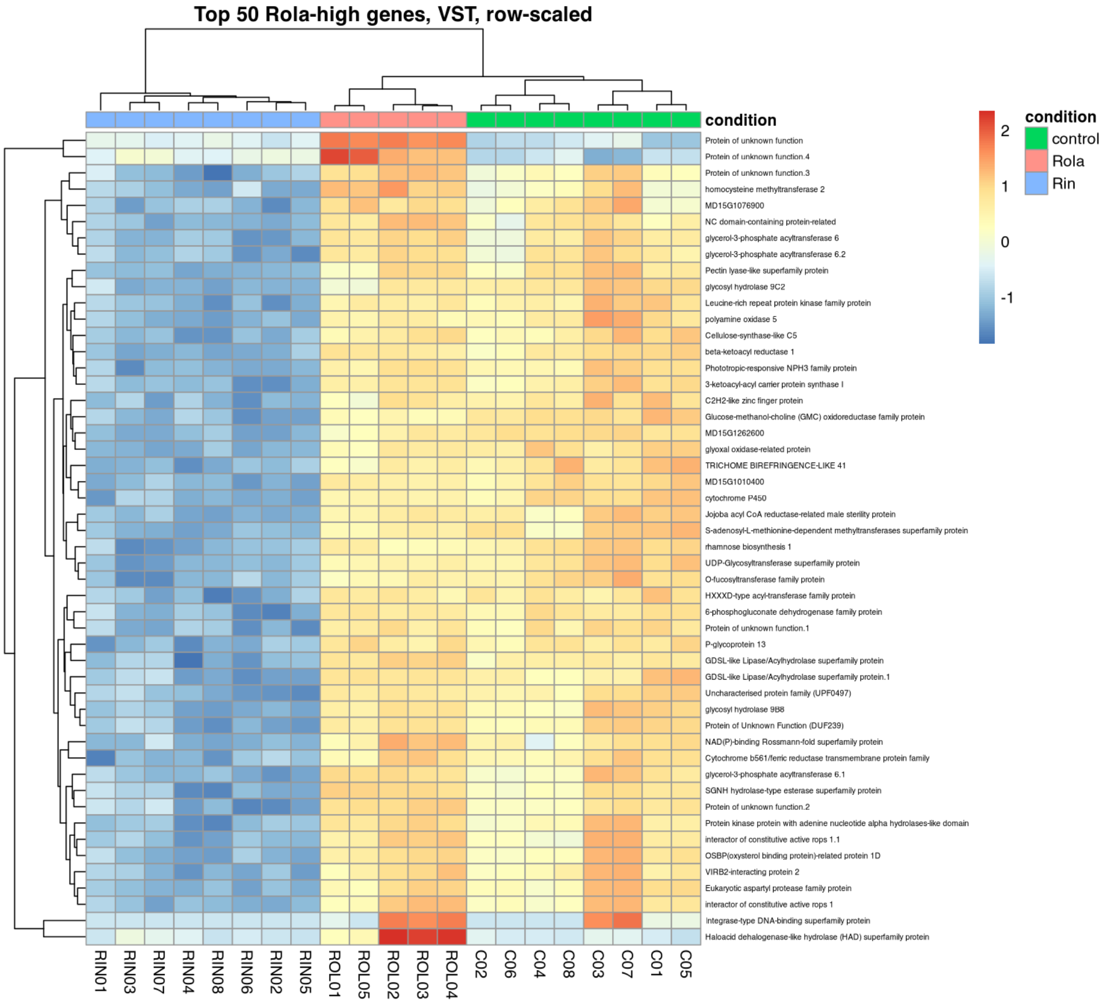


*Heatmap of top Rola-high candidate genes.*

### Figure 24

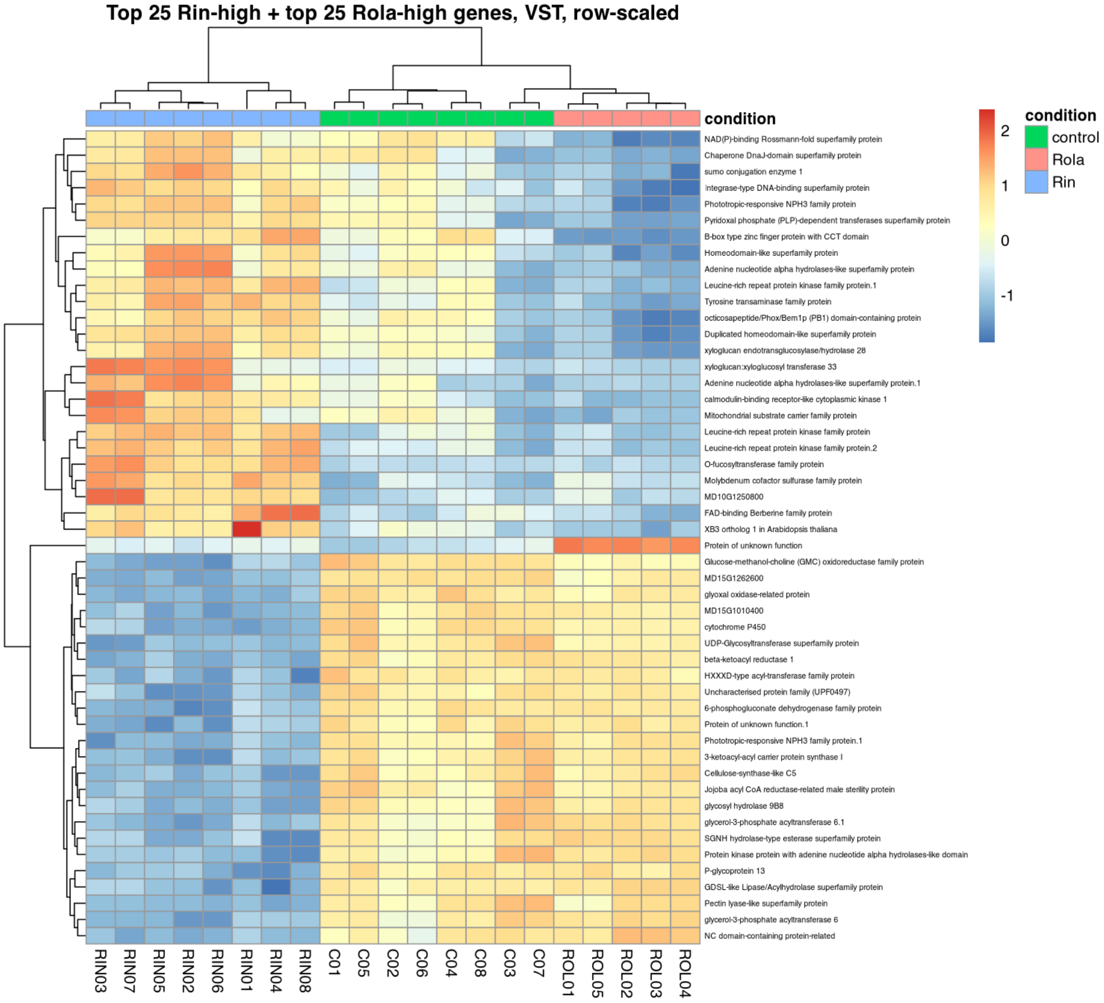


*Combined heatmap of top Rin-high and Rola-high candidate genes.*

### Figure 25

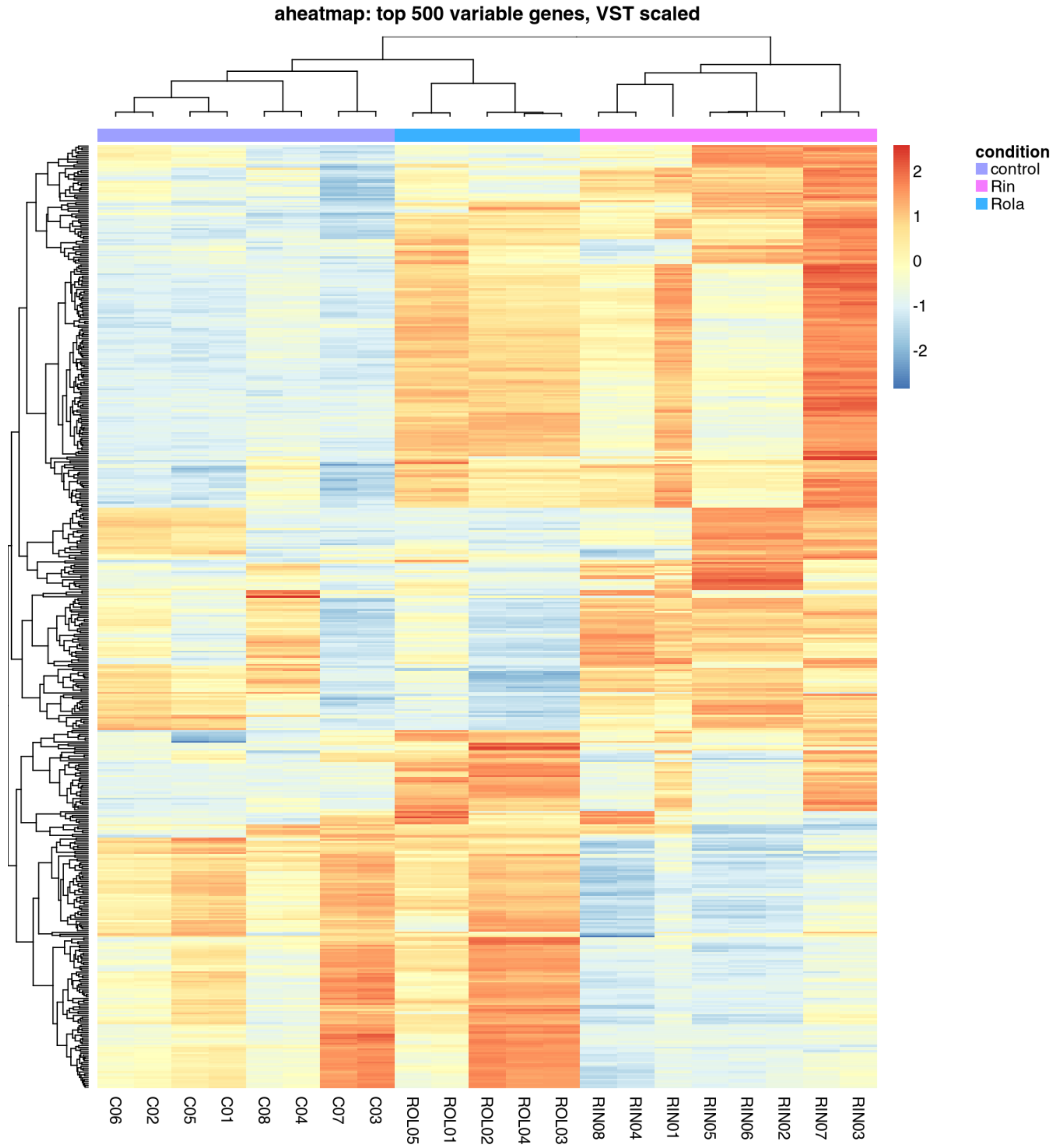


*NMF aheatmap of top variable genes, if NMF was available.*

#### Figure 26

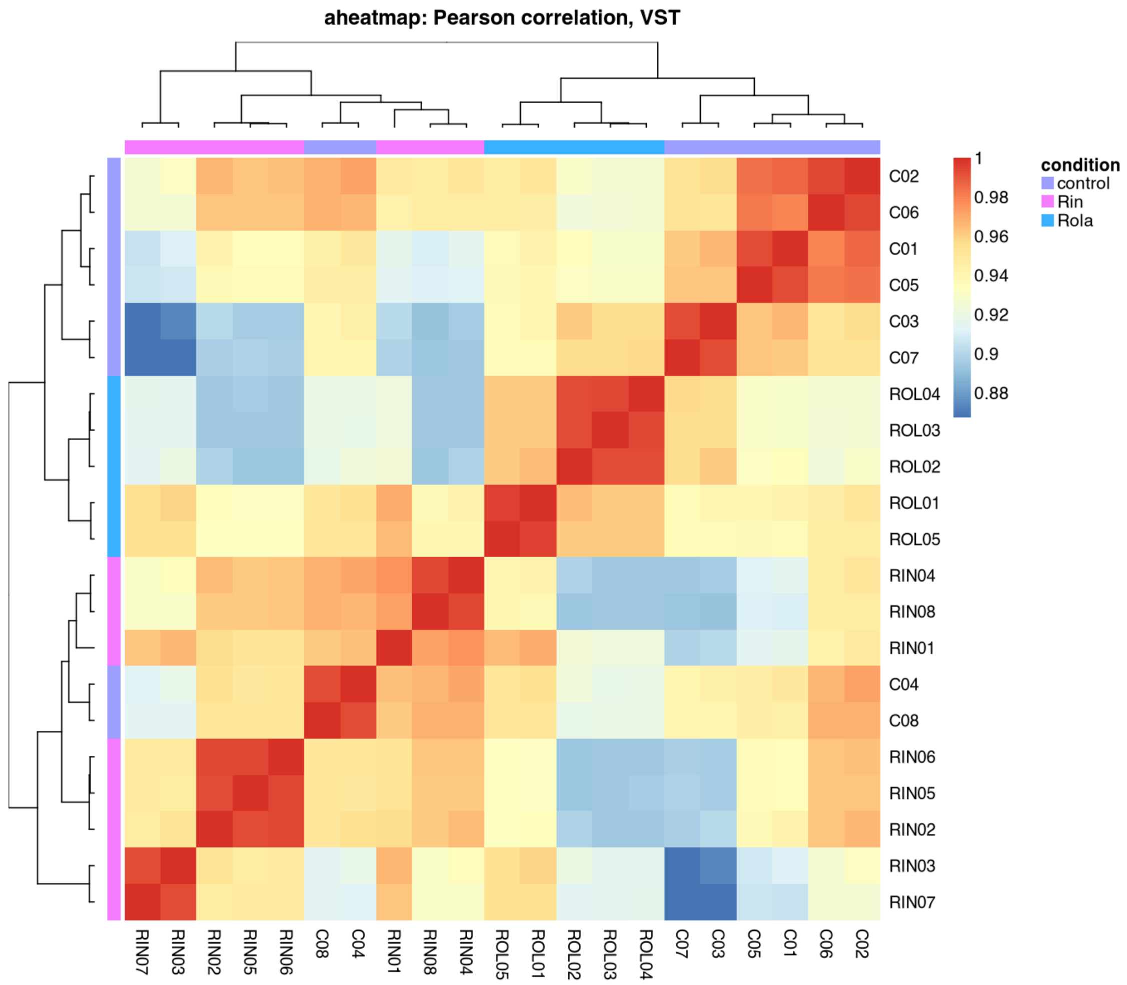


*NMF aheatmap of Pearson correlation matrix, if NMF was available.*

#### Figure 27

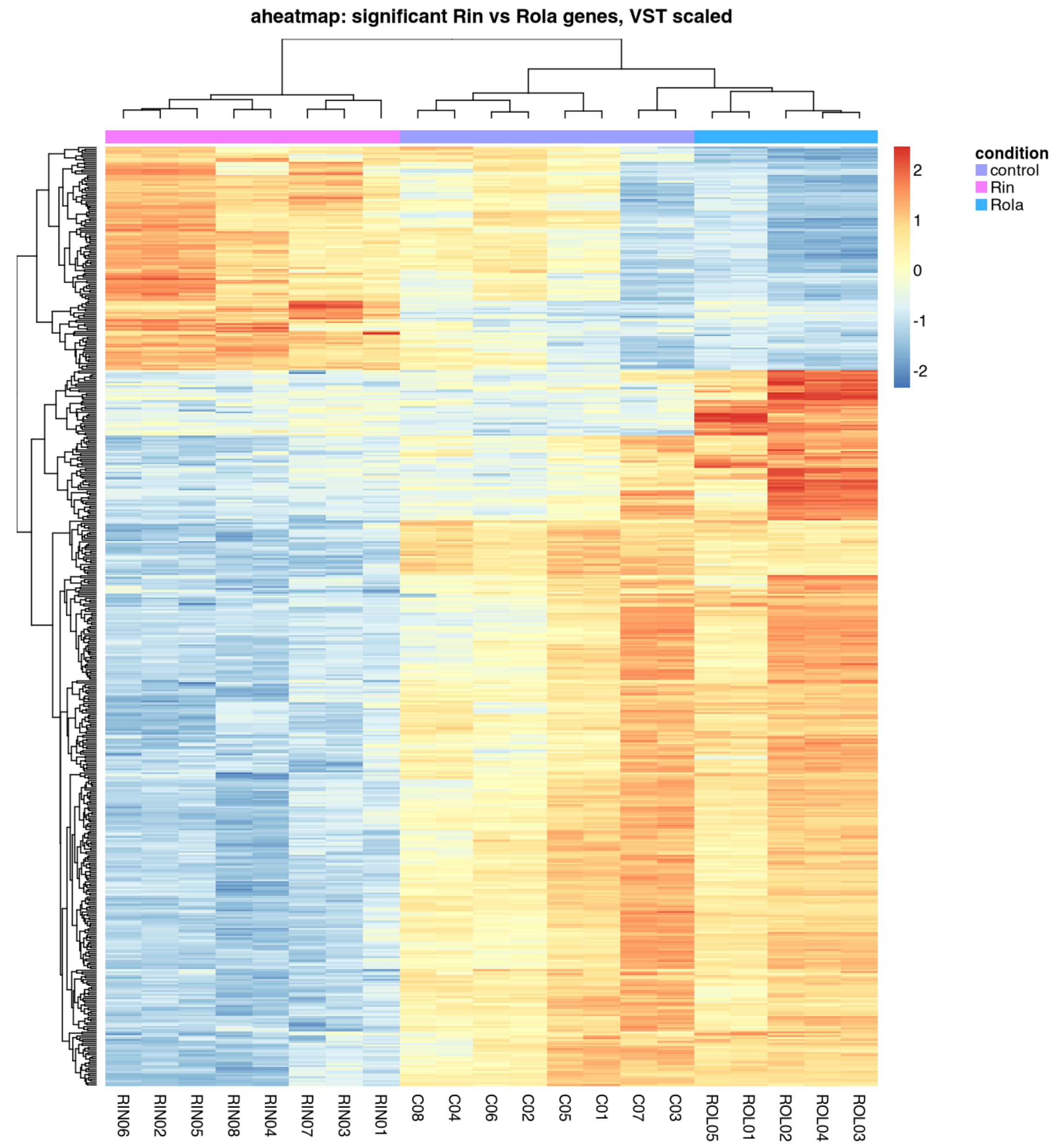


*NMF aheatmap of significant Rin vs Rola genes, if NMF and sufficient genes were available.*

#### Figure 28

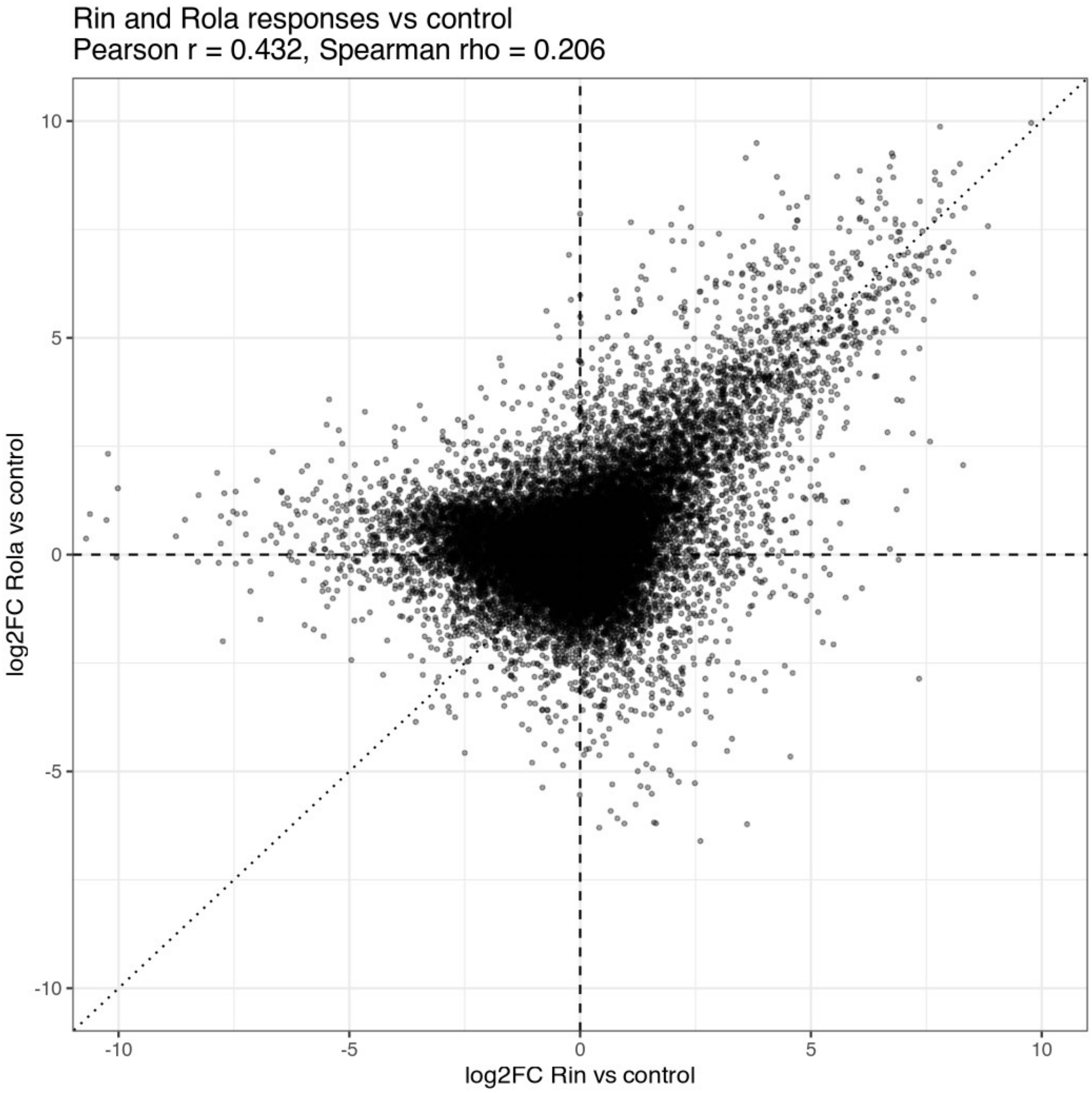


*Scatter plot comparing log2FC Rin vs control and Rola vs control.*

### Table previews

The full tables are written as CSV files in the output directories.
For readability, only the first 10 rows of each table are included in the knitted report.

#### Table preview 1

**Metadata used for DESeq2.**

| sample_id | condition | sample | batch |
|---|---|---|---|
| C01 | control | C01 | batch2_IAB |

| sample_id | condition | sample | batch |
|---|---|---|---|
| C02 | control | C02 | batch2_IAB |
| C03 | control | C03 | batch2_IAB |
| C04 | control | C04 | batch2_IAB |
| C05 | control | C05 | batch1_Eurofins |
| C06 | control | C06 | batch1_Eurofins |
| C07 | control | C07 | batch1_Eurofins |
| C08 | control | C08 | batch1_Eurofins |
| RIN01 | Rin | RIN01 | batch2_IAB |
| RIN02 | Rin | RIN02 | batch2_IAB |

#### Table preview 2

**Rin vs Rola DESeq2 results with gene descriptions.**

| gene_id | gene_name | baseMean | log2FoldChange | lfcSE | stat | pvalue | padj | Description |
|---|---|---|---|---|---|---|---|---|
| MD07G1003800 | MD07G1003800 | 266.7919 | 4.318660 | 0.2682259 | 16.10083 | 0 | 0 | O-fucosyltransferase family protein |
| MD15G1262600 | MD15G1262600 | 2196.0749 | -1.577524 | 0.1014381 | -15.55160 | 0 | 0 | MD15G1262600 |
| MD13G1087700 | MD13G1087700 | 369.9144 | -2.001719 | 0.1384273 | -14.46043 | 0 | 0 | glyoxal oxidase-related protein |
| MD15G1010400 | MD15G1010400 | 970.5317 | -1.438880 | 0.1059982 | -13.57457 | 0 | 0 | MD15G1010400 |
| MD15G1010500 | MD15G1010500 | 385.4605 | -1.605449 | 0.1190456 | -13.48600 | 0 | 0 | P-glycoprotein 13 |

| gene_id | gene_name | baseMean | log2FoldChange | lfcSE | stat | pvalue | padj | Description |
|---|---|---|---|---|---|---|---|---|
| 0 | 0 | | | | | | | |
| MD09G1148500 | MD09G1148500 | 140.6160 | -2.503914 | 0.1902764 | -13.15935 | 0 | 0 | GDSL-like Lipase/Acylhydrolase superfamily protein |
| MD11G1062600 | MD11G1062600 | 258.5170 | -2.849692 | 0.2241246 | -12.71477 | 0 | 0 | UDP-Glycosyltransferase superfamily protein |
| MD16G1107100 | MD16G1107100 | 1696.8125 | -1.050777 | 0.0826495 | -12.71366 | 0 | 0 | beta-ketoacyl reductase 1 |
| MD10G1169100 | MD10G1169100 | 286.2707 | -5.954488 | 0.4746279 | -12.54559 | 0 | 0 | Pectin lyase-like superfamily protein |
| MD12G1082400 | MD12G1082400 | 779.6653 | -2.015843 | 0.1634273 | -12.33480 | 0 | 0 | 6-phosphogluconate dehydrogenase family protein |

#### Table preview 3

**Rin vs control DESeq2 results with gene descriptions.**

| gene_id | gene_name | baseMean | log2FoldChange | lfcSE | stat | pvalue | padj | Description |
|---|---|---|---|---|---|---|---|---|
| MD15G1262600 | MD15G1262600 | 2196.0749 | -1.941514 | 0.0892034 | -21.76503 | 0 | 0 | MD15G1262600 |
| MD07G1003800 | MD07G1003800 | 266.7919 | 4.398128 | 0.2342835 | 18.77267 | 0 | 0 | O-fucosyltransferase family protein |

| gene_id | gene_name | baseMean | log2FoldChange | lfcSE | stat | pvalue | padj | Description |
|---|---|---|---|---|---|---|---|---|
| MD13G1087700 | MD13G1087700 | 369.9144 | -2.304956 | 0.1230072 | -18.73839 | 0 | 0 | glyoxal oxidase-related protein |
| MD04G1069000 | MD04G1069000 | 3981.3492 | -2.358464 | 0.1267419 | -18.60841 | 0 | 0 | Glucose-methanol-choline (GMC) oxidoreductase family protein |
| MD17G1034800 | MD17G1034800 | 2315.2122 | 1.529323 | 0.0842213 | 18.15837 | 0 | 0 | formate dehydrogenase |
| MD11G1062600 | MD11G1062600 | 258.5170 | -3.449746 | 0.1995429 | -17.28824 | 0 | 0 | UDP-Glycosyltransferase superfamily protein |
| MD15G1010400 | MD15G1010400 | 970.5317 | -1.597362 | 0.0935097 | -17.08231 | 0 | 0 | MD15G1010400 |
| MD07G1068700 | MD07G1068700 | 1708.2082 | -2.057509 | 0.1317623 | -15.61531 | 0 | 0 | cytochrome P450 |
| MD17G1140800 | MD17G1140800 | 1990.5436 | -2.690541 | 0.1776100 | -15.14859 | 0 | 0 | rhamnose biosynthesis 1 |
| MD08G1262600 | MD08G1262600 | 1286.1338 | -2.105991 | 0.1412659 | -14.90799 | 0 | 0 | Cellulose-synthase-like C5 |

#### Table preview 4

**Rola vs control DESeq2 results with gene descriptions.**

| gene_id | gene_name | baseMean | log2FoldChange | lfcSE | stat | pvalue | padj | Description |
|---|---|---|---|---|---|---|---|---|
| MD02G1080400 | MD02G1080400 | 59.01762 | 6.347044 | 0.4854996 | 13.07322 | 0 | 0 | Protein of unknown function |
| MD15G1211600 | MD15G1211600 | 260.95269 | 3.109105 | 0.2409064 | 12.90586 | 0 | 0 | Protein of unknown function |
| MD03G1152000 | MD03G1152000 | 226.04955 | 3.603033 | 0.3039801 | 11.85286 | 0 | 0 | Polynucleotidyl transferase |
| MD05G1002900 | MD05G1002900 | 448.53685 | 2.327532 | 0.2030502 | 11.46284 | 0 | 0 | Leucine-rich repeat (LRR) family protein |
| MD09G1084600 | MD09G1084600 | 46.42252 | 7.241337 | 0.6401450 | 11.31203 | 0 | 0 | allene oxide cyclase 4 |
| MD05G1311400 | MD05G1311400 | 1032.53935 | 2.111644 | 0.2016304 | 10.47284 | 0 | 0 | erf domain protein 9 |
| MD15G1032200 | MD15G1032200 | 441.31502 | 2.297057 | 0.2191460 | 10.48185 | 0 | 0 | NAC domain containing protein 28 |
| MD17G1281200 | MD17G1281200 | 197.10879 | 2.968891 | 0.2860598 | 10.37857 | 0 | 0 | tolB protein-related |
| MD08G1086500 | MD08G1086500 | 338.97160 | 4.701015 | 0.4583110 | 10.25726 | 0 | 0 | salt tolerance zinc finger |
| MD08G1175700 | MD08G1175700 | 73.21173 | 1.908834 | 0.1861483 | 10.25437 | 0 | 0 | MD08G1175700 |

#### Table preview 5

**Rin vs Rola DESeq2 results, backward-compatible filename.**

| gene_id | gene_name | baseMean | log2FoldChange | lfcSE | stat | pvalue | padj | Description |
|---|---|---|---|---|---|---|---|---|
| MD07G1003800 | MD07G1003800 | 266.7919 | 4.318660 | 0.2682259 | 16.10083 | 0 | 0 | O-fucosyltransferase family protein |
| MD15G1262600 | MD15G1262600 | 2196.0749 | -1.577524 | 0.1014381 | -15.55160 | 0 | 0 | MD15G1262600 |
| MD13G1087700 | MD13G1087700 | 369.9144 | -2.001719 | 0.1384273 | -14.46043 | 0 | 0 | glyoxal oxidase-related protein |
| MD15G1010400 | MD15G1010400 | 970.5317 | -1.438880 | 0.1059982 | -13.57457 | 0 | 0 | MD15G1010400 |
| MD15G1010500 | MD15G1010500 | 385.4605 | -1.605449 | 0.1190456 | -13.48600 | 0 | 0 | P-glycoprotein 13 |
| MD09G1148500 | MD09G1148500 | 140.6160 | -2.503914 | 0.1902764 | -13.15935 | 0 | 0 | GDSL-like Lipase/Acylhydrolase superfamily protein |
| MD11G1062600 | MD11G1062600 | 258.5170 | -2.849692 | 0.2241246 | -12.71477 | 0 | 0 | UDP-Glycosyltransferase superfamily protein |
| MD16G1107100 | MD16G1107100 | 1696.8125 | -1.050777 | 0.0826495 | -12.71366 | 0 | 0 | beta-ketoacyl reductase 1 |
| MD10G1169100 | MD10G1169100 | 286.2707 | -5.954488 | 0.4746279 | -12.54559 | 0 | 0 | Pectin lyase-like superfamily protein |
| MD12G10 | MD12G10 | 779.66 | -2.015843 | 0.1634 | -12.334 | 0 | 0 | 6-phosphogluconate dehydrogenase family |

| gene_id | gene_name | baseMean | log2FoldChange | lfcSE | stat | pvalue | padj | Description |
|---|---|---|---|---|---|---|---|---|
| 82400 | 82400 | 53 | | 273 | 80 | | | protein |

#### Table preview 6

**Rin vs control DESeq2 results, backward-compatible filename.**

| gene_id | gene_name | baseMean | log2FoldChange | lfcSE | stat | pvalue | padj | Description |
|---|---|---|---|---|---|---|---|---|
| MD15G1262600 | MD15G1262600 | 2196.0749 | -1.941514 | 0.0892034 | -21.76503 | 0 | 0 | MD15G1262600 |
| MD07G1003800 | MD07G1003800 | 266.7919 | 4.398128 | 0.2342835 | 18.77267 | 0 | 0 | O-fucosyltransferase family protein |
| MD13G1087700 | MD13G1087700 | 369.9144 | -2.304956 | 0.1230072 | -18.73839 | 0 | 0 | glyoxal oxidase-related protein |
| MD04G1069000 | MD04G1069000 | 3981.3492 | -2.358464 | 0.1267419 | -18.60841 | 0 | 0 | Glucose-methanol-choline (GMC) oxidoreductase family protein |
| MD17G1034800 | MD17G1034800 | 2315.2122 | 1.529323 | 0.0842213 | 18.15837 | 0 | 0 | formate dehydrogenase |
| MD11G1062600 | MD11G1062600 | 258.5170 | -3.449746 | 0.1995429 | -17.28824 | 0 | 0 | UDP-Glycosyltransferase superfamily protein |
| MD15G1010400 | MD15G1010400 | 970.531 | -1.597362 | 0.09350 | -17.0823 | 0 | 0 | MD15G1010400 |

| gene_id | gene_name | baseMean | log2FoldChange | lfcSE | stat | pvalue | padj | Description |
|---|---|---|---|---|---|---|---|---|
| 00 | 00 | 7 | | 97 | 1 | | | |
| MD07G106870000 | MD07G106870000 | 1708.2082 | -2.057509 | 0.1317623 | -15.61531 | 0 | 0 | cytochrome P450 |
| MD17G114080000 | MD17G114080000 | 1990.5436 | -2.690541 | 0.1776100 | -15.14859 | 0 | 0 | rhamnose biosynthesis 1 |
| MD08G112620000 | MD08G112620000 | 1286.1338 | -2.105991 | 0.1412659 | -14.90799 | 0 | 0 | Cellulose-synthase-like C5 |

#### Table preview 7

**Rola vs control DESeq2 results, backward-compatible filename.**

| gene_id | gene_name | baseMean | log2FoldChange | lfcSE | stat | pvalue | padj | Description |
|---|---|---|---|---|---|---|---|---|
| MD02G1080400 | MD02G1080400 | 59.01762 | 6.347044 | 0.4854996 | 13.07322 | 0 | 0 | Protein of unknown function |
| MD15G1211600 | MD15G1211600 | 260.95269 | 3.109105 | 0.2409064 | 12.90586 | 0 | 0 | Protein of unknown function |
| MD03G1152000 | MD03G1152000 | 226.04955 | 3.603033 | 0.3039801 | 11.85286 | 0 | 0 | Polynucleotidyl transferase |
| MD05G1002900 | MD05G1002900 | 448.53685 | 2.327532 | 0.2030502 | 11.46284 | 0 | 0 | Leucine-rich repeat (LRR) family protein |
| MD09G1084600 | MD09G1084600 | 46.42252 | 7.241337 | 0.6401450 | 11.31203 | 0 | 0 | allene oxide cyclase 4 |
| MD05G131 | MD05G131 | 1032.539 | 2.111644 | 0.20163 | 10.472 | 0 | 0 | erf domain protein 9 |

| gene_id | gene_name | baseMean | log2FoldChange | lfcSE | stat | pvalue | padj | Description |
|---|---|---|---|---|---|---|---|---|
| 1400 | 1400 | 35 | | 04 | 84 | | | |
| MD15G1032200 | MD15G1032200 | 441.31502 | 2.297057 | 0.2191460 | 10.48185 | 0 | 0 | NAC domain containing protein 28 |
| MD17G1281200 | MD17G1281200 | 197.10879 | 2.968891 | 0.2860598 | 10.37857 | 0 | 0 | tolB protein-related |
| MD08G1086500 | MD08G1086500 | 338.97160 | 4.701015 | 0.4583110 | 10.25726 | 0 | 0 | salt tolerance zinc finger |
| MD08G1175700 | MD08G1175700 | 73.21173 | 1.908834 | 0.1861483 | 10.25437 | 0 | 0 | MD08G1175700 |

#### Table preview 8

**Summary counts of significant DE genes.**

| contrast | padj_0.05 | padj_0.05_absLFC1 |
|---|---|---|
| Rin_vs_Rola | 14233 | 7438 |
| Rin_vs_control | 13975 | 7029 |
| Rola_vs_control | 8296 | 4123 |

#### Table preview 9

**DESeq2 normalized count matrix.**

| | C01 | C02 | C03 | C04 | C05 | C06 | C07 | C08 | RIN01 | RIN02 | RIN03 | RIN04 | RIN05 | RIN06 | RIN07 | RIN08 | ROL01 | ROL02 | ROL03 | ROL04 | ROL05 |
|---|---|---|---|---|---|---|---|---|---|---|---|---|---|---|---|---|---|---|---|---|---|
| MD00G1000 | 1.571840 | 2.012678 | 0.000000 | 0.7977415 | 3.485247 | 0.000000 | 0.000000 | 0.000000 | 0.8491195 | 1.2604314 | 1.048409 | 1.485409 | 0.000000 | 0.00000 | 0.000000 | 0.000000 | 0.985396 | 1.831184 | 3.22820 | 5.752355 | 0.000000 |

| | C001 | C002 | C003 | C004 | C005 | C006 | C007 | C008 | RIN001 | RIN002 | RIN003 | RIN004 | RIN005 | RIN006 | RIN007 | RIN008 | ROL001 | ROL002 | ROL003 | ROL004 | ROL005 |
|---|---|---|---|---|---|---|---|---|---|---|---|---|---|---|---|---|---|---|---|---|---|
| 100 | | | | | | | | | | | | | | | | | | | | | |
| MD00G1000200 | 163.995253 | 124.786040 | 138.073177 | 112.4815506 | 139.409899 | 92.395917 | 115.517352 | 81.316097 | 73.0242777 | 121.6316301 | 76.533863 | 71.299633 | 89.248866 | 97.03338 | 90.652899 | 61.588553 | 107.408159 | 123.604923 | 61.33581 | 116.485188 | 80.902936 |
| MD00G1000300 | 8.907090 | 9.057051 | 4.950831 | 21.5390203 | 5.227871 | 6.372132 | 1.650248 | 9.680488 | 5.9438366 | 10.0834512 | 10.484091 | 14.111386 | 15.191296 | 10.21404 | 0.000000 | 14.322919 | 3.613119 | 6.409144 | 0.00000 | 5.752355 | 4.412887 |
| MD00G1000500 | 8.383144 | 7.044373 | 23.653971 | 4.7864490 | 6.970495 | 14.337298 | 24.753718 | 11.616585 | 7.6420756 | 0.6302157 | 12.580909 | 4.456227 | 3.797824 | 3.40468 | 9.250296 | 4.296876 | 11.824751 | 19.227432 | 19.36920 | 18.695154 | 10.296737 |

| | C001 | C002 | C003 | C004 | C005 | C006 | C007 | C008 | RIN001 | RIN002 | RIN003 | RIN004 | RIN005 | RIN006 | RIN007 | RIN008 | ROL001 | ROL002 | ROL003 | ROL004 | ROL005 |
|---|---|---|---|---|---|---|---|---|---|---|---|---|---|---|---|---|---|---|---|---|---|
| MD00G1000600 | 1225.510853 | 2376.972789 | 1967.130208 | 2985.1486680 | 1164.072661 | 2383.177447 | 2079.312335 | 2896.401926 | 2362.2504721 | 2338.7304628 | 1339.866808 | 2181.323144 | 2690.758363 | 2306.67062 | 1313.542009 | 2277.344173 | 1498.458787 | 1648.523440 | 1815.86266 | 1704.135151 | 1509.207489 |
| MD00G1000700 | 135.702142 | 102.646581 | 103.417360 | 66.2125439 | 132.439405 | 111.512314 | 143.571566 | 83.252194 | 89.1575484 | 71.8445898 | 110.607158 | 69.814224 | 102.541250 | 98.73572 | 101.753254 | 67.317721 | 91.970289 | 118.111371 | 156.56771 | 113.609010 | 80.902936 |
| MD00G1000800 | 539.664890 | 610.847791 | 513.236153 | 811.3030987 | 543.698608 | 659.515684 | 567.685272 | 931.262919 | 797.3232183 | 764.4516441 | 699.813063 | 823.659301 | 770.958289 | 743.92255 | 797.375501 | 807.812650 | 727.550682 | 553.017583 | 694.06306 | 591.054470 | 685.468508 |
| MD00 | 16.2 | 12.0 | 18.1 | 11.96 | 13.9 | 6.37 | 21.4 | 7.74 | 12.73 | 10.71 | 13.6 | 8.91 | 7.59 | 15.3 | 20.3 | 10.0 | 14.1 | 17.8 | 12.9 | 17.2 | 16.1 |

| | C01 | C02 | C03 | C04 | C05 | C06 | C07 | C08 | RIN01 | RIN02 | RIN03 | RIN04 | RIN05 | RIN06 | RIN07 | RIN08 | ROL01 | ROL02 | ROL03 | ROL04 | ROL05 |
|---|---|---|---|---|---|---|---|---|---|---|---|---|---|---|---|---|---|---|---|---|---|
| G1000900 | 42341 | 76068 | 53047 | 61224 | 40990 | 2132 | 53222 | 4390 | 67926 | 36669 | 29318 | 2454 | 5648 | 2106 | 50651 | 26044 | 24009 | 54045 | 1280 | 57065 | 80587 |
| MD00G1001000 | 85.927225 | 70.946902 | 102.867268 | 77.3809249 | 102.814801 | 111.512314 | 90.763634 | 52.274634 | 87.4593094 | 67.4330799 | 80.727499 | 93.580768 | 91.147778 | 76.60530 | 66.602130 | 88.802100 | 95.583408 | 118.111371 | 124.28571 | 109.294744 | 77.961011 |
| MD00G1001100 | 5.239465 | 2.515848 | 11.001847 | 6.3819319 | 5.227871 | 0.000000 | 16.502479 | 3.872195 | 3.3964780 | 4.4115099 | 7.863068 | 5.941636 | 3.797824 | 3.40468 | 11.100355 | 11.458336 | 8.211633 | 13.733880 | 11.29870 | 12.942799 | 8.825775 |

#### Table preview 10

**PCA scores from the condition-only analysis.**

| PC1 | PC2 | PC3 | PC4 | PC5 | sample | condition |
|---|---|---|---|---|---|---|
| -42.72776 | -63.6047 | 51.9539 | -31.3525 | 18.551937 | C01 | control |

| PC1 | PC2 | PC3 | PC4 | PC5 | sample | condition |
|---|---|---|---|---|---|---|
| 7 | 6 | 3 | 44 | 2 | | |
| 2.801694 | -57.95744 | 26.63570 | -20.194813 | 12.2506980 | C02 | control |
| -104.166323 | -48.42400 | -11.30855 | 8.058678 | 0.3251004 | C03 | control |
| 21.449902 | -44.69204 | -60.91020 | -21.664989 | -24.3528448 | C04 | control |
| -41.619235 | -66.86930 | 53.72664 | -27.620786 | 14.4232980 | C05 | control |
| 4.660833 | -60.58712 | 27.22625 | -15.827726 | 7.9569760 | C06 | control |
| -103.367441 | -51.44279 | -12.33202 | 14.887297 | -5.9350796 | C07 | control |
| 23.926012 | -46.68314 | -62.99652 | -16.503150 | -34.6290671 | C08 | control |
| 53.323039 | 47.70861 | -40.61522 | -25.728398 | 11.9473724 | RIN01 | Rin |
| 82.583695 | -20.18922 | 36.12928 | 35.785237 | -12.4318905 | RIN02 | Rin |

#### Table preview 11

**VST expression matrix.**

| | C01 | C02 | C03 | C04 | C05 | C06 | C07 | C08 | RIN01 | RIN02 | RIN03 | RIN04 | RIN05 | RIN06 | RIN07 | RIN08 | ROL01 | ROL02 | ROL03 | ROL04 | ROL05 |
|---|---|---|---|---|---|---|---|---|---|---|---|---|---|---|---|---|---|---|---|---|---|
| MD00G1000100 | 4.657673 | 4.711721 | 4.243505 | 4.539058 | 4.857698 | 4.243505 | 4.243505 | 4.243505 | 4.548392 | 4.614635 | 4.582140 | 4.646201 | 4.243505 | 4.243505 | 4.243505 | 4.243505 | 4.571851 | 4.690286 | 4.834939 | 5.028809 | 4.243505 |
| MD00G | 7.64 | 7.32 | 7.44 | 7.20 | 7.45 | 6.99 | 7.23 | 6.85 | 6.74 | 7.29 | 6.79 | 6.72 | 6.95 | 7.04 | 6.97 | 6.58 | 7.15 | 7.31 | 6.57 | 7.24 | 6.85 |

| | C001 | C002 | C003 | C004 | C005 | C006 | C007 | C008 |
|---|---|---|---|---|---|---|---|---|
| 1000200 | 4475 | 5830 | 2061 | 8868 | 3239 | 3955 | 8647 | 9344 |
| MD00G1000300 | 5.214388 | 5.222229 | 4.973268 | 5.717039 | 4.992973 | 5.068968 | 4.667805 | 5.254081 |
| MD00G1000500 | 5.186402 | 5.110214 | 5.781780 | 4.961299 | 5.105788 | 5.462108 | 5.814037 | 5.346265 |
| MD00G1000600 | 10.302765 | 11.237633 | 10.969268 | 11.561724 | 10.230810 | 11.241335 | 11.047824 | 11.518733 |
| MD00G1000700 | 7.422011 | 7.107782 | 7.115975 | 6.651441 | 7.393947 | 7.199225 | 7.487486 | 6.883838 |
| MD | 9. | 9. | 9. | 9. | 9. | 9. | 9. | 9. |

| | RIN001 | RIN002 | RIN003 | RIN004 | RIN005 | RIN006 | RIN007 | RIN008 | ROL001 | ROL002 | ROL003 | ROL004 | ROL005 |
|---|---|---|---|---|---|---|---|---|---|---|---|---|---|
| 1000200 | 9167 | 6764 | 6893 | 5084 | 7041 | 6612 | 3632 | 0809 | 7639 | 5017 | 6839 | 8008 | 4060 |
| MD00G1000300 | 5.041454 | 5.274065 | 5.293496 | 5.452998 | 5.495811 | 5.280445 | 4.243505 | 5.461531 | 4.868694 | 5.071298 | 4.243505 | 5.028809 | 4.933262 |
| MD00G1000500 | 5.145126 | 4.506294 | 5.388943 | 4.936577 | 4.884224 | 4.850662 | 5.232229 | 4.924302 | 5.355643 | 5.641673 | 5.646448 | 5.623553 | 5.284462 |
| MD00G1000600 | 11.228810 | 11.214600 | 10.427826 | 11.115723 | 11.413898 | 11.195004 | 10.399983 | 11.176842 | 10.585065 | 10.719555 | 10.856077 | 10.766374 | 10.595126 |
| MD00G1000700 | 6.955955 | 6.732740 | 7.190160 | 6.703993 | 7.106658 | 7.065450 | 7.098225 | 6.667785 | 6.989020 | 7.263598 | 7.589377 | 7.220002 | 6.854060 |
| MD | 9. | 9. | 9. | 9. | 9. | 9. | 9. | 9. | 9. | 9. | 9. | 9. | 9. |

| | C01 | C02 | C03 | C04 | C05 | C06 | C07 | C08 | RIN01 | RIN02 | RIN03 | RIN04 | RIN05 | RIN06 | RIN07 | RIN08 | ROL01 | ROL02 | ROL03 | ROL04 | ROL05 |
|---|---|---|---|---|---|---|---|---|---|---|---|---|---|---|---|---|---|---|---|---|---|
| 00G1000800 | 172244 | 340246 | 104515 | 729220 | 182307 | 444776 | 240734 | 920020 | 705247 | 647257 | 525932 | 750081 | 658922 | 609820 | 705338 | 723272 | 579251 | 205288 | 514628 | 295464 | 497564 |
| MD00G1000900 | 5.535807 | 5.366842 | 5.604778 | 5.361959 | 5.446071 | 5.068968 | 5.714332 | 5.150952 | 5.395665 | 5.304443 | 5.433273 | 5.214669 | 5.142469 | 5.500835 | 5.678941 | 5.271245 | 5.453509 | 5.594279 | 5.403199 | 5.573000 | 5.533500 |
| MD00G1001000 | 6.916976 | 6.720105 | 7.110132 | 6.808165 | 7.109574 | 7.199225 | 6.974932 | 6.426023 | 6.935601 | 6.669480 | 6.851810 | 7.007600 | 6.979432 | 6.797847 | 6.657225 | 6.951720 | 7.030362 | 7.263598 | 7.321260 | 7.176909 | 6.815831 |
| MD00G1001100 | 4.993785 | 4.766419 | 5.318000 | 5.069586 | 4.992973 | 4.243505 | 5.545470 | 4.890364 | 4.849941 | 4.933156 | 5.157656 | 5.041310 | 4.884224 | 4.850662 | 5.322588 | 5.339071 | 5.177034 | 5.437586 | 5.331757 | 5.404477 | 5.210105 |

Table preview 12

**rlog expression matrix.**

| | C001 | C002 | C003 | C004 | C005 | C006 | C007 | C008 | RIN001 | RIN002 | RIN003 | RIN004 | RIN005 | RIN006 | RIN007 | RIN008 | ROL001 | ROL002 | ROL003 | ROL004 | ROL005 |
|---|---|---|---|---|---|---|---|---|---|---|---|---|---|---|---|---|---|---|---|---|---|
| MD00G1000100 | 0.1734883 | 0.2372915 | -0.0959648 | 0.0533309 | 0.3633872 | -0.051234 | -0.0494144 | -0.0409955 | 0.0618681 | 0.1252508 | 0.092149 | 0.1574828 | -0.0420318 | -0.0477989 | -0.0434187 | -0.0566175 | 0.0811472 | 0.2128587 | 0.3425362 | 0.5851213 | -0.0552841 |
| MD00G1000200 | 7.1812916 | 6.8796259 | 6.9905867 | 6.7657466 | 6.9959423 | 6.555342 | 6.7926042 | 6.4235224 | 6.3068083 | 6.8512250 | 6.353941 | 6.2811891 | 6.5195873 | 6.6069454 | 6.5357599 | 6.1384364 | 6.7164950 | 6.8693626 | 6.1364948 | 6.8020924 | 6.4164531 |
| MD00G1000300 | 2.9874333 | 2.9991588 | 2.6302955 | 3.6489334 | 2.6892430 | 2.782855 | 2.3253587 | 3.0184135 | 2.7371447 | 3.0709010 | 3.101081 | 3.3140008 | 3.3274662 | 3.0565756 | 2.1438946 | 3.3000494 | 2.4579537 | 2.7758161 | 2.1097042 | 2.7290920 | 2.6078310 |
| MD00 | 3.15 | 3.04 | 3.98 | 2.83 | 3.06 | 3.53 | 3.97 | 3.37 | 3.10 | 2.20 | 3.45 | 2.79 | 2.80 | 2.75 | 3.23 | 2.82 | 3.41 | 3.80 | 3.76 | 3.74 | 3.29 |

| | C01 | C02 | C03 | C04 | C05 | C06 | C07 | C08 | RIN01 | RIN02 | RIN03 | RIN04 | RIN05 | RIN06 | RIN07 | RIN08 | ROL01 | ROL02 | ROL03 | ROL04 | ROL05 |
|---|---|---|---|---|---|---|---|---|---|---|---|---|---|---|---|---|---|---|---|---|---|
| G1000500 | 97872 | 36186 | 06120 | 31756 | 71552 | 3031 | 66998 | 69883 | 34462 | 92542 | 6174 | 29925 | 50013 | 38088 | 07166 | 37876 | 17425 | 40016 | 94401 | 61486 | 99959 |
| MD00G1000600 | 10.4023513 | 11.1535732 | 10.9366101 | 11.4169030 | 10.3460776 | 11.156385 | 10.9999317 | 11.3815601 | 11.1463649 | 11.1348903 | 10.502091 | 11.0548568 | 11.2963090 | 11.1188572 | 10.4806026 | 11.1042014 | 10.6277717 | 10.7356658 | 10.8454920 | 10.7734129 | 10.6361919 |
| MD00G1000700 | 6.9754542 | 6.6714458 | 6.6791956 | 6.2092832 | 6.9440789 | 6.758964 | 7.0324896 | 6.4520631 | 6.5206198 | 6.2924457 | 6.751969 | 6.2635129 | 6.6693976 | 6.6294857 | 6.6612649 | 6.2322754 | 6.5528140 | 6.8235409 | 7.1278731 | 6.7792334 | 6.4207287 |
| MD00G100 | 9.152551 | 9.291729 | 9.096477 | 9.613804 | 9.161881 | 9.378456 | 9.210015 | 9.771161 | 9.593937 | 9.545993 | 9.445542 | 9.631102 | 9.555289 | 9.514776 | 9.593652 | 9.608633 | 9.489728 | 9.179877 | 9.436152 | 9.255083 | 9.422056 |

| | C001 | C002 | C003 | C004 | C005 | C006 | C007 | C008 | RIN01 | RIN02 | RIN03 | RIN04 | RIN05 | RIN06 | RIN07 | RIN08 | ROL01 | ROL02 | ROL03 | ROL04 | ROL05 |
|---|---|---|---|---|---|---|---|---|---|---|---|---|---|---|---|---|---|---|---|---|---|
| 0800 | 7 | 1 | 0 | 4 | 9 | | 1 | 5 | 3 | 7 | | 4 | 7 | 6 | 0 | 6 | 8 | 9 | 8 | 1 | 8 |
| MD00G1000900 | 3.8891479 | 3.6554553 | 3.9806864 | 3.6506109 | 3.7627106 | 3.283133 | 4.0977203 | 3.3980830 | 3.6970945 | 3.5681249 | 3.748810 | 3.4426559 | 3.3865287 | 3.8318290 | 4.0503935 | 3.5365971 | 3.7777041 | 3.9683929 | 3.7084914 | 3.9257363 | 3.8752487 |
| MD00G1001000 | 6.4321015 | 6.2294851 | 6.6252076 | 6.3216640 | 6.6214679 | 6.708751 | 6.4898458 | 5.9354457 | 6.4509594 | 6.1771556 | 6.365742 | 6.5230959 | 6.4941026 | 6.3136629 | 6.1722295 | 6.4669829 | 6.5464180 | 6.7757086 | 6.8257372 | 6.6877305 | 6.3311528 |
| MD00G1001100 | 2.6224800 | 2.2853790 | 3.0932091 | 2.7367542 | 2.6463284 | 2.063646 | 3.3546991 | 2.5306151 | 2.4304802 | 2.5364945 | 2.864565 | 2.6958594 | 2.5224168 | 2.4766076 | 3.0630262 | 3.0956612 | 2.8954751 | 3.2616561 | 3.0810431 | 3.1818992 | 2.9247363 |

#### Table preview 13

**Mean-SD summary by transformation.**

| transformation | mean | sd |
|---|---:|---:|
| log2 raw counts + 1 | 0.9947903 | 0.9474391 |
| log2 raw counts + 1 | 6.6408970 | 1.0471173 |
| log2 raw counts + 1 | 2.9288680 | 1.3709868 |
| log2 raw counts + 1 | 3.3039948 | 1.2586028 |
| log2 raw counts + 1 | 10.9192264 | 0.8666686 |
| log2 raw counts + 1 | 6.6578352 | 0.8655160 |
| log2 raw counts + 1 | 9.4317689 | 0.8131938 |
| log2 raw counts + 1 | 3.8306292 | 0.9570156 |
| log2 raw counts + 1 | 6.4721698 | 0.9092603 |
| log2 raw counts + 1 | 2.9082861 | 1.1575396 |

#### Table preview 14

**Pearson correlation matrix based on VST values.**

| | C001 | C002 | C003 | C004 | C005 | C006 | C007 | C008 | RIN001 | RIN002 | RIN003 | RIN004 | RIN005 | RIN006 | RIN007 | RIN008 | ROL001 | ROL002 | ROL003 | ROL004 | ROL005 |
|---|---|---|---|---|---|---|---|---|---|---|---|---|---|---|---|---|---|---|---|---|---|
| C001 | 1.0000000 | 0.9859258 | 0.9665673 | 0.9495962 | 0.9932205 | 0.9795358 | 0.9610784 | 0.9437210 | 0.9168634 | 0.9410046 | 0.9113075 | 0.9151802 | 0.9346891 | 0.9354720 | 0.9044681 | 0.9096098 | 0.9421246 | 0.9348092 | 0.9285482 | 0.9287521 | 0.9365982 |

| | C01 | C02 | C03 | C04 | C05 | C06 | C07 | C08 | RIN01 | RIN02 | RIN03 | RIN04 | RIN05 | RIN06 | RIN07 | RIN08 | ROL01 | ROL02 | ROL03 | ROL04 | ROL05 |
|---|---|---|---|---|---|---|---|---|---|---|---|---|---|---|---|---|---|---|---|---|---|
| C02 | 0.9859258 | 1.0000000 | 0.9558671 | 0.9715143 | 0.9830528 | 0.9937882 | 0.9527449 | 0.9675979 | 0.9476911 | 0.9669655 | 0.9309231 | 0.9504492 | 0.9629185 | 0.9635154 | 0.9261575 | 0.9469666 | 0.9508380 | 0.9292563 | 0.9253734 | 0.9259271 | 0.9473804 |
| C03 | 0.9665673 | 0.9558671 | 1.0000000 | 0.9437425 | 0.9624850 | 0.9502757 | 0.9929675 | 0.9385694 | 0.9014423 | 0.9005151 | 0.8728743 | 0.8963642 | 0.8957712 | 0.8955611 | 0.8675519 | 0.8918373 | 0.9404421 | 0.9613575 | 0.9559219 | 0.9567215 | 0.9359287 |
| C04 | 0.9495962 | 0.9715143 | 0.9437425 | 1.0000000 | 0.9461828 | 0.9663911 | 0.9400202 | 0.9927288 | 0.9642017 | 0.9546524 | 0.9177571 | 0.9708738 | 0.9498351 | 0.9504197 | 0.9124065 | 0.9665595 | 0.9546693 | 0.9232571 | 0.9186315 | 0.9191123 | 0.9506945 |
| C05 | 0.9932205 | 0.9830528 | 0.9624850 | 0.9461828 | 1.0000000 | 0.9816587 | 0.9626218 | 0.9453597 | 0.9140847 | 0.9378658 | 0.9076527 | 0.9127502 | 0.9368053 | 0.9369244 | 0.9061771 | 0.9111122 | 0.9388516 | 0.9306717 | 0.9298593 | 0.9299796 | 0.9376731 |
| C06 | 0.9 | 0.9 | 0.9 | 0.9 | 0.9 | 1.0 | 0.9 | 0.9 | 0.9 | 0.9 | 0.9 | 0.9 | 0.9 | 0.9 | 0.9 | 0.9 | 0.9 | 0.9 | 0.9 | 0.9 | 0.9 |

| | C01 | C02 | C03 | C04 | C05 | C06 | C07 | C08 | RIN01 | RIN02 | RIN03 | RIN04 | RIN05 | RIN06 | RIN07 | RIN08 | ROL01 | ROL02 | ROL03 | ROL04 | ROL05 |
|---|---|---|---|---|---|---|---|---|---|---|---|---|---|---|---|---|---|---|---|---|---|
| | 795358 | 937882 | 502757 | 663911 | 816587 | 000000 | 524763 | 676301 | 434657 | 621302 | 257288 | 466039 | 632138 | 632433 | 263567 | 468845 | 459743 | 236107 | 249789 | 254363 | 466627 |
| C07 | 0.9610784 | 0.9527449 | 0.9929675 | 0.9400202 | 0.9626218 | 0.9524763 | 1.0000000 | 0.9403099 | 0.8982742 | 0.8971214 | 0.8685497 | 0.8940895 | 0.8979088 | 0.8972103 | 0.8688556 | 0.8937768 | 0.9364842 | 0.9561913 | 0.9565868 | 0.9573920 | 0.9366221 |
| C08 | 0.9437210 | 0.9675979 | 0.9385694 | 0.9927288 | 0.9453597 | 0.9676301 | 0.9403099 | 1.0000000 | 0.9608281 | 0.9506762 | 0.9135359 | 0.9675253 | 0.9513534 | 0.9509729 | 0.9136460 | 0.9672294 | 0.9505156 | 0.9182574 | 0.9191030 | 0.9194546 | 0.9508192 |
| RIN01 | 0.9168634 | 0.9476911 | 0.9014423 | 0.9642017 | 0.9140847 | 0.9434657 | 0.8982742 | 0.9608281 | 1.0000000 | 0.9555014 | 0.9669404 | 0.9758654 | 0.9516496 | 0.9519129 | 0.9627022 | 0.9723688 | 0.9687984 | 0.9251107 | 0.9216944 | 0.9222609 | 0.9656882 |
| RIN02 | 0.9410 | 0.9669 | 0.9005 | 0.9546 | 0.9378 | 0.9621 | 0.8971 | 0.9506 | 0.9555 | 1.0000 | 0.9525 | 0.9654 | 0.9925 | 0.9934 | 0.9472 | 0.9615 | 0.9355 | 0.8985 | 0.8941 | 0.8948 | 0.9319 |

| | C01 | C02 | C03 | C04 | C05 | C06 | C07 | C08 | RIN01 | RIN02 | RIN03 | RIN04 | RIN05 | RIN06 | RIN07 | RIN08 | ROL01 | ROL02 | ROL03 | ROL04 | ROL05 |
|---|---|---|---|---|---|---|---|---|---|---|---|---|---|---|---|---|---|---|---|---|---|
| | 046 | 655 | 151 | 524 | 658 | 302 | 214 | 762 | 014 | 000 | 312 | 437 | 009 | 768 | 445 | 038 | 820 | 645 | 718 | 729 | 024 |

#### Table preview 15

**Pearson correlation matrix based on rlog values.**

| | C01 | C02 | C03 | C04 | C05 | C06 | C07 | C08 | RIN01 | RIN02 | RIN03 | RIN04 | RIN05 | RIN06 | RIN07 | RIN08 | ROL01 | ROL02 | ROL03 | ROL04 | ROL05 |
|---|---|---|---|---|---|---|---|---|---|---|---|---|---|---|---|---|---|---|---|---|---|
| C01 | 1.0000000 | 0.9941266 | 0.9862006 | 0.9797401 | 0.9969541 | 0.9915719 | 0.9846969 | 0.9777326 | 0.9639465 | 0.9745169 | 0.9610287 | 0.9641001 | 0.9734130 | 0.9735623 | 0.9593781 | 0.9631027 | 0.9743148 | 0.9713816 | 0.9699364 | 0.9699881 | 0.9727105 |
| C02 | 0.9941266 | 1.0000000 | 0.9811475 | 0.9883292 | 0.9931630 | 0.9971856 | 0.9808775 | 0.9869082 | 0.9761335 | 0.9851383 | 0.9690754 | 0.9783522 | 0.9844098 | 0.9845414 | 0.9679967 | 0.9777854 | 0.9773526 | 0.9684295 | 0.9681214 | 0.9682575 | 0.9762249 |
| C03 | 0.9862006 | 0.9811475 | 1.0000000 | 0.9753450 | 0.9845838 | 0.9791088 | 0.9968880 | 0.9737718 | 0.9559023 | 0.9569144 | 0.9445403 | 0.9544370 | 0.9570992 | 0.9568097 | 0.9440073 | 0.9542487 | 0.9724664 | 0.9818588 | 0.9801343 | 0.9804870 | 0.9711479 |

| | C01 | C02 | C03 | C04 | C05 | C06 | C07 | C08 | RIN01 | RIN02 | RIN03 | RIN04 | RIN05 | RIN06 | RIN07 | RIN08 | ROL01 | ROL02 | ROL03 | ROL04 | ROL05 |
|---|---|---|---|---|---|---|---|---|---|---|---|---|---|---|---|---|---|---|---|---|---|
| C04 | 0.9797401 | 0.9883292 | 0.9753450 | 1.0000000 | 0.9791811 | 0.9867443 | 0.9752101 | 0.9967806 | 0.9837151 | 0.9815874 | 0.9654924 | 0.9872886 | 0.9805364 | 0.9806702 | 0.9643254 | 0.9861512 | 0.9795015 | 0.9661973 | 0.9656760 | 0.9657987 | 0.9785949 |
| C05 | 0.9969541 | 0.9931630 | 0.9845838 | 0.9791811 | 1.0000000 | 0.9928458 | 0.9855308 | 0.9793291 | 0.9650926 | 0.9743928 | 0.9620587 | 0.9646189 | 0.9754339 | 0.9753173 | 0.9626099 | 0.9651757 | 0.9749617 | 0.9713864 | 0.9721487 | 0.9720975 | 0.9751224 |
| C06 | 0.9915719 | 0.9971856 | 0.9791088 | 0.9867443 | 0.9928458 | 1.0000000 | 0.9811451 | 0.9875479 | 0.9761019 | 0.9838875 | 0.9689196 | 0.9777189 | 0.9853098 | 0.9852036 | 0.9700523 | 0.9786711 | 0.9770749 | 0.9678424 | 0.9696295 | 0.9697199 | 0.9780352 |
| C07 | 0.9846969 | 0.9808775 | 0.9968880 | 0.9752101 | 0.9855308 | 0.9811451 | 1.0000000 | 0.9759006 | 0.9575273 | 0.9576169 | 0.9461711 | 0.9556697 | 0.9599637 | 0.9594635 | 0.9478658 | 0.9571144 | 0.9731851 | 0.9814715 | 0.9821394 | 0.9824488 | 0.9737212 |
| C08 | 0.9 | 0.9 | 0.9 | 0.9 | 0.9 | 0.9 | 0.9 | 1.0 | 0.9 | 0.9 | 0.9 | 0.9 | 0.9 | 0.9 | 0.9 | 0.9 | 0.9 | 0.9 | 0.9 | 0.9 | 0.9 |

| | C001 | C002 | C003 | C004 | C005 | C006 | C007 | C008 | RIN01 | RIN02 | RIN03 | RIN04 | RIN05 | RIN06 | RIN07 | RIN08 | ROL01 | ROL02 | ROL03 | ROL04 | ROL05 |
|---|---|---|---|---|---|---|---|---|---|---|---|---|---|---|---|---|---|---|---|---|---|
| | 777326 | 869082 | 737718 | 967806 | 793291 | 875479 | 759006 | 000000 | 831468 | 803830 | 649750 | 862360 | 816255 | 813407 | 660425 | 867357 | 788350 | 655243 | 671442 | 671728 | 796526 |
| RIN01 | 0.9639465 | 0.9761335 | 0.9559023 | 0.9837151 | 0.9650926 | 0.9761019 | 0.9575273 | 0.9831468 | 1.0000000 | 0.9805937 | 0.9870371 | 0.9888647 | 0.9800117 | 0.9800164 | 0.9857454 | 0.9878488 | 0.9866483 | 0.9682774 | 0.9679585 | 0.9681139 | 0.9857966 |
| RIN02 | 0.9745169 | 0.9851383 | 0.9569144 | 0.9815874 | 0.9743928 | 0.9838875 | 0.9576169 | 0.9803830 | 0.9805937 | 1.0000000 | 0.9783607 | 0.9859203 | 0.9965965 | 0.9970279 | 0.9768702 | 0.9846485 | 0.9713683 | 0.9553324 | 0.9551891 | 0.9553463 | 0.9708367 |

#### Table preview 16

**Input table for the custom MA plot.**

| gene_id | gene_name | baseMean | log2FoldChange | lfcSE | stat | pvalue | padj | Description | significant | baseMean_for_plot |
|---|---|---|---|---|---|---|---|---|---|---|
| MD07G1003800 | MD07G1003800 | 266.79219 | 4.318660 | 0.2682259 | 16.10083 | 0 | 0 | O-fucosyltransferase family protein | TRUE | 266.7919 |

| gene_id | gene_name | baseMean | log2FoldChange | lfcSE | stat | pvalue | padj | Description | significant | baseMean_for_plot |
|---|---|---|---|---|---|---|---|---|---|---|
| MD15G1262600 | MD15G1262600 | 2196.0749 | -1.577524 | 0.1014381 | -15.55160 | 0 | 0 | MD15G1262600 | TRUE | 2196.0749 |
| MD13G1087700 | MD13G1087700 | 369.9144 | -2.001719 | 0.1384273 | -14.46043 | 0 | 0 | glyoxal oxidase-related protein | TRUE | 369.9144 |
| MD15G1010400 | MD15G1010400 | 970.5317 | -1.438880 | 0.1059982 | -13.57457 | 0 | 0 | MD15G1010400 | TRUE | 970.5317 |
| MD15G1010500 | MD15G1010500 | 385.4605 | -1.605449 | 0.1190456 | -13.48600 | 0 | 0 | P-glycoprotein 13 | TRUE | 385.4605 |
| MD09G1148500 | MD09G1148500 | 140.6160 | -2.503914 | 0.1902764 | -13.15935 | 0 | 0 | GDSL-like Lipase/Acylhydrolase superfamily protein | TRUE | 140.6160 |
| MD11G1062600 | MD11G1062600 | 258.5170 | -2.849692 | 0.2241246 | -12.71477 | 0 | 0 | UDP-Glycosyltransferase superfamily protein | TRUE | 258.5170 |
| MD16G1107100 | MD16G1107100 | 1696.8125 | -1.050777 | 0.0826495 | -12.71366 | 0 | 0 | beta-ketoacyl reductase 1 | TRUE | 1696.8125 |
| MD10G116 | MD10G116 | 286.27 | -5.954488 | 0.4746 | -12.54 | 0 | 0 | Pectin lyase-like superfamily protein | TRUE | 286.2707 |

| gene_id | gene_name | baseMean | log2FoldChange | lfcSE | stat | pvalue | padj | Description | significant | baseMean_for_plot |
|---|---|---|---|---|---|---|---|---|---|---|
| 9100 | 9100 | 07 | | 279 | 559 | | | | | |
| MD12G1082400 | MD12G1082400 | 779.6653 | -2.015843 | 0.1634273 | -12.33480 | 0 | 0 | 6-phosphogluconate dehydrogenase family protein | TRUE | 779.6653 |

#### Table preview 17

**Input table for the volcano plot.**

| gene_id | gene_name | baseMean | log2FoldChange | lfcSE | stat | pvalue | padj | Description | direction | neg_log10_padj |
|---|---|---|---|---|---|---|---|---|---|---|
| MD07G1003800 | MD07G1003800 | 266.7919 | 4.318660 | 0.2682259 | 16.10083 | 0 | 0 | O-fucosyltransferase family protein | Rin-high | 53.03121 |
| MD15G1262600 | MD15G1262600 | 2196.0749 | -1.577524 | 0.1014381 | -15.55160 | 0 | 0 | MD15G1262600 | Rola-high | 49.54229 |
| MD13G1087700 | MD13G1087700 | 369.9144 | -2.001719 | 0.1384273 | -14.46043 | 0 | 0 | glyoxal oxidase-related protein | Rola-high | 42.57591 |
| MD15G1010400 | MD15G1010400 | 970.5317 | -1.438880 | 0.1059982 | -13.57457 | 0 | 0 | MD15G1010400 | Rola-high | 37.28080 |
| MD | MD | 38 | -1.60 | 0. | -1 | 0 | 0 | P-glycoprotein 13 | Ro | 36.8 |

| gene_id | gene_name | baseMean | log2FoldChange | lfcSE | stat | pvalue | padj | Description | direction | neg_log10_padj |
|---|---|---|---|---|---|---|---|---|---|---|
| 15G1010500 | 15G1010500 | 5.4605 | 5449 | 1190456 | 3.48600 | | | | la-high | 5444 |
| MD09G1148500 | MD09G1148500 | 140.6160 | -2.503914 | 0.1902764 | -13.15935 | 0 | 0 | GDSL-like Lipase/Acylhydrolase superfamily protein | Rola-high | 35.03309 |
| MD11G1062600 | MD11G1062600 | 258.5170 | -2.849692 | 0.2241246 | -12.71477 | 0 | 0 | UDP-Glycosyltransferase superfamily protein | Rola-high | 32.63926 |
| MD16G1107100 | MD16G1107100 | 1696.8125 | -1.050777 | 0.0826495 | -12.71366 | 0 | 0 | beta-ketoacyl reductase 1 | Rola-high | 32.63926 |
| MD10G1169100 | MD10G1169100 | 286.2707 | -5.954488 | 0.4746279 | -12.54559 | 0 | 0 | Pectin lyase-like superfamily protein | Rola-high | 31.76283 |
| MD12G1082400 | MD12G1082400 | 779.6653 | -2.015843 | 0.1634273 | -12.33480 | 0 | 0 | 6-phosphogluconate dehydrogenase family protein | Rola-high | 30.66246 |

#### Table preview 18

**Candidate Rin-high genes using padj < 0.05, |LFC| >= 1, baseMean >= 50.**

| gene_id | gene_name | baseMean | log2FoldChange | lfcSE | stat | pvalue | padj | Description |
|---|---|---|---|---|---|---|---|---|
| MD07G1003800 | MD07G1003800 | 266.7919 | 4.318660 | 0.2682259 | 16.100828 | 0 | 0 | O-fucosyltransferase family protein |
| MD07G1236800 | MD07G1236800 | 663.6725 | 3.979268 | 0.4010596 | 9.921888 | 0 | 0 | Leucine-rich repeat protein kinase family protein |
| MD07G1236500 | MD07G1236500 | 1385.3689 | 6.061322 | 0.6362824 | 9.526151 | 0 | 0 | Leucine-rich repeat protein kinase family protein |
| MD06G1211700 | MD06G1211700 | 1501.0921 | 2.314120 | 0.2442776 | 9.473318 | 0 | 0 | Duplicated homeodomain-like superfamily protein |
| MD04G1183000 | MD04G1183000 | 188.3291 | 4.901930 | 0.5270531 | 9.300638 | 0 | 0 | Leucine-rich repeat protein kinase family protein |
| MD13G1101300 | MD13G1101300 | 1785.6485 | 4.125229 | 0.4437685 | 9.295903 | 0 | 0 | octicosapeptide/Phox/Bem1p (PB1) domain-containing protein |
| MD16G1091200 | MD16G1091200 | 1546.4820 | 6.482363 | 0.7225875 | 8.971042 | 0 | 0 | xyloglucan:xyloglucosyl transferase 33 |
| MD13G1100800 | MD13G1100800 | 1563.8671 | 1.159618 | 0.1310225 | 8.850526 | 0 | 0 | Phototropic-responsive NPH3 family protein |
| MD17G1234900 | MD17G1234900 | 1671.7365 | 1.117557 | 0.1282586 | 8.713313 | 0 | 0 | Tyrosine transaminase family protein |
| MD14G1 | MD14G1 | 4059. | 1.796706 | 0.209 | 8.581 | 0 | 0 | B-box type zinc finger protein with CCT domain |

| gene_id | gene_name | baseMean | log2FoldChange | lfcSE | stat | pvalue | padj | Description |
|---|---|---|---|---|---|---|---|---|
| 226400 | 226400 | 2290 | | 3725 | 382 | | | |

#### Table preview 19

**Candidate Rola-high genes using padj < 0.05, |LFC| >= 1, baseMean >= 50.**

| gene_id | gene_name | baseMean | log2FoldChange | lfcSE | stat | pvalue | padj | Description |
|---|---|---|---|---|---|---|---|---|
| MD15G1262600 | MD15G1262600 | 2196.0749 | -1.577524 | 0.1014381 | -15.55160 | 0 | 0 | MD15G1262600 |
| MD13G1087700 | MD13G1087700 | 369.9144 | -2.001719 | 0.1384273 | -14.46043 | 0 | 0 | glyoxal oxidase-related protein |
| MD15G1010400 | MD15G1010400 | 970.5317 | -1.438880 | 0.1059982 | -13.57457 | 0 | 0 | MD15G1010400 |
| MD15G1010500 | MD15G1010500 | 385.4605 | -1.605449 | 0.1190456 | -13.48600 | 0 | 0 | P-glycoprotein 13 |
| MD09G1148500 | MD09G1148500 | 140.6160 | -2.503914 | 0.1902764 | -13.15935 | 0 | 0 | GDSL-like Lipase/Acylhydrolase superfamily protein |
| MD11G1062600 | MD11G1062600 | 258.5170 | -2.849692 | 0.2241246 | -12.71477 | 0 | 0 | UDP-Glycosyltransferase superfamily protein |
| MD16G11 | MD16G11 | 1696.8 | -1.050777 | 0.0826 | -12.713 | 0 | 0 | beta-ketoacyl reductase 1 |

| gene_id | gene_name | baseMean | log2FoldChange | lfcSE | stat | pvalue | padj | Description |
|---|---|---|---|---|---|---|---|---|
| 07100 | 07100 | 125 | | 495 | 66 | | | |
| MD10G1169100 | MD10G1169100 | 286.2707 | -5.954488 | 0.4746279 | -12.54559 | 0 | 0 | Pectin lyase-like superfamily protein |
| MD12G1082400 | MD12G1082400 | 779.6653 | -2.015843 | 0.1634273 | -12.33480 | 0 | 0 | 6-phosphogluconate dehydrogenase family protein |
| MD15G1386800 | MD15G1386800 | 137.6311 | -2.089000 | 0.1735969 | -12.03362 | 0 | 0 | HXXXD-type acyl-transferase family protein |

#### Table preview 20

**Rin vs Rola DESeq2 results using lfcThreshold = 1.**

| gene_id | gene_name | baseMean | log2FoldChange | lfcSE | stat | pvalue | padj | Description |
|---|---|---|---|---|---|---|---|---|
| MD07G1003800 | MD07G1003800 | 266.79193 | 4.318660 | 0.2682259 | 16.100828 | 0 | 0 | O-fucosyltransferase family protein |
| MD10G1169100 | MD10G1169100 | 286.27071 | -5.954488 | 0.4746279 | -12.545591 | 0 | 0 | Pectin lyase-like superfamily protein |
| MD15G1400200 | MD15G1400200 | 35.38328 | -24.534513 | 2.3086322 | -10.627294 | 0 | 0 | GDSL-like Lipase/Acylhydrolase superfamily protein |
| MD01G121080 | MD01G121080 | 38.79491 | -24.127114 | 2.3623828 | -10.213042 | 0 | 0 | ABC-2 type transporter family protein |

| gene_id | gene_name | baseMean | log2FoldChange | lfcSE | stat | pvalue | padj | Description |
|---|---|---:|---:|---:|---:|---:|---:|---|
| 0 | 0 | | | | | | | |
| MD12G1067600 | MD12G1067600 | 64.51832 | -25.197381 | 2.5683866 | -9.810587 | 0 | 0 | Integrase-type DNA-binding superfamily protein |
| MD02G1080400 | MD02G1080400 | 59.01762 | -4.922775 | 0.4295451 | -11.460437 | 0 | 0 | Protein of unknown function |
| MD17G1103100 | MD17G1103100 | 457.12378 | -4.807139 | 0.4383697 | -10.965947 | 0 | 0 | glycosyl hydrolase 9B8 |
| MD04G1030500 | MD04G1030500 | 1849.24772 | -8.376991 | 0.8901764 | -9.410485 | 0 | 0 | MD04G1030500 |
| MD11G1062600 | MD11G1062600 | 258.51698 | -2.849692 | 0.2241246 | -12.714768 | 0 | 0 | UDP-Glycosyltransferase superfamily protein |
| MD09G1084600 | MD09G1084600 | 46.42252 | -5.246255 | 0.5188396 | -10.111517 | 0 | 0 | allene oxide cyclase 4 |

#### Table preview 21

**Rin vs Rola lfcThreshold = 1 results sorted by log2FC.**

| gene_id | gene_name | baseMean | log2FoldChange | lfcSE | stat | pvalue | padj | Description |
|---|---|---:|---:|---:|---:|---:|---:|---|
| MD02G1138900 | MD02G1138900 | 114.21657 | 10.207431 | 1.8901696 | 5.400273 | 0.0000006 | 0.0000593 | Peroxidase superfamily protein |

| gene_id | gene_name | baseMean | log2FoldChange | lfcSE | stat | pvalue | padj | Description |
|---|---|---|---|---|---|---|---|---|
| MD11G1315600 | MD11G1315600 | 1955.11524 | 9.835361 | 1.2393802 | 7.935709 | 0.0000000 | 0.0000000 | Adenine nucleotide alpha hydrolases-like superfamily protein |
| MD01G1198900 | MD01G1198900 | 71.94180 | 9.214582 | 1.7731461 | 5.196741 | 0.0000018 | 0.0001540 | MD01G1198900 |
| MD07G1266700 | MD07G1266700 | 402.56427 | 9.211046 | 1.3338630 | 6.905542 | 0.0000000 | 0.0000001 | MD07G1266700 |
| MD13G1003800 | MD13G1003800 | 714.71655 | 7.840833 | 1.2676303 | 6.185425 | 0.0000000 | 0.0000062 | DNAJ heat shock N-terminal domain-containing protein |
| MD07G1299100 | MD07G1299100 | 3094.63193 | 7.780463 | 1.0828739 | 7.185013 | 0.0000000 | 0.0000001 | pseudo-response regulator 5 |
| MD12G1178600 | MD12G1178600 | 319.70567 | 7.756025 | 0.9828706 | 7.891196 | 0.0000000 | 0.0000000 | Mitochondrial substrate carrier family protein |
| MD00G1033200 | MD00G1033200 | 14.40147 | 7.714226 | 1.3701021 | 5.630402 | 0.0000005 | 0.0000524 | MD00G1033200 |

| gene_id | gene_name | baseMean | log2FoldChange | lfcSE | stat | pvalue | padj | Description |
|---|---|---|---|---|---|---|---|---|
| MD06G1089700 | MD06G1089700 | 31.02944 | 7.560113 | 1.2692494 | 5.956365 | 0.0000001 | 0.0000171 | myb domain protein 103 |
| MD10G1255800 | MD10G1255800 | 20.48919 | 7.535694 | 1.9613775 | 3.842042 | 0.0004376 | 0.0117134 | MD10G1255800 |

#### Table preview 22

**Rin-high candidates in Rin vs Rola using lfcThreshold = 1 and padj < 0.01.**

| gene_id | gene_name | baseMean | log2FoldChange | lfcSE | stat | pvalue | padj | Description |
|---|---|---|---|---|---|---|---|---|
| MD02G1138900 | MD02G1138900 | 114.21657 | 10.207431 | 1.8901696 | 5.400273 | 6.0e-07 | 5.93e-05 | Peroxidase superfamily protein |
| MD11G1315600 | MD11G1315600 | 1955.11524 | 9.835361 | 1.2393802 | 7.935709 | 0.0e+00 | 0.00e+00 | Adenine nucleotide alpha hydrolases-like superfamily protein |
| MD01G1198900 | MD01G1198900 | 71.94180 | 9.214582 | 1.7731461 | 5.196741 | 1.8e-06 | 1.54e-04 | MD01G1198900 |
| MD07G1266700 | MD07G1266700 | 402.56427 | 9.211046 | 1.3338630 | 6.905542 | 0.0e+00 | 1.00e-07 | MD07G1266700 |

| gene_id | gene_name | baseMean | log2FoldChange | lfcSE | stat | pvalue | padj | Description |
|---|---|---|---|---|---|---|---|---|
| MD13G1003800 | MD13G1003800 | 714.71655 | 7.840833 | 1.2676303 | 6.185425 | 00.0e+00 | 6.20e-06 | DNAJ heat shock N-terminal domain-containing protein |
| MD07G1299100 | MD07G1299100 | 3094.63193 | 7.780463 | 1.0828739 | 7.185013 | 0.0e+00 | 1.00e-07 | pseudo-response regulator 5 |
| MD12G1178600 | MD12G1178600 | 319.70567 | 7.756025 | 0.9828706 | 7.891196 | 0.0e+00 | 0.00e+00 | Mitochondrial substrate carrier family protein |
| MD00G1033200 | MD00G1033200 | 14.40147 | 7.714226 | 1.3701021 | 5.630402 | 5.0e-07 | 5.24e-05 | MD00G1033200 |
| MD06G1089700 | MD06G1089700 | 31.02944 | 7.560113 | 1.2692494 | 5.956365 | 1.0e-07 | 1.71e-05 | myb domain protein 103 |
| MD15G1082300 | MD15G1082300 | 419.76829 | 7.378332 | 1.1918653 | 6.190576 | 0.0e+00 | 7.50e-06 | cold regulated gene 27 |

#### Table preview 23

**Rola-high candidates in Rin vs Rola using lfcThreshold = 1 and padj < 0.01.**

| gene_id | gene_name | baseMean | log2FoldChange | lfcSE | stat | pvalue | padj | Description |
|---|---|---|---|---|---|---|---|---|
| MD12G1067600 | MD12G1067600 | 64.51832 | -25.197381 | 2.568387 | -9.810587 | 0e+00 | 0.000e+00 | Integrase-type DNA-binding superfamily protein |
| MD15G1400200 | MD15G1400200 | 35.38328 | -24.534513 | 2.308632 | -10.627294 | 0e+00 | 0.000e+00 | GDSL-like Lipase/Acylhydrolase superfamily protein |
| MD01G1210800 | MD01G1210800 | 38.79491 | -24.127114 | 2.362383 | -10.213042 | 0e+00 | 0.000e+00 | ABC-2 type transporter family protein |
| MD12G1252200 | MD12G1252200 | 8353.64251 | -13.226343 | 1.637601 | -8.076660 | 0e+00 | 0.000e+00 | GDSL-like Lipase/Acylhydrolase superfamily protein |
| MD05G1159700 | MD05G1159700 | 480.06173 | -12.555504 | 1.665455 | -7.538784 | 0e+00 | 0.000e+00 | Bifunctional inhibitor/lipid-transfer protein/seed storage 2S albumin superfamily protein |
| MD04G1 | MD04G1 | 3084.3 | -11.553662 | 1.32 | -8.7357 | 0e+ | 0.00 | GDSL-like Lipase/Acylhydrolase superfamily protein |

| gene_id | gene_name | baseMean | log2FoldChange | lfcSE | stat | pvalue | padj | Description |
|---|---|---|---|---|---|---|---|---|
| 234300 | 234300 | 9032 | | 2566 | 89 | 00 | e+00 | |
| MD00G1174400 | MD00G1174400 | 423.66967 | -11.545330 | 1.543794 | -7.478541 | 0e+00 | 0.000e+00 | fatty acid desaturase 5 |
| MD14G1081900 | MD14G1081900 | 384.01231 | -11.073301 | 1.735217 | -6.381510 | 0e+00 | 9.000e-07 | Gibberellin-regulated family protein |
| MD17G1140600 | MD17G1140600 | 262.49148 | -11.059716 | 1.952257 | -5.665094 | 1e-07 | 1.83e-05 | protodermal factor 1 |
| MD09G1154000 | MD09G1154000 | 162.97381 | -9.976635 | 1.743651 | -5.721693 | 1e-07 | 1.86e-05 | protodermal factor 1 |

#### Table preview 24

**Rin vs Rola candidates sorted by absolute log2FC.**

| gene_id | gene_name | baseMean | log2FoldChange | lfcSE | stat | pvalue | padj | direction | Description |
|---|---|---|---|---|---|---|---|---|---|
| MD12G1067600 | MD12G1067600 | 64.51832 | -25.19738 | 2.568387 | -9.810587 | 0e+00 | 0.00e+00 | Rola-high | Integrase-type DNA-binding superfamily protein |
| MD15G1400200 | MD15G1400200 | 35.38328 | -24.53451 | 2.308632 | -10.627294 | 0e+00 | 0.00e+00 | Rola-high | GDSL-like Lipase/Acylhydrolase superfamily protein |
| MD01G1210800 | MD01G1210800 | 38.79491 | -24.12711 | 2.362383 | -10.213042 | 0e+00 | 0.00e+00 | Rola-high | ABC-2 type transporter family protein |
| MD12G1252200 | MD12G1252200 | 8353.64251 | -13.22634 | 1.637601 | -8.076660 | 0e+00 | 0.00e+00 | Rola-high | GDSL-like Lipase/Acylhydrolase superfamily protein |
| MD05G1159700 | MD05G1159700 | 480.06173 | -12.55550 | 1.665455 | -7.538784 | 0e+00 | 0.00e+00 | Rola-high | Bifunctional inhibitor/lipid-transfer protein/seed storage 2S albumin superfamily protein |
| MD04G | MD04G | 3084.3 | -11.55366 | 1.32 | -8.7357 | 0e+ | 0.00 | Rola- | GDSL-like Lipase/Acylhydrolase superfamily protein |

| gene_id | gene_name | baseMean | log2FoldChange | lfcSE | stat | pvalue | padj | direction | Description |
|---|---|---|---|---|---|---|---|---|---|
| 1234300 | 1234300 | 9032 | | 2566 | 89 | 00 | e+00 | high | |
| MD00G1174400 | MD00G1174400 | 423.66967 | -11.54533 | 1.5437994 | -7.478541 | 0e+00 | 0.000e+00 | Rola-high | fatty acid desaturase 5 |
| MD14G1081900 | MD14G1081900 | 384.01231 | -11.07330 | 1.7352117 | -6.381510 | 0e+00 | 9.000e-07 | Rola-high | Gibberellin-regulated family protein |
| MD17G1140600 | MD17G1140600 | 262.49148 | -11.05972 | 1.9522557 | -5.665094 | 1e-07 | 1.83e-05 | Rola-high | protodermal factor 1 |
| MD02G1138900 | MD02G1138900 | 114.21657 | 10.20743 | 1.8901770 | 5.400273 | 6e-07 | 5.93e-05 | Rin-high | Peroxidase superfamily protein |

#### Table preview 25

**Rin vs control DESeq2 results using lfcThreshold = 1.**

| gene_id | gene_name | baseMean | log2FoldChange | lfcSE | stat | pvalue | padj | Description |
|---|---|---|---|---|---|---|---|---|
| MD07G1003800 | MD07G1003800 | 266.79193 | 4.398128 | 0.2342835 | 18.77267 | 0 | 0 | O-fucosyltransferase family protein |
| MD11G1062600 | MD11G1062600 | 258.51698 | -3.449746 | 0.1995429 | -17.28824 | 0 | 0 | UDP-Glycosyltransferase superfamily protein |
| MD10G1169100 | MD10G1169100 | 286.27071 | -6.123964 | 0.4275772 | -14.32248 | 0 | 0 | Pectin lyase-like superfamily protein |
| MD04G1069000 | MD04G1069000 | 3981.34923 | -2.358464 | 0.1267419 | -18.60841 | 0 | 0 | Glucose-methanol-choline (GMC) oxidoreductase family protein |
| MD11G1161800 | MD11G1161800 | 929.52476 | -3.643816 | 0.2462125 | -14.79947 | 0 | 0 | Jojoba acyl CoA reductase-related male sterility protein |
| MD15G1400200 | MD15G1400200 | 35.38328 | -22.969079 | 2.0556327 | -11.17373 | 0 | 0 | GDSL-like Lipase/Acylhydrolase superfamily protein |
| MD11G1169600 | MD11G1169600 | 74.33378 | 8.564226 | 0.7098082 | 12.06555 | 0 | 0 | F-box family protein |
| MD13G1087700 | MD13G1087700 | 369.91439 | -2.304956 | 0.1230072 | -18.73839 | 0 | 0 | glyoxal oxidase-related protein |
| MD15G1262600 | MD15G1262600 | 2196.07495 | -1.941514 | 0.0892034 | -21.76503 | 0 | 0 | MD15G1262600 |
| MD12G1 | MD12G1 | 64.518 | -23.98427 | 2.279 | -10.52 | 0 | 0 | Integrase-type DNA-binding superfamily protein |

| gene_id | gene_name | baseMean | log2FoldChange | lfcSE | stat | pvalue | padj | Description |
|---|---|---|---|---|---|---|---|---|
| 067600 | 067600 | 32 | 9 | 6057 | 124 | | | |

#### Table preview 26

**Rin vs control lfcThreshold = 1 results sorted by log2FC.**

| gene_id | gene_name | baseMean | log2FoldChange | lfcSE | stat | pvalue | padj | Description |
|---|---|---|---|---|---|---|---|---|
| MD02G1028600 | MD02G1028600 | 176.28388 | 9.771910 | 1.2297158 | 7.946478 | 0.00e+00 | 0.000000 | blue-copper-binding protein |
| MD15G1131700 | MD15G1131700 | 279.91793 | 9.111016 | 1.5932583 | 5.718481 | 2.00e-07 | 0.0000114 | Chalcone and stilbene synthase family protein |
| MD08G1048800 | MD08G1048800 | 100.15939 | 8.834990 | 1.1049926 | 7.995520 | 0.00e+00 | 0.000000 | MD08G1048800 |
| MD11G1169600 | MD11G1169600 | 74.33378 | 8.564226 | 0.7098082 | 12.065550 | 0.00e+00 | 0.000000 | F-box family protein |
| MD | MD | 49 | 8.51 | 1. | 5. | 9. | 0. | sodium/calcium exchanger family |

| gene_id | gene_name | baseMean | log2FoldChange | lfcSE | stat | pvalue | padj | Description |
|---|---|---|---|---|---|---|---|---|
| 03G1238300 | 03G1238300 | .54020 | 0665 | 5732120 | 409738 | 00e-07 | 0000457 | protein / calcium-binding EF hand family protein |
| MD15G1132100 | MD15G1132100 | 112.38722 | 8.332038 | 1.3447346 | 6.196047 | 0.000e+00 | 0.0000021 | Chalcone and stilbene synthase family protein |
| MD12G1075400 | MD12G1075400 | 84.48708 | 8.298809 | 0.9334089 | 8.890862 | 0.000e+00 | 0.0000000 | Glycosyltransferase family 61 protein |
| MD16G1160100 | MD16G1160100 | 52.64405 | 8.229399 | 1.8109680 | 4.544199 | 3.29e-05 | 0.0010433 | MLP-like protein 423 |
| MD02G1303900 | MD02G1303900 | 50.12084 | 8.095013 | 1.5698875 | 5.156429 | 3.10e-06 | 0.0001351 | NAD(P)-binding Rossmann-fold superfamily protein |
| MD00G1013000 | MD00G1013000 | 609.15576 | 8.091411 | 0.8052716 | 10.048052 | 0.000e+00 | 0.0000000 | Glucose-methanol-choline (GMC) oxidoreductase family protein |

#### Table preview 27

**Rin-high candidates in Rin vs control.**

| gene_id | gene_name | baseMean | log2FoldChange | lfcSE | stat | pvalue | padj | Description |
|---|---|---|---|---|---|---|---|---|
| MD02G1028600 | MD02G1028600 | 176.28388 | 9.771910 | 1.2297158 | 7.946478 | 0.00e+00 | 0.0000000 | blue-copper-binding protein |
| MD15G1131700 | MD15G1131700 | 279.91793 | 9.111016 | 1.5932583 | 5.718481 | 2.00e-07 | 0.0000114 | Chalcone and stilbene synthase family protein |
| MD08G1048800 | MD08G1048800 | 100.15939 | 8.834990 | 1.1049926 | 7.995520 | 0.00e+00 | 0.0000000 | MD08G1048800 |
| MD11G1169600 | MD11G1169600 | 74.33378 | 8.564226 | 0.7098082 | 12.065550 | 0.00e+00 | 0.0000000 | F-box family protein |
| MD03G1238300 | MD03G1238300 | 49.54020 | 8.510665 | 1.5732120 | 5.409738 | 9.00e-07 | 0.0000457 | sodium/calcium exchanger family protein / calcium-binding EF hand family protein |
| MD15G1 | MD15G1 | 112.38 | 8.33203 | 1.3447 | 6.1960 | 0.00 | 0.0000 | Chalcone and stilbene synthase family protein |

| gene_id | gene_name | baseMean | log2FoldChange | lfcSE | stat | pvalue | padj | Description |
|---|---|---|---|---|---|---|---|---|
| 132100 | 132100 | 722 | 8 | 346 | 47 | e+00 | 021 | |
| MD12G1075400 | MD12G1075400 | 84.48708 | 8.298809 | 0.9334089 | 8.890862 | 0.00e+00 | 0.0000000 | Glycosyltransferase family 61 protein |
| MD16G1160100 | MD16G1160100 | 52.64405 | 8.229399 | 1.8109680 | 4.544199 | 3.29e-05 | 0.0010433 | MLP-like protein 423 |
| MD02G1303900 | MD02G1303900 | 50.12084 | 8.095013 | 1.5698875 | 5.156429 | 3.10e-06 | 0.0001351 | NAD(P)-binding Rossmann-fold superfamily protein |
| MD00G1013000 | MD00G1013000 | 609.15576 | 8.091411 | 0.8052716 | 10.048052 | 0.00e+00 | 0.0000000 | Glucose-methanol-choline (GMC) oxidoreductase family protein |

#### Table preview 28

**Control-high candidates in Rin vs control.**

| gene_id | gene_name | baseMean | log2FoldChange | lfcSE | stat | pvalue | padj | Description |
|---|---|---|---|---|---|---|---|---|
| MD12G1067600 | MD12G1067600 | 64.51832 | -23.98428 | 2.279606 | -10.521240 | 0e+00 | 0.0e+00 | Integrase-type DNA-binding superfamily protein |
| MD15G1400200 | MD15G1400200 | 35.38328 | -22.96908 | 2.055633 | -11.173727 | 0e+00 | 0.0e+00 | GDSL-like Lipase/Acylhydrolase superfamily protein |
| MD01G1210800 | MD01G1210800 | 38.79491 | -22.11585 | 2.102506 | -10.518806 | 0e+00 | 0.0e+00 | ABC-2 type transporter family protein |
| MD12G1252200 | MD12G1252200 | 8353.64251 | -12.77706 | 1.450641 | -8.807874 | 0e+00 | 0.0e+00 | GDSL-like Lipase/Acylhydrolase superfamily protein |
| MD14G1081900 | MD14G1081900 | 384.01231 | -10.70403 | 1.559470 | -6.863891 | 0e+00 | 0.0e+00 | Gibberellin-regulated family protein |
| MD04G1 | MD04G1 | 3084.3 | -10.61915 | 1.17 | -9.0551 | 0e+ | 0.0 | GDSL-like Lipase/Acylhydrolase superfamily protein |

| gene_id | gene_name | baseMean | log2FoldChange | lfcSE | stat | pvalue | padj | Description |
|---|---|---|---|---|---|---|---|---|
| 234300 | 234300 | 9032 | | 2726 | 04 | 00 | e+00 | |
| MD17G1140600 | MD17G1140600 | 262.49148 | -10.26288 | 1.746936 | -5.874792 | 1e-07 | 4.2e-06 | protodermal factor 1 |
| MD05G1159700 | MD05G1159700 | 480.06173 | -10.22893 | 1.501495 | -6.812495 | 0e+00 | 1.0e-07 | Bifunctional inhibitor/lipid-transfer protein/seed storage 2S albumin superfamily protein |
| MD09G1154000 | MD09G1154000 | 162.97381 | -10.04298 | 1.567985 | -6.405027 | 0e+00 | 4.0e-07 | protodermal factor 1 |
| MD00G1174400 | MD00G1174400 | 423.66967 | -10.01521 | 1.394745 | -7.180672 | 0e+00 | 0.0e+00 | fatty acid desaturase 5 |

#### Table preview 29

**Rin vs control candidates sorted by absolute log2FC.**

| gene_id | gene_name | baseMean | log2FoldChange | lfcSE | stat | pvalue | padj | direction | Description |
|---|---|---|---|---|---|---|---|---|---|
| MD12G1067600 | MD12G1067600 | 64.51832 | -23.98428 | 2.2796006 | -10.52121240 | 0e+00 | 0.0e+00 | C-high | Integrase-type DNA-binding superfamily protein |
| MD15G1400200 | MD15G1400200 | 35.38328 | -22.96908 | 2.0556633 | -11.17373727 | 0e+00 | 0.0e+00 | C-high | GDSL-like Lipase/Acylhydrolase superfamily protein |
| MD01G1210800 | MD01G1210800 | 38.79491 | -22.11585 | 2.1025006 | -10.51888806 | 0e+00 | 0.0e+00 | C-high | ABC-2 type transporter family protein |
| MD12G1252200 | MD12G1252200 | 8353.64251 | -12.77706 | 1.4506641 | -8.807874 | 0e+00 | 0.0e+00 | C-high | GDSL-like Lipase/Acylhydrolase superfamily protein |
| MD14G1081900 | MD14G1081900 | 384.01231 | -10.70403 | 1.5594 70 | -6.863891 | 0e+00 | 0.0e+00 | C-high | Gibberellin-regulated family protein |
| MD04G | MD04G | 3084.3 | -10.61915 | 1.17 | -9.0551 | 0e+ | 0.0 | C-hig | GDSL-like Lipase/Acylhydrolase superfamily protein |

| gene_id | gene_name | baseMean | log2FoldChange | lfcSE | stat | pvalue | padj | direction | Description |
|---|---|---|---|---|---|---|---|---|---|
| 1234300 | 1234300 | 9032 | | 2726 | 04 | 00 | e+00 | h | |
| MD17G1140600 | MD17G1140600 | 262.49148 | -10.26288 | 1.746936 | -5.874792 | 1e-07 | 4.2e-06 | C-high | protodermal factor 1 |
| MD05G1159700 | MD05G1159700 | 480.06173 | -10.22893 | 1.501495 | -6.812495 | 0e+00 | 1.0e-07 | C-high | Bifunctional inhibitor/lipid-transfer protein/seed storage 2S albumin superfamily protein |
| MD09G1154000 | MD09G1154000 | 162.97381 | -10.04298 | 1.567985 | -6.405027 | 0e+00 | 4.0e-07 | C-high | protodermal factor 1 |
| MD00G1174400 | MD00G1174400 | 423.66967 | -10.01521 | 1.394745 | -7.180672 | 0e+00 | 0.0e+00 | C-high | fatty acid desaturase 5 |

#### Table preview 30

**Rola vs control DESeq2 results using lfcThreshold = 1.**

| gene_id | gene_name | baseMean | log2FoldChange | lfcSE | stat | pvalue | padj | Description |
|---|---|---|---|---|---|---|---|---|
| MD02G1080400 | MD02G1080400 | 59.017619 | 6.347044 | 0.4854996 | 13.073222 | 0 | 0 | Protein of unknown function |
| MD09G1084600 | MD09G1084600 | 46.422522 | 7.241337 | 0.6401450 | 11.312026 | 0 | 0 | allene oxide cyclase 4 |
| MD15G1211600 | MD15G1211600 | 260.952687 | 3.109105 | 0.2409064 | 12.905861 | 0 | 0 | Protein of unknown function |
| MD09G1233000 | MD09G1233000 | 9.009476 | 7.611130 | 0.7681272 | 9.908684 | 0 | 0 | basic helix-loop-helix (bHLH) DNA-binding superfamily protein |
| MD03G1152000 | MD03G1152000 | 226.049551 | 3.603033 | 0.3039801 | 11.852857 | 0 | 0 | Polynucleotidyl transferase |
| MD04G1147900 | MD04G1147900 | 105.581686 | 7.227458 | 0.7538334 | 9.587606 | 0 | 0 | HXXXD-type acyl-transferase family protein |
| MD09G1168300 | MD09G1168300 | 102.970272 | 5.947692 | 0.5991953 | 9.926134 | 0 | 0 | Terpenoid cyclases family protein |
| MD12G1040900 | MD12G1040900 | 16.595471 | 5.825247 | 0.5914281 | 9.849458 | 0 | 0 | plant U-box 23 |
| MD10G1293600 | MD10G1293600 | 175.553522 | 7.995223 | 0.8594692 | 9.302512 | 0 | 0 | Protein of unknown function (DUF506) |
| MD08G1 | MD08G1 | 338.97 | 4.701015 | 0.458 | 10.25 | 0 | 0 | salt tolerance zinc finger |

| gene_id | gene_name | baseMean | log2FoldChange | lfcSE | stat | pvalue | padj | Description |
|---|---|---|---|---|---|---|---|---|
| 086500 | 086500 | 1601 | | 3110 | 7260 | | | |

#### Table preview 31

**Rola vs control lfcThreshold = 1 results sorted by log2FC.**

| gene_id | gene_name | baseMean | log2FoldChange | lfcSE | stat | pvalue | padj | Description |
|---|---|---|---|---|---|---|---|---|
| MD15G1131700 | MD15G1131700 | 279.91793 | 11.158248 | 1.777807 | 6.276411 | 0.00e+00 | 0.0000015 | Chalcone and stilbene synthase family protein |
| MD02G1028600 | MD02G1028600 | 176.28388 | 9.958051 | 1.348671 | 7.383603 | 0.00e+00 | 0.00000 | blue-copper-binding protein |
| MD16G1056600 | MD16G1056600 | 113.16617 | 9.870722 | 1.762060 | 5.601807 | 2.00e-07 | 0.000340 | cytochrome P450 |
| MD14G1010300 | MD14G1010300 | 53.98611 | 9.491851 | 1.152815 | 8.233625 | 0.00e+00 | 0.00000 | Peroxidase superfamily protein |

| gene_id | gene_name | baseMean | log2FoldChange | lfcSE | stat | pvalue | padj | Description |
|---|---|---|---|---|---|---|---|---|
| MD08G1242800 | MD08G1242800 | 59.07963 | 9.255027 | 1.418584 | 6.524131 | 0.00e+00 | 0.0000009 | S-adenosyl-L-methionine-dependent methyltransferases superfamily protein |
| MD07G1292500 | MD07G1292500 | 33.28207 | 9.185255 | 1.905110 | 4.821377 | 8.70e-06 | 0.0006608 | cyclic nucleotide gated channel 1 |
| MD15G1132000 | MD15G1132000 | 51.14695 | 9.150195 | 1.636464 | 5.591442 | 3.00e-07 | 0.0000430 | Chalcone and stilbene synthase family protein |
| MD16G1160100 | MD16G1160100 | 52.64405 | 9.011560 | 2.026111 | 4.447712 | 3.88e-05 | 0.0023192 | MLP-like protein 423 |
| MD07G1291400 | MD07G1291400 | 29.02242 | 8.944365 | 1.632491 | 5.478969 | 6.00e-07 | 0.0000666 | cyclic nucleotide gated channel 1 |
| MD06G109440 | MD06G109440 | 32.19416 | 8.856732 | 1.310 | 6.758 | 0.00e+ | 0.000000 | Late embryogenesis abundant (LEA) hydroxyproline-rich glycoprotein family |

| gene_id | gene_name | baseMean | log2FoldChange | lfcSE | stat | pvalue | padj | Description |
|---|---|---|---|---|---|---|---|---|
| 0 | 0 | | | 473 | 425 | 00 | 4 | |

#### Table preview 32

**Rola-high candidates in Rola vs control.**

| gene_id | gene_name | baseMean | log2FoldChange | lfcSE | stat | pvalue | padj | Description |
|---|---|---|---|---|---|---|---|---|
| MD15G1131700 | MD15G1131700 | 279.91793 | 11.158248 | 1.777807 | 6.276411 | 0.00e+00 | 0.0000015 | Chalcone and stilbene synthase family protein |
| MD02G1028600 | MD02G1028600 | 176.28388 | 9.958051 | 1.348671 | 7.383603 | 0.00e+00 | 0.0000000 | blue-copper-binding protein |
| MD16G1056600 | MD16G1056600 | 113.16617 | 9.870722 | 1.762060 | 5.601807 | 2.00e-07 | 0.0000340 | cytochrome P450 |
| MD14G1010300 | MD14G1010300 | 53.98611 | 9.491851 | 1.152812 | 8.233620 | 0.00e+00 | 0.0000000 | Peroxidase superfamily protein |

| gene_id | gene_name | baseMean | log2FoldChange | lfcSE | stat | pvalue | padj | Description |
|---|---|---|---|---|---|---|---|---|
| | | | | 5 | 5 | | | |
| MD08G1242800 | MD08G1242800 | 59.07963 | 9.255027 | 1.418584 | 6.524131 | 0.00e+00 | 0.0000009 | S-adenosyl-L-methionine-dependent methyltransferases superfamily protein |
| MD07G1292500 | MD07G1292500 | 33.28207 | 9.185255 | 1.905110 | 4.821377 | 8.70e-06 | 0.0006608 | cyclic nucleotide gated channel 1 |
| MD15G1132000 | MD15G1132000 | 51.14695 | 9.150195 | 1.636464 | 5.591442 | 3.00e-07 | 0.0000430 | Chalcone and stilbene synthase family protein |
| MD16G1160100 | MD16G1160100 | 52.64405 | 9.011560 | 2.026111 | 4.447712 | 3.88e-05 | 0.0023192 | MLP-like protein 423 |
| MD07G1291400 | MD07G1291400 | 29.02242 | 8.944365 | 1.632491 | 5.478969 | 6.00e-07 | 0.0000666 | cyclic nucleotide gated channel 1 |
| MD06G109 | MD06G109 | 32.194 | 8.856732 | 1.31 | 6.75 | 0.00e | 0.000000 | Late embryogenesis abundant (LEA) hydroxyproline-rich glycoprotein family |

| gene_id | gene_name | baseMean | log2FoldChange | lfcSE | stat | pvalue | padj | Description |
|---|---|---|---|---|---|---|---|---|
| 4400 | 4400 | 16 | | 0473 | 8425 | +00 | 004 | |

#### Table preview 33

**Control-high candidates in Rola vs control.**

| gene_id | gene_name | baseMean | log2FoldChange | lfcSE | stat | pvalue | padj | Description |
|---|---|---|---|---|---|---|---|---|
| MD07G1266700 | MD07G1266700 | 402.564273 | -6.606136 | 1.3344916 | -4.950301 | 0.0000133 | 0.0009506 | MD07G1266700 |
| MD04G1140600 | MD04G1140600 | 16.342142 | -6.296526 | 1.1476020 | -5.486681 | 0.0000020 | 0.0001841 | HSP20-like chaperones superfamily protein |
| MD11G1315600 | MD11G1315600 | 1955.115241 | -6.218783 | 1.2396675 | -5.016493 | 0.0000128 | 0.0009224 | Adenine nucleotide alpha hydrolases-like superfamily protein |
| MD13G1003800 | MD13G1003800 | 714.716554 | -6.196723 | 1.2678044 | -4.887759 | 0.0000208 | 0.0013787 | DNAJ heat shock N-terminal domain-containing protein |
| MD07G1299100 | MD07G1299100 | 3094.631934 | -6.178670 | 1.0829187 | -5.705571 | 0.0000009 | 0.0000945 | pseudo-response regulator 5 |
| MD16G | MD16G | 1006.9 | -6.08 | 1.128 | -5.39 | 0.000 | 0.000 | flavin-binding |

| gene_id | gene_name | baseMean | log2FoldChange | lfcSE | stat | pvalue | padj | Description |
|---|---|---|---|---|---|---|---|---|
| 1018000 | 1018000 | 27461 | 4471 | 6093 | 1123 | 0033 | 2907 | |
| MD05G1028500 | MD05G1028500 | 1006.504967 | -5.914399 | 0.9153269 | -6.461515 | 0.0000000 | 0.0000077 | MD05G1028500 |
| MD13G1279400 | MD13G1279400 | 69.298535 | -5.762100 | 1.2568667 | -4.584496 | 0.0000757 | 0.0040109 | Transmembrane amino acid transporter family protein |
| MD08G1036100 | MD08G1036100 | 22.037949 | -5.542078 | 1.1604434 | -4.775828 | 0.0000454 | 0.0026170 | Protein of unknown function (DUF607) |
| MD00G1134100 | MD00G1134100 | 7.839481 | -5.371505 | 1.2239059 | -4.388822 | 0.0001774 | 0.0081034 | Protein of unknown function (DUF607) |

#### Table preview 34

**Rola vs control candidates sorted by absolute log2FC.**

| gene_id | gene_name | baseMean | log2FoldChange | lfcSE | stat | pvalue | padj | direction | Description |
|---|---|---|---|---|---|---|---|---|---|
| MD15G1131700 | MD15G1131700 | 279.91793 | 11.158248 | 1.777807 | 6.276411 | 0.00e+00 | 0.0000015 | Rola-high | Chalcone and stilbene synthase family protein |
| MD | MD | 17 | 9.95 | 1. | 7. | 0. | 0. | Ro | blue-copper-binding protein |

| gene_id | gene_name | baseMean | log2FoldChange | lfcSE | stat | pvalue | padj | direction | Description |
|---|---|---|---|---|---|---|---|---|---|
| 02G1028600 | 02G1028600 | 6.283888 | 8051 | 348671 | 383603 | 00e+00 | 0000000 | la-high | |
| MD16G1056600 | MD16G1056600 | 113.16617 | 9.870722 | 1.7620600 | 5.6018087 | 2.00e-07 | 0.0000340 | Rola-high | cytochrome P450 |
| MD14G1010300 | MD14G1010300 | 53.98611 | 9.491851 | 1.1528150 | 8.2336225 | 0.00e+00 | 0.0000000 | Rola-high | Peroxidase superfamily protein |
| MD08G1242800 | MD08G1242800 | 59.07963 | 9.255027 | 1.4185840 | 6.5241310 | 0.00e+00 | 0.0000009 | Rola-high | S-adenosyl-L-methionine-dependent methyltransferases superfamily protein |
| MD07G1292500 | MD07G1292500 | 33.28207 | 9.185255 | 1.9051110 | 4.8213777 | 8.70e-06 | 0.0006608 | Rola-high | cyclic nucleotide gated channel 1 |
| MD15G1132000 | MD15G1132000 | 51.14695 | 9.150195 | 1.6364 | 5.5914 | 3.00e-0 | 0.0000430 | Rola-high | Chalcone and stilbene synthase family protein |

| gene_id | gene_name | baseMean | log2FoldChange | lfcSE | stat | pvalue | padj | direction | Description |
|---|---|---|---|---|---|---|---|---|---|
| | | | | 64 | 42 | 7 | | | |
| MD16G1160100 | MD16G1160100 | 52.644405 | 9.011560 | 2.0261111 | 4.4477712 | 3.88e-05 | 0.0023192 | Rola-high | MLP-like protein 423 |
| MD07G1291400 | MD07G1291400 | 29.02242 | 8.944365 | 1.6324991 | 5.4789669 | 6.00e-07 | 0.0000666 | Rola-high | cyclic nucleotide gated channel 1 |
| MD06G1094400 | MD06G1094400 | 32.19416 | 8.856732 | 1.3104473 | 6.7584425 | 0.00e+00 | 0.0000004 | Rola-high | Late embryogenesis abundant (LEA) hydroxyproline-rich glycoprotein family |

#### Table preview 35

**Combined comparison table across Rin vs control, Rola vs control, and Rin vs Rola.**

| gene_id | gene_name | Description | baseMean_Rin_vs_C | log2FC_Rin_vs_C | lfcSE_Rin_vs_C | stat_Rin_vs_C | pvalue_Rin_vs_C | padj_Rin_vs_C | baseMean_Rola_vs_C | log2FC_Rola_vs_C | lfcSE_Rola_vs_C | stat_Rola_vs_C | pvalue_Rola_vs_C | padj_Rola_vs_C | baseMean_Rin_vs_Rola | log2FC_Rin_vs_Rola | lfcSE_Rin_vs_Rola | stat_Rin_vs_Rola | pvalue_Rin_vs_Rola | padj_Rin_vs_Rola | Rin_up_vs_C | Rin_down_vs_C | Rola_up_vs_C | Rola_down_vs_C | class |
|---|---|---|---|---|---|---|---|---|---|---|---|---|---|---|---|---|---|---|---|---|---|---|---|---|---|
| MD00G1000100 | MD00G1000100 | RNA-binding (RRM/RBD/RNP motifs) family protein | 1.157524 | -0.4299435 | 0.9807485 | -0.4383830 | 0.7918814 | NA | 1.157524 | 1.0819829 | 0.9587068 | 1.1285858 | 0.4808671 | 1 | 1.157524 | -1.5119264 | 1.0029187 | -1.507526 | 0.3110020 | 1 | FALSE | FALSE | FALSE | FALSE | not_strong_DE |
| MD00G1000200 | MD00G1000200 | Ribosomal protein L35 | 101.844066 | -0.5225327 | 0.177649 | -2.99394590 | 0.9963838 | 1 | 101.844066 | -0.2986476 | 0.2000608 | -1.4927841 | 0.9997723 | 1 | 101.844066 | -0.2238851 | 0.2028969 | -1.103443 | 0.9999347 | 1 | FALSE | FALSE | FALSE | FALSE | not_strong_DE |

| gene_id | gene_name | Description | baseMean_Rin_vs_C | log2FC_Rin_vs_C | lfcSE_Rin_vs_C | stat_Rin_vs_C | pvalue_Rin_vs_C | padj_Rin_vs_C | baseMean_Rola_vs_C | log2FC_Rola_vs_C | lfcSE_Rola_vs_C | stat_Rola_vs_C | pvalue_Rola_vs_C | padj_Rola_vs_C | baseMean_Rin_vs_Rola | log2FC_Rin_vs_Rola | lfcSE_Rin_vs_Rola | stat_Rin_vs_Rola | pvalue_Rin_vs_Rola | padj_Rin_vs_Rola | Rin_up_vs_C | Rin_down_vs_C | Rola_up_vs_C | Rola_down_vs_C | class |
|---|---|---|---|---|---|---|---|---|---|---|---|---|---|---|---|---|---|---|---|---|---|---|---|---|---|
| MD00G1000300 | MD00G1000300 | MD00G1000300 | 7.996346 | 0.2158363 | 0.4371772 | 0.4937044 | 0.9662780 | 1 | 7.996346 | -1.0313521 | 0.5380072 | -1.9169855 | 0.4768448 | 1 | 7.996346 | 1.2471884 | 0.5354766 | 2.329118 | 0.3221894 | 1 | FALSE | FALSE | FALSE | FALSE | not_strong_DE |
| MD00G1000400 | MD00G1000400 | alpha/beta-Hydrolases superfamily protein | NA | NA | NA | NA | NA | NA | NA | NA | NA | NA | NA | NA | NA | NA | NA | NA | NA | NA | FALSE | FALSE | FALSE | FALSE | not_strong_DE |

| gene_id | gene_name | Description | baseMean_Rin_vs_C | log2FC_Rin_vs_C | lfcSE_Rin_vs_C | stat_Rin_vs_C | pvalue_Rin_vs_C | padj_Rin_vs_C | baseMean_Rola_vs_C | log2FC_Rola_vs_C | lfcSE_Rola_vs_C | stat_Rola_vs_C | pvalue_Rola_vs_C | padj_Rola_vs_C | baseMean_Rin_vs_Rola | log2FC_Rin_vs_Rola | lfcSE_Rin_vs_Rola | stat_Rin_vs_Rola | pvalue_Rin_vs_Rola | padj_Rin_vs_Rola | Rin_up_vs_C | Rin_down_vs_C | Rola_up_vs_C | Rola_down_vs_C | class |
|---|---|---|---|---|---|---|---|---|---|---|---|---|---|---|---|---|---|---|---|---|---|---|---|---|---|
| MD00G1000500 | MD00G1000500 | Homeodomain-like superfamily protein | 10.810401 | -1.0915613 | 0.4580468 | -2.3803782 | 0.4207838 | 1 | 10.810401 | 0.3336363 | 0.4787807 | 0.6968458 | 0.9206799 | 1 | 10.810401 | -1.4251976 | 0.5052308 | -2.8200884 | 0.2000093 | 1 | FALSE | FALSE | FALSE | FALSE | not_strong_DE |
| MD00G1000600 | MD00G1000600 | Domain of unknown function (DUF1995) | 2003.066689 | -0.0227470 | 0.197773 | -0.1150130 | 0.9999997 | 1 | 2003.066689 | -0.3846142 | 0.2255758 | -1.7050328 | 0.9968147 | 1 | 2003.066689 | 0.3618672 | 0.2255923 | 1.604076 | 0.9976631 | 1 | FALSE | FALSE | FALSE | FALSE | not_strong_DE |

| gene_id | gene_name | Description | baseMean_Rin_vs_C | log2FC_Rin_vs_C | lfcSE_Rin_vs_C | stat_Rin_vs_C | pvalue_Rin_vs_C | padj_Rin_vs_C | baseMean_Rola_vs_C | log2FC_Rola_vs_C | lfcSE_Rola_vs_C | stat_Rola_vs_C | pvalue_Rola_vs_C | padj_Rola_vs_C | baseMean_Rin_vs_Rola | log2FC_Rin_vs_Rola | lfcSE_Rin_vs_Rola | stat_Rin_vs_Rola | pvalue_Rin_vs_Rola | padj_Rin_vs_Rola | Rin_up_vs_C | Rin_down_vs_C | Rola_up_vs_C | Rola_down_vs_C | class |
|---|---|---|---|---|---|---|---|---|---|---|---|---|---|---|---|---|---|---|---|---|---|---|---|---|---|
| MD00G1000700 | MD00G1000700 | Protein of unknown function | 102.461280 | -0.3065771 | 0.1783031 | -1.7194158 | 0.9999497 | 1 | 102.461280 | 0.0261734 | 0.1998102 | 0.1309915 | 0.9999996 | 1 | 102.461280 | -0.3327505 | 0.2016675 | -1.649995 | 0.9995313 | 1 | FALSE | FALSE | FALSE | FALSE | not_strong_DE |
| MD00G1000800 | MD00G1000800 | Mob1/phocein family protein | 696.842140 | 0.2649626 | 0.1156451 | 2.2911704 | 1.0000000 | 1 | 696.842140 | 0.0103931 | 0.1318994 | 0.0787956 | 1.0000000 | 1 | 696.842140 | 0.2545695 | 0.1316665 | 1.933442 | 1.0000000 | 1 | FALSE | FALSE | FALSE | FALSE | not_strong_DE |

| gene_id | gene_name | Description | baseMean_Rin_vs_C | log2FC_Rin_vs_C | lfcSE_Rin_vs_C | stat_Rin_vs_C | pvalue_Rin_vs_C | padj_Rin_vs_C | baseMean_Rola_vs_C | log2FC_Rola_vs_C | lfcSE_Rola_vs_C | stat_Rola_vs_C | pvalue_Rola_vs_C | padj_Rola_vs_C | baseMean_Rin_vs_Rola | log2FC_Rin_vs_Rola | lfcSE_Rin_vs_Rola | stat_Rin_vs_Rola | pvalue_Rin_vs_Rola | padj_Rin_vs_Rola | Rin_up_vs_C | Rin_down_vs_C | Rola_up_vs_C | Rola_down_vs_C | class |
|---|---|---|---|---|---|---|---|---|---|---|---|---|---|---|---|---|---|---|---|---|---|---|---|---|---|
| MD00G1000900 | MD00G1000900 | MD00G1000900 | 13.598212 | -0.1997888 | 0.2517512 | -0.7935960 | 0.9992609 | 1 | 13.598212 | 0.1730295 | 0.2664079 | 0.6494010 | 0.9990512 | 1 | 13.598212 | -0.3728183 | 0.2740095 | -1.3600604 | 0.9889578 | 1 | FALSE | FALSE | FALSE | FALSE | not_strong_DE |
| MD00G1100100 | MD00G1100100 | growth-regulating factor 2 | 89.146758 | -0.0876748 | 0.1423394 | -0.6159561 | 1.000000 | 1 | 89.146758 | 0.2767905 | 0.1569239 | 1.7638522 | 0.9999980 | 1 | 89.146758 | -0.3644654 | 0.1581021 | -2.305254 | 0.9999709 | 1 | FALSE | FALSE | FALSE | FALSE | not_strong_DE |

#### Table preview 36

**Classification summary of expression-response classes.**

| class | n |
|---|---|
| not_strong_DE | 50651 |
| Rin_specific_repressed | 708 |
| Rin_specific_induced | 508 |
| common_infection_induced | 494 |
| Rola_specific_induced | 332 |
| Rola_specific_repressed | 48 |
