## Supplementary tables and figures for "Transcriptome of apple cv. Ametyst in *Rvi6*-effective and *Rvi6*-breaking *Venturia inaequalis* interactions identifies defense candidates": 23_SupplementaryData_S14_DESeq2_Trinity.pdf

### DESeq2 analysis of Ametyst RNA-Seq data against Trinity de novo transcriptome

Mila

2026-09-02

#### Table of Contents

#### Analysis overview

This R Markdown document performs DESeq2 analysis of Ametyst RNA-Seq data quantified against the Trinity de novo transcriptome.

The Trinity input is a gene-level representative transcriptome: one representative sequence per Trinity gene. Therefore Salmon quantification is imported with `tximport` using a one-to-one `TXNAME -> GENEID` mapping, and DESeq2 is performed at the Trinity gene level.

The biological comparison is between three conditions:

- control
- `Rin`, representing the response to the *Venturia inaequalis* isolate from Rubin, which does not break Rvi6 resistance
- `Rola`, representing the response to the isolate from Rubinola, which breaks Rvi6 resistance

For the key contrast `Rin` versus `Rola`, positive log2 fold change means higher expression in `Rin`, while negative log2 fold change means higher expression in `Rola`.

The sample set is intentionally the same as in the corresponding GDDH-guided analysis:

- C01 to C08
- RIN01 to RIN08
- ROL01 to ROL05

The final DESeq2 design follows the attached GDDH-guided R Markdown script:

```
~ condition
```

Annotation is read from the OmicsBox-generated Trinity GFF3 file. The parser is adapted to OmicsBox attributes such as ID, Description, Gene, Ontology_term, Ontology_id, Enzyme_code, and Enzyme_name.

### Load packages

```
suppressPackageStartupMessages({
  library(tximport)
  library(DESeq2)
  library(SummarizedExperiment)
  library(dplyr)
  library(ggplot2)
  library(pheatmap)
  library(limma)
})
```

### 0. Paths and settings

```
# The project has lived on more than one volume. Resolving the root from a
candidate
# list instead of one hard-coded path is what stops a knit from half-running: the
knit
# of 12 Aug 2026 silently skipped section 16 because base_dir pointed at a volume
that
# was not mounted. Add new locations to the front of this list, do not edit them in.
base_dir_candidates <- c(
  "/Volumes/Data_Disk/Ametyst_re-analysis",
  "/Volumes/Mac_Data/Ametyst_re-analysis",
  getwd(),
  dirname(getwd()),
  dirname(dirname(getwd()))
)
found_root <- base_dir_candidates[dir.exists(file.path(base_dir_candidates,
"Trinity"))]
if (!length(found_root)) {
  stop("Project root not found. Tried:\n  ",
      paste(base_dir_candidates, collapse = "\n  "))
}
base_dir <- found_root[1]
cat("Project root resolved to:", base_dir, "\n")
```

```
## Project root resolved to: /Volumes/Data_Disk/Ametyst_re-analysis
```

```
trinity_dir <- file.path(base_dir, "Trinity")

trinity_fasta <- file.path(
  trinity_dir,
  "Trinity_ALL_Gene_Representatives_from_Transcripts.fasta"
)
```

```r
gff_file <- file.path(
  trinity_dir,
  "Trinity_Ametyst.gff"
)

annotation_file <- file.path(
  trinity_dir,
  "Trinity_Ametyst_OmicsBox_gene_annotation.tsv"
)

# Salmon quantification directory.
# The script checks several likely locations and uses the first existing one.
salmon_quant_candidates <- c(
  file.path(trinity_dir, "Salmon", "quant"),
  file.path(trinity_dir, "salmon", "quant"),
  file.path(trinity_dir, "quant"),
  file.path(base_dir, "salmon", "quant")
)

salmon_quant_dir <-
salmon_quant_candidates[dir.exists(salmon_quant_candidates)][1]

if (is.na(salmon_quant_dir)) {
  salmon_quant_dir <- file.path(trinity_dir, "Salmon", "quant")
  message("No Salmon quantification directory found automatically. Expected
location: ", salmon_quant_dir)
  message("Edit salmon_quant_dir in chunk 0 if your quant.sf files are elsewhere.")
}

# Optional metadata file. If it does not exist, metadata is generated from the fixed
sample list below.
metadata_file <- file.path(trinity_dir, "metadata", "samples_Trinity.tsv")

outdir <- file.path(trinity_dir, "DESeq2")
qcdir <- file.path(outdir, "QC_extra_plots")

dir.create(outdir, recursive = TRUE, showWarnings = FALSE)
dir.create(qcdir, recursive = TRUE, showWarnings = FALSE)

min_total_count <- 10
candidate_baseMean_cutoff <- 50
candidate_padj_cutoff <- 0.05
candidate_abs_lfc_cutoff <- 1

# Batch-design sensitivity analysis (section 16).
# Libraries were prepared in two batches about six months apart and sequenced
by two
# providers, which are perfectly aliased and therefore collapse into one batch
```

```
# rlog can be slow for large Trinity gene sets. Set TRUE only if needed.
run_rlog <- FALSE

has_vsn <- requireNamespace("vsn", quietly = TRUE)
has_NMF <- requireNamespace("NMF", quietly = TRUE)
has_ashr <- requireNamespace("ashr", quietly = TRUE)

cat("Base directory:\n", base_dir, "\n\n")

## Base directory:
##  /Volumes/Data_Disk/Ametyst_re-analysis

cat("Trinity FASTA:\n", trinity_fasta, "\n\n")

## Trinity FASTA:
##
/Volumes/Data_Disk/Ametyst_re-analysis/Trinity/Trinity_ALL_Gene_Representatives
_from_Transcripts.fasta

cat("OmicsBox GFF:\n", gff_file, "\n\n")

## OmicsBox GFF:
##  /Volumes/Data_Disk/Ametyst_re-analysis/Trinity/Trinity_Ametyst.gff

cat("Salmon quantification directory:\n", salmon_quant_dir, "\n\n")

## Salmon quantification directory:
##  /Volumes/Data_Disk/Ametyst_re-analysis/Trinity/Salmon/quant

cat("Output directory:\n", outdir, "\n\n")

## Output directory:
##  /Volumes/Data_Disk/Ametyst_re-analysis/Trinity/DESeq2
```

### 1. Helper functions

```
read_fasta_ids <- function(fasta_file) {
  if (!file.exists(fasta_file)) {
    stop("FASTA file does not exist: ", fasta_file)
  }
  headers <- grep("^>", readLines(fasta_file, warn = FALSE), value = TRUE)
  ids <- sub("^>", "", headers)
  ids <- sub("[[:space:]].*$", "", ids)
  unique(ids)
}

parse_omicsbox_attributes <- function(x) {
  items <- strsplit(x, ";", fixed = TRUE)[[1]]
  out <- list()

  for (item in items) {
```

```r
    if (!nzchar(item) || !grepl("=", item, fixed = TRUE)) next
    key <- sub("=.*$", "", item)
    value <- sub("^[^=]*=", "", item)
    value <- utils::URLdecode(value)
    value <- trimws(value)
    if (!nzchar(value) || value == "---NA---") next

    if (is.null(out[[key]])) {
      out[[key]] <- value
    } else {
      out[[key]] <- c(out[[key]], value)
    }
  }

  out
}

collapse_unique <- function(x, sep = "; ") {
  x <- unlist(x, use.names = FALSE)
  x <- trimws(x)
  x <- x[!is.na(x) & nzchar(x) & x != "---NA---"]
  if (length(x) == 0) return(NA_character_)
  paste(unique(x), collapse = sep)
}

create_gene_annotation_from_omicsbox_gff <- function(gff_file, annotation_file) {
  if (!file.exists(gff_file)) {
    stop("OmicsBox GFF file does not exist: ", gff_file)
  }

  message("Creating Trinity annotation from OmicsBox GFF: ", gff_file)

  gff_lines <- readLines(gff_file, warn = FALSE)
  gff_lines <- gff_lines[!grepl("^#", gff_lines) & nzchar(gff_lines)]

  parts <- strsplit(gff_lines, "\t", fixed = TRUE)
  parts <- parts[vapply(parts, length, integer(1)) >= 9]

  if (length(parts) == 0) {
    stop("No valid GFF records found in: ", gff_file)
  }

  records <- lapply(parts, function(p) {
    attrs <- parse_omicsbox_attributes(p[[9]])

    id <- collapse_unique(attrs[["ID"]])
    if (is.na(id) || !nzchar(id)) id <- p[[1]]

    gene_symbols <- collapse_unique(attrs[["Gene"]])
```

```
    description <- collapse_unique(attrs[["Description"]])
    ontology_terms <- collapse_unique(attrs[["Ontology_term"]], sep = ",")
    ontology_ids <- collapse_unique(attrs[["Ontology_id"]], sep = ",")
    enzyme_codes <- collapse_unique(attrs[["Enzyme_code"]], sep = ",")
    enzyme_names <- collapse_unique(attrs[["Enzyme_name"]], sep = ",")

    if (is.na(gene_symbols) || !nzchar(gene_symbols)) gene_symbols <- id
    if (is.na(description) || !nzchar(description)) description <- gene_symbols

    data.frame(
      gene_id = id,
      gene_name = gene_symbols,
      Description = description,
      chr = p[[1]],
      start = suppressWarnings(as.integer(p[[4]])),
      end = suppressWarnings(as.integer(p[[5]])),
      strand = p[[7]],
      type = p[[3]],
      Ontology_term = ontology_terms,
      Ontology_id = ontology_ids,
      Enzyme_code = enzyme_codes,
      Enzyme_name = enzyme_names,
      stringsAsFactors = FALSE
    )
  })

  annot_df <- dplyr::bind_rows(records)

  annotation <- annot_df |>
    dplyr::group_by(gene_id) |>
    dplyr::summarise(
      gene_name = dplyr::first(gene_name[!is.na(gene_name) & gene_name != ""]),
      Description = dplyr::first(Description[!is.na(Description) & Description !=
""]),
      chr = dplyr::first(chr),
      start = suppressWarnings(min(start, na.rm = TRUE)),
      end = suppressWarnings(max(end, na.rm = TRUE)),
      strand = dplyr::first(strand),
      Ontology_term = collapse_unique(Ontology_term, sep = ","),
      Ontology_id = collapse_unique(Ontology_id, sep = ","),
      Enzyme_code = collapse_unique(Enzyme_code, sep = ","),
      Enzyme_name = collapse_unique(Enzyme_name, sep = ","),
      .groups = "drop"
    ) |>
    dplyr::mutate(
      gene_name = ifelse(is.na(gene_name) | gene_name == "", gene_id,
gene_name),
      Description = ifelse(is.na(Description) | Description == "", gene_name,
Description)
```

```r
    ) |>
    dplyr::arrange(gene_id)

  write.table(
    annotation,
    annotation_file,
    sep = "\t",
    quote = FALSE,
    row.names = FALSE
  )

  message("Annotation written to: ", annotation_file)
  message("Annotated Trinity genes: ", nrow(annotation))

  invisible(annotation)
}

save_deseq_result <- function(res, filename, gene_annot, outdir) {
  df <- as.data.frame(res)
  df$gene_id <- rownames(df)

  df <- df |>
    left_join(gene_annot, by = "gene_id") |>
    mutate(
      gene_name = ifelse(is.na(gene_name) | gene_name == "", gene_id,
gene_name),
      Description = ifelse(is.na(Description) | Description == "", gene_name,
Description)
    ) |>
    select(
      gene_id,
      gene_name,
      baseMean,
      log2FoldChange,
      lfcSE,
      stat,
      pvalue,
      padj,
      Description,
      dplyr::any_of(c("Ontology_term", "Ontology_id", "Enzyme_code",
"Enzyme_name"))
    ) |>
    arrange(padj)

  write.csv(df, file.path(outdir, filename), row.names = FALSE)
  return(df)
}

write_png <- function(filename, width, height, expr, res = 150) {
```

```r
  png(
    filename = filename,
    width = width,
    height = height,
    units = "in",
    res = res,
    type = "cairo-png"
  )
  on.exit(dev.off(), add = TRUE)
  force(expr)
}

plot_candidate_heatmap <- function(genes, filename, title, vsd_mat, ann, res_df) {
  genes <- intersect(genes, rownames(vsd_mat))

  if (length(genes) < 2) {
    message("Skipping ", filename, ": fewer than 2 genes after intersecting with
VST matrix.")
    return(invisible(NULL))
  }

  mat <- vsd_mat[genes, , drop = FALSE]
  mat_scaled <- t(scale(t(mat)))

  labels <- res_df$Description[match(genes, res_df$gene_id)]
  labels[is.na(labels) | labels == ""] <- res_df$gene_name[match(genes,
res_df$gene_id)]
  labels[is.na(labels) | labels == ""] <- genes
  rownames(mat_scaled) <- make.unique(labels)

  write_png(file.path(qcdir, filename), 10, 9, {
    pheatmap(
      mat_scaled,
      annotation_col = ann,
      show_rownames = TRUE,
      fontsize_row = 5,
      main = title,
      cluster_cols = TRUE,
      cluster_rows = TRUE
    )
  })
}

mean_sd_df <- function(mat, label) {
  data.frame(
    transformation = label,
    mean = rowMeans(mat, na.rm = TRUE),
    sd = apply(mat, 1, sd, na.rm = TRUE)
  )
```

```r
}

show_png <- function(path, caption = NULL, width = "100%") {
  if (!file.exists(path)) {
    cat("\n\nMissing PNG file: `", path, "`\n\n", sep = "")
    return(invisible(NULL))
  }

  img_path <- path
  if (requireNamespace("xfun", quietly = TRUE)) {
    img_path <- xfun::relative_path(path, dir = getwd())
  }

  if (!is.null(caption) && nzchar(caption)) {
    cat("\n\n", "![", caption, "](", img_path, "){width=", width, "}", "\n\n", sep = "")
  } else {
    cat("\n\n", "![](", img_path, "){width=", width, "}", "\n\n", sep = "")
  }

  invisible(img_path)
}

show_table_head <- function(path, n = 10, caption = NULL) {
  if (!file.exists(path)) {
    cat("\n\nMissing table file: `", path, "`\n\n", sep = "")
    return(invisible(NULL))
  }

  if (!is.null(caption)) {
    cat("\n\n**", caption, "**\n\n", sep = "")
  }

  ext <- tolower(tools::file_ext(path))

  if (ext == "csv") {
    x <- read.csv(path, stringsAsFactors = FALSE, check.names = FALSE)
  } else {
    x <- read.delim(path, stringsAsFactors = FALSE, check.names = FALSE)
  }

  print(knitr::kable(head(x, n), format = "pipe"))
  invisible(x)
}
```

#### 2. Read or create metadata

This analysis uses the same sample set as the GDDH analysis:

- C01 to C08
- RIN01 to RIN08
- ROL01 to ROL05

If Trinity/metadata/samples_Trinity.tsv exists, it is used. Otherwise, the metadata table is created automatically.

```
if (file.exists(metadata_file)) {
  samples <- read.delim(metadata_file, stringsAsFactors = FALSE, check.names =
FALSE)
  samples[] <- lapply(samples, function(x) if (is.character(x)) trimws(x) else x)

  if (!"sample" %in% colnames(samples) && "sample_id" %in%
colnames(samples)) {
    samples$sample <- samples$sample_id
  }
} else {
  samples <- data.frame(
    sample = c(
      sprintf("C%02d", 1:8),
      sprintf("RIN%02d", 1:8),
      sprintf("ROL%02d", 1:5)
    ),
    condition = c(
      rep("control", 8),
      rep("Rin", 8),
      rep("Rola", 5)
    ),
    stringsAsFactors = FALSE
  )

  dir.create(dirname(metadata_file), recursive = TRUE, showWarnings = FALSE)
  write.table(samples, metadata_file, sep = "\t", quote = FALSE, row.names =
FALSE)
  message("Metadata file did not exist, so it was created: ", metadata_file)
}

if (!"sample" %in% colnames(samples)) {
  stop("Metadata must contain column 'sample' or 'sample_id'.")
}

if (!"condition" %in% colnames(samples)) {
  stop("Metadata must contain column 'condition'.")
}

samples$condition[samples$condition == "Rol"] <- "Rola"
samples$condition[samples$condition == "C"] <- "control"
samples$condition[samples$condition == "Control"] <- "control"
```

```r
samples$condition <- factor(samples$condition, levels = c("control", "Rola",
"Rin"))
samples <- droplevels(samples)

if (any(is.na(samples$condition))) {
  stop(
    "Some samples have condition values outside expected levels control/Rola/Rin."
  )
}

# Attach the library-preparation batch, used only by the sensitivity analysis in
# section 16. The main model remains ~ condition.
normalise_id <- function(x) {
  out <- toupper(trimws(x))
  for (p in c("RIN", "ROL", "C")) {
    hit <- startsWith(out, p)
    num <- suppressWarnings(as.integer(sub(paste0("^", p), "", out[hit])))
    out[hit] <- ifelse(is.na(num), out[hit], paste0(p, num))
  }
  out
}

# Inline fallback, a verbatim copy of metadata/samples_adjusted.tsv
# (created 5 Jun 2026; CORRECTED 12 Aug 2026 — ROL3 and ROL4 were
recorded as IAB
#  but their reads come from Eurofins originals, confirmed by read-ID matching
and
#  by file size, so Rola is 3 Eurofins / 2 IAB, not 1 / 4).
# It is embedded so section 16 runs on any machine, including one that holds the
counts but
# not that metadata file. The external file wins whenever it is present; this copy is
only
# used when it is not, and the report states which source was used. C9 and C10
belong to a
# different sample set and ROL6-ROL8 failed during library preparation; none of
them appear
# in the count matrix, so they are dropped automatically by the match below.
batch_meta_inline <- data.frame(
  sample_id = c(
    "C1", "C2", "C3", "C4", "C5", "C6", "C7", "C8", "C9", "C10", "RIN1", "RIN2",
"RIN3",
    "RIN4", "RIN5", "RIN6", "RIN7", "RIN8", "ROL1", "ROL2", "ROL3", "ROL4",
"ROL5", "ROL6",
    "ROL7", "ROL8"
  ),
  Sequencing = c(
    "IAB", "IAB", "IAB", "IAB", "EF", "EF", "EF", "EF", "EF", "EF", "IAB", "IAB",
"IAB",
    "IAB", "EF", "EF", "EF", "EF", "IAB", "IAB", "EF", "EF", "EF", "EF", "EF", "EF"
  ),
```

```r
  stringsAsFactors = FALSE
)

samples$batch <- NA_character_
batch_meta <- NULL
batch_source <- NA_character_
if (file.exists(batch_metadata_file)) {
  cand <- read.delim(batch_metadata_file, stringsAsFactors = FALSE,
check.names = FALSE)
  if (all(c("sample_id", "Sequencing") %in% colnames(cand))) {
    batch_meta <- cand
    batch_source <- paste("external file:", batch_metadata_file)
  } else {
    warning("Batch metadata lacks sample_id/Sequencing columns, using the inline
copy: ",
            batch_metadata_file)
  }
}
if (is.null(batch_meta)) {
  batch_meta <- batch_meta_inline
  batch_source <- "inline copy embedded in this notebook"
}
cat("Library-preparation batch taken from", batch_source, "\n")
```

```
## Library-preparation batch taken from external file:
/Volumes/Data_Disk/Ametyst_re-analysis/metadata/samples_adjusted.tsv
```

```r
idx <- match(normalise_id(samples$sample),
normalise_id(batch_meta$sample_id))
provider <- batch_meta$Sequencing[idx]
samples$batch <- dplyr::case_when(
  provider == "EF"  ~ "batch1_Eurofins",
  provider == "IAB" ~ "batch2_IAB",
  TRUE ~ NA_character_
)

# If both sources are available, they must agree, or the provenance is ambiguous.
if (identical(batch_source, paste("external file:", batch_metadata_file))) {
  ref <- batch_meta_inline$Sequencing[
    match(normalise_id(samples$sample),
normalise_id(batch_meta_inline$sample_id))]
  if (!isTRUE(all.equal(provider, ref))) {
    warning("The external batch metadata disagrees with the inline copy in this
notebook. ",
            "Resolve this before reporting the sensitivity analysis.")
  } else {
    cat("External file agrees with the inline copy for all", length(provider),
"libraries.\n")
  }
}
```

```r
write.csv(samples, file.path(outdir, "metadata_used_for_DESeq2.csv"), row.names
= FALSE)

cat("\nSamples used:\n")

##
## Samples used:

print(samples)

##    sample condition          batch
## 1     C01   control     batch2_IAB
## 2     C02   control     batch2_IAB
## 3     C03   control     batch2_IAB
## 4     C04   control     batch2_IAB
## 5     C05   control batch1_Eurofins
## 6     C06   control batch1_Eurofins
## 7     C07   control batch1_Eurofins
## 8     C08   control batch1_Eurofins
## 9   RIN01       Rin     batch2_IAB
## 10  RIN02       Rin     batch2_IAB
## 11  RIN03       Rin     batch2_IAB
## 12  RIN04       Rin     batch2_IAB
## 13  RIN05       Rin batch1_Eurofins
## 14  RIN06       Rin batch1_Eurofins
## 15  RIN07       Rin batch1_Eurofins
## 16  RIN08       Rin batch1_Eurofins
## 17  ROL01      Rola     batch2_IAB
## 18  ROL02      Rola     batch2_IAB
## 19  ROL03      Rola batch1_Eurofins
## 20  ROL04      Rola batch1_Eurofins
## 21  ROL05      Rola batch1_Eurofins

cat("\nCondition table:\n")

##
## Condition table:

print(table(samples$condition))

##
## control    Rola     Rin
##       8       5       8

mm <- model.matrix(~ condition, data = samples)
cat("\nDesign matrix rank check:\n")

##
## Design matrix rank check:

cat("rank =", qr(mm)$rank, "\n")
```

```
## rank = 3
```

```r
cat("columns =", ncol(mm), "\n")
```

```
## columns = 3
```

```r
if (qr(mm)$rank < ncol(mm)) {
  stop("Design matrix is not full rank for ~ condition.")
}
```

### 3. Import Salmon quantification with tximport

This section imports Salmon quant.sf files and creates gene-level counts for DESeq2. Because the Trinity FASTA contains one representative sequence per Trinity gene, the transcript-to-gene table is one-to-one.

```r
if (!dir.exists(salmon_quant_dir)) {
  stop(
    "Salmon quantification directory does not exist: ", salmon_quant_dir,
    "\nEdit salmon_quant_dir in chunk 0."
  )
}

if (!file.exists(trinity_fasta)) {
  stop("Trinity representative FASTA does not exist: ", trinity_fasta)
}

trinity_ids <- read_fasta_ids(trinity_fasta)

tx2gene <- data.frame(
  TXNAME = trinity_ids,
  GENEID = trinity_ids,
  stringsAsFactors = FALSE
)

write.table(
  tx2gene,
  file.path(outdir, "tx2gene_one_to_one_Trinity_gene_representatives.tsv"),
  sep = "\t",
  quote = FALSE,
  row.names = FALSE
)

sample_to_quant_path <- function(sample, quant_dir) {
  candidates <- unique(c(
    sample,
    sub("^C0", "C", sample),
```

```
    sub("^RIN0", "RIN", sample),
    sub("^ROL0", "ROL", sample),
    ifelse(grepl("^C[0-9]$", sample), sub("^C", "C0", sample), sample),
    ifelse(grepl("^RIN[0-9]$", sample), sub("^RIN", "RIN0", sample), sample),
    ifelse(grepl("^ROL[0-9]$", sample), sub("^ROL", "ROL0", sample), sample)
  ))

  paths <- file.path(quant_dir, candidates, "quant.sf")
  existing <- paths[file.exists(paths)]

  if (length(existing) == 0) {
    return(NA_character_)
  }

  existing[1]
}

quant_files <- vapply(samples$sample, sample_to_quant_path, character(1),
quant_dir = salmon_quant_dir)
names(quant_files) <- samples$sample

missing_quant <- names(quant_files)[is.na(quant_files)]
if (length(missing_quant) > 0) {
  stop(
    "Missing Salmon quant.sf files for samples: ", paste(missing_quant, collapse = ",
"),
    "\nChecked under: ", salmon_quant_dir,
    "\nExpected subdirectories such as C01 or C1, RIN01 or RIN1, ROL01 or
ROL1."
  )
}

cat("\nSalmon quant.sf files used:\n")

##
## Salmon quant.sf files used:

print(quant_files)

##                                                                     C01
##   "/Volumes/Data_Disk/Ametyst_re-analysis/Trinity/Salmon/quant/C01/quant.sf"
##                                                                     C02
##   "/Volumes/Data_Disk/Ametyst_re-analysis/Trinity/Salmon/quant/C02/quant.sf"
##                                                                     C03
##   "/Volumes/Data_Disk/Ametyst_re-analysis/Trinity/Salmon/quant/C03/quant.sf"
##                                                                     C04
##   "/Volumes/Data_Disk/Ametyst_re-analysis/Trinity/Salmon/quant/C04/quant.sf"
##                                                                     C05
##   "/Volumes/Data_Disk/Ametyst_re-analysis/Trinity/Salmon/quant/C05/quant.sf"
##                                                                     C06
```

```
##   "/Volumes/Data_Disk/Ametyst_re-analysis/Trinity/Salmon/quant/C06/quant.sf"
##                                                                      C07
##   "/Volumes/Data_Disk/Ametyst_re-analysis/Trinity/Salmon/quant/C07/quant.sf"
##                                                                      C08
##   "/Volumes/Data_Disk/Ametyst_re-analysis/Trinity/Salmon/quant/C08/quant.sf"
##                                                                    RIN01
## "/Volumes/Data_Disk/Ametyst_re-analysis/Trinity/Salmon/quant/RIN01/
quant.sf"
##                                                                    RIN02
## "/Volumes/Data_Disk/Ametyst_re-analysis/Trinity/Salmon/quant/RIN02/
quant.sf"
##                                                                    RIN03
## "/Volumes/Data_Disk/Ametyst_re-analysis/Trinity/Salmon/quant/RIN03/
quant.sf"
##                                                                    RIN04
## "/Volumes/Data_Disk/Ametyst_re-analysis/Trinity/Salmon/quant/RIN04/
quant.sf"
##                                                                    RIN05
## "/Volumes/Data_Disk/Ametyst_re-analysis/Trinity/Salmon/quant/RIN05/
quant.sf"
##                                                                    RIN06
## "/Volumes/Data_Disk/Ametyst_re-analysis/Trinity/Salmon/quant/RIN06/
quant.sf"
##                                                                    RIN07
## "/Volumes/Data_Disk/Ametyst_re-analysis/Trinity/Salmon/quant/RIN07/
quant.sf"
##                                                                    RIN08
## "/Volumes/Data_Disk/Ametyst_re-analysis/Trinity/Salmon/quant/RIN08/
quant.sf"
##                                                                    ROL01
## "/Volumes/Data_Disk/Ametyst_re-analysis/Trinity/Salmon/quant/ROL01/
quant.sf"
##                                                                    ROL02
## "/Volumes/Data_Disk/Ametyst_re-analysis/Trinity/Salmon/quant/ROL02/
quant.sf"
##                                                                    ROL03
## "/Volumes/Data_Disk/Ametyst_re-analysis/Trinity/Salmon/quant/ROL03/
quant.sf"
##                                                                    ROL04
## "/Volumes/Data_Disk/Ametyst_re-analysis/Trinity/Salmon/quant/ROL04/
quant.sf"
##                                                                    ROL05
## "/Volumes/Data_Disk/Ametyst_re-analysis/Trinity/Salmon/quant/ROL05/
quant.sf"
```

```
cat("\nImporting Salmon quantification with tximport.\n")
```

```
##
## Importing Salmon quantification with tximport.
```

```r
txi <- tximport(
  files = quant_files,
  type = "salmon",
  tx2gene = tx2gene,
  ignoreTxVersion = FALSE,
  countsFromAbundance = "lengthScaledTPM",
  importer = function(x) read.delim(x, stringsAsFactors = FALSE, check.names =
FALSE)
)
```

```
## 1 2 3 4 5 6 7 8 9 10 11 12 13 14 15 16 17 18 19 20 21
## summarizing abundance
## summarizing counts
## summarizing length
```

```r
cat("\nImported gene-level count matrix dimensions:\n")
```

```
##
## Imported gene-level count matrix dimensions:
```

```r
print(dim(txi$counts))
```

```
## [1] 249983     21
```

### 4. Create/read OmicsBox gene annotation

```r
if (!file.exists(annotation_file)) {
  gene_annot <- create_gene_annotation_from_omicsbox_gff(
    gff_file = gff_file,
    annotation_file = annotation_file
  )
} else {
  gene_annot <- read.delim(
    annotation_file,
    stringsAsFactors = FALSE,
    check.names = FALSE
  )
}

required_annot_cols <- c("gene_id", "gene_name", "Description")
missing_annot_cols <- setdiff(required_annot_cols, colnames(gene_annot))

if (length(missing_annot_cols) > 0) {
  stop(
    "Annotation file is missing required columns: ",
    paste(missing_annot_cols, collapse = ", "),
    "\nDelete the old annotation file and rerun:\n",
    annotation_file
  )
}
```

```r
# Add placeholder annotations for genes present in tximport but missing from
OmicsBox GFF.
missing_from_annotation <- setdiff(rownames(txi$counts), gene_annot$gene_id)
if (length(missing_from_annotation) > 0) {
  message("Genes missing from OmicsBox annotation: ",
length(missing_from_annotation))
  gene_annot <- bind_rows(
    gene_annot,
    data.frame(
      gene_id = missing_from_annotation,
      gene_name = missing_from_annotation,
      Description = missing_from_annotation,
      chr = NA_character_,
      start = NA_integer_,
      end = NA_integer_,
      strand = ".",
      Ontology_term = NA_character_,
      Ontology_id = NA_character_,
      Enzyme_code = NA_character_,
      Enzyme_name = NA_character_,
      stringsAsFactors = FALSE
    )
  )
}

gene_annot <- gene_annot |>
  dplyr::distinct(gene_id, .keep_all = TRUE)

save_result <- function(res, filename) {
  save_deseq_result(res, filename, gene_annot, outdir)
}
```

### 5. DESeq2 model

```r
dds <- DESeqDataSetFromTximport(
  txi = txi,
  colData = samples,
  design = ~ condition
)

## using just counts from tximport

dds <- dds[rowSums(counts(dds)) >= min_total_count, ]
dds <- DESeq(dds)

## estimating size factors

## estimating dispersions
```

```
## gene-wise dispersion estimates

## mean-dispersion relationship

## final dispersion estimates

## fitting model and testing

## -- replacing outliers and refitting for 133 genes
## -- DESeq argument 'minReplicatesForReplace' = 7
## -- original counts are preserved in counts(dds)

## estimating dispersions

## fitting model and testing

saveRDS(dds, file.path(outdir, "dds_Trinity_Salmon_condition_only.rds"))
writeLines(resultsNames(dds), file.path(outdir, "DESeq2_results_names.txt"))
```

### 6. DESeq2 contrasts and result tables

```
res_Rin_vs_Rola <- results(
  dds,
  contrast = c("condition", "Rin", "Rola"),
  alpha = candidate_padj_cutoff
)

res_Rin_vs_control <- results(
  dds,
  contrast = c("condition", "Rin", "control"),
  alpha = candidate_padj_cutoff
)

res_Rola_vs_control <- results(
  dds,
  contrast = c("condition", "Rola", "control"),
  alpha = candidate_padj_cutoff
)

df_Rin_vs_Rola <- save_deseq_result(
  res_Rin_vs_Rola,
  "DESeq2_Trinity_condition_only_Rin_vs_Rola_with_description.csv",
  gene_annot,
  outdir
)

df_Rin_vs_control <- save_deseq_result(
  res_Rin_vs_control,
  "DESeq2_Trinity_condition_only_Rin_vs_control_with_description.csv",
  gene_annot,
  outdir
```

```
)

df_Rola_vs_control <- save_deseq_result(
  res_Rola_vs_control,
  "DESeq2_Trinity_condition_only_Rola_vs_control_with_description.csv",
  gene_annot,
  outdir
)

write.csv(df_Rin_vs_Rola, file.path(outdir,
"DESeq2_Trinity_condition_only_Rin_vs_Rola.csv"), row.names = FALSE)
write.csv(df_Rin_vs_control, file.path(outdir,
"DESeq2_Trinity_condition_only_Rin_vs_control.csv"), row.names = FALSE)
write.csv(df_Rola_vs_control, file.path(outdir,
"DESeq2_Trinity_condition_only_Rola_vs_control.csv"), row.names = FALSE)

summary_table <- data.frame(
  contrast = c("Rin_vs_Rola", "Rin_vs_control", "Rola_vs_control"),
  padj_0.05 = c(
    sum(res_Rin_vs_Rola$padj < 0.05, na.rm = TRUE),
    sum(res_Rin_vs_control$padj < 0.05, na.rm = TRUE),
    sum(res_Rola_vs_control$padj < 0.05, na.rm = TRUE)
  ),
  padj_0.05_absLFC1 = c(
    sum(res_Rin_vs_Rola$padj < 0.05 & abs(res_Rin_vs_Rola$log2FoldChange) >=
1, na.rm = TRUE),
    sum(res_Rin_vs_control$padj < 0.05 &
abs(res_Rin_vs_control$log2FoldChange) >= 1, na.rm = TRUE),
    sum(res_Rola_vs_control$padj < 0.05 &
abs(res_Rola_vs_control$log2FoldChange) >= 1, na.rm = TRUE)
  )
)

write.csv(summary_table, file.path(outdir,
"DESeq2_Trinity_condition_only_summary_counts.csv"), row.names = FALSE)
print(summary_table)
```

```
##          contrast padj_0.05 padj_0.05_absLFC1
## 1     Rin_vs_Rola     18121             13382
## 2  Rin_vs_control     20107             14939
## 3 Rola_vs_control     15987             12860
```

### 7. Normalized/transformed matrices

```
raw_counts <- counts(dds, normalized = FALSE)
norm_counts <- counts(dds, normalized = TRUE)
log2_raw_counts <- log2(raw_counts + 1)
log2_norm_counts <- log2(norm_counts + 1)
```

```r
vsd <- vst(dds, blind = FALSE)
vsd_mat <- SummarizedExperiment::assay(vsd)

if (run_rlog) {
  rld <- rlog(dds, blind = FALSE)
  rlog_mat <- SummarizedExperiment::assay(rld)
} else {
  rld <- NULL
  rlog_mat <- NULL
  message("run_rlog is FALSE: skipping rlog transformation and rlog-specific
plots.")
}
```

```
## run_rlog is FALSE: skipping rlog transformation and rlog-specific plots.
```

```r
write.csv(as.data.frame(norm_counts), file.path(outdir,
"DESeq2_Trinity_condition_only_normalized_counts.csv"))
write.csv(as.data.frame(vsd_mat), file.path(qcdir, "VST_expression_matrix.csv"))
if (!is.null(rlog_mat)) {
  write.csv(as.data.frame(rlog_mat), file.path(qcdir,
"rlog_expression_matrix.csv"))
}

ann <- as.data.frame(colData(dds)[, c("condition"), drop = FALSE])
rownames(ann) <- colnames(vsd_mat)
```

### 8. PCA and sample distance QC

```r
write_png(file.path(outdir,
"PCA_Trinity_condition_only_raw_vst_by_condition.png"), 7, 6, {
  print(plotPCA(vsd, intgroup = "condition"))
})
```

```
## using ntop=500 top features by variance
```

```r
sampleDistMatrix <- as.matrix(dist(t(vsd_mat)))
rownames(sampleDistMatrix) <- samples$sample
colnames(sampleDistMatrix) <- samples$sample

write_png(file.path(outdir,
"Sample_distance_heatmap_Trinity_condition_only_raw_vst.png"), 9, 8, {
  pheatmap(sampleDistMatrix, annotation_col = ann, annotation_row = ann)
})

pca <- prcomp(t(vsd_mat))
percentVar <- round(100 * (pca$sdev^2 / sum(pca$sdev^2)), 1)

n_pc <- min(5, ncol(pca$x))
pca_df <- as.data.frame(pca$x[, seq_len(n_pc), drop = FALSE])
pca_df$sample <- rownames(pca_df)
```

```r
pca_df$condition <- samples$condition
write.csv(pca_df, file.path(outdir, "PCA_scores_condition_only.csv"), row.names =
FALSE)

for (grp in c("condition")) {
  write_png(file.path(outdir, paste0("PCA_Trinity_condition_only_by_", grp,
".png")), 7, 6, {
    print(
      ggplot(pca_df, aes(.data[["PC1"]], .data[["PC2"]], color = .data[[grp]], label =
sample)) +
        geom_point(size = 3) +
        geom_text(size = 2.5, vjust = -0.6, show.legend = FALSE) +
        xlab(paste0("PC1: ", percentVar[1], "% variance")) +
        ylab(paste0("PC2: ", percentVar[2], "% variance")) +
        coord_fixed() +
        theme_bw()
    )
  })
}
```

### 9. Extra QC: boxplots, mean-SD, correlations

```r
write_png(file.path(qcdir, "Boxplot_raw_counts.png"), 11, 6, {
  boxplot(raw_counts, las = 2, outline = FALSE, main = "Raw Trinity gene counts",
ylab = "Raw counts")
})

write_png(file.path(qcdir, "Boxplot_log2_raw_counts_plus1.png"), 11, 6, {
  boxplot(log2_raw_counts, las = 2, outline = FALSE, main = "log2(raw counts +
1)", ylab = "log2(raw counts + 1)")
})

write_png(file.path(qcdir, "Boxplot_normalized_counts.png"), 11, 6, {
  boxplot(norm_counts, las = 2, outline = FALSE, main = "DESeq2 normalized
counts", ylab = "Normalized counts")
})

write_png(file.path(qcdir, "Boxplot_log2_normalized_counts_plus1.png"), 11, 6, {
  boxplot(log2_norm_counts, las = 2, outline = FALSE, main = "log2(DESeq2
normalized counts + 1)", ylab = "log2(normalized counts + 1)")
})

write_png(file.path(qcdir, "Boxplot_VST_counts.png"), 11, 6, {
  boxplot(vsd_mat, las = 2, outline = FALSE, main = "Variance-stabilized counts",
ylab = "VST expression")
})

if (!is.null(rlog_mat)) {
  write_png(file.path(qcdir, "Boxplot_rlog_counts.png"), 11, 6, {
```

```r
    boxplot(rlog_mat, las = 2, outline = FALSE, main = "rlog-transformed counts",
ylab = "rlog expression")
  })
}

msd <- bind_rows(
  mean_sd_df(log2_raw_counts, "log2 raw counts + 1"),
  mean_sd_df(log2_norm_counts, "log2 normalized counts + 1"),
  mean_sd_df(vsd_mat, "VST")
)

if (!is.null(rlog_mat)) {
  msd <- bind_rows(msd, mean_sd_df(rlog_mat, "rlog"))
}

write.csv(msd, file.path(qcdir, "Mean_SD_by_transformation.csv"), row.names =
FALSE)

# geom_smooth() is deliberately not used here. ggplot2 resolves the smoother's
# default method through mgcv in StatSmooth$setup_params even when "loess" is
named
# explicitly, and loading mgcv segfaulted during the knit of 2 Sep 2026
# (dyn.load, cause 'memory not mapped') with the Trinity object in memory.
# stat_summary_bin is core ggplot2, loads nothing extra, and shows the same
trend.
# The scatter is drawn from a sample when the gene set is large - 119,509 genes
over
# three transformations is 358,527 overplotted points - while the median line is
# always computed on every gene.
msd_points <- msd
n_per_facet <- 40000
if (nrow(msd_points) > 3 * n_per_facet) {
  set.seed(1)
  keep <- unlist(lapply(
    split(seq_len(nrow(msd_points)), msd_points$transformation),
    function(ix) if (length(ix) > n_per_facet) sample(ix, n_per_facet) else ix
  ))
  msd_points <- msd_points[sort(keep), ]
  message("Mean-SD scatter drawn from ", n_per_facet,
          " genes per transformation; the median line uses all genes.")
}

## Mean-SD scatter drawn from 40000 genes per transformation; the median line
uses all genes.

write_png(file.path(qcdir, "Heteroskedasticity_mean_SD_plots.png"), 10, 8, {
  print(
    ggplot(msd_points, aes(x = mean, y = sd)) +
      geom_point(alpha = 0.15, size = 0.4) +
      stat_summary_bin(data = msd, fun = median, geom = "line", bins = 100,
```

```
                 colour = "#2166ac", linewidth = 0.8, na.rm = TRUE) +
    facet_wrap(~ transformation, scales = "free") +
    theme_bw() +
    xlab("Gene-wise mean expression") +
    ylab("Gene-wise standard deviation") +
    ggtitle("Mean-SD relationship / heteroskedasticity check")
  )
})

if (has_vsn) {
  write_png(file.path(qcdir, "vsn_meanSdPlot_log2_normalized_counts.png"), 7, 6,
{
    vsn::meanSdPlot(log2_norm_counts, ranks = FALSE)
  })
  write_png(file.path(qcdir, "vsn_meanSdPlot_VST.png"), 7, 6, {
    vsn::meanSdPlot(vsd_mat, ranks = FALSE)
  })
  if (!is.null(rlog_mat)) {
    write_png(file.path(qcdir, "vsn_meanSdPlot_rlog.png"), 7, 6, {
      vsn::meanSdPlot(rlog_mat, ranks = FALSE)
    })
  }
} else {
  message("Package 'vsn' not installed: skipping vsn::meanSdPlot outputs.")
}

## Package 'vsn' not installed: skipping vsn::meanSdPlot outputs.

pearson_cor_vst <- cor(vsd_mat, method = "pearson")
write.csv(pearson_cor_vst, file.path(qcdir,
"Pearson_correlation_VST_matrix.csv"))

write_png(file.path(qcdir, "Pearson_correlation_heatmap_VST.png"), 9, 8, {
  pheatmap(pearson_cor_vst, annotation_col = ann, annotation_row = ann, main =
"Pearson correlation: VST")
})

if (!is.null(rlog_mat)) {
  pearson_cor_rlog <- cor(rlog_mat, method = "pearson")
  write.csv(pearson_cor_rlog, file.path(qcdir,
"Pearson_correlation_rlog_matrix.csv"))
  write_png(file.path(qcdir, "Pearson_correlation_heatmap_rlog.png"), 9, 8, {
    pheatmap(pearson_cor_rlog, annotation_col = ann, annotation_row = ann, main
= "Pearson correlation: rlog")
  })
}
```

### 10. MA and volcano plots for Rin vs Rola

```
write_png(file.path(qcdir, "MA_plot_Rin_vs_Rola_DESeq2.png"), 7, 6, {
  DESeq2::plotMA(res_Rin_vs_Rola, ylim = c(-8, 8), main = "MA plot: Rin vs Rola")
})

ma_df <- df_Rin_vs_Rola |>
  mutate(
    significant = !is.na(padj) & padj < candidate_padj_cutoff,
    baseMean_for_plot = ifelse(baseMean <= 0, NA_real_, baseMean)
  )

write.csv(ma_df, file.path(qcdir, "MA_input_Rin_vs_Rola_DESeq2.csv"),
row.names = FALSE)

write_png(file.path(qcdir, "MA_plot_Rin_vs_Rola_custom_ggplot.png"), 7, 6, {
  print(
    ggplot(ma_df, aes(x = baseMean_for_plot, y = log2FoldChange)) +
      geom_point(aes(color = significant), alpha = 0.5, size = 0.6, na.rm = TRUE) +
      scale_x_log10() +
      geom_hline(yintercept = 0, linetype = "dashed") +
      coord_cartesian(ylim = c(-8, 8)) +
      theme_bw() +
      xlab("Mean of normalized counts: baseMean, log10 scale") +
      ylab("log2 fold change: Rin vs Rola") +
      ggtitle("MA plot: Rin vs Rola")
  )
})

if (has_ashr) {
  res_shrunk <- lfcShrink(dds, contrast = c("condition", "Rin", "Rola"), type =
"ashr")
  write_png(file.path(qcdir, "MA_plot_Rin_vs_Rola_LFC_shrunk_ashr.png"), 7, 6, {
    DESeq2::plotMA(res_shrunk, ylim = c(-8, 8), main = "MA plot: Rin vs Rola, LFC
shrinkage (ashr)")
  })
} else {
  message("Package 'ashr' not installed: skipping LFC-shrunk MA plot for the Rin
vs Rola contrast.")
}

## Package 'ashr' not installed: skipping LFC-shrunk MA plot for the Rin vs Rola
contrast.

volcano_df <- df_Rin_vs_Rola |>
  mutate(
    direction = case_when(
      !is.na(padj) & padj < candidate_padj_cutoff & log2FoldChange >=
candidate_abs_lfc_cutoff ~ "Rin-high",
      !is.na(padj) & padj < candidate_padj_cutoff & log2FoldChange <=
```

```r
-candidate_abs_lfc_cutoff ~ "Rola-high",
      TRUE ~ "not significant"
    ),
    neg_log10_padj = -log10(padj),
    neg_log10_padj = ifelse(is.infinite(neg_log10_padj), NA_real_, neg_log10_padj)
  )

write.csv(volcano_df, file.path(qcdir, "Volcano_input_Rin_vs_Rola.csv"),
row.names = FALSE)

write_png(file.path(qcdir, "Volcano_plot_Rin_vs_Rola.png"), 7, 6, {
  print(
    ggplot(volcano_df, aes(x = log2FoldChange, y = neg_log10_padj, color =
direction)) +
      geom_point(alpha = 0.6, size = 0.8, na.rm = TRUE) +
      geom_vline(xintercept = c(-candidate_abs_lfc_cutoff,
candidate_abs_lfc_cutoff), linetype = "dashed") +
      geom_hline(yintercept = -log10(candidate_padj_cutoff), linetype = "dashed")
+
      theme_bw() +
      xlab("log2FC: Rin vs Rola") +
      ylab("-log10 adjusted P value") +
      ggtitle("Volcano plot: Rin vs Rola")
  )
})
```

### 11. Heatmaps

```r
top_n <- 500
gene_var_vst <- apply(vsd_mat, 1, var)
top_var_genes <- names(sort(gene_var_vst, decreasing = TRUE))
[seq_len(min(top_n, length(gene_var_vst)))]
mat_top_vst_scaled <- t(scale(t(vsd_mat[top_var_genes, , drop = FALSE])))

write_png(file.path(qcdir,
"Heatmap_top500_variable_genes_VST_pheatmap.png"), 9, 10, {
  pheatmap(
    mat_top_vst_scaled,
    annotation_col = ann,
    show_rownames = FALSE,
    main = "Top 500 variable Trinity genes, VST, row-scaled"
  )
})

sig_genes <- volcano_df |>
  filter(!is.na(padj), padj < candidate_padj_cutoff, abs(log2FoldChange) >=
candidate_abs_lfc_cutoff) |>
  arrange(padj) |>
  slice_head(n = 500) |>
```

```r
  pull(gene_id)

if (length(sig_genes) >= 2) {
  mat_sig_vst_scaled <- t(scale(t(vsd_mat[sig_genes, , drop = FALSE])))
  write_png(file.path(qcdir,
"Heatmap_Rin_vs_Rola_significant_genes_VST_pheatmap.png"), 9, 10, {
    pheatmap(
      mat_sig_vst_scaled,
      annotation_col = ann,
      show_rownames = FALSE,
      main = "Significant Rin vs Rola Trinity genes, VST, row-scaled"
    )
  })
} else {
  message("Fewer than 2 significant Rin vs Rola genes: skipping significant-gene
heatmap.")
}

rin_high <- df_Rin_vs_Rola |>
  filter(!is.na(padj), padj < candidate_padj_cutoff,
         log2FoldChange >= candidate_abs_lfc_cutoff,
         baseMean >= candidate_baseMean_cutoff) |>
  arrange(padj)

rola_high <- df_Rin_vs_Rola |>
  filter(!is.na(padj), padj < candidate_padj_cutoff,
         log2FoldChange <= -candidate_abs_lfc_cutoff,
         baseMean >= candidate_baseMean_cutoff) |>
  arrange(padj)

write.csv(rin_high, file.path(outdir,
"Candidate_genes_Rin_high_padj0.05_LFC1_baseMean50.csv"), row.names =
FALSE)
write.csv(rola_high, file.path(outdir,
"Candidate_genes_Rola_high_padj0.05_LFC1_baseMean50.csv"), row.names =
FALSE)

plot_candidate_heatmap(
  head(rin_high$gene_id, 50),
  "Heatmap_top50_Rin_high_genes_VST_row_scaled.png",
  "Top 50 Rin-high Trinity genes, VST, row-scaled",
  vsd_mat,
  ann,
  df_Rin_vs_Rola
)

plot_candidate_heatmap(
  head(rola_high$gene_id, 50),
  "Heatmap_top50_Rola_high_genes_VST_row_scaled.png",
```

```
  "Top 50 Rola-high Trinity genes, VST, row-scaled",
  vsd_mat,
  ann,
  df_Rin_vs_Rola
)

plot_candidate_heatmap(
  c(head(rin_high$gene_id, 25), head(rola_high$gene_id, 25)),
  "Heatmap_top25_Rin_high_top25_Rola_high_VST_row_scaled.png",
  "Top 25 Rin-high + top 25 Rola-high Trinity genes, VST, row-scaled",
  vsd_mat,
  ann,
  df_Rin_vs_Rola
)

if (has_NMF) {
  ann_colors <- data.frame(condition = as.character(colData(dds)$condition))
  rownames(ann_colors) <- colnames(vsd_mat)

  write_png(file.path(qcdir, "aheatmap_top500_variable_genes_VST_scaled.png"),
9, 10, {
    NMF::aheatmap(
      mat_top_vst_scaled,
      annCol = ann_colors,
      Rowv = TRUE,
      Colv = TRUE,
      distfun = "euclidean",
      hclustfun = "complete",
      labRow = NA,
      main = "aheatmap: top 500 variable Trinity genes, VST scaled"
    )
  })

  write_png(file.path(qcdir, "aheatmap_Pearson_correlation_VST.png"), 9, 8, {
    NMF::aheatmap(
      pearson_cor_vst,
      annCol = ann_colors,
      annRow = ann_colors,
      Rowv = TRUE,
      Colv = TRUE,
      distfun = function(x) as.dist(1 - x),
      hclustfun = "complete",
      main = "aheatmap: Pearson correlation, VST"
    )
  })

  if (exists("mat_sig_vst_scaled") && nrow(mat_sig_vst_scaled) >= 2) {
    write_png(file.path(qcdir,
"aheatmap_Rin_vs_Rola_significant_genes_VST_scaled.png"), 9, 10, {
      NMF::aheatmap(
```

```
      mat_sig_vst_scaled,
      annCol = ann_colors,
      Rowv = TRUE,
      Colv = TRUE,
      distfun = "euclidean",
      hclustfun = "complete",
      labRow = NA,
      main = "aheatmap: significant Rin vs Rola Trinity genes, VST scaled"
    )
  })
 }
} else {
  message("Package 'NMF' not installed: skipping NMF::aheatmap outputs.")
}
```

### 12. Session info

```
sink(file.path(outdir, "sessionInfo.txt"))
sessionInfo()
sink()

cat("\nMain analysis done. Results written to:\n")

##
## Main analysis done. Results written to:

cat(outdir, "\n")

## /Volumes/Data_Disk/Ametyst_re-analysis/Trinity/DESeq2

cat("\nExtra QC plots written to:\n")

##
## Extra QC plots written to:

cat(qcdir, "\n")

## /Volumes/Data_Disk/Ametyst_re-analysis/Trinity/DESeq2/QC_extra_plots
```

### 13. Extra selection: Rin vs Rola, Rin vs control, Rola vs control

```
res_Rin_vs_Rola_LFC1 <- results(
  dds,
  contrast = c("condition", "Rin", "Rola"),
  alpha = 0.01,
  lfcThreshold = 1,
  altHypothesis = "greaterAbs"
)
```

```r
df_Rin_vs_Rola_LFC1 <- save_result(
  res_Rin_vs_Rola_LFC1,
  "DESeq2_Trinity_condition_only_Rin_vs_Rola_lfcThreshold1_with_description.csv"
)

df_Rin_vs_Rola_LFC1_sorted_by_LFC <- df_Rin_vs_Rola_LFC1 |>
  dplyr::arrange(dplyr::desc(log2FoldChange))

write.csv(
  df_Rin_vs_Rola_LFC1_sorted_by_LFC,
  file.path(outdir,
"DESeq2_Rin_vs_Rola_lfcThreshold1_padj0.01_with_description_by_log2FC.csv"),
  row.names = FALSE
)

rin_vs_rola_Rin_high <- df_Rin_vs_Rola_LFC1 |>
  dplyr::filter(!is.na(padj), padj < 0.01, log2FoldChange > 1) |>
  dplyr::arrange(dplyr::desc(log2FoldChange))

rin_vs_rola_Rola_high <- df_Rin_vs_Rola_LFC1 |>
  dplyr::filter(!is.na(padj), padj < 0.01, log2FoldChange < -1) |>
  dplyr::arrange(log2FoldChange)

write.csv(
  rin_vs_rola_Rin_high,
  file.path(outdir,
"Candidate_Rin_vs_Rola_Rin_high_lfcThreshold1_padj0.01_by_log2FC.csv"),
  row.names = FALSE
)

write.csv(
  rin_vs_rola_Rola_high,
  file.path(outdir,
"Candidate_Rin_vs_Rola_Rola_high_lfcThreshold1_padj0.01_by_log2FC.csv"),
  row.names = FALSE
)

candidate_Rin_vs_Rola_LFC1_by_absLFC <- df_Rin_vs_Rola_LFC1 |>
  dplyr::filter(!is.na(padj), padj < 0.01, abs(log2FoldChange) > 1) |>
  dplyr::mutate(
    direction = dplyr::case_when(
      log2FoldChange > 1  ~ "Rin-high",
      log2FoldChange < -1 ~ "Rola-high",
      TRUE ~ "none"
    )
  ) |>
  dplyr::select(
    gene_id, gene_name, baseMean, log2FoldChange, lfcSE, stat, pvalue, padj,
```

```
direction, Description,
    dplyr::any_of(c("Ontology_term", "Ontology_id", "Enzyme_code",
"Enzyme_name"))
  ) |>
  dplyr::arrange(dplyr::desc(abs(log2FoldChange)))

write.csv(
  candidate_Rin_vs_Rola_LFC1_by_absLFC,
  file.path(outdir,
"Candidate_Rin_vs_Rola_lfcThreshold1_padj0.01_by_abs_log2FC.csv"),
  row.names = FALSE
)

res_Rin_vs_C_LFC1 <- results(
  dds,
  contrast = c("condition", "Rin", "control"),
  alpha = 0.01,
  lfcThreshold = 1,
  altHypothesis = "greaterAbs"
)

df_Rin_vs_C_LFC1 <- save_result(
  res_Rin_vs_C_LFC1,
  "DESeq2_Trinity_condition_only_Rin_vs_C_lfcThreshold1_with_description.csv"
)

df_Rin_vs_C_LFC1_sorted_by_LFC <- df_Rin_vs_C_LFC1 |>
  dplyr::arrange(dplyr::desc(log2FoldChange))

write.csv(
  df_Rin_vs_C_LFC1_sorted_by_LFC,
  file.path(outdir,
"DESeq2_Rin_vs_C_lfcThreshold1_padj0.01_with_description_by_log2FC.csv"),
  row.names = FALSE
)

rin_vs_C_Rin_high <- df_Rin_vs_C_LFC1 |>
  dplyr::filter(!is.na(padj), padj < 0.01, log2FoldChange > 1) |>
  dplyr::arrange(dplyr::desc(log2FoldChange))

rin_vs_C_C_high <- df_Rin_vs_C_LFC1 |>
  dplyr::filter(!is.na(padj), padj < 0.01, log2FoldChange < -1) |>
  dplyr::arrange(log2FoldChange)

write.csv(
  rin_vs_C_Rin_high,
  file.path(outdir,
"Candidate_Rin_vs_C_Rin_high_lfcThreshold1_padj0.01_by_log2FC.csv"),
  row.names = FALSE
```

```
)

write.csv(
  rin_vs_C_C_high,
  file.path(outdir,
"Candidate_Rin_vs_C_C_high_lfcThreshold1_padj0.01_by_log2FC.csv"),
  row.names = FALSE
)

candidate_Rin_vs_C_LFC1_by_absLFC <- df_Rin_vs_C_LFC1 |>
  dplyr::filter(!is.na(padj), padj < 0.01, abs(log2FoldChange) > 1) |>
  dplyr::mutate(
    direction = dplyr::case_when(
      log2FoldChange > 1  ~ "Rin-high",
      log2FoldChange < -1 ~ "C-high",
      TRUE ~ "none"
    )
  ) |>
  dplyr::select(
    gene_id, gene_name, baseMean, log2FoldChange, lfcSE, stat, pvalue, padj,
direction, Description,
    dplyr::any_of(c("Ontology_term", "Ontology_id", "Enzyme_code",
"Enzyme_name"))
  ) |>
  dplyr::arrange(dplyr::desc(abs(log2FoldChange)))

write.csv(
  candidate_Rin_vs_C_LFC1_by_absLFC,
  file.path(outdir,
"Candidate_Rin_vs_C_lfcThreshold1_padj0.01_by_abs_log2FC.csv"),
  row.names = FALSE
)

res_Rola_vs_C_LFC1 <- results(
  dds,
  contrast = c("condition", "Rola", "control"),
  alpha = 0.01,
  lfcThreshold = 1,
  altHypothesis = "greaterAbs"
)

df_Rola_vs_C_LFC1 <- save_result(
  res_Rola_vs_C_LFC1,
  "DESeq2_Trinity_condition_only_Rola_vs_C_lfcThreshold1_with_description.csv"
)

df_Rola_vs_C_LFC1_sorted_by_LFC <- df_Rola_vs_C_LFC1 |>
  dplyr::arrange(dplyr::desc(log2FoldChange))
```

```r
write.csv(
  df_Rola_vs_C_LFC1_sorted_by_LFC,
  file.path(outdir,
"DESeq2_Rola_vs_C_lfcThreshold1_padj0.01_with_description_by_log2FC.csv"),
  row.names = FALSE
)

rola_vs_C_Rola_high <- df_Rola_vs_C_LFC1 |>
  dplyr::filter(!is.na(padj), padj < 0.01, log2FoldChange > 1) |>
  dplyr::arrange(dplyr::desc(log2FoldChange))

rola_vs_C_C_high <- df_Rola_vs_C_LFC1 |>
  dplyr::filter(!is.na(padj), padj < 0.01, log2FoldChange < -1) |>
  dplyr::arrange(log2FoldChange)

write.csv(
  rola_vs_C_Rola_high,
  file.path(outdir,
"Candidate_Rola_vs_C_Rola_high_lfcThreshold1_padj0.01_by_log2FC.csv"),
  row.names = FALSE
)

write.csv(
  rola_vs_C_C_high,
  file.path(outdir,
"Candidate_Rola_vs_C_C_high_lfcThreshold1_padj0.01_by_log2FC.csv"),
  row.names = FALSE
)

candidate_Rola_vs_C_LFC1_by_absLFC <- df_Rola_vs_C_LFC1 |>
  dplyr::filter(!is.na(padj), padj < 0.01, abs(log2FoldChange) > 1) |>
  dplyr::mutate(
    direction = dplyr::case_when(
      log2FoldChange > 1  ~ "Rola-high",
      log2FoldChange < -1 ~ "C-high",
      TRUE ~ "none"
    )
  ) |>
  dplyr::select(
    gene_id, gene_name, baseMean, log2FoldChange, lfcSE, stat, pvalue, padj,
direction, Description,
    dplyr::any_of(c("Ontology_term", "Ontology_id", "Enzyme_code",
"Enzyme_name"))
  ) |>
  dplyr::arrange(dplyr::desc(abs(log2FoldChange)))

write.csv(
  candidate_Rola_vs_C_LFC1_by_absLFC,
  file.path(outdir,
```

```
"Candidate_Rola_vs_C_lfcThreshold1_padj0.01_by_abs_log2FC.csv"),
  row.names = FALSE
)
```

### 14. Comparison table and classification

```
make_compact_result <- function(res, prefix) {
  df <- as.data.frame(res)
  df$gene_id <- rownames(df)

  df |>
    dplyr::select(
      gene_id, baseMean, log2FoldChange, lfcSE, stat, pvalue, padj
    ) |>
    dplyr::rename(
      !!paste0("baseMean_", prefix) := baseMean,
      !!paste0("log2FC_", prefix) := log2FoldChange,
      !!paste0("lfcSE_", prefix) := lfcSE,
      !!paste0("stat_", prefix) := stat,
      !!paste0("pvalue_", prefix) := pvalue,
      !!paste0("padj_", prefix) := padj
    )
}

rin_c <- make_compact_result(res_Rin_vs_C_LFC1, "Rin_vs_C")
rola_c <- make_compact_result(res_Rola_vs_C_LFC1, "Rola_vs_C")
rin_rola <- make_compact_result(res_Rin_vs_Rola_LFC1, "Rin_vs_Rola")

comparison <- gene_annot |>
  dplyr::select(
    gene_id,
    gene_name,
    Description,
    dplyr::any_of(c("Ontology_term", "Ontology_id", "Enzyme_code",
"Enzyme_name"))
  ) |>
  dplyr::left_join(rin_c, by = "gene_id") |>
  dplyr::left_join(rola_c, by = "gene_id") |>
  dplyr::left_join(rin_rola, by = "gene_id")

is_up <- function(lfc, padj) {
  !is.na(padj) & padj < 0.01 & !is.na(lfc) & lfc > 1
}

is_down <- function(lfc, padj) {
  !is.na(padj) & padj < 0.01 & !is.na(lfc) & lfc < -1
}

comparison <- comparison |>
```

```r
  dplyr::mutate(
    Rin_up_vs_C = is_up(log2FC_Rin_vs_C, padj_Rin_vs_C),
    Rin_down_vs_C = is_down(log2FC_Rin_vs_C, padj_Rin_vs_C),

    Rola_up_vs_C = is_up(log2FC_Rola_vs_C, padj_Rola_vs_C),
    Rola_down_vs_C = is_down(log2FC_Rola_vs_C, padj_Rola_vs_C),

    class = dplyr::case_when(
      Rin_up_vs_C & Rola_up_vs_C ~ "common_infection_induced",
      Rin_down_vs_C & Rola_down_vs_C ~ "common_infection_repressed",

      Rin_up_vs_C & !Rola_up_vs_C & !Rola_down_vs_C ~ "Rin_specific_induced",
      Rola_up_vs_C & !Rin_up_vs_C & !Rin_down_vs_C ~ "Rola_specific_induced",

      Rin_down_vs_C & !Rola_up_vs_C & !Rola_down_vs_C ~
"Rin_specific_repressed",
      Rola_down_vs_C & !Rin_up_vs_C & !Rin_down_vs_C ~
"Rola_specific_repressed",

      Rin_up_vs_C & Rola_down_vs_C ~ "opposite_Rin_up_Rola_down",
      Rin_down_vs_C & Rola_up_vs_C ~ "opposite_Rin_down_Rola_up",

      TRUE ~ "not_strong_DE"
    )
  )

write.csv(
  comparison,
  file.path(outdir,
"Comparison_Rin_vs_C_Rola_vs_C_Rin_vs_Rola_lfcThreshold1_padj0.01.csv"),
  row.names = FALSE
)

class_summary <- comparison |>
  dplyr::count(class, sort = TRUE)

write.csv(
  class_summary,
  file.path(outdir, "Comparison_class_summary_lfcThreshold1_padj0.01.csv"),
  row.names = FALSE
)

print(class_summary)

##                         class      n
## 1               not_strong_DE 242655
## 2       Rola_specific_induced   3352
## 3    common_infection_induced   2313
## 4        Rin_specific_induced    864
```

```
## 5     Rin_specific_repressed    728
## 6    Rola_specific_repressed     69
## 7 common_infection_repressed      2
```

### 15. Scatter plot: similarity of Rin and Rola vs control

```
plot_df <- comparison |>
  dplyr::filter(
    !is.na(log2FC_Rin_vs_C),
    !is.na(log2FC_Rola_vs_C)
  )

cor_pearson <- cor(
  plot_df$log2FC_Rin_vs_C,
  plot_df$log2FC_Rola_vs_C,
  method = "pearson",
  use = "complete.obs"
)

cor_spearman <- cor(
  plot_df$log2FC_Rin_vs_C,
  plot_df$log2FC_Rola_vs_C,
  method = "spearman",
  use = "complete.obs"
)

png(filename = file.path(outdir, "Scatter_log2FC_Rin_vs_C_vs_Rola_vs_C.png"),
width = 7, height = 7, units = "in", res = 150, type = "cairo-png")

print(
  ggplot(plot_df, aes(x = log2FC_Rin_vs_C, y = log2FC_Rola_vs_C)) +
    geom_point(alpha = 0.35, size = 0.7) +
    geom_hline(yintercept = 0, linetype = "dashed") +
    geom_vline(xintercept = 0, linetype = "dashed") +
    geom_abline(slope = 1, intercept = 0, linetype = "dotted") +
    coord_cartesian(xlim = c(-10, 10), ylim = c(-10, 10)) +
    theme_bw() +
    xlab("log2FC Rin vs control") +
    ylab("log2FC Rola vs control") +
    ggtitle(
      paste0(
        "Rin and Rola responses vs control\n",
        "Pearson r = ", round(cor_pearson, 3),
        ", Spearman rho = ", round(cor_spearman, 3)
      )
    )
)
```

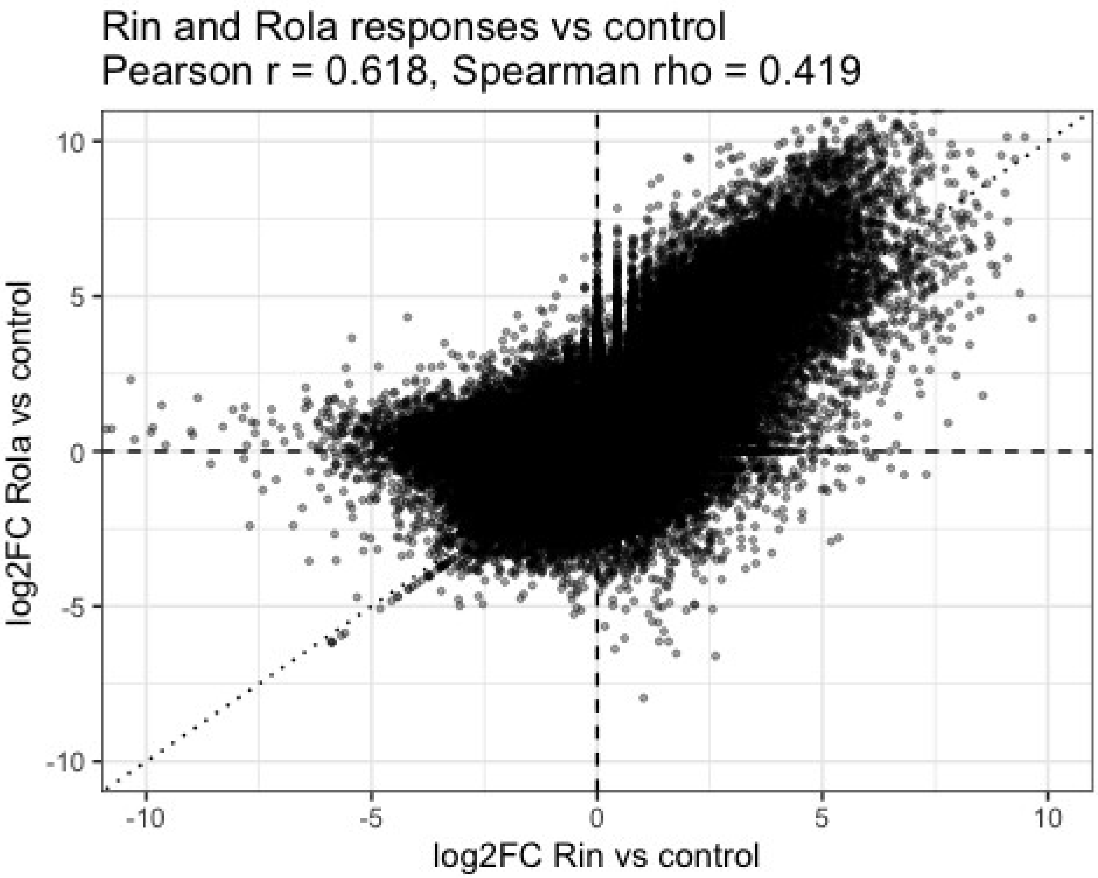


```r
dev.off()

## null device
##           1

cat("\nFull Trinity analysis completed successfully.\n")

##
## Full Trinity analysis completed successfully.
```

### 16. Batch-design sensitivity analysis

Libraries were prepared in two batches about six months apart and sequenced by two providers. Because the prep batch and the provider are perfectly aliased they cannot be separated and are treated as a single batch factor. All leaves were harvested at the same time and all RNA extracted together, so this batch effect is technical, not biological.

```
# NOTE: the Trinity gene set is large, so the refit takes appreciably longer
# than the GDDH13 one. Set run_batch_sensitivity <- FALSE in chunk 0 to skip it.
run_sensitivity <- isTRUE(run_batch_sensitivity) &&
  "batch" %in% colnames(samples) &&
  nlevels(droplevels(samples$batch)) == 2

if (!run_sensitivity) {
  cat(strrep("!", 78), "\n")
  cat("BATCH-DESIGN SENSITIVITY ANALYSIS SKIPPED - NO RESULT IN THIS
REPORT.\n")
  cat("Diagnosis:\n")
  cat("  run_batch_sensitivity is TRUE       :", isTRUE(run_batch_sensitivity), "\n")
  cat("  a 'batch' column reached section 16:", "batch" %in% colnames(samples),
"\n")
  if ("batch" %in% colnames(samples)) {
```

```r
    cat("  levels of that batch factor       :",
        nlevels(droplevels(as.factor(samples$batch))), "\n")
  } else {
    cat("  batch metadata file used          :", batch_metadata_file, "\n")
    cat("  that file exists                  :", file.exists(batch_metadata_file), "\n")
  }
  cat("The manuscript and Supplementary Table S10 both cite this comparison, so
this\n")
  cat("report is NOT usable as a supplement until the lines above all read TRUE.
\n")
  cat(strrep("!", 78), "\n")
} else {
  # The Trinity object is large; release the QC matrices before the refit.
  suppressWarnings(rm(list = intersect(
    c("msd", "msd_points", "log2_raw_counts", "log2_norm_counts", "rlog_mat"),
    ls())))
  invisible(gc())

  # Refit on the same object, so the gene filter and size factors are comparable and
  # the two designs differ only in the model formula.
  dds_batch <- dds
  design(dds_batch) <- ~ batch + condition
  dds_batch <- DESeq(dds_batch)

  saveRDS(dds_batch, file.path(outdir,
"dds_Trinity_Salmon_batch_plus_condition.rds"))
  writeLines(resultsNames(dds_batch),
             file.path(outdir,
"DESeq2_Trinity_batch_plus_condition_results_names.txt"))

  cat("\nGenes tested under both designs:", nrow(dds_batch), "\n")

  thresholded <- function(object, contrast) {
    r <- results(object, contrast = contrast, alpha = 0.01,
                 lfcThreshold = 1, altHypothesis = "greaterAbs")
    r[!is.na(r$padj), ]
  }

  contrast_list <- list(
    Rin_vs_Rola     = c("condition", "Rin", "Rola"),
    Rin_vs_control  = c("condition", "Rin", "control"),
    Rola_vs_control = c("condition", "Rola", "control")
  )

  sens_rows <- list()

  for (nm in names(contrast_list)) {
    r_cond  <- thresholded(dds,       contrast_list[[nm]])
    r_batch <- thresholded(dds_batch, contrast_list[[nm]])
```

```r
  g_cond  <- rownames(r_cond)[r_cond$padj < 0.01]
  g_batch <- rownames(r_batch)[r_batch$padj < 0.01]
  shared  <- intersect(g_cond, g_batch)
  common  <- intersect(rownames(r_cond), rownames(r_batch))

  sens_rows[[nm]] <- data.frame(
    contrast                  = nm,
    genes_with_padj_in_both  = length(common),
    candidates_condition     = length(g_cond),
    candidates_batch         = length(g_batch),
    shared                   = length(shared),
    only_condition           = length(setdiff(g_cond, g_batch)),
    only_batch               = length(setdiff(g_batch, g_cond)),
    jaccard                  = round(length(shared) / length(union(g_cond, g_batch)),
4),
    pearson_log2FC           = round(cor(r_cond[common, "log2FoldChange"],
                                   r_batch[common, "log2FoldChange"]), 4),
    spearman_log2FC          = round(cor(r_cond[common, "log2FoldChange"],
                                   r_batch[common, "log2FoldChange"],
                                   method = "spearman"), 4),
    stringsAsFactors = FALSE
  )

  # Gene-level detail for EVERY contrast, not just Rin vs Rola. The 151-gene set is
  # defined by Rin-vs-control induction with baseMean >= 50 and the ABSENCE
of a
  # thresholded Rola-vs-control call, so testing whether those genes still meet
their
  # own defining criteria under the batch model needs the per-gene tables for all
  # three contrasts. baseMean is carried for the same reason.
  #
  # lfcSE is carried for both models because the standard error is what the extra
  # batch parameter actually changes; comparing it gene by gene shows whether
a
  # candidate was dropped because its effect estimate moved or because its
  # uncertainty grew.
  #
  # Genes evaluable under only one model are KEPT and labelled, not dropped.
The
  # earlier version intersected on non-NA padj, so candidates the batch model
could
  # not evaluate vanished from the file: for Rin vs Rola the summary counted 340
  # condition-only losses while the per-gene table explained only 313, the missing
27
  # being exactly those cases. A file used to audit disagreements must not hide
any.
  all_ids <- union(rownames(r_cond), rownames(r_batch))
  grab <- function(res, ids, col) {
    out <- rep(NA_real_, length(ids))
```

```
    hit <- ids %in% rownames(res)
    out[hit] <- res[ids[hit], col]
    out
  }
  detail <- data.frame(
    contrast           = nm,
    gene_id            = all_ids,
    baseMean_condition = grab(r_cond,  all_ids, "baseMean"),
    log2FC_condition   = grab(r_cond,  all_ids, "log2FoldChange"),
    lfcSE_condition    = grab(r_cond,  all_ids, "lfcSE"),
    padj_condition     = grab(r_cond,  all_ids, "padj"),
    baseMean_batch     = grab(r_batch, all_ids, "baseMean"),
    log2FC_batch       = grab(r_batch, all_ids, "log2FoldChange"),
    lfcSE_batch        = grab(r_batch, all_ids, "lfcSE"),
    padj_batch         = grab(r_batch, all_ids, "padj"),
    stringsAsFactors   = FALSE
  )
  detail$lfcSE_ratio <- detail$lfcSE_batch / detail$lfcSE_condition
  detail$evaluable_condition <- detail$gene_id %in% rownames(r_cond)
  detail$evaluable_batch     <- detail$gene_id %in% rownames(r_batch)
  detail$candidate_condition <- detail$gene_id %in% g_cond
  detail$candidate_batch     <- detail$gene_id %in% g_batch
  detail$agreement <- dplyr::case_when(
    !detail$evaluable_batch     & detail$candidate_condition ~
"condition_only_not_evaluable_in_batch",
    !detail$evaluable_condition & detail$candidate_batch     ~
"batch_only_not_evaluable_in_condition",
    detail$candidate_condition  &  detail$candidate_batch    ~ "both",
    detail$candidate_condition  & !detail$candidate_batch    ~
"only_condition_design",
    !detail$candidate_condition &  detail$candidate_batch    ~
"only_batch_design",
    TRUE ~ "neither"
  )
  detail <- detail |>
    dplyr::left_join(
      gene_annot |> dplyr::select(gene_id, gene_name, Description),
      by = "gene_id"
    ) |>
    dplyr::filter(agreement != "neither") |>
    dplyr::arrange(agreement, dplyr::desc(abs(log2FC_condition)))

  write.csv(
    detail,
    file.path(outdir, paste0("DESeq2_Trinity_batch_sensitivity_", nm,
"_genes.csv")),
    row.names = FALSE
  )
  cat("\n", nm, ": candidate genes written to the per-gene file: ", nrow(detail),
      " rows\n", sep = "")
```

```
    print(table(detail$agreement))
    stopifnot(sum(detail$candidate_condition) == length(g_cond),
              sum(detail$candidate_batch)     == length(g_batch))
  }

  sensitivity_summary <- do.call(rbind, sens_rows)

  write.csv(
    sensitivity_summary,
    file.path(outdir, "DESeq2_Trinity_batch_design_sensitivity_summary.csv"),
    row.names = FALSE
  )

  cat("\nBatch-design sensitivity, de novo Trinity (Salmon + tximport) layer:\n")
  print(sensitivity_summary)

  cat("\nHow to read this. Pearson and Spearman r near 1 mean the two models
agree on\n")
  cat("effect sizes and on gene rankings. The candidate columns are a separate
question.\n")
  cat("Where the two candidate sets differ, this design cannot say which model
is\n")
  cat("right: Rola is 2 IAB / 3 Eurofins, so batch and condition are mildly
correlated\n")
  cat("for Rola contrasts, though far less so than the 1/4 split assumed before
the\n")
  cat("assignment was corrected on 12 Aug 2026. The design is full rank, so
any\n")
  cat("residual effect is imprecision, not confounding in the DESeq2 sense of a\n")
  cat("non-identifiable model. A gene called under one model and not\n")
  cat("the other may reflect the power an additional term costs, or a call the
batch\n")
  cat("term removed because batch partly explained it. Note that the contrast
which\n")
  cat("changes least is Rin vs control, where both groups are balanced 4/4, and
the\n")
  cat("contrasts that change most both involve Rola. The per-gene table carries
lfcSE\n")
  cat("for both models and their ratio, which separates the two readings gene by
gene:\n")
  cat("an lfcSE_ratio well above 1 with a near-identical log2FC means the call was
lost\n")
  cat("to added uncertainty, whereas a shifted log2FC at similar lfcSE means the
batch\n")
  cat("term moved the estimate itself. Report both models and the per-gene\n")
  cat("disagreements; do not read the direction of the asymmetry as evidence
either\n")
  cat("way.\n")
}
```

```
## using pre-existing size factors

## estimating dispersions

## found already estimated dispersions, replacing these

## gene-wise dispersion estimates

## mean-dispersion relationship

## final dispersion estimates

## fitting model and testing

##
## Genes tested under both designs: 119509
##
## Rin_vs_Rola: candidate genes written to the per-gene file: 1902 rows
##
##                                both condition_only_not_evaluable_in_batch
##                                1626                                   28
##                  only_batch_design                only_condition_design
##                                  85                                  163
##
## Rin_vs_control: candidate genes written to the per-gene file: 4015 rows
##
## batch_only_not_evaluable_in_condition                                 both
##                                     1                                 3712
## condition_only_not_evaluable_in_batch                    only_batch_design
##                                    34                                  107
##                 only_condition_design
##                                   161
##
## Rola_vs_control: candidate genes written to the per-gene file: 5810 rows
##
## batch_only_not_evaluable_in_condition                                 both
##                                     3                                 5506
## condition_only_not_evaluable_in_batch                    only_batch_design
##                                    19                                   71
##                 only_condition_design
##                                   211
##
## Batch-design sensitivity, de novo Trinity (Salmon + tximport) layer:
##                        contrast genes_with_padj_in_both candidates_condition
## Rin_vs_Rola         Rin_vs_Rola                   68271                 1817
## Rin_vs_control   Rin_vs_control                   79822                 3907
## Rola_vs_control Rola_vs_control                   89117                 5736
##                 candidates_batch shared only_condition only_batch jaccard
## Rin_vs_Rola                 1711   1626            191         85  0.8549
## Rin_vs_control              3820   3712            195        108  0.9245
## Rola_vs_control             5580   5506            230         74  0.9477
##                 pearson_log2FC spearman_log2FC
```

```
## Rin_vs_Rola            0.9964         0.9947
## Rin_vs_control          0.9972         0.9964
## Rola_vs_control          0.9977         0.9922
##
## How to read this. Pearson and Spearman r near 1 mean the two models agree
on
## effect sizes and on gene rankings. The candidate columns are a separate
question.
## Where the two candidate sets differ, this design cannot say which model is
## right: Rola is 2 IAB / 3 Eurofins, so batch and condition are mildly correlated
## for Rola contrasts, though far less so than the 1/4 split assumed before the
## assignment was corrected on 12 Aug 2026. The design is full rank, so any
## residual effect is imprecision, not confounding in the DESeq2 sense of a
## non-identifiable model. A gene called under one model and not
## the other may reflect the power an additional term costs, or a call the batch
## term removed because batch partly explained it. Note that the contrast which
## changes least is Rin vs control, where both groups are balanced 4/4, and the
## contrasts that change most both involve Rola. The per-gene table carries lfcSE
## for both models and their ratio, which separates the two readings gene by
gene:
## an lfcSE_ratio well above 1 with a near-identical log2FC means the call was
lost
## to added uncertainty, whereas a shifted log2FC at similar lfcSE means the
batch
## term moved the estimate itself. Report both models and the per-gene
## disagreements; do not read the direction of the asymmetry as evidence either
## way.
```

### Output overview

In addition to the DESeq2 result tables, candidate-gene tables, QC plots and sessionInfo.txt, section 16 writes DESeq2_Trinity_batch_design_sensitivity_summary.csv and one per-gene file per contrast, DESeq2_Trinity_batch_sensitivity_{Rin_vs_Rola,Rin_vs_control,Rola_vs_control}_genes.csv, to outdir.

```
## Main output directory:
##  /Volumes/Data_Disk/Ametyst_re-analysis/Trinity/DESeq2
##
##  [1] "build_table2_final.js"
##  [2] "Candidate_genes_Rin_high_padj0.05_LFC1_baseMean50.csv"
##  [3] "Candidate_genes_Rola_high_padj0.05_LFC1_baseMean50.csv"
##  [4] "Candidate_Rin_vs_C_C_high_lfcThreshold1_padj0.01_by_log2FC.csv"
##  [5] "Candidate_Rin_vs_C_lfcThreshold1_padj0.01_by_abs_log2FC.csv"
##  [6] "Candidate_Rin_vs_C_Rin_high_lfcThreshold1_padj0.01_by_log2FC.csv"
##  [7]
"Candidate_Rin_vs_Rola_lfcThreshold1_padj0.01_by_abs_log2FC_Trinity.csv"
##  [8] "Candidate_Rin_vs_Rola_lfcThreshold1_padj0.01_by_abs_log2FC.csv"
```

```
## [9] "Candidate_Rin_vs_Rola_Rin_high_lfcThreshold1_padj0.01_by_log2FC.csv"
## [10]
"Candidate_Rin_vs_Rola_Rola_high_lfcThreshold1_padj0.01_by_log2FC.csv"
## [11] "Candidate_Rola_vs_C_C_high_lfcThreshold1_padj0.01_by_log2FC.csv"
## [12] "Candidate_Rola_vs_C_lfcThreshold1_padj0.01_by_abs_log2FC.csv"
## [13] "Candidate_Rola_vs_C_Rola_high_lfcThreshold1_padj0.01_by_log2FC.csv"
## [14] "candidate.js"
## [15] "Comparison_class_summary_lfcThreshold1_padj0.01.csv"
## [16]
"Comparison_Rin_vs_C_Rola_vs_C_Rin_vs_Rola_lfcThreshold1_padj0.01.csv"
## [17] "dds_Trinity_Salmon_batch_plus_condition.rds"
## [18] "dds_Trinity_Salmon_condition_only.rds"
## [19] "DESeq2_results_names.txt"
## [20]
"DESeq2_Rin_vs_C_lfcThreshold1_padj0.01_with_description_by_log2FC.csv"
## [21]
"DESeq2_Rin_vs_Rola_lfcThreshold1_padj0.01_with_description_by_log2FC.csv"
## [22]
"DESeq2_Rola_vs_C_lfcThreshold1_padj0.01_with_description_by_log2FC.csv"
## [23] "DESeq2_Trinity_batch_design_sensitivity_summary.csv"
## [24] "DESeq2_Trinity_batch_plus_condition_results_names.txt"
## [25] "DESeq2_Trinity_batch_sensitivity_Rin_vs_control_genes.csv"
## [26] "DESeq2_Trinity_batch_sensitivity_Rin_vs_Rola_genes.csv"
## [27] "DESeq2_Trinity_batch_sensitivity_Rola_vs_control_genes.csv"
## [28] "DESeq2_Trinity_condition_only_normalized_counts.csv"
## [29]
"DESeq2_Trinity_condition_only_Rin_vs_C_lfcThreshold1_with_description.csv"
## [30] "DESeq2_Trinity_condition_only_Rin_vs_control_with_description.csv"
## [31] "DESeq2_Trinity_condition_only_Rin_vs_control.csv"
## [32]
"DESeq2_Trinity_condition_only_Rin_vs_Rola_lfcThreshold1_with_description.csv"
## [33] "DESeq2_Trinity_condition_only_Rin_vs_Rola_with_description.csv"
## [34] "DESeq2_Trinity_condition_only_Rin_vs_Rola.csv"
## [35]
"DESeq2_Trinity_condition_only_Rola_vs_C_lfcThreshold1_with_description.csv"
## [36] "DESeq2_Trinity_condition_only_Rola_vs_control_with_description.csv"
## [37] "DESeq2_Trinity_condition_only_Rola_vs_control.csv"
## [38] "DESeq2_Trinity_condition_only_summary_counts.csv"
## [39] "DESeq2_Trinity_OmicsBox.docx"
## [40] "DESeq2_Trinity_OmicsBox.pdf"
## [41] "make_table_extended.js"
## [42] "make_table.js"
## [43] "make_table.R"
## [44] "metadata_used_for_DESeq2.csv"
## [45] "node_modules"
## [46] "package-lock.json"
## [47] "package.json"
## [48] "PCA_scores_condition_only.csv"
## [49] "PCA_Trinity_condition_only_by_condition.png"
## [50] "PCA_Trinity_condition_only_raw_vst_by_condition.png"
```

```
## [51] "QC_extra_plots"
## [52] "Sample_distance_heatmap_Trinity_condition_only_raw_vst.png"
## [53] "Scatter_log2FC_Rin_vs_C_vs_Rola_vs_C.png"
## [54] "sessionInfo.txt"
## [55] "Table2_DEG_summary.docx"
## [56] "Table2_trinity_counts.csv"
## [57] "tx2gene_one_to_one_Trinity_gene_representatives.tsv"

##
## QC output directory:
##  /Volumes/Data_Disk/Ametyst_re-analysis/Trinity/DESeq2/QC_extra_plots
##
##  [1] "aheatmap_Pearson_correlation_VST.png"
##  [2] "aheatmap_Rin_vs_Rola_significant_genes_VST_scaled.png"
##  [3] "aheatmap_top500_variable_genes_VST_scaled.png"
##  [4] "Boxplot_log2_normalized_counts_plus1.png"
##  [5] "Boxplot_log2_raw_counts_plus1.png"
##  [6] "Boxplot_normalized_counts.png"
##  [7] "Boxplot_raw_counts.png"
##  [8] "Boxplot_VST_counts.png"
##  [9] "Heatmap_Rin_vs_Rola_significant_genes_VST_pheatmap.png"
## [10] "Heatmap_top25_Rin_high_top25_Rola_high_VST_row_scaled.png"
## [11] "Heatmap_top50_Rin_high_genes_VST_row_scaled.png"
## [12] "Heatmap_top50_Rola_high_genes_VST_row_scaled.png"
## [13] "Heatmap_top500_variable_genes_VST_pheatmap.png"
## [14] "Heteroskedasticity_mean_SD_plots.png"
## [15] "MA_input_Rin_vs_Rola_DESeq2.csv"
## [16] "MA_plot_Rin_vs_Rola_custom_ggplot.png"
## [17] "MA_plot_Rin_vs_Rola_DESeq2.png"
## [18] "MA_plot_Rin_vs_Rola_LFC_shrunk_ashr.png"
## [19] "Mean_SD_by_transformation.csv"
## [20] "Pearson_correlation_heatmap_VST.png"
## [21] "Pearson_correlation_VST_matrix.csv"
## [22] "Volcano_input_Rin_vs_Rola.csv"
## [23] "Volcano_plot_Rin_vs_Rola.png"
## [24] "vsn_meanSdPlot_log2_normalized_counts.png"
## [25] "vsn_meanSdPlot_VST.png"
## [26] "VST_expression_matrix.csv"
```

### Report appendix: inline figures and table previews

This section embeds saved PNG figures directly into the knitted report and shows the first 10 rows of each generated CSV table.

#### Inline graphical output

##### Figure 1

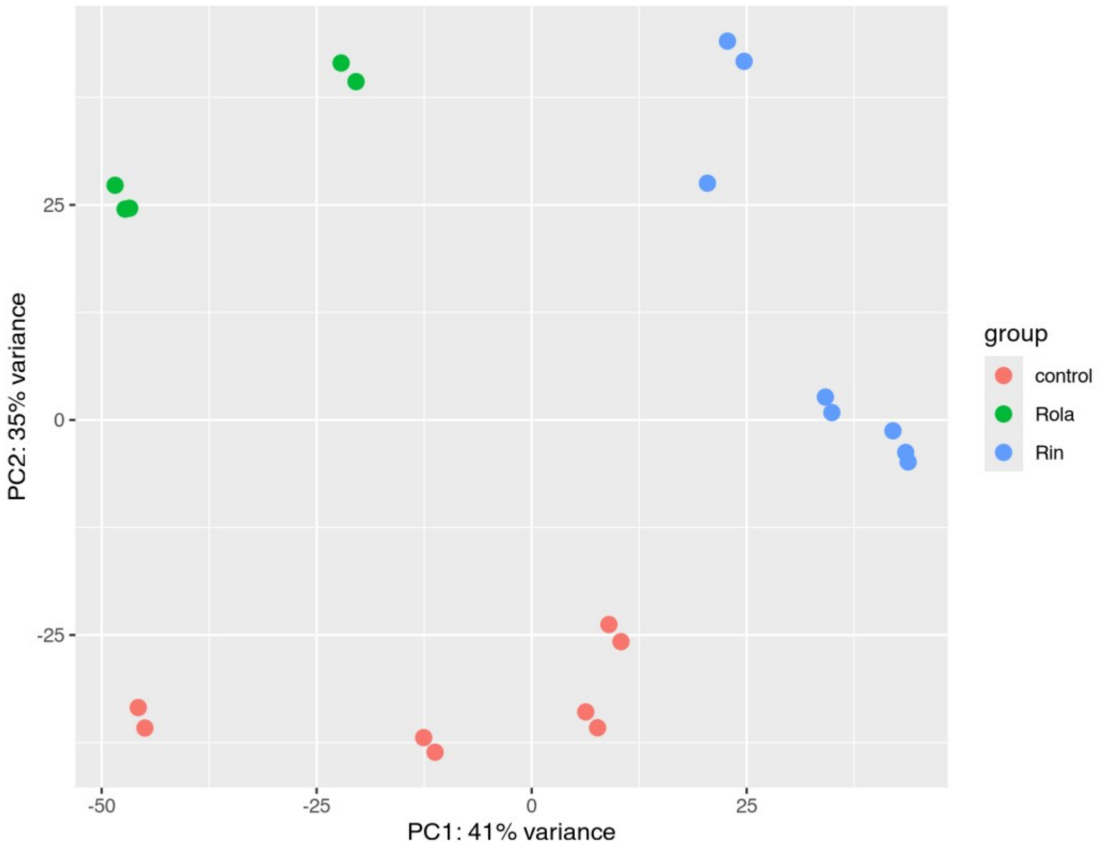


*PCA of VST-transformed Trinity gene expression values, colored by condition.*

Figure 2

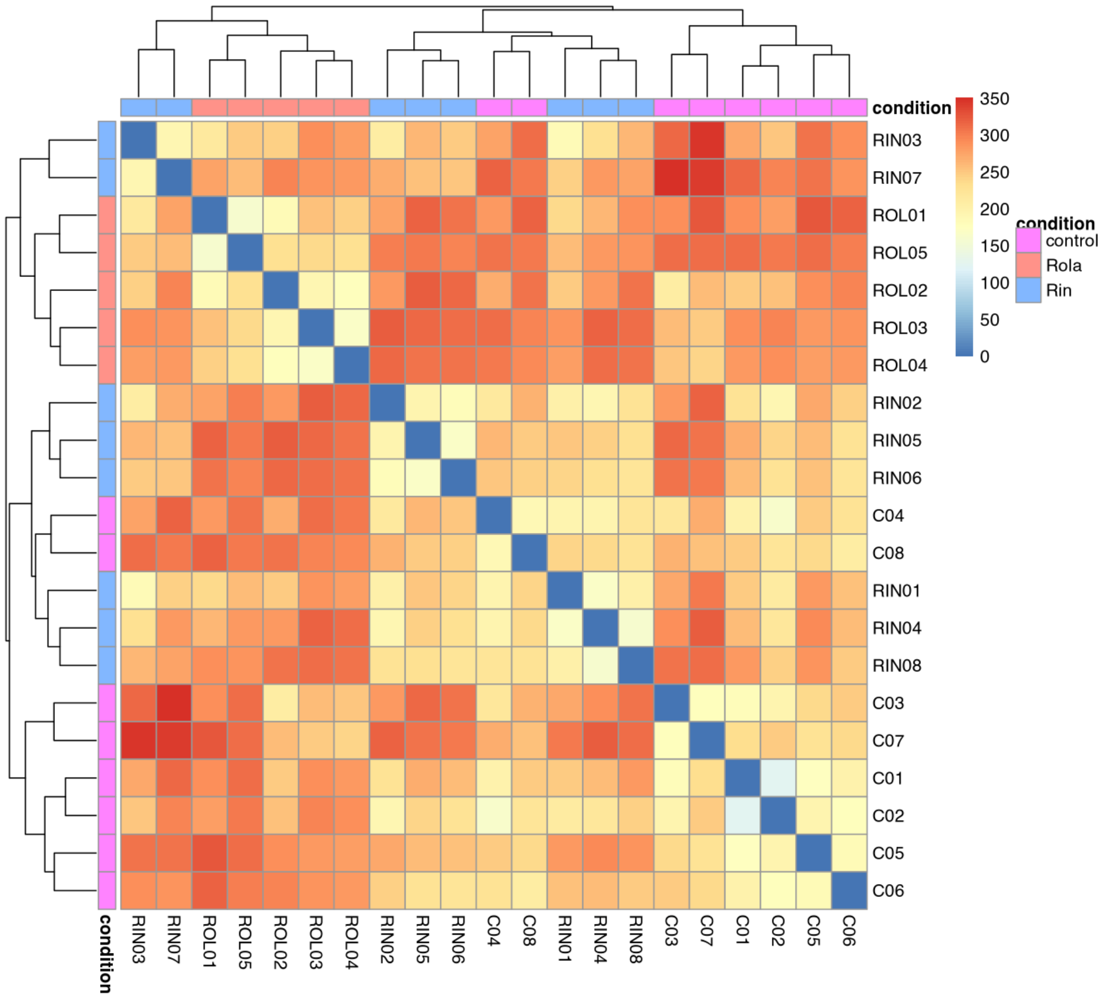


*Sample-to-sample distance heatmap based on VST-transformed Trinity gene expression values.*

#### Figure 3

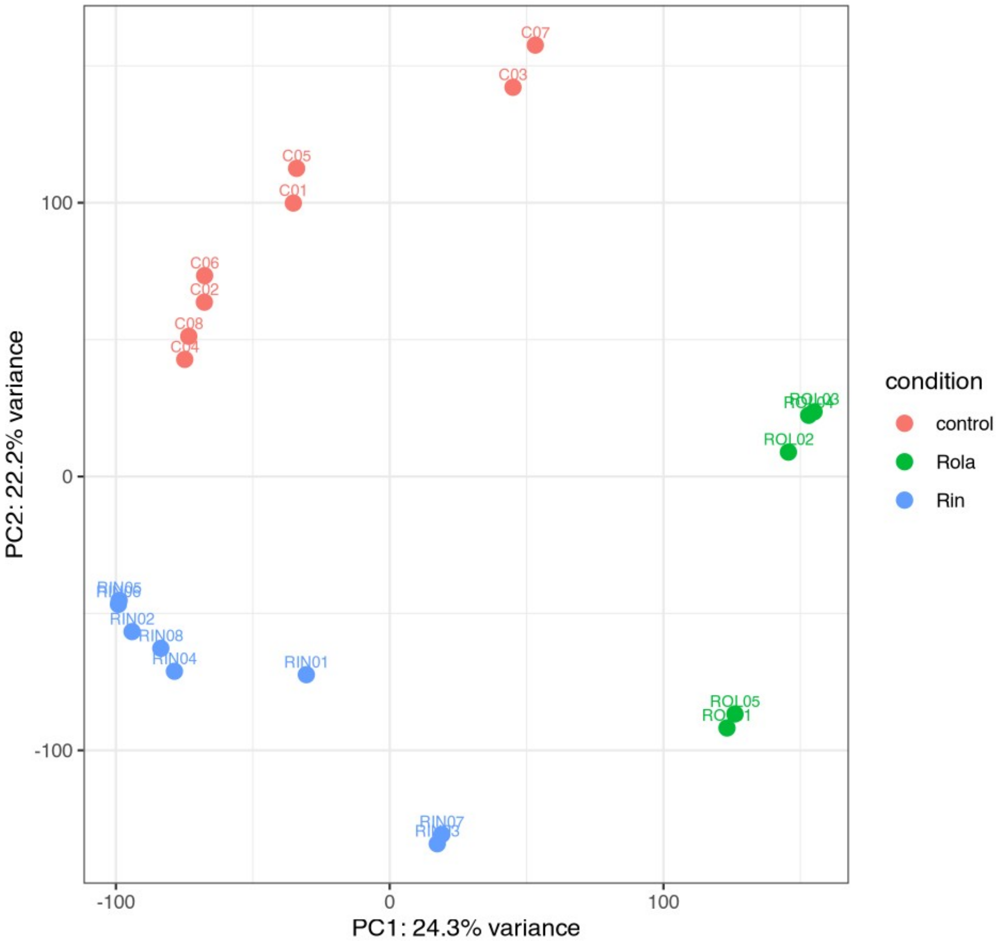


*Manual PCA of VST-transformed Trinity gene expression values, colored by condition.*

#### Figure 4

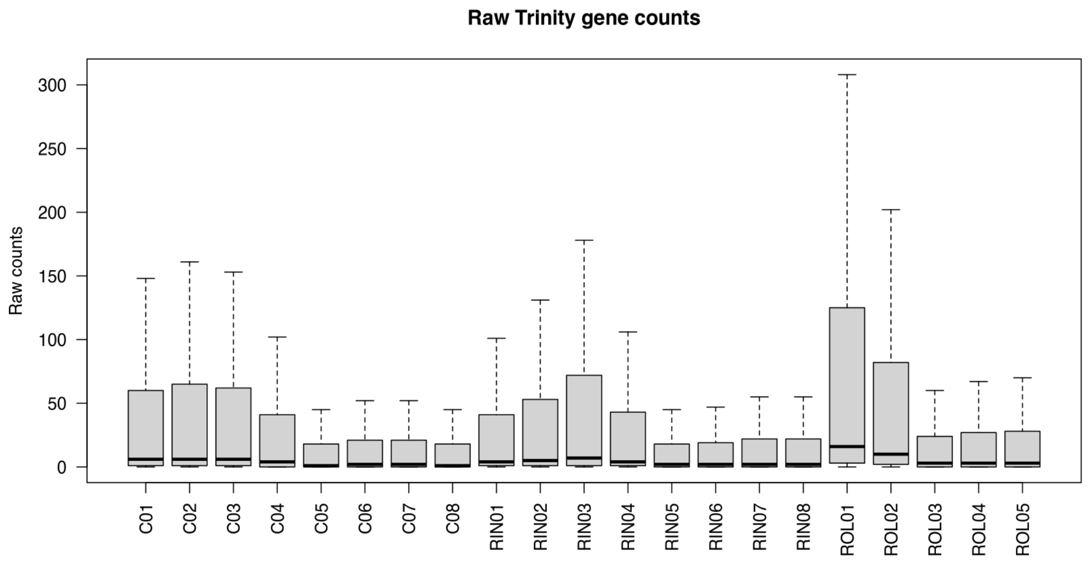


*Distribution of raw Trinity gene counts across samples.*

#### Figure 5

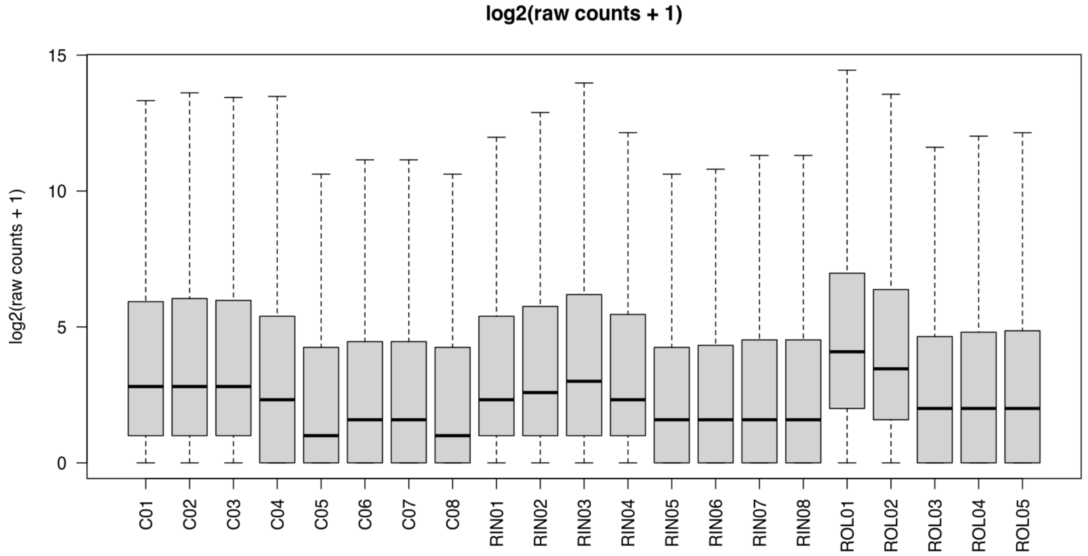


*Distribution of log2(raw counts + 1) across samples.*

#### Figure 6

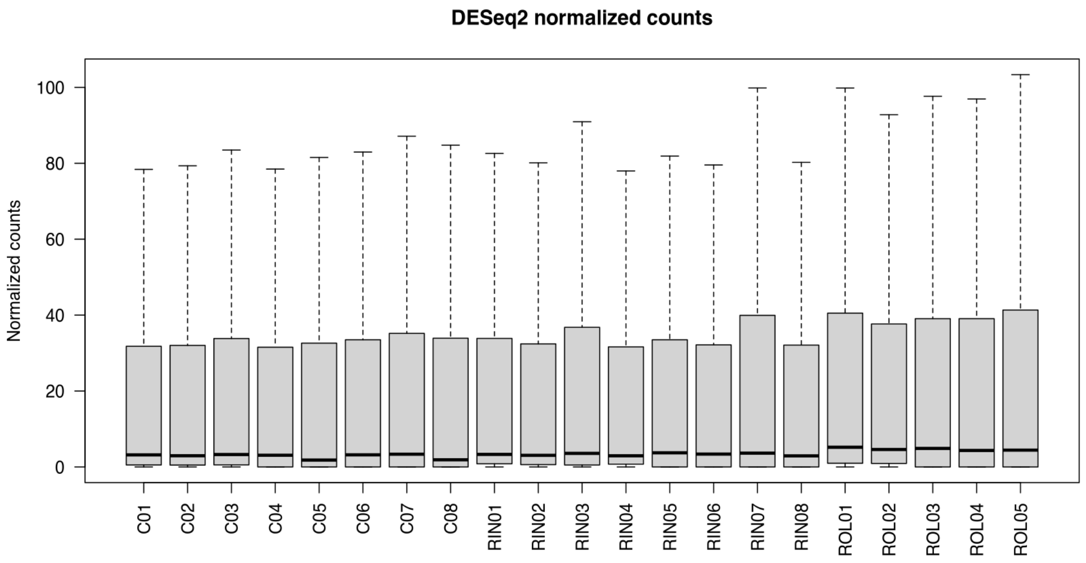


*Distribution of DESeq2 normalized counts across samples.*

#### Figure 7

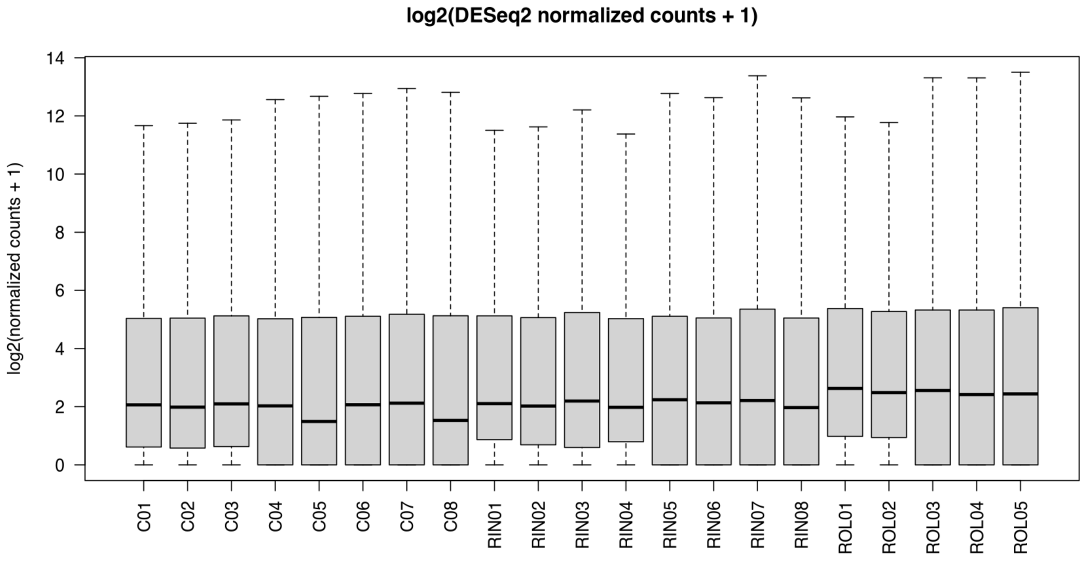


*Distribution of log2(normalized counts + 1) across samples.*

#### Figure 8

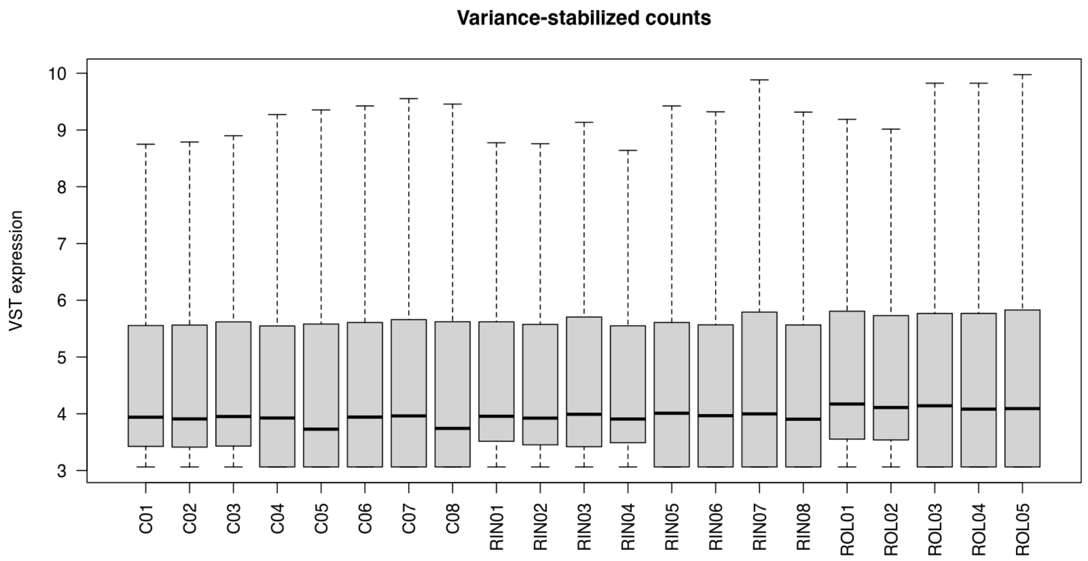


*Distribution of VST-transformed expression values across samples.*

#### Figure 9

Missing PNG file:
/Volumes/Data_Disk/Ametyst_re-analysis/Trinity/DESeq2/QC_extra_plots/
Boxplot_rlog_counts.png

#### Figure 10

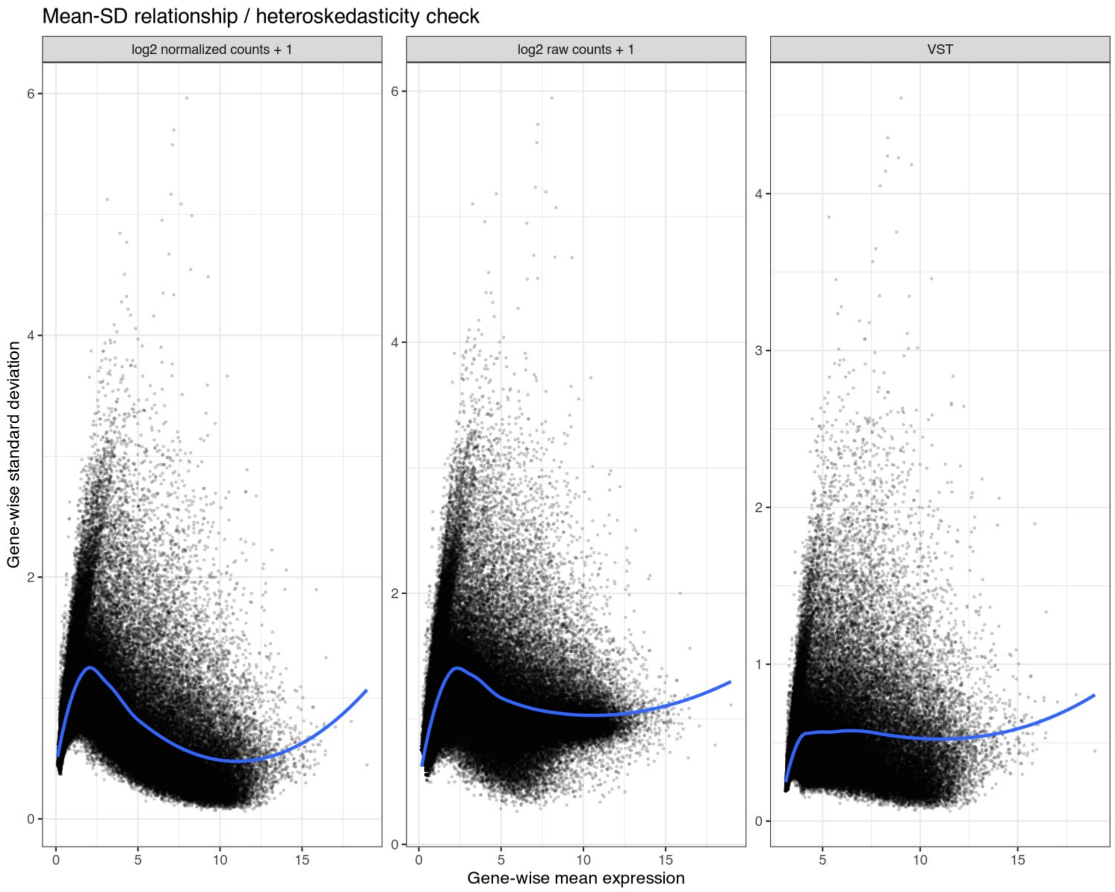


*Mean-SD relationship for raw, normalized, VST, and optional rlog transformations.*

#### Figure 11

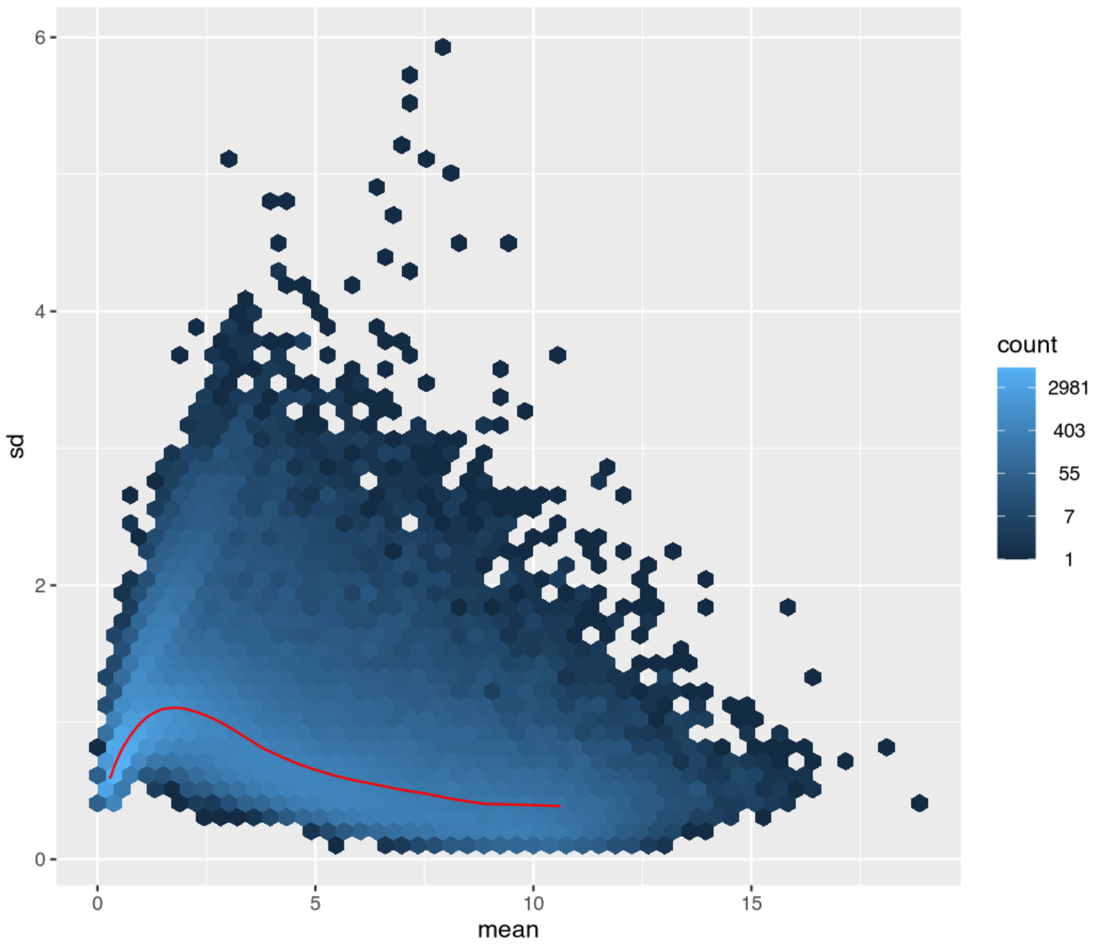


*vsn mean-SD plot for log2 normalized counts.*

#### Figure 12

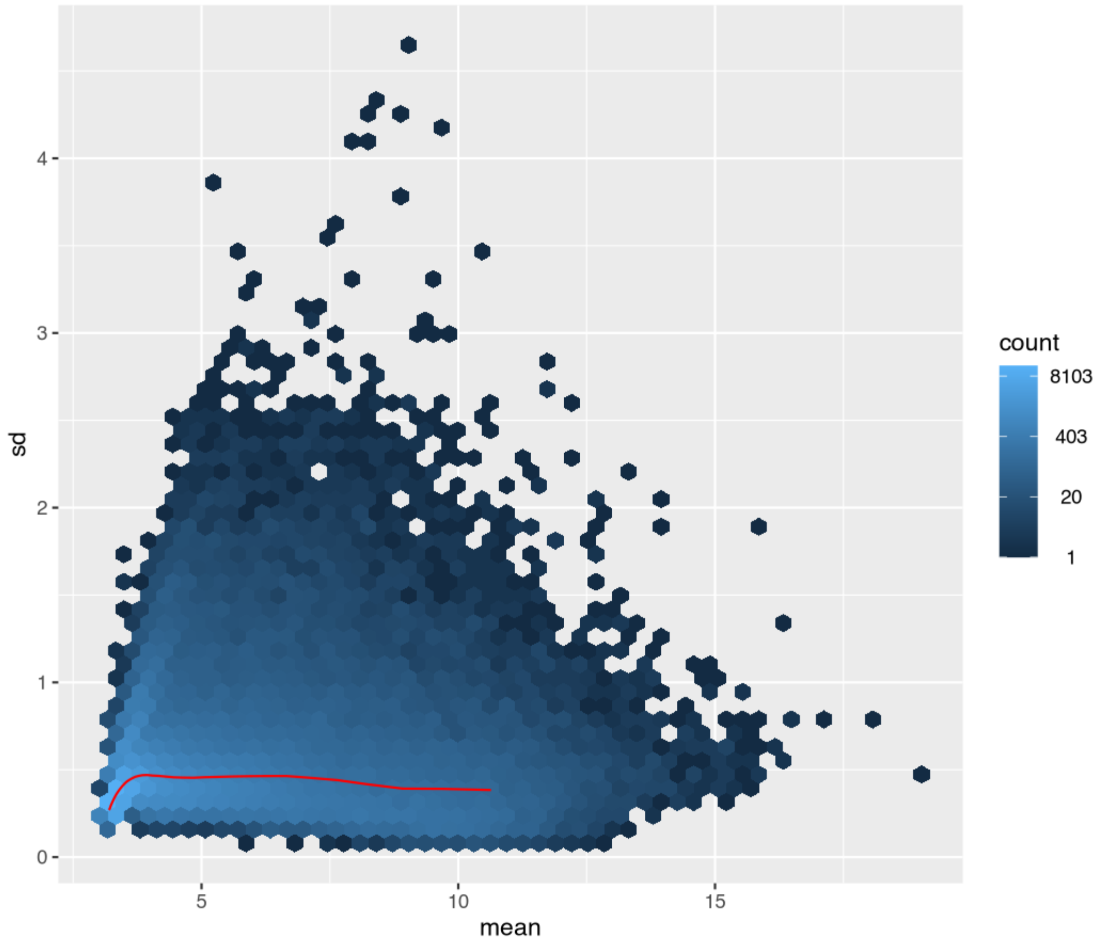


*vsn mean-SD plot for VST values.*

#### Figure 13

Missing PNG file:
/Volumes/Data_Disk/Ametyst_re-analysis/Trinity/DESeq2/QC_extra_plots/vsn_meanSdPlot_rlog.png

#### Figure 14

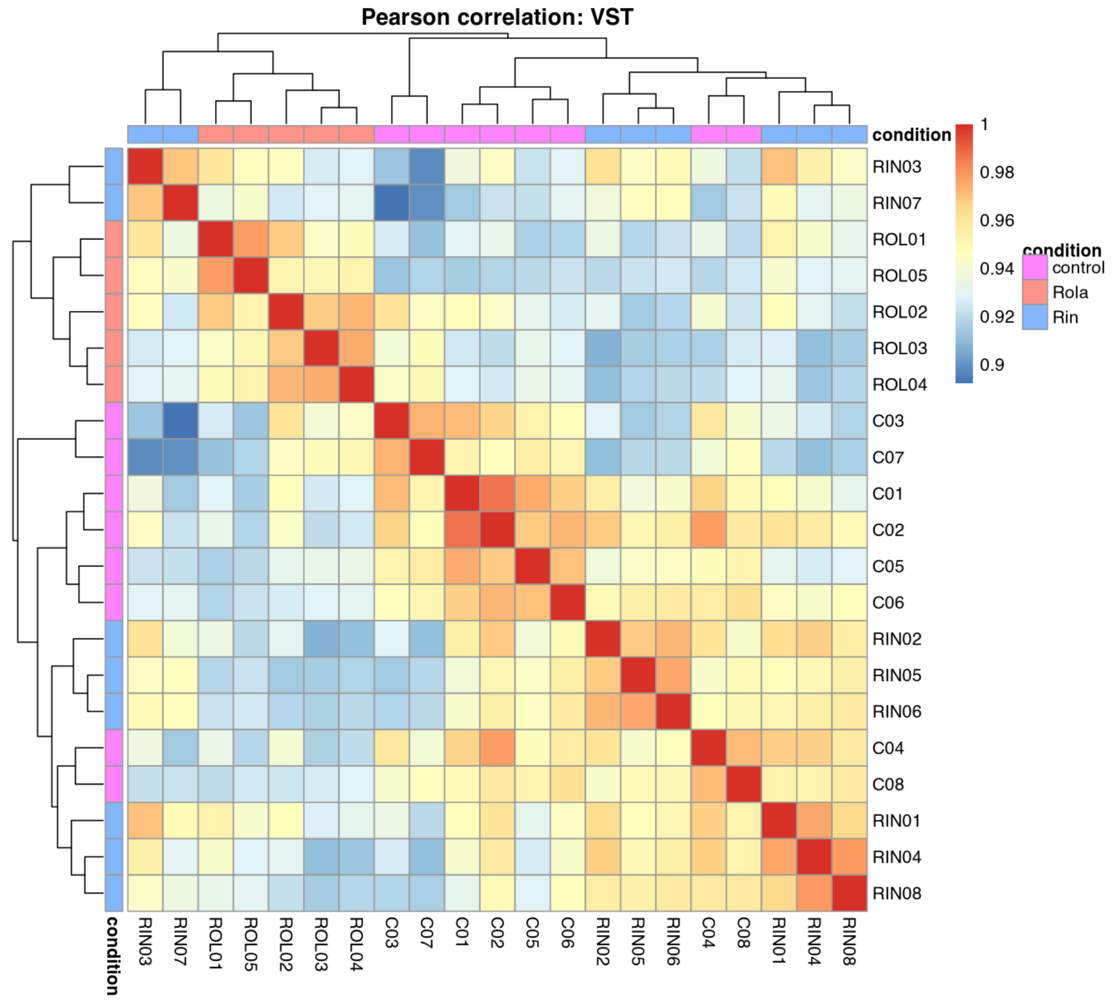


*Pearson correlation heatmap based on VST-transformed expression values.*

#### Figure 15

Missing PNG file:
/Volumes/Data_Disk/Ametyst_re-analysis/Trinity/DESeq2/QC_extra_plots/
Pearson_correlation_heatmap_rlog.png

Figure 16

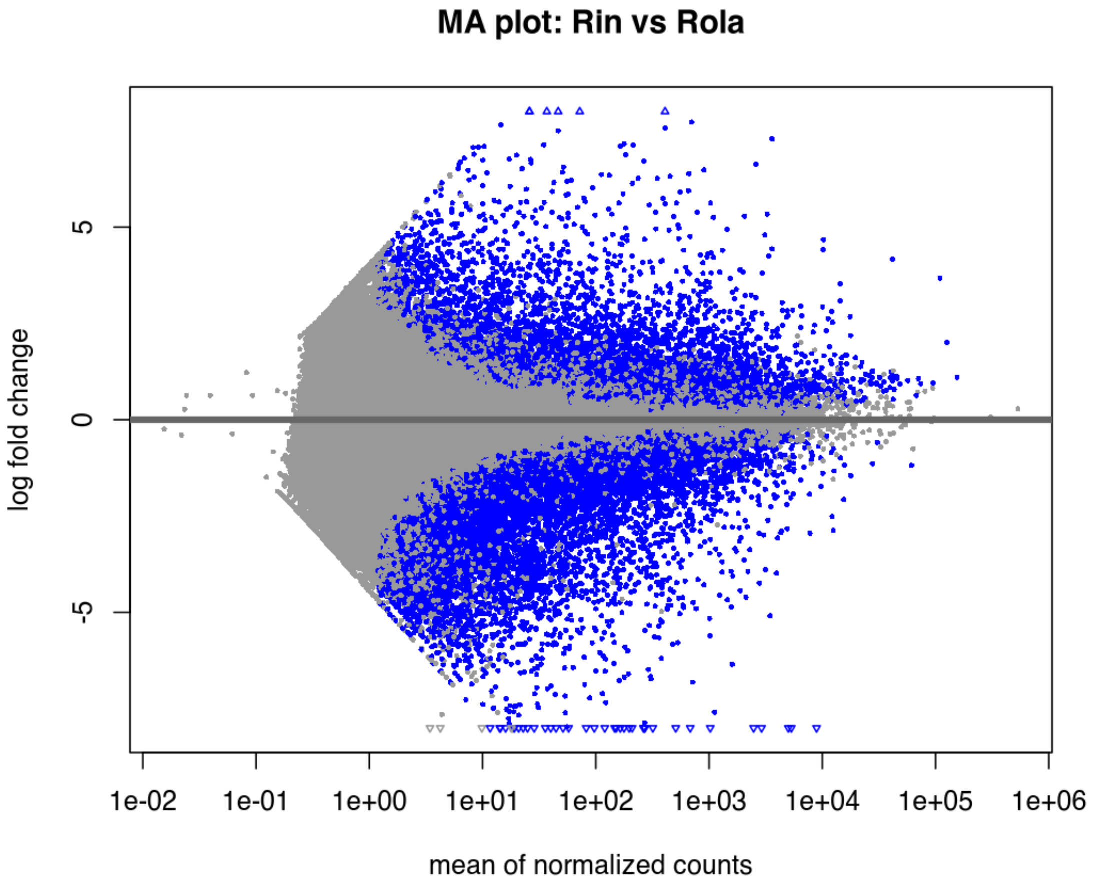


*DESeq2 MA plot for Rin vs Rola.*

#### Figure 17

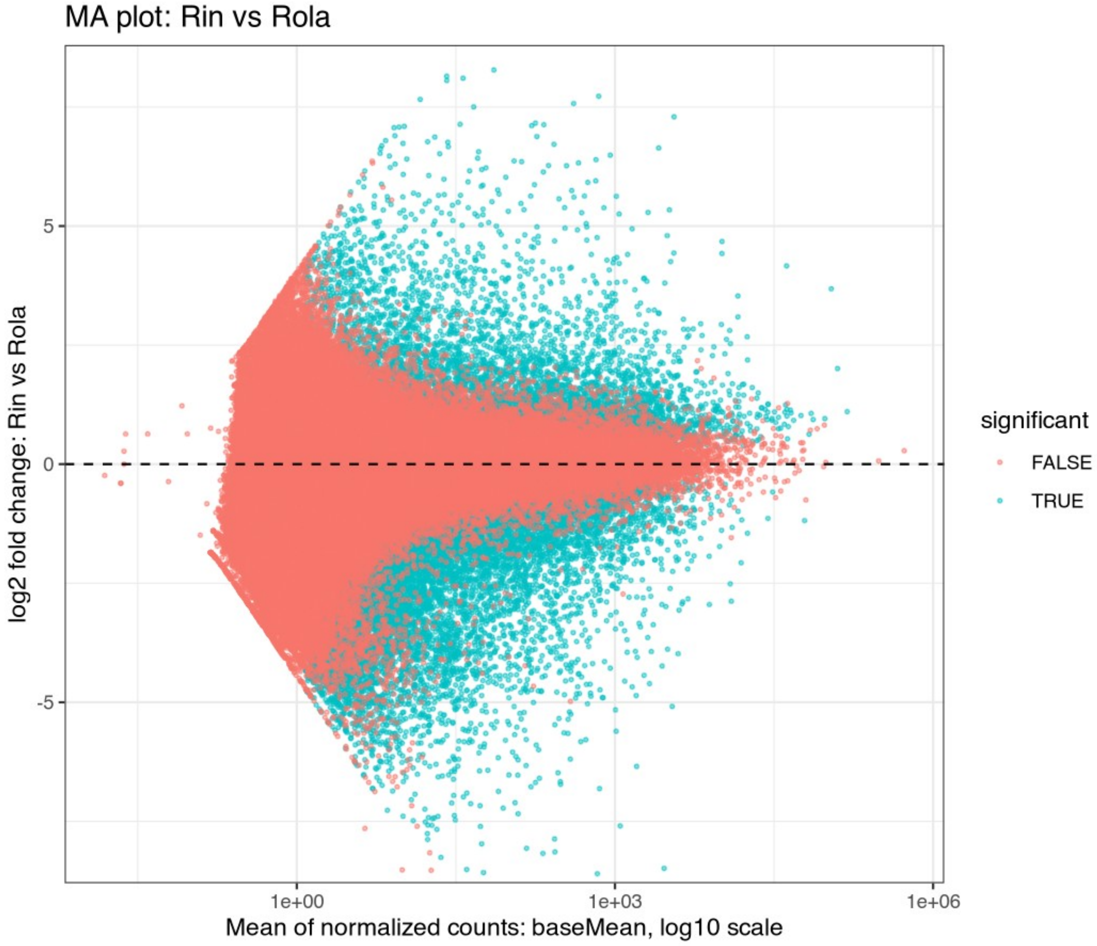


*Custom ggplot MA plot for Rin vs Rola.*

Figure 18

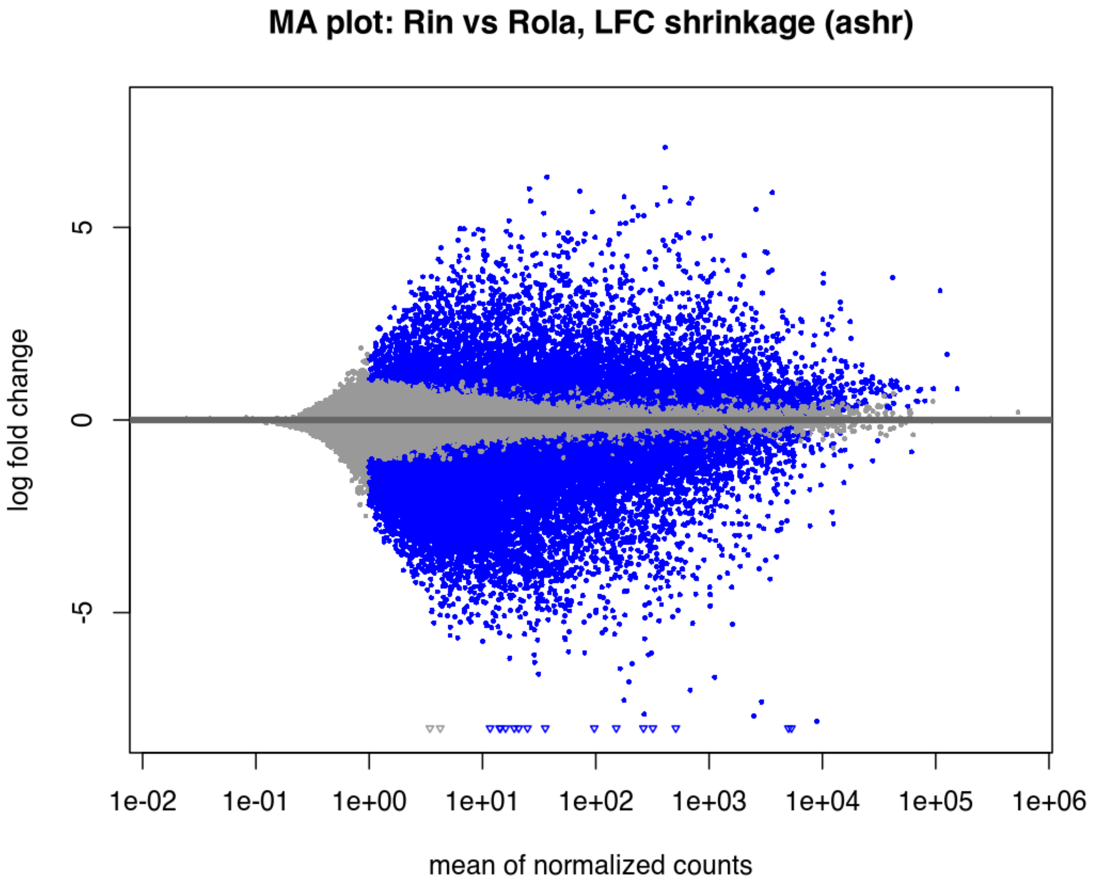


*MA plot using shrunken log2 fold changes for Rin vs Rola, if ashr was available.*

#### Figure 19

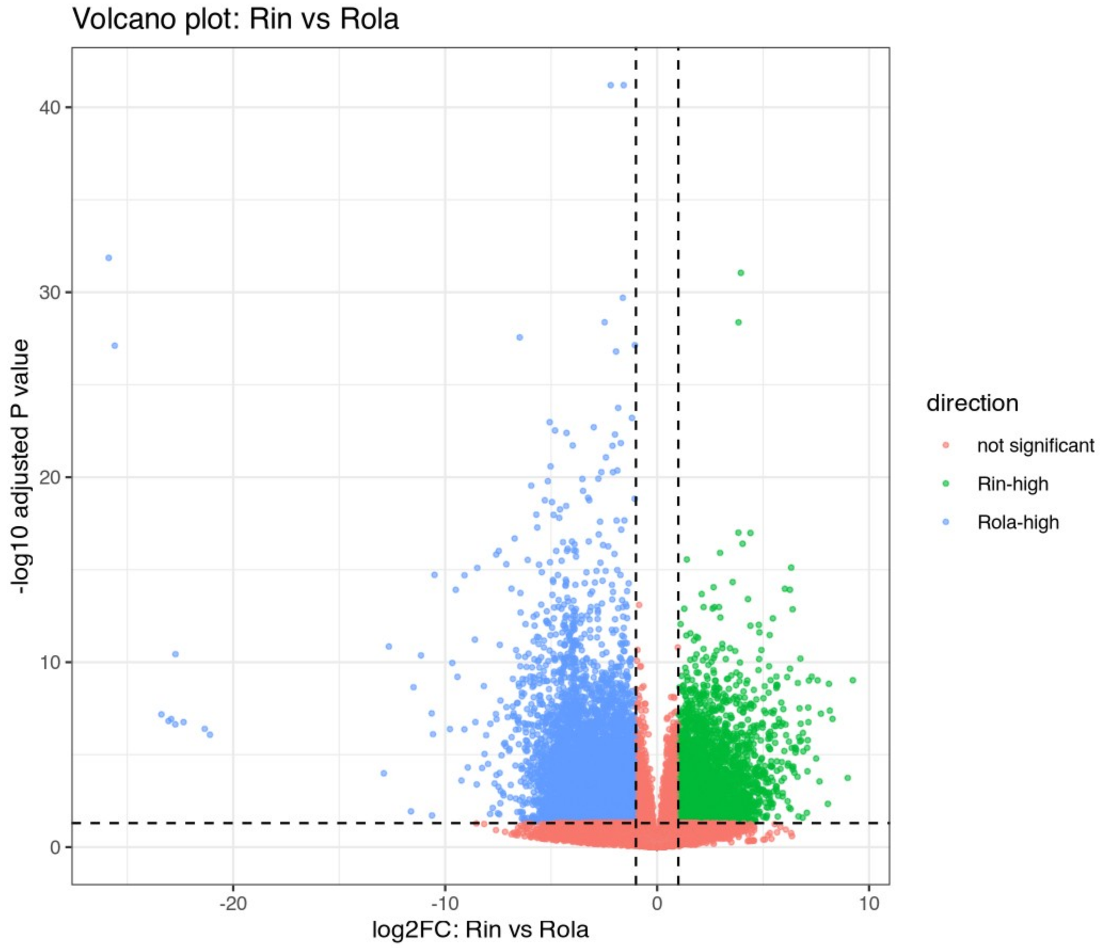


*Volcano plot for Rin vs Rola.*

#### Figure 20

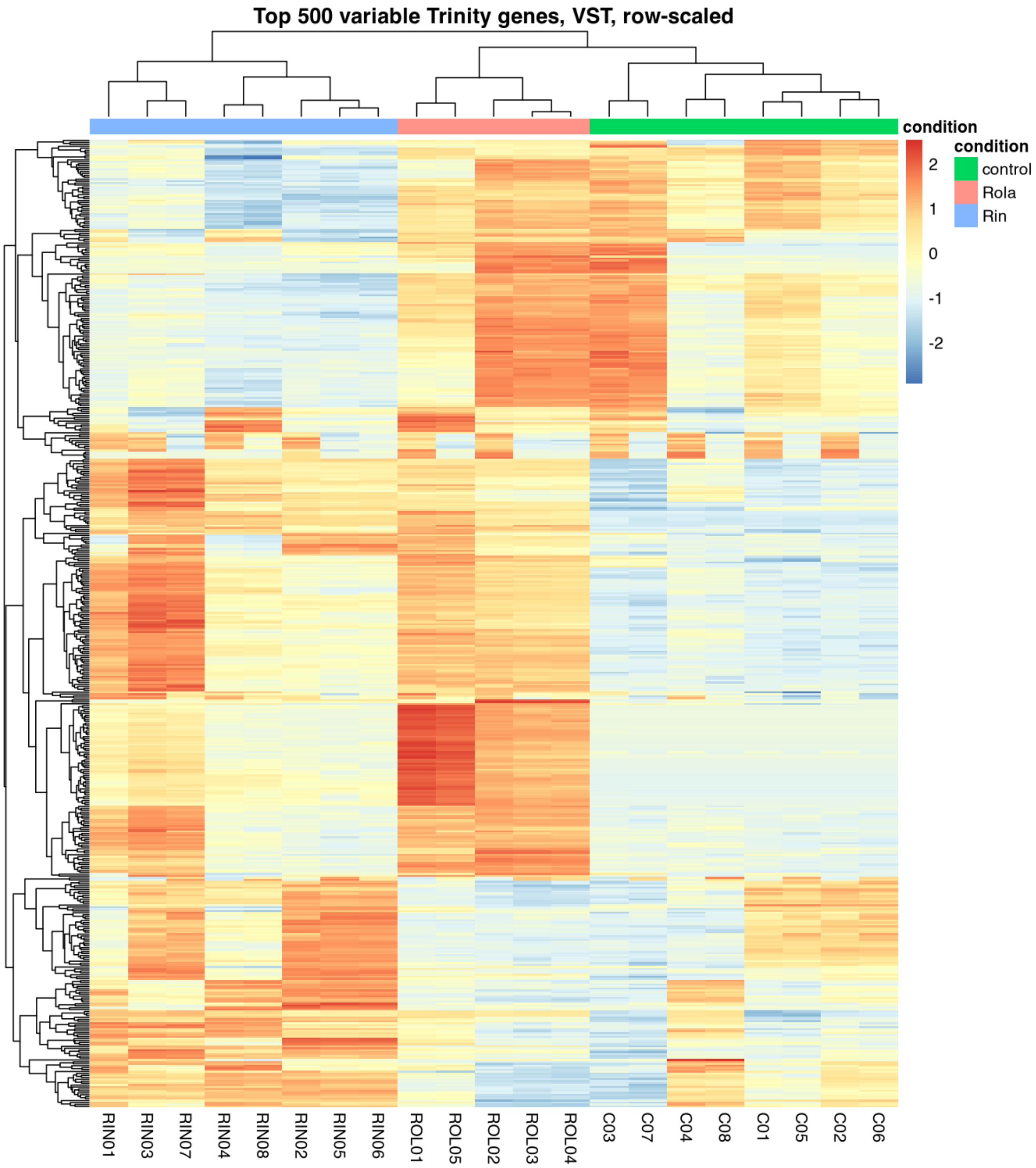


*Heatmap of top 500 most variable Trinity genes, VST row-scaled.*

#### Figure 21

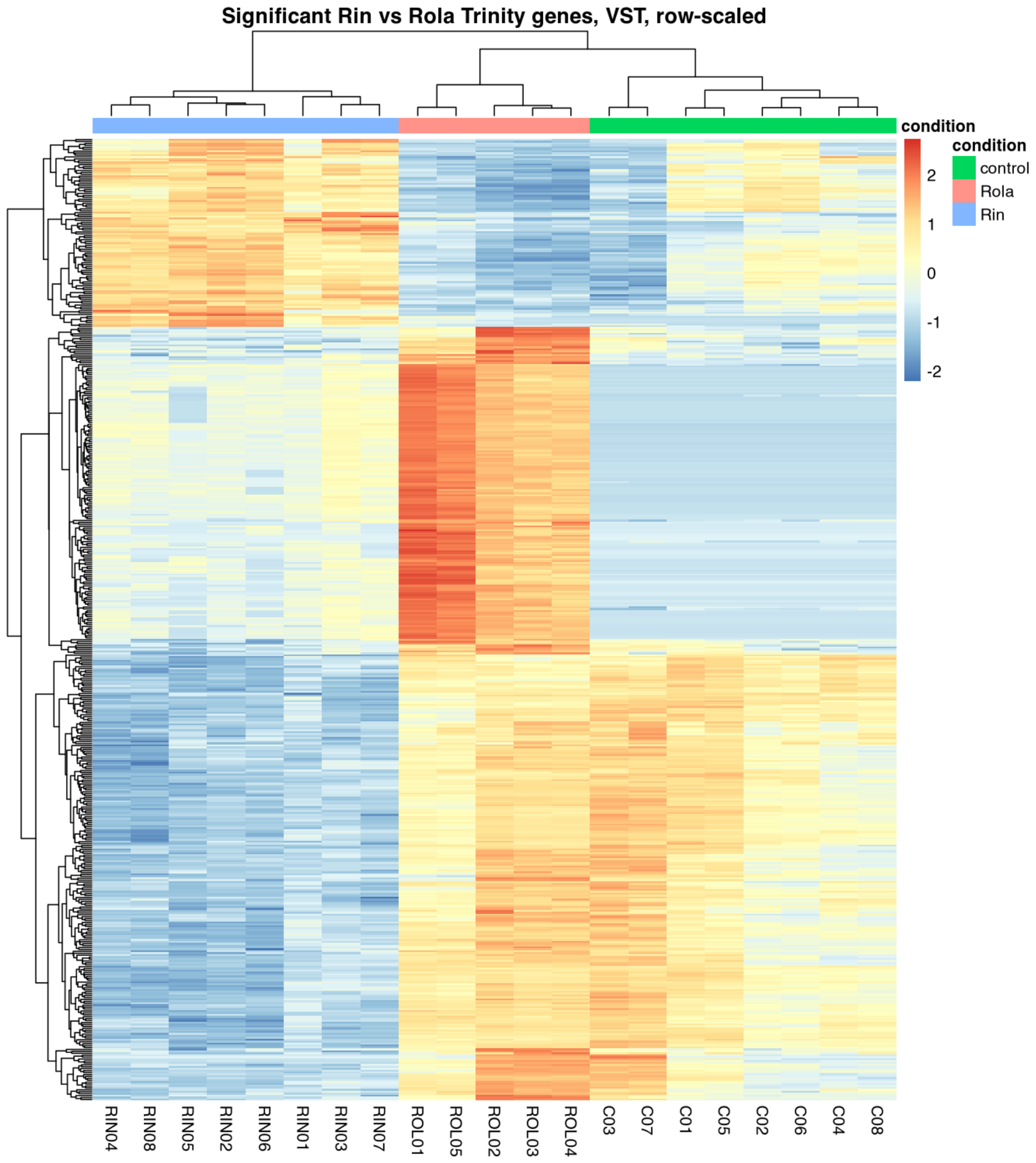


*Heatmap of significant Rin vs Rola Trinity genes, if enough genes passed the filter.*

### Figure 22

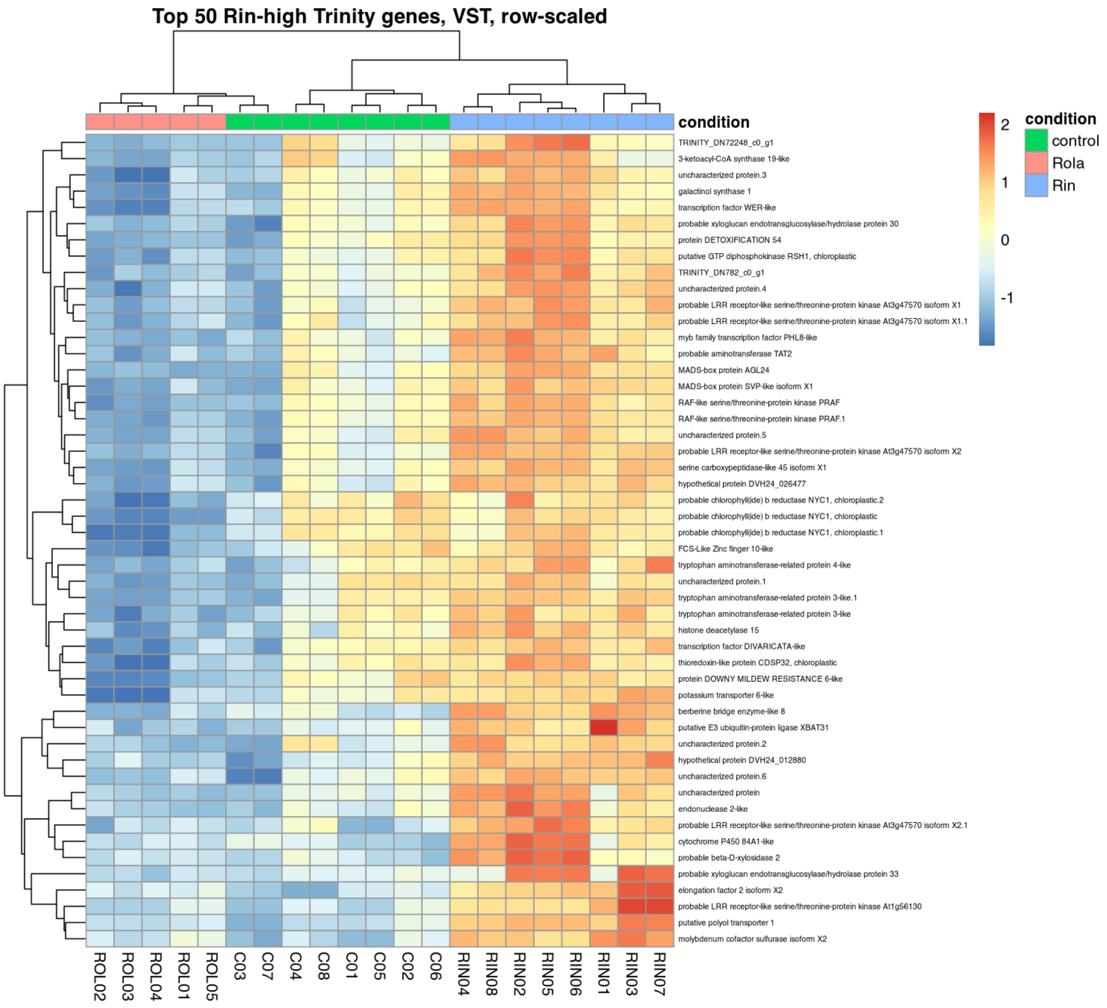


*Heatmap of top Rin-high candidate Trinity genes.*

### Figure 23

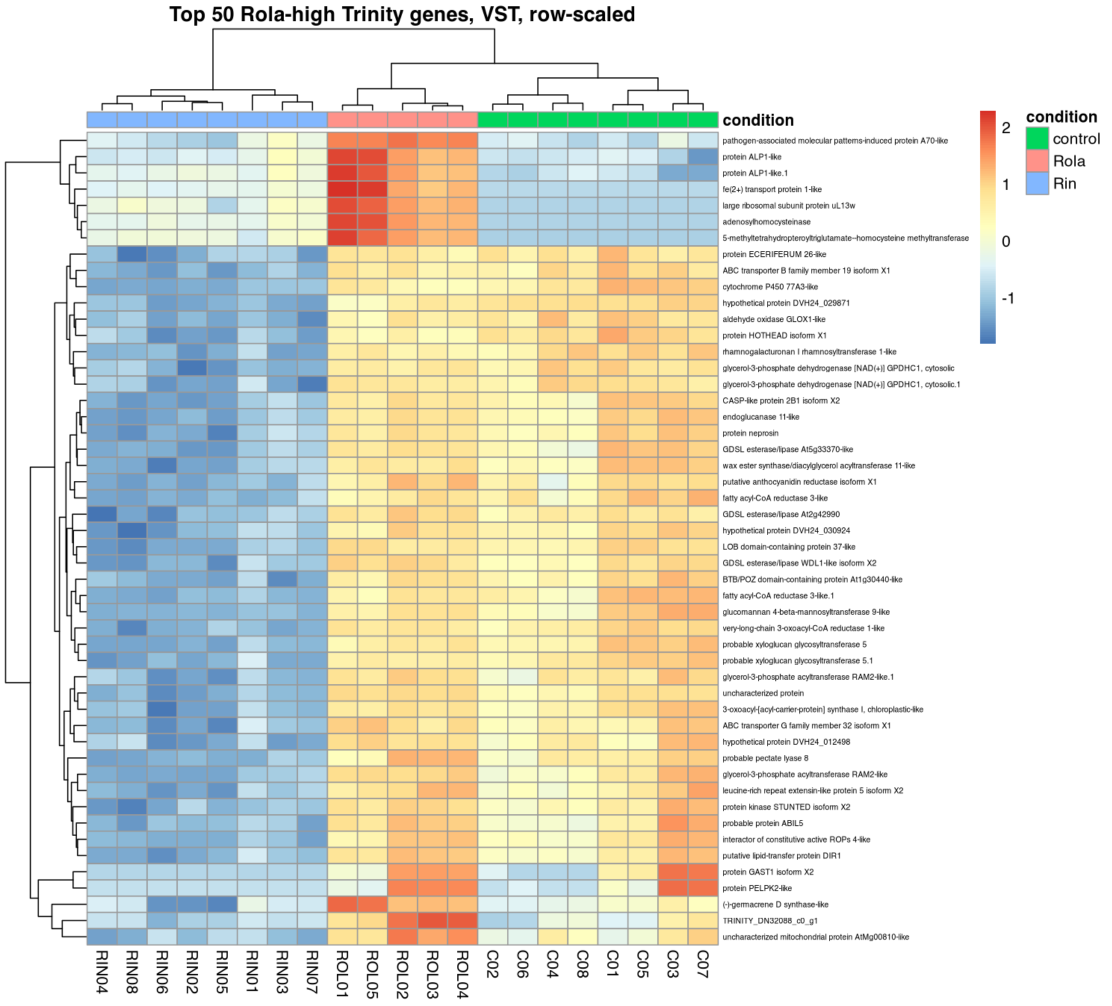


*Heatmap of top Rola-high candidate Trinity genes.*

### Figure 24

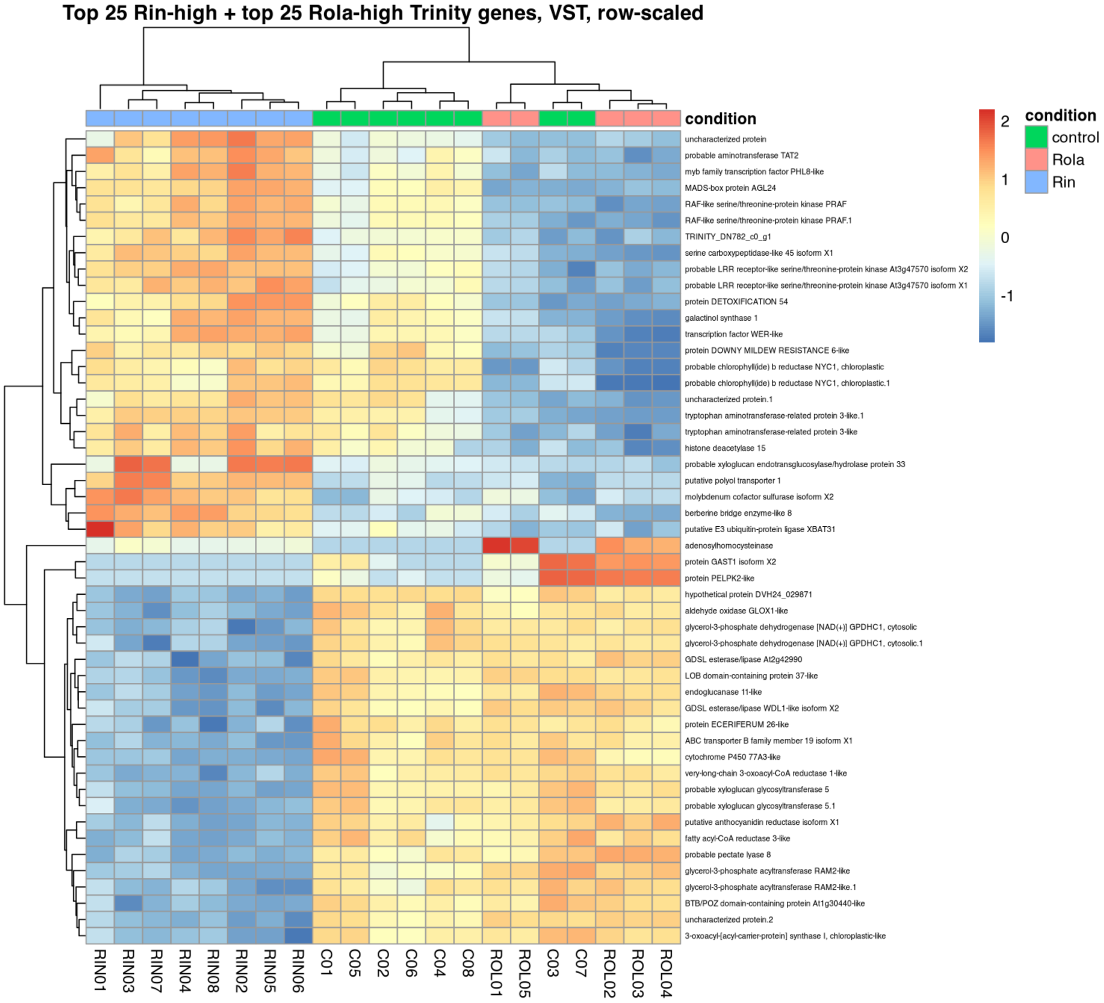


*Combined heatmap of top Rin-high and Rola-high Trinity candidate genes.*

#### Figure 25

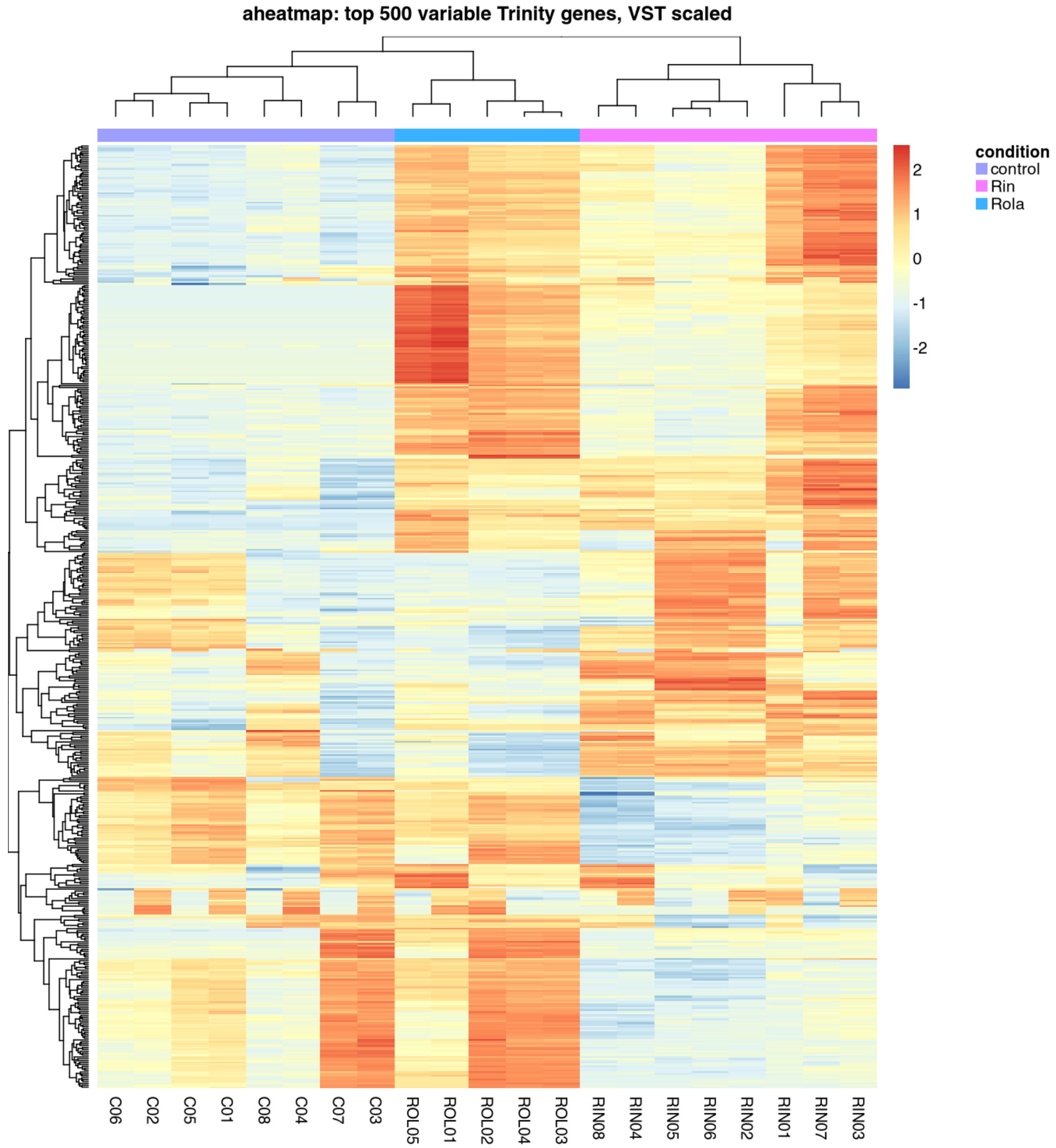


*NMF aheatmap of top variable Trinity genes, if NMF was available.*

### Figure 26

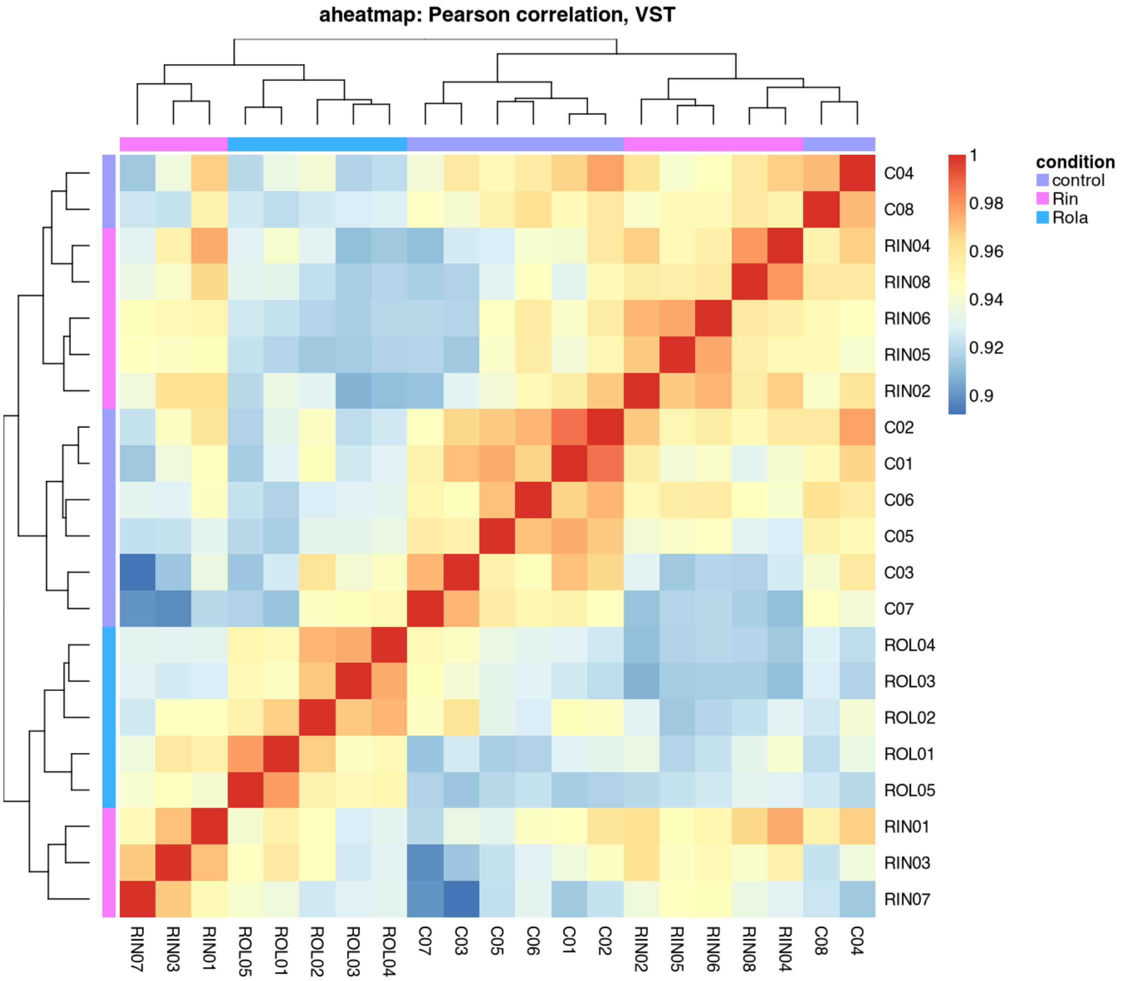


*NMF aheatmap of Pearson correlation matrix, if NMF was available.*

### Figure 27

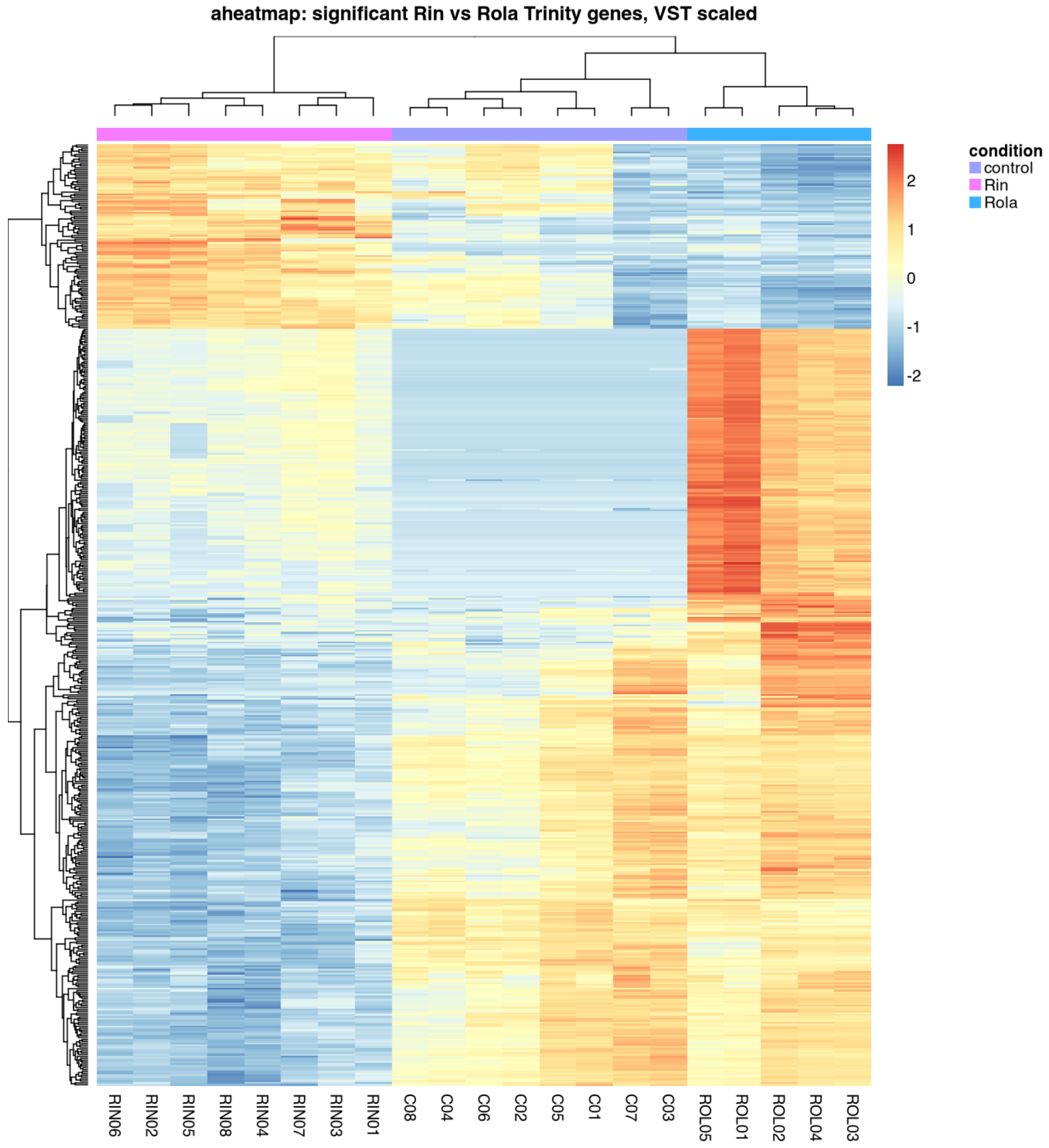


*NMF aheatmap of significant Rin vs Rola Trinity genes, if NMF and sufficient genes were available.*

#### Figure 28

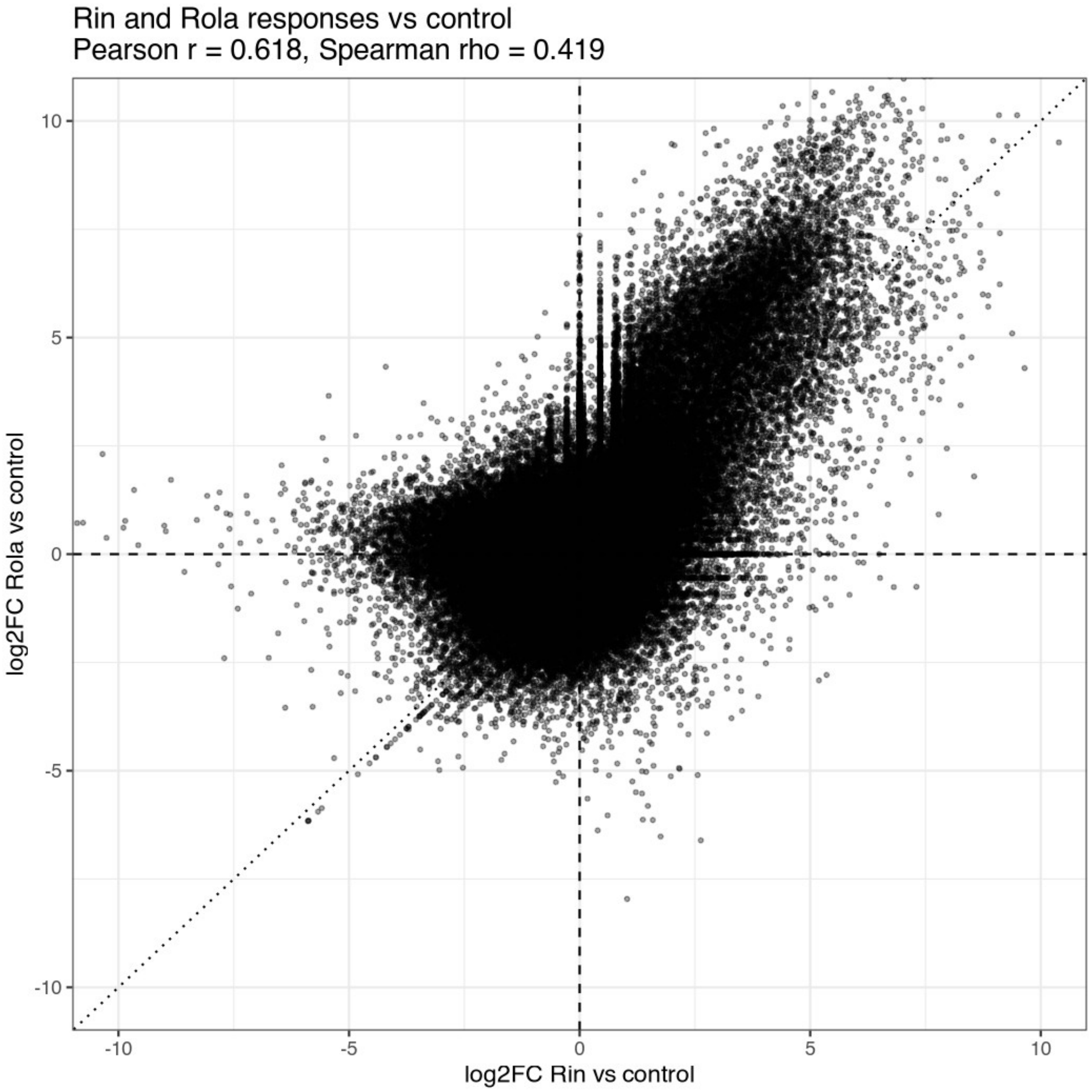


*Scatter plot comparing log2FC Rin vs control and Rola vs control.*

### Table previews

#### Table preview 1

**Metadata used for DESeq2.**

| sample | condition | batch |
| --- | --- | --- |
| C01 | control | batch2_IAB |
| C02 | control | batch2_IAB |
| C03 | control | batch2_IAB |
| C04 | control | batch2_IAB |

| sample | condition | batch |
|---|---|---|
| C05 | control | batch1_Eurofins |
| C06 | control | batch1_Eurofins |
| C07 | control | batch1_Eurofins |
| C08 | control | batch1_Eurofins |
| RIN01 | Rin | batch2_IAB |
| RIN02 | Rin | batch2_IAB |

#### Table preview 2

**One-to-one tx2gene table for Trinity gene representative sequences.**

| TXNAME | GENEID |
|---|---|
| TRINITY_DN4869_c0_g1 | TRINITY_DN4869_c0_g1 |
| TRINITY_DN4869_c1_g1 | TRINITY_DN4869_c1_g1 |
| TRINITY_DN4812_c0_g1 | TRINITY_DN4812_c0_g1 |
| TRINITY_DN4812_c1_g1 | TRINITY_DN4812_c1_g1 |
| TRINITY_DN4812_c1_g2 | TRINITY_DN4812_c1_g2 |
| TRINITY_DN4812_c2_g1 | TRINITY_DN4812_c2_g1 |
| TRINITY_DN4843_c0_g1 | TRINITY_DN4843_c0_g1 |
| TRINITY_DN4843_c0_g2 | TRINITY_DN4843_c0_g2 |
| TRINITY_DN4894_c0_g1 | TRINITY_DN4894_c0_g1 |
| TRINITY_DN4894_c1_g1 | TRINITY_DN4894_c1_g1 |

#### Table preview 3

**Rin vs Rola DESeq2 results with OmicsBox descriptions.**

| gene_id | gene_name | baseMean | log2FoldChange | lfcSE | stat | Description | Ontology_term | Ontology_id | Enzyme_code | Enzyme_name |
|---|---|---|---|---|---|---|---|---|---|---|
| T | DVH24_0 | 1 | - | 0 | - | hypoth | membrane | GO:00 | N | NA |

| gene_id | gene_name | baseMean | log2FoldChange | lfcSE | stat | Description | Ontology_term | Ontology_id | Enzyme_code | Enzyme_name |
|---|---|---|---|---|---|---|---|---|---|---|
| RINITY_DN13723_c0_g1 | 29871; DVH24_026477 | 951.74202 | 1.577304 | .1101007 | 14.32601 | etical protein DVH24_029871 | | 16020 | A | |
| TRINITY_DN51_c0_g1 | TRINITY_DN51_c0_g1 | 618.78604 | -2.187825 | 0.1525061 | -14.34582 | uncharacterized protein | NA | NA | NA | NA |
| TRINITY_ | TRINITY_DN4249_c0_g2 | 151.4 | -2.8 | 2.035 | -12.7 | protein GAST1 isoform X2 | NA | NA | NA | NA |

| gene_id | gene_name | baseMean | log2FoldChange | lfcSE | stat | Description | Ontology_term | Ontology_id | Enzyme_code | Enzyme_name |
|---|---|---|---|---|---|---|---|---|---|---|
| DN42449_c0_g2 | | 7730 | 76213 | 1337 | 1475 | | | | | |
| TRINITY_DN189936_c0_g1 | DVH24_011177; DVH24_011179 | 622.44398 | 3.954495 | 0.3152162 | 12.54534 | berberine bridge enzyme-like 8 | oxidoreductase activity,FAD binding | GO:0016491,GO:0071949 | EC:1 | Oxidoreductases |
| TRINITY_DN | DVH24_010981; DVH24_020628; DVH24_026348; DVH24_0 | 1184.77 | -1.6189 | 0.13184 | -12.278 | LOB domain-containing protein 37-like | cytoplasm,membrane,glycosyltransferase activity,regulation of gene expression | GO:0005737,GO:0016020,GO:0016757,GO | EC:2.4 | Glycosyltransferases |

| gene_id | gene_name | baseMean | log2FoldChange | lfcSE | stat | Description | Ontology_term | Ontology_id | Enzyme_code | Enzyme_name |
|---|---|---|---|---|---|---|---|---|---|---|
| 8741_c0_g1 | 12538 | 078 | 17 | 63 | 82 | | | :0010468 | | |
| TRINITY_DN31025_c0_g1 | DVH24_003302; DVH24_032401; DVH24_032402; DVH24_032651 | 140.87969 | -2.478467 | 0.2062848 | -12.01478 | GDSL esterase/lipase At2g42990 | hydrolase activity, acting on ester bonds | GO:0016788 | EC:3.1 | Acting on ester bonds |
| TRINITY_DN14 | DVH24_034924; DVH24_042310; DVH24_021501; DVH24_042312; MdSOT3 | 1999.4594 | 3.839685 | 0.3199470 | 12.00100 | putative polyol transporter 1 | membrane,carbohydrate:proton symporter activity,carbohydrate transmembrane transport,proton transmembrane transport | GO:0016020,GO:0005351,GO:0034219,GO:1902600 | NA | NA |

| gene_id | gene_name | baseMean | log2FoldChange | lfcSE | stat | Description | Ontology_term | Ontology_id | Enzyme_code | Enzyme_name |
|---|---|---|---|---|---|---|---|---|---|---|
| 66_c1_g1 | | 9 | | | | | | | | |
| TRINITY_DN6594_c1_g1 | DVH24_002005; DVH24_018718; DVH24_010436; DVH24_028654 | 290.43930 | -6.481784 | 0.5477278 | -11.83395 | probable pectate lyase 8 | pectate lyase activity,metal ion binding,pectin catabolic process | GO:0030570,GO:0046872,GO:0045490 | EC:4.2.2.2 | pectate lyase |
| TRINITY_DN53886_c0 | DVH24_033743; DVH24_033745; DVH24_040960 | 1023.00655 | -1.050814 | 0.0894867 | -11.74269 | very-long-chain 3-oxoacyl-CoA reductase 1-like | endoplasmic reticulum,membrane,ketoreductase activity | GO:0005783,GO:0016020,GO:0045703 | EC:1.1 | Acting on the CH-OH group of donors |

| gene_id | gene_name | baseMean | log2FoldChange | lfcSE | stat | Description | Ontology_term | Ontology_id | Enzyme_code | Enzyme_name |
|---|---|---|---|---|---|---|---|---|---|---|
| _g1 | | | | | | | | | | |
| TRINITY_DN36061_c1_g1 | TRINITY_DN36061_c1_g1 | 96.95591 | -25.590027 | 2.1818836 | -11.72841 | protein PELPK2-like | NA | NA | NA | NA |

#### Table preview 4

**Rin vs control DESeq2 results with OmicsBox descriptions.**

| gene_id | gene_name | Description | Ontology_term | Ontology_id | Enzyme_code | Enzyme_name |
|---|---|---|---|---|---|---|
| TRINITY_DN137233_c0_g1 | DVH24_029871; DVH24_026477 | hypothetical protein DVH24_029871 | membrane | GO:0016020 | NA | NA |
| TRINITY_D | TRINITY_DN18768_ | TRINITY_D | NA | NA | NA | NA |

| gene_id | gene_name | Description | Ontology_term | Ontology_id | Enzyme_code | Enzyme_name |
|---|---|---|---|---|---|---|
| N18768_c0_g1 | c0_g1 | N18768_c0_g1 | | | | |
| TRINITY_DN23990_c0_g1 | TRINITY_DN23990_c0_g1 | TRINITY_DN23990_c0_g1 | NA | NA | NA | NA |
| T | T | T | NA | NA | N | NA |

| gene_id | gene_name | Description | Ontology_term | Ontology_id | Enzyme_code | Enzyme_name |
|---|---|---|---|---|---|---|
| RINITY_DN16515_c0_g1 | RINITY_DN16515_c0_g1 | RINITY_DN16515_c0_g1 | | | A | |
| TRINITY_DN1724 | TRINITY_DN17241_c0_g1 | TRINITY_DN1724 | NA | NA | NA | NA |

| gene_id | gene_name | Description | Ontology_term | Ontology_id | Enzyme_code | Enzyme_name |
|---|---|---|---|---|---|---|
| 1_c0_g1 | | 1_c0_g1 | | | | |
| TRINITY_DN22704_c0_g1 | TRINITY_DN22704_c0_g1 | aldehydeoxidaseGLOX1-like | NA | NA | NA | NA |
| TRINIT | DVH24_013 | cytochr | regulation of ER to Golgi vesicle-mediated transport,iron ion binding,retrograde vesicle-mediated transport, Golgi to endoplasmic reticulum,oxidoreductase | GO:0060628,GO:0005506,GO:0006890,GO: | EC:1.1 | Acting on paired donors, with incorporation or reduction of |

| gene_id | gene_name | Description | Ontology_term | Ontology_id | Enzyme_code | Enzyme_name |
|---|---|---|---|---|---|---|
| Y_DN8783_c0_g1 | 674;DVH24_025666;DVH24_009719 | omeP450077A3-like | activity, acting on paired donors, with incorporation or reduction of molecular oxygen,Dsl1/NZR complex,membrane,monooxygenase activity,heme binding | 0016705,GO:0070939,GO:0016020,GO:0004497,GO:0020037 | 4 | molecular oxygen. The oxygen incorporated need not be derived from O2 |
| TRINITY_DN11126_c | DVH24_032726;DVH24_000363; | proteinHOTHEADisofor | oxidoreductase activity, acting on CH-OH group of donors,mandelonitrile lyase activity,flavin adenine dinucleotide binding | GO:0016614,GO:0046593,GO:0050660 | EC:1.1,EC:4.1.2. | Acting on the CH-OH group of donors,(R)-mandelonitrile lyase |

| gene_id | gene_name | Description | Ontology_term | Ontology_id | Enzyme_code | Enzyme_name |
|---|---|---|---|---|---|---|
| 3_g1 | DVH24_001284 | mX1 | | | 10 | |
| TRINITY_DN4249_c0_g2 | TRINITY_DN4249_c0_g2 | protein GAST1 isoform X2 | NA | NA | NA | NA |
| TRINIT | TRINITY_DN | unchar | NA | NA | NA | NA |

| gene_id | gene_name | Description | Ontology_term | Ontology_id | Enzyme_code | Enzyme_name |
|---|---|---|---|---|---|---|
| Y_DN51_c0_g1 | 51_c0_g1 | acterized protein | | | | |

#### Table preview 5

**Rola vs control DESeq2 results with OmicsBox descriptions.**

| gene_id | gene_name | baseMean | log2FoldChange | Description | Ontology_term | Ontology_id | Enzyme_code | Enzyme_name |
|---|---|---|---|---|---|---|---|---|
| TRIN | TRINITY_DN18768_c0_g1 | 4375 | 12.5 | TRINITY_DN18768_c0_g1 | NA | NA | NA | NA |

| gene_id | gene_name | baseMean | log2FoldChange | Description | Ontology_term | Ontology_id | Enzyme_code | Enzyme_name |
|---|---|---|---|---|---|---|---|---|
| ITY_DN18768_c0_g1 | | .061331 | 51520 | | | | | |
| TRINITY_DN16515_c0 | TRINITY_DN16515_c0_g1 | 2205.329440 | 12.976204 | TRINITY_DN16515_c0_g1 | NA | NA | NA | NA |

| gene_id | gene_name | baseMean | log2FoldChange | Description | Ontology_term | Ontology_id | Enzyme_code | Enzyme_name |
|---|---|---|---|---|---|---|---|---|
| _g1 | | | | | | | | |
| TRINITY_DN23990_c0_g1 | TRINITY_DN23990_c0_g1 | 2005.1322820 | 12.3819028 | TRINITY_DN23990_c0_g1 | NA | NA | NA | NA |
| TRINITY_DN | TRINITY_DN17241_c0_g1 | 940.1062621 | 10.6603870 | TRINITY_DN17241_c0_g1 | NA | NA | NA | NA |

| gene_id | gene_name | baseMean | log2FoldChange | Description | Ontology_term | Ontology_id | Enzyme_code | Enzyme_name |
|---|---|---|---|---|---|---|---|---|
| 172241_c0_g1 | | | | | | | | |
| TRINITY_DN81098_c0_g1 | DVH24_016150; DVH24_003921; DVH24_016152; DVH24_016153 | 124.28616 | 11.336094 | ADP,ATP carrier protein 1, mitochondrial-like | mitochondrial inner membrane,ATP:ADP antiporter activity,mitochondrial ADP transmembrane transport,mitochondrial ATP transmembrane transport | GO:0005743,GO:0005471,GO:0140021,GO:1990544 | NA | NA |
| TR | atp9; ATP9; | 97 | 10 | ATP synthase | membrane | GO:00160 | NA | NA |

| gene_id | gene_name | baseMean | log2FoldChange | Description | Ontology_term | Ontology_id | Enzyme_code | Enzyme_name |
|---|---|---|---|---|---|---|---|---|
| INITY_DN275399_c0_g1 | DVH24_038917 | .274770 | .9771022 | F0 subunit 9 | | 20 | | |
| TRINITY_DN480440 | DVH24_000798; DVH24_034180 | 59.871331 | 10.353032 | adenosylhomocysteinase | NA | NA | NA | NA |

| gene_id | gene_name | baseMean | log2FoldChange | Description | Ontology_term | Ontology_id | Enzyme_code | Enzyme_name |
|---|---|---|---|---|---|---|---|---|
| _c0_g1 | | | | | | | | |
| TRINITY_DN7445_c0_g1 | TRINITY_DN7445_c0_g1 | 281.414479 | 3.1436618 | protein ALP1-like | NA | NA | NA | NA |
| TRINITY_D | DVH24_005516; DVH24_035080; DVH24_01903 | 63.966999 | 10.420996 | 5-methyltetrahydropteroyltriglutamate–homocysteine methyltra | methyltransferase activity | GO:0008168 | EC:2.1.1 | Transferring one-carbon |

| gene_id | gene_name | baseMean | log2FoldChange | Description | Ontology_term | Ontology_id | Enzyme_code | Enzyme_name |
|---|---|---|---|---|---|---|---|---|
| N245488_c1_g1 | 5 | | 2 | nsferase | | | | groups |
| TRINITY_DN284883_c1_g1 | DVH24_038846; DVH24_042792; DVH24_005917 | 146.152273 | 11.5078692 | protein disulfide-isomerase-like | NA | NA | NA | NA |

#### Table preview 6

**Rin vs Rola DESeq2 results, backward-compatible filename.**

| gene_id | gene_name | baseMean | log2FoldChange | lfcSE | stat | Description | Ontology_term | Ontology_id | Enzyme_code | Enzyme_name |
|---|---|---|---|---|---|---|---|---|---|---|
| TRINITY_DN137723_c0_g1 | DVH24_029871; DVH24_026477 | 1951.774202 | -1.577304 | 0.1101007 | -14.32601 | hypothetical protein DVH24_029871 | membrane | GO:0016020 | NA | NA |
| TRINITY_DN51_c0_g1 | TRINITY_DN51_c0_g1 | 618.786604 | -2.187825 | 0.1525061 | -14.34582 | uncharacterized protein | NA | NA | NA | NA |

| gene_id | gene_name | baseMean | log2FoldChange | lfcSE | stat | Description | Ontology_term | Ontology_id | Enzyme_code | Enzyme_name |
|---|---|---|---|---|---|---|---|---|---|---|
| TRINITY_DN4249_c0_g2 | TRINITY_DN4249_c0_g2 | 151.47730 | -25.876213 | 2.0351337 | -12.71475 | protein GAST1 isoform X2 | NA | NA | NA | NA |
| TRINITY_DN189336_c0_g1 | DVH24_011177; DVH24_011179 | 622.44398 | 3.954495 | 0.3152162 | 12.54534 | berberine bridge enzyme-like 8 | oxidoreductase activity,FAD binding | GO:0016491,GO:0071949 | EC:1 | Oxidoreductases |
| TRI | DVH24_010981; | 11 | -1 | 0. | -1 | LOB domai | cytoplasm,membrane,glycosyltransferas | GO:0005737 | EC | Glycosylt |

| gene_id | gene_name | baseMean | log2FoldChange | lfcSE | stat | Description | Ontology_term | Ontology_id | Enzyme_code | Enzyme_name |
|---|---|---|---|---|---|---|---|---|---|---|
| NITY_DN8741_c0_g1 | DVH24_020628; DVH24_026348; DVH24_012538 | 84.7707078 | .6189917 | 13.1287463 | 2.1278882 | n-containing protein 37-like | e activity,regulation of gene expression | ,GO:0016020,GO:0016757,GO:0010468 | :2.4 | ransferases |
| TRINITY_DN3102 5_c0_g1 | DVH24_003302; DVH24_032401; DVH24_032402; DVH24_032651 | 140.8796 69 | -2.4784 67 | 0.2206218 48 | -12.201478 | GDSL esterase/lipase At2g42990 | hydrolase activity, acting on ester bonds | GO:0016788 | EC:3.1 | Acting on ester bonds |
| TRINIT | DVH24_034924; DVH24_042310; | 1999 | 3.83 | 0.31 | 12.0 | putative polyol transp | membrane,carbohydrate:proton symporter activity,carbohydrat | GO:0016020,GO:000535 | NA | NA |

| gene_id | gene_name | baseMean | log2FoldChange | lfcSE | stat | Description | Ontology_term | Ontology_id | Enzyme_code | Enzyme_name |
|---|---|---|---|---|---|---|---|---|---|---|
| Y_DN14 66_c1_g1 | DVH24_021501; DVH24_042312; MdSOT3 | .459 49 | 9685 | 99470 0 | 0100 | orter 1 | e transmembrane transport,proton transmembrane transport | 1,GO:0034219,GO:1902600 | | |
| TRINITY_DN6594_c1_g1 | DVH24_002005; DVH24_018718; DVH24_010436; DVH24_028654 | 290.4439 30 | -6.4817 84 | 0.5477 278 | -11.833 95 | probable pectate lyase 8 | pectate lyase activity,metal ion binding,pectin catabolic process | GO:0030570,GO:0046872,GO:0045490 | EC:4.2.2.2 | pectate lyase |
| TRINITYDN | DVH24_033743; DVH24_033745; DVH24_040960 | 1023.00 | -1.05 08 | 0.08 948 | -11.742 | very-long-chain 3-oxoacyl-CoA reduct | endoplasmic reticulum,membrane,ketoreductase activity | GO:0005783,GO:0016020,GO:0045703 | EC:1.1 | Acting on the CH-OH grou |

| gene_id | gene_name | baseMean | log2FoldChange | lfcSE | stat | Description | Ontology_term | Ontology_id | Enzyme_code | Enzyme_name |
|---|---|---|---|---|---|---|---|---|---|---|
| 53886_c0_g1 | | 655 | 14 | 67 | 69 | ase 1-like | | | | p of donors |
| TRINITY_DN36061_c1_g1 | TRINITY_DN36061_c1_g1 | 96.95591 | -25.590027 | 2.1818836 | -11.72841 | protein PELPK 2-like | NA | NA | NA | NA |

#### Table preview 7

**Rin vs control DESeq2 results, backward-compatible filename.**

| gene_id | gene_name | Description | Ontology_term | Ontology_id | Enzyme_code | Enzyme_name |
|---|---|---|---|---|---|---|
| TRINITY_DN13723_c0_g1 | DVH24_029871;DVH24_026477 | hypothetical protein DVH24_029871 | membrane | GO:0016020 | NA | NA |
| TRINITY_D | TRINITY_DN18768_ | TRINITY_D | NA | NA | NA | NA |

| gene_id | gene_name | Description | Ontology_term | Ontology_id | Enzyme_code | Enzyme_name |
|---|---|---|---|---|---|---|
| N187688_c0_g1 | c0_g1 | N187688_c0_g1 | | | | |
| TRINITY_DN23990_c0_g1 | TRINITY_DN23990_c0_g1 | TRINITY_DN23990_c0_g1 | NA | NA | NA | NA |
| T | T | T | NA | NA | N | NA |

| gene_id | gene_name | Description | Ontology_term | Ontology_id | Enzyme_code | Enzyme_name |
|---|---|---|---|---|---|---|
| RINITY_DN16515_c0_g1 | RINITY_DN16515_c0_g1 | RINITY_DN16515_c0_g1 | | | A | |
| TRINITY_DN1724 | TRINITY_DN17241_c0_g1 | TRINITY_DN1724 | NA | NA | NA | NA |

| gene_id | gene_name | Description | Ontology_term | Ontology_id | Enzyme_code | Enzyme_name |
|---|---|---|---|---|---|---|
| 1_c0_g1 | | 1_c0_g1 | | | | |
| TRINITY_DN22704_c0_g1 | TRINITY_DN22704_c0_g1 | aldehydeoxidaseGLOX1-like | NA | NA | NA | NA |
| TRINIT | DVH24_013 | cytochr | regulation of ER to Golgi vesicle-mediated transport,iron ion binding,retrograde vesicle-mediated transport, Golgi to endoplasmic reticulum,oxidoreductase | GO:0060628,GO:0005506,GO:0006890,GO: | EC:1.1 | Acting on paired donors, with incorporation or reduction of |

| gene_id | gene_name | Description | Ontology_term | Ontology_id | Enzyme_code | Enzyme_name |
|---|---|---|---|---|---|---|
| Y_DN8783_c0_g1 | 674;DVH24_025666;DVH24_009719 | omeP45077A3-like | activity, acting on paired donors, with incorporation or reduction of molecular oxygen,Dsl1/NZR complex,membrane,monooxygenase activity,heme binding | 0016705,GO:0070939,GO:0016020,GO:0004497,GO:0020037 | 4 | molecular oxygen. The oxygen incorporated need not be derived from O2 |
| TRINITY_DN11126_c | DVH24_032726;DVH24_000363; | proteinHOTHEADisofor | oxidoreductase activity, acting on CH-OH group of donors,mandelonitrile lyase activity,flavin adenine dinucleotide binding | GO:0016614,GO:0046593,GO:0050660 | EC:1.1,EC:4.1.2. | Acting on the CH-OH group of donors,(R)-mandelonitrile lyase |

| gene_id | gene_name | Description | Ontology_term | Ontology_id | Enzyme_code | Enzyme_name |
|---|---|---|---|---|---|---|
| 3_g1 | DVH24_00128 4 | mX1 | | | 10 | |
| TRINITY_DN4249_c0_g2 | TRINITY_DN4249_c0_g2 | proteinGAST1isoformX2 | NA | NA | NA | NA |
| TRINIT | TRINITY_DN | unchar | NA | NA | NA | NA |

| gene_id | gene_name | Description | Ontology_term | Ontology_id | Enzyme_code | Enzyme_name |
|---|---|---|---|---|---|---|
| Y_DN51_c0_g1 | 51_c0_g1 | acterized protein | | | | |

#### Table preview 8

**Rola vs control DESeq2 results, backward-compatible filename.**

| gene_id | gene_name | baseMean | log2FoldChange | Description | Ontology_term | Ontology_id | Enzyme_code | Enzyme_name |
|---|---|---|---|---|---|---|---|---|
| TRIN | TRINITY_DN18768_c0_g1 | 4375 | 12.5 | TRINITY_DN18768_c0_g1 | NA | NA | NA | NA |

| gene_id | gene_name | baseMean | log2FoldChange | Description | Ontology_term | Ontology_id | Enzyme_code | Enzyme_name |
|---|---|---|---|---|---|---|---|---|
| ITY_DN187668_c0_g1 | | .061331 | 51520 | | | | | |
| TRINITY_DN16515_c0 | TRINITY_DN16515_c0_g1 | 2205.329440 | 12.976204 | TRINITY_DN16515_c0_g1 | NA | NA | NA | NA |

| gene_id | gene_name | baseMean | log2FoldChange | Description | Ontology_term | Ontology_id | Enzyme_code | Enzyme_name |
|---|---|---|---|---|---|---|---|---|
| _g1 | | | | | | | | |
| TRINITY_DN23990_c0_g1 | TRINITY_DN23990_c0_g1 | 2005.132220 | 12.381908 | TRINITY_DN23990_c0_g1 | NA | NA | NA | NA |
| TRINITY_DN | TRINITY_DN17241_c0_g1 | 940.10621 | 10.660387 | TRINITY_DN17241_c0_g1 | NA | NA | NA | NA |

| gene_id | gene_name | baseMean | log2FoldChange | Description | Ontology_term | Ontology_id | Enzyme_code | Enzyme_name |
|---|---|---|---|---|---|---|---|---|
| 172441_c0_g1 | | | | | | | | |
| TRINITY_DN81098_c0_g1 | DVH24_016150; DVH24_003921; DVH24_016152; DVH24_016153 | 124.328616 | 11.336094 | ADP,ATP carrier protein 1, mitochondrial-like | mitochondrial inner membrane,ATP:ADP antiporter activity,mitochondrial ADP transmembrane transport,mitochondrial ATP transmembrane transport | GO:0005743,GO:0005471,GO:0140021,GO:1990544 | NA | NA |
| TR | atp9; ATP9; | 97 | 10 | ATP synthase | membrane | GO:00160 | NA | NA |

| gene_id | gene_name | baseMean | log2FoldChange | Description | Ontology_term | Ontology_id | Enzyme_code | Enzyme_name |
|---|---|---|---|---|---|---|---|---|
| INITY_DN27539_c0_g1 | DVH24_038917 | .274770 | .9771002 | F0 subunit 9 | | 20 | | |
| TRINITY_DN48040 | DVH24_000798; DVH24_034180 | 59.871331 | 10.353032 | adenosylhomocysteinase | NA | NA | NA | NA |

| gene_id | gene_name | baseMean | log2FoldChange | Description | Ontology_term | Ontology_id | Enzyme_code | Enzyme_name |
|---|---|---|---|---|---|---|---|---|
| _c0_g1 | | | | | | | | |
| TRINITY_DN7445_c0_g1 | TRINITY_DN7445_c0_g1 | 281.414789 | 3.143618 | protein ALP1-like | NA | NA | NA | NA |
| TRINITY_D | DVH24_005516; DVH24_035080; DVH24_01903 | 63.966999 | 10.420996 | 5-methyltetrahydropteroyltriglutamate–homocysteine methyltra | methyltransferase activity | GO:0008168 | EC:2.1.1 | Transferring one-carbon |

| gene_id | gene_name | baseMean | log2FoldChange | Description | Ontology_term | Ontology_id | Enzyme_code | Enzyme_name |
|---|---|---|---|---|---|---|---|---|
| N245548_c1_g1 | 5 | | 2 | nsferase | | | | groups |
| TRINITY_DN284883_c1_g1 | DVH24_038846; DVH24_042792; DVH24_005917 | 146.152273 | 11.507869 | protein disulfide-isomerase-like | NA | NA | NA | NA |

#### Table preview 9

**Summary counts of significant DE Trinity genes.**

| contrast | padj_0.05 | padj_0.05_absLFC1 |
|---|---|---|
| Rin_vs_Rola | 18121 | 13382 |
| Rin_vs_control | 20107 | 14939 |
| Rola_vs_control | 15987 | 12860 |

#### Table preview 10

**DESeq2 normalized count matrix.**

| | C01 | C02 | C03 | C04 | C05 | C06 | C07 | C08 | RIN01 | RIN02 | RIN03 | RIN04 | RIN05 | RIN06 | RIN07 | RIN08 | ROL01 | ROL02 | ROL03 | ROL04 | ROL05 |
|---|---|---|---|---|---|---|---|---|---|---|---|---|---|---|---|---|---|---|---|---|---|
| TRINITY_DN0_c0_g1 | 11.652359 | 14.289025 | 18.007192 | 10.001155 | 0.00000 | 6.381117 | 5.027082 | 0.000000 | 14.865941 | 9.174574 | 6.130714 | 5.8849381 | 0.00000 | 1.692673 | 1.815308 | 5.835163 | 6.158155 | 11.942571 | 9.764733 | 5.787301 | 4.428652 |
| TRINITY_DN0_c0_g2 | 1.588958 | 2.956350 | 1.637018 | 2.307959 | 5.43535 | 3.190558 | 0.000000 | 3.767386 | 1.651771 | 3.058191 | 1.021786 | 0.7356173 | 0.00000 | 5.007018 | 0.000000 | 1.458791 | 1.944681 | 1.377989 | 3.254911 | 0.000000 | 0.000000 |
| TRINITY_DN1_c0_g1 | 1809.82316 | 2720.33470 | 1138.27282 | 2227.94962 | 1683.14658 | 2638.59190 | 1075.79565 | 2194.50227 | 2566.02668 | 4269.84677 | 2102.32392 | 3249.221473 | 4077.78291 | 4223.21863 | 2207.41495 | 3441.28758 | 1382.34373 | 911.769389 | 745.374634 | 811.669015 | 1347.78654 |

| | C01 | C02 | C03 | C04 | C05 | C06 | C07 | C08 | RIN01 | RIN02 | RIN03 | RIN04 | RIN05 | RIN06 | RIN07 | RIN08 | ROL01 | ROL02 | ROL03 | ROL04 | ROL05 |
|---|---|---|---|---|---|---|---|---|---|---|---|---|---|---|---|---|---|---|---|---|---|
| | 5 | 4 | 5 | 9 | | 5 | 6 | 4 | 8 | 0 | 0 | 0 | | 1 | 1 | 6 | 7 | | | | 0 |
| TRINITY_DN10_c0_g1 | 409.421512 | 567.619196 | 430.535598 | 646.997803 | 221.03755 | 330.222808 | 283.192315 | 331.529957 | 480.665443 | 633.045611 | 347.918005 | 592.1719008 | 329.42381 | 353.768615 | 216.021693 | 366.156500 | 380.185042 | 229.205504 | 148.098454 | 162.044438 | 253.909403 |
| TRINITY_DN100_c0_g1 | 38.664645 | 40.403450 | 13.641812 | 166.942358 | 34.42388 | 43.072540 | 5.027082 | 145.044356 | 160.221814 | 121.104378 | 303.470330 | 72.8261095 | 76.30721 | 59.243548 | 230.544160 | 74.398333 | 53.154601 | 13.320560 | 6.509822 | 5.787301 | 35.429219 |
| TRINITY_DN100_c1_g1 | 86.333385 | 64.546975 | 259.194435 | 523.137344 | 52.54171 | 52.644216 | 192.704829 | 408.761368 | 126.360503 | 64.833657 | 18.392141 | 341.3264124 | 67.00145 | 32.160783 | 12.707158 | 259.664769 | 134.831183 | 135.961581 | 118.804254 | 88.256346 | 97.430352 |
| TRINITY_DN1000_c0_g1 | 33.89777 | 67.99604 | 13.09614 | 136.1695 | 21.74140 | 43.07254 | 8.378471 | 114.9052 | 434.4158 | 253.8298 | 1156.661 | 294.98252 | 254.9777 | 223.4328 | 1054.694 | 268.4175 | 391.8531 | 183.7318 | 139.9611 | 166.3849 | 349.8635 |

| | C01 | C02 | C03 | C04 | C05 | C06 | C07 | C08 | RIN01 | RIN02 | RIN03 | RIN04 | RIN05 | RIN06 | RIN07 | RIN08 | ROL01 | ROL02 | ROL03 | ROL04 | ROL05 |
|---|---|---|---|---|---|---|---|---|---|---|---|---|---|---|---|---|---|---|---|---|---|
| | 1 | 9 | 0 | 73 | | 0 | | 69 | 47 | 82 | 326 | 45 | 5 | 09 | 150 | 14 | 25 | 67 | 76 | 14 | 38 |
| TRINITY_DN10000_c0_g1 | 33.368118 | 28.578050 | 39.834092 | 26.156867 | 12.68248 | 46.263099 | 23.459718 | 22.604315 | 32.209540 | 22.630616 | 33.208033 | 36.7808634 | 18.61151 | 23.697419 | 19.968392 | 11.670327 | 24.632620 | 30.315758 | 27.666744 | 18.808729 | 23.619479 |
| TRINITY_DN10000_c0_g2 | 78.388595 | 70.459675 | 69.846079 | 53.852374 | 132.26017 | 132.408179 | 93.838873 | 105.486805 | 37.990739 | 51.989253 | 57.730888 | 49.2863570 | 81.89067 | 118.487096 | 127.071584 | 59.810424 | 58.016302 | 58.334868 | 118.804254 | 125.873805 | 110.716309 |
| TRINITY_DN100005_c0_g1 | 11.122706 | 4.434525 | 3.819707 | 2.307959 | 19.92961 | 9.571676 | 6.702777 | 7.534772 | 1.651771 | 4.893106 | 3.065357 | 4.4137036 | 11.16691 | 5.078018 | 3.630617 | 2.917582 | 2.592907 | 4.593297 | 14.647100 | 0.000000 | 4.428652 |

#### Table preview 11

**PCA scores from the condition-only analysis.**

| PC1 | PC2 | PC3 | PC4 | PC5 | sample | condition |
|---|---|---|---|---|---|---|
| -35.20076 | 99.89176 | 43.46334 | 84.748291 | 29.022414 | C01 | control |

| PC1 | PC2 | PC3 | PC4 | PC5 | sample | condition |
|---:|---:|---:|---:|---:|---|---|
| -67.71105 | 63.68873 | 51.04690 | 54.646482 | 16.337187 | C02 | control |
| 45.03216 | 142.16021 | 67.16305 | 18.926714 | -37.910416 | C03 | control |
| -74.86571 | 42.78950 | 90.27091 | -40.529300 | 10.592668 | C04 | control |
| -34.00420 | 112.54325 | -64.11731 | 48.136238 | 66.334497 | C05 | control |
| -67.61183 | 73.37044 | -63.23596 | 8.826754 | 49.795486 | C06 | control |
| 53.18053 | 157.57396 | -40.49545 | -34.254773 | -25.732191 | C07 | control |
| -73.39816 | 51.21549 | -24.39734 | -108.793844 | 40.572974 | C08 | control |
| -30.45531 | -72.37507 | 64.91484 | -21.473088 | -3.435924 | RIN01 | Rin |
| -94.11309 | -56.63379 | 32.75128 | 60.771807 | -52.951222 | RIN02 | Rin |

#### Table preview 12

**VST expression matrix.**

| | C01 | C02 | C03 | C04 | C05 | C06 | C07 | C08 | RIN01 | RIN02 | RIN03 | RIN04 | RIN05 | RIN06 | RIN07 | RIN08 | ROL01 | ROL02 | ROL03 | ROL04 | ROL05 |
|---|---|---|---|---|---|---|---|---|---|---|---|---|---|---|---|---|---|---|---|---|---|
| TRINITY_DN0_c0_g1 | 4.680617 | 4.835919 | 5.025911 | 4.571699 | 3.063259 | 4.286799 | 4.155800 | 3.063259 | 4.867382 | 4.512967 | 4.263872 | 4.240836 | 3.063259 | 3.707126 | 3.729650 | 4.236104 | 4.266410 | 4.698742 | 4.555213 | 4.231531 | 4.091490 |
| TRINITY_DN0_c0_g2 | 3.687 | 3.909 | 3.696 | 3.812 | 4.197 | 3.940 | 3.063 | 4.014 | 3.699 | 3.923 | 3.565 | 3.489 | 3.063 | 4.161 | 3.063 | 3.661 | 3.752 | 3.645 | 3.949 | 3.063 | 3.063 |

| | C001 | C002 | C003 | C004 | C005 | C006 | C007 | C008 | RIN001 | RIN002 | RIN003 | RIN004 | RIN005 | RIN006 | RIN007 | RIN008 | ROL001 | ROL002 | ROL003 | ROL004 | ROL005 |
|---|---|---|---|---|---|---|---|---|---|---|---|---|---|---|---|---|---|---|---|---|---|
| | 400 | 090 | 622 | 890 | 223 | 995 | 259 | 498 | 425 | 126 | 139 | 695 | 259 | 069 | 259 | 665 | 560 | 083 | 537 | 259 | 259 |
| TRINITY_DN1_c0_g1 | 10.834868 | 11.418394 | 10.173589 | 11.132266 | 10.731169 | 11.374650 | 10.093350 | 11.110606 | 11.334674 | 12.065601 | 11.049173 | 11.673272 | 11.999466 | 12.049821 | 11.119006 | 11.755716 | 10.450192 | 9.858620 | 9.573647 | 9.694013 | 10.414107 |
| TRINITY_DN10_c0_g1 | 8.734624 | 9.190360 | 8.804443 | 9.374191 | 7.891544 | 8.437704 | 8.227261 | 8.443136 | 8.957805 | 9.343530 | 8.509551 | 9.249764 | 8.434374 | 8.532540 | 7.860704 | 8.580037 | 8.631999 | 7.940391 | 7.360984 | 7.478728 | 8.078781 |
| TRINITY_DN100_c0_g1 | 5.757294 | 5.804005 | 4.799624 | 7.517892 | 5.636228 | 5.872765 | 4.155800 | 7.333857 | 7.463878 | 7.101322 | 8.321789 | 6.471775 | 6.527745 | 6.229387 | 7.948241 | 6.497327 | 6.105534 | 4.781195 | 4.298381 | 4.231531 | 5.665912 |
| TRINITY_DN100_c1_g1 | 6.677622 | 6.329036 | 8.106737 | 9.076081 | 6.092440 | 6.094638 | 7.707966 | 8.732385 | 7.155706 | 6.334227 | 5.044028 | 8.483204 | 6.372876 | 5.566941 | 4.745198 | 8.109198 | 7.239225 | 7.250012 | 7.076857 | 6.704646 | 6.826957 |
| TRINITY_DN10 | 5.6 | 6.3 | 4.7 | 7.2 | 5.1 | 5.8 | 4.4 | 7.0 | 8.8 | 8.0 | 10. | 8.2 | 8.0 | 7.9 | 10. | 8.1 | 8.6 | 7.6 | 7.2 | 7.5 | 8.5 |

| | C001 | C002 | C003 | C004 | C005 | C006 | C007 | C008 | RIN001 | RIN002 | RIN003 | RIN004 | RIN005 | RIN006 | RIN007 | RIN008 | ROL001 | ROL002 | ROL003 | ROL004 | ROL005 |
|---|---|---|---|---|---|---|---|---|---|---|---|---|---|---|---|---|---|---|---|---|---|
| 00_c0_g1 | 20432 | 90260 | 68151 | 51988 | 91677 | 72765 | 53277 | 34408 | 16914 | 78357 | 196380 | 82974 | 84476 | 06041 | 065209 | 54258 | 73836 | 44577 | 87538 | 13489 | 17236 |
| TRINITY_DN10000_c0_g1 | 5.604343 | 5.449440 | 5.788884 | 5.363778 | 4.743727 | 5.950717 | 5.261356 | 5.227133 | 5.568471 | 5.228199 | 5.599443 | 5.704832 | 5.054238 | 5.270709 | 5.115633 | 4.681747 | 5.306876 | 5.507710 | 5.417839 | 5.063347 | 5.267650 |
| TRINITY_DN10000_c0_g2 | 6.560186 | 6.432423 | 6.422039 | 6.120294 | 7.214386 | 7.215828 | 6.780316 | 6.926350 | 5.738756 | 6.080531 | 6.199623 | 6.020757 | 6.613151 | 7.073451 | 7.162907 | 6.240381 | 6.205287 | 6.211583 | 7.076857 | 7.150756 | 6.987353 |
| TRINITY_DN100005_c0_g1 | 4.646792 | 4.092144 | 4.020850 | 3.812890 | 5.113919 | 4.541569 | 4.315497 | 4.386177 | 3.699425 | 4.141793 | 3.924104 | 4.089823 | 4.649653 | 4.161069 | 3.997661 | 3.903679 | 3.856749 | 4.109646 | 4.855543 | 3.063259 | 4.091490 |

#### Table preview 13

Missing table file: /Volumes/Data_Disk/Ametyst_re-analysis/Trinity/DESeq2/QC_extra_plots/rlog_expression_matrix.csv

#### Table preview 14

**Mean-SD summary by transformation.**

| transformation | mean | sd |
|---|---|---|
| log2 raw counts + 1 | 2.775882 | 1.6767883 |
| log2 raw counts + 1 | 1.434499 | 0.9431727 |
| log2 raw counts + 1 | 10.948517 | 1.1108990 |
| log2 raw counts + 1 | 8.424697 | 1.2713204 |
| log2 raw counts + 1 | 5.637903 | 1.9065101 |
| log2 raw counts + 1 | 6.693845 | 1.6823348 |
| log2 raw counts + 1 | 7.181445 | 2.1062472 |
| log2 raw counts + 1 | 4.740053 | 1.1190431 |
| log2 raw counts + 1 | 6.348150 | 0.5875451 |
| log2 raw counts + 1 | 2.576938 | 0.9556311 |

#### Table preview 15

**Pearson correlation matrix based on VST values.**

| | C01 | C02 | C03 | C04 | C05 | C06 | C07 | C08 | RIN01 | RIN02 | RIN03 | RIN04 | RIN05 | RIN06 | RIN07 | RIN08 | ROL01 | ROL02 | ROL03 | ROL04 | ROL05 |
|---|---|---|---|---|---|---|---|---|---|---|---|---|---|---|---|---|---|---|---|---|---|
| C01 | 1.0000000 | 0.9865386 | 0.9711644 | 0.9663213 | 0.9750045 | 0.9669222 | 0.9520649 | 0.9483964 | 0.9463787 | 0.9555020 | 0.9365323 | 0.9408829 | 0.9382327 | 0.9427406 | 0.9145569 | 0.9313024 | 0.9292413 | 0.9469902 | 0.9249156 | 0.9286184 | 0.9150429 |
| C02 | 0.9 | 1.0 | 0.9 | 0.9 | 0.9 | 0.9 | 0.9 | 0.9 | 0.9 | 0.9 | 0.9 | 0.9 | 0.9 | 0.9 | 0.9 | 0.9 | 0.9 | 0.9 | 0.9 | 0.9 | 0.9 |

| | C001 | C002 | C003 | C004 | C005 | C006 | C007 | C008 | RIN001 | RIN002 | RIN003 | RIN004 | RIN005 | RIN006 | RIN007 | RIN008 | ROL001 | ROL002 | ROL003 | ROL004 | ROL005 |
|---|---|---|---|---|---|---|---|---|---|---|---|---|---|---|---|---|---|---|---|---|---|
| | 865386 | 000000 | 656328 | 773835 | 681049 | 730279 | 463643 | 585284 | 603058 | 681268 | 445118 | 575594 | 507394 | 553371 | 227927 | 485544 | 323882 | 437463 | 207151 | 247939 | 177627 |
| C03 | 0.9711644 | 0.9656328 | 1.0000000 | 0.9580215 | 0.9536251 | 0.9469357 | 0.9733264 | 0.9409609 | 0.9350719 | 0.9298389 | 0.9128132 | 0.9261912 | 0.9139700 | 0.9180865 | 0.8922500 | 0.9177178 | 0.9260114 | 0.9610673 | 0.9395599 | 0.9437976 | 0.9127533 |
| C04 | 0.9663213 | 0.9773835 | 0.9580215 | 1.0000000 | 0.9479654 | 0.9566787 | 0.9387018 | 0.9713044 | 0.9672013 | 0.9605679 | 0.9360279 | 0.9674896 | 0.9421613 | 0.9464143 | 0.9142860 | 0.9578254 | 0.9341307 | 0.9400082 | 0.9168973 | 0.9203866 | 0.9191143 |
| C05 | 0.9750045 | 0.9681049 | 0.9536251 | 0.9479654 | 1.0000000 | 0.9709060 | 0.9564050 | 0.9521140 | 0.9316094 | 0.9383836 | 0.9226057 | 0.9266124 | 0.9429924 | 0.9448406 | 0.9217608 | 0.9299506 | 0.9166879 | 0.9319982 | 0.9321314 | 0.9341948 | 0.9196704 |
| C06 | 0.9669 | 0.9730 | 0.9469 | 0.9566 | 0.9709 | 1.0000 | 0.9511 | 0.9627 | 0.9445 | 0.9491 | 0.9292 | 0.9420 | 0.9557 | 0.9572 | 0.9308 | 0.9465 | 0.9174 | 0.9263 | 0.9286 | 0.9306 | 0.9226 |

| | C01 | C02 | C03 | C04 | C05 | C06 | C07 | C08 | RIN01 | RIN02 | RIN03 | RIN04 | RIN05 | RIN06 | RIN07 | RIN08 | ROL01 | ROL02 | ROL03 | ROL04 | ROL05 |
|---|---|---|---|---|---|---|---|---|---|---|---|---|---|---|---|---|---|---|---|---|---|
| | 222 | 279 | 357 | 787 | 060 | 000 | 366 | 779 | 090 | 926 | 891 | 129 | 934 | 890 | 711 | 369 | 851 | 948 | 914 | 593 | 476 |
| C07 | 0.9520649 | 0.9463643 | 0.9733264 | 0.9387018 | 0.9564050 | 0.9511366 | 1.0000000 | 0.9452141 | 0.9192608 | 0.9115054 | 0.8976617 | 0.9109538 | 0.9183861 | 0.9198009 | 0.8997852 | 0.9160436 | 0.9118280 | 0.9444879 | 0.9472825 | 0.9496870 | 0.9171875 |
| C08 | 0.9483964 | 0.9585284 | 0.9409609 | 0.9713044 | 0.9521140 | 0.9627779 | 0.9452141 | 1.0000000 | 0.9528305 | 0.9427091 | 0.9223344 | 0.9530131 | 0.9488809 | 0.9497534 | 0.9234607 | 0.9575038 | 0.9205983 | 0.9243165 | 0.9265419 | 0.9279936 | 0.9249658 |
| RIN01 | 0.9463787 | 0.9603058 | 0.9350719 | 0.9672013 | 0.9316094 | 0.9445090 | 0.9192608 | 0.9528305 | 1.0000000 | 0.9636573 | 0.9703963 | 0.9752324 | 0.9469021 | 0.9512398 | 0.9486078 | 0.9652892 | 0.9531901 | 0.9461324 | 0.9272211 | 0.9310482 | 0.9412736 |
| RIN02 | 0.9555020 | 0.9681268 | 0.9298389 | 0.9605679 | 0.9383836 | 0.9491926 | 0.9115054 | 0.9427091 | 0.9636573 | 1.0000000 | 0.9619220 | 0.9676302 | 0.9681223 | 0.9724150 | 0.9376846 | 0.9559254 | 0.9348628 | 0.9301528 | 0.9075593 | 0.9113953 | 0.9197219 |

#### Table preview 16

Missing table file:
/Volumes/Data_Disk/Ametyst_re-analysis/Trinity/DESeq2/QC_extra_plots/Pearson_correlation_rlog_matrix.csv

#### Table preview 17

**Input table for the custom MA plot.**

| gene_id | gene_name | baseMean | log2FoldChange | lfcSE | stat | Description | Ontology_term | Ontology_id | Enzyme_code | Enzyme_name | significant | baseMean_for_plot |
|---|---|---|---|---|---|---|---|---|---|---|---|---|
| TRINITY_DN137723_c0_g1 | DVH24_029871; DVH24_026477 | 1951.74202 | -1.577304 | 0.1101007 | -14.32601 | hypothetical protein DVH24_029871 | membrane | GO:0016020 | NA | NA | TRUE | 1951.74202 |

| gene_id | gene_name | baseMean | log2FoldChange | lfcSE | stat | Description | Ontology_term | Ontology_id | Enzyme_code | Enzyme_name | significant | baseMean_for_plot |
|---|---|---|---|---|---|---|---|---|---|---|---|---|
| TRINITY_DN51_c0_g1 | TRINITY_DN51_c0_g1 | 618.786604 | -2.1878225 | 0.15250661 | -14.34582 | uncharacterized protein | NA | NA | NA | NA | TRUE | 618.786604 |
| TRINITY_DN424 | TRINITY_DN4249_c0_g2 | 151.477730 | -25.8762013 | 2.03513377 | -12.7147 5 | protein GAST1 isoform X2 | NA | NA | NA | NA | TRUE | 151.477730 |

| gene_id | gene_name | baseMean | log2FoldChange | lfcSE | stat | Description | Ontology_term | Ontology_id | Enzyme_code | Enzyme_name | significant | baseMean_for_plot |
|---|---|---|---|---|---|---|---|---|---|---|---|---|
| 9_c0_g2 | | | | | | | | | | | | |
| TRINITY_DN18936_c0_g1 | DVH24_011177; DVH24_011179 | 622.44398 | 3.954495 | 0.3152162 | 12.54534 | berberine bridge enzyme-like 8 | oxidoreductase activity,FAD binding | GO:0016491,GO:0071949 | EC:1 | Oxidoreductases | TRUE | 622.44398 |
| TRI | DVH24_010981; DVH24_0 | 118 | -1. | 0.1 | -12 | LOB domain- | cytoplasm,membrane,glycosyltransferase | GO:0005737,G | EC: | Glycosyltran | TRU | 118 |

| gene_id | gene_name | baseMean | log2FoldChange | lfcSE | stat | Description | Ontology_term | Ontology_id | Enzyme_code | Enzyme_name | significant | baseMean_for_plot |
|---|---|---|---|---|---|---|---|---|---|---|---|---|
| NITY_DN8741_c0_g1 | 20628; DVH24_026348; DVH24_012538 | 4.770778 | 618917 | 318463 | .27882 | containing protein 37-like | activity,regulation of gene expression | O:0016020, GO:0016757,GO:0010468 | 2.4 | sferases | E | 4.770778 |
| TRINITY_DN3102 | DVH24_003302; DVH24_032401; DVH24_032402; DVH24_032651 | 140.87969 | -2.478467 | 0.2062848 | -12.01478 | GDSL esterase/lipase At2g42990 | hydrolase activity, acting on ester bonds | GO:0016788 | EC:3.1 | Acting on ester bonds | TRUE | 140.87969 |

| gene_id | gene_name | baseMean | log2FoldChange | lfcSE | stat | Description | Ontology_term | Ontology_id | Enzyme_code | Enzyme_name | significant | baseMean_for_plot |
|---|---|---|---|---|---|---|---|---|---|---|---|---|
| 5_c0_g1 | | | | | | | | | | | | |
| TRINITY_DN1466_c1_g1 | DVH24_034924; DVH24_042310; DVH24_021501; DVH24_042312; MdSOT3 | 1999.45949 | 3.839685 | 0.3199470 | 12.00100 | putative polyol transporter 1 | membrane,carbohydrate:proton symporter activity,carbohydrate transmembrane transport,proton transmembrane transport | GO:0016020,GO:0005351,GO:0034219,GO:1902600 | NA | NA | TRUE | 1999.45949 |
| TRIN | DVH24_002005; DVH24_018718; | 290. | -6.4 | 0.54 | -11. | probable pectate | pectate lyase activity,metal ion binding,pectin catabolic process | GO:0030570,GO:004 | EC:4 | pectate lyase | TRUE | 290. |

| gene_id | gene_name | baseMean | log2FoldChange | lfcSE | stat | Description | Ontology_term | Ontology_id | Enzyme_code | Enzyme_name | significant | baseMean_for_plot |
|---|---|---|---|---|---|---|---|---|---|---|---|---|
| ITY_DN65594_c1_g1 | DVH24_010436; DVH24_028654 | 43930 | 81784 | 77278 | 83395 | lyase 8 | | 6872, GO:0045490 | .2.2.2 | | | 43930 |
| TRINITY_DN53886_c | DVH24_033743; DVH24_033745; DVH24_040960 | 1023.00655 | -1.050814 | 0.0894867 | -11.74269 | very-long-chain 3-oxoacyl-CoA reductase 1-like | endoplasmic reticulum,membrane,ketoreductase activity | GO:0005783,GO:0016020,GO:0045703 | EC:1.1 | Acting on the CH-OH group of donors | TRUE | 1023.00655 |

| gene_id | gene_name | baseMean | log2FoldChange | lfcSE | stat | Description | Ontology_term | Ontology_id | Enzyme_code | Enzyme_name | significant | baseMean_for_plot |
|---|---|---|---|---|---|---|---|---|---|---|---|---|
| 0_g1 | | | | | | | | | | | | |
| TRINITY_DN36061_c1_g1 | TRINITY_DN36061_c1_g1 | 96.95591 | -25.590027 | 2.1818836 | -11.72841 | protein PELP K2-like | NA | NA | NA | NA | TRUE | 96.95591 |

#### Table preview 18

**Input table for the volcano plot.**

| gene_id | gene_name | baseMean | log2FoldChange | lfcSE | stat | Description | Ontology_term | Ontology_id | Enzyme_code | Enzyme_name | direction | neg_log10_padj |
|---|---|---|---|---|---|---|---|---|---|---|---|---|
| TRINITY_DN137233_c0_g1 | DVH24_029871; DVH24_026477 | 1951.7742022 | -15.7730043 | 0.1410120079 | -14.032601 | hypothetical protein DVH24_029871 | membrane | GO:0016020 | NA | NA | Rola-high | 41.19543 |
| TRINITY_DN51_c | TRINITY_DN51_c0_g1 | 618.7868604 | -12.1878825 | 0.1452505861 | -14.3458 [illegible]2 | uncharacterized protein | NA | NA | NA | NA | Rola-high | 41.19543 |

| gene_id | gene_name | baseMean | log2FoldChange | lfcSE | stat | Description | Ontology_term | Ontology_id | Enzyme_code | Enzyme_name | direction | neg_log10_padj |
|---|---|---|---|---|---|---|---|---|---|---|---|---|
| 0_g1 | | | | | | | | | | | | |
| TRINITY_DN4249_c0_g2 | TRINITY_DN4249_c0_g2 | 151.47730 | -25.876213 | 2.0351337 | -12.71475 | protein GAST1 isoform X2 | NA | NA | NA | NA | Rola-high | 31.85919 |
| TRINITY_DN | DVH24_011177;DVH24_011179 | 622.44398 | 3.954495 | 0.3152162 | 12.54534 | berberine bridge enzyme-like 8 | oxidoreductase activity,FAD binding | GO:0016491,GO:0071949 | EC:1 | Oxidoreductases | Rin-high | 31.04916 |

| gene_id | gene_name | baseMean | log2FoldChange | lfcSE | stat | Description | Ontology_term | Ontology_id | Enzyme_code | Enzyme_name | direction | neg_log10_padj |
|---|---|---|---|---|---|---|---|---|---|---|---|---|
| 18936_c0_g1 | | | | | | | | | | | | |
| TRINITY_DN87441_c0_g1 | DVH24_010981; DVH24_020628; DVH24_026348; DVH24_012538 | 1184.77078 | -1.618917 | 0.1318463 | -12.27882 | LOB domain-containing protein 37-like | cytoplasm,membrane,glycosyltransferase activity,regulation of gene expression | GO:0005737,GO:0016020,GO:0016757,GO:0010468 | EC:2.4 | Glycosyltransferases | Rola-high | 29.700019 |
| TRI | DVH24_003302; DVH24_0 | 140 | -2. | 0.12 | -12 | GDSL esterase/lipa | hydrolase activity, acting on ester bonds | GO:0016788 | EC: | Acting on | Rol | 28. |

| gene_id | gene_name | baseMean | log2FoldChange | lfcSE | stat | Description | Ontology_term | Ontology_id | Enzyme_code | Enzyme_name | direction | neg_log10_padj |
|---|---|---|---|---|---|---|---|---|---|---|---|---|
| NITY_DN31025_c0_g1 | 32401; DVH24_032402; DVH24_032651 | .87969 | 478467 | 062848 | .01478 | se At2g42990 | | | 3.1 | ester bonds | a-high | 377116 |
| TRINITY_DN1466_c1 | DVH24_034924; DVH24_042310; DVH24_021501; DVH24_042312; MdSOT3 | 1999.459549 | 3.839685 | 0.3199470 | 12.00100 | putative polyol transporter 1 | membrane,carbohydrate:proton symporter activity,carbohydrate transmembrane transport,proton transmembrane transport | GO:0016020,GO:0005351, GO:0034219,GO:1902600 | NA | NA | Rin-high | 28.37176 |

| gene_id | gene_name | baseMean | log2FoldChange | lfcSE | stat | Description | Ontology_term | Ontology_id | Enzyme_code | Enzyme_name | direction | neg_log10_padj |
|---|---|---|---|---|---|---|---|---|---|---|---|---|
| _g1 | | | | | | | | | | | | |
| TRINITY_DN6594_c1_g1 | DVH24_002005; DVH24_018718; DVH24_010436; DVH24_028654 | 290.43930 | -6.481784 | 0.5477278 | -11.83395 | probable pectate lyase 8 | pectate lyase activity,metal ion binding,pectin catabolic process | GO:0030570,GO:0046872,GO:0045490 | EC:4.2.2.2 | pectate lyase | Rola-high | 27.55914 |
| TRINITY_DN5 | DVH24_033743; DVH24_033745; DVH24_040960 | 1023.006555 | -1.050814 | 0.1089486 | -11.74269 | very-long-chain 3-oxoacyl-CoA reductase 1-like | endoplasmic reticulum,membrane,ketoreductase activity | GO:0005783,GO:0016020,GO:0045703 | EC:1.1.1 | Acting on the CH-OH group of donors | Rola-high | 27.13974 |

| gene_id | gene_name | baseMean | log2FoldChange | lfcSE | stat | Description | Ontology_term | Ontology_id | Enzyme_code | Enzyme_name | direction | neg_log10_padj |
|---|---|---|---|---|---|---|---|---|---|---|---|---|
| 386_c0_g1 | | | | | | | | | | | | |
| TRINITY_DN36061_c1_g1 | TRINITY_DN36061_c1_g1 | 96.95591 | -25.590027 | 2.1818836 | -11.72841 | protein PELP K2-like | NA | NA | NA | NA | Rola-high | 27.11221 |

#### Table preview 19

**Candidate Rin-high Trinity genes using padj < 0.05, |LFC| >= 1, baseMean >= 50.**

| gene_id | gene_name | log2FoldChange | Description | Ontology_term | Ontology_id | Enzyme_code | Enzyme_name |
|---|---|---|---|---|---|---|---|
| TRINITY_DN18936_c0_g1 | DVH24_011177;DVH24_011179 | 3.95444995 | berberine bridge enzyme-like 8 | oxidoreductase activity,FAD binding | GO:0016491,GO:0071949 | EC:1 | Oxidoreductases |
| TRINITY_DN | DVH24_034924;DVH24_042310;DVH24_021501;DVH24 | 3.8396685 | putative polyol transporter 1 | membrane,carbohydrate:proton symporter activity,carbohydrate transmembrane transport,proton transmembrane transport | GO:0016020,GO:0005351,GO:0034219,GO:1902 | NA | NA |

| gene_id | gene_name | log2FoldChange | Description | Ontology_term | Ontology_id | Enzyme_code | Enzyme_name |
|---|---|---|---|---|---|---|---|
| 1466_c1_g1 | _042312; MdSOT3 | | | | 600 | | |
| TRINITY_DN76668_c2_g1 | DVH24_022240 | 3.83698989 | RAF-like serine/threonine-protein kinase PRAF | NA | NA | NA | NA |
| T | DVH24 | 4 | RAF-like | negative regulation | GO:0 | N | NA |

| gene_id | gene_name | log2FoldChange | Description | Ontology_term | Ontology_id | Enzyme_code | Enzyme_name |
|---|---|---|---|---|---|---|---|
| RINITY_DN14750_c0_g1 | _022240; DVH24_025811; DVH24_001430 | .4028 74 | serine/threonine-protein kinase PRAF | of DNA-templated transcription,transcription corepressor activity | 045892,GO:0003714 | A | |
| TRINITY_DN2 | DVH24_029261; DVH24_017909 | 2.968774 | probable chlorophyll(ide) b reductase NYC1, chloroplastic | membrane,chlorophyll(ide) b reductase activity,PSII associated light-harvesting complex II catabolic process,chlorophyll catabolic process | GO:0016020,GO:0034256,GO:0010304,GO:0015996 | EC:1.1.1.29 | chlorophyll(ide) b reductase |

| gene_id | gene_name | log2FoldChange | Description | Ontology_term | Ontology_id | Enzyme_code | Enzyme_name |
|---|---|---|---|---|---|---|---|
| 33357_c0_g1 | | | | | | 4 | |
| TRINITY_DN499_c0_g1 | DVH24_007191; DVH24_000721; MYBR7 | 1.401382 | myb family transcription factor PHL8-like | oxidoreductase activity, acting on CH-OH group of donors,mandelonitrile lyase activity,flavin adenine dinucleotide binding | GO:0016614,GO:0046593,GO:0050660 | EC:1.1,EC:4.1.2.10 | Acting on the CH-OH group of donors, (R)-mandelonitrile lyase |
| TR | TRINITY_DN24 | 6. | probable LRR | NA | NA | NA | NA |

| gene_id | gene_name | log2FoldChange | Description | Ontology_term | Ontology_id | Enzyme_code | Enzyme_name |
|---|---|---|---|---|---|---|---|
| INITY_DN24806_c0_g1 | 806_c0_g1 | 325222 | receptor-like serine/threonine-protein kinase At3g47570 isoform X2 | | | | |
| TRINITY_DN43 | TRINITY_DN4310_c0_g1 | 3.562830 | uncharacterized protein | NA | NA | NA | NA |

| gene_id | gene_name | log2FoldChange | Description | Ontology_term | Ontology_id | Enzyme_code | Enzyme_name |
|---|---|---|---|---|---|---|---|
| 10_c0_g1 | | | | | | | |
| TRINITY_DN22261_c0_g2 | DVH24_006306; DVH24_029261; DVH24_017909 | 2.664371 | probable chlorophyll(ide) b reductase NYC1, chloroplastic | membrane,chlorophyll(ide) b reductase activity,PSII associated light-harvesting complex II catabolic process,chlorophyll catabolic process | GO:0016020,GO:0034256,GO:0010304,GO:0015996 | EC:1.1.1.294 | chlorophyll(ide) b reductase |
| TRI | DVH24_024453 | 6.0 | probable LRR receptor- | membrane | GO:0016020 | NA | NA |

| gene_id | gene_name | log2FoldChange | Description | Ontology_term | Ontology_id | Enzyme_code | Enzyme_name |
|---|---|---|---|---|---|---|---|
| NITY_DN4777_c8_g3 | | 25652 | like serine/threonine-protein kinase At3g47570 isoform X1 | | | | |

#### Table preview 20

**Candidate Rola-high Trinity genes using padj < 0.05, |LFC| >= 1, baseMean >= 50.**

| gene_id | gene_name | Description | Ontology_term | Ontology_id | Enzyme_code | Enzyme_name |
|---|---|---|---|---|---|---|
| TRINITY_DN13723_c0_g1 | DVH24_029871; DVH24_026477 | hypothetical protein DVH24_029871 | membrane | GO:0016020 | NA | NA |
| TRINITY_DN51_ | TRINITY_DN51_c0_g1 | uncharacterized protein | NA | NA | NA | NA |

| gene_id | gene_name | Description | Ontology_term | Ontology_id | Enzyme_code | Enzyme_name |
|---|---|---|---|---|---|---|
| c0_g1 | | | | | | |
| TRINITY_DN4249_c0_g2 | TRINITY_DN4249_c0_g2 | protein GAST1 isoform X2 | NA | NA | NA | NA |
| TRINITY_D | DVH24_010981;DVH24_0 | LOB domain-containing protein | cytoplasm,membrane,glycosyltransferase activity,regulation of gene expression | GO:0005737,GO:0016020,GO:0016757,GO:0010468 | EC:2.4 | Glycosyltransferases |

| gene_id | gene_name | Description | Ontology_term | Ontology_id | Enzyme_code | Enzyme_name |
|---|---|---|---|---|---|---|
| N8741_c0_g1 | 20628;DVH24_026348;DVH24_012538 | 37-like | | | | |
| TRINITY_DN31025_c0_ | DVH24_003302;DVH24_032401;DVH24_032402;DV | GDSL esterase/lipase At2g42990 | hydrolase activity, acting on ester bonds | GO:0016788 | EC:3.1 | Acting on ester bonds |

| gene_id | gene_name | Description | Ontology_term | Ontology_id | Enzyme_code | Enzyme_name |
|---|---|---|---|---|---|---|
| g1 | H24_0326 51 | | | | | |
| TRINITY_DN6594_c1_g1 | DVH24_0020 05;DVH24_0187 18;DVH24_0104 36;DVH24_0286 54 | probable pectate lyase 8 | pectate lyase activity,metal ion binding,pectin catabolic process | GO:0030570,GO:0046872,GO:0045490 | EC:4.2.2.2 | pectate lyase |
| TRINI | DVH24_0333 74 | very-long-chai | endoplasmic reticulum,membrane,ketoreductase activity | GO:0005783,GO:0016020,GO:0045703 | EC:1. | Acting on the CH-OH group |

| gene_id | gene_name | Description | Ontology_term | Ontology_id | Enzyme_code | Enzyme_name |
|---|---|---|---|---|---|---|
| TY_DN5386_c0_g1 | 3;DVH24_033745;DVH24_040960 | n 3-oxoacyl-CoA reductase 1-like | | | 1 | of donors |
| TRINITY_DN36061_c1_ | TRINITY_DN36061_c1_g1 | protein PELPK2-like | NA | NA | NA | NA |

| gene_id | gene_name | Description | Ontology_term | Ontology_id | Enzyme_code | Enzyme_name |
|---|---|---|---|---|---|---|
| g1 | | | | | | |
| TRINITY_DN22704_c0_g1 | TRINITY_DN22704_c0_g1 | aldehyde oxidase GLOX1-like | NA | NA | NA | NA |
| TRINITY_DN1 | DVH24_00495 5; DVH24_00657 | glycerol-3-phosphate dehydrogenase | nucleus,cytosol,DNA-binding transcription factor activity, RNA polymerase II-specific,cis-regulatory region sequence-specific DNA binding,protein dimerization activity,NAD binding,glycerol-3-phosphate dehydrogenase (NAD+) activity,carbohydrate metabolic process,positive regulation of transcription by RNA polymerase | GO:0005634,GO:0005829,GO:0000981,GO:0000987,GO:0046983,GO:0051287,GO:0141152, | EC:1.1.1.8, | glycerol-3-phosphate dehydrogenase (NAD(+)),glycerol-3- |

| gene_id | gene_name | Description | Ontology_term | Ontology_id | Enzyme_code | Enzyme_name |
|---|---|---|---|---|---|---|
| 0440_c0_g1 | 5;DVH24_016043;DVH24_019631 | [NAD(+)] GPDHC1, cytosolic | II,glycerol-3-phosphate catabolic process | GO:0005975,GO:0045944,GO:0046168 | EC:1.1.1.94 | phosphate dehydrogenase [NAD(P)(+)] |

#### Table preview 21

**Rin vs Rola DESeq2 results using lfcThreshold = 1.**

| gene_id | gene_name | Description | Ontology_term | Ontology_id | Enzyme_code | Enzyme_name |
|---|---|---|---|---|---|---|
| TRINITY_DN | TRINITY_DN4249_c0_g2 | protein GAST1 isoform X2 | NA | NA | NA | NA |

| gene_id | gene_name | Description | Ontology_term | Ontology_id | Enzyme_code | Enzyme_name |
|---|---|---|---|---|---|---|
| 4249_c0_g2 | | | | | | |
| TRINITY_DN36061_c1_g1 | TRINITY_DN36061_c1_g1 | protein PELPK2-like | NA | NA | NA | NA |
| TRINITY | DVH24_02005; DVH24_01871 | probable pectate lya | pectate lyase activity,metal ion binding,pectin catabolic process | GO:0030570,GO:0046872,GO:0045490 | EC:4.2.2.2 | pectate lyase |

| gene_id | gene_name | Description | Ontology_term | Ontology_id | Enzyme_code | Enzyme_name |
|---|---|---|---|---|---|---|
| _DN65994_c1_g1 | 8;DVH24_010436;DVH24_028654 | se8 | | | | |
| TRINITY_DN18936_c0_g1 | DVH24_011177;DVH24_011179 | berberine bridge enzyme-like 8 | oxidoreductase activity,FAD binding | GO:0016491,GO:0071949 | EC:1 | Oxidoreductases |
| TRIN | DVH24_034924; | putative pol | membrane,carbohydrate:proton symporter activity,carbohydr | GO:0016020,GO:000535 | NA | NA |

| gene_id | gene_name | Description | Ontology_term | Ontology_id | Enzyme_code | Enzyme_name |
|---|---|---|---|---|---|---|
| ITY_DN14666_c1_g1 | DVH24_04231 0;DVH24_02150 1;DVH24_04231 2;MdSOT3 | yoltransporter1 | ate transmembrane transport,proton transmembrane transport | 1,GO:0034219,GO:1902600 | | |
| TRINITY_DN48040_c0_g1 | DVH24_00798;DVH24_03418 0 | adenosylhomocysteinase | NA | NA | NA | NA |
| T | DVH | en | extracellular | GO:00 | EC:3.2. | cellulase |

| gene_id | gene_name | Description | Ontology_term | Ontology_id | Enzyme_code | Enzyme_name |
|---|---|---|---|---|---|---|
| RINITY_DN1459_c1_g1 | 24_04092;DVH24_32378;DVH24_40086;DVH24_28125 | doglucanase 11-like | region,cellulase activity,carbohydrate binding,cellulose catabolic process | 05576,GO:0008810,GO:0030246,GO:0030245 | 1.4 | |
| TRINITY_DN25548_c0_ | DVH24_01106 7;DVH24_11013;DVH24_40555;DVH24_12188 | fe(2+) transport protein 1-like | NA | NA | NA | NA |

| gene_id | gene_name | Description | Ontology_term | Ontology_id | Enzyme_code | Enzyme_name |
|---|---|---|---|---|---|---|
| g1 | | | | | | |
| TRINITY_DN10437_c0_g1 | DVH24_03213 9;DVH24_04048 7;DVH24_07375 | allene oxide cyclase, chloroplastic-like | chloroplast,initiator methionyl aminopeptidase activity,allene-oxide cyclase activity,metal ion binding,metalloaminopeptidase activity,proteolysis,jasmonic acid biosynthetic process | GO:0009507,GO:0004239,GO:0046423,GO:0046872,GO:0070006,GO:0006508,GO:0009695 | EC:3.4.11.18,EC:3.4.11.21,EC:3.4.11.22,EC:3.4.11.6,EC:3.4.11.5,EC:5.3.99.6,EC:3.4.11.9 | methionyl aminopeptidase,aspartyl aminopeptidase,aminopeptidase I,aminopeptidase B,prolyl aminopeptidase,allene-oxide cyclase,Xaa-Pro aminopeptidase |
| TRINITY_DN21977 | DVH24_02658 9;DVH24_03609 7;DVH24_03716 2;DVH24_00710 | glycerol-3-phosphate acyltransferase RA | membrane,phosphatase activity,glycerol-3-phosphate 2-O-acyltransferase activity,cutin biosynthetic process,protein homooligomerization | GO:0016020,GO:0016791,GO:0090447,GO:0010143,GO:0051260 | EC:2.3.1.198,EC:3.1.3 | glycerol-3-phosphate 2-O-acyltransferase,Acting on ester bonds |

| gene_id | gene_name | Description | Ontology_term | Ontology_id | Enzyme_code | Enzyme_name |
|---|---|---|---|---|---|---|
| _c0_g1 | 6;DVH24_014245 | M2-like | | | | |

#### Table preview 22

**Rin vs Rola lfcThreshold = 1 results sorted by log2FC.**

| gene_name | Description | Ontology_term | Ontology_id | Enzyme_code | Enzyme_name |
|---|---|---|---|---|---|
| DVH24_024718;DVH2 | uncharacterizedpro | response to cold,regulation of circadian rhythm | GO:0009409,GO:0042752 | NA | NA |

| gene_name | Description | Ontology_term | Ontology_id | Enzyme_code | Enzyme_name |
|---|---|---|---|---|---|
| 4_022573 | tein | | | | |
| DVH24_042242 | hypothetical protein DVH24_04 | nucleus,3-phosphoinositide-dependent protein kinase activity,DNA-dependent protein kinase activity,AMP-activated protein kinase activity,eukaryotic translation initiation factor 2alpha kinase activity,ribosomal protein S6 kinase activity,ATP binding,histone H3S10 kinase activity,histone H3T11 kinase activity,histone H3T6 kinase activity,histone H2AXS139 kinase activity,histone H3S28 kinase activity,histone H4S1 kinase activity,histone H2AS1 kinase activity,histone H2BS14 kinase activity,histone H3T3 kinase activity,histone H2AS121 kinase activity,Rho-dependent protein serine/threonine kinase activity,histone H2BS36 kinase activity,histone H3S57 kinase activity,histone H3T45 kinase activity,histone H2AT120 kinase activity,nuclear-transcribed mRNA catabolic process, nonsense-mediated decay,chromatin remodeling | GO:0005634,GO:0004676,GO:0004677,GO:0004679,GO:0004694,GO:0004711,GO:0005524,GO:0035175,GO:0035402,GO:0035403,GO:0035979,GO:0044022,GO:0044023,GO:0044024,GO:0044025,GO:0072354,GO:0072371,GO:0072518,GO:0140823,GO:0140855,GO:01408 | EC:2.7.11.1,EC:2.7.11.29,E | non-specific serine/threonine protein kinase, [low-density-lipoprotein receptor] kinase, dephospho-[reductase kinase] kinase, tropomyosin kinase, [Goodp |

| gene_name | Description | Ontology_term | Ontology_id | Enzyme_code | Enzyme_name |
|---|---|---|---|---|---|
| | 2242 | | 57,GO:1990244,GO:0000184,GO:0006338 | C:2.7.11.3,EC:2.7.11.28,EC:2.7.11. | asture-antigen-binding protein] kinase |

| gene_name | Description | Ontology_term | Ontology_id | Enzyme_code | Enzyme_name |
| --- | --- | --- | --- | --- | --- |
| DVH24_02235 0;DVH24_024986 | two-component response regulator-like | nucleus,phosphorelay signal transduction system,cytokinin-activated signaling pathway,rhythmic process | GO:0005634,GO:0000160,GO:0009736,GO:0048511 | NA | NA |

| gene_name | Description | Ontology_term | Ontology_id | Enzyme_code | Enzyme_name |
|---|---|---|---|---|---|
| DVH24_03022 2;DVH24_02 6133;C0I | APRR5 probable xyloglucan endotransgluc | apoplast,hydrolase activity, hydrolyzing O-glycosyl compounds,xyloglucan:xyloglucosyl transferase activity,xyloglucan metabolic process,cell wall biogenesis,cell wall organization | GO:0048046,GO:0004553,GO:0016762,GO:0010411,GO:0042546,GO:0071555 | EC:2.4.1.207,EC:3.2.1 | xyloglucan:xyloglucosyl transferase,Glycosylases |

| gene_name | Description | Ontology_term | Ontology_id | Enzyme_code | Enzyme_name |
|---|---|---|---|---|---|
| RH7 | osylase/hydrolaseprotein6precursor | | | | |
| D | m | nucleus | GO:00056 | N | NA |

| gene_name | Description | Ontology_term | Ontology_id | Enzyme_code | Enzyme_name |
|---|---|---|---|---|---|
| VH24_0199555;DVH24_0199551;DVH24_008 | yb-relatedprotein308-like | | 34 | A | |

| gene_name | Description | Ontology_term | Ontology_id | Enzyme_code | Enzyme_name |
|---|---|---|---|---|---|
| 114;DVH24_00811 3 | | | | | |
| DVH24_026532 | peroxidase72 | extracellular region,heme binding,metal ion binding,lactoperoxidase activity,response to oxidative stress,hydrogen peroxide catabolic process,cellular oxidant detoxification | GO:0005576,GO:0020037,GO:0046872,GO:0140825,GO:0006979,GO:0042744,GO:0098869 | EC:1.11.1.7 | peroxidase |
| DVH24_ | chaper | membrane | GO:0016020 | NA | NA |

| gene_name | Description | Ontology_term | Ontology_id | Enzyme_code | Enzyme_name |
|---|---|---|---|---|---|
| 006097;DVH24_030860;DVH24_040197 | one protein dnaJ C76, chloroplastic | | | | |
| TR | TR | NA | NA | NA | NA |

| gene_name | Description | Ontology_term | Ontology_id | Enzyme_code | Enzyme_name |
|---|---|---|---|---|---|
| INITY_DN21285_c1_g1 | INITY_DN21285_c1_g1 | | | | |
| TRINITY_DN57958 | senescence-speci | NA | NA | NA | NA |

| gene_name | Description | Ontology_term | Ontology_id | Enzyme_code | Enzyme_name |
|---|---|---|---|---|---|
| _c1_g1 | fic cysteine protease SAG12-like | | | | |
| DVH24_0 | cold-re | Golgi cisterna membrane,fucosyltransferase activity | GO:0032580,GO:0008417 | EC:2.4. | Glycosyltransferases |

| gene_name | Description | Ontology_term | Ontology_id | Enzyme_code | Enzyme_name |
|---|---|---|---|---|---|
| 124556;DVH24_020724;DVH24_021640 | gulated protein 27 | | | 1 | |

#### Table preview 23

**Rin-high candidates in Rin vs Rola using lfcThreshold = 1 and padj < 0.01.**

| gene_name | Description | Ontology_term | Ontology_id | Enzyme_code | Enzyme_name |
|---|---|---|---|---|---|
| DVH24_02471 8;DVH24_02257 3 | uncharacterized protein | response to cold,regulation of circadian rhythm | GO:0009409,GO:0042752 | NA | NA |
| DVH24 | hypot | nucleus,3-phosphoinositide-dependent protein kinase activity,DNA-dependent protein kinase activity,AMP-activated protein kinase activity,eukaryotic translation initiation factor 2alpha | GO:0005634,GO:0004676,GO:0004677,GO:0004679, | EC:2. | non-specific serine/threoni |

| gene_name | Description | Ontology_term | Ontology_id | Enzyme_code | Enzyme_name |
|---|---|---|---|---|---|
| _042242 | hetical protein DVH24_042242 | kinase activity,ribosomal protein S6 kinase activity,ATP binding,histone H3S10 kinase activity,histone H3T11 kinase activity,histone H3T6 kinase activity,histone H2AXS139 kinase activity,histone H3S28 kinase activity,histone H4S1 kinase activity,histone H2AS1 kinase activity,histone H2BS14 kinase activity,histone H3T3 kinase activity,histone H2AS121 kinase activity,Rho-dependent protein serine/threonine kinase activity,histone H2BS36 kinase activity,histone H3S57 kinase activity,histone H3T45 kinase activity,histone H2AT120 kinase activity,nuclear-transcribed mRNA catabolic process, nonsense-mediated decay,chromatin remodeling | GO:0004694,GO:0004711,GO:0005524,GO:0035175,GO:0035402,GO:0035403,GO:0035979,GO:0044022,GO:0044023,GO:0044024,GO:0044025,GO:0072354,GO:0072371,GO:0072518,GO:0140823,GO:0140855,GO:0140857,GO:1990244,GO:0000184,GO:0006338 | 7.11.1,EC:2.7.11.29,EC:2.7.11.3,EC | ne protein kinase,[low-density-lipoprotein receptor] kinase,dephospho-[reductase kinase] kinase,tropomyosin kinase,[Goodpasture-antigen-binding protein] kinase |

| gene_name | Description | Ontology_term | Ontology_id | Enzyme_code | Enzyme_name |
|---|---|---|---|---|---|
| | | | | :2.7.11.28,EC:2.7.11.9 | |
| DVH24_0223500 | two-componen | nucleus,phosphorelay signal transduction system,cytokinin-activated signaling pathway,rhythmic process | GO:0005634,GO:0000160,GO:0009736,GO:0048511 | NA | NA |

| gene_name | Description | Ontology_term | Ontology_id | Enzyme_code | Enzyme_name |
|---|---|---|---|---|---|
| ;DVH24_02498 6 | tresponseregulator-likeAPRR5 | | | | |
| DVH24_03 | probable | apoplast,hydrolase activity, hydrolyzing O-glycosyl compounds,xyloglucan:xyloglucosyl transferase activity,xyloglucan metabolic process,cell wall biogenesis,cell wall organization | GO:0048046,GO:0004553,GO:0016762,GO:0010411,GO:0042546,GO:0071555 | EC:2.4.1 | xyloglucan:xyloglucosyl transferase,Glycosylases |

| gene_name | Description | Ontology_term | Ontology_id | Enzyme_code | Enzyme_name |
|---|---|---|---|---|---|
| 0222;DVH24_026133;C0IRH7 | xyloglucanendotransglucosylase/hydrol | | | .207,EC:3.2.1 | |

| gene_name | Description | Ontology_term | Ontology_id | Enzyme_code | Enzyme_name |
|---|---|---|---|---|---|
| | aseprotein6precursor | | | | |
| DVH24_0199555;D | myb-relatedpro | nucleus | GO:0005634 | NA | NA |

| gene_name | Description | Ontology_term | Ontology_id | Enzyme_code | Enzyme_name |
| --- | --- | --- | --- | --- | --- |
| VH24_019951;DVH24_008114;DVH24_008 | tein308-like | | | | |

| gene_name | Description | Ontology_term | Ontology_id | Enzyme_code | Enzyme_name |
|---|---|---|---|---|---|
| 113 | | | | | |
| DVH24_00060097;DVH24_03086 0;DVH24 | chaperone protein dnaJC76, chlorop | membrane | GO:0016020 | NA | NA |

| gene_name | Description | Ontology_term | Ontology_id | Enzyme_code | Enzyme_name |
|---|---|---|---|---|---|
| _040197 | lastic | | | | |
| TRINITY_DN21285_c1_g1 | TRINITY_DN21285_c1_g1 | NA | NA | NA | NA |
| TRINIT | senesc | NA | NA | NA | NA |

| gene_name | Description | Ontology_term | Ontology_id | Enzyme_code | Enzyme_name |
| --- | --- | --- | --- | --- | --- |
| Y_DN57958_c1_g1 | ence-specific cysteine protease SAG12-li | | | | |

| gene_name | Description | Ontology_term | Ontology_id | Enzyme_code | Enzyme_name |
|---|---|---|---|---|---|
| DVH24_012456;DVH24_020724;DVH24_ | kecold-regulatedprotein27 | Golgi cisterna membrane,fucosyltransferase activity | GO:0032580,GO:0008417 | EC:2.4.1 | Glycosyltransferases |

| gene_name | Description | Ontology_term | Ontology_id | Enzyme_code | Enzyme_name |
|---|---|---|---|---|---|
| 02164 0 | | | | | |
| DVH24_00539 1;DVH24_02235 0;DV | two-component response regulat | nucleus,phosphorelay signal transduction system,cytokinin-activated signaling pathway,rhythmic process | GO:0005634,GO:0000160,GO:0009736,GO:0048511 | NA | NA |

| gene_name | Description | Ontology_term | Ontology_id | Enzyme_code | Enzyme_name |
|---|---|---|---|---|---|
| H24_024986 | or-like APRR5 | | | | |

#### Table preview 24

**Rola-high candidates in Rin vs Rola using lfcThreshold = 1 and padj < 0.01.**

| gene_id | gene_name | baseMean | log2FoldChange | lfcSE | stat | pvalue | padj | Description | Ontology_term | Ontology_id | Enzyme_code | Enzyme_name |
|---|---|---|---|---|---|---|---|---|---|---|---|---|
| TRINITY_DN4249_c0_g2 | TRINITY_DN4249_c0_g2 | 151.4773 | -25.87621 | 2.0351334 | -12.7147 | 0 | 0.0e+00 | protein GAST1 isoform X2 | NA | NA | NA | NA |

| gene_id | gene_name | baseMean | log2FoldChange | lfcSE | stat | pvalue | padj | Description | Ontology_term | Ontology_id | Enzyme_code | Enzyme_name |
|---|---|---|---|---|---|---|---|---|---|---|---|---|
| | | 0 | | | 48 | | | | | | | |
| TRINITY_DN36061_c1_g1 | TRINITY_DN36061_c1_g1 | 96.9559591 | -25.590003 | 2.1818884 | -11.728411 | 0 | 0.0e+00 | protein PELPK2-like | NA | NA | NA | NA |
| TRINITY_DN11246_c0_g1 | TRINITY_DN11246_c0_g1 | 20.8436 8 | -23.38642 | 3.7954223 | -6.1617743 | 0 | 8.0e-07 | TRINITY_DN11246_c0_g1 | NA | NA | NA | NA |
| TRINITY_DN139024_c0_g1 | TRINITY_DN139024_c0_g1 | 24.944417 | -23.053375 | 3.8389884 | -6.0051771 | 0 | 1.6e-06 | TRINITY_DN139024_c0_g1 | NA | NA | NA | NA |
| TRINITY_D | TRINITY_DN6336_c0_g1 | 19. | -22.92 | 3.7 | -6. | 0 | 1.3 | TRINITY_DN6336_c0_g1 | NA | NA | NA | NA |

| gene_id | gene_name | baseMean | log2FoldChange | lfcSE | stat | pvalue | padj | Description | Ontology_term | Ontology_id | Enzyme_code | Enzyme_name |
|---|---|---|---|---|---|---|---|---|---|---|---|---|
| N6336_c0_g1 | | 07138 | 457 | 87636 | 052473 | | e-06 | | | | | |
| TRINITY_DN229006_c0_g1 | TRINITY_DN229006_c0_g1 | 35.817112 | -22.733337 | 3.8393385 | -5.921096 | 0 | 2.4e-06 | TRINITY_DN229006_c0_g1 | NA | NA | NA | NA |
| TRINITY_DN18390_c0_g1 | DVH24_033020; DVH24_038538; DVH24_017268 | 14.378668 | -22.724416 | 3.0625554 | -7.420001 | 0 | 0.0e+000 | alpha-glucan phosphorylase, H isozyme-like | 1,4-alpha-oligoglucan phosphorylase activity | GO:0004645 | EC:2.4.1.1 | glycogen phosphorylase |
| TRINITY_DN21772_c0_g1 | TRINITY_DN21772_c0_g1 | 15.981171 | -22.342267 | 3.7378444 | -5.977422 | 0 | 1.9e-06 | TRINITY_DN21772_c0_g1 | NA | NA | NA | NA |

| gene_id | gene_name | baseMean | log2FoldChange | lfcSE | stat | pvalue | padj | Description | Ontology_term | Ontology_id | Enzyme_code | Enzyme_name |
|---|---|---|---|---|---|---|---|---|---|---|---|---|
| TRINITY_DN81325_c0_g2 | TRINITY_DN81325_c0_g2 | 11.663340 | -21.34459 | 3.672315 | -5.8122297 | 0 | 3.9e-06 | TRINITY_DN81325_c0_g2 | NA | NA | NA | NA |
| TRINITY_DN138657_c0_g1 | TRINITY_DN138657_c0_g1 | 14.178807 | -21.10062 | 3.723414 | -5.6670070 | 0 | 7.8e-06 | TRINITY_DN138657_c0_g1 | NA | NA | NA | NA |

#### Table preview 25

**Rin vs Rola candidates sorted by absolute log2FC.**

| gene_id | gene_name | baseMean | log2FoldChange | lfcSE | stat | pvalue | padj | direction | Description | Ontology_term | Ontology_id | Enzyme_code | Enzyme_name |
|---|---|---|---|---|---|---|---|---|---|---|---|---|---|
| TRI | TRINITY_ | 1 | -2 | 2 | - | 0 | 0 | R | protein | NA | N | N | NA |

| gene_id | gene_name | baseMean | log2FoldChange | lfcSE | stat | pvalue | padj | direction | Description | Ontology_term | Ontology_id | Enzyme_code | Enzyme_name |
|---|---|---|---|---|---|---|---|---|---|---|---|---|---|
| NITY_DN4249_c0_g2 | DN4249_c0_g2 | 51.47730 | 5.87621 | .035134 | 12.714748 | | .0e+00 | ola-high | GAST1 isoform X2 | | A | A | |
| TRINITY_DN36061_c1_g1 | TRINITY_DN36061_c1_g1 | 96.955591 | -25.59003 | 2.181884 | -11.728411 | 0 | 0.0e+00 | Rola-high | protein PELPK2-like | NA | NA | NA | NA |
| TRINITY_DN11246_c0_g1 | TRINITY_DN11246_c0_g1 | 20.84368 | -23.38642 | 3.795423 | -6.161743 | 0 | 8.0e-07 | Rola-high | TRINITY_DN11246_c0_g1 | NA | NA | NA | NA |
| TRINITY_DN139024_c0 | TRINITY_DN139024_c0_g1 | 24.944 | -23.05375 | 3.8389 | -6.005 | 0 | 1.6e-0 | Rola-h | TRINITY_DN139024_c0_g1 | NA | NA | NA | NA |

| gene_id | gene_name | baseMean | log2FoldChange | lfcSE | stat | pvalue | padj | direction | Description | Ontology_term | Ontology_id | Enzyme_code | Enzyme_name |
|---|---|---|---|---|---|---|---|---|---|---|---|---|---|
| _g1 | | 17 | | 84 | 171 | | 6 | igh | | | | | |
| TRINITY_DN6336_c0_g1 | TRINITY_DN6336_c0_g1 | 19.0713 8 | -22.92457 | 3.787636 | -6.052473 | 0 | 1.3e-06 | Rola-high | TRINITY_DN6336_c0_g1 | NA | NA | NA | NA |
| TRINITY_DN229006_c0_g1 | TRINITY_DN229006_c0_g1 | 35.81712 | -22.73337 | 3.839385 | -5.921096 | 0 | 2.4e-06 | Rola-high | TRINITY_DN229006_c0_g1 | NA | NA | NA | NA |
| TRINITY_DN18390_c0_g1 | DVH24_033020; DVH24_038538; DVH24_017268 | 14.37868 | -22.72416 | 3.062554 | -7.420001 | 0 | 0.00e+00 | Rola-high | alpha-glucan phosphorylase, H isozyme-like | 1,4-alpha-oligoglucan phosphorylase activity | GO:0004645 | EC:2.4.1.1 | glycogen phosphorylase |
| TRINITY_DN21772 | TRINITY_DN21772 | 15 | -22. | 3. | -5 | 0 | 1. | Ro | TRINITY_DN21772 | NA | NA | NA | NA |

| gene_id | gene_name | baseMean | log2FoldChange | lfcSE | stat | pvalue | padj | direction | Description | Ontology_term | Ontology_id | Enzyme_code | Enzyme_name |
|---|---|---|---|---|---|---|---|---|---|---|---|---|---|
| Y_DN21772_c0_g1 | _c0_g1 | .98171 | 34267 | 737844 | .977422 | | 9e-06 | la-high | _c0_g1 | | | | |
| TRINITY_DN81325_c0_g2 | TRINITY_DN81325_c0_g2 | 11.66340 | -21.34459 | 3.672315 | -5.812297 | 0 | 3.9e-06 | Rola-high | TRINITY_DN81325_c0_g2 | NA | NA | NA | NA |
| TRINITY_DN138657_c0_g1 | TRINITY_DN138657_c0_g1 | 14.17807 | -21.10062 | 3.723414 | -5.667007 | 0 | 7.8e-06 | Rola-high | TRINITY_DN138657_c0_g1 | NA | NA | NA | NA |

#### Table preview 26

**Rin vs control DESeq2 results using lfcThreshold = 1.**

| gene_id | gene_name | baseMean | log2FoldChange | lfcSE | stat | Description | Ontology_term | Ontology_id | Enzyme_code | Enzyme_name |
|---|---|---|---|---|---|---|---|---|---|---|
| TRINITY_DN18768_c0_g1 | TRINITY_DN18768_c0_g1 | 4375.061331 | 12.555326 | 0.6590477 | 19.05071 | TRINITY_DN18768_c0_g1 | NA | NA | NA | NA |
| TRINITY_DN16515_c0_g1 | TRINITY_DN16515_c0_g1 | 2205.32940 | 13.340787 | 0.8122674 | 16.42413 | TRINITY_DN16515_c0_g1 | NA | NA | NA | NA |
| TRINITY_DN23 | TRINITY_DN23990_c0_g1 | 2005.13 | 13.0169 | 0.79128 | 16.4505 | TRINITY_DN23990_c0_g1 | NA | NA | NA | NA |

| gene_id | gene_name | baseMean | log2FoldChange | lfcSE | stat | Description | Ontology_term | Ontology_id | Enzyme_code | Enzyme_name |
|---|---|---|---|---|---|---|---|---|---|---|
| 990_c0_g1 | | 2220 | 87 | 23 | 0 | | | | | |
| TRINITY_DN17241_c0_g1 | TRINITY_DN17241_c0_g1 | 940.10621 | 11.525007 | 0.7480258 | 15.40723 | TRINITY_DN17241_c0_g1 | NA | NA | NA | NA |
| TRINITY_DN4249_c0_g2 | TRINITY_DN4249_c0_g2 | 151.47730 | -25.563385 | 1.8182855 | -14.05906 | protein GAST1 isoform X2 | NA | NA | NA | NA |
| TRINIT | TRINITY_DN36061_c1_g1 | 96. | -24 | 1.9 | -12 | protein PELP | NA | NA | NA | NA |

| gene_id | gene_name | baseMean | log2FoldChange | lfcSE | stat | Description | Ontology_term | Ontology_id | Enzyme_code | Enzyme_name |
|---|---|---|---|---|---|---|---|---|---|---|
| Y_DN36061_c1_g1 | | 95591 | .792725 | 449776 | .74705 | K2-like | | | | |
| TRINITY_DN48909_c2_g1 | TRINITY_DN48909_c2_g1 | 125.91551 | 10.392925 | 0.8482170 | 12.25267 | hypothetical protein RGQ29_032212 | NA | NA | NA | NA |
| TRINITY_DN2203_c0_ | TRINITY_DN2203_c0_g1 | 48.08191 | 9.379273 | 0.8205472 | 11.43051 | TRINITY_DN2203_c0_g1 | NA | NA | NA | NA |

| gene_id | gene_name | baseMean | log2FoldChange | lfcSE | stat | Description | Ontology_term | Ontology_id | Enzyme_code | Enzyme_name |
|---|---|---|---|---|---|---|---|---|---|---|
| g1 | | | | | | | | | | |
| TRINITY_DN1466_c1_g1 | DVH24_034924; DVH24_042310; DVH24_021501; DVH24_042312; MdSOT3 | 1999.45949 | 3.860717 | 0.2804527 | 13.76602 | putative polyol transporter 1 | membrane,carbohydrate:proton symporter activity,carbohydrate transmembrane transport,proton transmembrane transport | GO:0016020, GO:0005351, GO:0034219, GO:1902600 | NA | NA |
| TRINITY_DN13723_c0_g1 | DVH24_029871; DVH24_026477 | 1951.74202 | -1.981992 | 0.0967896 | -20.47733 | hypothetical protein DVH24_029871 | membrane | GO:0016020 | NA | NA |

#### Table preview 27

**Rin vs control lfcThreshold = 1 results sorted by log2FC.**

| gene_id | gene_name | baseMean | log2FoldChange | lfcSE | stat | pvalue | Description | Ontology_term | Ontology_id | Enzyme_code | Enzyme_name |
|---|---|---|---|---|---|---|---|---|---|---|---|
| TRINITY_DN3584_c4_g2 | DVH24_014699; DVH24_011850; DVH24_018955; DVH24_025615; DVH24_015416; DVH24_009787; DVH24_002387 | 9.860234 | 13.735854 | 3.4593991 | 3.970589 | NA | hypothetical protein DVH24_025615 | regulation of cellular respiration | GO:0043457 | NA | NA |
| TRINITY_DN14899_c1_g1 | DVH24_023837; DVH24_023281; DVH24_006985 | 4.381821 | 13.484050 | 3.4593991 | 3.897801 | NA | hypothetical protein DVH24_023837, partial | NA | NA | NA | NA |
| TRINITY_DN16515_c0_g1 | TRINITY_DN16515_c0_g1 | 2205.329400 | 13.340787 | 0.8122674 | 16.424132 | 0 | TRINITY_DN16515_c0_g1 | NA | NA | NA | NA |
| TRINITY_D | TRINITY_DN23990_c0_g1 | 200 | 13.0 | 0.7 | 16. | 0 | TRINITY_DN2399 | NA | NA | NA | NA |

| gene_id | gene_name | baseMean | log2FoldChange | lfcSE | stat | pvalue | Description | Ontology_term | Ontology_id | Enzyme_code | Enzyme_name |
|---|---|---|---|---|---|---|---|---|---|---|---|
| N23990_c0_g1 | | 5.132204 | 16987 | 912823 | 450496 | | 0_c0_g1 | | | | |
| TRINITY_DN18768_c0_g1 | TRINITY_DN18768_c0_g1 | 4375.061310 | 12.5555326 | 0.6590477 | 19.050709 | 0 | TRINITY_DN18768_c0_g1 | NA | NA | NA | NA |
| TRINITY_DN17241_c0_g1 | TRINITY_DN17241_c0_g1 | 940.106214 | 11.525007 | 0.7480258 | 15.407232 | 0 | TRINITY_DN17241_c0_g1 | NA | NA | NA | NA |
| TRINITY_DN4 | TRINITY_DN48909_c2_g1 | 125. | 10.39 | 0.84 | 12.2 | 0 | hypothetical protein RGQ29_0 | NA | NA | NA | NA |

| gene_id | gene_name | baseMean | log2FoldChange | lfcSE | stat | pvalue | Description | Ontology_term | Ontology_id | Enzyme_code | Enzyme_name |
|---|---|---|---|---|---|---|---|---|---|---|---|
| 8909_c2_g1 | | 915511 | 2925 | 82170 | 52672 | | 32212 | | | | |
| TRINITY_DN10247_c0_g1 | TRINITY_DN10247_c0_g1 | 57.060087 | 9.649154 | 0.94670882 | 10.1923321 | 0 | TRINITY_DN10247_c0_g1 | NA | NA | NA | NA |
| TRINITY_DN6850_c0_g1 | TRINITY_DN6850_c0_g1 | 99.328697 | 9.487431 | 0.9142169 | 10.377658 | 0 | hypothetical protein R6Q59_010160 | NA | NA | NA | NA |
| TRINITY_DN2203_c0_g1 | TRINITY_DN2203_c0_g1 | 48.081907 | 9.379273 | 0.8205472 | 11.430511 | 0 | TRINITY_DN2203_c0_g1 | NA | NA | NA | NA |

#### Table preview 28

**Rin-high candidates in Rin vs control.**

| gene_id | gene_name | baseMean | log2FoldChange | lfcSE | stat | pvalue | Description | Ontology_term | Ontology_id | Enzyme_code | Enzyme_name |
|---|---|---|---|---|---|---|---|---|---|---|---|
| TRINITY_DN16515_c0_g1 | TRINITY_DN16515_c0_g1 | 2205.32940 | 13.340787 | 0.8122674 | 16.424132 | 0 | TRINITY_DN16515_c0_g1 | NA | NA | NA | NA |
| TRINITY_DN23990_c0_g1 | TRINITY_DN23990_c0_g1 | 2005.13220 | 13.016987 | 0.7912823 | 16.450496 | 0 | TRINITY_DN23990_c0_g1 | NA | NA | NA | NA |
| TRINITY_DN18768_c0_g1 | TRINITY_DN18768_c0_g1 | 4375.06131 | 12.555326 | 0.6590477 | 19.050709 | 0 | TRINITY_DN18768_c0_g1 | NA | NA | NA | NA |
| TRINITY_DN17241_c0_ | TRINITY_DN17241_c0_g1 | 940.10 | 11.525007 | 0.748 | 15.40 | 0 | TRINITY_DN17241_c0_g1 | NA | NA | NA | NA |

| gene_id | gene_name | baseMean | log2FoldChange | lfcSE | stat | pvalue | Description | Ontology_term | Ontology_id | Enzyme_code | Enzyme_name |
|---|---|---|---|---|---|---|---|---|---|---|---|
| g1 | | 621 | | 0258 | 7232 | | | | | | |
| TRINITY_DN48909_c2_g1 | TRINITY_DN48909_c2_g1 | 125.91551 | 10.392925 | 0.84882170 | 12.252672 | 0 | hypothetical protein RGQ29_032212 | NA | NA | NA | NA |
| TRINITY_DN10247_c0_g1 | TRINITY_DN10247_c0_g1 | 57.06009 | 9.649154 | 0.9467082 | 10.192321 | 0 | TRINITY_DN10247_c0_g1 | NA | NA | NA | NA |
| TRINITY_DN6850_c0_g1 | TRINITY_DN6850_c0_g1 | 99.32870 | 9.487431 | 0.9142169 | 10.377658 | 0 | hypothetical protein R6Q59_010160 | NA | NA | NA | NA |
| TRINITY_DN2203_ | TRINITY_DN2203_c0_g1 | 48.08 | 9.379273 | 0.82 | 11.4 | 0 | TRINITY_DN2203_c0_g1 | NA | NA | NA | NA |

| gene_id | gene_name | baseMean | log2FoldChange | lfcSE | stat | pvalue | Description | Ontology_term | Ontology_id | Enzyme_code | Enzyme_name |
|---|---|---|---|---|---|---|---|---|---|---|---|
| c0_g1 | | 191 | | 05472 | 30511 | | | | | | |
| TRINITY_DN14467_c0_g1 | DVH24_017761; DVH24_029144 | 181.779277 | 9.271246 | 1.18391118 | 7.83310281 | 0 | uclacyanin 1-like | plasma membrane,electron transfer activity | GO:0005886,GO:0009055 | NA | NA |
| TRINITY_DN38134_c0_g1 | TRINITY_DN38134_c0_g1 | 46.18077 | 9.112993 | 0.85957766 | 10.6001724 | 0 | TRINITY_DN38134_c0_g1 | NA | NA | NA | NA |

#### Table preview 29

**Control-high candidates in Rin vs control.**

| gene_id | gene_name | baseMean | log2FoldChange | lfcSE | stat | pvalue | Description | Ontology_term | Ontology_id | Enzyme_code | Enzyme_name |
|---|---|---|---|---|---|---|---|---|---|---|---|
| TRINITY_DN4249_c0_g2 | TRINITY_DN4249_c0_g2 | 151.477730 | -25.56339 | 1.818285 | -14.059060 | 0 | protein GAST1 isoform X2 | NA | NA | NA | NA |
| TRINITY_DN36061_c1_g1 | TRINITY_DN36061_c1_g1 | 96.95591 | -24.79272 | 1.944978 | -12.747049 | 0 | protein PELPK2-like | NA | NA | NA | NA |
| TRINITY_DN229006_c0_g1 | TRINITY_DN229006_c0_g1 | 35.817112 | -24.13405 | 3.383951 | -7.131914 | 0 | TRINITY_DN229006_c0_g1 | NA | NA | NA | NA |
| TRINITY_DN139024_c | TRINITY_DN139024_c0_g1 | 24.94 | -23.3102 | 3.384 | -6.88 | 0 | TRINITY_DN139024_c0_g1 | NA | NA | NA | NA |

| gene_id | gene_name | baseMean | log2FoldChange | lfcSE | stat | pvalue | Description | Ontology_term | Ontology_id | Enzyme_code | Enzyme_name |
|---|---|---|---|---|---|---|---|---|---|---|---|
| 0_g1 | | 417 | 4 | 403 | 7547 | | | | | | |
| TRINITY_DN138657_c0_g1 | TRINITY_DN138657_c0_g1 | 14.178807 | -22.95078 | 3.2795503 | -6.9982249 | 0 | TRINITY_DN138657_c0_g1 | NA | NA | NA | NA |
| TRINITY_DN11246_c0_g1 | TRINITY_DN11246_c0_g1 | 20.84368 | -22.77082 | 3.3471894 | -6.8029669 | 0 | TRINITY_DN11246_c0_g1 | NA | NA | NA | NA |
| TRINITY_DN6336_c0_g1 | TRINITY_DN6336_c0_g1 | 19.07138 | -22.66635 | 3.3400660 | -6.7862122 | 0 | TRINITY_DN6336_c0_g1 | NA | NA | NA | NA |
| TRINITY | TRINITY_DN81325 | 11 | -22. | 3. | -6 | 0 | TRINITY_DN81325_ | NA | NA | NA | NA |

| gene_id | gene_name | baseMean | log2FoldChange | lfcSE | stat | pvalue | Description | Ontology_term | Ontology_id | Enzyme_code | Enzyme_name |
|---|---|---|---|---|---|---|---|---|---|---|---|
| _DN81325_c0_g2 | _c0_g2 | .663340 | 59586 | 236148 | .9823332 | | c0_g2 | | | | |
| TRINITY_DN18390_c0_g1 | DVH24_033020; DVH24_038538; DVH24_017268 | 14.378688 | -22.42292 | 2.708501 | -8.2788721 | 0 | alpha-glucan phosphorylase, H isozyme-like | 1,4-alpha-oligoglucan phosphorylase activity | GO:0004645 | EC:2.4.1.1 | glycogen phosphorylase |
| TRINITY_DN21772_c0_g1 | TRINITY_DN21772_c0_g1 | 15.981171 | -22.09229 | 3.296637 | -6.7014462 | 0 | TRINITY_DN21772_c0_g1 | NA | NA | NA | NA |

#### Table preview 30

**Rin vs control candidates sorted by absolute log2FC.**

| gene_id | gene_name | baseMean | log2FoldChange | lfcSE | stat | pvalue | direction | Description | Ontology_term | Ontology_id | Enzyme_code | Enzyme_name |
|---|---|---|---|---|---|---|---|---|---|---|---|---|
| TRINITY_DN4249_c0_g2 | TRINITY_DN4249_c0_g2 | 151.477730 | -25.56339 | 1.818285 | -14.059060 | 0 | C-high | protein GAST1 isoform X2 | NA | NA | NA | NA |
| TRINITY_DN36061_c1_g1 | TRINITY_DN36061_c1_g1 | 96.95591 | -24.79272 | 1.94978 | -12.747049 | 0 | C-high | protein PELPK2-like | NA | NA | NA | NA |
| TRINITY_DN229006_c0_g1 | TRINITY_DN229006_c0_g1 | 35.81712 | -24.13405 | 3.383951 | -7.131914 | 0 | C-high | TRINITY_DN229006_c0_g1 | NA | NA | NA | NA |
| TRINITY_DN13902 | TRINITY_DN139024_c0_g1 | 24.94 | -23.3102 | 3.384 | -6.88 | 0 | C-hig | TRINITY_DN139024_c0_g1 | NA | NA | NA | NA |

| gene_id | gene_name | baseMean | log2FoldChange | lfcSE | stat | pvalue | direction | Description | Ontology_term | Ontology_id | Enzyme_code | Enzyme_name |
|---|---|---|---|---|---|---|---|---|---|---|---|---|
| 4_c0_g1 | | 417 | 4 | 403 | 7547 | | h | | | | | |
| TRINITY_DN138657_c0_g1 | TRINITY_DN138657_c0_g1 | 14.17807 | -22.95078 | 3.279503 | -6.998249 | 0 | C-high | TRINITY_DN138657_c0_g1 | NA | NA | NA | NA |
| TRINITY_DN11246_c0_g1 | TRINITY_DN11246_c0_g1 | 20.84368 | -22.77082 | 3.347189 | -6.802969 | 0 | C-high | TRINITY_DN11246_c0_g1 | NA | NA | NA | NA |
| TRINITY_DN6336_c0_g1 | TRINITY_DN6336_c0_g1 | 19.07138 | -22.66635 | 3.340060 | -6.786212 | 0 | C-high | TRINITY_DN6336_c0_g1 | NA | NA | NA | NA |
| TRINIT | TRINITY_DN81325 | 11 | -22. | 3. | -6 | 0 | C- | TRINITY_DN81325_ | NA | NA | NA | NA |

| gene_id | gene_name | baseMean | log2FoldChange | lfcSE | stat | pvalue | direction | Description | Ontology_term | Ontology_id | Enzyme_code | Enzyme_name |
|---|---|---|---|---|---|---|---|---|---|---|---|---|
| Y_DN81325_c0_g2 | _c0_g2 | .663340 | 59586 | 236148 | .982332 | | high | c0_g2 | | | | |
| TRINITY_DN18390_c0_g1 | DVH24_033020; DVH24_038538; DVH24_017268 | 14.378868 | -2.42292 | 2.708501 | -8.278721 | 0 | C-high | alpha-glucan phosphorylase, H isozyme-like | 1,4-alpha-oligoglucan phosphorylase activity | GO:0004645 | EC:2.4.1.1 | glycogen phosphorylase |
| TRINITY_DN21772_c0_g1 | TRINITY_DN21772_c0_g1 | 15.981771 | -2.09229 | 3.296637 | -6.701462 | 0 | C-high | TRINITY_DN21772_c0_g1 | NA | NA | NA | NA |

#### Table preview 31

**Rola vs control DESeq2 results using lfcThreshold = 1.**

| gene_id | gene_name | baseMean | log2FoldChange | Description | Ontology_term | Ontology_id | Enzyme_code | Enzyme_name |
|---|---|---|---|---|---|---|---|---|
| TRINITY_DN18768_c0_g1 | TRINITY_DN18768_c0_g1 | 4375.06131 | 12.55152 | TRINITY_DN18768_c0_g1 | NA | NA | NA | NA |
| TRINITY_DN165 | TRINITY_DN16515_c0_g1 | 2205.32940 | 12.97620 | TRINITY_DN16515_c0_g1 | NA | NA | NA | NA |

| gene_id | gene_name | baseMean | log2FoldChange | Description | Ontology_term | Ontology_id | Enzyme_code | Enzyme_name |
|---|---|---|---|---|---|---|---|---|
| 15_c0_g1 | | | | | | | | |
| TRINITY_DN23990_c0_g1 | TRINITY_DN23990_c0_g1 | 2005.132220 | 12.381991 | TRINITY_DN23990_c0_g1 | NA | NA | NA | NA |
| TRINIT | TRINITY_DN17241_c0_g1 | 940.1 | 10.66 | TRINITY_DN17241_c0_g1 | NA | NA | NA | NA |

| gene_id | gene_name | baseMean | log2FoldChange | Description | Ontology_term | Ontology_id | Enzyme_code | Enzyme_name |
|---|---|---|---|---|---|---|---|---|
| Y_DN172441_c0_g1 | | 0621 | 039 | | | | | |
| TRINITY_DN81098_c0_ | DVH24_016150; DVH24_003921; DVH24_016152; DVH24_016153 | 124.28616 | 11.33609 | ADP,ATP carrier protein 1, mitochondrial-like | mitochondrial inner membrane,ATP:ADP antiporter activity,mitochondrial ADP transmembrane transport,mitochondrial ATP transmembrane transport | GO:0005743,GO:0005471,GO:0140021,GO:1990544 | NA | NA |

| gene_id | gene_name | baseMean | log2FoldChange | Description | Ontology_term | Ontology_id | Enzyme_code | Enzyme_name |
|---|---|---|---|---|---|---|---|---|
| g1 | | | | | | | | |
| TRINITY_DN27539_c0_g1 | atp9; ATP9; DVH24_038917 | 97.27470 | 10.97710 | ATP synthase F0 subunit 9 | membrane | GO:0016020 | NA | NA |
| TRINITY_DN2 | DVH24_038846; DVH24_042792; DVH24_005917 | 146.15273 | 11.50787 | protein disulfide-isomerase-like | NA | NA | NA | NA |

| gene_id | gene_name | baseMean | log2FoldChange | Description | Ontology_term | Ontology_id | Enzyme_code | Enzyme_name |
|---|---|---|---|---|---|---|---|---|
| 84833_c1_g1 | | | | | | | | |
| TRINITY_DN48040_c0_g1 | DVH24_000798; DVH24_034180 | 59.8713031 | 10.3553033 | adenosylhomocysteinase | NA | NA | NA | NA |
| TRI | DVH24_005516; | 63. | 10. | 5-methyltetrahydropte | methyltransferase activity | GO:0008168 | EC: | Transferi |

| gene_id | gene_name | baseMean | log2FoldChange | Description | Ontology_term | Ontology_id | Enzyme_code | Enzyme_name |
|---|---|---|---|---|---|---|---|---|
| NITY_DN24548_c1_g1 | DVH24_035080; DVH24_019035 | 96699 | 42096 | royltriglutamate–homocysteine methyltransferase | | | 2.1.1 | ng one-carbon groups |
| TRINITY_DN21847_c | DVH24_040656 | 153.181381388 | 11.2715155 | eukaryotic translation initiation factor 5A-2 | NA | NA | NA | NA |

| gene_id | gene_name | baseMean | log2FoldChange | Description | Ontology_term | Ontology_id | Enzyme_code | Enzyme_name |
|---|---|---|---|---|---|---|---|---|
| 0_g1 | | | | | | | | |

#### Table preview 32

**Rola vs control lfcThreshold = 1 results sorted by log2FC.**

| gene_id | gene_name | baseMean | log2FoldChange | lfcSE | stat | pvalue | Description | Ontology_term | Ontology_id | Enzyme_code | Enzyme_name |
|---|---|---|---|---|---|---|---|---|---|---|---|
| TRINITY_DN3584_c4_g2 | DVH24_014699; DVH24_011850; DVH24_018955; DVH24_025615; DVH24_015416; DVH24_009787; DVH24_002387 | 9.860234 | 22.25660 | 3.8387321 | 5.797904 | NA | hypothetical protein DVH24_025615 | regulation of cellular respiration | GO:0043457 | NA | NA |
| TRI | DVH24_014699; | 4 | 2 | 3 | 5 | N | hypotheti | NA | N | N | N |

| gene_id | gene_name | baseMean | log2FoldChange | lfcSE | stat | pvalue | Description | Ontology_term | Ontology_id | Enzyme_code | Enzyme_name |
|---|---|---|---|---|---|---|---|---|---|---|---|
| NITY_DN8803_c7_g1 | DVH24_023281; DVH24_015416; DVH24_006985; DVH24_009787; DVH24_002387; DVH24_037524 | .236892 | 1.60487 | .8405733 | .625429 | A | cal protein DVH24_033647 | | A | A | A |
| TRINITY_DN3584_c1_g2 | TRINITY_DN3584_c1_g2 | 3.4340366 | 21.33084 | 3.8413289 | 5.552985 | NA | Regulator of rDNA transcription protein 15 | NA | NA | NA | NA |
| TRINITY_DN14899_c1_g1 | DVH24_023837; DVH24_023281; DVH24_006985 | 4.3818221 | 21.13457 | 3.8404877 | 5.503095 | NA | hypothetical protein DVH24_023837, partial | NA | NA | NA | NA |
| TRINITY_DN1674_c0_g1 | TRINITY_DN1674_c0_g1 | 593.861 | 13.65050 | 1.83093 | 7.45548 | 0 | TRINITY_DN1674_c0_g1 | NA | NA | NA | NA |

| gene_id | gene_name | baseMean | log2FoldChange | lfcSE | stat | pvalue | Description | Ontology_term | Ontology_id | Enzyme_code | Enzyme_name |
|---|---|---|---|---|---|---|---|---|---|---|---|
| | | 570 | | 51 | 0 | | | | | | |
| TRINITY_DN16515_c0_g1 | TRINITY_DN16515_c0_g1 | 2205.329400 | 12.976220 | 0.8502320 | 15.261957 | 0 | TRINITY_DN16515_c0_g1 | NA | NA | NA | NA |
| TRINITY_DN18768_c0_g1 | TRINITY_DN18768_c0_g1 | 4375.061310 | 12.55152 | 0.7081131 | 17.725305 | 0 | TRINITY_DN18768_c0_g1 | NA | NA | NA | NA |
| TRINITY_DN23990_c0_g1 | TRINITY_DN23990_c0_g1 | 2005.132 | 12.38191 | 0.831843 | 14.88490 | 0 | TRINITY_DN23990_c0_g1 | NA | NA | NA | NA |

| gene_id | gene_name | baseMean | log2FoldChange | lfcSE | stat | pvalue | Description | Ontology_term | Ontology_id | Enzyme_code | Enzyme_name |
|---|---|---|---|---|---|---|---|---|---|---|---|
| | | 204 | | 2 | 7 | | | | | | |
| TRINITY_DN25322_c0_g1 | TRINITY_DN25322_c0_g1 | 206.141378 | 12.10057 | 1.3700964 | 8.831910 | 0 | TRINITY_DN25322_c0_g1 | NA | NA | NA | NA |
| TRINITY_DN7837_c0_g1 | ycf1 | 190.616264 | 12.05235 | 1.6022115 | 7.522323 | 0 | hypothetical chloroplast RF1 | NA | NA | NA | NA |

#### Table preview 33

**Rola-high candidates in Rola vs control.**

| gene_id | gene_name | baseMean | log2FoldChange | lfcSE | stat | pvalue | padj | Description | Ontology_term | Ontology_id | Enzyme_code | Enzyme_name |
|---|---|---|---|---|---|---|---|---|---|---|---|---|
| TRINITY_DN1674_c0_g1 | TRINITY_DN1674_c0_g1 | 593.8616 | 13.65050 | 1.83093351 | 7.455480 | 0 | 0.0e+00 | TRINITY_DN1674_c0_g1 | NA | NA | NA | NA |
| TRINITY_DN16515_c0_g1 | TRINITY_DN16515_c0_g1 | 2205.3294 | 12.97620 | 0.85023220 | 15.261957 | 0 | 0.0e+00 | TRINITY_DN16515_c0_g1 | NA | NA | NA | NA |
| TRINITY_DN18768_c0_g1 | TRINITY_DN18768_c0_g1 | 4375.0613 | 12.55152 | 0.70811331 | 17.725305 | 0 | 0.0e+00 | TRINITY_DN18768_c0_g1 | NA | NA | NA | NA |
| TRINITY_DN23990_c0_g1 | TRINITY_DN23990_c0_g1 | 2005.13 | 12.38191 | 0.831184 | 14.8849 | 0 | 0.0e+0 | TRINITY_DN23990_c0_g1 | NA | NA | NA | NA |

| gene_id | gene_name | baseMean | log2FoldChange | lfcSE | stat | pvalue | padj | Description | Ontology_term | Ontology_id | Enzyme_code | Enzyme_name |
|---|---|---|---|---|---|---|---|---|---|---|---|---|
| | | 22 | | 32 | 07 | | 0 | | | | | |
| TRINITY_DN25322_c0_g1 | TRINITY_DN25322_c0_g1 | 2206.1414 | 12.10057 | 1.37009664 | 8.831910 | 0 | 0.00e+00 | TRINITY_DN25322_c0_g1 | NA | NA | NA | NA |
| TRINITY_DN7837_c0_g1 | ycf1 | 1900.6163 | 12.05235 | 1.60221115 | 7.522323 | 0 | 0.00e+00 | hypothetical chloroplast RF1 | NA | NA | NA | NA |
| TRINITY_DN96438_c0_g1 | HYH03_006855; KSW81_004590; WJX72_006455; C2E20_7425; MNEG_1885; KSW81_004591 | 1700.2185 | 11.85476 | 1.34840052 | 8.7916690 | 0 | 0.00e+00 | fasciclin-like protein | NA | NA | NA | NA |
| TRINITY_DN4597 | TRINITY_DN45973_c0_g1 | 170. | 11.83907 | 1.99 | 5.93 | 0 | 1.3e | TRINITY_DN45973_c0_g1 | NA | NA | NA | NA |

| gene_id | gene_name | baseMean | log2FoldChange | lfcSE | stat | pvalue | padj | Description | Ontology_term | Ontology_id | Enzyme_code | Enzyme_name |
|---|---|---|---|---|---|---|---|---|---|---|---|---|
| 3_c0_g1 | | 5588 | | 40685 | 7142 | | -06 | | | | | |
| TRINITY_DN2665_c0_g1 | TRINITY_DN2665_c0_g1 | 155.7880 | 11.6921 | 1.2487385 | 9.344795 | 0 | 0.00e+00 | TRINITY_DN2665_c0_g1 | NA | NA | NA | NA |
| TRINITY_DN6785_c0_g1 | TRINITY_DN6785_c0_g1 | 145.0172 | 11.6516 | 1.9609385 | 5.948762 | 0 | 1.3e-06 | TRINITY_DN6785_c0_g1 | NA | NA | NA | NA |

#### Table preview 34

**Control-high candidates in Rola vs control.**

| gene_name | Description | Ontology_term | Ontology_id | Enzyme_code | Enzyme_name |
|---|---|---|---|---|---|
| DVH24_042242 | hypothetical protein DVH24_042242 | nucleus,3-phosphoinositide-dependent protein kinase activity,DNA-dependent protein kinase activity,AMP-activated protein kinase activity,eukaryotic translation initiation factor 2alpha kinase activity,ribosomal protein S6 kinase activity,ATP binding,histone H3S10 kinase activity,histone H3T11 kinase activity,histone H3T6 kinase activity,histone H2AXS139 kinase activity,histone H3S28 kinase activity,histone H4S1 kinase activity,histone H2AS1 kinase activity,histone H2BS14 kinase activity,histone H3T3 kinase activity,histone H2AS121 kinase activity,Rho-dependent protein serine/threonine kinase activity,histone H2BS36 kinase activity,histone H3S57 kinase activity,histone H3T45 kinase activity,histone H2AT120 kinase activity,nuclear-transcribed mRNA catabolic process, nonsense-mediated decay,chromatin remodeling | GO:0005634,GO:0004676,GO:0004677,GO:0004679,GO:0004694,GO:0004711,GO:0005524,GO:0035175,GO:0035402,GO:0035403,GO:0035979,GO:0044022,GO:0044023,GO:0044024,GO:0044025,GO:0072354,GO:0072371,GO:0072518,GO:0140823,GO:0140855,GO:0140857,GO:1990244,GO:0000184,GO:0006338 | EC:2.7.11.1,EC:2.7.11.29,EC:2.7.11 | non-specific serine/threonine protein kinase,[low-density-lipoprotein receptor] kinase,dephospho-[reductase kinase] kinase,tropomyosin kinase,[Goodpasture-antigen-binding protein] kinase |

| gene_name | Description | Ontology_term | Ontology_id | Enzyme_code | Enzyme_name |
| --- | --- | --- | --- | --- | --- |
| | | | | .3,EC:2.7.11.28,EC:2.7.11.9 | |
| DVH24_0 | unchara | response to cold,regulation of circadian rhythm | GO:0009409,GO:0042752 | NA | NA |

| gene_name | Description | Ontology_term | Ontology_id | Enzyme_code | Enzyme_name |
| --- | --- | --- | --- | --- | --- |
| 24718; DVH24_022573 | cterized protein | | | | |
| DVH24_022350; D | two-component r | nucleus,phosphorelay signal transduction system,cytokinin-activated signaling pathway,rhythmic process | GO:0005634,GO:0000160,GO:0009736,GO:0048511 | NA | NA |

| gene_name | Description | Ontology_term | Ontology_id | Enzyme_code | Enzyme_name |
|---|---|---|---|---|---|
| VH24_0249886 | esponse regulator-like APRR5 | | | | |
| TRINIT | 18.1 k | NA | NA | NA | NA |

| gene_name | Description | Ontology_term | Ontology_id | Enzyme_code | Enzyme_name |
|---|---|---|---|---|---|
| Y_DN182589_c0_g1 | Da class I heat shock protein-like | | | | |

| gene_name | Description | Ontology_term | Ontology_id | Enzyme_code | Enzyme_name |
|---|---|---|---|---|---|
| DVH24_00609 7; DVH24_030860; DVH24_ | chaperone protein dnaJ C76, chloro | membrane | GO:0016020 | NA | NA |

| gene_name | Description | Ontology_term | Ontology_id | Enzyme_code | Enzyme_name |
|---|---|---|---|---|---|
| 0401977 | plastic | | | | |
| DVH24_01245 6;DVH24_020724; | cold-regulated protein 27 | Golgi cisterna membrane,fucosyltransferase activity | GO:0032580,GO:0008417 | EC:2.4.1 | Glycosyltransferases |

| gene_name | Description | Ontology_term | Ontology_id | Enzyme_code | Enzyme_name |
| --- | --- | --- | --- | --- | --- |
| DVH24_021640 TRINITY_DN64355_c0_g1 | protein argonaute 18- | NA | NA | NA | NA |

| gene_name | Description | Ontology_term | Ontology_id | Enzyme_code | Enzyme_name |
|---|---|---|---|---|---|
| DVH24_005391; DVH24_022350; DV | like two-component response regulat | nucleus,phosphorelay signal transduction system,cytokinin-activated signaling pathway,rhythmic process | GO:0005634,GO:0000160,GO:0009736,GO:0048511 | NA | NA |

| gene_name | Description | Ontology_term | Ontology_id | Enzyme_code | Enzyme_name |
| --- | --- | --- | --- | --- | --- |
| H24_0249886 | or-like APRR5 | | | | |
| DVH24_0053991; DVH24_0 | two-component respons | nucleus,phosphorelay signal transduction system,cytokinin-activated signaling pathway,rhythmic process | GO:0005634,GO:0000160,GO:0009736,GO:0048511 | NA | NA |

| gene_name | Description | Ontology_term | Ontology_id | Enzyme_code | Enzyme_name |
|---|---|---|---|---|---|
| 22350;DVH24_02498 6 | e regulator-like APRR5 | | | | |
| DVH24_022350 | two-componen | nucleus,phosphorelay signal transduction system,cytokinin-activated signaling pathway,rhythmic process | GO:0005634,GO:0000160,GO:0009736,GO:0048511 | NA | NA |

| gene_name | Description | Ontology_term | Ontology_id | Enzyme_code | Enzyme_name |
|---|---|---|---|---|---|
| ; DVH24_024986 | t response regulator-like APRR5 | | | | |

#### Table preview 35

**Rola vs control candidates sorted by absolute log2FC.**

| gene_id | gene_name | baseMean | log2FoldChange | lfcSE | stat | pvalue | padj | direction | Description | Ontology_term | Ontology_id | Enzyme_code | Enzyme_name |
|---|---|---|---|---|---|---|---|---|---|---|---|---|---|
| TRINITY_DN1674_c0_g1 | TRINITY_DN1674_c0_g1 | 593.8616 | 13.65050 | 1.8309351 | 7.455480 | 0 | 0.0e+00 | Rola-high | TRINITY_DN1674_c0_g1 | NA | NA | NA | NA |
| TRINITY_DN16515_c0_g1 | TRINITY_DN16515_c0_g1 | 2205.3294 | 12.97620 | 0.8502320 | 15.261957 | 0 | 0.0e+00 | Rola-high | TRINITY_DN16515_c0_g1 | NA | NA | NA | NA |
| TRINITY_DN18768_c0_g1 | TRINITY_DN18768_c0_g1 | 4375.0613 | 12.55152 | 0.7081131 | 17.725305 | 0 | 0.0e+00 | Rola-high | TRINITY_DN18768_c0_g1 | NA | NA | NA | NA |
| TRINITY_DN23990_c0_g1 | TRINITY_DN23990_c0_g1 | 2005.13 | 12.38191 | 0.83184 | 14.8849 | 0 | 0.0e+0 | Rola-hi | TRINITY_DN23990_c0_g1 | NA | NA | NA | NA |

| gene_id | gene_name | baseMean | log2FoldChange | lfcSE | stat | pvalue | padj | direction | Description | Ontology_term | Ontology_id | Enzyme_code | Enzyme_name |
|---|---|---|---|---|---|---|---|---|---|---|---|---|---|
| | | 22 | | 32 | 07 | | 0 | gh | | | | | |
| TRINITY_DN25322_c0_g1 | TRINITY_DN25322_c0_g1 | 206.1414 | 12.10057 | 1.3700964 | 8.831910 | 0 | 0.00e+00 | Rola-high | TRINITY_DN25322_c0_g1 | NA | NA | NA | NA |
| TRINITY_DN7837_c0_g1 | ycf1 | 190.6163 | 12.05235 | 1.6022115 | 7.522323 | 0 | 0.00e+00 | Rola-high | hypothetical chloroplast RF1 | NA | NA | NA | NA |
| TRINITY_DN96438_c0_g1 | HYH03_006855; KSW81_004590; WJX72_006455; C2E20_7425; MNEG_1885; KSW81_004591 | 170.2185 | 11.85476 | 1.3484052 | 8.791690 | 0 | 0.00e+00 | Rola-high | fasciclin-like protein | NA | NA | NA | NA |
| TRINITY_DN45 | TRINITY_DN45973_c0_g1 | 170. | 11.83907 | 1.999 | 5.93 | 0 | 1.3e | Rola | TRINITY_DN45973_c0_g1 | NA | NA | NA | NA |

| gene_id | gene_name | baseMean | log2FoldChange | lfcSE | stat | pvalue | padj | direction | Description | Ontology_term | Ontology_id | Enzyme_code | Enzyme_name |
|---|---|---|---|---|---|---|---|---|---|---|---|---|---|
| 973_c0_g1 | | 5588 | | 40685 | 7142 | | -06 | -high | | | | | |
| TRINITY_DN2665_c0_g1 | TRINITY_DN2665_c0_g1 | 155.7880 | 11.6921 | 1.2487385 | 9.344795 | 0 | 0.0e+00 | Rola-high | TRINITY_DN2665_c0_g1 | NA | NA | NA | NA |
| TRINITY_DN6785_c0_g1 | TRINITY_DN6785_c0_g1 | 145.0172 | 11.6516 | 1.9609385 | 5.948762 | 0 | 1.3e-06 | Rola-high | TRINITY_DN6785_c0_g1 | NA | NA | NA | NA |

#### Table preview 36

**Combined comparison table across Rin vs control, Rola vs control, and Rin vs Rola.**

| gene_id | gene_name | Description | Ontology_term | Ontology_id | baseMean_Rola_vs_C | baseMean_Rin_vs_Rola | log2FC_Rin_vs_Rola | pvalue_Rin_vs_Rola |
|---|---|---|---|---|---|---|---|---|
| TRINITY_DN0_c0_g1 | TRINITY_DN0_c0_g1 | TRINITY_DN0_c0_g1 | NA | NA | 7.087555 | 7.087555 | -0.3416879 | 0.8659393 |
| TRINITY_ | DVH24_00 | photosystem II | chloroplast,membrane | GO:0009507,GO:0016020 | 1.926921 | 1.926921 | 0.206221 | 0.905444 |

| gene_id | gene_name | Description | Ontology_term | Ontology_id | baseMean_Rola_vs_C | baseMean_Rin_vs_Rola | log2FC_Rin_vs_Rola | pvalue_Rin_vs_Rola |
|---|---|---|---|---|---|---|---|---|
| DN0_c0_g2 | 6911;DVH24_042775 | 22kDa protein, chloroplastic | | | | | 7 | 4 |
| TRINITY_D | TRINITY_DN | TRINITY_DN1000 | NA | NA | NA | NA | NA | NA |

| gene_id | gene_name | Description | Ontology_term | Ontology_id | baseMean_Rola_vs_C | baseMean_Rin_vs_Rola | log2FC_Rin_vs_Rola | pvalue_Rin_vs_Rola |
|---|---|---|---|---|---|---|---|---|
| N100003_c0_g1 | 100003_c0_g1 | 03_c0_g1 | | | | | | |
| TRINITY_DN1010 | DVH24_031902; | large ribosomal subuni | ribosome,ribonucleoprotein complex,DNA binding,structural constituent of ribosome,regulation of DNA-templated transcription,translation | GO:0005840,GO:1990904,GO:0003677,GO:0003735,GO:0006355,GO:00 | 6.119179 | 6.119179 | -0.1572277 | 0.9445742 |

| gene_id | gene_name | Description | Ontology_term | Ontology_id | baseMean_Rola_vs_C | baseMean_Rin_vs_Rola | log2FC_Rin_vs_Rola | pvalue_Rin_vs_Rola |
|---|---|---|---|---|---|---|---|---|
| 0005_c0_g1 | DVH24_007180;DVH24_037118 | t protein eL34-like | | 06412 | | | | |
| T | T | T | NA | NA | 2 | 2 | 0 | 0 |

| gene_id | gene_name | Description | Ontology_term | Ontology_id | baseMean_Rola_vs_C | baseMean_Rin_vs_Rola | log2FC_Rin_vs_Rola | pvalue_Rin_vs_Rola |
|---|---|---|---|---|---|---|---|---|
| RINITY_DN10000_c0_g1 | RINITY_DN10000_c0_g1 | RINITY_DN10000_c0_g1 | | | 6.512704 | 6.512704 | .0254975 | .9998474 |
| TRINI | TRINIT | TRINITY_ | NA | NA | 85.35 | 85.35 | -0.37 | 0.976 |

| gene_id | gene_name | Description | Ontology_term | Ontology_id | baseMean_Rola_vs_C | baseMean_Rin_vs_Rola | log2FC_Rin_vs_Rola | pvalue_Rin_vs_Rola |
|---|---|---|---|---|---|---|---|---|
| TY_DN10000_c0_g2 | Y_DN10000_c0_g2 | DN10000_c0_g2 | | | 9205 | 9205 | 46438 | 1327 |
| TRINITY_D | FH972_023 | hypotheticalprotei | NA | NA | 0.556098 | 0.556098 | -0.735503 | 0.761842 |

| gene_id | gene_name | Description | Ontology_term | Ontology_id | baseMean_Rola_vs_C | baseMean_Rin_vs_Rola | log2FC_Rin_vs_Rola | pvalue_Rin_vs_Rola |
|---|---|---|---|---|---|---|---|---|
| N1000010_c0_g1 | 986 | nFH972_023986 | | | | | 7 | |
| TRINITY_DN10 | DVH24_013063; | dehydrin COR47-like | cytosol,membrane,response to water deprivation,cold acclimation,response to abscisic acid | GO:0005829,GO:0016020,GO:0009414,GO:0009631,GO:0009737 | 443.33930 | 443.33930 | -1.7230259 | 0.0391361 |

| gene_id | gene_name | Description | Ontology_term | Ontology_id | baseMean_Rola_vs_C | baseMean_Rin_vs_Rola | log2FC_Rin_vs_Rola | pvalue_Rin_vs_Rola |
|---|---|---|---|---|---|---|---|---|
| 0010_c4_g1 | DVH24_020054;DHN8;DHN9 | | | | | | | |
| TRINI | DVH24 | proteinTI | TOC-TIC supercomplex I,heme binding,metal ion binding,lactopero | GO:0061927,GO:0020037,GO: | 3219. | 3219. | 0.299 | 0.999 |

| gene_id | gene_name | Description | Ontology_term | Ontology_id | baseMean_Rola_vs_C | baseMean_Rin_vs_Rola | log2FC_Rin_vs_Rola | pvalue_Rin_vs_Rola |
|---|---|---|---|---|---|---|---|---|
| TY_DN10001_c0_g1 | _038405;DVH24_041706 | C110, chloroplastic-like | xidase activity,response to oxidative stress,hydrogen peroxide catabolic process,protein import into chloroplast stroma,cellular oxidant detoxification | 0046872,GO:0140825,GO:0006979,GO:0042744,GO:0045037,GO:0098869 | 040665 | 040665 | 7496 | 9999 |
| TRINIT | DVH24_ | protein TIC | TOC-TIC supercomplex I,heme binding,metal ion binding,lactoperoxidase | GO:0061927,GO:0020037,GO:00468 | 2.0741 | 2.0741 | 0.5416 | 0.7378 |

| gene_id | gene_name | Description | Ontology_term | Ontology_id | baseMean_Rola_vs_C | baseMean_Rin_vs_Rola | log2FC_Rin_vs_Rola | pvalue_Rin_vs_Rola |
|---|---|---|---|---|---|---|---|---|
| Y_DN10001_c1_g1 | 038405;DVH24_041706 | 110,chloroplastic | activity,response to oxidative stress,hydrogen peroxide catabolic process,protein import into chloroplast stroma,cellular oxidant detoxification | 72,GO:0140825,GO:0006979,GO:0042744,GO:0045037,GO:0098869 | 44 | 44 | 216 | 206 |

#### Table preview 37

**Classification summary of expression-response classes.**

| class | n |
|---|---|
| not_strong_DE | 242655 |
| Rola_specific_induced | 3352 |

| class | n |
|---|---|
| common_infection_induced | 2313 |
| Rin_specific_induced | 864 |
| Rin_specific_repressed | 728 |
| Rola_specific_repressed | 69 |
| common_infection_repressed | 2 |
