## Supplementary tables and figures for "Transcriptome of apple cv. Ametyst in *Rvi6*-effective and *Rvi6*-breaking *Venturia inaequalis* interactions identifies defense candidates": 24_SupplementaryTable_S15_screening_summary.pdf

**Supplementary Table S15. Contamination and adaptor screening of the *de novo* gene-representative set.**

Summary of the NCBI FCS-GX and FCS-adaptor screens described in Materials and methods, gathered here because the individual figures are otherwise distributed across Table 1 and Supplementary Tables S5, S10 and S11. FCS-GX 0.5.5 was run against the GX reference database build of 24 January 2023 with *Malus* × *domestica* (taxonomy identifier 3750) declared as the expected organism. Only EXCLUDE calls identify sequences the tool assigns confidently to another organism; REVIEW refers a sequence for manual assessment and TRIM marks a contaminant interval within an otherwise acceptable sequence. Screening followed the differential-expression analysis, so no sequence was removed before quantification and no reported statistic was recomputed; the flagged fractions are reported to show where non-plant sequence falls, not to correct any test. Absence of a flag is not positive evidence of apple origin. Per-sequence calls are in the Supplementary Table S11 workbook and, for the Rin-vs-Rola candidates, in Supplementary Table S5.

| Item | Sequences | *Note* |
|---|---|---|
| **FCS-GX actions on the 249,983 gene-representative sequences** | | |
| EXCLUDE | *99,047* | 50,397,640 bp; assigned confidently to another organism |
| REVIEW | *281* | 182,995 bp; referred for manual assessment, not a confirmed contaminant |
| TRIM | *4* | 11,321 bp flagged within otherwise acceptable sequences |
| total actioned | *99,332* | |
| **Taxonomic division of the 99,332 actioned sequences** | | |
| animals | *51,340* | primates 18,745; insects 16,362; rodents 15,534 |
| fungi | *27,752* | ascomycetes 21,775; basidiomycetes 5,956 |
| bacteria | *19,831* | |
| viruses, archaea, protists, synthetic | *409* | 367 / 38 / 3 / 1 |
| **Most abundant taxa** | | |
| *Venturia inaequalis* | *18,480* | median length 617 bp; 33.5 % below 400 bp |
| *Homo sapiens* | *17,451* | median length 239 bp; 97.6 % below 400 bp — very short, warranting caution in biological interpretation |
| *Trialeurodes vaporariorum* | *16,183* | greenhouse whitefly |
| *Mus musculus* | *14,517* | |
| *Pelomonas aquatica* | *5,537* | |
| *Filobasidium wieringae* | *4,479* | |
| *Cutibacterium acnes* | *2,453* | |
| *Aureobasidium pullulans* | *1,188* | |
| **Sequences through the cleaning pipeline** | | |
| gene-representative set as analyzed | *249,983* | deposited in GEO with the per-sequence calls |
| after removing EXCLUDE | *150,936* | |
| after also removing REVIEW | *150,655* | conservative choice for a public deposit |
| after FCS-adaptor | *150,629* | 26 excluded, 159 trimmed |
| after removing internal TruSeq read-through | *150,621* | 20 trimmed, 8 fell below the 200 bp minimum length; this is the set archived at Zenodo |
| **Flagged fraction of the thresholded-Wald candidate sets (*de novo* layer)** | | |
| Rin vs Rola | *612 of 1,817 (33.7 %)* | 97.1 % of the flagged are *V. inaequalis* |
| Rin vs control | *2,001 of 3,907 (51.2 %)* | 93.4 % |
| Rola vs control | *4,571 of 5,736 (79.7 %)* | 98.1 % |
| **Flagged fraction by cross-layer class, Rin vs Rola (Supplementary Table S5)** | | |
| candidate in both layers, same direction | *2 of 815 (0.2 %)* | |
| Trinity signal stronger, GDDH13 match | *219 of 330 (66.4 %)* | |
| GDDH13 match, no clean ortholog | *162 of 274 (59.1 %)* | |
| Trinity-only, unannotated | *223 of 385 (57.9 %)* | |
| Trinity-only, plant-supported | *2 of 4 (50.0 %)* | |
| opposite direction | *4 of 9 (44.4 %)* | |
