## Supplementary figures and images for "Transcriptome of apple cv. Ametyst in *Rvi6*-effective and *Rvi6*-breaking *Venturia inaequalis* interactions identifies defense candidates"

### 02_SupplementaryFigure_S1_workflow.pdf

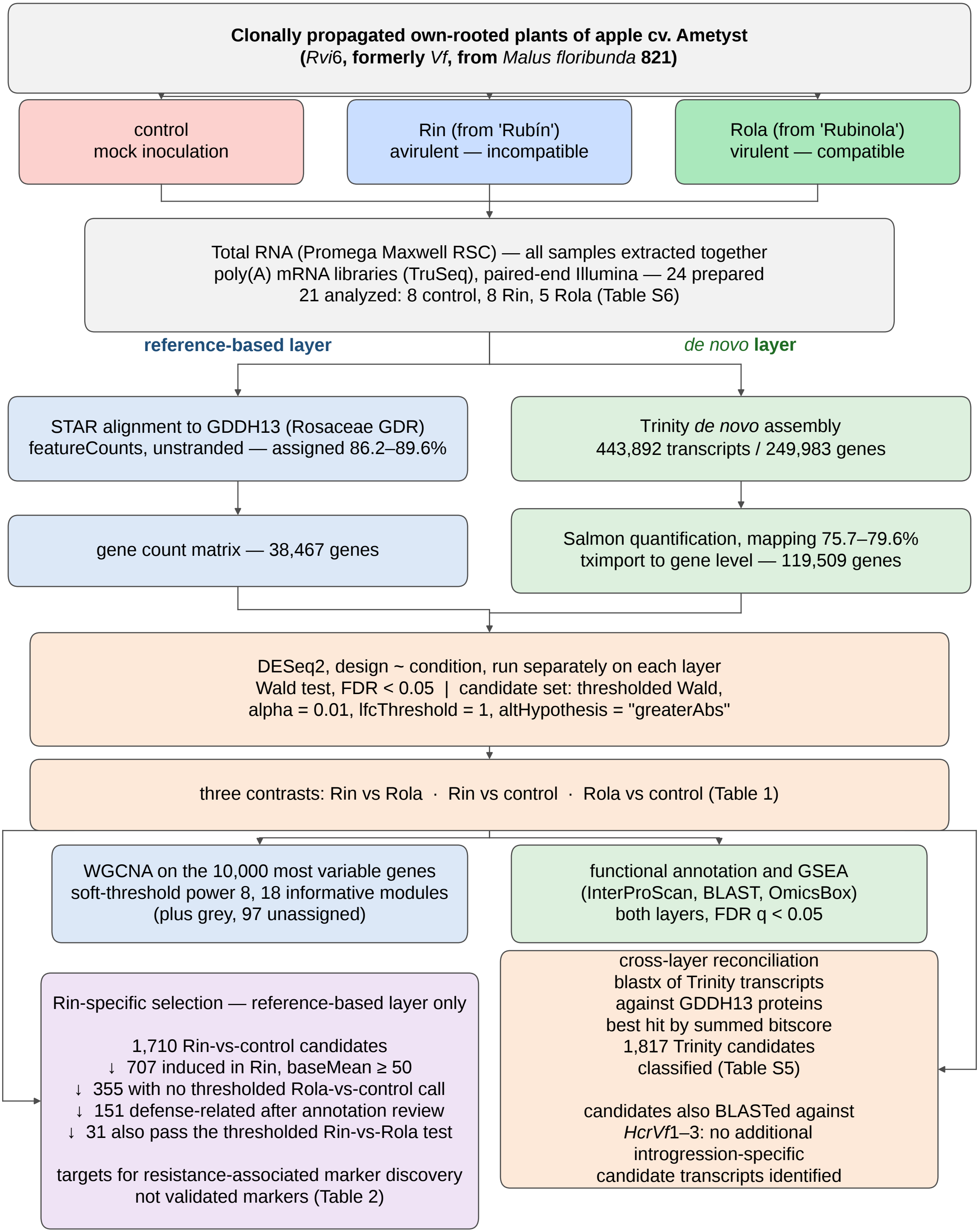

### 03_SupplementaryFigure_S2_heatmap.pdf

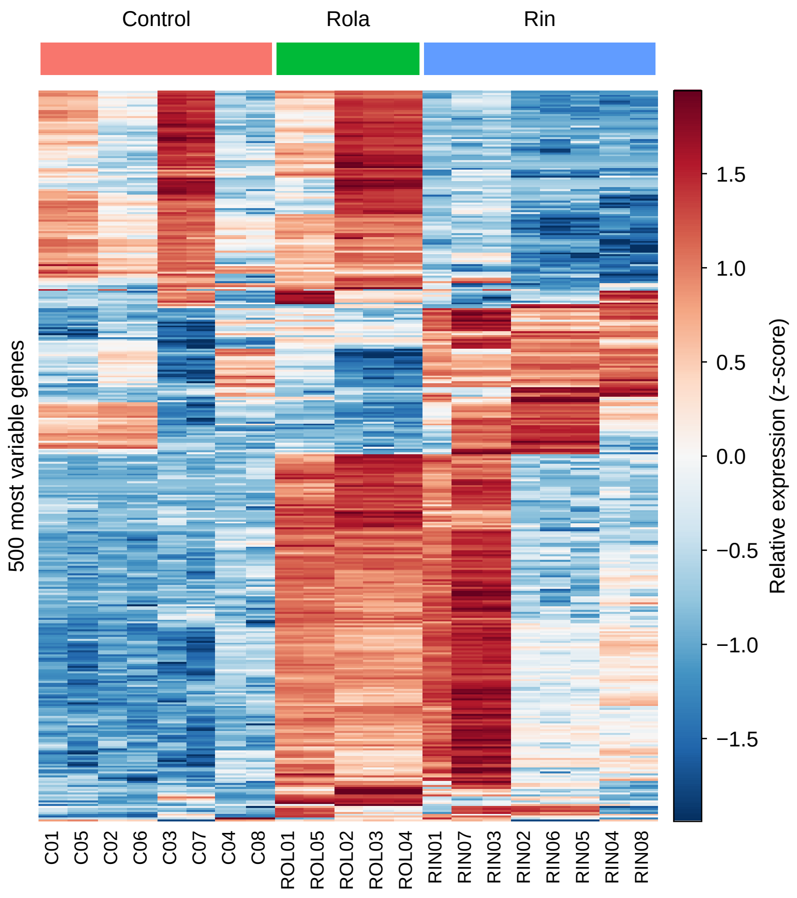

### 04_SupplementaryFigure_S3_trinity_pca.pdf

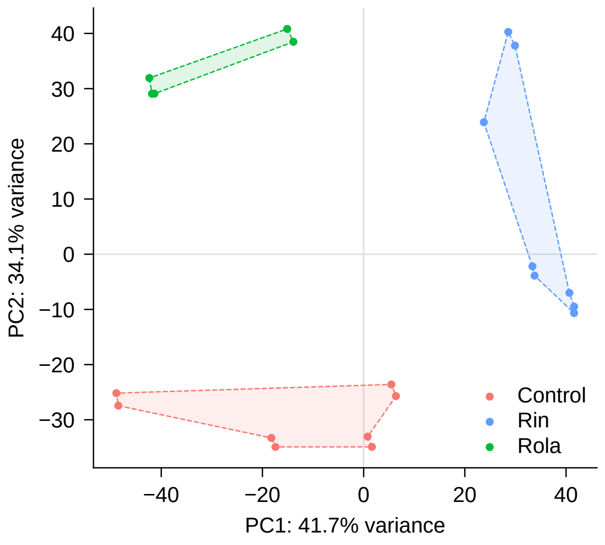

### 05_SupplementaryFigure_S4_trinity_gsea.pdf

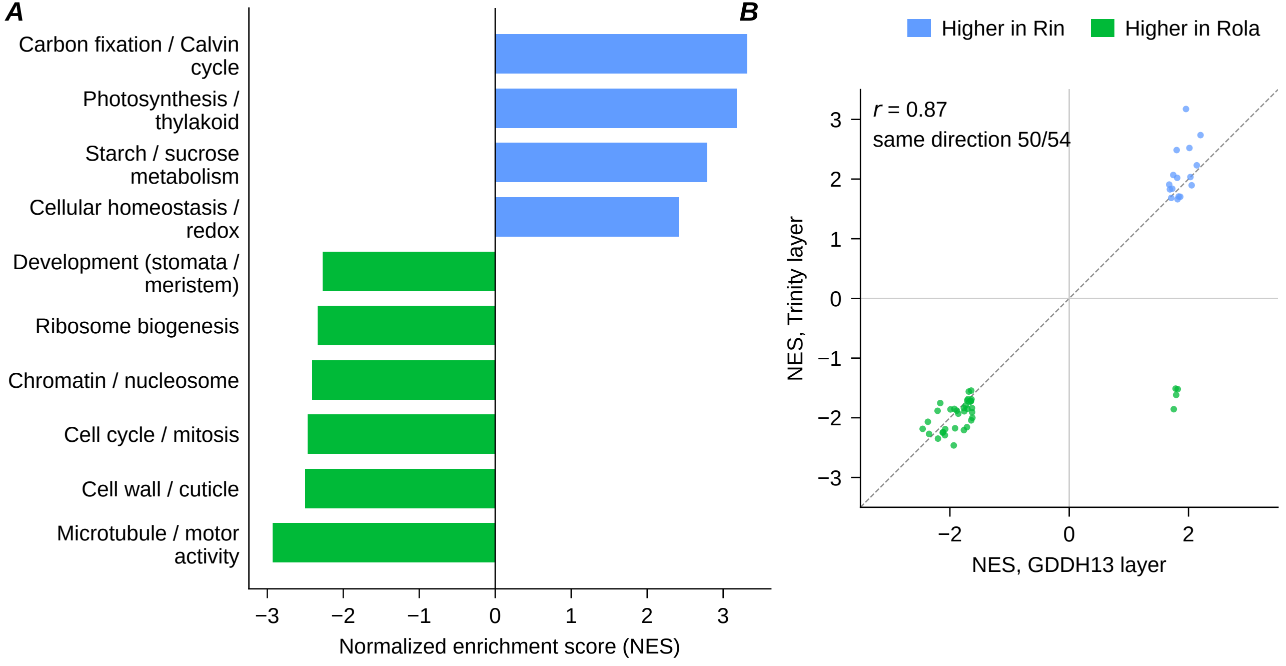

### 06_SupplementaryFigure_S5_qc.pdf

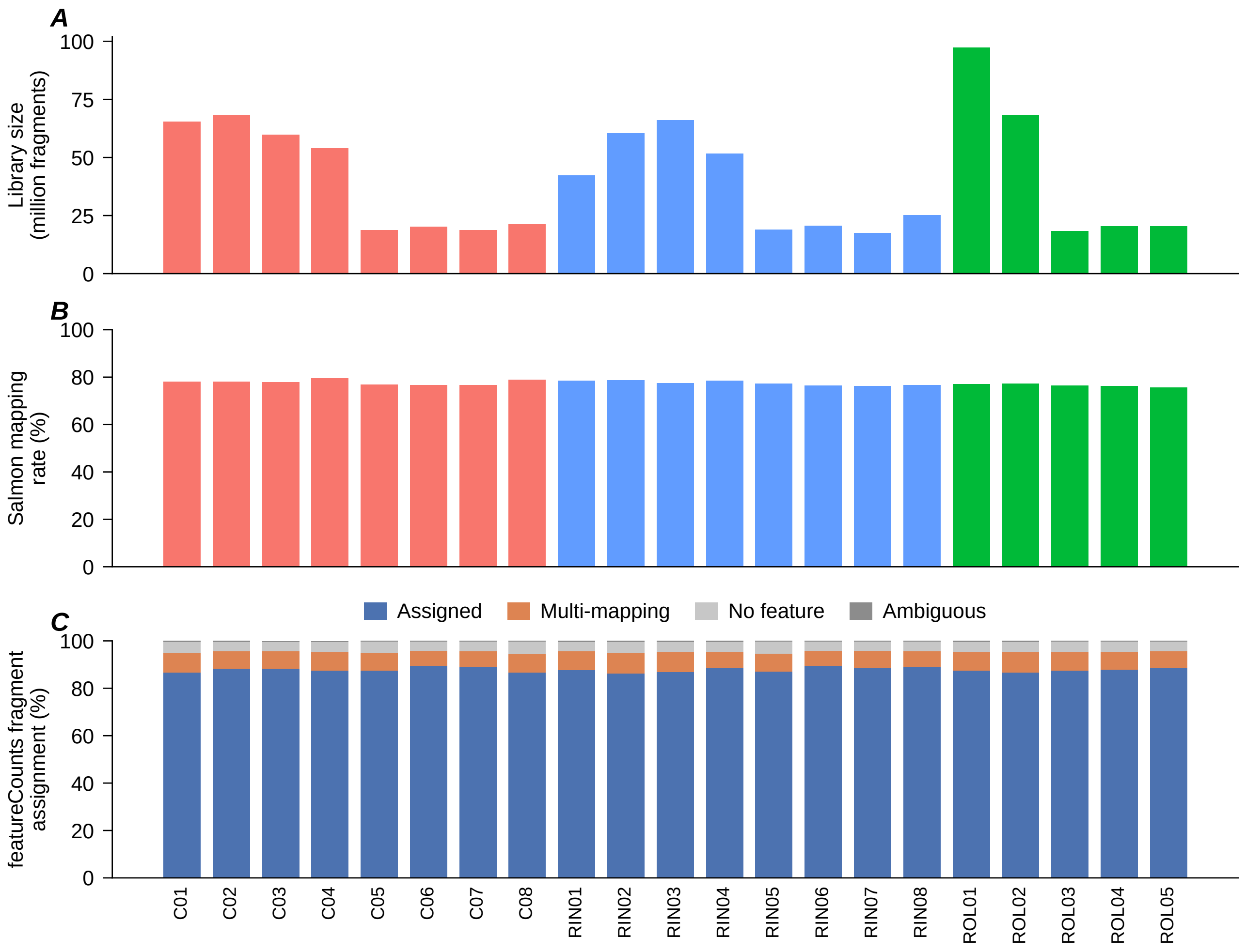
